# IF1 Protein Controls Aging Rate

**DOI:** 10.1101/2021.10.28.466310

**Authors:** Michael David Forrest

## Abstract

IF1 protein inhibits F_1_F_0_ ATP hydrolysis (and not F_1_F_0_ ATP synthesis). Across investigated species more IF1 protein, and less F_1_F_0_ ATP hydrolysis, correlates with greater maximal lifespan. Increased IF1 protein, and decreased F_1_F_0_ ATP hydrolysis, safely reduces a biomarker of aging in mice. Body temperature decrease, in mice administered with a small molecule drug that selectively inhibits F_1_F_0_ ATP hydrolysis (which doesn’t inhibit F_1_F_0_ ATP synthesis), is evidence that F_1_F_0_ ATP hydrolysis is used for metabolic heat generation *in vivo*. Instrumental to homeothermy, which is a new fundamental discovery. A further discovery is that cancer cells subvert F_1_F_0_ ATP hydrolysis to drive their distinctive Warburg metabolism and so selective drug inhibition of F_1_F_0_ ATP hydrolysis exerts potent anticancer activity. When the body is in an ambient temperature of 37°C (or more), no metabolic heat generation is needed for the body to be at 37°C, and so a large dose of a F_1_F_0_ ATP hydrolysis inhibiting anticancer drug may be administered, which may slow aging. So, here might be an entirely new class of anticancer drugs that may (when appropriately used) help, instead of harm, normal cells. Distinct from present anticancer drugs, which greatly harm normal cells, causing horrific side-effects, which kill many and cause many others to abandon cancer treatment.

In short, this paper teaches how mammals metabolically generate heat, why different mammal species have different maximal lifespans, and new anticancer drugs, that are predicted to slow aging.

**SIGNIFICANCE:** Has nature taught us how to slow aging? Different mammal species age at different rates, conferring different maximal lifespans. For example, the maximal lifespan of a mouse is 4 years, while that of a bowhead whale is 211 years. So, aging is modifiable. But how? A clue might be body size: smaller mammal species tend to age faster than larger ones. In geometry, by its square-cube law, smaller objects have a greater surface-area to volume ratio than larger objects. Meaning smaller mammal species more readily lose their metabolically generated heat. And so, per unit time, each gram of a smaller mammal species needs to generate more metabolic heat than each gram of a larger mammal species, to keep their body temperature around 37°C. The chemical reactions that the body uses to obtain energy from food (e.g., to keep the body warm) produce harmful by-products: Reactive Oxygen Species (ROS), which cause molecular damage. The accumulation of which might be aging. Per unit time, each gram of a smaller mammal species generates more metabolic heat, uses more food, produces more ROS, and ages more.

Newly reported herein is a chemical reaction that homeotherms use to generate heat (F_1_F_0_ ATP hydrolysis). By the 2^nd^ Law of Thermodynamics, whenever energy converts from one form to another, some of this energy must be dissipated as heat (no energy conversion can be 100% efficient). I’ve discovered, in homeotherms, ATP synthase enzyme hydrolyses some of the ATP it synthesizes (i.e., performs F_1_F_0_ ATP hydrolysis). Causing futile cycling between ATP synthesis and ATP hydrolysis, conditional upon passing and pumping protons along a concentration gradient respectively. So, cyclically interconverting between potential and chemical energies, which (by the inefficiency of energy conversions) generates heat to maintain body temperature.

Across a set of mammal species: per unit time, each gram of smaller (shorter-living) mammal species do more of this heat generating reaction (F_1_F_0_ ATP hydrolysis) than each gram of larger (longer-living) mammal species. Because they have less IF1 protein (activity per unit mass), where IF1 protein selectively inhibits F_1_F_0_ ATP hydrolysis (*doesn’t inhibit F_1_F_0_ ATP synthesis*). Across these mammal species, maximal lifespan is inversely proportional to the use (per unit time per unit mass) of F_1_F_0_ ATP hydrolysis. That drives the inverse proportionality between metabolic rate per unit mass and maximal lifespan, which causes the inverse proportionality between heart rate and maximal lifespan, observed across these mammal species. Increased IF1 protein, and decreased F_1_F_0_ ATP hydrolysis, safely reduces a biomarker of aging in mice. So, correlational and interventional data.

My interpretation of data herein is that different mammal species have different maximal lifespans because of different IF1 protein activity (per unit mass). Where more IF1 protein activity (per unit mass) confers longer lifespan.

A small-molecule drug that selectively inhibits F_1_F_0_ ATP hydrolysis, which doesn’t inhibit F_1_F_0_ ATP synthesis, is shown to dose-dependently reduce metabolic heat generation (and metabolic rate thereby) in mice. Higher dose reduces it more. Such a drug is predicted to slow aging. Indeed, its mechanism of action (selectively inhibiting F_1_F_0_ ATP hydrolysis) is shown to safely decrease intracellular ROS concentration in mice.

Less metabolic heat generation doesn’t necessarily mean lower body temperature. Body temperature can be the same with less metabolic heat generation by proportionally greater body insulation, such as wearing more or better clothing, and/or a conducive ambient temperature. A human, in typical clothing, is most comfortable at an ambient temperature around 20.3°C. But much of the world is hotter, at least for part of the year, especially when close to the equator (43% of the world’s population lives in the tropics). Such a drug might, by dose-dependently reducing metabolic heat generation, increase thermal comfort in hot places, possibly slowing aging. To illustrate: a relatively small drug dose might increase a clothed person’s preferred ambient temperature to 23°C, a higher dose to 27°C, an even higher dose to 32°C, and so on. When metabolic heat generation is low, the preferred ambient temperature is close to 37°C. When the ambient temperature is 37°C or more, no metabolic heat generation is needed for the body to be at 37°C.

I predict when such a drug is applied topically to a small body part, such as to the face in a cosmetic cream, it will reduce metabolic heat generation at that location, reducing metabolic rate and thereby slow aging there. Wherein heat transfer from the rest of the body, via blood flow, maintains this body part at around 37°C, because topical use can’t reduce body temperature at any ambient temperature. Less F_1_F_0_ ATP hydrolysis, enough predicted to slow aging by two-thirds, has been proven safe in mice, at least when localized to a body part.

Slowing the aging of even just a small part of the body has cosmetic and - because many diseases of aging are highly localized (*for example, to the eyes: e.g., Age-Related Macular Degeneration*) - medical applications. Probably the incidence and progression of age-related diseases correlates with age/aging because aging is causal to them, and so a single drug that slows aging might confer therapeutic benefit for many, varied diseases of aging. Such diseases *must* be beaten to avert the otherwise coming demographic/economic crisis in which too much of the population suffers, and is debilitated by, at least one of them. A drug to slow aging is a desperate want and has been since the dawn of mankind.

**GRAPHICAL ABSTRACT:** 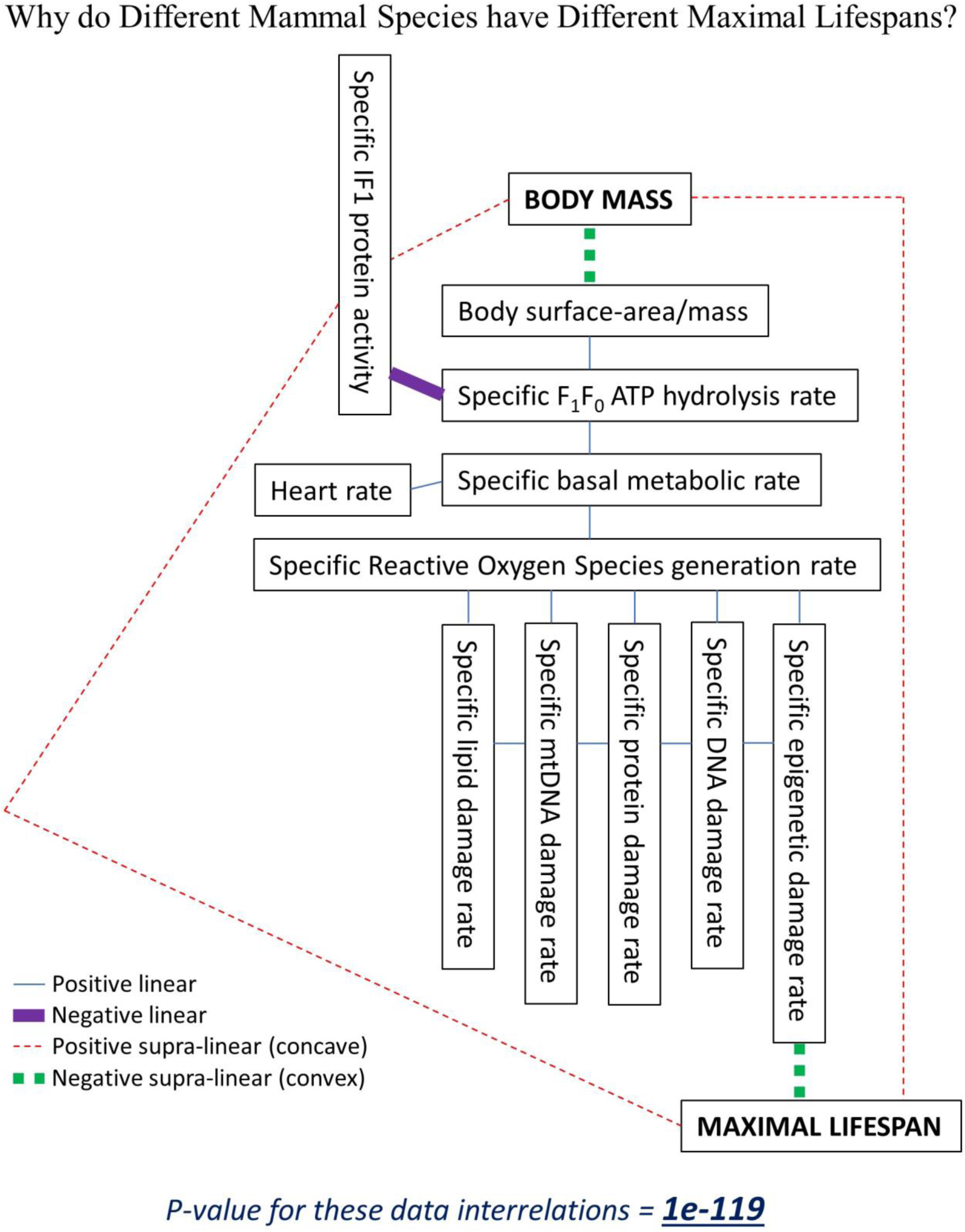

## INTRODUCTION

Different species age at different rates and have different maximal lifespans [1]. If we could discover why, elucidating how nature modulates aging, we might be able to harness this power for ourselves.

At the time of writing, a widely dismissed theory of aging is that chemical reactions in the body, those for obtaining energy from food, produce harmful by-products (Reactive Oxygen Species, ROS), and these cause molecular damage, which accumulates over time, which is aging [2–5]. This paper adds to this theory, with new experimental data, analysis, discoveries, and drugs, which I argue resuscitates it. Problems that luminaries in the field presently have with this theory, such as the underwhelming effect of antioxidants on lifespan, are addressed herein. Commonly cited [6] is reinterpreted upon collinearity I’ve discovered in its data.

Aerobic respiration converts the chemical energy of food into the chemical energy of ATP [7]. An intermediate in this process is the generation of an electrochemical gradient of protons, a proton motive force (pmf), comprising ΔpH and voltage (Ψ_IM_) components, across the inner mitochondrial membrane. ATP synthase is in this membrane. It can use the pmf to generate ATP from ADP and inorganic phosphate (Pi). But ATP synthase is reversible. Depending on its substrate and product concentrations, and the pmf value, operating forwards (passing protons, making ATP) or backwards (pumping protons, consuming ATP). Its forward and reverse modes respectively. Which may also be termed F_1_F_0_ ATP synthesis and F_1_F_0_ ATP hydrolysis respectively.

IF1 protein is an endogenous protein that selectively blocks the reverse mode of ATP synthase (F_1_F_0_ ATP hydrolysis), which does ***<u>not</u>*** block the forward mode of ATP synthase (F_1_F_0_ ATP synthesis) [8–14] (Figure 1). It is widely considered irrelevant during aerobic respiration because (*i*) ATP synthase is solely performing F_1_F_0_ ATP synthesis, and (*ii*) IF1 protein activity is pH sensitive [15], being low at the normal mitochondrial matrix pH (∼pH 8). But it is considered significant upon matrix acidification. Caused by collapse of the pmf across the mitochondrial inner membrane. Which causes ATP synthase to reverse its operation, performing F_1_F_0_ ATP hydrolysis instead, consuming cellular ATP. This can occur, for example, during ischemia [16–17]. When a tissue, or part(s) thereof, is completely/partially cut off from blood flow, foregoing respiratory substrates and O_2_ delivery. So, the present consensus is that IF1 protein is a safety device. Important in mitigating pathology, but irrelevant normally.

**Figure 1.**
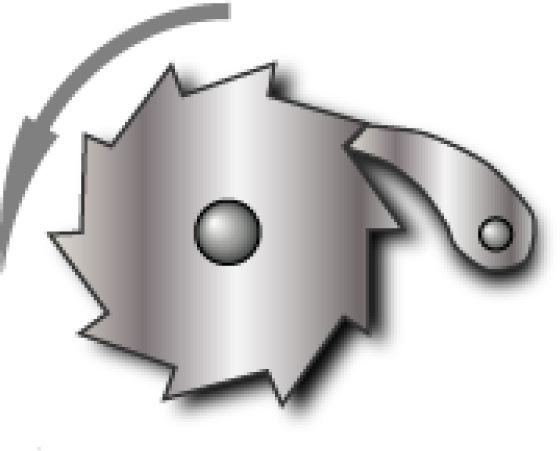
A ratchet and pawl mechanism, wherein the pawl ensures that the ratchet cog can only rotate in the direction shown by the arrow [55]. IF1 protein’s inhibition is unidirectional. Only of F_1_F_0_ ATP hydrolysis, it not inhibiting F_1_F_0_ ATP synthesis [8–14]. ATP synthase comprises a molecular rotor, which rotates one way when performing ATP synthesis [56]. And the opposite way when performing ATP hydrolysis. But F_1_F_0_ ATP hydrolysis is not simply the reverse of F_1_F_0_ ATP synthesis, because they follow different molecular pathways [57–59]. Reflected, for example, by azide inhibiting F_1_F_0_ ATP hydrolysis and not F_1_F_0_ ATP synthesis [60–61] (*N.B., azide also inhibits Complex IV* [62] *and so is a metabolic poison*). IF1 protein can only bind ATP synthase when it is hydrolysing ATP. When bound it acts analogously to a pawl, blocking the rotational direction of ATP hydrolysis, but not that of ATP synthesis. It being driven off/disassociated by the action/rotation of ATP synthesis. IF1 protein binding to ATP synthase blocks the rotational direction of ATP hydrolysis directly (by its N-terminal section directly binding ATP synthase’s central shaft γ subunit [13]). And indirectly by preventing ATP hydrolysis (by its binding imposing at one of ATP synthase’s three catalytic centres the structure and properties of the β_TP_-α_TP_ interface upon the β_DP_-α_DP_ interface that it binds, thereby preventing this centre hydrolysing its bound ATP [10–13]).

By contrast, I herein report that F_1_F_0_ ATP hydrolysis ***<u>does</u>*** occur normally. It being important for generating metabolic heat. Whilst IF1 protein activity, although subdued by the normal alkaline pH of the mitochondrial matrix, is *usefully* non-zero. Wherein the amount of IF1 protein activity constrains the amount of F_1_F_0_ ATP hydrolysis (metabolic heat generation).

Why does F_1_F_0_ ATP hydrolysis occur? A chemical reaction away from its equilibrium is a form of potential energy, that releases energy as it moves towards its equilibrium, which may be utilized to do work. In a cell the ATP ⇌ ADP + Pi reaction is maintained at great disequilibrium, with a very high ATP:ADP ratio, meaning every ATP hydrolysis (i.e., towards equilibrium) releases energy, which can be harnessed to do work [7] (with remainder energy dissipated as heat). It is the magnitude of this disequilibrium that confers the amount of energy available upon each ATP hydrolysis in a cell. And (*I’ve interpreted experimental data herein to teach that*) because this disequilibrium is maintained so high, there are inherently fluctuating microdomains in the mitochondrial matrix where the ATP:ADP ratio is temporarily so great that the ATP synthase molecule(s) in that microdomain reverses its operation and hydrolyses, instead of synthesizes, ATP for a period. All be it this occurs less than it would if it were not for IF1 protein activity, which - by unidirectionally inhibiting the reverse mode of ATP synthase - enables a greater ATP:ADP ratio in the cell, which increases the energy available from each ATP hydrolysis. However, at least in homeotherms, IF1 protein activity is delimited because (as aforementioned) F_1_F_0_ ATP hydrolysis has an actual purpose, that of metabolic heat generation. Futile (heat generating) cycling of ATP synthesis and hydrolysis.

I interpret that the different maximal lifespans, in different mammal species, are (at least partially) because of different specific F_1_F_0_ ATP hydrolysis rates, set by different specific IF1 protein activities. Herein I show, in a species set, (larger) longer-living mammal species have more specific IF1 protein activity, and a lower specific F_1_F_0_ ATP hydrolysis rate. Moreover, that increased IF1 protein, and decreased F_1_F_0_ ATP hydrolysis, safely reduces a biomarker of aging in mice. Consistent with IF1 protein activity being a molecular determinant of aging rate and maximal lifespan. Teaching that a drug mimicking IF1 protein activity, selectively inhibiting F_1_F_0_ ATP hydrolysis (*without inhibiting F_1_F_0_ ATP synthesis*), might slow aging and extend the lifespan of mammals. If administered when ambient temperature (and/or body insulation) ensures no, or only a safe, body temperature drop. Such drugs, biologics (IF1 protein derivatives) and small molecules, are taught in my international (Patent Cooperation Treaty, PCT) patent applications [18–20]. Some small-molecule examples therefrom are used in experiments reported herein.

To repeat, a theory of aging is that metabolic rate, that is oxygen consumption rate, enables life but causes damaging byproducts and accumulating damage, which is aging, that ultimately takes life away. This paper teaches drugs that slow metabolic rate which may, if this theory is correct, slow aging. Indeed, they usefully confer means to test this theory. Promisingly, others have shown that reducing metabolic heat generation (and metabolic rate thereby) increases mice lifespan, wherein they intervened by making a genetic change in the brain that made the body produce less metabolic heat [21–22]. Moreover, reducing metabolic rate increases the lifespan of worms [23], flies [24–26], wasps [27], and fish [28]. Humans with lower metabolic rate per unit mass tend to live longer, over subsequent decades of monitoring [29]. Humans with lower body temperature (perhaps at least in part because of less metabolic heat generation) tend to live longer, over subsequent decades of monitoring [30]. Calorie restriction, which is well-known to increase the lifespan of at least model organisms [31–32], reduces (*including in humans*) metabolic heat generation and metabolic rate [33–39]. So, in the literature, there is already data - including interventional and human data – to suggest that lifespan is inversely proportional to metabolic rate per unit mass.

In this paper’s Supplementary Material it is shown, across 517 mammal and bird species, that maximal lifespan inversely correlates with (basal) metabolic rate per unit mass. At great statistical significance.

Reactive Oxygen Species (ROS) concentration increases with age/aging [40–43]. As a correlate of aging, this is a “biomarker” of aging. Reduction (reversal) of a biomarker of aging indicates reducing (reversing) of aging [44]. Herein, it is shown that inhibiting F_1_F_0_ ATP hydrolysis (*without inhibiting F_1_F_0_ ATP synthesis*) greatly decreases ROS concentration in mice. So, demonstrated *in vivo* reduction (reversal) of a biomarker of aging. Supporting, shown in mice herein, inhibiting F_1_F_0_ ATP hydrolysis (*without inhibiting F_1_F_0_ ATP synthesis*) reduces metabolic rate, where reducing metabolic rate extends lifespan in mice [21–22], and other species [23–28].

Instrumental variable analysis can infer causality [45]. Presented instrumental variable analysis reports causality between lower specific F_1_F_0_ ATP hydrolysis rate and higher maximal lifespan, and causality between lower specific basal metabolic rate and higher maximal lifespan.

A theory of aging, which leaders in the field presently favour (e.g., [46]), is that aging is caused by epigenetic changes. But although these correlate with age/aging (i.e., they are a biomarker of aging) there is presently no evidence for causality. Are these changes caused by, or causal to, aging? Moreover, no one knows what causes these epigenetic changes, dubbed the “ticking” of the “epigenetic clock”. Herein, I present novel analysis that suggests that these epigenetic changes are a form of metabolic damage, downstream of metabolic rate cause. For example, showing across species that epigenetic change rate linearly correlates with specific basal metabolic rate. Presented instrumental variable analysis reports causality between specific basal metabolic rate and epigenetic change rate.

A specific F_1_F_0_ ATP hydrolysis inhibiting drug (*that doesn’t inhibit F_1_F_0_ ATP synthesis*) used for anticancer purpose has the side-effect of decreasing metabolic heat generation but, unlike the side-effects of present cancer drugs (which kill many [47–49]), this side-effect is mitigable, by conducive ambient temperature and/or body insulation, and may even confer benefit (slower aging).

Preprint versions of this present paper were published on bioRxiv in 2021 and 2022, but with different titles than the present title [50–53]. This new version uses data from more species, now >500 different species, and uses a greater diversity of statistical techniques, to support its teaching. It is simultaneously shorter, but with more support, by the introduction of a Supplementary Material (>300 pages) section. Earlier, this paper’s teaching was disclosed, with much more detail on at least some aspects, in some of this author’s international (PCT) patent applications, which published from 2018 onwards [18–20]. That F_1_F_0_ ATP hydrolysis is a novel cancer drug target was first suggested, from computational modelling, by this author in their bioRxiv preprint published in 2015 [54], which put forward a biophysical explanation for the longstanding mystery of why cancer cells tend to have a more hyperpolarised mitochondrial membrane potential than normal cells.

Some findings of this paper in brief: F_1_F_0_ ATP hydrolysis is used for metabolic heat generation, maximal lifespan inversely correlates with it across mammal species, inhibiting it reduces a biomarker of aging in mice, and it is a new cancer drug target.

## RESULTS

### New experimental data

If F_1_F_0_ ATP hydrolysis is a drive of metabolic heat generation, then systemic, selective drug inhibition of F_1_F_0_ ATP hydrolysis (without inhibition of F_1_F_0_ ATP synthesis) should reduce metabolic heat generation in a subject. Which should, *if* the ambient temperature is lower than their normal body temperature, reduce their body temperature towards the ambient temperature. This is experimentally observed in Figure 2.

**Figure 2.**
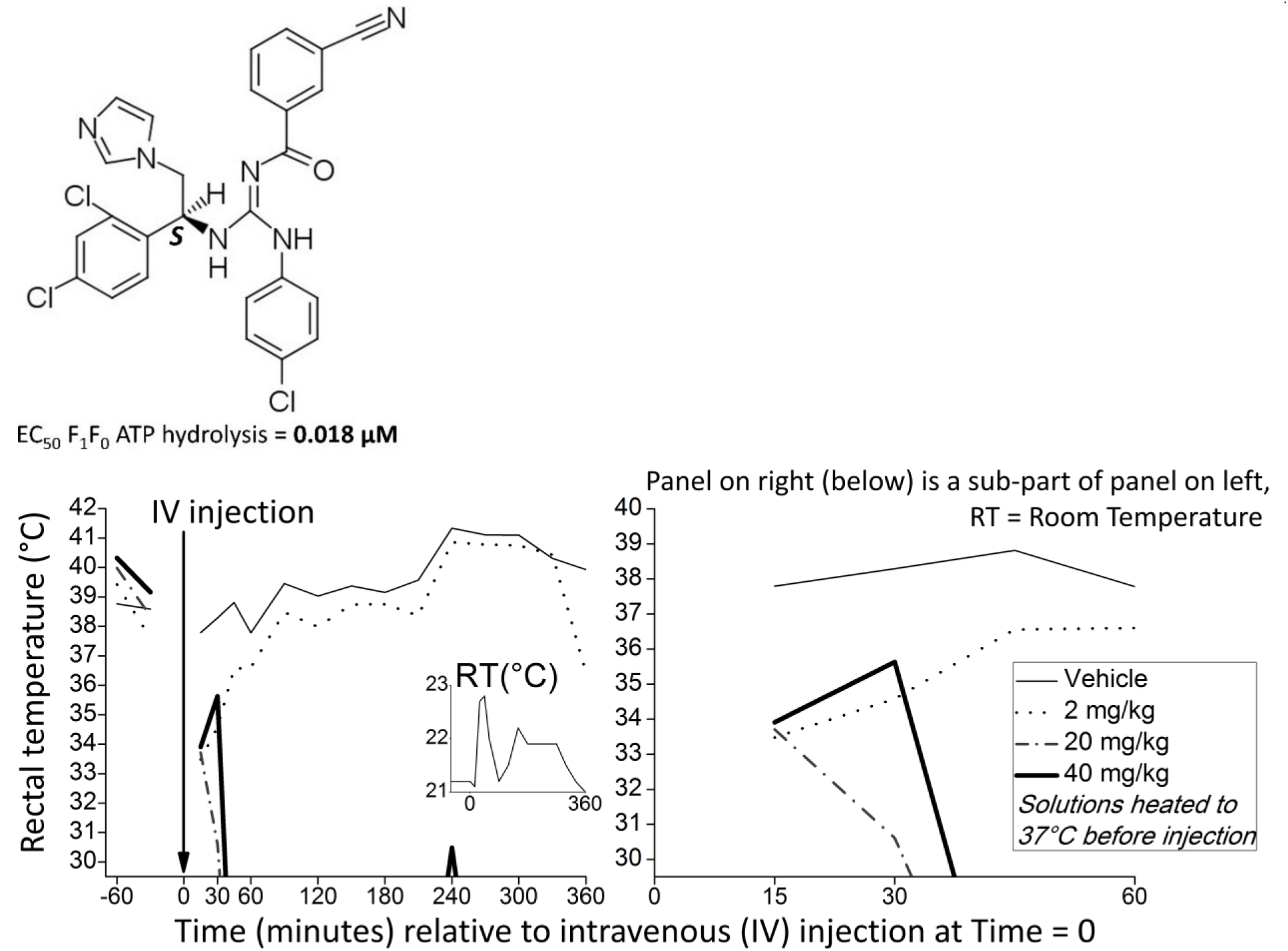
Selectively inhibiting F_1_F_0_ ATP hydrolysis (without inhibition of F_1_F_0_ ATP synthesis) in mice reduces their rectal temperature towards room temperature. Room temperature (RT) was around 22°C. Drug structure shown (with enantiomeric excess (ee) of *S* stereoisomer ≥97%) was systemically (intravenously) administered to mice, whose rectal temperatures were monitored. This drug selectively, and potently, inhibits F_1_F_0_ ATP hydrolysis (EC_50_ = 0.018 μM [=18 nM], in a Sub-Mitochondrial Particle {SMP} assay {in which *<u>no</u>*inhibition of F_1_F_0_ ATP synthesis by this drug was observed}) [63–64]. Four mice were used. One was administered vehicle (control), and the other three differing doses of the drug: 2 mg/kg, 20 mg/kg, and 40 mg/kg (all solutions heated to 37°C before administration). Rectal thermistor used couldn’t measure rectal temperatures lower than 30°C. Both 20 and 40 mg/kg (but not 2 mg/kg) drug doses made rectal temperature fall below 30°C and out of range. Although the ensuing uncertainty is bounded because rectal temperature cannot fall below room temperature (RT): RT ≤ rectal temperature < 30°C. Rectal temperature was recorded every 15 minutes in the 1^st^ hour after dosing, and every 30 minutes in the later hours shown. It was also recorded at 60 and 30 minutes before dosing. For each mouse, the 1^st^ rectal temperature recording is typically of an atypically high body temperature. Which is associated with the stress of being handled, which a mouse typically becomes habituated to over the course of the experiment. This handling effect has been reported in other rectal thermistor studies of rodents, e.g. [66]. Statistics upon this data: For the 360 minutes (6 hours) after intravenous (IV) injection: comparing data for vehicle (mean rectal temperature = 39.50°C*, median = 39.41, standard deviation = 1.16, Standard Error of the Mean, SEM = 0.3104, data does not differ significantly from that which is normally distributed by the Kolmogorov-Smirnov test of normality*) and 2 mg/kg drug (mean rectal temperature = 38.06°C, *median = 38.43, standard deviation = 2.31, Standard Error of the Mean, SEM = 0.6175, data does not differ significantly from that which is normally distributed by the Kolmogorov-Smirnov test of normality*): independent t-test [degrees of freedom=(N1-1)+(N2-1)=(14-1)+(14-1)=26)]: t-value = 2.08873, one-tailed (because alternative hypothesis is directional) p-value = 0.023333, i.e. statistically significant at p<0.05 (Using Welch’s t-test instead: t-value=2.0887, one-tailed p-value= 0.023335. Mann-Whitney U Test: U-value = 58, z-score = 1.81493, one-tailed p-value = 0.03515). With a large effect size [67–71]: Cohen’s *d* = 0.787833, Hedges’ *g* = 0.787833, Glass’s *delta* = 1.241379. Given that the 20 mg/kg and 40 mg/kg data are even more distinct from the vehicle data (than the 2 mg/kg data is), they must also each be (by t testing) significantly different than vehicle data at p<0.05. And with, clearly (by visual inspection), a larger effect size. For the 360 minutes after intravenous (IV) injection: comparing the rectal temperature data of all the mice (abstracting rectal temperatures below 30°C to all be 29.5°C, which probably subtracts from the differences to vehicle control {and perhaps to each other}, attenuating statistical significance and effect size) using the Kruskal-Wallis H Test (*which, unlike one-way ANOVA, doesn’t assume normality of data*): H statistic = 39.0917, p-value = 0.00000002. With a large effect size: η^2^ (eta–squared, its classical formulation [72]) = 0.77. Under author’s direction, this mouse study was conducted by Crown Bioscience, a Contract Research Organization (CRO). Drug synthesis was by WuXi AppTec, another CRO.

The drug administered to mice to produce Figure 2 was developed decades ago by Bristol Myers Squibb (BMS), a leading pharmaceutical company, to treat ischemia [63–64] (*never tested in humans. Orally bioavailable, with good pharmacokinetics, in rats* [63]*. Abandoned when BMS business strategy switched away from ischemia because its acuteness deemed commercially unattractive [big pharmaceutical companies continually switch in and out of disease areas upon strategy changes, which can coincide with management staff changes]*). This drug potently inhibits F_1_F_0_ ATP hydrolysis. But ***<u>not</u>*** F_1_F_0_ ATP synthesis.

As shown by assays with Sub-Mitochondrial Particles (SMPs; formed by sonicating mitochondria). Where F_1_F_0_ ATP hydrolysis is assayed by a spectroscopic assay for NADH fluorescence, which incubates SMPs with reagents that include pyruvate kinase and lactate dehydrogenase enzymes (*F_1_F_0_ ATP hydrolysis confers ADP for pyruvate kinase to convert phosphoenolpyruvate to pyruvate, producing ATP thereby, whilst lactate dehydrogenase converts pyruvate to lactate, and NADH to NAD^+^. So, the amount of F_1_F_0_ ATP hydrolysis can be read out by the decrease of NADH fluorescence*). F_1_F_0_ ATP synthesis is assayed by a spectroscopic assay for NADPH fluorescence, which incubates SMPs with reagents that include hexokinase and glucose-6-phosphate dehydrogenase enzymes (*F_1_F_0_ ATP synthesis confers ATP for hexokinase to convert glucose to glucose 6-phosphate, producing ADP thereby, whilst glucose-6-phosphate dehydrogenase converts glucose 6-phosphate to 6- phosphogluconolactone, and NADP^+^ to NADPH. So, the amount of F_1_F_0_ ATP synthesis can be read out by the increase of NADPH fluorescence*).

A drug shown to inhibit F_1_F_0_ ATP hydrolysis, and not inhibit F_1_F_0_ ATP synthesis, in SMPs has been shown to do the same in isolated (*ex vivo*) rat heart [65]. Shown to conserve ATP during ischemia. Without reducing [ATP] under normal conditions. Thence shown to be distinct from drugs that inhibit both modes of ATP synthase (such as oligomycin) which, by contrast, are shown to reduce [ATP] under normal conditions.

Figure 2 indicates that F_1_F_0_ ATP hydrolysis is a major determinant of metabolic heat generation (and so of metabolic rate). Its data suggests that during normal aerobic respiration, ATP synthase synthesizes ATP, but also hydrolyses a fraction of the ATP synthesized, which means more ATP needs to be synthesized. Which is a futile cycle, dissipating proton motive force (pmf) to heat, because of the inherent inefficiency of any energy conversion (2^nd^ Law of Thermodynamics). And because it, by conferring a proportion of ATP synthase molecules not passing protons, but instead pumping protons back, increases the rate of proton leak. Wherein pumped protons return to the mitochondrial matrix outside of ATP synthase. Their potential (electrochemical) energy dissipated as heat rather than (partially) captured in ATP. All of this drives a higher rate of aerobic respiration. Where the F_1_F_0_ ATP hydrolysis rate, increasing the rate of aerobic respiration (metabolic rate), is constrained by the amount of IF1 protein activity (per the amount of ATP synthase).

A prediction atop of Figure 2, for future testing, is that if the ambient temperature was instead at the mouse’s normal body temperature (around 37°C [1]), or safely higher, then no body temperature drop would occur. Even at high drug doses. Because a mouse’s body temperature cannot be at a lower temperature than its surround (2^nd^ Law of Thermodynamics). To illustrate, by analogous example, anaesthetic can dramatically reduce a mouse’s body temperature, but *not* when the mouse is kept at an ambient temperature of 37°C [73]. At an ambient temperature of 37°C (or higher), the body doesn’t have to generate any heat to maintain a body temperature of 37°C. So, an ambient temperature of 37°C (or higher) counteracts any drop in metabolic heat production, of any magnitude.

Any drop in metabolic heat production is drug dose-dependent and is predicted inconsequential if the mice are kept at an accordingly higher ambient temperature to compensate. To illustrate: if after the selected drug dose is administered, body temperature drops by 3°C, increasing the ambient temperature by 3°C (or safely more) counterbalances, meaning body temperature remains 37°C. Or if after a *greater* selected drug dose is administered, body temperature drops by 7°C, increasing the ambient temperature by 7°C (or safely more) counterbalances, meaning body temperature remains 37°C. Where the ambient temperature never needs to exceed 37°C, no matter how large the administered drug dose. If the mouse were already at a higher ambient temperature (e.g., 35°C), for example because of a tropical location, the drug’s dose-dependent decrease in metabolic heat generation (thereby increase of thermoneutral temperature) could increase the mouse’s thermal comfort. Indeed, this drug may have utility in humans for increasing thermal comfort in hot climates (e.g., close to the equator) and in the summer of seasonally hot climates.

In the experiment reported in Figure 2, the drug-treated mice’s body temperature couldn’t have fallen to less than room temperature, which was ∼22°C. Mice can survive a body temperature as low as 16°C, which can occur during their torpor [74]. Torpor, characteristic of many small mammal species, is a period of metabolic depression. Where body temperature is below 31°C. And O_2_ consumption is less than 25% of that during normal, normothermic inactivity. With diminished responsiveness to stimuli. Prompted by low food intake and/or low ambient temperature (<20°C). Can occur frequently, even daily, depending on food regime and/or the time course of ambient temperature.

A prediction is that this drug can slow aging and increase lifespan. For example, regularly administering it to a mouse (at a dose that can slow its metabolic rate per unit mass to that of a human {slowing its heart rate from 600 to 60 beats per minute}), with the ambient temperature set accordingly, is predicted to make this mouse live much longer. Perhaps even as long as a human.

I’ve filed patent claims for methods of using this drug for various non-ischemic indications/applications. Moreover, I’ve invented better (potency) selective F_1_F_0_ ATP hydrolysis inhibiting drugs (*that don’t inhibit F_1_F_0_ ATP synthesis*). Disclosed in a series of patent applications: e.g., [18–20]. One of these applications has already been granted (i.e., determined novel and inventive) in Australia [75]. A novel drug therefrom is used later herein (Figure 19).

### Novel analysis and interpretation of publicly available data

#### Lower specific F_1_F_0_ ATP hydrolysis rate correlates with higher maximal lifespan

Figure 3 shows that, across twelve different mammal and bird species, lower specific F_1_F_0_ ATP hydrolysis rate (*in Sub-Mitochondrial Particles {SMPs} in vitro. In a nuanced assay described in the Methods section*) correlates with larger body mass, lower specific basal metabolic rate, slower heart rate, and longer maximal lifespan. Data of this figure is in Table S2 of the Supplementary Material, with data sources disclosed there.

**Figure 3.**
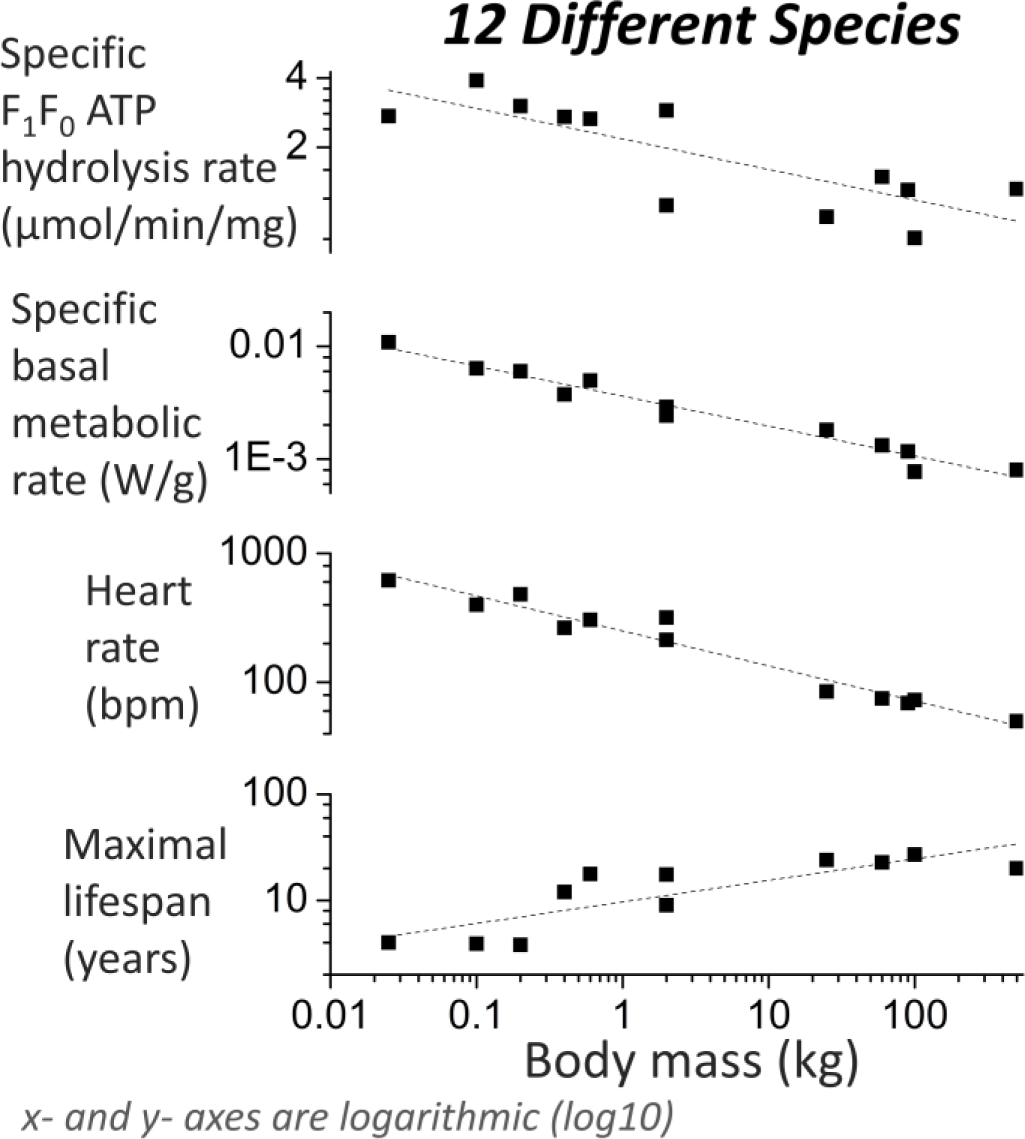
Across species, 1^st^ (top) panel shows negative correlation between specific F_1_F_0_ ATP hydrolysis rate (*mg is of mitochondrial protein*) and body mass, 2^nd^ panel shows negative correlation between specific basal metabolic rate and body mass, 3^rd^ panel shows negative correlation between heart rate (*bpm is beats per minute*) and body mass, and 4^th^ panel shows positive correlation between maximum lifespan and body mass.

Shown best-fit lines to the plots of Figure 3 are of the form:

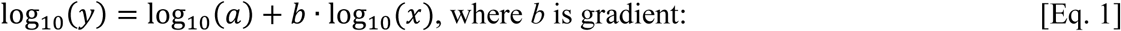

**Table 1.**

| | $\log_{10}(a)$ | $b$ | Adjusted $R^2$ |
| --- | --- | --- | --- |
| 1<sup>st</sup> panel | 0.33791 ($\pm 0.04411$) | -0.13213 ($\pm 0.03077$) | 0.61323 |
| 2<sup>nd</sup> panel | -2.44192 ($\pm 0.02381$) | -0.26591 ($\pm 0.01661$) $\approx -0.25$ | 0.95872 |
| 3<sup>rd</sup> panel | 2.39875 ($\pm 0.02593$) | -0.27123 ($\pm 0.01808$) $\approx -0.25$ | 0.95318 |
| 4<sup>th</sup> panel | 0.98595 ($\pm 0.06233$) | 0.20246 ($\pm 0.04528$) $\approx +0.25$ | 0.65512 |

Pearson correlation coefficient (R), and associated p-values (one-tailed, because each alternative hypothesis is directional), for the log_10_-log_10_ data of Figure 3:

**Table 2.**

| | | $\log_{10}(y1)$ | $\log_{10}(y2)$ | $\log_{10}(y3)$ | $\log_{10}(y4)$ |
| --- | --- | --- | --- | --- | --- |
| R | $\log_{10}(x)$ | -0.80523 | -0.98105 | -0.97849 | 0.83043 |
| p |  | 0.000788 | 0.0000000093 | 0.000000017 | 0.0007755 |

Where *x* = body mass, *y1* = specific F_1_F_0_ ATP hydrolysis rate, *y2* = specific basal metabolic rate, *y3* = heart rate, and *y4* = maximal lifespan.

Figure 4 recasts some data of Figure 3 (omitting body mass data).

**Figure 4.**
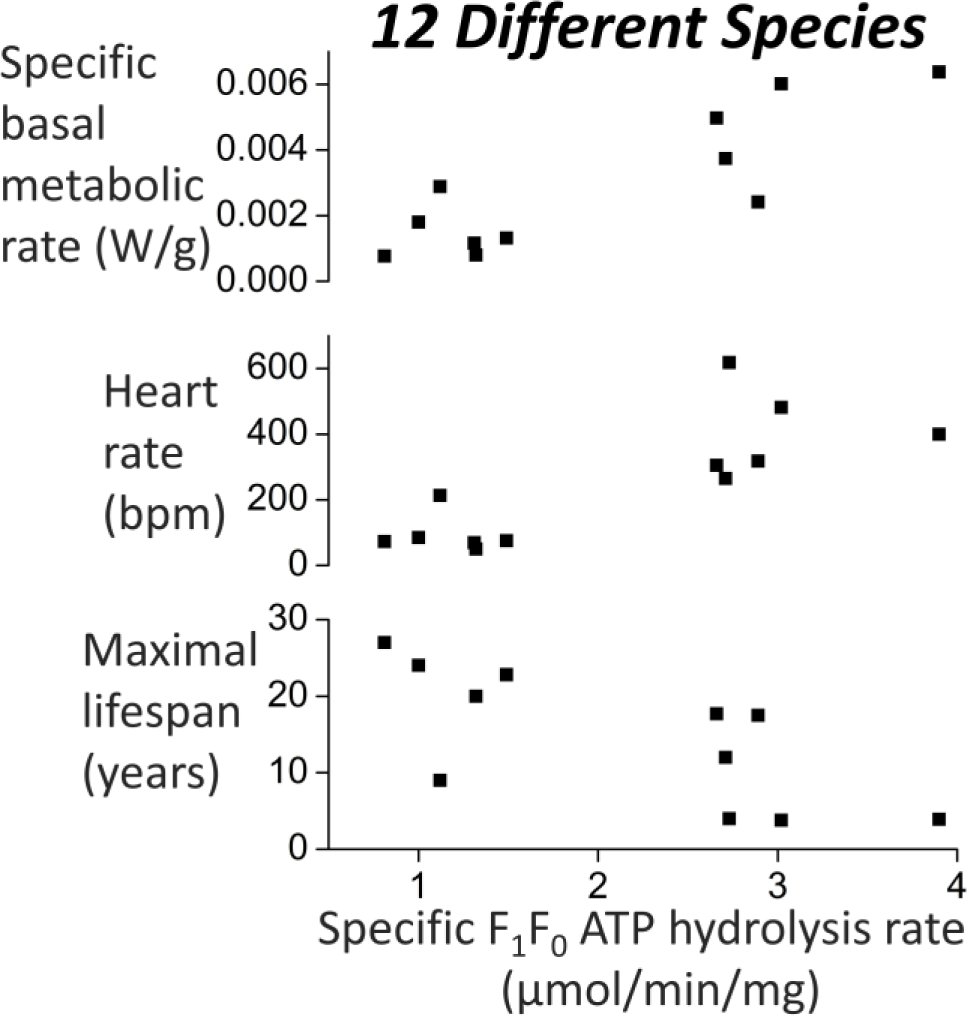
Across species, 1^st^ (top) panel shows positive correlation between specific basal metabolic rate and specific F_1_F_0_ ATP hydrolysis rate (*mg is of mitochondrial protein*), 2^nd^ panel shows positive correlation between heart rate and specific F_1_F_0_ ATP hydrolysis rate, and 3^rd^ panel shows negative correlation between maximal lifespan and specific F_1_F_0_ ATP hydrolysis rate. Herein proposed, the 1^st^ panel correlation drives the 2^nd^ and 3^rd^ panel correlations.

For the data used to produce Figure 4, Table 3 below presents Pearson correlation (R) coefficients, and associated p-values (one-tailed, because each alternative hypothesis is directional), for its inter-relations.

**Table 3.**

| Pearson correlation coefficient (R) |  |  |  |
| --- | --- | --- | --- |
|  | Specific basal metabolic rate | Heart rate | Maximal lifespan |
| Specific F<sub>1</sub>F<sub>0</sub> ATP hydrolysis rate | 0.7016 | 0.8007 | -0.7192 |
|  | Specific basal metabolic rate | 0.9476 | -0.8144 |
|  |  | Heart rate | -0.8572 |
| p-value (one-tailed) |  |  |  |
|  | Specific basal metabolic rate | Heart rate | Maximal lifespan |
| Specific F<sub>1</sub>F<sub>0</sub> ATP hydrolysis rate | 0.0054965 | 0.0008765 | 0.0063065 |
|  | Specific basal metabolic rate | 0.0000015 | 0.001136 |
|  |  | Heart rate | 0.0003725 |
| Asymptotically exact harmonic mean p-value for combining independent/dependent tests |  |  |  |
| Of all the p-values above (in this present table) = 0.000008933458 = **0.000009** |  |  |  |
| Fisher's combined probability test |  |  |  |
| Using p-values for [Specific F<sub>1</sub>F<sub>0</sub> ATP hydrolysis rate vs. Maximal lifespan] (=0.0063065) & |  |  |  |
| [Heart rate vs. Specific basal metabolic rate] (=0.0000015) = **0.0000002** |  |  |  |

The p-values are small and statistically significant, despite small values of *n*, as testament to the high *R* values. The asymptotically exact harmonic mean p-value was calculated according to the method of [76] (*its post-published corrected method, as corrected on 7 October 2019*). For comparison, for these species, the Pearson correlation coefficient between maximal lifespan and adult body mass is 0.3595 (one-tailed p-value = 0.1387675, not significant).

#### Local inhibition of F_1_F_0_ ATP hydrolysis

I suggest that when F_1_F_0_ ATP hydrolysis is inhibited locally in a subject (i.e., only in a body region) then there is no appreciable body temperature drop. Not even in the affected body region, because of heat transfer (especially via blood flow) from other body regions. Indeed, supporting, [77] has shown the safety of local inhibition of F_1_F_0_ ATP hydrolysis *in vivo*. Wherein, in forebrain neurons of mice, they reduced the amount of F_1_F_0_ ATP hydrolysis by ∼35%. By increasing their IF1 protein amount by 300%. Wherein the extra IF1 protein was (actually) human IF1 protein, with a single amino acid substitution that increases its inhibitory potency against F_1_F_0_ ATP hydrolysis at pH 8. These mice were “normal in appearance, home-cage behaviour, reproduction, and longevity up to 1-year follow-up”.

[78] showed the *in vivo* safety of locally decreasing F_1_F_0_ ATP hydrolysis, by locally increasing IF1 protein amount, in mouse liver (especially in perivenous hepatocytes): locally, extra IF1 protein decreased F_1_F_0_ ATP hydrolysis by 40% and decreased State 3 respiration rate by 44%. Wherein these mice had “no differences in weight, life span and cage behaviour when compared to controls after one year of follow up”.

[79] showed the *in vivo* safety of locally decreasing F_1_F_0_ ATP hydrolysis, by locally increasing IF1 protein amount, in mouse intestine: locally, this extra (human) IF1 protein decreased the F_1_F_0_ ATP hydrolysis capability by 35%, which decreased the oligomycin sensitive respiration rate by 60%. All safely.

#### Increased IF1 protein, and decreased F_1_F_0_ ATP hydrolysis, safely reduces a biomarker of aging in mice

Intracellular and extracellular Reactive Oxygen Species (ROS) concentrations increase with age/aging [40–43]. A “biomarker” of aging. Increasing specific IF1 protein activity, decreasing F_1_F_0_ ATP hydrolysis (*without inhibiting F_1_F_0_ ATP synthesis*), reduces (reverses) this biomarker of aging. Reduction/reversal of a biomarker of aging can be interpreted as reducing/reversing aging [44].

[77] interpret their data as showing that increased IF1 protein amount, and decreased F_1_F_0_ ATP hydrolysis, *increases* [ROS]. But this is in error. What their data (actually) shows is that increased IF1 protein amount, and decreased F_1_F_0_ ATP hydrolysis, *decreases* [ROS].

[77] transgenically increased IF1 protein amount, which decreased F_1_F_0_ ATP hydrolysis, in the forebrain neurons of mice. This manipulation furnished these cells with a lower respiration rate (lower O_2_ consumption rate). And a more hyperpolarized membrane potential across their mitochondrial inner membrane (Ψ_IM_; presumably because of less proton motive force dissipation to heat by futile cycling of F_1_F_0_ ATP synthesis and F_1_F_0_ ATP hydrolysis). Because of this greater hyperpolarisation (rendering a more negative mitochondrial matrix), these cells accumulate more positively charged ROS (superoxide) reporting MitoSOX^TM^ probe. Which means they have greater ROS signal. Which [77] mistakenly take at its face value, interpreting this as meaning greater [ROS]. However, once the Ψ_IM_ disparity is factored in, which is a contribution of the present paper, then one can see that these cells (actually) have *lower* [ROS].

[77] assay Ψ_IM_ by assaying TMRM accumulation. And assay [ROS] by MitoSOX probe fluorescence. But they don’t realize that MitoSOX, like TMRM, is also a Delocalized Lipophilic Cation (DLC). And so, its accumulation in the mitochondrial matrix is dependent upon Ψ_IM_. Indeed, to an even greater degree than TMRM:

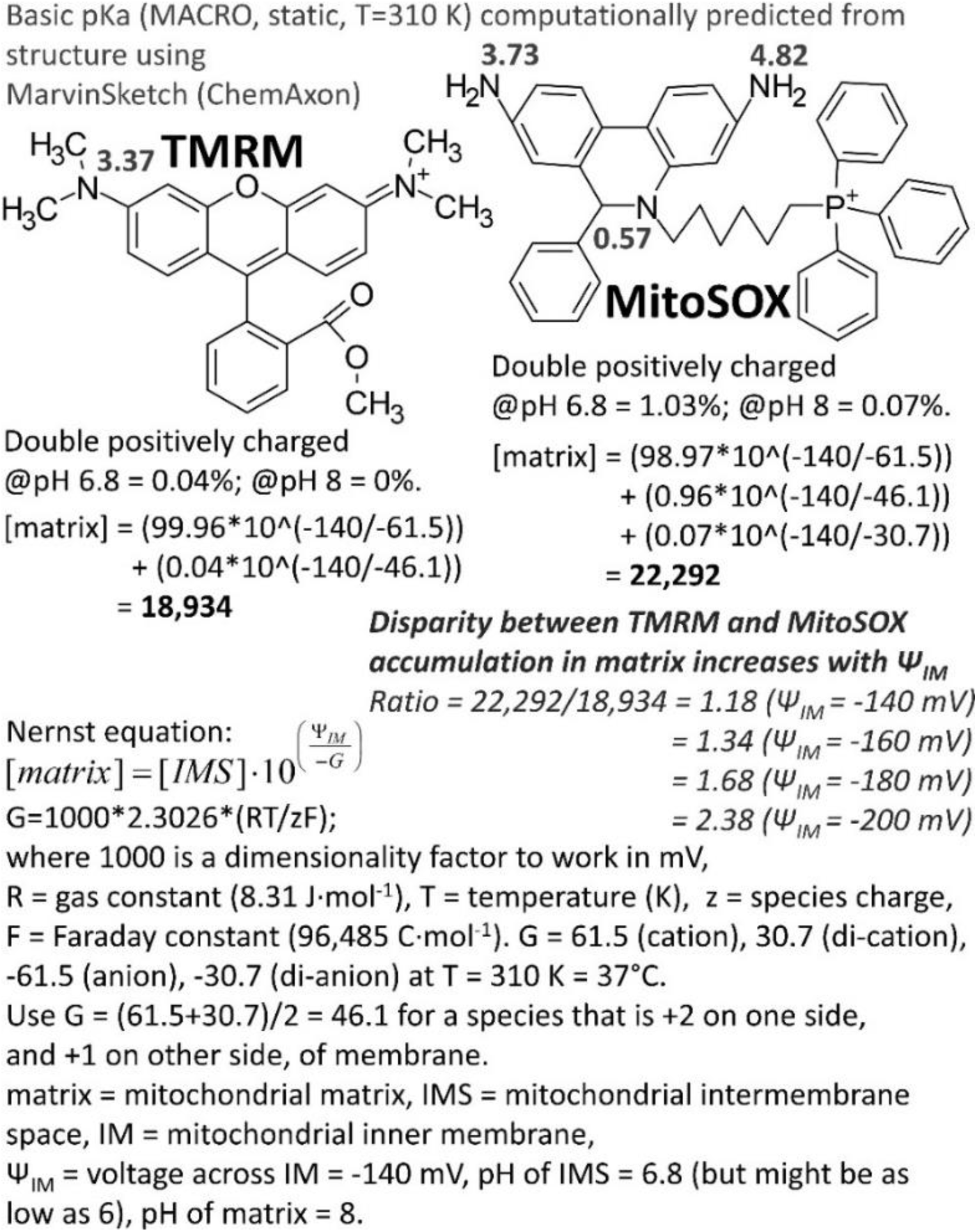

*References for voltage across IM* [80]*, pH of IMS* [81–82]*, and pH of mitochondrial matrix* [83]*. Constraint in the User Interface (UI) of MarvinSketch meant that pH 6.8, rather than 6.88 or 6.9* [81]*, was used for the IMS. But pH 6.8 is within the observed range, wherein* [81] *reports 6.88 ± 0.09*.

Figure 5 herein, using data from [77], shows

a. the disparity in TMRM accumulation (Ψ_IM_) between cells that have elevated IF1 protein and those that don’t; and
b. by my calculation, the predicted disparity in MitoSOX fluorescence signal that this disparity in Ψ_IM_ should cause; and
c. the observed disparity in MitoSOX fluorescence signal.

**Figure 5.**
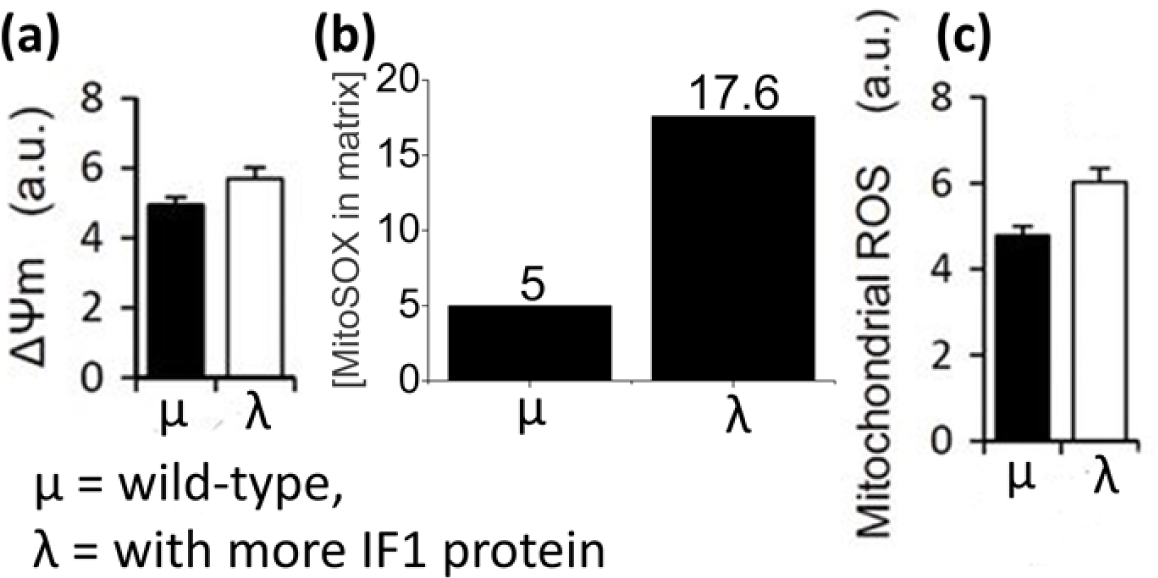
Compared to wild-type mouse cells (μ), transgenic mouse cells, with more IF1 protein (λ), have a lower Reactive Oxygen Species (ROS) concentration. **(a)** Experimental data [77]: membrane potential across the inner mitochondrial membrane, Ψ_IM_ (or Ψ_M_), as measured by TMRM accumulation. A typical value for Ψ_IM_ in normal mitochondria is -140 mV [80]. If we equate the 5 a.u. value for μ with -140 mV. Then the 6 a.u. value of λ is -168 mV. So, transgenic mouse cells, with more IF1 protein, have a more hyperpolarized membrane potential than wild-type cells. **(b)** Calculated data: predicted differential in MitoSOX accumulation in the mitochondrial matrix because of the different Ψ_IM_ values. **(c)** Experimental data [77]: Reported [ROS] by MitoSOX fluorescence. The MitoSOX fluorescence of transgenic mouse cells, with more IF1 protein, is 6 a.u. But wherein, because of their more hyperpolarized Ψ_IM_, the MitoSOX fluorescence is expected to be greater than this, at 17.6 a.u. The fact that they have less MitoSOX fluorescence than this expected amount shows that they must have less [ROS]: ({17.6-6}/17.6)*100 = 66%. So, [ROS] is 66% lower in transgenic mouse cells, with more IF1 protein, than wild-type.

Wherein the observed is 66% less than what is predicted by the disparity in Ψ_IM_. So, then the cells with elevated IF1 protein (and thence decreased F_1_F_0_ ATP hydrolysis) must have 66% lower [ROS].

#### Why don’t antioxidants work much?

Some past approaches in the literature have tried to decrease [ROS] by increasing ROS mitigation [84–86]. By administering an antioxidant compound(s). I suggest that the potential in this ROS mitigation approach is constrained. Because when administering an antioxidant compound into a biological system, there are inherently so many more biological molecules (some of which are incredibly large: e.g., DNA) than administered antioxidant molecules.

That a ROS is always more likely to collide with a biological molecule, causing damage, rather than collide with an exogenously introduced antioxidant molecule, and be mitigated. For an exogenous antioxidant to efficaciously outcompete at scale, for ROS collision, with the *huge* number of biological molecules, so much antioxidant compound would have to be administered that toxicity is likely.

Especially as at least some antioxidants, such as vitamin C (ascorbic acid, ascorbate), are pro-oxidants also [87], wherein their harmful pro-oxidant action may outweigh the benefit of their anti-oxidant action at higher concentrations (vitamin C mitigates ROS species by reduction [electron donation]. But this means that it can also reduce Fe^3+^ to Fe^2+^. Priming the Fenton reaction to produce, from H_2_O_2_, arguably the worst ROS species of all: the hydroxyl radical, OH. Fenton reaction: H_2_O_2_ + Fe^2+^→ ^•^OH + OH^-^ + Fe^3+^, thereafter OH^-^ collects a proton to become H_2_O).

In summary, ROS mitigation is a constrained/flawed approach. By contrast, the present paper proposes a different approach to decreasing [ROS]. By decreasing ROS generation. Where, as observed in the data herein (Figure 5), decreasing ROS generation can (unlike antioxidants) dramatically decrease [ROS].

[ROS] increases with body age [40–43]. Indeed, hair greying/whitening is reportedly because of increased [H_2_O_2_] (i.e., hair is bleached by the hydrogen peroxide) [88]. So, decreasing the [ROS] of an old animal, if only to the [ROS] of a young animal, is interesting. Moreover, keeping the [ROS] of a young animal at the same level throughout its life.

Any anti-aging utility notwithstanding, ROS have been implicated as contributory to many diseases, especially age-related diseases, and so a drug that can significantly reduce [ROS] might have utility for the prevention and treatment of all these different, varied diseases. For example, to mention just two (of many) diseases in which ROS have been implicated: Age-Related Macular Degeneration [89] and atherosclerosis [90].

#### IF1 protein does not inhibit F_1_F_0_ ATP synthesis, misreports that it does are hereby explained

[77–79] interpret increased IF1 protein amount decreasing respiration rate in mice as evidence that IF1 protein directly inhibits F_1_F_0_ ATP synthesis. This is misinterpreted. Newly revealed herein (*moreover in my corresponding patent applications, e.g.* [18–20]): in mice (especially; and in other mammals), substantial F_1_F_0_ ATP hydrolysis is occurring under normal conditions (Figure 2). Hydrolysing a fraction of the synthesized ATP, meaning more ATP needs to be synthesized. Which sets oxidative phosphorylation (OXPHOS) at high rate, to generate heat. Increased [IF1 protein] inhibits F_1_F_0_ ATP hydrolysis more, so less ATP needs to be made by F_1_F_0_ ATP synthesis, thence less OXPHOS is required, and less O_2_ is consumed. So, more IF1 protein *does* decrease F_1_F_0_ ATP synthesis. But this is by inhibiting F_1_F_0_ ATP hydrolysis. *<u>Not</u>* by inhibiting F_1_F_0_ ATP synthesis. This reinterpretation, of the data of [77–79], distinctively aligns with the interpretation of structural data by [10–13] (summarized in Figure 1 herein), which concludes that IF1 protein can only block F_1_F_0_ ATP hydrolysis, and *not* F_1_F_0_ ATP synthesis.

#### Equation for maximal lifespan in terms of specific F_1_F_0_ ATP hydrolysis rate

Recasting experimental data used earlier, Figure 6 shows (across different species) specific F_1_F_0_ ATP hydrolysis rate, specific basal metabolic rate, and heart rate for different values of maximal lifespan.

**Figure 6.**
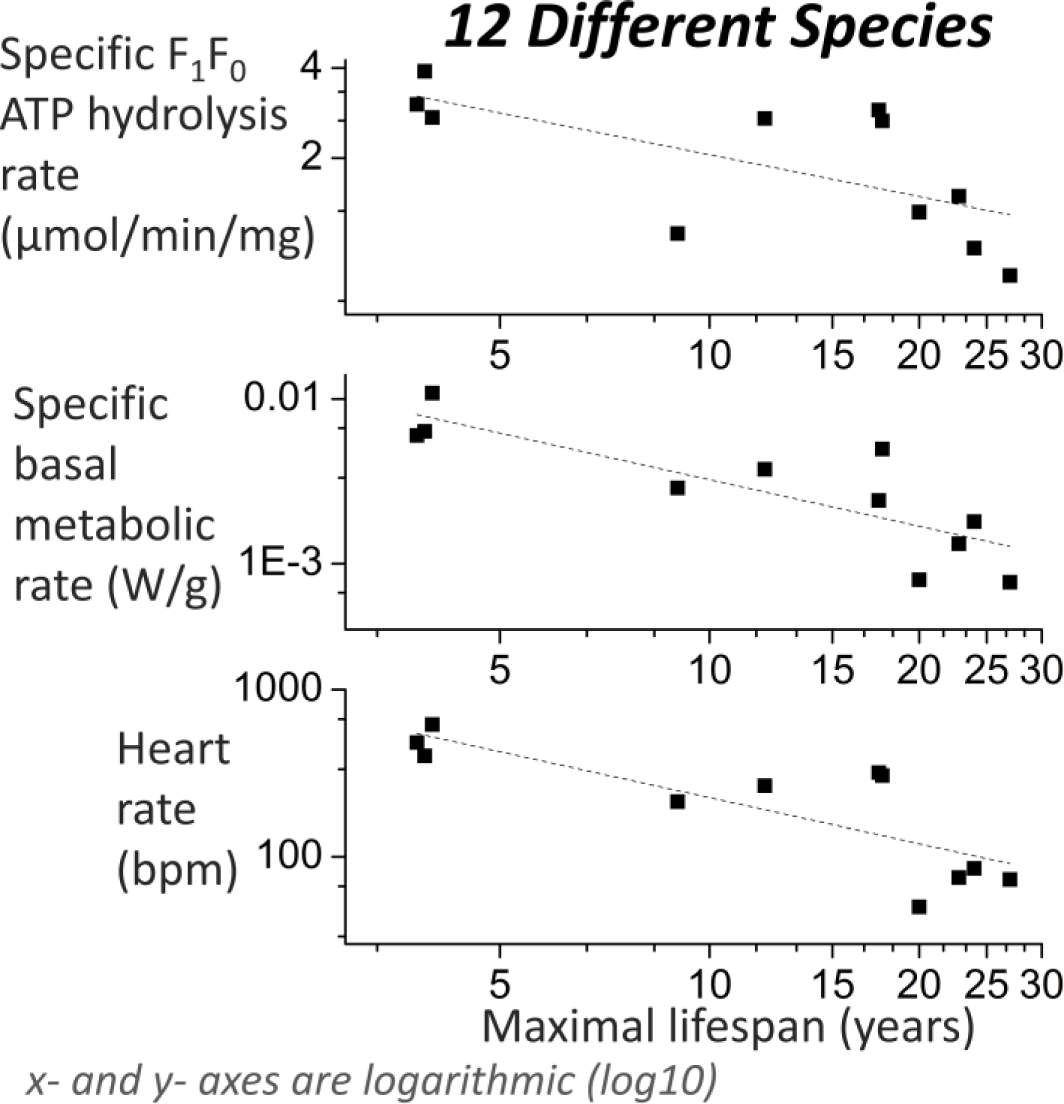
Across different species, specific F_1_F_0_ ATP hydrolysis rate, specific basal metabolic rate, and heart rate for different values of maximal lifespan.

Shown best-fit lines in Figure 6 are of the form:

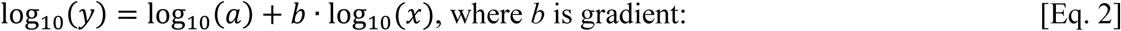

**Table 4.**
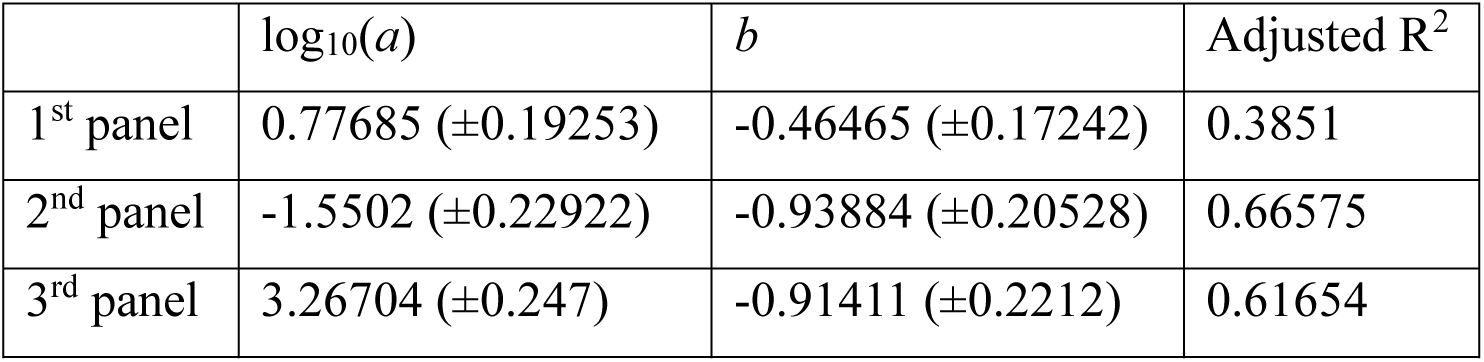

Rearranging the plot equation from the 1^st^ (top) panel in Figure 6, gives an equation for (at least across these species) maximal lifespan (*years*), *x*, in terms of specific F_1_F_0_ ATP hydrolysis rate (*µmol/min/mg*), *y*:

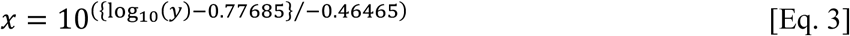

#### Locally, lower specific F_1_F_0_ ATP hydrolysis rate, predicted sufficient to slow aging locally by two-thirds, is safe in mice

As aforementioned, safely in mice, study [77] locally (*locally meaning that only some tissue(s) is affected, not all the body*) reduced F_1_F_0_ ATP hydrolysis rate by ∼35%. And study [79] by 35%, and study [78] by 44%.

Study [91] reports that specific F_1_F_0_ ATP hydrolysis rate for mice is 2.73 μmol/min/mg. 44% less is 1.5288 μmol/min/mg. Substituting this latter value into Eq. 3 outputs a maximal lifespan of 18.84 years. That is 3.5 times greater than 5.41 years, which it is with 2.73 μmol/min/mg. So, locally, the aging rate is (predicted to be) 1/3.5 (=0.29) of the normal aging rate: i.e., less than a third less. More than two-thirds slower. Safely.

At least safety is proven in mice. I expect that an even lower specific F_1_F_0_ ATP hydrolysis rate is safe, at least locally, but this is untested to date.

#### Mediation analysis reports that specific F_1_F_0_ ATP hydrolysis rate dictates maximal lifespan, via dictating specific basal metabolic rate

Mediation analysis is summarized in Figure 7. I performed mediation analysis in JASP software (version 0.14.1. https://jasp-stats.org/), which runs atop of R [92]. Selecting a bootstrap method called “Percentile” in JASP, which corresponds to “Percentile bootstrap (PC)” in [93]. Run with 10,000 replications (ten times greater than default).

**Figure 7.**
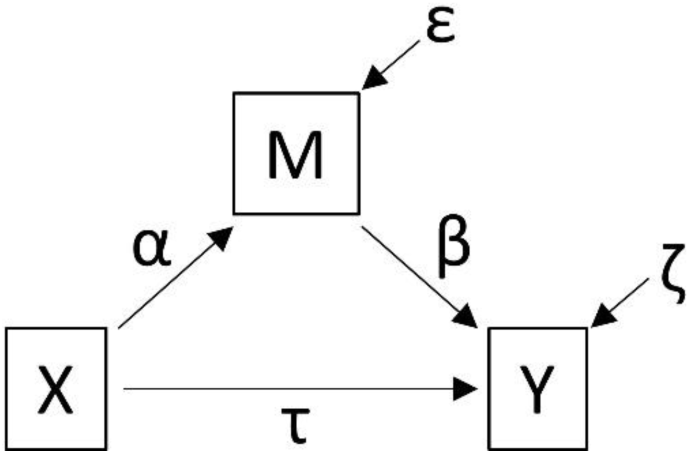
An indirect effect (mediation) exists when an independent variable (X) acts upon a dependent variable (Y) through an intervening/mediating variable (M). Represented by two regression equations [93]:

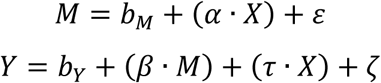

*b_M_* and *b_Y_* are regression intercepts. *ε* and *ζ* are random disturbance terms. *α*, *β* and *τ* are regression coefficients. **Indirect effect** (*effect of X upon Y, via M*) **= α·β**. Direct effect = τ. Total effect (*of X upon Y*) = (α·β) + τ.

For the data inputted, I multiplied all specific basal metabolic rate values by 10,000 (rescaled this variable) to prevent numerical underflow in the underlying computations performed by lavaan [94], an R package that JASP runs for mediation analysis. Needed because otherwise the very small numbers of specific basal metabolic rate in W/g causes even smaller numbers to occur in the internal computations, causing numerical underflow.

An alternative method to prevent numerical underflow is to select the option of “Standardized estimates” in JASP. To standardize (mean = 0, standard deviation = 1) all the variables before estimation (*independently for each variable: for each of its values: subtract value from this variable’s mean and then divide the output by this variable’s standard deviation*). But this method is (very slightly) inferior. Because it is manipulating each variable with quantities, the mean and standard deviation, which are *estimated*, with an error. Instead of manipulating a single variable with a constant, which is *precisely known* (10,000). Both methods produce consistent results. Those from using standardized estimates are presented afterwards, at the end of this section.

Using notation of Figure 7, Table 5 below reports results of mediation analysis in which *X* is specific F_1_F_0_ ATP hydrolysis rate, *M* is specific basal metabolic rate (*10,000), and *Y* is maximal lifespan (*data of these variables is as presented in Table S2 of the Supplementary Material*).

**Table 5.**

|  |  |  | 95% Confidence |  |
| --- | --- | --- | --- | --- |
|  |  |  | Interval (CI) |  |
| $\alpha$ | $\beta$ | Indirect effect ($\alpha \cdot \beta$) | Lower | Upper |
| 20 | -0.17 | -3.452 | -9.512 | -1.201 |
| | | Direct effect ($\tau$) | Lower | Upper |
|  |  | -2.554 | -6.266 | 4.124 |

Evidence for an effect is when zero (0) is not within the range of the 95% Confidence Interval (CI). The result communicated in Table 5 above reports that there is no direct effect between specific F_1_F_0_ ATP hydrolysis rate and maximal lifespan. That the effect of specific F_1_F_0_ ATP hydrolysis rate upon maximal lifespan is solely indirect, via specific basal metabolic rate.

Figure 8 presents the mediation that the data points to. Mediation data supporting its [specific F_1_F_0_ ATP hydrolysis rate → specific basal metabolic rate → maximal lifespan] arm has already been presented. Mediation data supporting its [specific F_1_F_0_ ATP hydrolysis rate → specific basal metabolic rate → heart rate] arm is in Table 6 below. From where *X* is specific F_1_F_0_ ATP hydrolysis rate, *M* is specific basal metabolic rate (*10,000), and *Y* is heart rate (*data of these variables is as presented in Table S2 of the Supplementary Material*).

**Figure 8.**
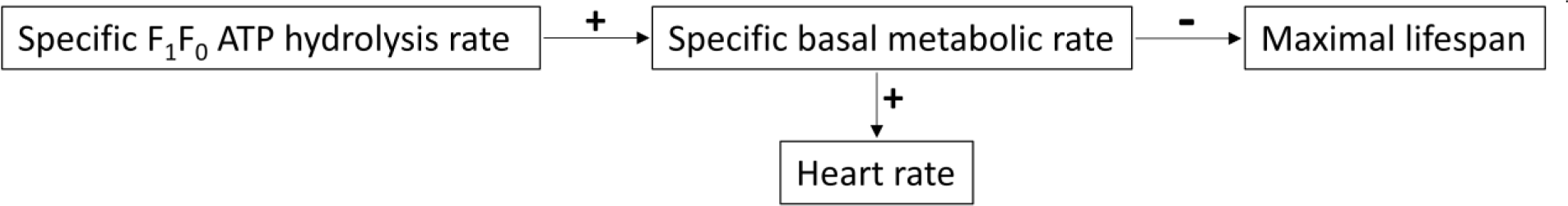
Interrelations reported by mediation analysis. Wherein specific F_1_F_0_ ATP hydrolysis rate dictates heart rate and maximal lifespan, via dictating specific basal metabolic rate.

**Table 6.**

|  |  |  | 95% Confidence |  |
| --- | --- | --- | --- | --- |
|  |  |  | Interval (CI) |  |
| $\alpha$ | $\beta$ | Indirect effect ($\alpha \cdot \beta$) | Lower | Upper |
| 20 | 4.6 | 94.363 | 22.578 | 191.3 |
| | | Direct effect ($\tau$) | Lower | Upper |
|  |  | 48.372 | -35.584 | 109.96 |

Table 7 below presents results from instead conducting the mediation analysis with standardized estimates. In this case, specific basal metabolic rate values are *not* multiplied by 10,000. Results are consistent with the earlier mediation analysis.

**Table 7.**

|  |  |  |  |  |
| --- | --- | --- | --- | --- |
|  |  |  | 95% Confidence |  |
|  |  |  | Interval (CI) |  |
| Specific F1F0 ATP hydrolysis rate → Specific basal metabolic rate → Maximal lifespan |  |  |  |  |
| α | β | Indirect effect (α·β) | Lower | Upper |
| 0.66 | -0.61 | -0.402 | -1.096 | -0.137 |
|  |  | Direct effect (τ) | Lower | Upper |
|  |  | -0.297 | -0.759 | 0.485 |
| Specific F1F0 ATP hydrolysis rate → Specific basal metabolic rate → Heart rate |  |  |  |  |
| α | β | Indirect effect (α·β) | Lower | Upper |
| 0.66 | 0.76 | 0.506 | 0.12 | 1.007 |
|  |  | Direct effect (τ) | Lower | Upper |
|  |  | 0.259 | -0.186 | 0.589 |

Table 8 below is as Table 7 above, but it presents results from using logged (log_10_) data variables. Results are consistent with the earlier mediation analysis.

**Table 8.**

|  |  |  |  |  |
| --- | --- | --- | --- | --- |
|  |  |  | 95% Confidence |  |
|  |  |  | Interval (CI) |  |
| Specific F1F0 ATP hydrolysis rate → Specific basal metabolic rate → Maximal lifespan |  |  |  |  |
| α | β | Indirect effect (α·β) | Lower | Upper |
| 3.4 | -0.82 | -2.761 | -4.878 | -0.575 |
|  |  | Direct effect (τ) | Lower | Upper |
|  |  | -0.102 | -2.744 | 1.901 |
| Specific F1F0 ATP hydrolysis rate → Specific basal metabolic rate → Heart rate |  |  |  |  |
| α | β | Indirect effect (α·β) | Lower | Upper |
| 3.4 | 0.78 | 2.649 | 0.936 | 4.857 |
|  |  | Direct effect (τ) | Lower | Upper |
|  |  | 0.902 | -0.717 | 2.799 |

#### Interventional experimental evidence for causality

Interventional mouse experiments report causality between lower F_1_F_0_ ATP hydrolysis rate and lower metabolic heat generation (lower metabolic rate; Figure 2), and causality between lower metabolic heat generation (lower metabolic rate) and longer lifespan [21–22].

#### Instrumental variable analysis reports causality between lower specific basal metabolic rate and higher maximal lifespan

Instrumental variable analysis can infer causality between an independent and a dependent variable, if the “instrumental variable” only impacts the dependent variable through its causal effect upon the independent variable [45]. Earlier presented mediation analysis (Table 5) reported that specific F_1_F_0_ ATP hydrolysis rate only affects maximal lifespan through specific basal metabolic rate, with *no* direct effect upon maximal lifespan.

R code in the Supplementary Material, modified from that of [45], using data from Table S2 of the Supplementary Material (human excluded by rationale in Methods), with maximal lifespan as the dependent variable, specific basal metabolic rate as the independent variable, and specific F_1_F_0_ ATP hydrolysis rate as the instrumental variable, reports causality between lower specific basal metabolic rate and higher maximal lifespan.

Results shown in Table 9 below. Coefficient for the effect of the independent variable on the dependent variable, *β*, is a Two-Stage Least Squares (TSLS) estimate (a type of *k*-class estimate). *E* is the standard error in *β*. *P* is the one-tailed (alternative hypothesis is directional) p-value for *β*. Causality can be inferred from it being significant (at *P* < 0.05).

**Table 9.**

| $\beta$ | E | P | A | C | A2 | C2 |
| --- | --- | --- | --- | --- | --- | --- |
| -2956.3935 | 860.<br>2659 | 0.003715 | 0.012622 | -9381.85990832096,<br>-1258.87418235506 | 0.012622 | -9381.85948762469,<br>-1258.87421171728 |

Incidentally, because there is only one independent variable, the results are the same (and observed to be the same, not shown) if we use the Limited Information Maximum Likelihood (LIML) estimate, which is a different type of *k*-class estimate, instead. *A* is the p-value for the Anderson-Rubin test (under F distribution), when significant (at *p* < 0.05) this rejects the null hypothesis of *β* = 0, and *C* is its 95% confidence interval. *A2* is the p-value for the conditional likelihood ratio test (under Normal approximation), when significant (at *p* < 0.05) this rejects the null hypothesis of *β* = 0, and *C2* is its 95% confidence interval.

*The R code used utilizes the ivmodel R package* [45]*. Same result was returned when using other R code, which is also in the Supplementary Material, that instead utilizes the AER R package. Same β, E, and P values (this other package doesn’t report A, C, A2, C2 values). Such consistency of results was observed with all instrumental variable analysis conducted herein*.

#### Instrumental variable analysis reports causality between lower specific F_1_F_0_ ATP hydrolysis rate and higher maximal lifespan

Instrumental variable analysis can infer causality [45]. IF1 protein is a selective inhibitor of F_1_F_0_ ATP hydrolysis (*doesn’t inhibit F_1_F_0_ ATP synthesis*) [8–14]. Instrumental variable analysis with maximal lifespan as the dependent variable, specific F_1_F_0_ ATP hydrolysis rate as the independent variable, and specific IF1 protein activity as the instrumental variable, reports causality between lower specific F_1_F_0_ ATP hydrolysis rate and higher maximal lifespan.

Results shown in Table 10 below (with table headings as in earlier Table 9). R code used is in the Supplementary Material, which uses data from Table S2 of the Supplementary Material, and specific IF1 protein activity data from [91] (human excluded by rationale in Methods).

**Table 10.**

| $\beta$ | E | P | A | C | A2 | C2 |
| --- | --- | --- | --- | --- | --- | --- |
| -9.6192 | 4.19<br>15 | 0.0237 | 0.039307 | (-infinity,<br>-1.44376813071093)<br>union<br>(21.3353360504571,<br>infinity) | 0.039307 | (-infinity,<br>-1.4437203851784)<br>union<br>(21.3346515903459,<br>infinity) |

#### Across species, higher specific F_1_F_0_ ATP hydrolysis rate correlates with higher specific Reactive Oxygen Species (ROS) generation rate

Table 4 below reports correlations between variables listed down its left-hand-side (*their data is presented in Table S2 of the Supplementary Material*) and superoxide (O_2_^•-^) and H_2_O_2_ detected (per unit mass per unit time) data from [95]. Where [95] is a review paper that consolidates data from different species and studies.

In Table 11 above, its lower section (for H_2_O_2_) has a different selection of capital letter headings than its upper section (for superoxide). Because it wasn’t possible to calculate columns H, I, J, K for superoxide, and columns A, E, F for H_2_O_2_, from the data available in [95] (or the primary papers it pulls data from).

**Table 11.**

| Pearson correlation coefficient (R) |  |  |  |  |  |  |  |  |
| --- | --- | --- | --- | --- | --- | --- | --- | --- |
|  | [SUPEROXIDE] DETECTED (nmol superoxide/minute/mg of protein) |  |  |  |  |  |  |  |
|  | A | B | C | D | E | F |  |  |
| Specific $F_1F_0$ ATP hydrolysis rate | 0.7711 | 0.6714 | 0.7724 | 0.7815 | 0.7995 | 0.7937 | | |
| Specific basal metabolic rate | 0.7489 | 0.9415 | 0.8734 | 0.9449 | 0.7904 | 0.7891 |  |  |
| Heart rate | 0.8573 | 0.9667 | 0.8327 | 0.9241 | 0.786 | 0.779 |  |  |
| Maximal lifespan | -0.7362 | -0.7372 | -0.6732 | -0.7322 | -0.7114 | -0.6662 |  |  |
| | [H2O2] DETECTED (nmol $H_2O_2$/minute/mg of protein) | | | | | | | |
|  | B | C | D | G | H | I | J | K |
| Specific $F_1F_0$ ATP hydrolysis rate | 0.865 | 0.8675 | 0.8226 | 0.712 | 0.7665 | 0.719 | 0.703 | 0.7945 |
| Specific basal metabolic rate | 0.8288 | 0.525 | 0.6127 | 0.5542 | 0.6065 | 0.5804 | 0.6945 | 0.6866 |
| Heart rate | 0.813 | 0.5486 | 0.627 | 0.5808 | 0.7646 | 0.621 | 0.6553 | 0.7057 |
| Maximal lifespan | -0.8191 | -0.7059 | -0.7755 | -0.7878 | -0.8073 | -0.8246 | -0.6642 | -0.7533 |

Table 12 below is a key to Table 11 above. Its capital letters correspond to columns of Table 11, and it refers to primary source papers as they are referred to in tables 1 and 2 of [95]. Its *n* is the number of species that overlap with the species that I have specific F_1_F_0_ ATP hydrolysis rate data for (from [91]), which is the number of species used to generate the Pearson correlation coefficients in that capital letter headed column of Table 11. The highest value of *n* is 8, wherein this consists of cow, pig, rabbit, pigeon, guinea pig, rat, hamster, and mouse. The data in [95] from “Sohal et al. 1993b” (as [95] refers to it) was excluded. Because its values are over 1,000 times different from the values of the other studies. That primary paper aside, all the other ROS data in [95], for species I have specific F_1_F_0_ ATP hydrolysis rate data for (from [91]), was included to make Table 11 above. If a primary source paper’s data, sourced from [95], had more than 4 species overlapping with the species I have specific F_1_F_0_ ATP hydrolysis rate data for, Pearson correlation coefficients were calculated. For primary source papers with 4 or less species of overlap, their data is only incorporated into the means. In total, via [95], ROS data from 7 different primary papers is used, from 3 different organs (liver, kidney, heart), from 3 different research groups, from studies separated by up to 18 years (by publication date).

**Table 12.**
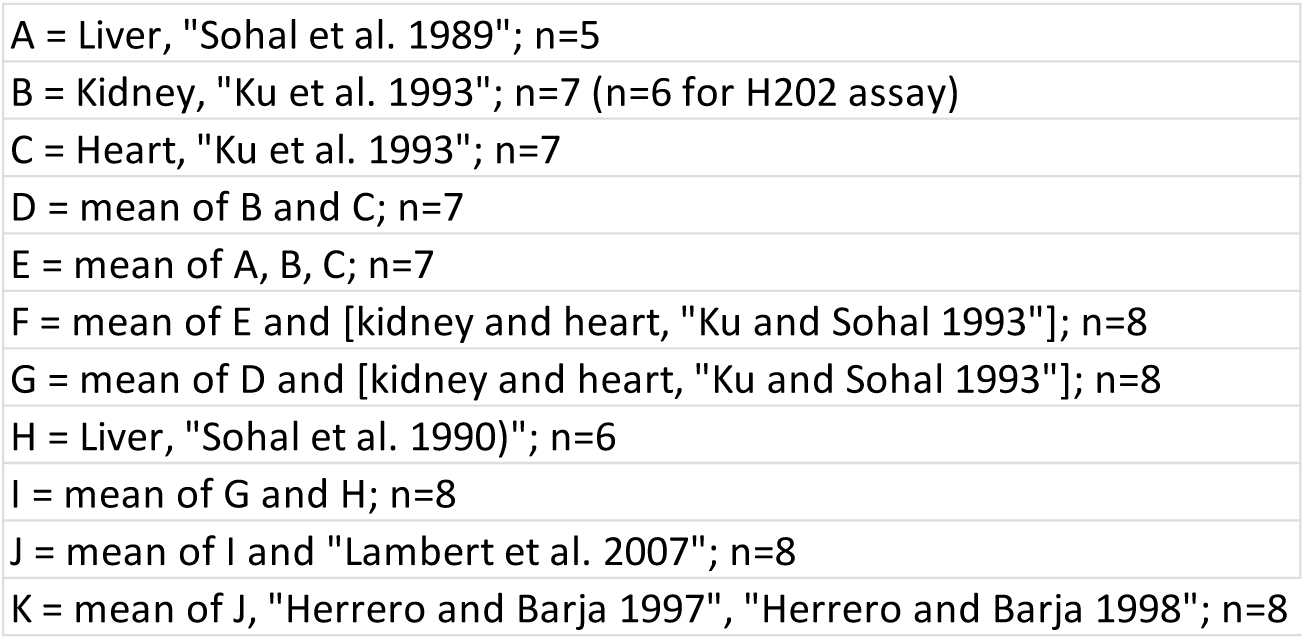

In Table 11, its Column F reports Pearson correlation coefficients using mean of *all* the superoxide data, across all the used studies, and its Column K reports Pearson correlation coefficients using mean of *all* the H_2_O_2_ data, across all the used studies. Table 13 below reports p-values for these columns (one-tailed, because each alternative hypothesis is directional). Which are small and statistically significant, despite the small value of *n* (8) for each, as testament to the high *R* values.

**Table 13.**

| p-value (one-tailed) |  |  |
| --- | --- | --- |
|  | Superoxide | H2O2 |
|  | F | K |
| Specific $F_1F_0$ ATP hydrolysis rate | 0.009347 | 0.0092445 |
| Specific basal metabolic rate | 0.009949 | 0.0300005 |
| Heart rate | 0.011355 | 0.0252435 |
| Maximal lifespan | 0.035629 | 0.015467 |

[95] only reports, and so earlier Table 11 only uses, data from isolated mitochondria and SMPs thereof. What is the situation in a more complete system? In whole cells respiring glucose? [43] shows that, in *ex vivo* brain slices respiring glucose, ROS (superoxide) production increases with age. And that the gradient of this increase is steeper in shorter living species. “The rate of age-related increases of superoxide dependent chemiluminescence was inversely related to the maximum lifespan of the animals” [43]. At least for the species they looked at. Wherein there weren’t enough shared species to calculate a Pearson correlation coefficient with data herein unfortunately.

#### Across species, higher specific F_1_F_0_ ATP hydrolysis rate correlates with more specific Reactive Oxygen Species (ROS) damage

Table 14 below reports correlations between variables along its top (*their data is presented in Table S2 of the Supplementary Material*) and data from [96]. Showing Pearson correlation (R) coefficients and p-values (one-tailed, because each alternative hypothesis is directional).

**Table 14.**

| | | Specific $F_1F_0$ ATP<br>hydrolysis rate | Specific basal<br>metabolic rate | Heart rate | Maximal<br>lifespan | Body mass |
| --- | --- | --- | --- | --- | --- | --- |
| R | **8-OxoGua** | **0.8455** | **0.9019** | **0.9488** | **-0.972** | **-0.8901** |
| p |  | 0.0169805 | 0.0069815 | 0.001933 | 0.002801 | 0.0087265 |
| R | **8-OxodG** | **0.838** | **0.9955** | **0.9626** | **-0.7675** | **-0.6149** |
| p |  | 0.01862 | 0.000015 | 0.001036 | 0.064891 | 0.0969485 |
| R | **5-HMUra** | **0.7053** | **0.9545** | **0.8704** | **-0.6204** | **-0.6532** |
| p |  | 0.0587375 | 0.001529 | 0.012053 | 0.13209 | 0.0797755 |
| R | **MEAN** | **0.8223** | **0.9779** | **0.9609** | **-0.8453** | **-0.8057** |
| p |  | 0.02228 | 0.0003635 | 0.001132 | 0.035661 | 0.0264805 |
| *Unit: nmol/mmol creatinine* |  |  |  |  |  |  |

[96] reports, across species, upon urinary excretion of repair products of oxidative DNA damage: 8-oxo-7,8-dihydroguanine (8-oxoGua), 8-oxo-7,8-dihydro-2’-deoxyguanosine (8-oxodG), and 5-(hydroxymethyl)uracil (5-HMUra). More such DNA repair products reflects more DNA repair, and thereby more ROS damage of DNA. Disparity in DNA amount in different species (because of different body sizes and therefore cell number) was washed out of the data by dividing by creatinine concentration in the urine.

*Oxidative damaged DNA increases with age* [97–98]*. In* [96]*, the “biological age” of the individuals from the different species was constant (around 20% of maximal lifespan for each species) except for pig (4% of maximal lifespan)*.

Table 15 below reports correlations between variables along its top (*their data is presented in Table S2 of the Supplementary Material*) and data from [99]. Showing Pearson correlation (R) coefficients and p-values (one-tailed because each alternative hypothesis is directional). [99] reports, across species, upon amount of a ROS damage product, 8-oxo-7,8-dihydro-2’-deoxyguanosine (8-oxodG) (standardized to the amount of deoxyguanosine, dG; per 10^5^ dG), in mitochondrial DNA (mtDNA).

**Table 15.**

| | 8-oxodG/$10^5$dG | Specific F1F0 ATP | Specific basal | | Maximal | |
| --- | --- | --- | --- | --- | --- | --- |
|  | in mtDNA | hydrolysis rate | metabolic rate | Heart rate | lifespan | Body mass |
| R **Brain** |  | **0.885** | **0.8837** | **0.9246** | **-0.7369** | **-0.4484** |
| p |  | 0.022999 | 0.0233855 | 0.0122855 | 0.077724 | 0.224417 |
| R **Heart** |  | **0.7257** | **0.9509** | **0.9406** | **-0.7438** | **-0.6871** |
| p |  | 0.0324335 | 0.0004995 | 0.0007995 | 0.027638 | 0.0440525 |
| R **MEAN** |  | **0.8212** | **0.9738** | **0.9867** | **-0.8533** | **-0.6689** |
| p |  | 0.044142 | 0.0025355 | 0.000919 | 0.032973 | 0.1084925 |

*In Table 15 above, the mean only includes species for which both brain and heart data is in the data set. Data was extracted from graphs in* [99] *using WebPlotDigitizer,* https://apps.automeris.io/wpd/. [99] *optimized more for a constant chronological, rather than “biological”, age. All the individuals, from the different species, were around one to two years old: “the mean age of the animals was 8 months (mice), 11 months (rat), 1.4 years (guinea pig), 1.5 years (rabbit), 1 year (pig), and 1.5–2.5 years (sheep, cow, and horse)”* [99].

Table 16 below reports correlations between variables along its top (*their data is presented in Table S2 of the Supplementary Material*) and data from [100] (*data sourced from its supplementary table 4*). Showing Pearson correlation (R) coefficients and p-values (one-tailed because each alternative hypothesis is directional). [100] reports, across species, DNA substitution (SBS1, SBSB, and SBSC types), indel (somatic insertions and deletions), and mtDNA mutation rates per year. Per genome or, in the latter case, per mtDNA copy number. [Table 16]

**Table 16.**

|  | Specific F1F0 ATP | Specific basal |  | Maximal |  |
| --- | --- | --- | --- | --- | --- |
|  | hydrolysis rate | metabolic rate | Heart rate | lifespan | Body mass |
| R **DNA substitution rate per year** | **0.9155** | **0.9332** | **0.9595** | **-0.7817** | **-0.3861** |
| p | 0.0052045 | 0.003272 | 0.0012135 | 0.0591735 | 0.2248145 |
| R **Indel rate per year** | **0.9109** | **0.9569** | **0.9667** | **-0.7511** | **-0.4532** |
| p | 0.0057775 | 0.001373 | 0.0008225 | 0.071684 | 0.1833705 |
| R **mtDNA mutation rate per year** | **0.906** | **0.9619** | **0.9592** | **-0.7238** | **-0.4272** |
| p | 0.0064195 | 0.001075 | 0.0012315 | 0.0834205 | 0.199091 |
| *N.B. Substitution rate per year = SBS1 + SBSB + SBSC rate per year* |  |  |  |  |  |

Table 17 below reports correlations between variables along its top (*their data is presented in Table S2 of the Supplementary Material*) and data from [101]. Showing Pearson correlation (R) coefficients and p-values (one-tailed because each alternative hypothesis is directional). [101] reports, across species, amount of lipid peroxidation product malondialdehyde (MDA) per unit tissue mass.

**Table 17.**

|  | Specific F1F0 ATP | Specific basal |  | Maximal |  |
| --- | --- | --- | --- | --- | --- |
|  | hydrolysis rate | metabolic rate | Heart rate | lifespan | Body mass |
| LP R | **0.8122** | **0.9678** | **0.9291** | **-0.7594** | **-0.3096** |
| p | 0.0132285 | 0.0001755 | 0.001237 | 0.02384 | 0.249615 |
| LP = in vivo Lipid Peroxidation (mmol MDA/g tissue) |  |  |  |  |  |

*Data was extracted from Fig. 5 in* [101] *using WebPlotDigitizer. “All animals used were young adults with an age within 15–30% of their MLSP”* [101]*, wherein MLSP is maximum lifespan*.

Table 18 below reports correlations between variables along its top (*their data is presented in Table S2 of the Supplementary Material*) and data from [102]. Showing Pearson correlation (R) coefficients and p-values (one-tailed because each alternative hypothesis is directional). [102] reports, across species, protein degradation rate constants (K_deg_) for some different proteins, the median of which was taken for each species. K_deg_ data was obtained by personal communication with the first author of [102] (ages of the animals used were unknown, except for mouse [6 months old female]).

**Table 18.**

|  | Specific F1F0 ATP | Specific basal |  | Maximal |  |
| --- | --- | --- | --- | --- | --- |
|  | hydrolysis rate | metabolic rate | Heart rate | lifespan | Body mass |
| R **Median Kdeg** | **0.9465** | **0.8369** | **0.9065** | **-0.9884** | **-0.7601** |
| p | 0.0021085 | 0.0188665 | 0.0063525 | 0.0007485 | 0.0397125 |

So, shorter living species have faster protein turnover (greater median K_deg_). I interpret this is because, as a function of greater specific F_1_F_0_ ATP hydrolysis rate (driving greater specific basal metabolic rate), they have greater intracellular [ROS], and so a greater rate of ROS damage of proteins, and so their rate of protein replacement needs to be greater.

Compounding because protein replacement itself requires ATP energy, requiring ATP synthesis, (if ATP is synthesized aerobically) generating ROS, increasing the requirement for protein replacement.

To summarize this headed section, at least across these species, statistically significant correlation has been shown between specific F_1_F_0_ ATP hydrolysis rate and damage to DNA, mtDNA, lipids, and (by inference) proteins.

#### Does F_1_F_0_ ATP hydrolysis (at least partially) turn the hands of the epigenetic clock?

Epigenetic clocks are reviewed in [103]. Carbonyl cyanide m-chlorophenylhydrazone (CCCP), an uncoupler drug, which is well known to increase metabolic rate, increases the “ticking” rate of the epigenetic clock [104]. Symmetrically, I predict a drug that decreases metabolic rate (*e.g., a drug that selectively inhibits F_1_F_0_ ATP hydrolysis, which doesn’t inhibit F_1_F_0_ ATP synthesis* [18–20]) can slow the ticking rate of the epigenetic clock.

Below is a series of equations, implicit from “Clock 1” in [105] (note that “log” in Clock 1 of [105] refers to the natural logarithm, ln), which relates *DNAmAge* (DNA methylation age, also known as epigenetic clock age) to chronological age, for mammal species.

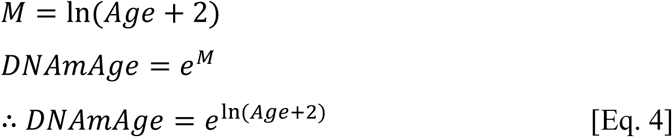

Where *Age* is chronological age, *DNAmAge* is predicted age, *M* is a linear combination of DNAm (=β) values for selected DNA cytosines, and the +2 offset is to avoid negative numbers with any prenatal data.

The natural logarithm (ln) is required because *M* changes (“ticks”) slower in mammal species with a higher maximal lifespan. Indeed, [106] reports the rate of epigenetic ticking is inversely proportional to maximal lifespan across 26 bat species. [107] reports the rate of epigenetic ticking is inversely proportional to maximal lifespan across 3 species (mouse, monkey, human). Moreover, [108] reports, using a 6 mammal species set, slower epigenetic ticking in longer-lived species. And (using a transchromosomic mouse strain, *Tc1*, that harbours a largely intact functional human chromosome 21) shows much faster epigenetic ticking upon a human chromosome when it is in mice (instead of human) cells.

Why this slower ticking of the epigenetic clock in longer-living mammals? What even causes the ticking of the epigenetic clock? I hypothesize that Reactive Oxygen Species (ROS) cause, at least partially, directly and/or indirectly, the ticking of the epigenetic clock. And, as taught herein, shorter-living mammals have greater specific F_1_F_0_ ATP hydrolysis rate, greater specific basal metabolic rate [so faster heart rate], and greater specific ROS generation rate.

In support, if cells *in vitro* are cultured in a lower O_2_ concentration, their epigenetic clock ticks slower [109] ([ROS] in cells inherently being a function of [O_2_] in cells). Calorie restriction, which can reduce ROS [110], slows the epigenetic clock in mice [107]. Men have a greater specific basal/resting metabolic rate than women [29, 111–114] and their epigenetic clock ticks faster [115]. Metabolism slows, and the epigenetic clock ticks slower, during hibernation [116]. ROS cause DNA methylation changes by a variety of mechanisms, reviewed in [117–118]. For example, 5-methyl-cytosine (5mC) can be oxidized by the hydroxyl radical (a ROS) to 5-hydroxymethyl-cytosine (5hmC), which when repaired results in demethylation [119–120]. At least mechanistically, it has been reported how superoxide (a ROS) could promote DNA and histone methylation [121]. A theory is that epigenetic change is driven by DNA damage [122], it adamant that ROS aren’t involved in aging, but it overlooks that ROS are a cause of DNA damage: e.g., DNA double-strand breaks [123–124]. Indeed, there is experimental data showing that ROS-caused DNA damage can cause epigenetic changes [125]. So, ROS may at least cause epigenetic changes indirectly, by causing DNA damage. And possibly directly as well.

The degree to which a length of DNA is wound up in chromatin presumably dictates ROS access to it. Different cell-types, as a function of different patterns of gene expression, tend to have different parts of the genome unwound to different extents. Possibly (at least partially) explaining why some epigenetic clocks (which use a subset of DNA cytosines) are tissue specific. Whilst pan-tissue (e.g. [126]; and/or pan-species, e.g. [105]) epigenetic clocks, which use different subsets of DNA cytosines, might be solely/predominantly using DNA stretches that tend to be reasonably unwound across multiple cell-types (and/or species). When a randomly damaging agent has non-random access to a damageable substrate then the ensuing pattern of damage is non-random. Preeminent leaders in the field interpret, I think misinterpret, the non-randomness of epigenetic changes as aging being programmed rather than damage [46, 105].

#### Equations interrelating epigenetic clock ticking rate (*ΔM*) and maximal lifespan

*ΔM* is the average change in (aforementioned) variable *M* per year in a mammal species:

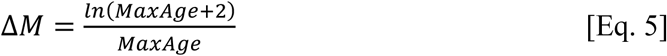

Where *MaxAge* is the maximal lifespan (in years) of the species. What this new equation does, in its numerator, is calculate the maximal value of *M* in this species: i.e., at its maximal lifespan. And then divide that by the number of years to get to that point (the maximal lifespan), to get the average change in *M* per year in that species. As earlier, a +2 offset is used. In an equation variant, not used herein, this offset is omitted, which has equivalent utility for postnatal data:

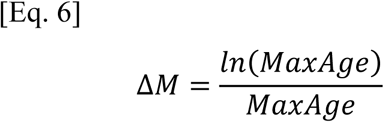

Clock 1 of [105] was fitted to data from many more eutherian species than marsupial and monotreme species and so I (arguably overly cautiously) restrict to using eutherian species data here (*N.B., in the first version of* [105] *on bioRxiv, its Clock 1 was only fitted to eutherian species data*). For eutherian species probed by the mammalian methylation array [127], the array used by [105], those 112 species thereof whose maximal lifespan is available in [1], including human, the Pearson correlation coefficient (R) between *ΔM* (calculated by Eq. 5) and maximal lifespan (values sourced from [1]) is -0.7388. With a one-tailed (alternative hypothesis is directional) p-value of 7.14744E-21 (*if human omitted: R = - 0.7775, one-tailed p-value = 5.38374E-24*). Corresponding Spearman’s rank correlation coefficient (*with human included or excluded*) = -1, one-tailed p-value = 0.

So, the greater the maximal lifespan of a eutherian species, the slower its epigenetic clock ticks. Form of this relationship is shown in Figure 9 (includes human). With the caveat that these values of *ΔM* are calculated from an equation (Eq. 5) discovered implicit to a fitted function to experimental data, rather than being directly experimental data. The fitted function being “Clock 1” in, and the experimental data being from, [105]. I first published this discovered relation in September 2022 in a bioRxiv preprint (.v3) precursor to the present paper [52], which has since been later reported by others.

**Figure 9.**
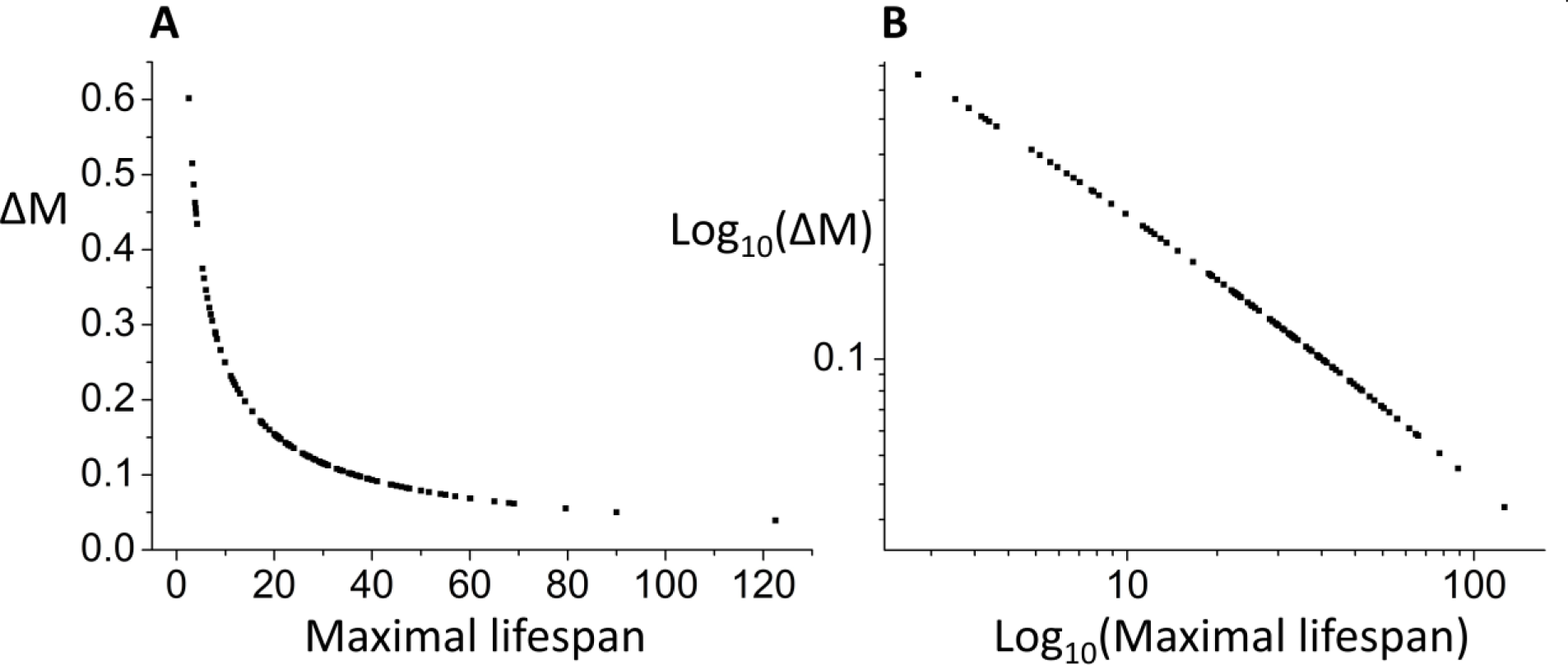
**(A)** Across 112 eutherian species (different points are for different species): average epigenetic change per year (*ΔM*) is inversely proportional to maximal lifespan. **(B)** Log_10_-log_10_ plot of the data of (A).

For eutherians, a predictive equation to calculate maximal lifespan from *ΔM* is below. Derived by log_10_-log_10_ plotting maximal lifespan (as *y*) versus *ΔM* (as *x*), using the same data from 112 eutherian species used to generate Figure 9, which includes human, which is a linear plot, and finding its best-fit line equation using OriginPro 8 software (OriginLab Corporation, Northampton, MA, USA. Using its “Fit Linear” functionality, which uses linear regression, with least squares estimation. Adjusted R^2^ = 0.99855):

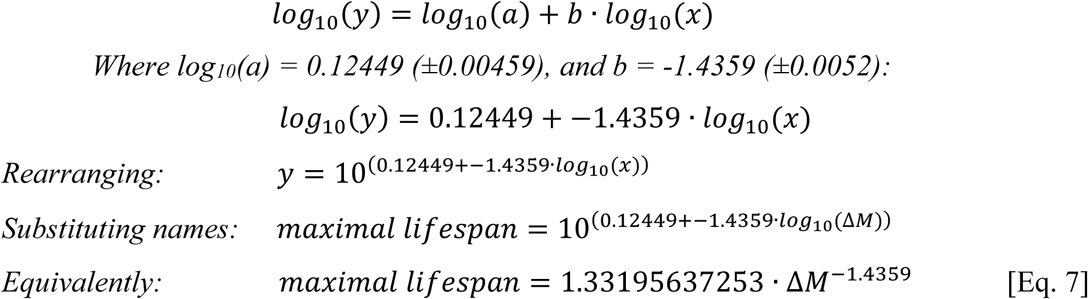

Because of the, when unlogged, non-linearity (negative convex shaped plot) between maximal lifespan and *ΔM* (which can be seen with *ΔM* on the *y*-axis, as in Figure 9, or on the *x*-axis instead, not shown) a given absolute decrease in *ΔM* can correspond with a very different absolute increase in maximal lifespan depending on what the value of *ΔM* is, which this new equation captures: an absolute decrease amount in *ΔM* corresponds with a greater absolute increase amount in maximal lifespan the smaller that *ΔM* is. Although this equation was derived from interspecies data it might have utility for intraspecies use. For example, to predict a new higher maximal lifespan if *ΔM* is successfully reduced to a lower value by a drug intervention. This equation suggests that a certain absolute decrease in *ΔM* corresponds with a greater increase in maximal lifespan in a human than in a mouse.

A different equation for maximal lifespan from *ΔM* comes from rearranging Eq. 5, where *W* is the Lambert *W* function:

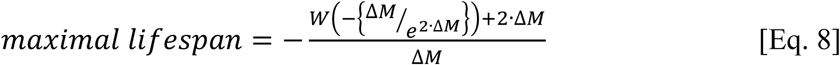

Rearranging Eq. 6 gives the simpler: [Eq. 9]:

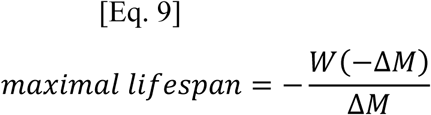

#### Specific F_1_F_0_ ATP hydrolysis rate dictates epigenetic clock ticking rate (*ΔM*), via dictating specific basal metabolic rate

Table 19 below correlates *ΔM* to the variables listed along its top (*data of which, except surface-area per unit body mass, is presented in Table S2 of the Supplementary Material. Omitting birds here, restricting to mammals because the ΔM equation was derived solely from mammal data, and omitting human by rationale in the Methods section. Surface-area estimated from body mass by the Meeh equation with k = 0.1 m^2^/kg^0.67^ and b = 0.67* [128]*. Then divided by body mass to calculate body surface-area per unit body mass*). P-values are one-tailed, because each alternative hypothesis is directional.

**Table 19.**

|  | Surface-area per | Specific F1F0 ATP | Specific basal |  | Maximal |  |
| --- | --- | --- | --- | --- | --- | --- |
|  | unit body mass | hydrolysis rate | metabolic rate | Heart rate | lifespan | Body mass |
| R $\Delta M$ | 0.8829 | 0.8476 | 0.8908 | 0.944 | -0.943 | -0.4246 |
| p | 0.0008055 | 0.0019535 | 0.0006355 | 0.000065 | 0.000069 | 0.1273265 |

Note how weakly $\Delta M$ (negatively) correlates with body mass as compared to how strongly (positively or negatively) it correlates with the other variables shown. For example, to surface-area per unit body mass. And specific basal metabolic rate.

Note how weakly *ΔM* (negatively) correlates with body mass as compared to how strongly (positively or negatively) it correlates with the other variables shown. For example, to surface-area per unit body mass. And specific basal metabolic rate.

When the surface-area per unit body mass variable is populated with values from [128] that use experimentally determined, rather than (Meeh equation) calculated, body surface-area values, then its Pearson correlation coefficient (R) to *ΔM* is 0.8328 (*sample size, n, is one less than used to make Table 19, because* [128] *doesn’t have data for hamster, one-tailed p-value = 0.0051345*).

Using data used to produce Table 19 above, the results of mediation analysis between *ΔM* (as the dependent variable) and specific F_1_F_0_ ATP hydrolysis rate, with the Mediator variable being specific basal metabolic rate, are in Table 20 below. On its first line of data. On its second line of data, are results of mediation analysis between maximal lifespan (as the dependent variable) and specific basal metabolic rate, with the Mediator variable being *ΔM*. Conducted in JASP software (version 0.14.1). When inputting this data into JASP, I multiplied specific basal metabolic rate values (W/g) by 10,000 and values of *ΔM* by 100 (rescaled these variables) to prevent numerical underflow in the underlying computations performed. Bootstrap method, called “Percentile” in JASP, was used with 10,000 replications.

**Table 20.**
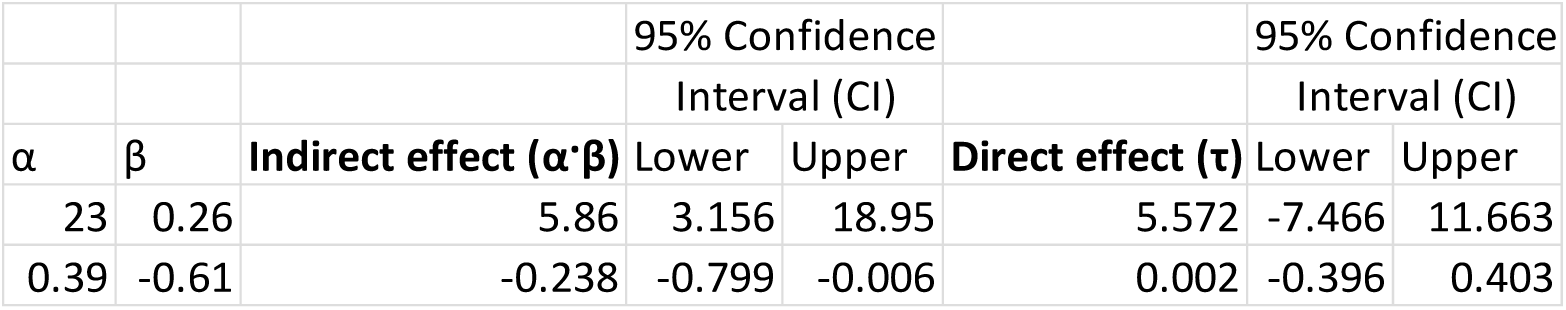

In both cases mediation is shown. Because, for both, there *is* a zero (0) within the 95% Confidence Interval (CI) for a direct effect and *no* zero within the 95% CI for an indirect effect. So, this data is consistent with the following mediation cascade:

Specific F_1_F_0_ ATP hydrolysis rate → Specific basal metabolic rate → *ΔM* → Maximal lifespan. Where all arrows signify positive effect, except the last, which signifies negative effect. This result is consistent with the ticking of the epigenetic clock being at least a partial drive of aging.

#### Instrumental variable analysis reports causality between specific basal metabolic rate and the ticking rate of the epigenetic clock (*ΔM*)

Instrumental variable analysis can infer causality between an independent and a dependent variable, if the “instrumental variable” only impacts the dependent variable through its causal effect upon the independent variable [45]. Earlier presented mediation analysis (Table 20) reported that specific F_1_F_0_ ATP hydrolysis rate only affects *ΔM* through specific basal metabolic rate, with *no* direct effect upon *ΔM*.

Using data used to generate earlier Table 20, with *ΔM* as the dependent variable, specific basal metabolic rate as the independent variable, and specific F_1_F_0_ ATP hydrolysis rate as the instrumental variable, using R code that does instrumental variable analysis (code modified from that of [45] and shown in the Supplementary Material), causality is reported between specific basal metabolic rate and *ΔM.* That specific basal metabolic rate is a causal drive of *ΔM*.

Result shown in Table 21 below, with column headings as specified earlier for Table 9.

**Table 21.**
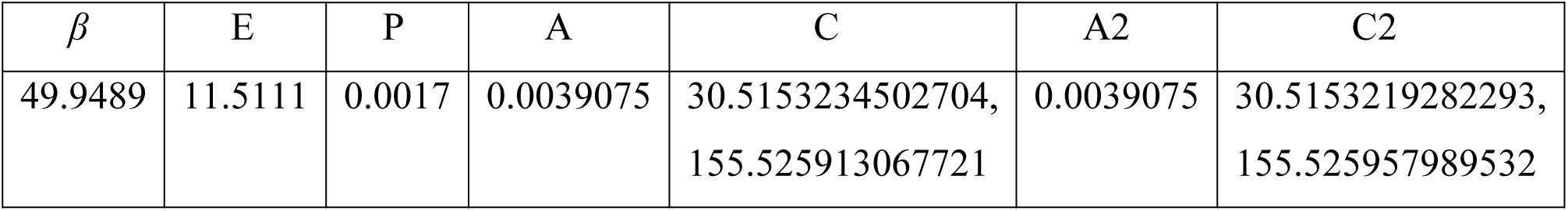

| $\beta$ | E | P | A | C | A2 | C2 |
| --- | --- | --- | --- | --- | --- | --- |
| 49.9489 | 11.5111 | 0.0017 | 0.0039075 | 30.5153234502704,<br>155.525913067721 | 0.0039075 | 30.5153219282293,<br>155.525957989532 |

#### Epigenetic clock ticking rate (*ΔM*) is linear function of specific basal metabolic rate

Using the data used to produce Figure 9, delimited to data from those 51 species thereof that have a specific basal metabolic rate value available in [1], the Pearson correlation coefficient (R) between *ΔM* and specific basal metabolic rate is 0.6669, one-tailed (alternative hypothesis is directional) p-value = 4.56278E-08 *(if human omitted: R = 0.6639)*.

For these 51 species, when log_10_ specific basal metabolic rate is plotted against log_10_ adult body mass, the specific basal metabolic rate value for *Sorex araneus* (common shrew) is anomalously high (not shown). Probably because it is not a true specific basal metabolic rate value, as it might not be possible to get such a value for shrews (family of Soricidae) because they become hyperactive when postabsorptive [129]. So, postabsorptive and inactive, both requirements for a basal metabolic rate value, might be mutually exclusive in shrews.

When *Sorex araneus* is excluded, rendering a 50 species set, the correlation is better: Pearson correlation coefficient between *ΔM* and specific basal metabolic rate = 0.7269, one-tailed (alternative hypothesis is directional) p-value = 1.13693E-09 *(if human omitted: R = 0.7202)*.

*ΔM* correlates with specific basal metabolic rate much better than it (inversely) correlates with adult body mass. For eutherian species probed by the mammalian methylation array [127] (array used by [105]), those 111 species thereof whose maximal lifespan (required for calculation of *ΔM*) and adult body masses are available in [1]: Pearson correlation coefficient between *ΔM* and adult body mass = -0.237, one-tailed (alternative hypothesis is directional) p-value = 0.0061315 *(if human omitted: R = -0.2409)*.

Comparing Pearson correlation coefficients, *ΔM* correlates with specific basal metabolic rate much better than it (inversely) correlates to adult body mass (0.7269 vs. -0.237) because *ΔM* is a linear function of specific basal metabolic rate (Figure 10) and *ΔM* is a negative supra-linear (convex) function of adult body mass (Figure 14).

**Figure 10.**
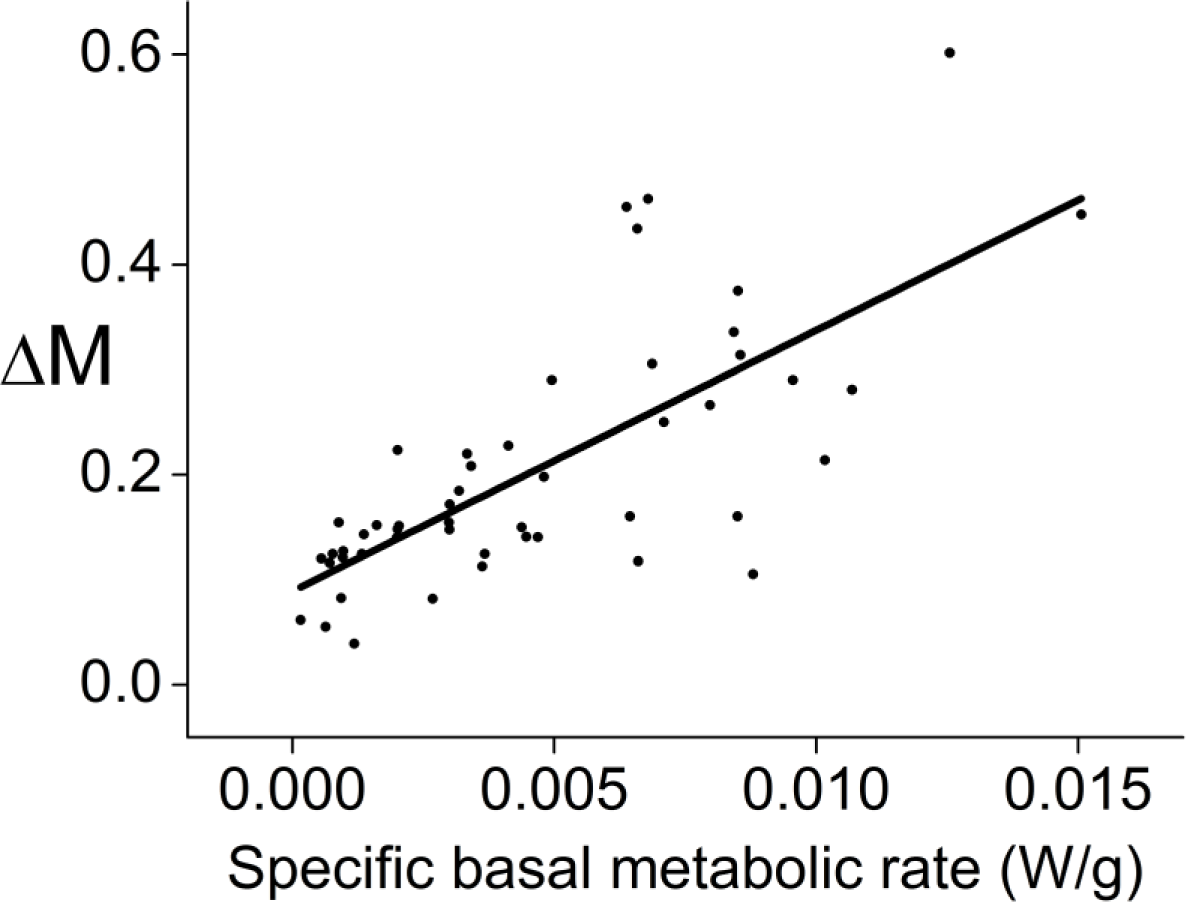
Across 50 eutherian species (different points are for different species): average epigenetic change per year (*ΔM*) is linearly proportional to specific basal metabolic rate.

In the Supplementary Material it is shown that the statistically significant correlation, across species, between *ΔM* and specific basal metabolic rate is maintained when phylogeny is considered.

#### Maximal lifespan is the same function of standardized epigenetic clock ticking rate (*ΔM*) as it is of standardized specific basal metabolic rate

Figure 11 shows, across 52 eutherian species (including human), maximal lifespan vs. standardized (to be between 0-100; by Eq. S1 in the Supplementary Material) specific basal metabolic rate, and maximal lifespan vs. standardized (by same Eq. S1) *ΔM*. Note the overlap between these plots. Further information is later herein, and in the Supplementary Material.

**Figure 11.**
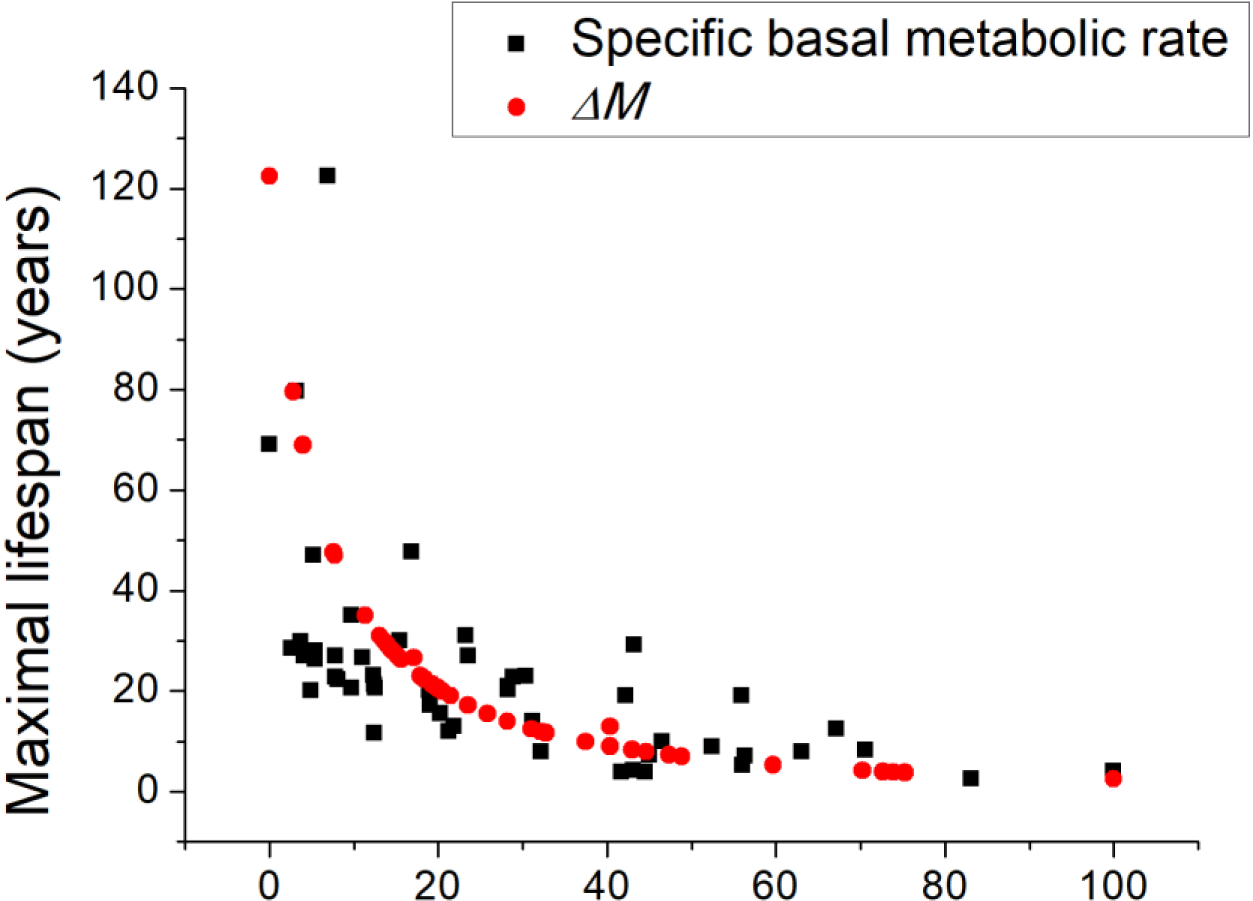
Maximal lifespan is the same function of standardized epigenetic clock ticking rate (*ΔM*) as it is of standardized specific basal metabolic rate. Across 52 eutherian species: maximal lifespan vs. standardized specific basal metabolic rate; and maximal lifespan vs. standardized *ΔM*. Both standardized (to be between 0-100) by Eq. S1 in the Supplementary Material. *To remind, ΔM is data calculated from an equation (Eq. 5) implicit to a fitted function to experimental data. The fitted function being “Clock 1” in, and the experimental data being from,* [105].

#### Epigenetic clock ticking rate (*ΔM*) is linear function of body surface-area per unit body mass

For eutherian species probed by the mammalian methylation array [127] (array used by [105]), those 111 species thereof whose maximal lifespan (required for calculation of *ΔM* by Eq. 5) and adult body masses are available in [1], delimited to the 99 non-flying species, their body surface-area was estimated from their adult body mass [1] by the Meeh equation (using *k* = 0.1 m^2^/kg^0.67^ and *b* = 0.67 [128], parameter values that aren’t suitable for flying mammal species). And then their body surface-area per unit body mass was calculated, which *ΔM* correlates with: Pearson correlation coefficient = 0.8052, one-tailed (alternative hypothesis is directional) p-value = 4.8409E-24 *(if human omitted: R = 0.8048)*.

As available, using experimentally determined, instead of Meeh equation calculated, body surface-area values with body masses from [128] (*values for “Peromyscus” and “rhinoceros” in* [128] *ignored because I’m unsure which species they are referring to*), the Pearson correlation coefficient between *ΔM* and body surface-area per unit body mass = 0.8681 (*n* = 21, one-tailed p-value = 1.70338E-07).

So, when comparing Pearson correlation coefficients (0.8681 vs. -0.237), *ΔM* correlates with body surface-area per unit body mass much better than it (inversely) correlates with adult body mass.

Out of the earlier used 99 non-flying species, for the 45 species that have specific basal metabolic rate data available in [1], 44 species after outlier *Sorex araneus* (common shrew) is removed: for 10 of these species, experimentally-determined body surface-area per unit body mass values are used, using data from [128], and for the other 34 species, absent in [128], they are estimated using body surface-area estimates from calculation, using the Meeh equation, with *k* = 0.1 m^2^/kg^0.67^ and *b* = 0.67 [128], with adult body mass input values from [1]: Pearson correlation coefficient between specific basal metabolic rate and surface-area per unit body mass = 0.8989, one-tailed (alternative hypothesis is directional) p-value = 6.00614E-17.

#### Specific basal metabolic rate mediates effect of body surface-area per unit body mass on epigenetic clock ticking rate (*ΔM*), and *ΔM* is a mediator of specific basal metabolic rate’s effect on maximal lifespan

For these 44 species, which includes human, using JASP software (version 0.14.1, selecting its bootstrap method called “Percentile”, run with 10,000 replications, utilizing its standardized estimates option, which makes each variable mean = 0 and standard deviation = 1, used to prevent numerical underflow in the underlying computations), mediation is observed between *ΔM* (as the dependent variable) and body surface-area per unit body mass, with the Mediator variable being specific basal metabolic rate. And between maximal lifespan (as the dependent variable) and specific basal metabolic rate, with the Mediator variable being *ΔM* (*reaffirming, with more species, that shown in earlier Table 20*). Results shown in first and second lines respectively of Table 22 below.

**Table 22.**

|  |  | 95% Confidence |  |  | 95% Confidence |  |  |
| --- | --- | --- | --- | --- | --- | --- | --- |
|  |  | Interval (CI) |  |  | Interval (CI) |  |  |
| $\alpha$ | $\beta$ | Indirect effect ($\alpha \cdot \beta$) | Lower | Upper | Direct effect ($\tau$) | Lower | Upper |
| 8 | 0.87 | 7.003 | 0.945 | 11.543 | -0.625 | -4.459 | 5.424 |
| 225 | -0.64 | -143.902 | -290.281 | -73.089 | -8.516 | -61.183 | 100.961 |

In Table 22 above, mediation is shown on both lines because neither has zero (0) within the 95% Confidence Interval (CI) for an indirect effect, and both have zero within the 95% CI for a direct effect. Mediation also observed when human is excluded (not shown).

#### Multiple linear regression reports that specific basal metabolic rate is a significant predictor of epigenetic clock ticking rate (*ΔM*), including when adult body mass is considered

For these 44 species, which includes human, I used JASP software (version 0.14.1) to conduct multiple linear regression. The response variable being *ΔM*, with adult body mass (kg) and specific basal metabolic rate (W/g) as explanatory/predictor variables. All variables logged (log_10_). Appreciable R^2^ observed. But high collinearity also, indicating redundancy (entwinning) of predictors. Which is consistent with adult body mass dictating *ΔM* because it determines specific basal metabolic rate. Wherein larger eutherian species have lower specific basal metabolic rate because they need to generate less metabolic heat per unit mass per unit time to maintain their body temperature. Because (of the square-cube law of geometry) they have a lower body surface-area per unit body mass, and so lose their metabolically generated heat less readily. Where (to interpret) a metabolic rate (by-)product(s), directly and/or indirectly, at least partially, drives ticking of the epigenetic clock.

In multiple linear regression, if the predictors are not collinear then the Variance Inflation Factor (VIF) is one, and if they are collinear then VIF is greater than one, how much more depending upon the degree of collinearity. A common rule of thumb is that if VIF is greater than four this is notable. In the results below, VIF exceeds four.

Collinearity in a regression model can reduce the precision of the estimated coefficients: increasing their standard errors, which can decrease their statistical significance, possibly making important coefficients not statistically significant. It can even confer them improper sign: negative when should be positive, or vice-versa. Although collinearity corrupts a multiple linear regression model, reporting redundancy (entwinning) of predictors confers insight.

Using JASP’s default method, which it calls “Enter”, in which explanatory variables are forced into the regression model in a user specified order, wherein adult body mass was forced in first (results are the same when specific basal metabolic rate is forced in first):

R^2^ = 0.658, adjusted R^2^ = 0.642, Unstandardized coefficients: β_0_ (intercept) = 0.058 ±

0.295 (p-value = 0.845), β_1_ = -0.052 ± 0.035 (regarding adult body mass. p-value = 0.146. VIF = <u>4.977</u>), β_2_ = 0.311 ± 0.120 (regarding specific basal metabolic rate. p-value = 0.013. VIF = <u>4.977</u>).

*Standardized coefficients: β_1_ = -0.302, β_2_ = 0.530*.

*Durbin-Watson statistic = 1.651 (between 1 and 3 is desirable, ideally around 2). The higher the F statistic, the better the model. F statistic is significant: F (2, 41) = 39.479, p = 2.767e-10. Residuals vs. predicted plot shows a balanced random distribution of the residuals around the baseline suggesting that the assumption of homoscedasticity has not been violated. Q-Q plot shows that the standardized residuals fit nicely along the diagonal suggesting that both assumptions of normality and linearity have not been violated*.

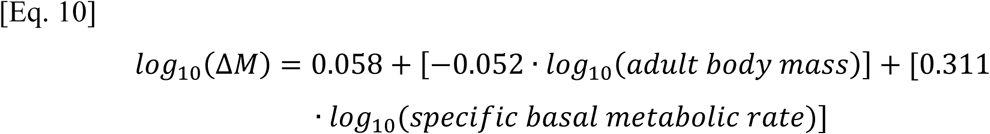

In the model above, specific basal metabolic rate *is*, and adult body mass is *not*, a significant predictor of *ΔM*. So, even with adult body mass included in the model, specific basal metabolic rate *is* a significant predictor of *ΔM*.

When using any of JASP’s other methods, called “Backward”, “Forward”, and “Stepwise”, in which a predictor can be removed (or not entered) if its presence is redundant in predicting the response variable (*ΔM*): adult body mass is excluded by the algorithm, and specific basal metabolic rate included. All these methods yield the same result, shown below. The adjusted R^2^ below is nearly as great as above: i.e., the regression model without adult body mass has very nearly the same predictive power for *ΔM* as with it, such is the collinearity, and the predictive power of specific basal metabolic rate for *ΔM*. (*N.B., when human is excluded, adjusted R^2^ is better at 0.664, not shown*).

R^2^ = 0.640, adjusted R^2^ = 0.631, Unstandardized coefficients: β_0_ (intercept) = 0.444 ± 0.140 (p-value = 0.003), β_1_ = 0.469 ± 0.054 (regarding specific basal metabolic rate. p-value = 7.325e-11). *Standardized coefficient = 0.8. Durbin-Watson statistic = 1.608. F (1, 42) = 74.626, p = 7.325e-11. Assumptions of homoscedasticity, normality, and linearity unviolated*.

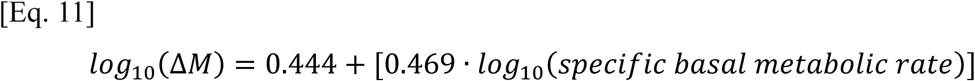

The regression analysis above probably underrepresents how much better specific basal metabolic rate is than adult body mass in predicting *ΔM* because it uses logged variables to make all the relations linear (because linearity is a requisite of multiple linear regression). But *ΔM* and specific basal metabolic rate, unlike *ΔM* and adult body mass, don’t require any logging to make their relation linear. Presumably, logging all these variables attenuates much of the superiority in how much more predictive specific basal metabolic rate is than adult body mass in predicting *ΔM*.

Unlike multiple linear regression, a Generalized Linear Model (GLM) doesn’t require linearity between the response variable and predictors (if such linearity can be conferred by a link function in the GLM). So, here it can be used with untransformed variables. Which are less collinear than their logged (log_10_) versions, solving the collinearity complication. I constructed a GLM (its random component = “Gamma” and systemic component = “Identity”) in JASP software (version 0.18.1.0) with *ΔM* as the response variable, and adult body mass (kg) and specific basal metabolic rate (W/g) as predictors (all unlogged). The model is significant (chi-squared p-value = 5.964e-12), wherein specific basal metabolic rate is a statistically significant predictor of *ΔM*, and adult body mass is nearly not, with p-values (for coefficients) of 8.118e-10 and 0.031 respectively (p-value for intercept = 1.819e-9). So, even with adult body mass included in the model, specific basal metabolic rate is a significant predictor of *ΔM*. Collinearity is low (VIF = 1.121). Model equation:

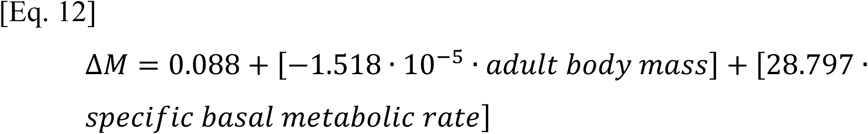

When adult body mass is omitted from the GLM, the model is *more* significant (chi-squared p-value = 1.277e-12), and specific basal metabolic rate is a more significant predictor (its p-value = 1.123e-10).

A Generalized Additive Model (GAM) is presented in the Supplementary Material, also using untransformed variables, which reports specific basal metabolic rate *is*, whereas adult body mass is *not*, a significant predictor of epigenetic clock ticking rate (*ΔM*). Arguably, GAMs are superior to GLMs.

#### Multiple linear regression with body surface-area per unit body mass as a predictor

*ΔM* is a linear function of specific basal metabolic rate and of body surface-area per unit body mass (which are themselves linearly interrelated). So, with these variables multiple linear regression can be performed with untransformed variables. No logging required. Below is the regression result, using the same 44 species as in the preceding section, wherein the response variable is *ΔM*, while body surface-area per unit body mass (m^2^/kg) and specific basal metabolic rate (W/g) are explanatory variables. Using JASP’s “Enter” method (result is the same whichever explanatory variable is forced in first):

R^2^ = 0.656, adjusted R^2^ = 0.639, Unstandardized coefficients: β_0_ (intercept) = 0.089 ± 0.018 (p-value = 1.529e-5), β_1_ = -0.078 ± 0.235 (regarding body surface-area per unit body mass. p-value = 0.740. VIF = <u>5.208</u>), β_2_ = 30.412 ± 7.296 (regarding specific basal metabolic rate. p-value = 1.545e-4. VIF = <u>5.208</u>). *Standardized coefficients: β_1_ = -0.070, β_2_ = 0.872. Durbin-Watson statistic = 1.541. F (2, 41) = 39.010, p = 3.250e-10. Assumptions of homoscedasticity, normality, and linearity unviolated*.

In this model, the collinearity is so severe that the standard error in the regression coefficient for body surface-area per unit body mass is so great that this coefficient is erroneously negative. Erroneous because *ΔM* positively correlates (e.g., in a Pearson correlation) with body surface-area per unit body mass. Specific basal metabolic rate is a significant predictor of *ΔM* in this model.

Using any of JASP’s other methods (Backward, Forward, Stepwise), body surface-area per unit body mass is excluded by the algorithm, and specific basal metabolic rate included, which increases adjusted R^2^:

R^2^ = 0.655, adjusted R^2^ = 0.646, Unstandardized coefficients: β_0_ (intercept) = 0.088 ± 0.018 (p-value = 1.140e-5), β_1_ = 28.218 ± 3.163 (regarding specific basal metabolic rate. p-value = 3.020e-11). *Standardized coefficient = 0.809. Durbin-Watson statistic = 1.559. F (1, 42) = 79.591, p = 3.020e-11. Assumptions of homoscedasticity, normality, and linearity unviolated*.

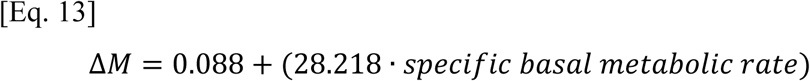

If the intercept is set as zero (which might be legitimate because if body surface-area per unit body mass is zero, then inherently both specific basal metabolic rate and *ΔM* must be zero), using JASP’s “Enter” method, now the coefficient for body surface-area per unit body mass is of the correct sign, but now the standard error is greater, probably because the collinearity is greater (VIF now equals 12.595):

R^2^ = 0.859, adjusted R^2^ = 0.853, Unstandardized coefficients: β_1_ = 0.133 ± 0.287 (regarding body surface-area per unit body mass. p-value = 0.645. VIF = <u>12.595</u>), β_2_ = 36.418 ± 8.951 (regarding specific basal metabolic rate. p-value = 2.038e-4. VIF = <u>12.595</u>). *Standardized coefficients: β_1_ = 0.119, β_2_ = 1.044. Durbin-Watson statistic = 1.294. F (2, 42) = 128.400, p = 1.274e-18. Assumptions of homoscedasticity, normality, and linearity unviolated*.

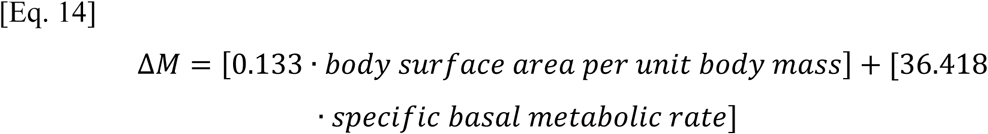

When using any of JASP’s other methods (Backward, Forward, Stepwise), body surface-area per unit body mass is algorithmically excluded, deemed redundant, given its collinearity with specific basal metabolic rate:

R^2^ = 0.859, adjusted R^2^ = 0.855, Unstandardized coefficient: β_1_ = 40.398 ± 2.499 (p-value = 6.934e-20), *Standardized coefficient = 1.158. Durbin-Watson statistic = 1.265. F (1, 43) = 261.358, p = 6.934e-20. Assumptions of homoscedasticity, normality, and linearity unviolated*.

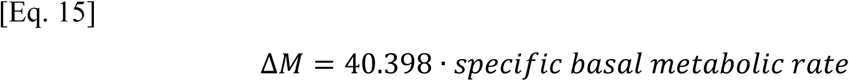

In the model above, note the high adjusted R^2^ value (0.855), and statistical significance of specific basal metabolic rate as sole predictor of *ΔM* (p-value = 6.934e-20).

#### Partial and semi-partial correlations with epigenetic clock ticking rate (*ΔM*)

Using Pearson: partial and semi-partial (part) correlations were performed in JASP (version 0.18.1.0) for these 44 species, which includes human. A partial correlation is the correlation between a dependent and an independent variable after controlling for the influence of one or more other variables on *both* the dependent and independent variable. Semi-partial (part) correlation is the correlation between a dependent and an independent variable after controlling for the influence of one or more other variables on the dependent *or* the independent variable (herein the independent variable).

In Table 23 below: for set *A*: *Y* is *ΔM*, *X* is specific basal metabolic rate (W/g), and *Z* is adult body mass (kg) (all logged, log_10_). For set *B*: *Y* is *ΔM*, *X* is specific basal metabolic rate (W/g), and *Z* is body surface-area per unit body mass (m^2^/kg) (all untransformed, because they are all linear relations of each other). P-values are one-tailed because each alternative hypothesis is directional.

**Table 23.** Pearson correlation coefficients:

| Set | **Y.X** | Partial<br>Y.X(Z) | Part<br>Y.X(Z) | **Y.Z** | Partial<br>Y.Z(X) | Part<br>Y.Z(X) | **Z.X** |
| --- | --- | --- | --- | --- | --- | --- | --- |
| A | 0.800 | 0.376 | 0.238 | -0.776 | -0.226 | -0.135 | -0.894 |
| B | 0.809 | 0.546 | 0.382 | 0.714 | -0.052 | -0.031 | 0.899 |

One-tailed p-values:
| Set | **Y.X** | Partial<br>Y.X(Z) | Part<br>Y.X(Z) | **Y.Z** | Partial<br>Y.Z(X) | Part<br>Y.Z(X) | **Z.X** |
| --- | --- | --- | --- | --- | --- | --- | --- |
| A | 3.63541E-11 | 0.0059455 | 0.059888 | 3.05166E-10 | 0.0700905 | 0.191136 | 1.54484E-16 |
| B | 1.52117E-11 | 0.000063 | 0.0052515 | 2.64147E-08 | 0.6297393 | 0.5782184 | 5.89924E-17 |

In Table 23 above, *ΔM* significantly correlates with specific basal metabolic rate, and *ΔM* significantly correlates with body surface-area per unit body mass, even when controlling for adult body mass. By contrast, when statistically excluding the effect of specific basal metabolic rate, there is no statistically significant relationship between *ΔM* and adult body mass, nor between *ΔM* and body surface-area per unit body mass. Beware that partial and semi-partial (part) correlations are, as explained in the Supplementary Material, interrelated with multiple linear regression. Just as collinearity can corrupt multiple linear regression, it can corrupt partial and semi-partial correlations. So, given the earlier reported collinearity, these partial and semi-partial correlation coefficients are likely (to an extent proportional to the *VIF* values) corrupted.

#### Growth rate as input into the ticking rate of the epigenetic clock

Preeminent leaders in the field think that aging isn’t damage but is development (e.g., [46, 105]). Programmed. Wherein the program is the ticking of the epigenetic clock (*ΔM*), and it is executed faster in shorter-living species. All be it this theory doesn’t explain what causes *ΔM*. Nor why faster development inexorably means shorter maximal lifespan. Why does the former limit the latter? Molecular mechanism(s)? If there is no such limitation, why is the program that way? Why age faster rather than slower?

My interpretation of data is that *ΔM* is (*at least partially; directly and/or indirectly*) metabolic damage. A stipulation of a specific basal metabolic rate value is that the animal is an adult. Development, which stops at adulthood, inextricably requires metabolic rate and I interpret that this is how development contributes to the ticking of the epigenetic clock. *ΔM* is the average change in (aforementioned) variable *M* per year, over the entire lifetime. *M* changes faster while a mammal develops [105]. This is consistent with the present thesis, wherein a metabolic (by-)product(s) drives change in *M*. When a postnatal mammal isn’t fully grown its body mass is smaller than an adult, and so its surface-area to body mass ratio is larger, and so it must produce more metabolic heat per unit mass per unit time (i.e., have a higher resting metabolic rate per unit mass than an adult) to maintain its body temperature. Moreover, while the juvenile mammal develops it also has a higher resting metabolic rate per unit mass (than an adult) to provide energy, and metabolic intermediates, for growth. Wherein, presumably, a higher growth rate inextricably requires a higher resting metabolic rate per unit mass. To hypothesize, resting specific metabolic rate isn’t only a drive of metabolic damage (≈aging) during adulthood, but during development also.

Newly reported herein, using data for 26 of the earlier used 44 eutherian species, those 26 species that have postnatal growth data available in [1]: *ΔM* is a linear correlate of postnatal growth rate (and specific basal metabolic rate), specific basal metabolic rate is a linear correlate of postnatal growth rate.

As *ΔM* is a linear function of both postnatal growth rate and specific basal metabolic rate (which are themselves linearly interrelated), with these variables multiple linear regression can be performed with untransformed data. No logging required. Below is a multiple linear regression model with response variable of *ΔM*, while postnatal growth rate (days^-1^) and specific basal metabolic rate (W/g) are predictors. Using JASP’s “Enter” method (result is the same whichever explanatory variable is forced in first) or equally its “Backward” method:

R^2^ = 0.700, adjusted R^2^ = 0.674, Unstandardized coefficients: β_0_ (intercept) = 0.082 ± 0.022 (p-value = 0.001), β_1_ = 1.968 ± 1.091 (regarding postnatal growth rate. p-value = 0.084. VIF = 2.379), β_2_ = 18.354 ± 5.680 (regarding specific basal metabolic rate. p-value = 0.004. VIF = 2.379). *Standardized coefficients: β_1_ =* 0.318*, β_2_ =* 0.569*. Durbin-Watson statistic = 1.462. F (2, 23) = 26.838, p = 9.690e-7. Assumptions of homoscedasticity, normality, and linearity unviolated*.

In this model, specific basal metabolic rate *is*, and postnatal growth rate is *not*, a significant predictor of *ΔM*.

When instead using JASP’s “Forward” or “Stepwise” methods: postnatal growth rate is excluded by the algorithm, and specific basal metabolic rate included (adjusted R^2^ = 0.643).

So, specific basal metabolic rate is more predictive of *ΔM* than postnatal growth rate. Possibly because, over their lifetime, eutherians spend longer as an adult than a juvenile.

Using Pearson: partial and semi-partial (part) correlation coefficients were obtained using JASP, results are in Table 24 below. Where *Y* is *ΔM*, *X* is specific basal metabolic rate (W/g), and *Z* is postnatal growth rate (1/days). All untransformed, as they are all linear relations of each other. P-values are one-tailed because each alternative hypothesis is directional.

**Table 24.**
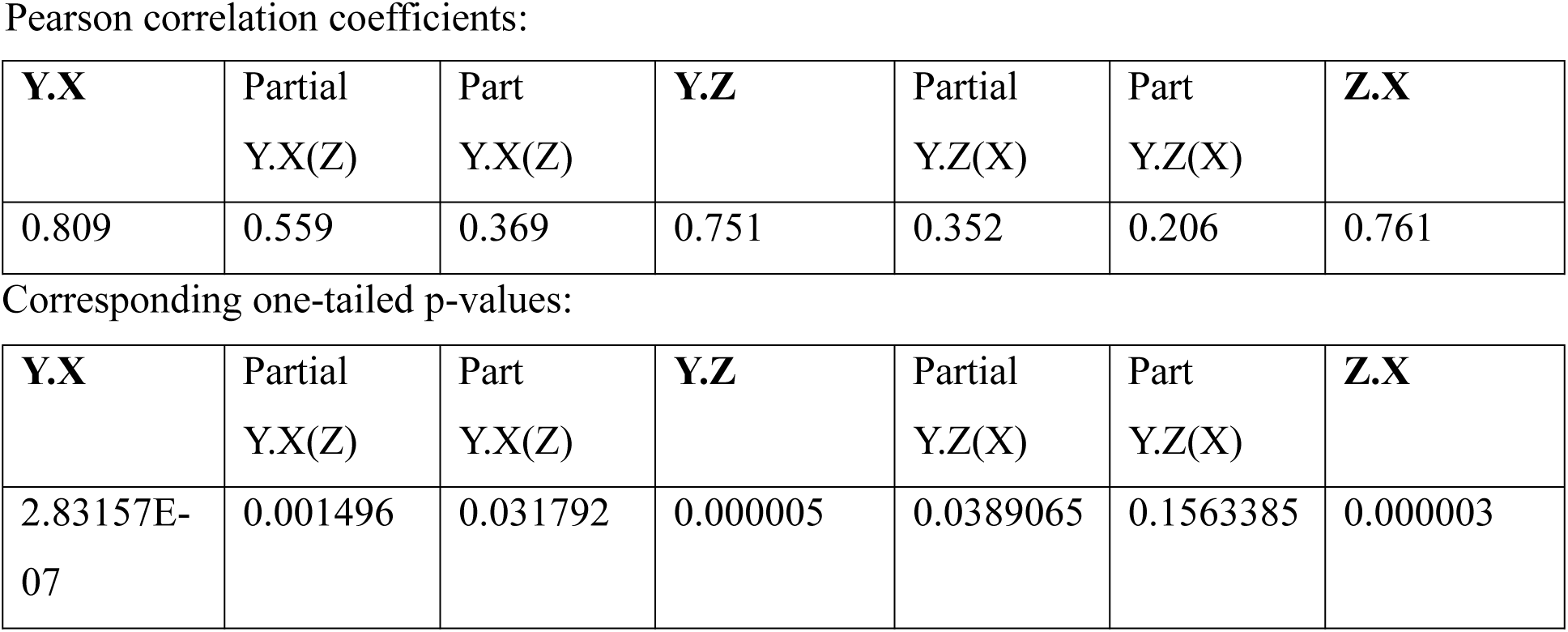

Analysing Table 24 above, when statistically excluding the effect of postnatal growth rate, there is still a statistically significant relationship between *ΔM* and specific basal metabolic rate. But when statistically excluding the effect of specific basal metabolic rate, there is not a statistically significant relationship between *ΔM* and postnatal growth rate for the part correlation. Although this analysis was conducted with a small sample size (*n*=26). Collinearity isn’t so high in this case (VIF = 2.379 in the correspondent multiple linear regression).

Average rate of methylation change, in developing (juvenile) versus developed (adult), correlates with a Pearson correlation coefficient of 0.78 [130]. In this present data, postnatal growth rate (which, presumably, is inextricably linked to resting metabolic rate per unit mass during postnatal growth) and specific basal metabolic rate (an adult specific measure) correlate with a Pearson correlation coefficient of 0.761 (one-tailed p-value = 3.1453e-6). Across 131 mammal and bird species (all those mammal and bird species with sufficient data available in [1]), the Pearson correlation coefficient between postnatal growth rate and specific basal metabolic rate is 0.759 (one-tailed p-value = 4.158e-26. Spearman = 0.770).

#### Is the epigenetic clock ticking rate (*ΔM*) a program or damage?

If aging is a deliberate program, it is presumably corruptible by mutation(s), and so out of the billions of humans that have ever lived there is likely to be a number in which this program is corrupted and so don’t age, but this hasn’t been reported.

Age-related decline of form and function is consistent with aging being accumulating damage. Why make the program mimic damage when actual damage can confer the same effect? Why make a machine without *net* damage, which can last indefinitely, that is then programmed to not last indefinitely, but for a short time, with declining form and function, in mimicry of accumulating damage? Why is damage (e.g., to DNA [97–98]) observed to accumulate with age?

In later Figure 14, *ΔM* is referred to as “specific epigenetic damage rate”, to correspond with my interpretation that *ΔM* is metabolic damage. Figure 14 shows that *ΔM* positively linearly correlates with rates of damage, such as DNA damage rate. Correlating with different damage rates and not being a damage rate itself, being programmed, isn’t the most parsimonious account for this fact pattern.

Programmed theory dismisses damage as contributory to aging rate, but then why do damage rates, such as DNA damage rate, inversely and stereotypically relate to maximal lifespan? By the same relation form that *ΔM* relates to maximal lifespan. Refer to Figure 14.

The programmed concept doesn’t molecularly specify what causes *ΔM*, not even by falsifiable hypothesis. Maximal lifespan relates to *ΔM* by the same equation (Eq. 18) that it relates to different damage rates, and to specific basal metabolic rate, which is consistent with *ΔM* being a metabolic damage rate. Refer to Eq. 18, Figure 11, and their accompanying texts.

#### Why do different mammal species have different maximal lifespans?

In short, (*my interpretation of data*) because of IF1 protein. Data in the Supplementary Material shows that in a species set, maximal lifespan positively correlates with specific IF1 protein activity. IF1 protein is an endogenous inhibitor of F_1_F_0_ ATP hydrolysis (*that doesn’t inhibit F_1_F_0_ ATP synthesis*) [8–14], and thereby of specific basal metabolic rate.

Figure 12 shows that, across 263 mammal species, maximal lifespan is inversely proportional to specific basal metabolic rate. Further such analysis is in the Supplementary Material.

**Figure 12.**
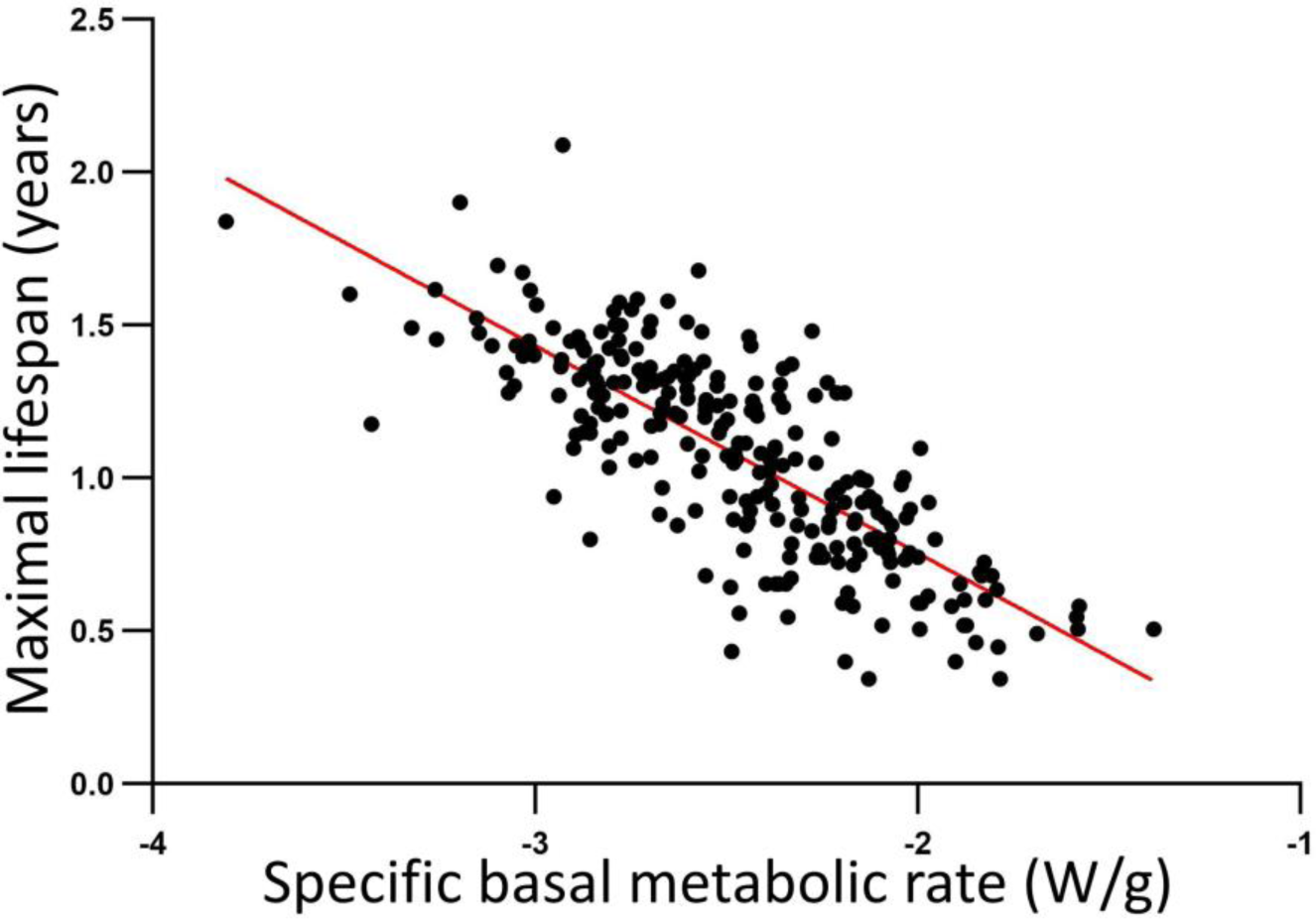
A log_10_-log_10_ plot showing that, across 263 mammal species, maximal lifespan (years) is inversely proportional to specific basal metabolic rate (W/g). Fitted line is *y* = m*x* + c, where *m* = -0.6780 (±0.0651), *c* = -0.6017 (±0.1631); and adjusted R^2^ = 0.6155. Pearson correlation coefficient = -0.7855, one-tailed (alternative hypothesis is directional) p-value = **1*10^-56^**.

Figure 13 is a log_10_-log_10_ plot showing, across 259 mammal species, that specific basal metabolic rate (W/g) negatively, and maximal lifespan (years) positively, scales with adult body mass (kg). Wherein I interpret this data to teach that, at least across these species, adult body mass dictates specific basal metabolic rate, which in turn (at least in part) dictates maximal lifespan.

**Figure 13.**
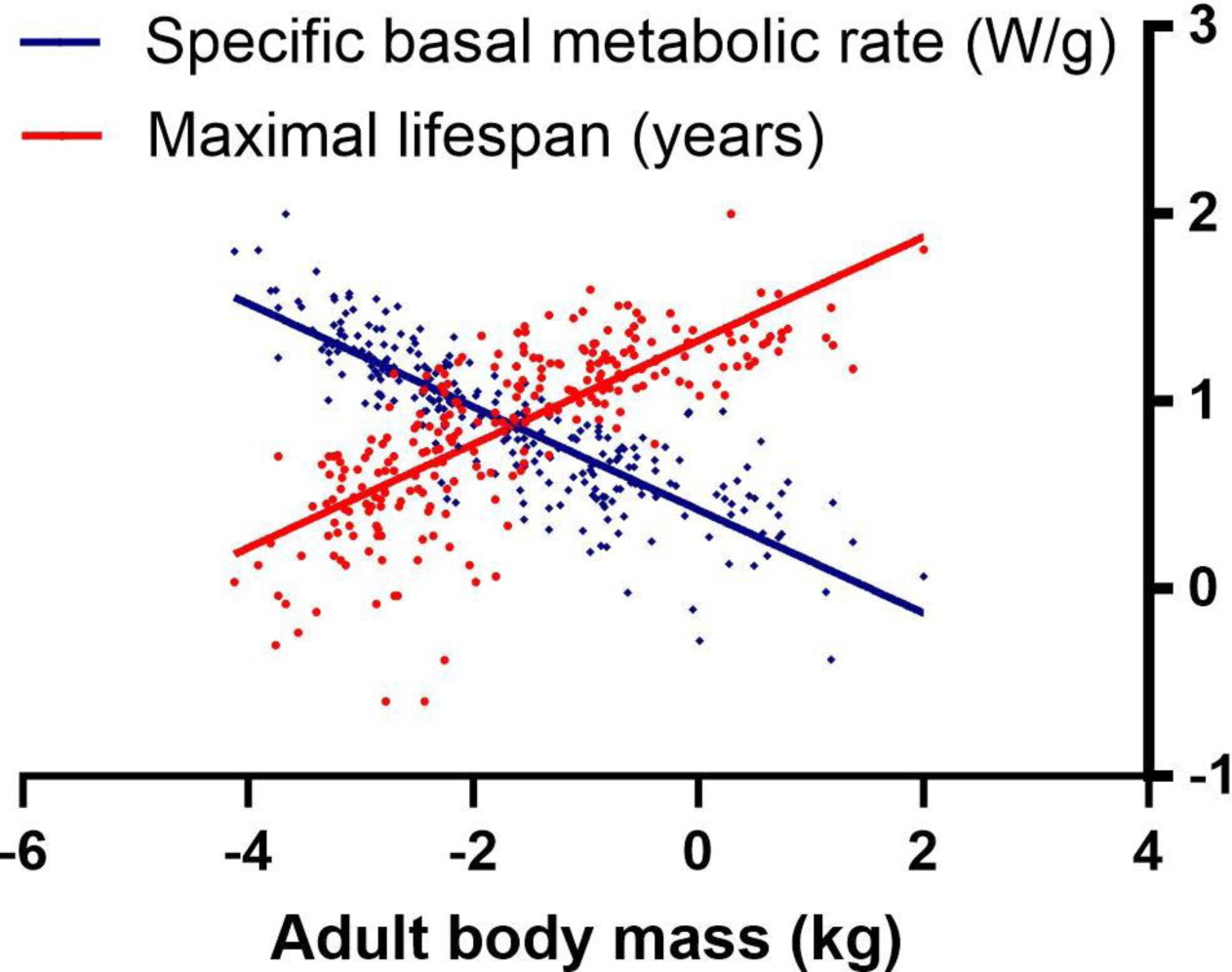
Log_10_-log_10_ plot showing, across 259 mammal species, that specific basal metabolic rate negatively, and maximal lifespan positively, scales with adult body mass. Plotted data is standardized by Eq. S1 and logged as explained in the Supplementary Material.

Figure 14 summarizes a pattern of interrelations I’ve discovered across mammal species, with all its boxes arranged as per (predicted) causality. For each interrelation shown, the higher box is the independent (*x*-axis), and the lower box is the dependent (*y*-axis), variable. When boxes have the same height in the schema then causality isn’t necessarily predicted, as they correlate with each other because of shared cause higher in the schema, and their horizontal ordering is arbitrary (*though DNA damage may drive epigenetic damage* [125]).

**Figure 14.**
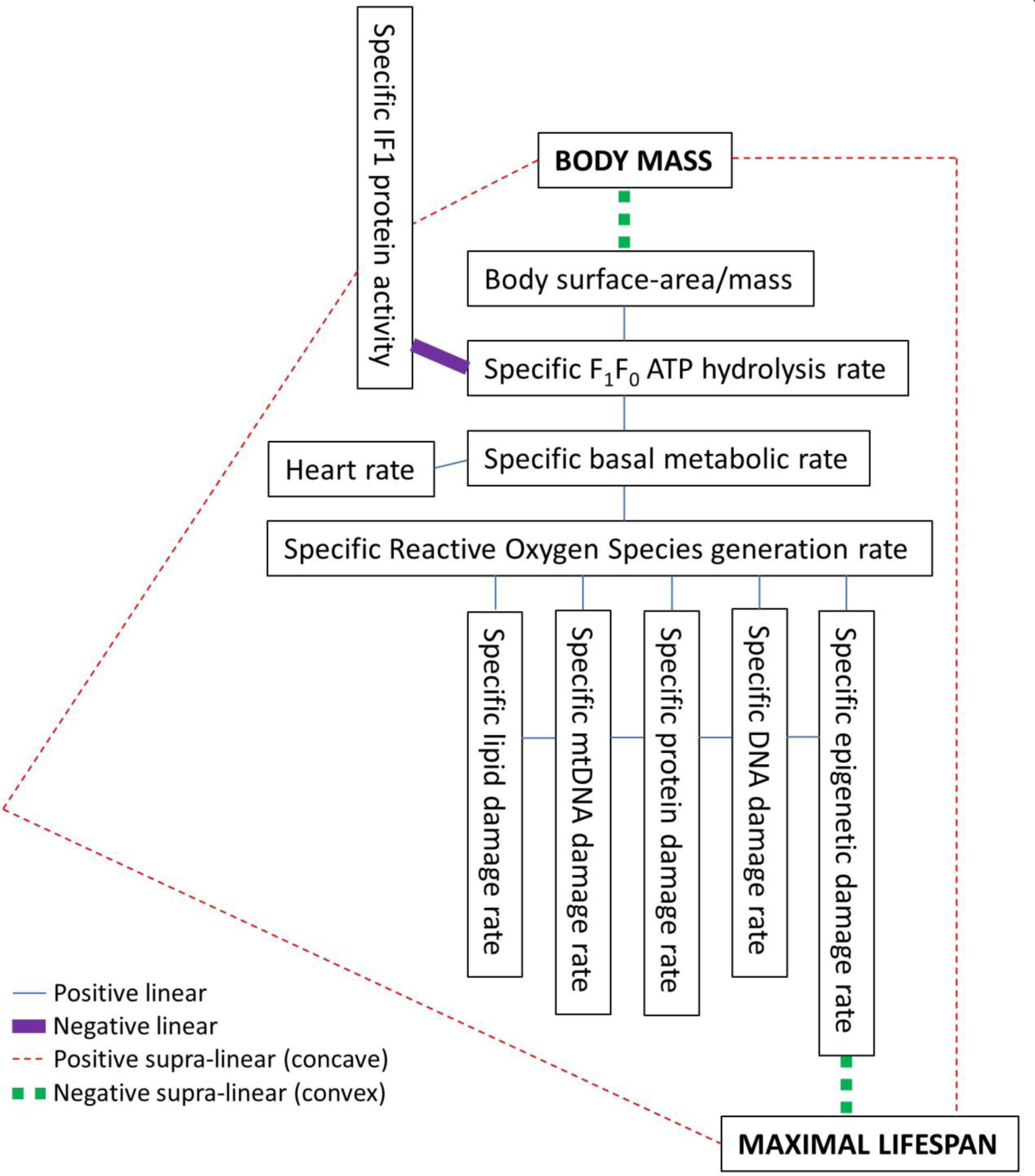
Using data from 353 mammal species, this pattern of interrelations is observed across mammal species. “Specific” in the figure means “per unit mass” (or otherwise controlled for body mass). “mtDNA” is mitochondrial DNA. “Body surface-area/mass” is body surface-area per unit body mass. For each interrelation shown, the higher box is the independent (*x*-axis), and the lower box is the dependent (*y*-axis), variable. When two boxes have the same height in the schema, the box on the right is the independent (*x*-axis), and the box on the left is the dependent (*y*-axis), variable (or equally vice versa when the interrelation is positive linear). I interpret this pattern as each higher box causes/determines its lower box (or plurality of the latter), wherein the highest box’s determination of the second is because of the square-cube law of geometry, from which the rest cascades, wherein a (by-)product(s) of metabolic rate causes damage, which is aging. The correlation coefficients that this diagram is a schematization of are disclosed in the Supplementary Material. Their asymptotically exact harmonic mean p-value [76] {*its post-published corrected method, as corrected on 7 October 2019*} = **1*10^-119^**. Extremely significant. Each supra-linear function here can be made linear by logging the *x*- and *y*-axis data. Further information is in the main text here, and much more in the Supplementary Material.

To illustrate an implicit feature of this schema, by using three imaginary illustrating variables called *v1*, *v2* and *v3*: if variable *v1* is some function (one of the four types shown in the schema) of another variable *v2*, wherein *v2* is a positive linear function of a third variable *v3*, then *v1* is the same form of function (that it is of *v2*) of *v3* also. For example, Body surface-area/mass is a negative supra-linear (convex) function of Body mass, wherein Specific F_1_F_0_ ATP hydrolysis rate is a positive linear function of Body surface-area/mass, meaning Specific F_1_F_0_ ATP hydrolysis rate is a negative supra-linear (convex) function of Body mass also.

So, because of this implicit character to the schema, the number of interrelations that it discloses (78) is much greater than the number of connecting lines it shows. Indeed, beware that a cursory look at Figure 14 might interpret it as disclosing only epigenetic damage rate as dictating maximal lifespan. But its implicit disclosure must be considered also. The other damage rates shown are (implicitly) disclosed as contributory factors also.

The damage rates correlate with each other because (to interpret) they share the same cause, which they also positively correlate with, which is ultimately specific basal metabolic rate, as largely set by specific F_1_F_0_ ATP hydrolysis rate (as constrained by specific IF1 protein activity), as per their body surface-area/mass ratio. Wherein I interpret that (at least) these damage rates are collectively the aging rate.

Data from 353 mammal species was used to produce Figure 14. The correlations that Figure 14 is a schema of are reported in the Supplementary Material. As shown there, the asymptotically exact harmonic mean p-value [76] (*its post-published corrected method, as corrected on 7 October 2019*) for the correlations that Figure 14 is a schema of = **1*10^-119^**. Extremely significant. Extremely. Disparate data interlocks like jigsaw pieces to produce Figure 14.

Instrumental variable analysis, mediation analysis, Structural Equation Modelling (SEM), and other analysis presented herein (and in the Supplementary Material) is consistent with Figure 14, conferring it further statistical basis.

Figure 14, and its underlying data, predicts that a selective F_1_F_0_ ATP hydrolysis inhibitor drug (*that doesn’t inhibit F_1_F_0_ ATP synthesis*) can slow aging. For example, a drug taught in one or more of my international (PCT) patent applications [18–20]. Note how such intervention interdicts high in the schema, critically before it forks into different branches, wherein interdicting any single one of those branches is likely sub-optimal as the other branches would be extant, and addressing each individually is probably *much* more complicated than a single point of intervention higher in the schema.

Figure 14 in equations: [Eq. 16]

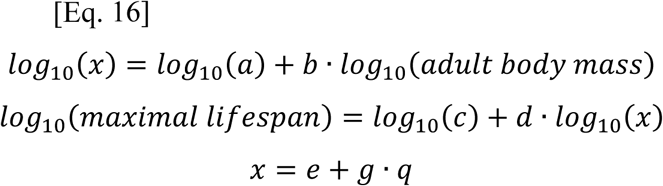

Where these scaffold equations are populated with different variables to get the full set of equations. Iterating, *x* is selected in turn from body surface-area/mass, specific IF1 protein activity, specific F_1_F_0_ ATP hydrolysis rate, specific basal metabolic rate, heart rate, specific Reactive Oxygen Species (ROS) generation rate, specific DNA damage rate, specific mtDNA damage rate, specific protein damage rate, specific lipid damage rate, specific epigenetic damage rate, *q* is a different selection of *x* (iterating, each different option of *x* selected in turn), and *a*, *b*, *c*, *d*, *e*, and *g* are constants, which can be different for different selections of *x*. Wherein *b* and *d* are positive when *x* is specific IF1 protein activity, and *g* is negative when *x* or *q* is specific IF1 protein activity.

Parameter values (*a*, *b*, *c*, *d*, *e*, *g*) are disclosed in the Supplementary Material.

#### Proposed basis to the interrelations of Figure 14

In short, because of the square-cube law of geometry, and because a metabolic rate (by-)product(s) causes aging.

*N.B. in this headed section, “specific damage rate” can refer to any one of specific DNA damage rate, specific mtDNA damage rate, specific protein damage rate, specific lipid damage rate, specific epigenetic damage rate, or combination of two or more thereof*.

Across mammal species, because of the square-cube law of geometry, body surface-area is a supra-linear (positive concave) function of body mass, which makes body surface-area per unit body mass a supra-linear (negative convex) function of body mass, which is why each of the following is observed to be a supra-linear (negative convex) function of body mass: specific F_1_F_0_ ATP hydrolysis rate, specific basal metabolic rate, heart rate, specific ROS generation rate, specific damage rate. Shared supra-linearity to body mass is why there is the observed linearity between each of body surface-area per unit body mass, specific F_1_F_0_ ATP hydrolysis rate, specific basal metabolic rate, heart rate, specific ROS generation rate, and specific damage rate. Maximal lifespan is a supra-linear (negative convex) function of each of these, and a supra-linear (positive concave) function of body mass.

Specific IF1 protein activity inhibits specific F_1_F_0_ ATP hydrolysis rate, wherein they have an inverse linear relationship. Specific IF1 protein activity (by inhibition) sets the F_1_F_0_ ATP hydrolysis rate, which (largely) sets the specific basal metabolic rate [which sets the heart rate], which sets the specific ROS generation rate, which sets the specific damage rate, which sets the maximal lifespan, wherein each species’ specific IF1 protein activity is inherently set in accordance with their body surface-area/mass, which is a function of their body mass, because this determines how much metabolic heat they need to produce per unit mass per unit time to maintain their body temperature.

#### Across mammal species, why is maximal lifespan a supra-linear and not a linear negative function of specific damage rate?

Maximal lifespan is observed to be a supra-linear (negative convex) function of each of the specific damage rates I considered, and of their (standardized, by Eq. S1) combination. Suggesting that specific damage rate synergistically detriments maximal lifespan. Instead of additively (which would be indicated by maximal lifespan being a negative linear relation of specific damage rate).

So, an increase in specific damage rate is more detrimental [to maximal lifespan] the higher the prior specific damage rate. So, detriment to maximal lifespan of increased specific damage rate is not simply the mathematical sum of the prior specific damage rate and the extra specific damage rate, it is synergistically more than that, how much more depending upon the prior specific damage rate. Suggesting that when the specific damage rate is less, decreasing specific damage rate by a certain absolute amount increases maximal lifespan more. This hints at increasingly increasing maximal lifespan benefit from decreasing specific damage rate more and more. Such that decreasing specific damage rate by a certain absolute amount increases maximal lifespan more in a human than a mouse.

At least in part, this synergy may come from interaction(s) between the same and/or different damage types, where more of one type(s) can increase the amount, and/or detriment of the amount, of the same and/or another damage type(s). For (non-limiting) possible example, some lipid peroxidation products are themselves Reactive Oxygen Species (ROS), which can attack further lipid molecules, generating more such ROS in a peroxidative chain reaction, any one or more of which can cause damage to a different macromolecule type [131]. For example, peroxidized cardiolipin, in the inner mitochondrial membrane, can inactivate Complex IV [132].

### Figure 14 as a graph

Refer to Figure 14: consider body mass as its “input variable”, maximal lifespan as its “output variable”, and all the other variables as “internal variables”.

After collapsing specific IF1 protein activity and specific F_1_F_0_ ATP hydrolysis rate into a single internal variable of the latter divided by the former (*needed because specific IF1 protein activity negatively correlates with the other internal variables*), standardizing all these internal variables (each internal variable standardized to be between 0-100, using Eq. S1 in the Supplementary Material), I took their average to create a single, combination internal variable, which I call *λ* (*which I interpret, given its components, to encompass, per unit body mass: basal metabolic drive, rate, and damage rate*).

Figure 15 shows, when reading its two plots together, body mass vs. maximal lifespan, via *λ*. You can see that *λ* is a negative supra-linear (convex) function of body mass, and that maximal lifespan is a negative supra-linear (convex) function of *λ*. An incredible result, which occurs because all these internal variables are so highly linearly correlated with each other, and because each of them too is a negative supra-linear (convex) function of body mass, and maximal lifespan is a negative supra-linear (convex) function of each of them too, and so when they are standardized (to each be between 0-100) and combined, this combination has the same relations as each of them individually. Further information, and further such analysis, is in the Supplementary Material.

**Figure 15.**
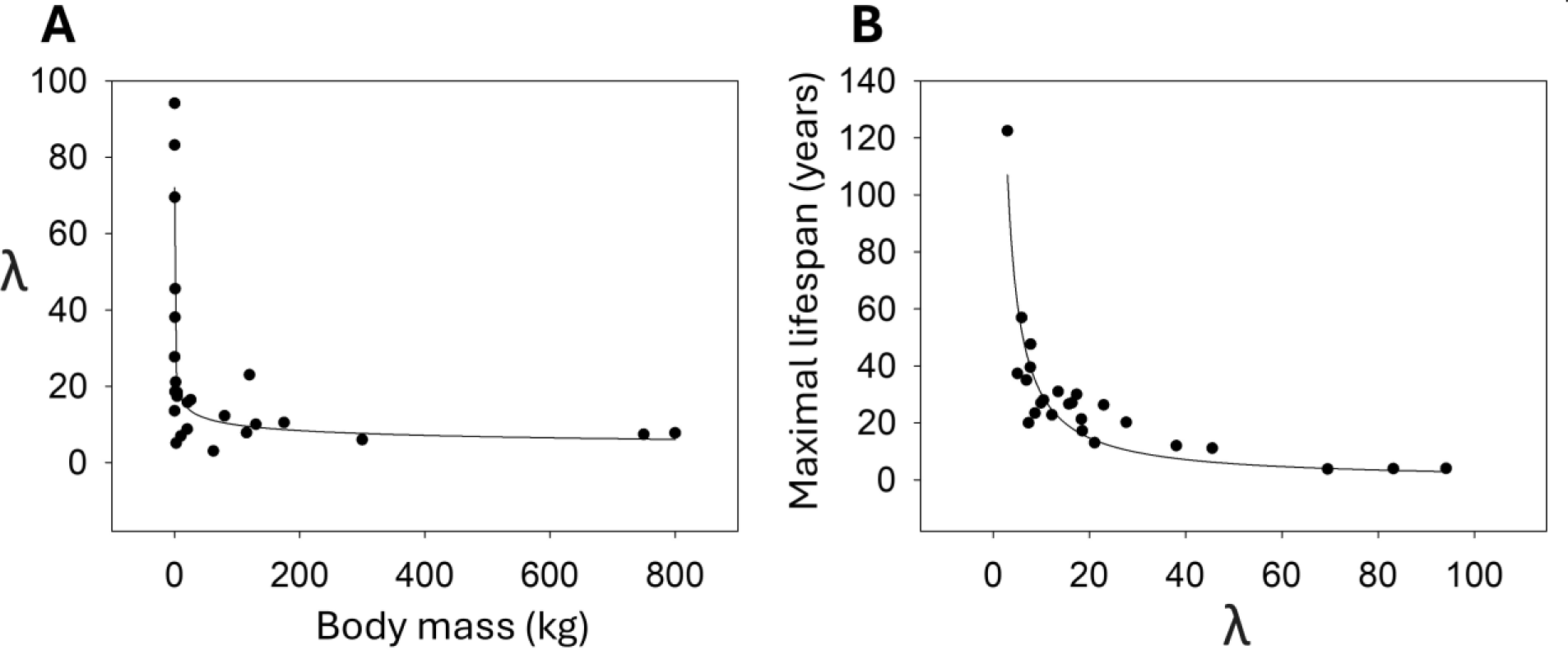
**(A)** *λ* vs. body mass (kg). Best-fit line equation: *λ* = 28.9609 ·body mass^−0.2344^. Adjusted R^2^ = 0.5220. **(B)** Maximal lifespan (years) vs. *λ*. Best-fit line equation: maximal lifespan = 335.3174 · *λ*^−1.0438^. Adjusted R^2^ = 0.8138. *Plots were done, and best-fit line equations found, using SigmaPlot 15.0 software (Systat Software, Inc., San Jose, California, USA). Human data included in both plots*.

Figure 16 is a collapsing of the two plots of Figure 15 into a single plot, wherein *λ* is as defined prior and *γ* is the average of standardized body mass and standardized maximal longevity, each standardized to be between 0-100 by (aforementioned) Eq. S1.

**Figure 16.**
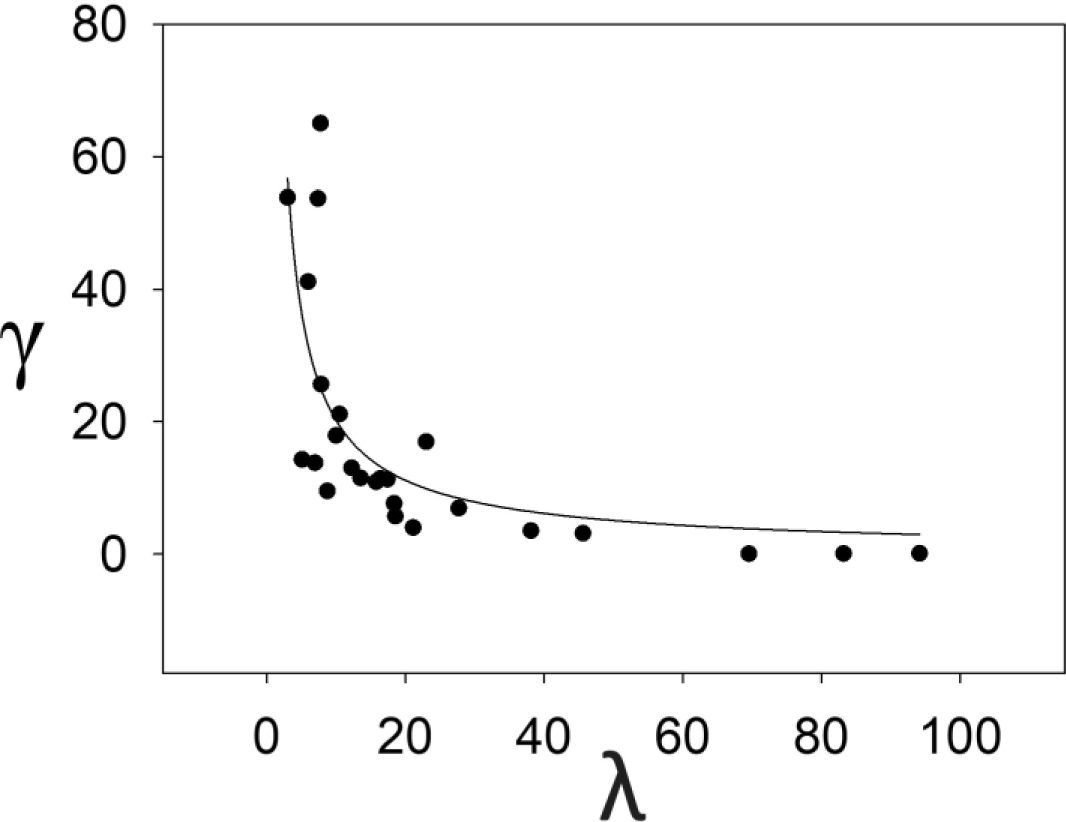
γ vs. λ. Best-fit line equation: *γ* = 145.0482 · *λ*^−0.8575^. Adjusted R^2^ = 0.5158. Plot done, and its best-fit line equation found, using SigmaPlot 15.0 software (Systat Software, Inc., San Jose, California, USA). Human data among that used.

### Figure 14 as an equation

Best-fit line equations of Figure 15: [Eq. 17] and [Eq. 18] respectively:

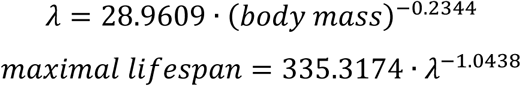

Where, to extrapolate, *λ* can be any of, or an average of multiple of, (each standardized, by Eq. S1): body surface-area/mass, specific F_1_F_0_ ATP hydrolysis rate/specific IF1 protein activity, specific F_1_F_0_ ATP hydrolysis rate, specific basal metabolic rate, heart rate, specific Reactive Oxygen Species (ROS) generation rate, specific DNA damage rate, specific mtDNA damage rate, specific protein damage rate, specific lipid damage rate, specific epigenetic damage rate.

Refer to earlier Figure 11, which shows plots of two standardized internal variables, specific basal metabolic rate, and specific epigenetic damage rate. Note the overlap of these two plots. This illustrates how these equations can have such interoperability of *λ*. *λ* being different standardized internal variables, or average of two or more thereof.

Incidentally, Eq. 18 rearranged:

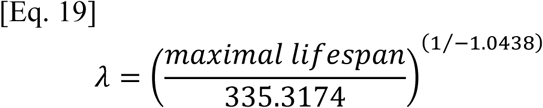

Eq. 17 substituted into Eq. 18, gives a single equation of Figure 14:

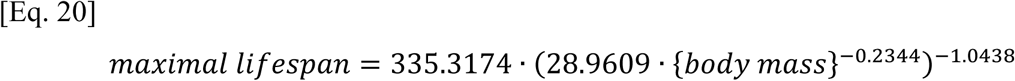

Where maximal lifespan is in years and body mass is in kg.

#### Predicted maximal lifespan by Eq. 20 versus Observed maximal lifespan

**Figure 17.**
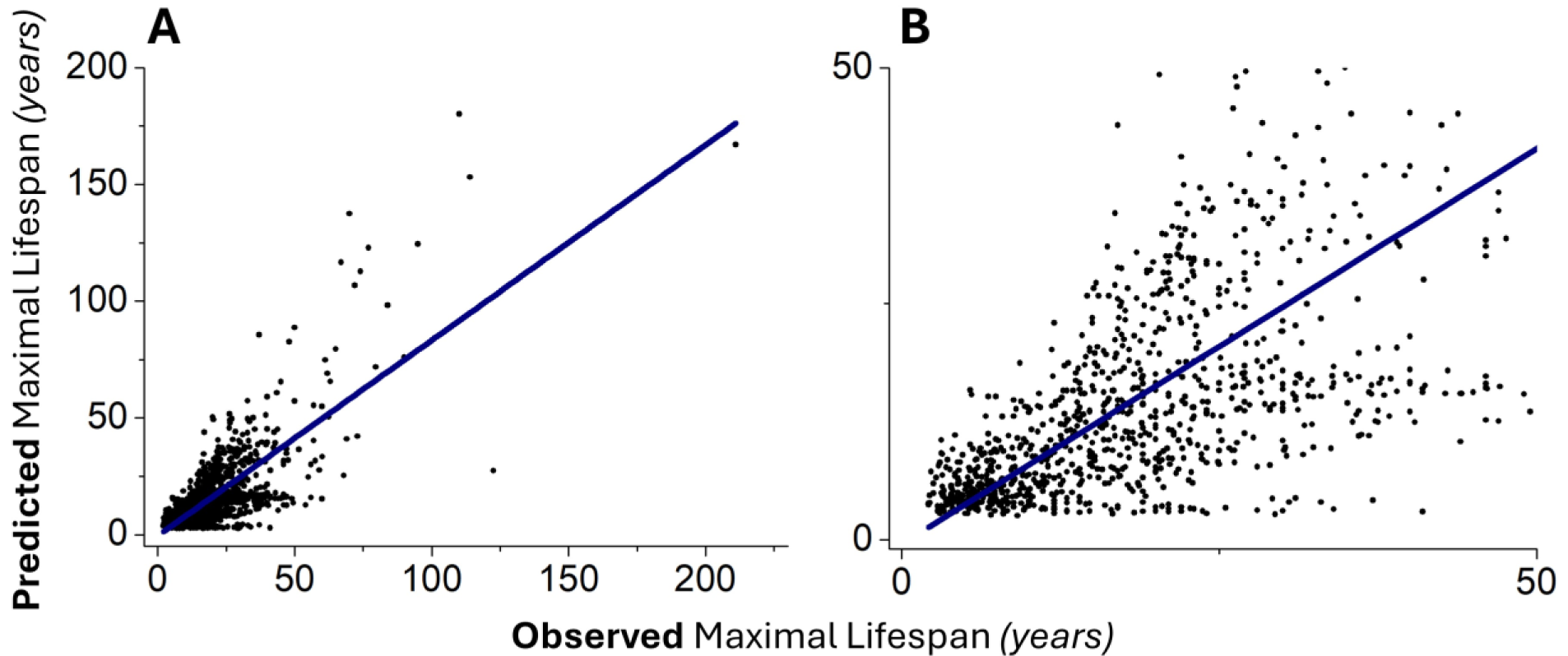
**(A)** Predicted maximal lifespan (years) by Eq. 20 plotted against observed maximal lifespan (years) for 1,023 mammal species. Testing this equation against a much larger species set than used to set its parameters. Adult body mass (input to Eq. 20) and maximal lifespan data sourced from [1]. All mammal species with such data available in [1] were used. Best-fit line slope = 0.83719 (±0.02298), intercept = -0.43544 (±0.57985), adjusted R^2^ = 0.56484, Root Mean Squared Error (RMSE) = 11.47562 years, Mean Absolute Error (MAE) = 8.25213 years. Pearson correlation coefficient = 0.7518, one-tailed (alternative hypothesis is directional) p-value = **3.6776*10^-187^**. *Omitting human, by rationale in the Methods, increases slope to 0.86932 (±0.02298) with 0.9 (i.e., close to optimal 1) within its standard error, and Pearson correlation coefficient to 0.7642 (not shown).* So, good out-of-sample prediction, conferring further statistical basis to Figure 14. **(B)** Part of (A) at higher resolution. At least some of the dispersion is likely due to noise in observed maximal lifespans, as most of these values are from extremely small sample sizes.

#### Linear regression of maximal lifespan on *λ* has a high adjusted R^2^ value of 0.841

In JASP (version 0.18.1.0), using data of Figure 15B, linear regression was performed, with the response variable being maximal lifespan (years) and the predictor variable being *λ*. Both logged (log_10_). Human included.

R^2^ = 0.847, adjusted R^2^ = 0.841, Unstandardized coefficients: β_0_ (intercept) = 2.344 ± 0.094 (p-value = 3.796e-18), β_1_ = -0.847 ± 0.075 (p-value = 7.327e-11). *Standardized coefficient: β_1_ = -0.920. Durbin-Watson statistic = 1.431. F (1, 23) = 127.577, p = 7.327e-11. Assumptions of homoscedasticity, normality, and linearity unviolated*.

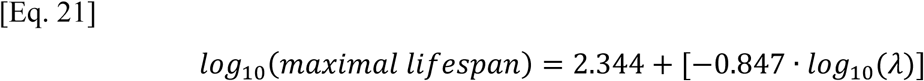

Pearson correlation coefficient between log_10_ maximal lifespan and log_10_ *λ* = **-0.9205**, one-tailed (alternative hypothesis is directional) p-value = 3.64435E-11.

If log_10_ adult body mass (kg) is added as a second predictor, the adjusted R^2^ goes down (to 0.833), wherein log_10_ *λ* is a highly significant predictor and log_10_ adult body mass is *not*, with unstandardized regression coefficient p-values of 8.028e-8 and 0.918 respectively (VIF = 2.018). So, even with adult body mass included in the model, *λ* is a significant predictor of maximal lifespan.

For log_10_ maximal lifespan on log_10_ *λ*, statistically excluding the effect of log_10_ adult body mass, the partial and semi-partial (part) correlation coefficients are -0.859 and -0.654 respectively. For log_10_ maximal lifespan on log_10_ adult body mass, statistically excluding the effect of log_10_ *λ*, the partial and semi-partial (part) correlation coefficients are -0.022 and - 0.009 respectively.

Further information, and further such analysis (e.g., using data from more species), is in the Supplementary Material. For example, showing (across species) that maximal lifespan *significantly* correlates with *λ* when the effects of phylogeny and adult body mass are both statistically excluded. For further example, presenting a Generalized Additive Model (GAM) of maximal lifespan, with *λ* and adult body mass as predictors, which has an adjusted R^2^ of **0.943**, with **95.4%** deviance explained.

#### Instrumental variable analysis reports causality between lower *λ* and higher maximal lifespan

Shown in the Supplementary Material, instrumental variable analysis reports causality between lower *λ* and higher maximal lifespan, at *incredible* statistical significance. With a p-value smaller than the smallest reportable by the statistical software used.

### New experimental data

#### F_1_F_0_ ATP hydrolysis is a new cancer drug target, and F_1_F_0_ ATP hydrolysis inhibiting drugs exert potent anticancer activity

I’ve discovered that cancers disproportionally use F_1_F_0_ ATP hydrolysis. Indeed, distinctively from normal cells, they absolutely require it. Not for heat generation (which isn’t an absolute need because exogenous heat {i.e., the ambient temperature} can substitute for its absence). But for operating their distinctive, abnormal, heavily glycolytic (“Warburg effect”) metabolism (some literature upon this metabolism: [133–138]). Indeed, I show that drugs, which inhibit F_1_F_0_ ATP hydrolysis, each exert potent anticancer activity (against 60 different cancers from 9 different tissues) *in vitro*: Figures 18 and 19. With analysis proving that this anticancer action is by inhibiting F_1_F_0_ ATP hydrolysis.

These cancer drugs might (unlike present cancer drugs) help rather than harm normal cells. Indeed, by making their metabolism more efficient (less chemical energy of food dissipated as heat), each of these drugs may combat cachexia (wasting/weight loss). Cachexia is prevalent in advanced cancer patients and is often the cause of death [139]. In support, transgenic mice with less metabolic heat generation have greater body weight (and longer lifespan) [21–22]. Incidentally, aging can also cause wasting (e.g., sarcopenia, frailty, etc.), especially in later life.

Figures 18 and 19 show results from the NCI-60 one-dose *in vitro* assay. Performed by the National Cancer Institute (NCI, USA). With compounds I submitted to them. This protocol tests the effect, if any, of a single test compound (10 µM) on the growth/survivability of 60 different cancer cell lines, originated from 9 different tissues, as compared to the no compound control [140]. A tested compound’s activity can range from negative % (cancer growth promotion), to 0% (no activity), to 100% (complete cancer growth inhibition), to 200% (all starting cancer cells are dead). NCI-60 tests are performed at a controlled temperature of 37°C. For 48 hours.

More specifically, Figures 18 and 19 show that compounds which potently inhibit F_1_F_0_ ATP hydrolysis exert potent anti-cancer activity. Figure 18 shows the *in vitro* anti-cancer activity of separated stereoisomers 6a and 6b. Figure 18A shows their structure. 6a is the *R* stereoisomer in high enantiomeric excess (>97% ee). 6b is the *S* stereoisomer in high enantiomeric excess (>97% ee; administered to mice in Figure 2). Figure 18B shows the anti-cancer activity of 6a (10 µM). Figure 18C shows the anti-cancer activity of 6b (10 µM).

As specified in Figure 18A, 6b potently inhibits F_1_F_0_ ATP hydrolysis (EC_50_ F_1_F_0_ ATP hydrolysis = 0.018 µM [=18 nM] in a Sub-Mitochondrial Particle {SMP} assay {in which *<u>no</u>* inhibition of F_1_F_0_ ATP synthesis by this drug was observed} [63–64]). Whilst 6a does not (EC_50_ F_1_F_0_ ATP hydrolysis > 100 µM in the SMP assay {in which *<u>no</u>* inhibition of F_1_F_0_ ATP synthesis by this drug was observed} [63–64]). In other words, the *S* stereoisomer potently inhibits F_1_F_0_ ATP hydrolysis, and the *R* stereoisomer does not. But the anti-cancer activity of 6a and 6b is similar in Figures 18B and 18C. Because, to interpret, they both undergo racemization in a biological system. Which erodes their enantiomeric excess during the 48 hours duration of the NCI-60 anti-cancer tests. So, they both converge towards/upon being the racemate, 19a (EC_50_ F_1_F_0_ ATP hydrolysis = 0.033 µM in the SMP assay [63–64], Figure 18A). Such that both samples ultimately contain a substantial proportion of *S* stereoisomer. And so, both exert anti-cancer activity by inhibiting F_1_F_0_ ATP hydrolysis. But racemization is not instantaneous, and so the 6b sample confers greater *S* stereoisomer exposure (“area under the curve”) to the cancer cells than the 6a sample. And thence confers greater anti-cancer activity: 66% vs. 57% mean (67% vs. 59% median) cancer growth inhibition, across all 60 cancer cell lines.

Observable in (aforementioned) Figure 18a, opposite stereoisomers, 6a and 6b, have hydrogen on their chiral carbon. Figure 19 discloses the anti-cancer activity of stereoisomers 7a and 7b. These have the same structure as 6a and 6b, except that they have deuterium (enrichment) instead of hydrogen on their chiral carbon (>99% molar percent deuterium incorporation at their chiral carbon). Their structure is shown in Figure 19A. 7a is the *R* stereoisomer in high enantiomeric excess (>97% ee). 7b is the *S* stereoisomer in high enantiomeric excess (>97% ee). Figure 19B shows the anti-cancer activity of 7a (10 µM). Figure 19C shows the anti-cancer activity of 7b (10 µM). This anti-cancer data is also from the National Cancer Institute’s standardized NCI-60 testing protocol. So, directly comparable to the (aforementioned) anti-cancer data for 6a and 6b. To summarize all the NCI-60 data, the mean and median % cancer growth inhibition conferred (by 10 µM) is shown in Table 25 below:

**Table 25.**

|  | 6a | 6b | (6b-6a) | 7a | 7b | (7b-7a) | (7b-7a)/(6b-6a) |
| --- | --- | --- | --- | --- | --- | --- | --- |
| Mean | 57.3 | 66.15 | 8.85 | 52.35 | 72.81 | 20.46 | 2.3 |
| Median | 58.62 | 66.9 | 8.28 | 52.09 | 74.51 | 22.42 | 2.7 |

7b exerts greater anti-cancer activity than 6b. 7a exerts less anti-cancer activity than 6a. Thence the difference between the anti-cancer activity of 7b and 7a is greater than that between 6b and 6a. To interpret, this is because, whereas 6a and 6b have hydrogen (H) attached to the chiral carbon, 7a and 7b have deuterium (D, ^2^H) attached to the chiral carbon. Deuterium slows the racemization rate by a Kinetic Isotope Effect (KIE [141]; C-^2^H bond is stronger than a C-H bond). So, because of a slower racemization rate, 7b maintains its enantiomeric excess (of *S*) better than 6b. Conferring 7b greater anti-cancer activity than 6b. Because of a slower racemization rate, 7a maintains its enantiomeric excess (of *R*) better than 6a. Conferring 7a less anti-cancer activity than 6a. The disparity in anti-cancer activity between 7b and 7a is 2-3 times greater than that between 6b and 6a. Which is the correct order of magnitude for a KIE (>1 and ≤10 [142], greater if tunnelling is very mechanistically relevant [143]).

If the *S* stereoisomer has anti-cancer activity, and the *R* stereoisomer does not, hypothesizing their enantiomerization in a biological system, the mean of 6a and 6b anti-cancer activity should equal the anti-cancer activity of the 6a/6b racemate. If this racemate was tested. Similarly, the mean of 7a and 7b anti-cancer activity should equal the anti-cancer activity of the 7a/7b racemate. If this racemate was tested. Supportively, the mean of 6a and 6b median anti-cancer activity is equal to the mean of 7a and 7b median anti-cancer activity: numbers below are drawn from earlier Table 25:

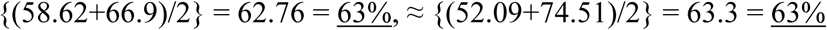

(*The mean of 6a and 6b mean anti-cancer activity is nearly equal to the mean of 7a and 7b mean anti-cancer activity: {(57.3+66.15)/2} = 61.73 = 62%, ≈ {(52.35+72.81)/2} = 62.58 = 63%*).

The following features of the data have already been herein addressed/explained, wherein each following sample name refers to the median of its anti-cancer activity (sourced from earlier Table 25): (7a+7b)/2=(6a+6b)/2, 7b>7a, 6b>6a, 7b>6b, 7a<6a. In NCI-60 compound testing, anti-cancer activity can range from 0% (no activity) to 100% (complete cancer growth inhibition) to 200% (all starting cancer cells are dead). So, the possible range is 0-200% (*if* the assumption is made that no administered compound will promote cancer growth, and so *if* the possibility of increased cancer growth upon compound administration is dismissed). To do some supporting theoretical analysis, using A, B, C, D, wherein each of these letters can independently be any integer in the range 0-200, the chance, by chance, of all of the following being true at the same time: (A+B)/2=(C+D)/2 (equivalently A+B=C+D), A>B, C>D, A>C, B<D, is 0.04% (125 times smaller than 5%, the most common significance threshold used by those of the art; its corresponding decimal p-value is 0.0004, which is <0.05). If A, B, C, D can be fractional also (i.e., not only integers) the chance is even smaller, much smaller (not shown). The chance, by chance, of all the following being true at the same time (delimited to integers): (A+B)/2=(C+D)/2=63, A>B, C>D, A>C, B<D, is 0.00025% (corresponding decimal p-value is 0.0000025, which is <0.05).

The anti-cancer activity of 6a, 6b, 7a, 7b are all highly correlated (in Table 26 below, p-values are one-tailed because each alternative hypothesis is directional). Because they tend to have greater, or lesser, anti-cancer activity against the same of the 60 cancer cell lines. This, by the rationale/basis of [144], indicates that they are exerting anti-cancer activity by the same molecular mechanism. Indeed, to interpret the data herein, by selectively inhibiting F_1_F_0_ ATP hydrolysis (*without inhibition of F_1_F_0_ ATP synthesis*).

**Table 26.**

| Pearson Correlation Coefficient (R): |  |  |  |  |
| --- | --- | --- | --- | --- |
|  | 6a | 6b | 7a | 7b |
| 6a | 1 |  |  |  |
| 6b | 0.7991 | 1 |  |  |
| 7a | 0.7693 | 0.5205 | 1 |  |
| 7b | 0.6614 | 0.6451 | 0.8049 | 1 |
| *N=60 for all samples; P-value (one-tailed):* |  |  |  |  |
|  | 6a | 6b | 7a | 7b |
| 6a | / |  |  |  |
| 6b | 9.85E-15 | / |  |  |
| 7a | 3.48E-13 | 1.01E-05 | / |  |
| 7b | 4.39E-09 | 1.32E-08 | 4.60E-15 | / |
| Fisher's combined probability test |  |  |  |  |
| Using p-values for [6a vs. 7a] and [6b vs. 7b] |  |  |  |  |
|  |  | 3.48E-13 | 1.32E-08 |  |
| p-value = **2.20E-19** = 0.0000000000000000002 |  |  |  |  |

So, Figures 18 and 19 show that compounds which potently inhibit F_1_F_0_ ATP hydrolysis exert potent anti-cancer activity. I predicted this result before it was observed. By rational design. Because I reasoned that the abnormally glycolytic metabolism of cancers (Warburg effect [133]), especially used by the most dangerous thereof [134–138], hinges upon abnormally high rates of F_1_F_0_ ATP hydrolysis. Consuming glycolytic ATP (imported into mitochondria by the Adenine Nucleotide Transporter, driven by higher [ATP], and lower [ADP], in the cytoplasm than in the mitochondrial matrix). Releasing glycolysis from ATP negative feedback inhibition (ATP allosterically exerts negative feedback inhibition upon the glycolytic enzymes phosphofructokinase and pyruvate kinase [145]). Yielding higher glycolytic rate. Thence more glycolytic intermediates are available to be shunted into biosynthesis (enabling faster cancer cell proliferation). And conferring more NADPH produced by the pentose phosphate pathway, enabling more Reactive Oxygen Species (ROS) mitigation. By the NADPH dependent glutathione, and NADPH dependent thioredoxin, ROS mitigation systems. And less ROS are produced because oxidative phosphorylation {OXPHOS} is disfavoured by the high proton motive force (pmf, comprising high {more hyperpolarized} Ψ_IM_ [54]) across the mitochondrial inner membrane created by F_1_F_0_ ATP hydrolysis. So, [ROS] is lower. Freeing them from aging, thereby conferring “limitless replicative potential” (i.e. immortality, which is distinctive trait of cancer [146], which normal somatic cells don’t share; {I propose} mutational load of cancer primarily acquired before, on the path to, cancer).

A selective F_1_F_0_ ATP hydrolysis inhibiting drug (*that doesn’t inhibit F_1_F_0_ ATP synthesis*) inhibits this cancer metabolic program, conferring anti-cancer activity. Whilst (prospectively) helping, instead of harming, normal cells *in vivo*. Increasing their metabolic efficiency, so less chemical energy of food is dissipated as heat (Figure 2). Thereby (*if* the ambient temperature, and/or body insulation, substitutes for lower metabolic heat production) possibly combating cachexia/wasting. Which is observed in many advanced cancer patients, often being the cause of their death [139].

Moreover, *in vivo*, to hypothesize, cachexia/wasting might be additionally combated by the compound conferred decrease in glycolytic rate of cancer cells. Decreasing their glucose influx and lactate efflux. Thence decreasing the energetically expensive conversion of lactate to glucose by the Cori cycle of the liver. Which costs 6 ATP for every lactate to glucose converted [145]. Elevated blood [lactate] correlates with worse cancer prognosis [147]. In this way, the anti-cancer benefit *in vivo* is expected to be greater than that shown herein *in vitro*. Especially in the case of cancer cells in a solid (hypoxic/anoxic) tumour, which might singly rely upon F_1_F_0_ ATP hydrolysis to maintain pmf (Ψ_IM_). Where loss of Ψ_IM_ is a well-known trigger for apoptosis [148]. The *in vitro* studies herein cannot report this (hypothesized) effect because they are conducted at atmospheric pO_2_ (at the elevation of the National Cancer Institute in Maryland, USA). Tumour hypoxia correlates with poor patient prognosis [149–154]. Tumour hypoxia is associated with radio- [156–157] and chemo- [157–158] resistant cancer.

In further supporting data, I’ve shown anti-cancer activity, in NCI-60 testing, by other compounds, of completely different scaffolds, which also selectively inhibit F_1_F_0_ ATP hydrolysis (*that don’t inhibit F_1_F_0_ ATP synthesis*). For example, please see the 45 pages of experimental data in my international (PCT) patent filing [18] (following my earlier PCT [19]) and further experimental data in my PCT [20]. My work, predominantly in the patent (rather than scholarly) literature, reports a new molecular target for exerting anti-cancer activity. F_1_F_0_ ATP hydrolysis. And novel drugs acting upon that target.

It is notable that selective F_1_F_0_ ATP hydrolysis inhibiting compounds (*that don’t inhibit F_1_F_0_ ATP synthesis*), hypothesized to slow aging herein, have anti-cancer activity (Figures 18 and 19). Because a worry/critique of trying to slow aging is the argument that, because cancer is immortal [146], conferring slower aging/immortality upon a normal cell is making it closer to being cancer. Such that even if an anti-aging approach could be found, because of it inextricably increasing the cancer risk, lifespan wouldn’t be very much extended, if at all. Conferring a practical block/limit to appreciable lifespan extension. For example, this is a concern, in at least some people’s minds, with expression of telomerase (in normal somatic cells) approaches [159–161]. However, by contrast, selectively inhibiting F_1_F_0_ ATP hydrolysis (*without inhibiting F_1_F_0_ ATP synthesis*) actively exerts potent anti-cancer activity (Figures 18 and 19). So, distinctively, it is a drive against, and not to, cancer. Arguably, anti-cancer activity is a highly desirable, if not a mandatory, trait of a longevity drug. Indeed, rapamycin (also known as sirolimus), an mTOR inhibiting drug, which can slow aging and extend lifespan in mice, has anti-cancer activity also [162]. In NCI-60 one-dose (10 µM) testing, it confers mean and median % cancer growth inhibition of 56.22% and 55.45% respectively (NSC 226080 in [163]). Temsirolimus, a prodrug of rapamycin (metabolized to rapamycin in the body), is approved by the U.S. Food and Drug Administration (FDA), and the European Medicines Agency (EMA), for the treatment of Renal Cell Carcinoma (RCC) [164].

At least some present cancer treatments (e.g., radiotherapy; probably because their Mechanism of Action {MOA} is, at least in part, to increase Reactive Oxygen Species, ROS [155–158]) accelerate aging in normal cells. Thence (presumably because aging/age is a major risk factor for cancer [165]) increasing subsequent cancer risk [166–167]. I propose selective F_1_F_0_ ATP hydrolysis inhibitor compounds do the inverse. Still conferring anti-cancer therapy (Figures 18 and 19. Partly, by aforementioned rationale, via increasing [ROS] in cancer cells). But whilst slowing aging in normal cells (by reducing their intracellular [ROS], Figure 5), which reduces the subsequent/secondary cancer risk.

To theorize, inhibiting F_1_F_0_ ATP hydrolysis slows both heavily glycolytic (cancer) and predominantly oxidative (normal cell) metabolisms. The former *increases* intracellular [ROS] in cancer cells. The latter *decreases* intracellular [ROS] in normal cells.

Note that cancer can increase body temperature [168]. Indeed, cancers can cause fever [169]. Especially some types thereof, such as Hodgkin’s and non-Hodgkin’s lymphoma, renal cell carcinoma, hepatocellular carcinoma, Acute Myeloid Leukaemia (AML), hairy cell leukaemia, glioblastoma multiforme, blast crisis of Chronic Myelogenous Leukemia (CML), and ovarian cancer. At low/moderate ambient temperature, this higher body temperature confers margin/benefit for a greater dose of a selective F_1_F_0_ ATP hydrolysis inhibiting drug (*that doesn’t inhibit F_1_F_0_ ATP synthesis*).

Compound 7b has the best anticancer activity. Conceivably, its anticancer utility confers margin to follow a *relatively* fast regulatory path to its clinical approval in humans. Afterward used/trialled as a treatment/preventative for other diseases of aging. Those many varied diseases whose incidence increases exponentially with age/aging. Such as Age-related Macular Degeneration (AMD) and Alzheimer’s disease. If aging is causal to age-related diseases, (conceivably) a single drug, which slows aging (predicted of compound 7b), can confer therapy for all these (many and varied) diseases.

So, here might be a drug class with anticancer activity but instead of harming normal cells, as present cancer drugs do, when appropriately used, it helps normal cells by making their metabolism more efficient (less chemical energy of food dissipated as heat), slowing their aging, and combating cachexia.

## Supplementary Material

Please don’t overlook this paper’s **>300 pages** of Supplementary Material, comprising Supplementary Results, Discussion, and Methods.

## DISCUSSION

### Discovery of the keystone of homeothermy

A contribution of this paper (*and its earlier, sister patent applications: e.g.,* [18–20]) is the new finding, from new experimental data, that F_1_F_0_ ATP hydrolysis is prevalent *in vivo*, and that it confers metabolic heat generation. It is the principal chemical reaction that homeotherms use for metabolic heat generation.

#### IF1 protein as the reason why different mammal species have different maximal lifespans

Mammals (and birds) are endothermic and metabolically generate heat to maintain their body temperature. In Euclidean geometry, smaller objects have a larger surface-area (A) to volume (V) ratio than larger objects. Because, where L is length, A∝L^2^ and V∝L^3^ (“square-cube law”). So, smaller mammal species have a larger surface-area-to-volume ratio. And so, they lose a greater fraction of their metabolically generated heat. Thus, they must produce more heat per unit mass per unit time, requiring them to have a greater metabolic rate per unit mass (serviced by a greater heart rate). Therefore, generating more Reactive Oxygen Species (ROS) per unit mass, per unit time. Accumulating ROS caused molecular damage faster. Interpreted to explain why smaller mammal species tend to age faster and have shorter maximal lifespans.

How is the metabolic rate per unit mass of different sized mammal species set differently? By the 2^nd^ Law of Thermodynamics, whenever energy converts from one form to another, some of this energy must be dissipated as heat (no energy conversion can be 100% efficient). I’ve discovered that mammals cyclically synthesize and hydrolyse ATP (F_1_F_0_ ATP synthesis and F_1_F_0_ ATP hydrolysis respectively). Conditional upon passing, and pumping, protons along their concentration gradient across the inner mitochondrial membrane, respectively. So, cyclically interconverting between chemical and potential energies. Which (by the inefficiency of energy conversions) generates heat to maintain body temperature.

F_1_F_0_ ATP synthesis exceeds F_1_F_0_ ATP hydrolysis, conferring net ATP production. Especially since ATP is consumed by cellular ATP demand, denying it to F_1_F_0_ ATP hydrolysis, and because IF1 protein only inhibits F_1_F_0_ ATP hydrolysis [8–14]. Wherein the amount of F_1_F_0_ ATP hydrolysis (amount of heat generated) is constrained by the amount of IF1 protein activity.

Per unit mass, per unit time, smaller (shorter lifespan) species run this “futile” (merely heat-generating) cycle more than larger (longer lifespan) species (*Figure 3*). Because they have less specific IF1 protein activity (*Supplementary Material*). Making IF1 protein a molecular determinant of lifespan (*interpretation*).

This futile cycle also generates heat by its increase of proton leak across the inner mitochondrial membrane (*Supplementary Material section headed “Numerical prediction observed”*). Because it pumps protons into the mitochondrial intermembrane space. Increasing the chance that protons cross the inner mitochondrial membrane, into the mitochondrial matrix, outside of ATP synthase. Dissipating their potential energy as heat.

### Mutually reinforcing support

This novel account is supported by mouse data of Figure 2. Which shows that F_1_F_0_ ATP hydrolysis is a causal input to metabolic heat generation, and so metabolic rate. Further mouse data shows that F_1_F_0_ ATP hydrolysis is a causal input to ROS generation (Figure 5).

At incredible statistical significance (asymptotically exact harmonic mean p-value of its constituent correlations = 1e-119), Figure 14 schematizes disparate experimental data from 353 mammal species to teach the following. Across mammal species, adult body mass (by the square-cube law of geometry) sets body surface-area per unit body mass, and thence metabolic heat generation per unit mass per unit time required, and therefore performed, which is set by specific IF1 protein activity determining specific F_1_F_0_ ATP hydrolysis rate, dictating specific basal metabolic rate, therefore specific ROS generation rate, and thence specific ROS damage rate, which determines maximal lifespan.

Where such ROS damage is varied, and includes epigenetic damage, as measured by the epigenetic clock. Instrumental variable analysis reports causality between specific basal metabolic rate and epigenetic clock “ticking” rate (Table 21). Mediation analysis reports that body surface-area per unit body mass dictates epigenetic clock ticking rate, via dictating specific basal metabolic rate (Table 22). Mediation analysis reports that specific F_1_F_0_ ATP hydrolysis rate dictates epigenetic clock ticking rate, via dictating specific basal metabolic rate (Table 20). The high correlation between specific basal metabolic rate and epigenetic clock ticking rate endures when phylogeny is accounted for (Supplementary Material). Linear regression, multiple linear regression, partial and semi-partial correlations, generalized linear modelling, and generalized additive modelling report that specific basal metabolic rate is a highly significant predictor of epigenetic clock ticking rate, including in cases where adult body mass is considered (herein and the Supplementary Material). Observed collinearity in multiple linear regressions is consistent with adult body mass impacting epigenetic clock ticking rate because it dictates (by the square-cube law of geometry) body surface-area per unit body mass, which dictates (metabolic heat generation per unit mass per unit time required to maintain body temperature) specific basal metabolic rate (Eq. 10-15, Table 23).

Instrumental variable analysis reports causality between lower specific ROS damage rate, comprising different damage rates combined, including epigenetic damage rate, and higher maximal lifespan (Supplementary Material). Mediation analysis reports that lower specific basal metabolic rate causes higher maximal lifespan, by causing lower specific ROS damage rate (Supplementary Material). Mediation analysis reports that lower specific basal metabolic rate causes higher maximal lifespan, by causing lower epigenetic damage rate (Tables 20 and 22).

Collapsing Figure 14 into equations teaches combination variable, *λ*, which is incredibly (statistically) explanatory for why different mammal species have different maximal lifespans (Figure 15, Eq. 18, Eq. 21, Supplementary Material). Even when the effects of adult body mass and phylogeny are statistically excluded. A Generalized Additive Model (GAM), with *λ* and adult body mass as predictors of mammal species’ maximal lifespan, has an adjusted R^2^ of 0.943, with 95.4% of deviance explained. Across mammal species, mediation analysis reports that adult body mass dictates maximal lifespan indirectly via *λ*, with no direct effect. Instrumental variable analysis reports causality between lower *λ* and higher maximal lifespan. At incredible statistical significance (with p-value smaller than the smallest reportable by the statistical software used). Eq. 20, derived from *λ*, can significantly predict the maximal lifespan of 1023 mammal species, including many species whose data wasn’t used to set its parameters (Figure 17). So, good out-of-sample prediction, conferring further statistical basis to the schema of Figure 14.

Specific basal metabolic rate is componentry to *λ*. Across 517 mammal and bird species, maximal lifespan inversely correlates with specific basal metabolic rate at incredible statistical significance (Supplementary Material). In a Generalized Linear Model (GLM), and a Generalized Additive Model (GAM), across 517 mammal and bird species, specific basal metabolic rate is a highly significant predictor of maximal lifespan, including when adult body mass is included in the model (Supplementary Material). Observed collinearity in multiple linear regressions, and corresponding partial and semi-partial correlations, is consistent with adult body mass impacting maximal lifespan because it dictates (by the square-cube law of geometry) body surface-area per unit body mass, which dictates (metabolic heat generation per unit mass per unit time required to maintain body temperature) specific basal metabolic rate (Supplementary Material).

Aging rate is the rate of change of a correlate/biomarker of aging. For example, DNA (or epigenetic) damage rate. Maximal lifespan relates to aging rate by the same equation, Eq. 18, that it relates to specific F_1_F_0_ ATP hydrolysis rate. Consistent with specific F_1_F_0_ ATP hydrolysis rate dictating aging rate, and so maximal lifespan. Maximal lifespan relates to specific basal metabolic rate by this same equation. Consistent with specific F_1_F_0_ ATP hydrolysis rate dictating, via dictating specific basal metabolic rate, aging rate. By contrast, for example, maximal lifespan doesn’t relate to adult body mass by this equation.

Mediation analysis reports that lower specific F_1_F_0_ ATP hydrolysis rate causes higher maximal lifespan, by causing lower specific basal metabolic rate (Table 5). Instrumental variable analysis reports causality between lower specific F_1_F_0_ ATP hydrolysis rate and higher maximal lifespan (Table 10), and causality between lower specific basal metabolic rate and higher maximal lifespan (Table 9). Interventional mouse experiments report causality between lower F_1_F_0_ ATP hydrolysis rate and lower metabolic heat generation (lower metabolic rate; Figure 2), and causality between lower metabolic heat generation (lower metabolic rate) and longer lifespan [21–22]. Increased IF1 protein, and decreased F_1_F_0_ ATP hydrolysis, safely reduces a biomarker of aging in mice (Figure 5). Reduction of a biomarker of aging indicates reducing/reversing of aging [44].

Highly statistically significant Structural Equation Modelling (SEM) of the schema of Figure 14 supports it, and confers it further statistical basis, especially as this model can faithfully out-of-sample predict (Supplementary Material). It is a mathematical model of aging and lifespan, completely parameterized by experimental data. All its parameter’s p-values are incredibly small, most are smaller than the smallest reportable by the statistical software used. It can be, and is, used for *in silico* experiments. It demonstrates robustness and teaches in sensitivity and scenario analysis, using one-way and multi-way methods thereof. The model is a concise, molecular resolution, experimentally derived, experimentally predictive, mathematical theory of aging. Collapsing much complexity to - capturing much complexity in - a short chain of equations (Eq. S11 in the Supplementary Material).

Key experimental data used has been replicated across different studies: e.g., specific F_1_F_0_ ATP hydrolysis rate data and specific IF1 protein activity data (Supplementary Material). Reproduced, reproducible data.

Selected examples have been listed above. There is further cumulative, corroborating, convergent, and consilient evidence in this paper, and its Supplementary Material (of >300 pages).

Triangulation by convergent findings from different methods and data. The wealth of disparate experimental data (including interventional data and replicated data) used from >500 different species, sourced by different groups using diverse experimental methods, and correlations analysis, patterns analysis, mediations analysis, bootstrapping, instrumental variables analysis, Structural Equation Models (SEMs), sensitivity analysis, scenario analysis, linear regressions, multiple linear regressions, collinearities analysis, partial and semi-partial correlations analysis, hierarchical regressions, ridge regressions, multiple non-linear regressions, Generalized Linear Models (GLMs), Generalized Additive Models (GAMs), Phylogenetically Independent Contrasts (PIC; multiple methods thereof; multiple linear regressions, Principal Component Analysis [PCA], partial and semi-partial correlations analysis with phylogenetically adjusted data), consolidation of disparate data into single variable analysis (and thereof: GAMs, linear regressions, multiple linear regressions, partial and semi-partial correlations analysis, mediation analysis, instrumental variable analysis, PIC, multiple linear regressions and partial and semi-partial correlations analysis with phylogenetically adjusted data), numerical modelling, disparate data sharing the same equation form to maximal lifespan (Eq. 18), equations predicting data not used in setting their parameters (out-of-sample prediction), statistical significance in significance testing (including advanced thereof, such as Fisher’s combined probability test), predictions made and observed in further (independently conducted) experiments (Figures 2, 18-19), all support, in mutually reinforcing unison, this study’s conclusions.

**Figure 18.**
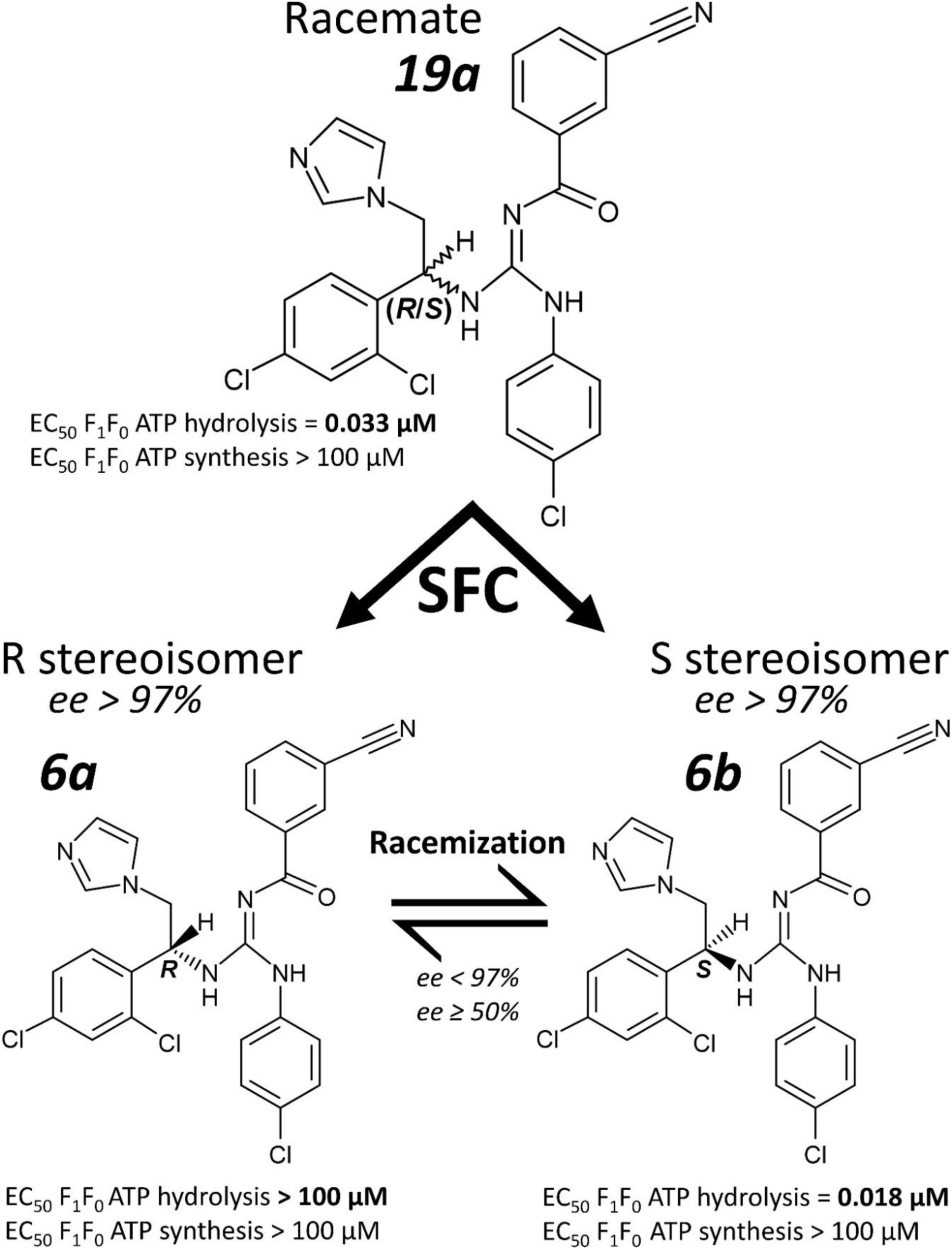
A Structure of racemate, 19a, and its separated stereoisomers (separated stereoisomer fractions) 6a and 6b. SFC is chiral supercritical fluid chromatography, ee is enantiomeric excess. EC_50_ values are from a Sub-Mitochondrial Particle (SMP) assay.

**Figure 18B.**
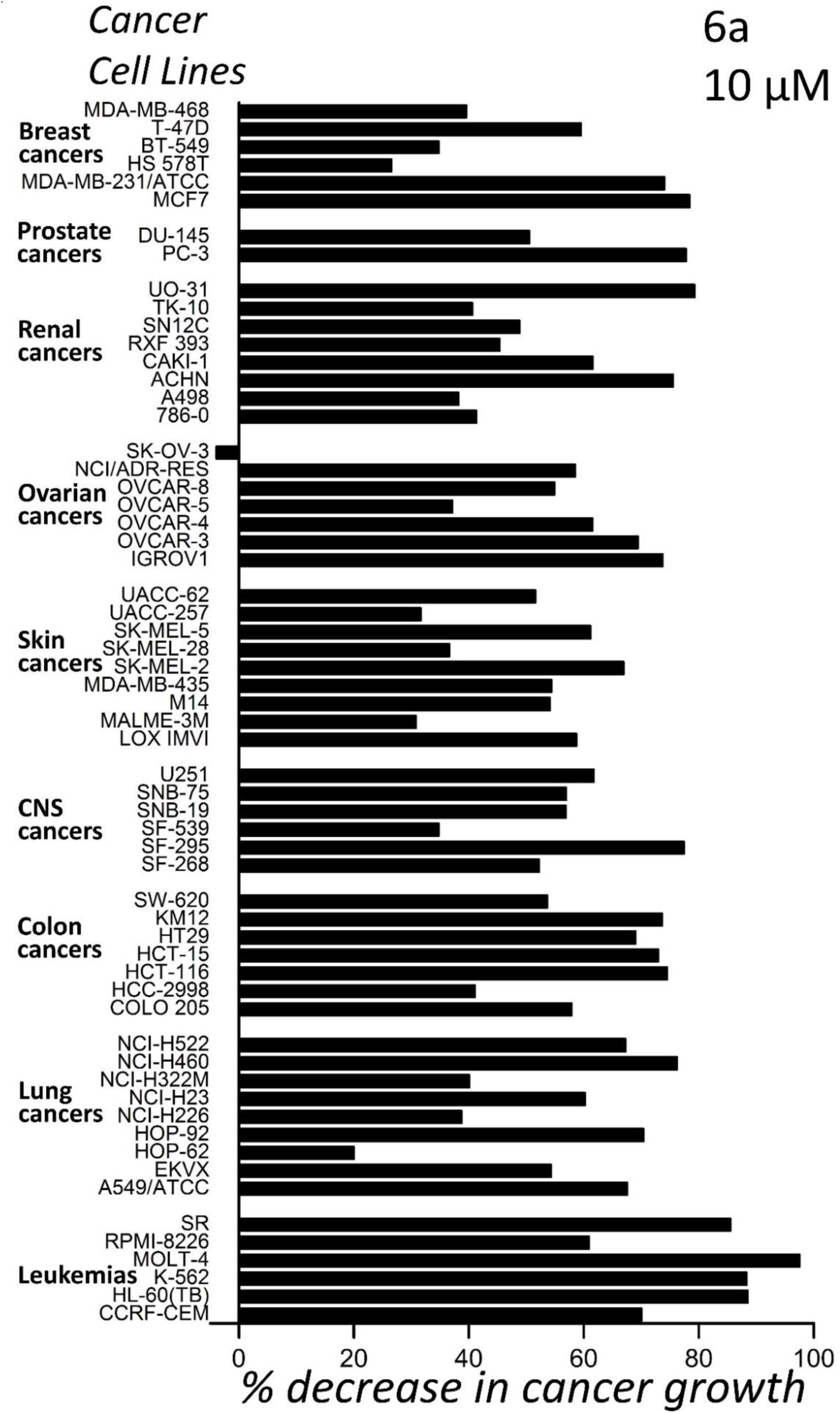
Anti-cancer activity of compound 6a (10 µM) in NCI-60 testing at the National Cancer Institute (NCI, USA).

**Figure 18C.**
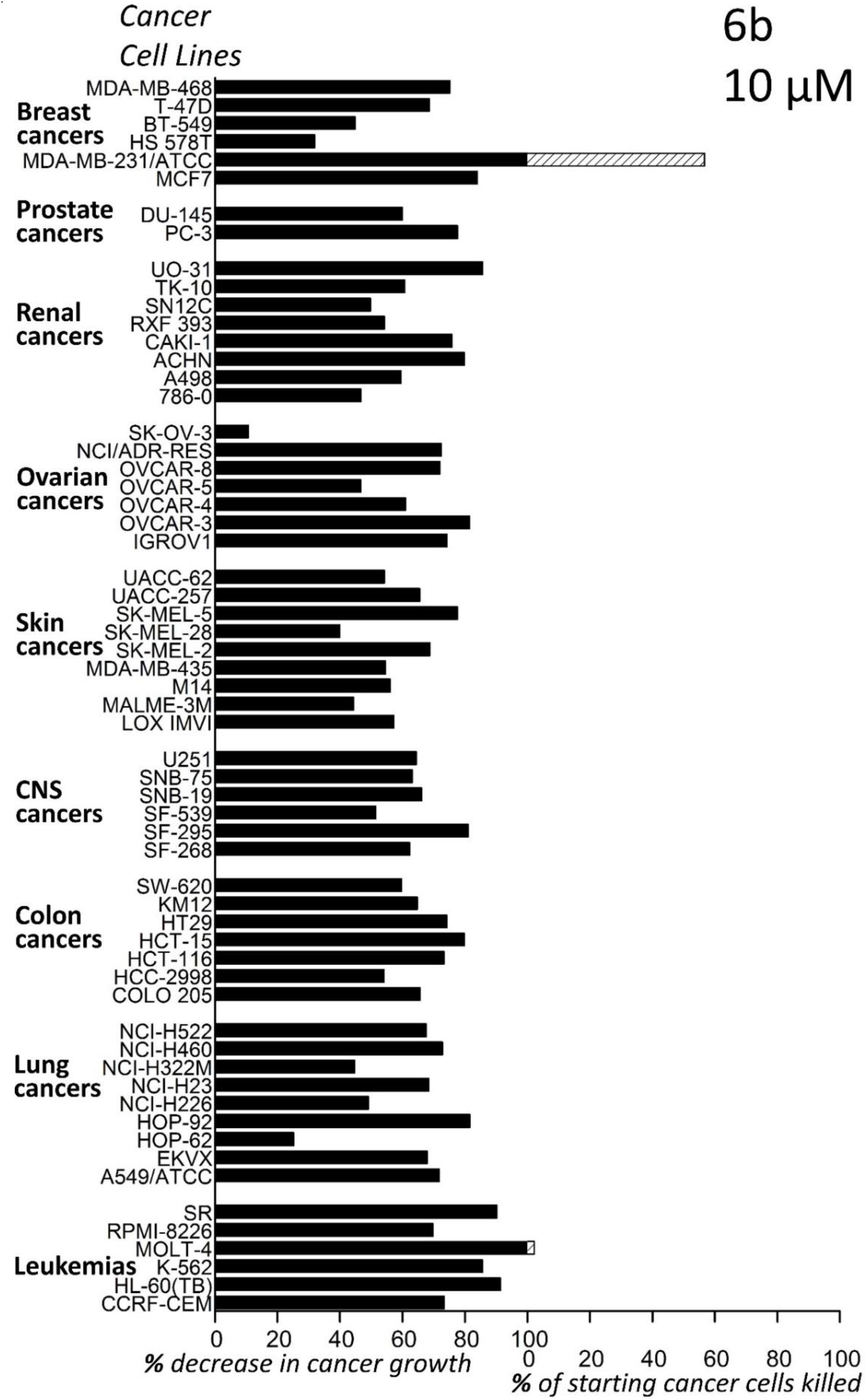
Anti-cancer activity of compound 6b (10 µM) in NCI-60 testing at the National Cancer Institute (NCI, USA).

The conclusions of this study align with the starting multi-pronged hypothesis of this study, which wasn’t generated from the data, it preceded obtaining the data, and so this data tests the hypothesis, validating the hypothesis. A confirmatory (hypothesis before data) rather than an exploratory (hypothesis from data) study. The chance that such a large amount of varied data, from different species, sources, and methods (including interventional data, including drug intervention data), and multifaceted analysis thereof, supports such a predefined multi-pronged hypothesis by chance is small.

### IF1 protein as the hourglass of life

Larger, longer-living mammals do less F_1_F_0_ ATP hydrolysis per unit mass per unit time, for a longer time (longer lifetime). Because they have less specific IF1 protein activity. Explaining why different mammal species have different maximal lifespans. Kingmaking IF1 protein, which selectively inhibits F_1_F_0_ ATP hydrolysis (*doesn’t inhibit F_1_F_0_ ATP synthesis*), as the molecular determinant of maximal lifespan. Predicting that a selective F_1_F_0_ ATP hydrolysis inhibitor drug (*which doesn’t inhibit F_1_F_0_ ATP synthesis*) slows aging. Such drugs are in hand [18–20].

### Prediction

Depending on

i. drug dose, and
ii. ambient temperature,

a systemically administered selective F_1_F_0_ ATP hydrolysis inhibitor drug (*that doesn’t inhibit F_1_F_0_ ATP synthesis*) can slow the subject’s aging by only the first, or both, of the following:

a. by decreasing specific F_1_F_0_ ATP hydrolysis rate (thereby decreasing specific basal metabolic rate, ROS generation rate, and ROS damage rate),
b. by slightly decreasing body temperature (as chemical reactions are slower at lower temperature: decreasing ROS generation and damage rate; decreasing ROS damage repair rate also, but as damage outruns repair, conferring *net* reduction of ROS damage rate).

*(a)* can occur in the absence of (*b*) (Figure 20). Or they can occur together, with (*a*) causing (*b*). That (*a*) can occur without (*b*) means a very high drug dose (conferring *much* slower aging) can be safely administered. Where higher ambient temperature permits more of (*a*) without (*b*). When ambient ≥ body temperature, (*a*) can be maximal and (*b*) cannot occur. An ambient temperature slightly below normal body temperature enables the maximum of (*a*), yet still with (*b*).

**Figure 19.**
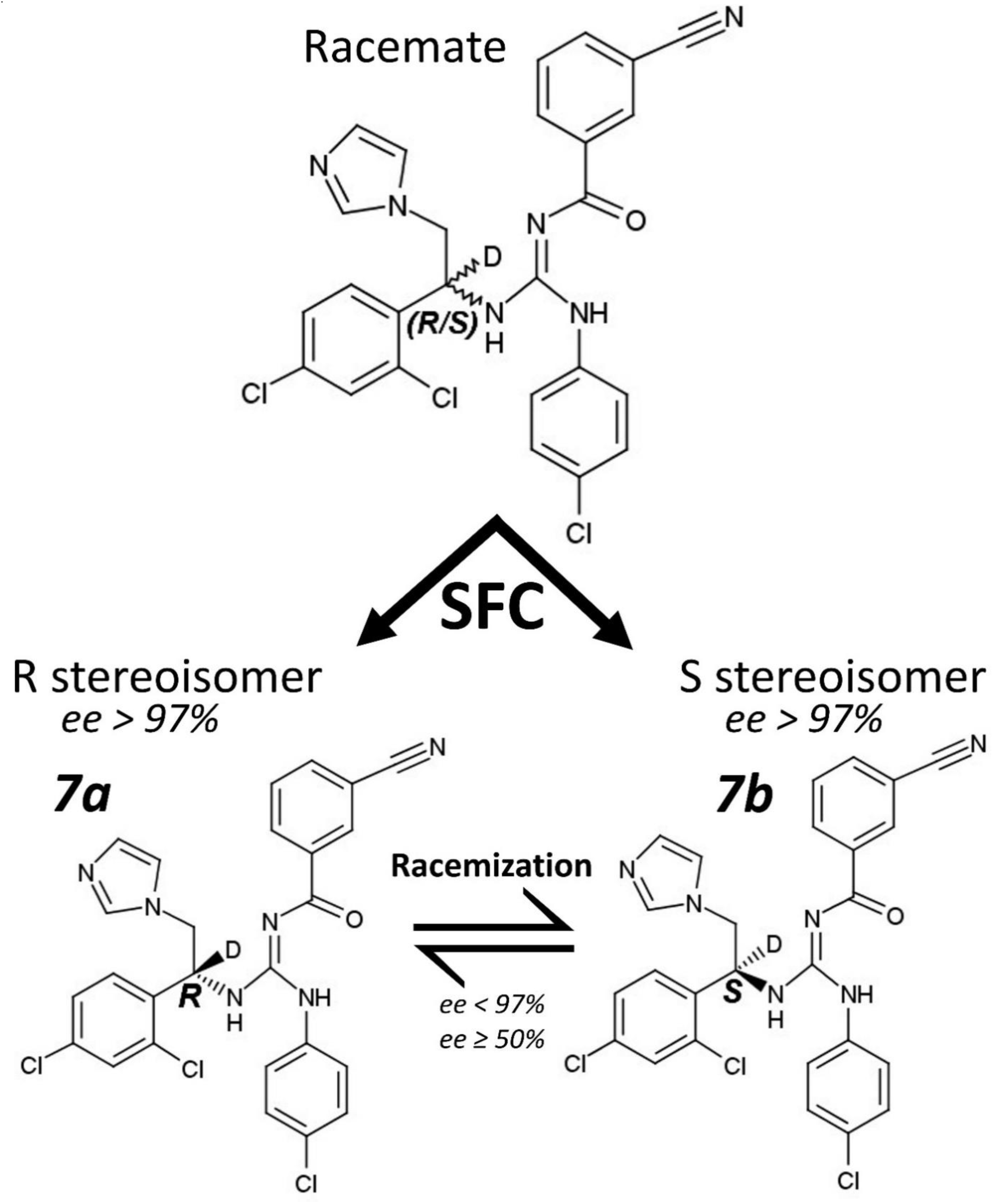
A Structure of separated stereoisomers (separated stereoisomer fractions) 7a and 7b. SFC is chiral supercritical fluid chromatography, ee is enantiomeric excess. D is deuterium (^2^H).

**Figure 19B.**
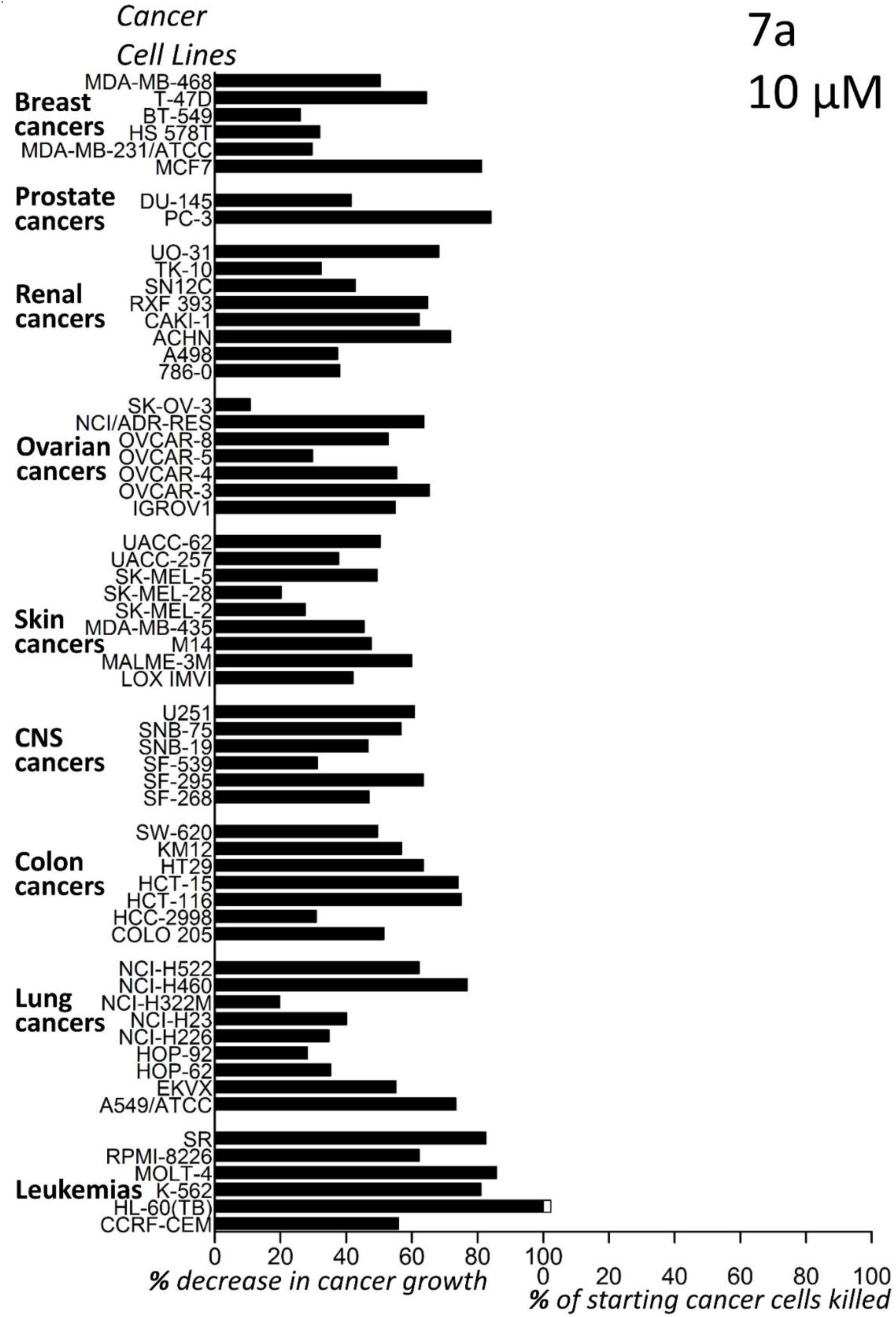
Anti-cancer activity of compound 7a (10 µM) in NCI-60 testing at the National Cancer Institute (NCI, USA).

**Figure 19C.**
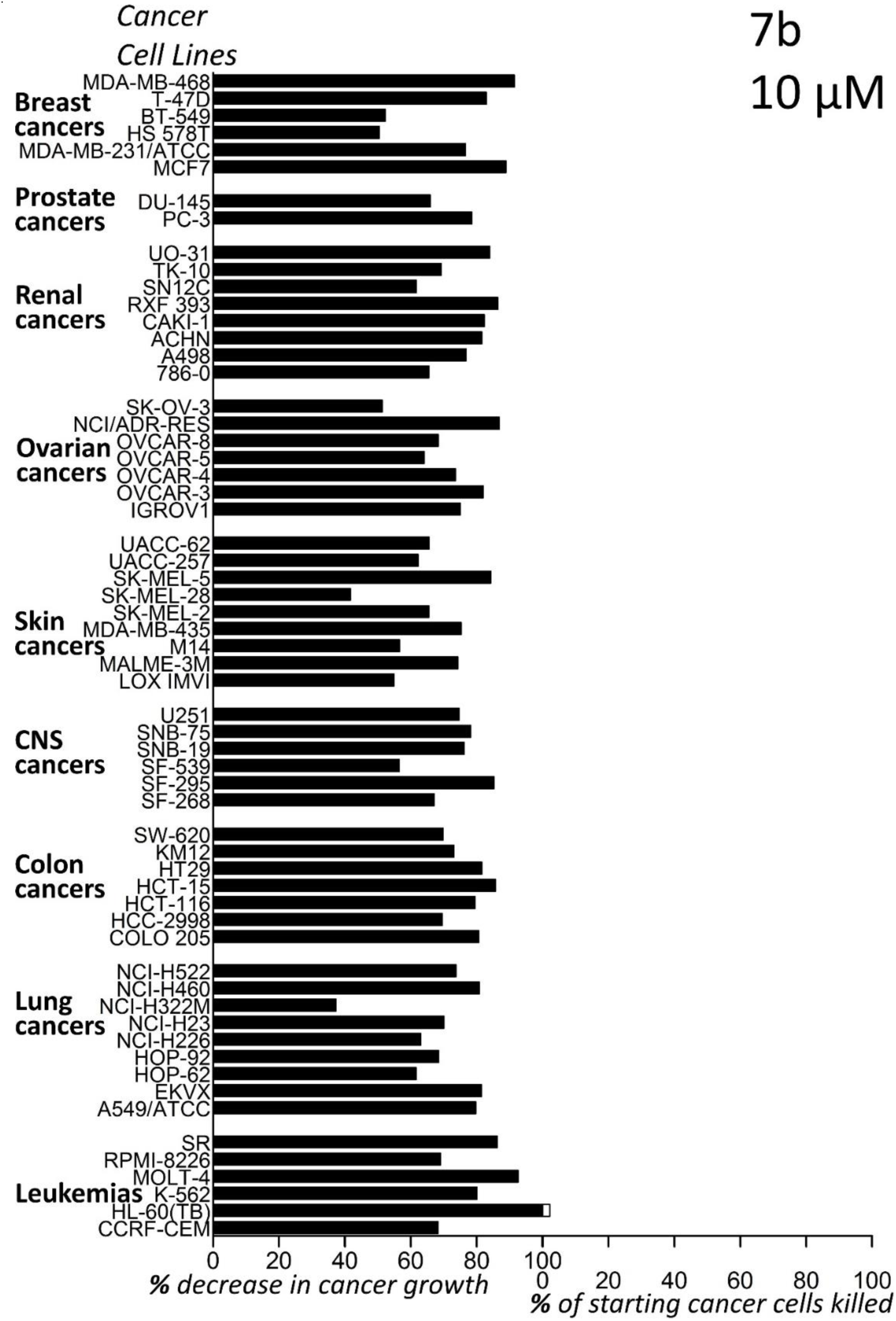
Anti-cancer activity of compound 7b (10 µM) in NCI-60 testing at the National Cancer Institute (NCI, USA).

**Figure 20.**
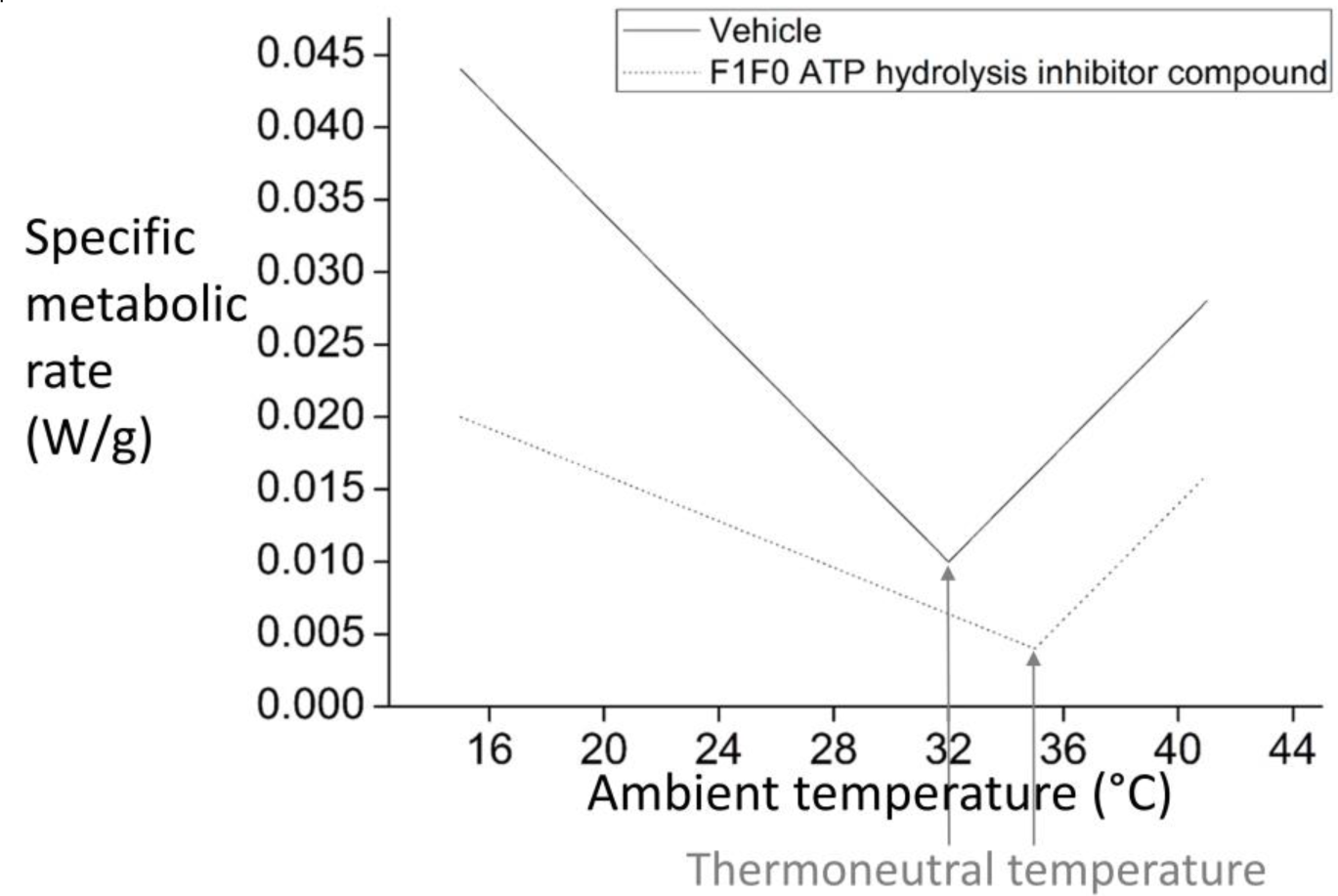
Diagram: Why systemic administration of a selective F_1_F_0_ ATP hydrolysis inhibitor drug (*that doesn’t inhibit F_1_F_0_ ATP synthesis*) doesn’t necessarily reduce body temperature. This diagram relates to a mouse. Mice have a natural body temperature of 37°C [1]. “Thermoneutral temperature” is the ambient temperature that the body is most comfortable at (*at which specific metabolic rate is the “specific basal metabolic rate”*). Above and below which, greater metabolic rate is required to maintain the body at 37°C [178]. At the mouse’s natural thermoneutral temperature of 32°C, its specific basal metabolic rate is 0.0092125 W/g [179] (*N.B. human thermoneutral temperature is less because humans are bigger, and moreover wear clothes. Thermoneutral temperature of a human in typical clothes is 20.3°C* [180]). Systemic administration of a selective F_1_F_0_ ATP hydrolysis inhibitor drug (*that doesn’t inhibit F_1_F_0_ ATP synthesis*) reduces the mouse’s specific basal metabolic rate and shifts its thermoneutral temperature higher. Illustratively to 35°C in this figure. Which makes the mouse more comfortable (lower metabolic rate) at equal or higher ambient temperatures. Furthermore, this figure anticipates that F_1_F_0_ ATP hydrolysis is integral to thermogenic metabolic rate, in addition to basal metabolic rate. And so, the gradient of the thermogenic metabolic rate increase is shallower, because of reduced F_1_F_0_ ATP hydrolysis. The specific basal metabolic rate at thermoneutral temperature = 35°C was selected by drawing a line from the specific basal metabolic rate at thermoneutral temperature = 32°C, = 0.0092125 W/g, which is an experimental data point from [179], to 37°C on the x-axis (thermoneutral temperature = 37°C, specific basal metabolic rate = 0 W/g) and selecting the corresponding specific basal metabolic rate for 35°C on this line (the equation of this line is *y* = *a* + *b* · *x* = 0.06817 + −0.00184 · *x*). Accordingly, the specific basal metabolic rate was 60% lower. And, in concordance, the gradient of the thermogenic plot was reduced by 60% also. Anticipating that F_1_F_0_ ATP hydrolysis contributes equally to basal and thermogenic metabolic rates. Although it probably contributes more to the thermogenic than basal metabolic rate, in which case the ascending thermogenic metabolic gradient should be shallower. Regardless, this diagram is here to show that decreased metabolic heat generation, by drug conferred selective F_1_F_0_ ATP hydrolysis inhibition, increases the thermoneutral temperature. Which does *not* cause body temperature drop unless the drug dose and ambient temperature are such that the updated thermoneutral temperature exceeds the ambient temperature. In which case, because of the shallower thermogenic gradient, mouse body temperature falls towards (but never less than) the ambient temperature. So, if for example the ambient temperature is 36°C, if the administered drug dose increases the thermoneutral temperature (from 32°C) to 35°C then there is ***<u>no</u>*** body temperature drop and the mouse is made more thermos-comfortable at this ambient temperature. But if the ambient temperature is still 36°C, and a greater drug dose increases the thermoneutral temperature to 36.5°C then there *is* a body temperature drop. Its magnitude being a function of thermoneutral minus ambient temperature. No matter the drug dose, thermoneutral temperature can never be made equal or greater than 37°C (because equal and greater would mean a specific basal metabolic rate of zero and negative respectively). Remember that body temperature drop can only occur when the thermoneutral exceeds the ambient temperature, wherein the thermoneutral temperature can’t be equal or greater than 37°C, and so if the ambient temperature is equal or greater than 37°C, no body temperature drop can occur, no matter how great the administered drug dose.

Decreasing metabolic heat generation increases the ambient temperature that the subject is most thermally comfortable at. It doesn’t necessarily cause body temperature to drop (Figure 20). With the drug in the body: if the ambient temperature is at or greater than (the drug-increased) preferred ambient temperature, then there is *no* body temperature drop, and the subject is more thermally comfortable. If the ambient temperature is lower than (the drug-increased) preferred ambient temperature: body temperature falls towards, but never less than, ambient temperature. The magnitude of the drop is a function of (the drug-increased) preferred ambient temperature minus the ambient temperature. Drop can be counteracted by wearing more and/or better clothing.

Reduced metabolic heat production is desirable in hotter parts of the world. Because it increases the ambient temperature that the body is most comfortable at. Its “thermoneutral temperature”. Above and below which, greater metabolic rate is required to maintain the body at 37°C (Figure 20).

An awake, typically clothed, human is most comfortable at an ambient temperature around 20.3°C [180]. This is their thermoneutral temperature. But much of the world is hotter than that, for at least part of the year. A selective F_1_F_0_ ATP hydrolysis inhibitor drug (*that doesn’t inhibit F_1_F_0_ ATP synthesis*), by decreasing metabolic heat generation, increases a subject’s thermoneutral temperature. The dose dictating by how much, and pharmacokinetics dictating for how long. For example, a drug dose can increase the thermoneutral temperature (of someone in typical clothes) to 23°C, a higher dose to 27°C, an even higher dose to 32°C, etc. Thereby it can increase thermal comfort in hot climates and seasons: i.e., upon much of the globe. 43% of the world’s population, 3.8 billion people, live in the tropics [170].

A subject may relocate to a warmer climate (e.g., to Florida) to take a higher drug dose, for example if they have cancer or another age-related disease, perhaps upon retirement. Ambient temperature to the subject, and/or their body insulation (e.g., clothing), only needs to be conducive to the drug dose when they have an effective amount of drug in their system. Optionally taking the drug before sleep (in a warm room) and, pharmacokinetics depending, it fractionally or totally clearing from the body by morning.

If ambient temperature is at typical room temperature, ∼20°C [171], then a *low* drug dose is permissible, moderately slowing aging by a relatively small amount of aforementioned (*a*), and by (*b*) atop.

*High* drug dose at a room temperature of ∼20°C, decreasing body temperature to (little above) this, will put the subject into hibernation, which can be maintained for an extended period by intravenous drug infusion, which has applications. For example, spaceflight [172]. Especially as this hibernation can be ended quickly by raising ambient temperature. There will be much slower aging during this hibernation. Hibernating species tend to have longer maximal lifespans [173–177]. Incidentally, there may be surgical applications as body temperature is made 20-25°C for some surgeries, presently very crudely.

### Some predicted use cases

[1] ***Systemically*** administer **high dose** of a selective F_1_F_0_ ATP hydrolysis inhibitor compound (*that doesn’t inhibit F_1_F_0_ ATP synthesis*), to greatly slow aging in the entirety of the body. Wherein the subject’s ambient temperature and/or body insulation offsets their decrease in metabolic heat generation (which scales with drug dose), so their body temperature remains the same. Possibly used for anticancer therapy in a heated hospital ward, given such compound’s anticancer activity: Figures 18-19.

Possibly used for increasing thermal comfort in hot climates/seasons. By (dose-dependently) reducing metabolic heat generation, increasing the ambient temperature that the subject is most thermally comfortable at. Concurrently slowing their aging.

[2] ***Systemically*** administer **low dose** of a selective F_1_F_0_ ATP hydrolysis inhibitor compound (*that doesn’t inhibit F_1_F_0_ ATP synthesis*). To slow aging in the entirety of the body, wherein the subject’s ambient temperature and/or body insulation does not offset their decrease in metabolic heat generation (which scales with drug dose). So, there is a slight, imperceptible, safe body temperature drop.

Lower body temperature corresponds with longer lifespan in mice [21] and humans [31, 168, 181–182], and longer healthspan in humans [183–184].

Lower body temperature occurs during calorie restriction [35–39], including in humans [37–39]. Calorie restriction, possibly by reducing body temperature [185], extends lifespan in model organisms [31–32]. A calorie restriction mimetic is a longstanding hope of the longevity field [186]. At least in mice, lower resting body temperature occurs after exercise [187].

Mice made to have *mean* body temperature 0.34°C lower have 20% longer median lifespan [21]. This lower *mean* body temperature is (presumably) imperceptible, as the daily imperceptible fluctuations in body temperature are greater than that in mice, and in humans [188].

Lower body temperature may be therapeutic for Alzheimer’s disease (refer Supplementary Material).

[3] ***Locally*** administer **high dose** of a F_1_F_0_ ATP hydrolysis inhibitor compound (*that doesn’t inhibit F_1_F_0_ ATP synthesis*) to slow aging in that locality of the body. No thermoregulatory concern because of heat transfer, especially via blood flow, from other body regions, which maintains this body part around 37°C. Local inhibition of F_1_F_0_ ATP hydrolysis (herein predicted sufficient to slow aging by two-thirds) is proven safe in mice [77–79].

Example: *local* administration to skin of the human face: e.g., administered around the eyes in a cosmetic cream.

The body’s natural F_1_F_0_ ATP hydrolysis inhibiting compound (*which doesn’t inhibit F_1_F_0_ ATP synthesis*) is IF1 protein [8–14]. Mammal species with more IF1 protein activity (per unit mass) have a greater maximal lifespan. Increasing IF1 protein amount safely reduces a biomarker of aging in mice. Reducing a biomarker of aging indicates reducing/reversing of aging [44].

My international (PCT) patent application [20] discloses and protects cosmetics containing a functional IF1 protein derivative. For example, a shorter but still functional human IF1 protein fragment: e.g., its residues 42-58 [189–192]. Concatenated, to its N-terminal end, is a Mitochondrial Import Sequence (MIS), as used by the complete IF1 protein. And concatenated, to its N-terminal end, is a Cell Penetrating Peptide (CPP) sequence. Once inside the cell, endogenous protein machinery in the mitochondrial matrix cleaves off the MIS. And inherently the, more N-terminal, CPP sequence with it.

A string of seven arginine residues is experimentally demonstrated to be an effective CPP sequence [193]. No one knows why. I think because, as the inside of the cell is negative [194], its concentration of seven positive arginine charges in a relatively small space outside the cell are collectively so attracted that, given that the membrane has “fluid” character [194], they punch through the membrane, pulling their cargo into the cell. Similarly, the CPP sequence that HIV (as a virus, it is much larger than a protein) uses to get into cells has concentrated positive charges [20].

This seven-arginine (CPP) sequence is a naturally occurring amino acid sequence within the human body, in multiple proteins findable by a BLAST [195] search. So, this cosmetic agent is a concatenation of amino acid sequences that naturally occur within the human body.

Being proteinaceous helps keep drug action local to the skin applied to, as such drugs don’t last long in the bloodstream, being cleaved by proteases in the blood. Although [20] also teaches sequence variants, such as retroinverse thereof, which should have enhanced half-life in blood, if this is desired.

Cosmetics typically contain antioxidants (e.g., vitamin C, vitamin E) because decreasing Reactive Oxygen Species (ROS) amount is a cosmetic aim, and marketable cosmetic claim. But antioxidants don’t decrease [ROS] by much. By contrast, increased IF1 protein, and less F_1_F_0_ ATP hydrolysis thereby, greatly decreases [ROS] (Figure 5). Hair greying/whitening is reportedly caused by ROS [88].

Premium brand cosmetics (>$100 per unit sale) typically contain patented, proprietary amino acid sequences.

Incidentally, Unity Biotechnology is a public company (NASDAQ: UBX), which has raised hundreds of millions of US dollars, whose candidate anti-aging compounds (“senolytics”) are systemically toxic, and can only be administered locally (e.g., to the eye or knee) [196]. So, drugs, and drug candidates, that can only be administered locally can be valuable. Especially for diseases of aging of the eye (e.g., Age-related Macular Degeneration, AMD), for which there is great unmet medical need.

### Anticipated objection: “But what about bats?”

At least some bat species have a much greater maximal lifespan than expected for their adult body mass [197]. For example, *Myotis brandtii* has a maximal lifespan of 41 years, despite only having a typical adult body mass of 7 g [1]. Using the applicable best-fit line equation of Figure 3 herein, an inputted body mass of 7 g predicts a maximal lifespan of only 3.6 years.

This discrepancy might be explained by the following. Bats, probably because they can fly and so can sleep in very safe places (e.g., hanging from the roof of caves), lower their metabolism to an atypically large extent during sleep. So much that their body temperature can fall to just above ambient temperature. Indeed, when the ambient temperature is low, at least some bat species cannot fly when they first wake up, having to metabolically generate heat to raise their body temperature first, which takes time (0.63°C per minute in greater, and slower than this in lesser, horseshoe bats) [198].

Quotes from [199]: “Bats tend to hibernate whenever they fall asleep, even in the middle of a summer night”, “body temperature of bats drops considerably during this period of torporific sleep”, “poikilothermic sleep”, “in bats the daytime drop in metabolism was as much as 90 per cent” (bats sleep during the day, and for much of the night). When awake, which is just for 4 hours per 24 hours in some bat species [200], they are typically flying and inherently generating heat thereby.

So, bats are anomalous amongst mammals in that metabolic heat generation isn’t as significant a component of their metabolic rate. A lower body temperature for most of the time, just above ambient temperature, doesn’t require as great a specific metabolic rate to maintain, even with a small body mass.

This feature then confers geography (latitude and altitude), setting the ambient temperature, as a determinant of what the bat’s metabolic rate is for most of its lifespan, thence (I would argue) is a determinant of its lifespan. Some bat species hibernate (living longer thereby) [201] - e.g., for up to 8 months of a year [202] - tending to hibernate for longer in places with longer (typically inherently colder) winters [203]. Greater latitude is a predictor of higher maximal lifespan in both non-hibernating and, to a greater extent, in hibernating bat species (table 2 of [173]). The *Myotis brandtii* individual that died at 41 years old lived in Siberia [203]. Some tropical bat species, or only some individuals within such species, are poikilothermic [204], but are often mischaracterised in the literature as universally homeothermic.

Moreover, as compared to *Blarina brevicauda* (short-tailed shrew) and *Peromyscus leucopus* (white-footed mouse), the respiratory chain of *Myotis lucifugus* (little brown bat) fumbles a lower percentage of the electrons it passages, producing fewer Reactive Oxygen Species (ROS) per unit of O_2_ consumed [205]. Incidentally, less electron fumbling has also been reported in a long-lived bird species [206].

### Anticipated objection: “But what about naked mole rats?”

Naked mole rats have a much greater maximal lifespan (31 years) than expected for their typical adult body mass (35 g) [1]. Naked mole rats are anomalous amongst mammals in that metabolic heat generation isn’t as significant a component of their metabolic rate. Firstly, because their body temperature is atypically low for a mammal, at around 32°C [1]. Secondly, because they generate very little metabolic heat [207–209] and rely on the ambient temperature to maintain their body temperature. Which they can do because their ambient temperature (∼30°C) is *stably* close to their body temperature (∼32°C). Because they live underground in sealed burrows, hardly ever leaving these, eating underground tubers, in a hot equatorial East African climate. Underground is buffered from short colder periods above ground. For example, during storms. So, to interpret, naked mole rats produce less metabolic heat, with a lower specific basal metabolic rate thereby, and live longer accordingly. And even longer still because of their low body temperature (lower body temperature can confer longer lifespan: refer Supplementary Material).

Their specific basal metabolic rate (*for mean body mass of 42.8 g, ambient temperature was in their thermoneutral zone {31-34°C}, O_2_ consumption of 1 ml/g/h* [207] *= 0.00558333333 W/g using 1 ml O_2_ = 20.1 Joules* [178, 210]) is atypically low for a mammal of their size. Much lower than the value predicted (*0.0083558914 W/g*) for an animal of 42.8 g body weight from the best-fit line for specific basal metabolic rate vs. body mass of Figure 3 herein.

*Data of* [209] *reports a specific metabolic rate value of 0.00307083333 W/g. But this isn’t a basal value, because it was sourced at an ambient temperature of 30°C, and the lower temperature of the naked mole rat’s thermoneutral zone is 31°C. The corresponding specific basal metabolic rate value, sourced at thermoneutrality, would be even lower*.

An interesting parallel is that systemic administration of a selective F_1_F_0_ ATP hydrolysis inhibiting drug (*that doesn’t inhibit F_1_F_0_ ATP synthesis*), at high dose, confers the naked mole rat profile. Low endogenous heat generation, conferring reliance on exogenous heat to maintain body temperature: i.e., outsourcing body temperature maintenance to ambient temperature. With lower specific basal metabolic rate, and (to predict) slower aging thereby.

Although, unlike naked mole rats, humans don’t live in stably hot burrows in equatorial East Africa, buffered from cold periods above. But humans in the tropics can pause their use of the drug (or reduce its dose) when a storm is predicted. And/or shelter in a building. And/or increase their body insulation with clothing. Don’t underestimate clothing: e.g., it permits people to live in Siberia, which is extremely cold in winter.

### Anticipated objection: “But what about marsupials?”

Some (especially smaller bodied) marsupial species seem to opt for life on a budget, with a lower body temperature [211] (e.g., 35°C in the North American opossum, *Didelphis virginiana* [1]), reducing their food demand and risk of starvation, but presumably increasing their risk of predation. As their nerve conduction (and so action/reaction) must then be slower than predators with a body temperature of 37°C (action potential characteristics, such as conduction velocity, are temperature dependent [212–217]). Indeed, the North American opossum is renowned for feigning death, “playing possum”, rather than running away.

So, perhaps because predation constrains their lifespan to short anyhow, and doubling down on their “life on a budget” ecological niche, these marsupial species tend not to allocate as much (NADPH) energy to antioxidant defences. Possibly, I suggest, explaining the longstanding mystery [197] of why these marsupial species tend to have short (for their adult body size) maximal lifespans.

Indeed, such a marsupial species, the Brazilian gracile opossum (*Gracilinanus microtarsus*), of similar size to a mouse, has a “poor NADPH dependent antioxidant capacity”, “both content and state of reduction (mainly of NADPH) were lower in the marsupial mitochondria than in mice mitochondria”, “a more oxidized state of the mitochondrial NADP was demonstrated by HPLC analysis which strongly supports the idea that the mitochondrial NADPH dependent antioxidant systems glutathione and thioredoxin peroxidases/reductases are less effective in the marsupial due to a lower reducing power provided by NADPH”, isolated liver mitochondria from *Gracilinanus microtarsus*, compared to isolated liver mitochondria from mice, “have a much lower constitutive antioxidant capacity represented by the NADPH/NADP^+^ content and redox potential” [218].

*Didelphis virginiana*’s maximal lifespan predicted from the best-fit line of maximal lifespan vs. body mass, in Figure 3, is 12.1 years. Whereas its observed maximal lifespan is lower at 6.6 years [1]. The maximal lifespan observed in an island population of *Didelphis virginiana* without any predators is 45% higher than that observed in a mainland population with predators [219]. By contrast, with some other (non-marsupial) species, a population with predators has the same [220] or a higher [221–222] lifespan than a population without predators (see also [223]).

### Anticipated objection: “But after correcting for body mass and phylogeny, maximal lifespan does not correlate with basal metabolic rate across mammals and birds”

Citing analysis of [6], preeminent leader in the longevity field Professor Steve Horvath (and coauthors) recently wrote in [130]: “the application of modern statistical methods which account for the effects of both body size and phylogeny. When these factors are appropriately adjusted for, there is no correlation between metabolic rate and lifespan in mammals or birds”.

I refute this with novel, extensive analysis in the Supplementary Material. [6] overlooked collinearity between adult body mass and basal metabolic rate across these species, a critical error, rectified in my analysis, which changes the conclusion.

To expand on this very briefly here. Collinearity breaches an assumption of multiple linear regression, that assumption being that there is no collinearity. Collinearity between adult body mass and basal metabolic rate (VIF > 16) increases standard errors of unstandardized regression coefficients in multiple linear regression, conferring the unstandardized regression coefficient for basal metabolic rate erroneous sign and magnitude, which confers partial and semi-partial (part) correlation coefficients erroneous sign and magnitude, which is also the case when using phylogenetically independent contrasts thereof.

Using specific basal metabolic rate instead of basal metabolic rate, which is (I argue) the more salient measure, in a Generalized Additive Model (GAM) that, distinct from multiple linear regression, doesn’t require inter-linearity and so can be used with unlogged variables which are less collinear, even with adult body mass as a predictor, specific basal metabolic rate is a highly significant predictor of maximal lifespan across 517 mammal and bird species. Moreover, phylogenetically Independent Contrasts (PIC) reports that, across 479 mammal and bird species, maximal lifespan significantly correlates with specific basal metabolic rate even when any effect of phylogeny is statistically excluded.

### Anticipated objection: “But predation explains why larger mammal species live longer”

I address this fully in the Supplementary Material. But to make a brief point here. If predation constrains and thereby sets maximal lifespan, why can’t apex predators, which have *no* predators, not live for *much* longer than they do? Indeed, the lion has a shorter maximal lifespan than its prey the zebra. Predation theory (of maximal lifespan) can’t explain this. But the present teaching can. Because the lion is smaller. Refer to Figure 14.

### Anticipated objection: “But what about ectotherms?”

Ectotherm data, analysis thereof, and discussion is in the Supplementary Material. Larger, longer-living ectotherm species have lower specific basal metabolic rate. Interpreted to be because they have lower proton leak rate per unit mass because of lower mitochondrial membrane surface-area per unit mass because moving a larger body requires less energy per unit mass. Refer Supplementary Material.

### Anticipated objection: “But what about exercise?”

Discussed in the Supplementary Material, but to say briefly here, it might not be exercise itself that confers benefit, but rather the body’s response to it, as exercise with insufficient rest (*to make and benefit from such a response*) is harmful. Data suggests that a response to exercise is lower specific basal metabolic rate.

### Anticipated objection: “But why do antioxidants only extend life modestly or not at all?”

To be expected. As explained earlier in the Results section headed “Why don’t antioxidants work much?”. Please read there before reading below.

To add a further point here: even an endogenous antioxidant enzyme can be pro-oxidizing. SOD mitigates superoxide (O_2_^•-^). But it can also perform a side-reaction, generating from H_2_O_2_ the hydroxyl radical (^•^OH), which is arguably the most damaging ROS: e.g., it can cause DNA double-strand breaks [123–124, 224–227]. This may become increasingly prevalent with age, because intracellular H_2_O_2_ increases with age [40–42, 88]. So, SOD activity is essential (mice without SOD2 die within 10 days of birth [228]) but more SOD isn’t necessarily beneficial [229]. This is especially the case with SOD2, which has manganese (Mn) in its active site. As ROS damage releases iron (Fe) from ferritin [230] and from iron-sulfur (e.g., Fe-S, Fe_2_-S_2_ etc.) centres of proteins [231–234], increasing intracellular [Fe], Fe increasingly outcompetes Mn to be in the active site of SOD2. Forming FeSOD2 instead of MnSOD2 [235], which readily produces ^•^OH from H_2_O_2_ by the Fenton reaction. Too much SOD2 or SOD1 can be lethal [236–237].

The endogenous antioxidant system converts superoxide to water and oxygen [4, 238–239]. Why does approximately halving the amount of SOD2 enzyme not significantly decrease mice lifespan (*3 and 4% decreased mean and maximal lifespan respectively, although not significant with the statistical power of the study*) [240]. And why does approximately doubling the amount of SOD2 not significantly increase mice lifespan (*2 and 3% increased mean and maximal lifespan respectively, although not significant with the statistical power of the study*) [241]? *{N.B.,* [242] *reports that increased SOD2 increases mean and maximal lifespan of mice by 4 and 10% respectively, and* [243] *reports that increased SOD2 increases maximal lifespan in flies by up to 37%}*.

I interpret that approximately halving or doubling the normal amount of SOD2 has little to no impact on lifespan, as reported in [240–241], because of the antioxidant system’s network character, causing non-linearity between SOD2 amount and lifespan, set out as follows. Because the antioxidant system is a network of interlocking/interdependent reactions, this dampens the benefit (to lifespan) of overexpressing any specific antioxidant enzyme. As a pathway can only go as fast as its rate-limiting (rate-determining) step [244], wherein if the rate-limiting step is sped it might not speed the pathway much if its rate-limiting character soon switches to another step. Overexpressing an antioxidant enzyme that isn’t the rate-limiting step has little to no effect on lifespan. Lessening an antioxidant enzyme that isn’t the rate-limiting step has little to no effect (on lifespan) until the lessening is sufficient to make it the rate-limiting step. Moreover, it isn’t only the amount of antioxidant enzymes that sets the rate of antioxidation, but also the rate of NADPH energy that this antioxidant system is afforded, wherein if this is limiting (and/or if some other component is limiting, such as glutathione and/or manganese) this is likely to further dampen the effect of overexpressing at least some types of antioxidant enzyme(s).

Alternatively, the antioxidant system may not have a single rate-limiting step but multiple flux-controlling steps, possibly that control to different degrees (that may vary depending on the conditions), which dampens the effect of modulating the amount of any single enzyme. Indeed, to illustrate, glycolysis is such a system [245–247], wherein 50-80% less expression of some glycolytic enzymes doesn’t appreciably slow glycolysis, as read out by glycolysis-dependent wing-beating frequency of flies [248].

The antioxidant system has parallelization (e.g., catalase, glutathione peroxidases, and peroxiredoxins, all reduce H_2_O_2_ to H_2_O [238]; e.g. thioredoxins [249], etc.). With this redundancy, presumably knock-out (or less) of one antioxidant enzyme can increase the substrate and so the rate (by Michaelis–Menten enzyme kinetics [145]) of a parallel antioxidant enzyme(s). Parallelization dampens the negative effect (on lifespan) of less/loss of a specific antioxidant enzyme(s), and it dampens the positive effect (on lifespan) from overexpression of any specific antioxidant enzyme(s), as parallelization reduces the extent to which any single antioxidant enzyme is rate-limiting. Note that we might not yet know all enzyme elements of the antioxidant system. Further dampening/buffering of effect (on lifespan) is afforded by ROS damage repair enzymes.

SOD2 is in the mitochondrial matrix [250, 239], so close to the respiratory chain, where [superoxide] is likely highest. Although the antioxidant system has SOD1 and SOD3 also [250, 239], outside the mitochondrial matrix, these may not substitute well for SOD2 loss because superoxide can’t easily cross the inner mitochondrial membrane [239]. So, this might be a point in the antioxidant system which isn’t well parallelized, which is perhaps why mice without SOD2 die within 18 days of birth [251] (within 10 days in [228]). This reflects how damaging ROS are, and how vital the antioxidant system is. So, endogenous antioxidant SOD2 extends mouse lifespan from ≤18 days (when it is absent) to years, which is a large effect size.

Mice without SOD1 have a ∼30% shorter lifespan [252], mice without methionine R sulfoxide reductase have a ∼40% shorter lifespan [252], and lack of any of phospholipid glutathione hydroperoxidase, thioredoxin 1, thioredoxin 2, thioredoxin reductase 1, and thioredoxin reductase 2 is embryonically lethal in mice [253–258].

Varied exogenous and endogenous antioxidants, at least at some dosage(s), extend lifespan: e.g., [84–86, 242–243, 259–279]. For which, a parsimonious interpretation is that ROS are a drive of aging. Indeed, to my knowledge, no one has been able to explain these lifespan extensions, shown by different groups (including the Interventions Testing Program, ITP [279]) with different and varied antioxidant interventions, otherwise. But their effect size is typically modest. To interpret, not least because there are ***<u>so</u>*** many more biological than added antioxidant molecules in a body, wherein the former tend to be larger (e.g., DNA is *much* larger), and so each ROS is *extremely* more likely to collide with, and damage, a biological molecule than collide with, and be mitigated, by an added antioxidant molecule. Where there is a limit to how many exogenous antioxidant molecules can be safely administered, because too much of *any* exogenous (or even endogenous) molecule is toxic, especially in this instance because at least some antioxidants can be pro-oxidizing at higher concentrations: e.g., they generate ^•^OH from H_2_O_2_ by the Fenton reaction [87].

Proverbially, I suggest that antioxidant administration is like closing the barn door after the horse has bolted. For us to meaningfully drug reduce ROS concentration, we should reduce ROS generation. This paper, and its sister patent applications [18–20], teaches drugs to do this. A novel approach.

## Supplementary Material

Much further discussion is in the Supplementary Material.

## CONCLUSION

How can an old dog be younger chronologically, but older biologically, than its human owner? Why do dogs age faster than humans? Why do different mammal species age at different rates, and have different maximal lifespans? This work might have answered these questions at molecular resolution. Mammal species with more IF1 protein activity (per unit mass) have a greater maximal lifespan. Moreover, increasing IF1 protein amount safely reduces a biomarker of aging in mice. IF1 protein constrains specific basal metabolic rate. Concordantly, there is, across and within species, correlational *and* interventional data, including (correlational) human data, pointing to maximal lifespan being inversely proportional to specific basal metabolic rate. For example, a genetic intervention in mice that *slightly* reduces metabolic rate *greatly* extends lifespan. Herein taught are drugs that, like IF1 protein, dose-dependently inhibit F_1_F_0_ ATP hydrolysis (*and not F_1_F_0_ ATP synthesis*) and thereby reduce specific basal metabolic rate, as demonstrated for one such drug in mice, which I therefore predict can at least slow aging. Both small-molecule and biologic drugs. The latter primarily being IF1 protein derivatives, particularly amenable to cosmetic use.

Herein disclosed is how mammals principally generate heat, F_1_F_0_ ATP hydrolysis, a fundamental discovery, filling a large gap in our understanding, which wasn’t even appreciated to be a gap previously. And this work contributes a new cancer drug target, F_1_F_0_ ATP hydrolysis, and a new class of anticancer drugs for further testing, which distinctly and valuably, may help instead of harm normal cells. Anticancer drugs that may slow aging.

## METHODS

Some definitions: “specific” (by convention) can refer to “per unit mass”, which is a convention used often herein. “Maximal lifespan” of a species is the longest lifespan ever recorded for a member of that species.

### 6b drug administered to mice (*generating Figure 2*)

Four female C57BL/6 strain mice (*Mus Musculus*) were sourced from Shanghai Lingchang Bio-Technology Co. Ltd. They were 12-14 weeks old on the day of the rectal temperature recording experiment. Their care and use were in strict accordance with the regulations of the Association for Assessment and Accreditation of Laboratory Animal Care (AAALAC). Some details of their care: *ad libitum* Co^60^ irradiation sterilized dry granule food, reverse osmosis autoclaved drinking water, corn cob bedding, 20-26°C room temperature, 12 hours light/dark cycle, polysulfone individually ventilated cage (IVC, containing up to 5 animals): 325 mm*210 mm*180 mm, 40-70% humidity.

StudyDirector^TM^ software (Studylog Systems, Inc., CA, USA) was used to allocate/randomize control/treatment groups. Mice were marked by ear coding (notch).

When recording rectal temperature, care was taken to ensure constant depth of probe insertion across different recordings. Time was afforded for rectal probe temperature to equilibrate with rectal temperature. And time was afforded between recordings for probe temperature to reset. The rectal temperature recording experiment was started between 8 and 9 am.

Intravenous (IV) administration (tail vein): dosing volume per weight of drug = 10 µl/g, solution (not suspension). Sterilised (0.22 µm filter), vortexed vehicle = 12.5% solutol, 12.5% ethanol, 75% water (admittedly, it was unwise to include a drug, ethanol, in the vehicle. All be it is commonly used for this purpose). IV solutions freshly prepared before injection.

The two mice intravenously injected with 20 and 40 mg/kg drug doses both exhibited hypoactivity and tachypnea. Which are both signs of hypothermia [280], coinciding with their rectal temperature drop to below 30°C. 210 minutes (3.5 hours) after the intravenous injection of 20 mg/kg, that mouse died (“choked when drinking water” was the reported observation by the experimenter. Reason/how is unknown). The mouse injected with 40 mg/kg was found dead the next day. The mouse injected with 2 mg/kg survived (and was sacrificed 54 days later). 24 hours post the IV injection, its rectal temperature was 39.99°C, compared to 39.48°C in the control (vehicle administered) mouse.

So, the (intravenous) Maximal Tolerated Dose (MTD) in mice is certainly more than 2 mg/kg, and possibly less than 20 mg/kg. Demonstrated, it is an active, potent drug *in vivo*. A prediction is that the MTD would be greater if the ambient temperature was safely higher. For example, it would be higher if the ambient temperature was instead 30°C. Even higher if it was 34°C, and higher still if 37°C. *Incidentally, it is widely unknown/underappreciated that, for any mouse experiment, laboratory mice should be kept at a higher than typical room temperature. Typical room temperature (∼20°C* [171]*) is thermoneutral for us humans (in clothes* [180]*) but it isn’t for mice, whose thermoneutral temperature is instead ∼32°C* [178, 281–283].

A possibility is that either or both the ≥20 mg/kg drug administered mice were mischaracterized as dead (if the 40 mg/kg drug administered mouse was mischaracterized, then the drug’s MTD is >40 mg/kg). And instead of dead, one or both were merely hypothermic due to the drug action. With reduced body temperature (at slightly above ambient temperature, which was ∼22°C), as a function of reduced metabolic rate. And unresponsive to external stimuli upon cursory inspection (rodent unresponsiveness during hypothermia was observed by [284]: “it did not arouse even when repeatedly disturbed”). Because of the drug’s action, unable to generate heat to arouse themselves. Unfortunately, the experimenter wasn’t forewarned to be vigilant to this possibility (e.g., not told to check for a heartbeat, all be it likely slowed). Rendering an inconclusive MTD. Because one or both these mice might have survived if given the opportunity. Returning to normal/normothermia as the drug cleared from the body. Which might have taken some time given that reaction rates are temperature dependent, and these mice had lower than normal body temperature. Data of [284] suggests that rodents can recover from hypothermia once the hypothermic drive is removed.

In this case the effect size is large (the drug-conferred changes in mouse body temperature are large: e.g., dropping body temperature to below 30°C). When an effect size is large it can be detected at statistical significance by a small sample size (observable by scenario testing with the Excel spreadsheet model of [285] or statistical power calculators such as [286]). For animal studies, an overpowered sample size (more subjects than required to hit statistical significance) is anathema to the 3Rs (Replacement, Reduction, Refinement) [287].

A (unfortunate, but mice sparing) confounding issue is that in the 43 days preceding the rectal temperature recording experiment (which was conducted on day 44) the mice were used in a different experiment. A dose escalation study of a different chemical entity. Of the same chemical formula, but the opposite stereoisomer thereof (*R*), in enantiomeric excess (≥97%): compound 6a in Figure 18A. Which, unlike compound 6b, does *not* inhibit F_1_F_0_ ATP hydrolysis (6b EC_50_ F_1_F_0_ ATP hydrolysis = 0.018 μM, whereas 6a EC_50_ F_1_F_0_ ATP hydrolysis > 100 μM [63–64]). This experiment sought the Maximal Tolerated Dose (MTD) in mice of compound 6a. Wherein the control mouse was always administered vehicle, and the three test mice were administered an escalating 6a dose amount every 3 days, ultimately reaching 40 mg/kg. The final dose of 6a was on day 39, wherein the highest dose given then was 2 mg/kg (ran out of 6a and so doses were low on that last day). So, the MTD was not successfully found for 6a.

More details of these experiments can be found in my international (PCT) patent application [18].

To give some data for comparison: in mice, the intravenous LD_50_ (dose that kills 50% of the cohort) values for the FDA approved anti-depressants, clomipramine HCl and imipramine HCl, are 22 and 27 mg/kg respectively (Register of Toxic Effects of Chemical Substances, RTECS). Some human patients take these drugs daily and safely for years.

### Generating Figure 3

For each species, specific F_1_F_0_ ATP hydrolysis rate data herein is from Sub-Mitochondrial Particles (SMPs). Produced by sonicating mitochondria, which were isolated from ischemic heart tissue. Ischemia, by collapsing the proton motive force (pmf), causing a lower pH in the mitochondrial matrix, increases the active fraction of IF1 protein. Which comprises IF1 protein monomers and dimers, which *can* bind ATP synthase [15]. As opposed to IF1 protein tetramers, and higher oligomers, which cannot. Ischemia ensures that a large proportion of IF1 protein complexes with the membrane bound ATP synthase. Which means that more IF1 protein is retained/available after the subsequent SMP generation protocol. SMPs are inside-out (inverted). With the mitochondrial matrix side of the membrane, and so the ATP hydrolysing and IF1 binding modules of ATP synthase, on their outer face. So, IF1 protein is wanted on the outside, not on the inside, of the SMPs. Prior ischemia, which combines IF1 protein with the mitochondrial matrix side of ATP synthase, assists this. Moreover, (at least some of) this activated IF1 protein fraction carries over into the start of the subsequent F_1_F_0_ ATP hydrolysis assay (conducted at pH 7.8, for 5 minutes). So best showcasing differences in IF1 protein activity between different species. Rendering them detectable by the resolution of this experiment. Where (to hypothesize) the differences in IF1 protein activity, driving differences in F_1_F_0_ ATP hydrolysis, comprise differences in IF1 protein amount (versus ATP synthase amount), IF1 protein inhibitory potency for F_1_F_0_ ATP hydrolysis, and IF1 protein tendency/self-affinity for tetramerization and higher oligomerization (self-sequesteration) at non-acidic pH.

This specific F_1_F_0_ ATP hydrolysis rate data was sourced from [91] (column titled “Ischemic” in its table 1), along with its heart rate and body mass data therein. Maximum lifespan data was sourced from the AnAge database [1]. Specific basal metabolic rate data herein, for each cited species, is the mean of values for that species sourced from AnAge [1] and the landmark papers by Max Kleiber in 1932 and 1947 [288–289], which set out Kleiber’s Law. But the specific basal metabolic rate of rat value from [289] was omitted from this mean, because its value is not from an adult. Further incorporated into the mean of mouse is the value of 0.0092125 W/g (sourced at ambient [thermoneutral] temperature of 32°C, for C57 mouse strain) from [179]. Further incorporated into the mean of rat is the value of 0.005695 W/g (sourced at ambient [thermoneutral] temperature of 28°C) from [210]. For converting O_2_ consumption into energy I used [1 ml O_2_ = 20.1 Joules], which is a well-known conversion factor [178, 210].

Best-fit lines were calculated by OriginPro 8 software (OriginLab Corporation, Northampton, MA, USA), using its “Fit Linear” functionality. Which uses linear regression, with least squares estimation.

### Marrying data

In Figure 3, body mass data and data in the 1^st^ (top) and 3^rd^ panels is from [91], data in the 2^nd^ and 4^th^ panels is from [1] (*2^nd^ panel incorporates further data from other sources, as specified earlier*). This data combination, and its interpretation here, is novel. There was some (enduring) margin for error in marrying these data sets because [91] uses imprecise terms such as sheep, hamster etc., wherein there are multiple species in [1] that can fall into these categories. But a common-sense alignment was applied in each case, by estimating which species [91] likely had easiest access to, so most likely used, and so most likely refers to. So, utilizing this *estimation*, the 12 species that Figure 3 herein refers to are: cow (domestic cattle, *Bos taurus*), mouse (house mouse, *Mus musculus*), rat (brown rat, *Rattus norvegicus*), hamster (golden hamster, *Mesocricetus auratus*), guinea pig (*Cavia porcellus*), pigeon (common wood-pigeon, *Columba palumbus*), chicken (*Gallus gallus*), rabbit (European rabbit, *Oryctolagus cuniculus*), sheep (domestic sheep, *Ovis aries*), pig (*Sus scrofa domesticus*), dog (*Canis lupus familiaris*) and human (*Homo sapiens*). Although an acknowledged, reasonable possibility is that the pigeon species that [91] used was instead the rock pigeon (*Columba livia*). Which I discuss in the Supplementary Material, in its section titled “Birds”.

Data from [1] was sourced primarily in August 2018. Which is when the kernel (e.g., Figure 3) of this work was done.

### Omitting human maximal lifespan

Apart from where stated otherwise, maximum lifespan of human (122.5 years [1]) isn’t shown in the figures nor utilized in calculations (e.g., not used in Pearson or Spearman’s rank correlation coefficient calculations to maximal lifespan). Because arguably this value isn’t comparable to the other maximal lifespan values used. As (*a*) modern medicine is disproportionally applied to humans, (*b*) many modern humans live in shelter and comfort (with complete freedom and stimulation, unlike an animal in a zoo for example, whose optimal husbandry might not even be known/performed), and (*c*) the verifiable lifespan data set for humans is hugely larger, with so many countries recording births and deaths (up to a presumable limit, the bigger the data set, the greater the chance a higher maximum lifespan will be found). So, compared to the human number, which is drawn from a *huge* sample size (probably of billions), the other numbers, drawn from small sample sizes (incredibly small in some cases), are likely to be an underestimate of species maximal lifespan. Human could perhaps be comparably incorporated by using a maximum lifespan record from a smaller human data set. To mirror the small data sets for the other species. Wherein this data set comes from humans living in the past. For example, from Germany in 1881, where life expectancy of men and women was 35.6 and 38.5 years respectively (Statistisches Bundesamt Deutschland, www.destatis.de), or perhaps even earlier, before the industrial and agricultural revolutions, before modern medicine (e.g., before antibiotics and vaccines). However, omission was typically chosen instead. But not always, because in larger species sets, the anomalous maximal lifespan value for human is diluted by the other data and so the imperative to omit it also.

### Chicken maximum lifespan

The AnAge database [1] states 30 years for the maximal lifespan of chicken (red junglefowl, *Gallus gallus*). But (*at least when referred to in August 2018*) in the “Observations” section of this database entry, AnAge says that this “remains unproven. For comparative analyses the use of a more conservative value for maximum longevity, such as 15 or 20 years, is recommended”. I split the difference, between 15 and 20, and use 17.5 years.

## Supplementary Material

Please refer to the Supplementary Material for much more pertaining to Methods.

## ACKNOWLEDGEMENTS

Under author’s direction, two Contract Research Organisation (CROs) were utilized to generate the data of Figure 2, and Figures S19-S53. WuXi AppTec synthesized the compounds. Crown Bioscience performed the experimentation in mice. National Cancer Institute (NCI, USA) did the anticancer testing, with compounds submitted to them by author’s instruction, generating the data of Figures 18-19. All of whom the author thanks greatly.

