## Supplementary Material for "IF1 Protein Controls Aging Rate"

Comprising:

[1] Supplementary Results

[2] Supplementary Discussion

[3] Supplementary Methods

[4] Supplementary Material References

#### SUPPLEMENTARY RESULTS

Normality of data

For data used to produce Figure 3:

[Table S1]

| **Kolmogorov-Smirnov Test of Normality (D)** | | p-value | Verdict |
|---|---|---|---|
| Specific F1F0 ATP hydrolysis rate | 0.22906 | 0.48489 | Normal |
| Specific basal metabolic rate | 0.17987 | 0.77037 | Normal |
| Heart rate | 0.23315 | 0.46279 | Normal |
| Maximal lifespan | 0.17922 | 0.81215 | Normal |
| Body mass | 0.34847 | 0.08319 | Normal |
| *Normal = data does not differ significantly from that which is normally distributed* | | | |

Raw (unlogged) data used to produce Figure 3

[Table S2]

| | Body mass | Specific F1F0 ATP hydrolysis rate | Specific basal metabolic rate | Heart rate | Maximum lifespan |
|---|---|---|---|---|---|
| Cow | 500 | 1.32 | 0.000798458 | 50 | 20 |
| Pig | 100 | 0.81 | 0.000771 | 73 | 27 |
| Human | 90 | 1.31 | 0.001164574 | 69 | Unused |
| Sheep | 60 | 1.49 | 0.001319925 | 75 | 22.8 |
| Dog | 25 | 1 | 0.001804232 | 85 | 24 |
| Rabbit | 2 | 1.12 | 0.002887816 | 213 | 9 |
| Chicken | 2 | 2.89 | 0.002418352 | 318 | 17.5 |
| Pigeon | 0.6 | 2.66 | 0.004975053 | 305 | 17.7 |
| Guinea Pig | 0.4 | 2.71 | 0.003740752 | 265 | 12 |
| Rat | 0.2 | 3.02 | 0.00601463 | 481 | 3.8 |
| Hamster | 0.1 | 3.9 | 0.006377 | 400 | 3.9 |
| Mouse | 0.025 | 2.73 | 0.010858546 | 618 | 4 |

Specific $F_1F_0$ ATP hydrolysis rate (µmol/min/mg) data is from [91] (column titled "Ischemic" in its table 1). Body mass (kg) and heart rate (bpm) data is also from [91] (table 3 therein). Maximal lifespan (years) data is from [1] (human maximal lifespan unused by rationale given in Methods). Specific basal metabolic rate (W/g) data is the mean of data from [1], [288-289], [179, 210]: a compilation of sources because no single source has data for all the investigated species. The mean specific basal metabolic rate data in Table S2 above is calculated from its source papers as shown in Table S3 below, where [α] is [1], [β] is [288], [γ] is [289], [δ] is [179], and [ε] is [210]. The specific basal metabolic rate of rat value from [289] is omitted, because its value is not from an adult. With data from [179] and [210], for converting $O_2$ consumption into energy I used [1 ml $O_2$ = 20.1 Joules], which is a well-known conversion factor: e.g., as used by [178, 210].

[Table S3]

| | [α] | [β] | [γ] | [δ] | [ε] | Mean |
|---|---|---|---|---|---|---|
| Cow | 0.000884 | 0.000759568 | 0.000751805 | | | 0.000798458 |
| Pig | 0.000771 | | | | | 0.000771 |
| Human | 0.001183 | 0.001195378 | 0.001115345 | | | 0.001164574 |
| Sheep | | 0.001296472 | 0.001343377 | | | 0.001319925 |
| Dog | | 0.001745968 | 0.001862497 | | | 0.001804232 |
| Rabbit | 0.003411 | | 0.002364633 | | | 0.002887816 |
| Chicken | 0.002216 | 0.002620705 | | | | 0.002418352 |
| Pigeon | | 0.004975053 | | | | 0.004975053 |
| Guinea Pig | 0.003333 | | 0.004148504 | | | 0.003740752 |
| Rat | 0.006786 | 0.00556289 | Omitted | | 0.005695 | 0.00601463 |
| Hamster | 0.006377 | | | | | 0.006377 |
| Mouse | 0.015056 | | 0.008307139 | 0.0092125 | | 0.010858546 |

Figures 4 and 6 use the same raw data as Figure 3, except without the body mass data.

My workings to get to the 0.0092125 W/g figure from [δ] [179]: from its table 3, ignoring error bars, for the C57 mouse strain, at an ambient temperature of 32°C (which is the ambient temperature with the lowest $O_2$ consumption on this line of table. And so, out of the ambient temperatures tested at, this one is most likely closest to, or possibly even at, the C57 mouse's thermoneutral temperature. The thermoneutral temperature of mice is known to be at/around 32°C: refer Fig. 5 in [178]), $O_2$ consumption = 1.65 cc/g/hour, where "cc" is cubic centimetre ($cm^3$), (*to get into kg, which is the SI unit of mass*) *1,000 = 1,650 $cm^3$/kg/hour = (*because 1 $cm^3$ = 1 ml, and 1 ml $O_2$ consumption = 20.1 Joules* [178]) 33,165 J/kg/hour = (*because* *power = energy/time = J/s; 60*60 seconds in 1 hour*) 33,165/(60*60) = 9.2125 W/kg = 0.0092125 W/g.

My workings to get to the 0.005695 W/g figure from [ε] [210]: from the first line of its table 1, which refers to BMR (=Basal Metabolic Rate), sourced at an ambient temperature of 28°C (at/around the thermoneutral temperature of a rat [210]), for the Simonsen (Sprague-Dawley derived) rat strain, $O_2$ consumption = 17 ml/kg/minute = (*because 1 ml $O_2$ consumption = 20.1 Joules* [178, 210]) 341.7 J/kg/minute = (*because power = energy/time = J/s; 60 seconds in 1 minute*) 341.7/60 = 5.695 W/kg = 0.005695 W/g. Which corresponds well with the value given for specific metabolic rate on the same line of the same table = 5.7 W/kg.

The conversion algorithm I used for data from [288] and [289]: starting with metabolic rate per day (Kcal) and body weight (kg): converted kcal to calories, converted calories to Joules, divided this by body weight (kg) to get in J/kg/day, (*because power = J/s*) divided by number of seconds in a day (24*60*60) to get in W/kg, then /1000 to get in W/g.

Data was drawn from table 1 of Kleiber's 1932 paper [288]: from its "W Average weight, kilo-grams" and "Cals. per 24 hrs. per animal" columns. In my calculations, an error in this paper was corrected: the table reports kilocalories (kcal), not calories as it purports. This error is evident, for example, when comparing the data (e.g., for woman) of Kleiber's 1932 [288] and 1947 [289] papers (which report independent data sets: "No data were used that were already incorporated in the earlier study (1932)" [289]). The latter's table 2, where the unit of its "Metabol. rate per day" column is correctly labelled kcal. The unit error is identified to be with the 1932, rather than the 1947 paper, because I know the order of magnitude of calories that a person needs per day, even if inactive, is thousands of kilocalories. For example, from the National Health Service (NHS) website: "As a guide, an average man needs around 2,500 kcal (10,500 kJ) a day to maintain a healthy body weight. For an average woman, that figure is around 2,000 kcal (8,400 kJ) a day" [290].

Note, to relay a possibly confounding issue in the data sourced from [288]: quoting [288]: "The relatively low value of the hen may be in connection with the fact that the determinations had been made in darkness".

Note that, in at least some cases, a specific basal metabolic rate value sourced from [1] is a mean of values sourced from multiple sources ("if more than one pair of values was available, the logarithmic average was used. If discrepancies were noticed between two or more pairs of values, then only the most recent pair was used" [291]). But none of these sources include any of [288], [289], [179], [210]. So, these are all independent data sources. At least for now. But these paper values may well get incorporated into the means of the [1] database, called the AnAge (Animal Ageing and Longevity) database, in the future. Be aware that [1] is periodically updated and so values in the present and/or future may be different than those that were sourced from it in the past.

Heart rate

Heart rate values were sourced from [91]. For species with a range of values disclosed therein, the middle value of the range was taken. This heart rate data comes from conscious animals, *except* for hamster: its values are from anesthetized animals. This disparity was ignored. But if instead it isn't, this can increase at least some of the correlations to heart rate, and their statistical significance. The anesthetized heart rate range for hamster in [91] is 375-425. And the value drawn was 400. But if the upper bound value is taken instead, 425, then at least some of the investigated Pearson correlation coefficient (R) values relating to heart rate are increased. And their associated (one-tailed) p-values decreased: i.e., increased statistical significance:

[Table S4]

| | | Specific F1F0 ATP hydrolysis rate | Specific basal metabolic rate | Maximal lifespan | log10(Body mass) |
|---|---|---|---|---|---|
| Heart rate | R | 0.8139 | 0.9476 | -0.8648 | -0.9439 |
| | p | 0.000637 | 0.0000015 | 0.000295 | 0.000002 |

Birds

Conceptually, a different way to decrease ROS generation is, instead of (or in addition to) reducing metabolic rate and $O_2$ consumption rate, modifying the respiratory chain such that it fumbles a lower percentage of the electrons it passages (i.e., increased efficiency) [292]. Some species, possibly at least some bird species, may have availed themselves of such increased efficiency. For example, there is data suggesting that the particularly long-lived (for its size) pigeon species, *Columba livia*, has such increased efficiency [206]. An observed strategy, for increased efficiency, is to reduce the amount of Complex I [293-296].

Moreover, long-lived (for their size) bird species' membranes (e.g., the mitochondrial inner membrane) tend to have less propensity to amplify ROS damage (wherein lipid peroxidation

PUFA products are themselves ROS, which can attack further PUFA molecules, a chain reaction, which can go on to damage other cellular components, e.g., membrane proteins, mtDNA, etc.) because their polyunsaturated fatty acids (PUFAs) tend to have less double bonds [131, 297-302].

To generate the principal correlative Results of this paper I used data from [91], which was (at the time of writing) published 34 years ago. Including its data from pigeon. But [91] doesn't specify which pigeon species it used. I made the estimation that [91] used the common wood pigeon (*Columba palumbus*), which has a maximal lifespan of 17.7 years [1]. Although I concede that [91] may instead have used the rock pigeon (*Columba livia*), which has a very different maximal lifespan. Of 35 years [1]. So, nearly twice as long. Given this uncertainty, I reason that a maximal lifespan of 17.7 years (that for *Columba palumbus*) is the best value to use for pigeon, by the balance of probabilities, as I explain later. And so, this is the maximal lifespan that I allocated to pigeon. Which is of the order of the maximal lifespan (in brackets after each name) of most other pigeon and dove {Columbidae family} species in [1]: *Leucosarcia melanoleuca* (16.3), *Patagioenas leucocephala* (14.4), *Columba oenas* (12.6), *Cepphus Columba* (15.2), *Caloenas nicobarica* (20.5), *Patagioenas fasciata* (18.5), *Streptopelia decaocto* (17.9), *Streptopelia risoria* (12), *Streptopelia turtur* (13.2), *Columbina passerina* (7.2), *Streptopelia chinensis* (9.8), *Geopelia striata* (14), *Leptotila verreauxi* (8.6), *Columbina inca* (13.3). *But* there are three longer-lived, outlier species: *Columba livia* (35), *Zenaida macroura* (31.3), *Zenaida asiatica* (25) [1]. The particularly long-lived *Columba livia* (*breed thereof unspecified*) has been shown to have low Complex I content, in the aforementioned [293]. And a low number of double bonds in its PUFAs, in the aforementioned [298-299].

If [91] used *Columba livia*, which is more of an outlier species, living longer (maximal lifespan of 35 years) for its size, instead of *Columba palumbus*, then the Pearson correlation coefficients (R, and their p-values, one-tailed because each alternative hypothesis is directional) are as shown in Table S5 below. Its second line of R is when pigeon is omitted from the data set, and its third line of R is when the other bird species (chicken) is omitted also, such that all the remaining species are mammals (*N.B., [1] didn't, and presently {July 2021} doesn't, have metabolic data for any pigeon, so the specific basal metabolic rate value of pigeon [particular species unknown] is singly from [288]*):
[Table S5]

| | | Specific F1F0 ATP hydrolysis rate | Specific basal metabolic rate | Heart rate | Body mass |
|---|---|---|---|---|---|
| Maximal lifespan | R | -0.5041 | -0.6008 | -0.66 | 0.224 |
| | p | 0.0569265 | 0.025307 | 0.013556 | 0.253941 |
| Maximal lifespan | R | -0.7535 | -0.8413 | -0.8744 | 0.3818 |
| | p | 0.005924 | 0.0011405 | 0.000467 | 0.13814 |
| Maximal lifespan | R | -0.8189 | -0.8387 | -0.8993 | 0.4078 |
| | p | 0.0034685 | 0.002361 | 0.000483 | 0.1379625 |

A complication with *Columba livia* is that there are hundreds of different breeds thereof. Different "fancy pigeons". Which can be very morphologically diverse. And which, at least in some cases, have had very different maximal lifespans reported, which *may* reflect true differences in maximal lifespan. Perhaps reflecting, deliberate and/or inadvertent, artificial selection for longer lifespan in at least some breeds. For example, discussing different breeds of *Columba livia*, a Cropper pigeon reportedly lived to 35 years old, but (from the *same* source paper) the "limit of the age of the Pouter Pigeon is only 10 to 12 years, the "great proportion dying in less than half that time"" [303] (*incidentally, [303] is the {once removed} upstream source of the 35 year maximal lifespan reported for Columba livia in [1], for which [303] cites a further upstream source, dating from 1911*). Whilst as regards the original/ancestor *Columba livia*, without any breeding/selecting by man: in captivity, in Egypt, at the Giza zoo, "two Rock-Pigeons [*Columba livia*] from the Dongola Province of the Sudan lived certainly for 8 years and possibly for 12 years 10 months" [303]. In captivity, at the same location, a hybrid cross pigeon, of *Columba livia* x *Columba lumbus*, lived for 5 years, 7 months, 17 days [303].

So, it *might* be that *some* breeds of *Columba livia* have a maximal lifespan that is less than, or comparable to, that of *Columba palumbus*. Given that we don't truly know which species of pigeon [91] used, nor which breed thereof (*if* it used *Columba livia*), nor definitively how different breeds of *Columba livia* map to maximal lifespan, moreover wherein at least some bird species tend to live much/anomalously longer than similarly sized mammal species, where mammals are my primary focus, there is a case for omitting pigeon from the data/investigation. But in the Results section of this present paper, wherever a pigeon maximal lifespan is utilized/presented, a relatively conservative maximal lifespan value (17.7 years), that of a very common pigeon species (*Columba lumbus*) [1], of the order of the maximal lifespan of most other pigeon species [1], and *possibly* of *some* breeds of *Columba livia*, is used. Instead of omission.

If omission is selected instead, as shown in Table S6 below, the Pearson correlation coefficients (R) between maximal lifespan and the presented variables are *greater*, and some of the (one-tailed) p-values are accordingly lower (despite the smaller value of *n*). "WITH pigeon" data is as presented in the Results section of the main text. "WithOUT pigeon" data is as presented in earlier Table S5.

[Table S6]

| | | Specific F1F0 ATP hydrolysis rate | Specific basal metabolic rate | Heart rate | Body mass |
|---|---|---|---|---|---|
| **WITH** pigeon | | | | | |
| Maximal lifespan | R | -0.7192 | -0.8144 | -0.8572 | 0.3595 |
| | p | 0.0063065 | 0.001136 | 0.0003725 | 0.1387675 |
| **WithOUT** pigeon | | | | | |
| Maximal lifespan | R | -0.7535 | -0.8413 | -0.8744 | 0.3818 |
| | p | 0.005924 | 0.0011405 | 0.0004665 | 0.13814 |

Replication of the specific $F_1F_0$ ATP hydrolysis rate data used in this study by other studies

In Table S7 below, Paper [α] is [91] (whose data is used in the Results section of this present paper), Paper [β] is [304], Paper [γ] is [305], Paper [δ] is [306], Paper [ε] is [307], and Paper [ψ] is [308]. All these papers are from the same laboratory. But with varied authorship. Probably (at least partly) because their publication dates span 9 years. Notably, the co-author of Paper [γ] is Maynard E. Pullman, who co-discovered the IF1 protein in 1963 [8].

There are some differences in protocol between at least some of the papers. For (non-exhaustive) example, to prepare Sub-Mitochondrial Particles (SMPs), Paper [α] uses sonication only. But Papers [γ] and [ψ] use sonication *and* at least one subsequent centrifugation step. Centrifugation is for mitochondrial debris clearing, e.g., of large mitochondrial remnants and membrane fragments. But it must be used judiciously, lest it remove some of the IF1 protein. Confusingly, at least some of these papers use an idiosyncratic terminology. Where they refer to mitochondrial preparations that have undergone both sonication and centrifugation as Sub-Mitochondrial Particles (SMPs). Whilst referring to those that have only undergone sonication as "sonicated mitochondria". This terminology is somewhat misleading because *both* these preparations contain SMPs (it is the sonication step that breaks up the mitochondria into SMPs). A constant for the data in Table S7 below is that the mitochondria, broken up to form SMPs, were sourced from ischemic heart tissue. If body mass is reported as a range in a source paper, I report its middle value in Table S7 below. An uncontrolled aspect is body mass differences, in some cases large, between the different (paper's) studies.

[Table S7]

| | **Paper [α]** | | **Paper [β]** | | **Paper [γ]** | | **Paper [δ]** | | **Paper [ε]** | | **Paper [ψ]** | |
|---|---|---|---|---|---|---|---|---|---|---|---|---|
| | [A] | [B] | [A] | [B] | [A] | [B] | [A] | [B] | [A] | [B] | [A] | [B] |
| Cow | 500 | 1.32 | | | | | | | | | | |
| Pig | 100 | 0.81 | 17.01 | 1.33 (±0.15) | | | | | | | | |
| Human | 90 | 1.31 | | | | | | | | | | |
| Sheep | 60 | 1.49 | | | | | | | | | | |
| Dog | 25 | 1 | | | NS | 0.87 (±0.11) | 30 | 1.12 (±0.12) | 10 | 1.75 (±0.08) | | |
| Rabbit | 2 | 1.12 | 1 | 1.32 (±0.18) | | | | | | | 1 | 1.18 (±0.08) |
| Chicken | 2 | 2.89 | | | | | | | | | | |
| Pigeon | 0.6 | 2.66 | 0.4 | 2.99 (±0.12) | | | | | | | 1 | 3.35 (±0.16) |
| Guinea Pig | 0.4 | 2.71 | 0.4 | 3.04 (±0.06) | | | | | | | | |
| Rat | 0.2 | 3.02 | 0.3 | 3.27 (±0.11) | | | 0.25 | 2.91 (±0.24) | 0.5 | 2.66 (±0.17) | | |
| Hamster | 0.1 | 3.9 | | | | | | | | | | |
| Mouse | 0.03 | 2.73 | 0.025 | 3.31 (±0.07) | | | | | | | | |
| | [A] = **Body mass (*kg*), **[B] = **specific F1F0 ATP hydrolysis rate (*μmol/min/mg*)**; NS= Not Specified | | | | | | | | | | | |

Firstly, to compare just Paper [α] and Paper [β]. Their source papers, [91] and [304] respectively, were published 7 years apart, in different peer-reviewed journals, by the same laboratory, wherein the latter paper has two additional authors. Paper [β] specifies its tested species more precisely: *Sus scrofa* ("farm pig"), *Oryctolagus cuniculus* (rabbit), *Columba livia* (rock pigeon, Indian Fantail breed), *Cavia porcellus* (guinea pig), *Rattus norvegicus* (brown rat), *Mus musculus* (mouse). Note the large difference in pig body mass between the two studies. Where Paper [β] presumably reports upon piglets/adolescents rather than adults (I took the middle value of its weight range: "thirty-five to 40-lb. farm pigs" = 17.01 kg). Which *might* be why it reports a greater specific $F_1F_0$ ATP hydrolysis rate for pig. Because, to speculate, piglets/adolescents are smaller than adult pigs, so have a higher surface-area to volume ratio, thence require greater heat generation per unit mass, and so a higher specific basal metabolic rate, driven by greater specific $F_1F_0$ ATP hydrolysis rate.

*From Table S7 above, only using data from the lines that present data from both studies:* The Pearson correlation coefficient (R) between the specific $F_1F_0$ ATP hydrolysis rate data of Paper [α] and Paper [β] is 0.98751 (one-tailed, because the alternative hypothesis is directional, p-value = 0.0001165). This is without applying the error bars. If these are instead applied, R is instead 0.97423. Ignoring the error bars, comparing the specific $F_1F_0$ ATP hydrolysis rate data of Paper [α] and Paper [β]: they are *not* significantly different from each other: independent (two-tailed) t-test statistic = -0.6706, p-value = 0.517661; Mann-Whitney U test (two-tailed): U-value = 9, z-score = -1.36109, p-value = 0.17384; Kruskal-Wallis test: H statistic = 2.0769, p-value = 0.14954; Levene's test of homogeneity of variance: f-ratio = 0.00078, p-value = 0.978277. If the error bar is added or subtracted (operation independently selected in each case) from each datum of Paper [β], whichever manipulation makes each value further from the corresponding value from Paper [α], again the data sets are *not* significantly different from each other: independent (two-tailed) t-test statistic = -0.89801, p-value = 0.390289; Mann-Whitney U test (two-tailed): U-value = 8, z-score = -1.52122, p-value = 0.12852; Kruskal-Wallis test: H statistic = 2.5641, p-value = 0.10931; Levene's test of homogeneity of variance: f-ratio = 0.02019, p-value = 0.889822.

Now to compare, as regards specific $F_1F_0$ ATP hydrolysis rate, for their shared species/genera, Paper [α] with the mean of Papers [β], [γ], [δ], [ε], [ψ]. Ignoring error bars. The Pearson correlation coefficient (R) = 0.9708 (one-tailed, because the alternative hypothesis is directional, p-value = 0.0001375). These two data sets are *not* significantly different from each other: independent (two-tailed) t-test statistic = -0.60998, p-value = 0.553253; Mann-Whitney U test (two-tailed): U-value = 13, z-score = -1.40553, p-value = 0.15854; Kruskal-Wallis test: H statistic = 2.1592, p-value = 0.14172; Levene's test of homogeneity of variance: f-ratio = 0.03268, p-value = 0.859571. Incidentally, the pigeon species used by Paper [ψ] is *Columba livia* (White Carneaux breed).

IF1 protein content

Under normal conditions *in vivo*, the inhibitory capability of a species' IF1 protein population is at least a function of (*a*) how much IF1 protein it has per ATP synthase molecule, (*b*) how potently (how much inhibition, per unit amount) its IF1 protein can inhibit *its* $F_1F_0$ ATP hydrolysis, and (*c*) it's IF1 protein's tendency/self-affinity for tetramerization and higher oligomerization (self-sequesteration) at the alkaline pH of the mitochondrial matrix.

As taught in the Methods section, assaying (at pH 7.8, for 5 minutes) $F_1F_0$ ATP hydrolysis in Sub-Mitochondrial Particles (SMPs), which are derived (by sonication) from mitochondria that have been isolated from an ischemic tissue of the species, reflects upon (*a*), (*b*) and (*c*). Their combination. Enabling species comparisons. Such data has been utilized up until this point (e.g., utilized in Figure 3).

By contrast, this present section uses a more limited (comparatively disfavoured) assay, which doesn't as fully reflect upon (*a*) and (*b*), and that doesn't reflect upon (*c*) at all. Only capturing the IF1 protein quantity (and not its ratio to ATP synthase) aspect of (*a*). And (aside from for rat) not capturing the "*its*" $F_1F_0$ ATP hydrolysis" aspect to (*b*).

This present assay reports the combination of (*i*) IF1 protein amount, with (*ii*) IF1 protein potency against rat $F_1F_0$ ATP hydrolysis. Although in the literature, the latter potency element is overlooked/discounted, and this assay is considered to simply report upon "IF1 protein amount" or "IF1 protein content". Hereafter I align with this terminology, with the caveat of what has been said (elsewhere herein I often refer to this data as "specific IF1 protein activity").

For different species, Table S8 below utilizes "IF1 protein content", which is the amount of mitochondrial extract {measured as mg of mitochondrial protein} from a species that can decrease $F_1F_0$ ATP hydrolysis rate of rat Sub-Mitochondrial Particles (SMPs) by 1 μmol/min. Assay performed at pH 6.2. This assay works because rat has very little endogenous IF1 protein, and so a high rate of $F_1F_0$ ATP hydrolysis. Moreover, because IF1 protein sequences, and ATP synthase protein sequences, are incredibly conserved across species. Such that IF1 proteins are typically interoperable between species: e.g., yeast IF1 protein can inhibit bovine $F_1F_0$ ATP hydrolysis [309].

In Table S8 below, IF1 protein content data for 12 different species is used from Paper [α] [91] (*from column titled "ATPase Inhibitor Content, IU/mg" in its table 1*), and from 6 different species from Paper [β] [304] (*from column titled "IF1 content (I.U./mg)" in its table III. From species: Sus scrofa*, *Oryctolagus cuniculus*, *Columba livia*, *Cavia porcellus*, *Rattus norvegicus*, *Mus musculus*). Pearson correlation coefficients reported, and one-tailed (because each alternative hypothesis is directional) p-values.
[Table S8]

| | | | Specific basal metabolic rate | Heart rate | Maximal lifespan | log10(Body mass) |
|---|---|---|---|---|---|---|
| **Paper [α]** | IF1 protein content | R | -0.6669 | -0.6051 | 0.6261 | 0.486 |
| | | p | 0.0089225 | 0.018549 | 0.019661 | 0.054583 |
| **Paper [β]** | IF1 protein content | R | -0.8211 | -0.9285 | 0.6677 | 0.7488 |
| | | p | 0.0225725 | 0.003743 | 0.073644 | 0.0433635 |

IF1 protein content data used to produce Table S8 above aligns with other data used in this study, supporting conclusions drawn therefrom.

Combining specific $F_1F_0$ ATP hydrolysis rate and IF1 protein content data

The Pearson correlation coefficient between specific $F_1F_0$ ATP hydrolysis rate and IF1 protein content data of [91] is -0.5292. With one-tailed (alternative hypothesis is directional) p-value of 0.038426. So, the greater the IF1 protein content, the lower the specific $F_1F_0$ ATP hydrolysis rate.

Earlier in this work, for specific basal metabolic rate, heart rate, and maximal lifespan: correlations to IF1 protein content were reported, and correlations to specific $F_1F_0$ ATP hydrolysis rate were separately reported. Reported on the row labelled "*Combined*", in Table S9 below, Fisher's combined probability test is used to combine p-values for these two sets of correlations. The independence requirement of Fisher's method is met because specific $F_1F_0$ ATP hydrolysis rate and IF1 protein content are different measures, assayed by different experimental protocols. On lower rows are some correlation p-values presented earlier. Lower in the table, p-values are combined by two different methods: the asymptotically exact harmonic mean p-value for combining independent/dependent tests [76] (*its post-published corrected method, as corrected on 7 October 2019*), and using Fisher's combined probability test.

[Table S9]

| **p-value (one-tailed)** | | | |
|---|---|---|---|
| | Specific basal metabolic rate | Heart rate | Maximal lifespan |
| *Combined* | 0.00053569 | 0.0001955 | 0.00123938 |
| | Specific basal metabolic rate | 0.0000015 | 0.001136 |
| | | Heart rate | 0.0003725 |
| **Asymptotically exact harmonic mean p-value for combining independent/dependent tests** | | | |
| Of all the p-values above (in this present table) = 0.000008850359 = **0.000009** | | | |
| **Fisher's combined probability test** | | | |
| Using p-values for [*Combined* vs. Maximal lifespan] (=0.00123938) & [Heart rate vs. Specific basal metabolic rate] (=0.0000015) = 0.00000004 | | | |

Consistent IF1 protein contents reported by different studies

Comparing corresponding species/genus data (*i.e., comparing corresponding pig, rabbit, guinea pig, rat, mouse, and pigeon data*), the Pearson correlation coefficient (R) between IF1 protein content data from Papers [α] and [β] is 0.8971, one-tailed (alternative hypothesis is directional) p-value is 0.007669.

Comparing corresponding species/genus data, IF1 protein content data from Papers [α] and [β] is *not* significantly different from each other: independent (two-tailed) t-test statistic = -0.42674, p-value = 0 .677802; Mann-Whitney U test (two-tailed): U-value = 14, z-score = 0.56045, p-value = 0.57548; Kruskal-Wallis test: H statistic = 0.4103, p-value = 0.52184; Levene's test of homogeneity of variance: f-ratio = 0.29322, p-value = 0.600018.

The IF1 protein content data of Paper [α] doesn't have error bars. But that of Paper [β] does. If the error bar is added or subtracted (operation independently selected in each case) from each datum of Paper [β], whichever manipulation makes each value further from the corresponding value from Paper [α], again these IF1 protein content data sets are *not* significantly different from each other: independent (two-tailed) t-test statistic = 0.35084, p-value = 0.73299; Mann-Whitney U test (two-tailed): U-value = 14, z-score = 0.56045, p-value = 0.57548; Kruskal-Wallis test: H statistic = 0.4103, p-value = 0.52184; Levene's test of homogeneity of variance: f-ratio = 0.32067, p-value = 0.58369.

Tyranny of the midwit

Of course, "correlation does not *necessarily* imply causation", but we mustn't succumb to the common fallacy of bastardizing this to be "correlation does not imply causation" and dismissing any correlation out of hand. The causal reason(s) why different species have different maximal lifespans is likely a positive or negative correlate of maximal lifespan across species. Dismissing a correlate for further study precisely because it is a correlate isn't wise.

Bradford Hill criteria

Historically, the Bradford Hill criteria (also known as Hill's criteria for causation) [309] were used to persuade that lung cancer *correlates* with smoking in epidemiological data, because smoking *causes* lung cancer. Incidentally, this was contested by the great statistician Ronald Fisher ("an error has been made, of an old kind, in arguing from correlation to causation"

[311]). The present work reports correlation between more IF1 protein activity (lower $F_1F_0$ ATP hydrolysis rate) and longer maximal lifespan. This correlation, although not epidemiological, satisfies (at least the spirit of) at least most of the Bradford Hill criteria. Moreover, distinctly, the present work is confirmatory rather than exploratory (*plus is supported by interventional as well as correlational data*).

This was a confirmatory study of a predefined hypothesis

The multi-pronged hypothesis, that IF1 protein activity (by curtailing $F_1F_0$ ATP hydrolysis rate, dictating specific basal metabolic rate [which sets heart rate]) is a molecular determinant of maximal lifespan, was set *before* the supporting data was collected. So, in statistical terminology, this study is “confirmatory” (hypothesis before data, so data tests the hypothesis, in this case validates the hypothesis). Rather than “exploratory” (data before hypothesis, and hypothesis generated from the data).

Predation as *the* determinant of maximal lifespan?

Why does the tiger, an apex predator, maximally live 26.3 years [1] and not >100 years, or >200 years? Why does the tiger have a shorter maximal lifespan than its water buffalo prey?

Larger mammal species may have a greater maximal lifespan because of less predation pressure, conferring margin for a longer maximal lifespan to be selected for. The teaching of this work isn’t contrary to that, as it acts in the same direction, conferring a (candidate) molecular mechanism to it. Being larger may simultaneously confer (*a*) margin for a longer maximal lifespan (by less predation pressure), and (*b)* the means of achieving it (by permitting a lower specific basal metabolic rate).

So, the present teaching isn’t inconsistent with predation pressure theory, but it isn’t dependent on it either. The theories aren’t mutually exclusive. Predation pressure theory has a mechanistic hollow, which can be filled by the present teaching, or this teaching can stand alone, *sui generis*, without predation pressure theory. Either way, it can explain observations that predation pressure theory can’t.

Mammalian apex predators have a constrained maximal lifespan, following the same relation to body mass as their prey/non-predators: e.g., maximal lifespan of a tiger, *Panthera tigris*, at 26.3 years [1], isn’t much greater than that predicted, 25.5 years, from its adult body mass,

119.7 kg [1], by the relevant best-fit line equation of Figure 3 in the main text, which was derived from mostly non-predator species. The appropriate best-fit line equation of Figure 3, appropriately rearranged, is shown below, where *y* is maximal lifespan (years), and *x* is body mass (kg):

$$y = 10^{0.98595+0.20246 \cdot log10(x)} \qquad \text{[Eq. S2]}$$

Moreover, the observed isn't any greater than prediction at all when the error bars in one, or both, of the equation's constants (0.98595±0.06233, 0.20246±0.04528) are factored in.

Table S13 below relates to some well-known mammalian apex predators, showing that their observed maximal lifespan [1] isn't much different than predicted, by this equation, from their adult body mass (adult body mass values sourced from [1]):

[Table S13]

| Species | Observed maximal lifespan [1] | Predicted maximal lifespan | Closest prediction given error bars in prediction |
|---|---|---|---|
| Tiger*, Panthera tigris* | 26.3 | 25.5 | 26.3 |
| Jaguar, *Panthera onca* | 28 | 23.6 | 28 |
| Lion, *Panthera leo* | 28 | 27.5 | 28 |
| Polar bear, *Ursus maritimus* | 43.8 | 33.7 | 43.8 |
| Black bear, *Ursus americanus* | 34 | 26.9 | 34 |
| Brown bear, *Ursus arctos* | 40 | 30.2 | 40 |
| Orca, *Orcinus orca* | 90 | 51.9 | 87.2 |
| Gray wolf, *Canis lupus* | 20.6 | 18.8 | 20.6 |
| Cheetah*, Acinonyx jubatus* | 20.5 | 20.3 | 20.5 |
| Leopard, *Panthera pardus* | 27.3 | 21.7 | 27.3 |
| Cougar, *Puma concolor* | 27 | 22.4 | 27 |

If maximal lifespan is set by predation pressure, one would expect predators, especially apex predators, to have a longer maximal lifespan than their prey. But this is not observed in at least the following cases. The maximal lifespan of a plains zebra (*Equus quagga*, 38 years) is greater than that of its predator the lion (*Panthera leo*, 28 years) [1]. While this is inconsistent with predation pressure setting maximal lifespan, it is consistent with the teaching of the present work. As a plains zebra (*Equus quagga*, 280 kg) is larger than a lion (*Panthera leo*,

175 kg) [1]. So, upon the data herein (Figure 14), I predict that it has less metabolic heat generation per unit mass per unit time, thence a lower specific basal metabolic rate, a prediction as there is no metabolic data for any species of zebra in [1] yet, which (to predict) is why it has a slower aging rate, and a longer maximal lifespan.

Another prey of lion, which is larger and has a longer maximal lifespan than the lion, is the African buffalo (*Syncerus caffer*), which has an adult body weight of 700 kg and a maximal lifespan of 32.8 years [1]. The giraffe can be prey of lion also, which again is larger (800 kg) and has a longer maximal lifespan (39.5 years) than the lion [1]. Similarly, the water buffalo (*Bubalus bubalis*) is larger (725 kg) and has a longer maximal lifespan (34.9 years) than its predator the tiger (119.7 kg, 26.3 years) [1]. Moreover, the red deer (*Cervus elaphus*) is larger (200 kg) and has a longer maximal lifespan (31.5 years) than its predator *Canis lupus*, the gray wolf (26.6 kg, 20.6 years) [1]. The red kangaroo (*Macropus rufus*) is larger (55 kg) and has a longer maximal lifespan (25 years) than its predator the *Canis lupus dingo* (26.6 kg, 20.6 years) [1]. The observed maximal lifespan of each of these species is as predicted from their adult body mass by (within its margin of error) the equation presented earlier in this section.

Generally, in an ecosystem, a larger predator has fewer individuals (across species) that can predate it, and it has more individuals (across species) that it can predate. In ecosystems, by Damuth's law, there tend to be fewer individuals of larger species: (unlogged) population density of a species is a negative convex (i.e., a supra-linear) function of adult body mass [312] (*possibly compounded by ecosystems tending to have more smaller species than larger species [313-315], but perhaps not because this may only be observed at large spatial scale*). So, predation pressure is increasingly less with greater adult body mass and therefore, if predation pressure sets maximal lifespan, maximal lifespan should be increasingly more with greater adult body mass: across mammal species, maximal lifespan should be a super-linear function of adult body mass. Especially because apex predators, which have no predators themselves, should (by this theory) have an extremely high maximal lifespan, because they have *no* predation constraint to their maximal lifespan. However, across mammal species, maximal lifespan is instead observed to be a supra-linear (concave) function of adult body mass. The present teaching offers explanation, at molecular resolution. Refer to Figure 14, and its associated text.

Another way of stating predation pressure theory is that smaller species age faster than larger species, but this theory doesn't explain why. Why don't they age at the same rate as larger species, or slower? Why does predation pressure select for faster aging? Why *must* they age faster?

An implicit assumption of predation pressure theory is that mammals die biologically young in the wild, but this isn't true, many mammal individuals die biologically old in the wild [316]. For example, by predation. Older and slower increases the risk of being predated, especially as a well-known predator strategy is to specifically target such individuals. This breached assumption of predation pressure theory gravely undermines it, perhaps fatally.

When the theory incorporates many prey individuals dying biologically old in the wild, especially via predation, it predicts runaway instability, as predation pressure drives faster aging (its core tenant), faster aging increases predation pressure, so aging rate increasingly increases, and maximal lifespan increasingly decreases over time. But this hasn't been observed.

Slower aging individuals of a species, because they can reproduce for longer because of longer fertility, vigour, and lifespan, should have more reproductive success (more, and better cared for, offspring than their peers. Not least for prey species because slower aging individuals face lower predation pressure, because predators favour, and have more success, catching old individuals), such that slower aging is selected for, until nearly all individuals *do* die biologically young. Why doesn't this happen? Predation pressure theory can't explain this, whereas the present teaching (conceivably) can. At molecular resolution.

Predation pressure theory of maximal lifespan in mammals lacks molecular basis. The present teaching has (drug-falsifiable) molecular basis, which isn't inconsistent with predation pressure theory, which can explain things that predation pressure theory can't. Such as why smaller species age faster than larger species, why apex predators (which have *no* predators, and so *no* predation constraint to their lifespan) don't live for *much* longer than they do, and why some predators, even some apex predators, have shorter maximal lifespans than their prey. Furthermore, why maximal lifespan is a supra- and not a super-linear function of species body mass, and why many mammals die biologically old in the wild. Predation pressure theory is, at least alone, insufficiently explanatory for the data. Indeed, compared to a mainland population with predators, an island population of the same species without predators has a maximal lifespan 1.2 years higher [219] (*why not much higher?*). By contrast, with some other species, a population with predators has the same [220] or a higher [221-222] lifespan than a population without predators (see also [223]).

Any theory of aging that (unlike the present teaching) can't explain why and how different mammal species have different maximal lifespans and unparsimoniously outsources this *key* observation completely to predation pressure theory inherits its insufficiency identified here.

Why does the lion, an apex predator, not have a high/limitless maximal lifespan, but instead a shorter maximal lifespan than its prey the zebra?

<u>Across mammal species, maximal lifespan inversely correlates with body surface-area per unit body mass</u>

As shown in Table S10 below, which shows Pearson correlation coefficients (R), and associated p-values (one-tailed, because each alternative hypothesis is directional). Where *A* = body surface-area ($m^2$), *B* = body mass (kg), *C* = body surface-area per unit body mass ($m^2$/kg), *D* = maximal lifespan (years), *E* = specific $F_1F_0$ ATP hydrolysis rate (μmol/min/mg), *F* = specific basal metabolic rate (W/g), *G* = heart rate (beats per minute, bpm), where *A*, *B* and *C* use data from [128], other variables are as disclosed prior in Table S2, for those species overlapping with those in [128]. *E*, *F* and *G* positively correlate with *C*, which is perhaps why *D* negatively correlates with *C*.

[Table S10]

| **R values** | | | | |
|---|---|---|---|---|
| | D | E | F | G |
| A | 0.4389 | -0.3883 | -0.4886 | -0.5283 |
| B | 0.331 | -0.2999 | -0.4114 | -0.4445 |
| C | -0.7855 | 0.7416 | 0.9877 | 0.9431 |
| **p-values** | | | | |
| | D | E | F | G |
| A | 0.138319 | 0.170906 | 0.109619 | 0.089158 |
| B | 0.211608 | 0.235247 | 0.155621 | 0.134918 |
| C | 0.010437 | 0.017603 | 2.5E-06 | 0.000221 |

Table S11 below is the same as Table S10 above except the correlation values are, instead of Pearson, Spearman's rank correlation coefficients (ρ).

[Table S11]

| ρ values | | | | |
|---|---|---|---|---|
| | D | E | F | G |
| A | 0.7143 | -0.6667 | -0.9524 | -0.9762 |
| B | 0.7857 | -0.7143 | -0.9762 | -1 |
| C | -0.7381 | 0.619 | 0.9286 | 0.9762 |
| **p-values** | | | | |
| | D | E | F | G |
| A | 0.023261 | 0.035484 | 0.00013 | 1.65E-05 |
| B | 0.01041 | 0.023261 | 1.65E-05 | 0 |
| C | 0.018276 | 0.050884 | 0.000431 | 1.65E-05 |

In Figure S1 below, across 45 non-flying mammal species, maximal lifespan (in years, values sourced from [1]) is a supra-linear (negative convex) function of body surface-area per unit body mass ($m^2$/kg, experimental data of [128]). *{N.B. Where [128] states "Cavia" I assume they refer to Cavia porcellus. I didn't use the "rhinoceros" and "Peromyscus" data of [128] because it is ambiguous which species they are referring to by those terms. These points apply at all points that I use [128] data herein}.*

Pearson correlation coefficient = -0.3663, one-tailed (alternative hypothesis is directional) p-value = 0.0066665. Corresponding Spearman's rank correlation coefficient = -0.8705, one-tailed p-value = 3.98053E-15.

*If outlier Mustela frenata (the long-tailed weasel, that has a very long tail compared to its body length, which makes its surface-area anomalously large for its body mass) is omitted: Pearson = -0.5109 (0.0001975), Spearman = -0.8678 (1.22658E-14).*

*if all the non-terrestrial mammal species, which live in water all or some of the time, are omitted also: Pearson = -0.5934 (0.0003455), Spearman = -0.8375 (7.33747E-09).*

*For all the 45 species, using unlogged data, with GraphPad Prism 10.0.3 software with its default settings (using least square regression. With chosen starting values: a=1, b=0.25): supra-linear equation $y = a \cdot x^b$ was curve-fitted, which best-fitted with a = 5.978, b = -0.4431, wherein adjusted $R^2$ = 0.8105. A linear equation (which does not fit as well as the [aforementioned] supra-linear equation with same number of parameters): $y = m \cdot x + c$, where m = -41.02, c = 32.28, and adjusted $R^2$ = 0.1140. When x and y are both logged, a linear equation fits well: $log_{10}(y) = m \cdot log_{10}(x) + log_{10}(c)$, where m = -0.5533, $log_{10}(c)$ = 0.5544, and adjusted $R^2$ = 0.7215.*

*When outlier Mustela frenata is omitted, rendering 44 species: $y = a \cdot x^b$, wherein a = 5.954, b = -0.4438; adjusted $R^2$ = 0.8084.*

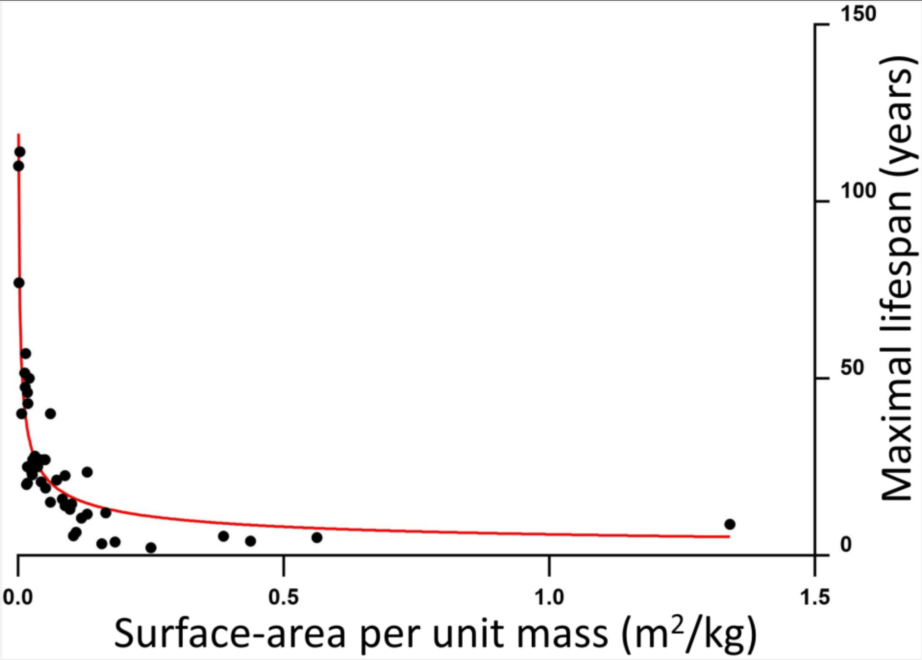

Figure S1 – Across 45 mammal species, maximal lifespan (years) is inversely proportional to (a negative convex function of) body surface-area per unit body mass ($m^2$/kg).

The outlier data point, with very high surface-area per unit mass, is for the long-tailed weasel (*Mustela frenata*), which has a very long tail compared to its body length.

*Fitted line is of the form* $y = a \cdot x^b$, *wherein a = 5.978, b = -0.4431; adjusted* $R^2$ *= 0.8105.*

Using body surface-area per unit body mass ($m^2$/kg) data for non-flying mammal species from [128], those 27 species thereof that I could source specific basal metabolic rate (W/g) data from [1]: specific basal metabolic rate is a linear function of body surface-area per unit body mass, wherein Pearson correlation coefficient = 0.4269, one-tailed (alternative hypothesis is directional) p-value = 0.013183. Corresponding Spearman's rank correlation coefficient = 0.7448, one-tailed p-value = 0.000004.

When outlier *Mustela frenata* is omitted (rendering 26 species): Figure S2: Pearson correlation coefficient = 0.7134, one-tailed (alternative hypothesis is directional) p-value = 0.0000215. Corresponding Spearman's rank correlation coefficient = 0.7292, one-tailed p-value = 0.000012.

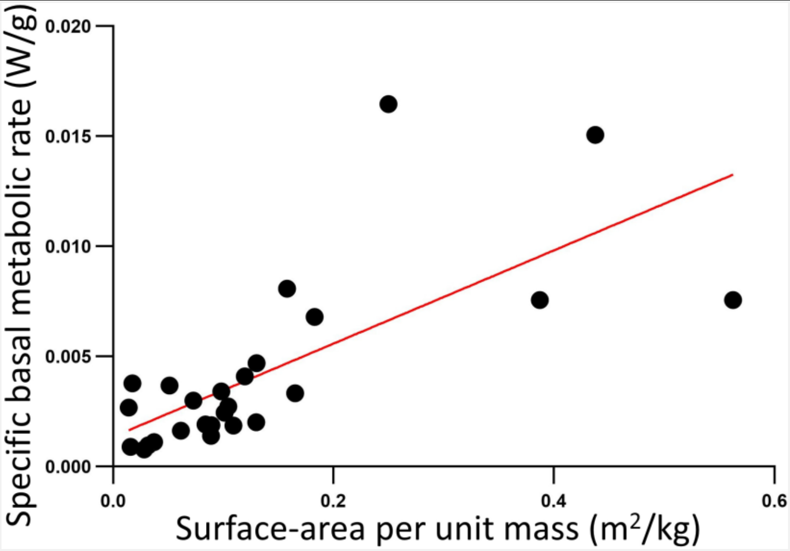

Figure S2 – Across 26 mammal species, specific basal metabolic rate (W/g) is a linear function of body surface-area per unit body mass ($m^2$/kg).

*Fitted line is of the form* $y = m \cdot x + c$, *where m = 0.02117, c = 0.001347; adjusted* $R^2$ *= 0.4885.*

Across homethermic species, maximal lifespan is inversely proportional to specific basal metabolic rate

The Pearson correlation coefficient can discriminate a supra-linear from a linear function, by conferring the former with a lower value, whereas Spearman's rank correlation coefficient cannot and doesn't. When the Spearman is much more than the corresponding Pearson coefficient, this might indicate that the relation is supra-linear. Although that Spearman can be less impacted (positively or negatively) by one or more outliers is another factor to consider. Below, each of the maximal lifespan (*y*) vs. adult body mass (*x*) plots are supra-linear (positive concave), each of the specific basal metabolic rate (*y*) vs. adult body mass (x) plots are supra-linear (negative convex), and each of the maximal lifespan (*y*) vs. specific basal metabolic rate (*x*) plots are supra-linear (negative convex), which is probably why in each case the Spearman is much greater than the Pearson. For the analysis of this section, units are maximal lifespan (years), adult body mass (kg), and specific basal metabolic rate (W/g).

For *all* the **MAMMAL** species that presently have adult body mass, specific basal metabolic rate, and maximal lifespan data in [1], which is 350 mammal species: (*a*) maximal lifespan is proportional to adult body mass, which might be because across these same mammal species: (*b*) specific basal metabolic rate is inversely proportional to adult body mass, and thence (*c*) maximal lifespan is inversely proportional to specific basal metabolic rate. The correlations in turn for (*a*), (*b*), and (*c*) are listed below, all statistically significant, at least some incredibly so:

**(*a*)** Supra-linear (positive concave) shaped plot.

Pearson correlation coefficient = 0.3834, one-tailed (alternative hypothesis is directional) p-value = 5.30633E-14.

Spearman's rank correlation coefficient = 0.6694, one-tailed p-value = 3.86456E-47.

*Using unlogged data with GraphPad Prism 10.0.3 software with its default settings (using least square regression. With selected starting values: a = 1, b = 0.25, which were used for other fittings of this equation herein also): supra-linear equation* $y = a \cdot x^b$ *was curve-fitted, which best-fitted with a = 15.02, b = 0.1659, wherein adjusted* $R^2$ *= 0.4189. Simplifying this equation worsened the* $R^2$*, making it negative (!): (starting value a=0.25) supra-linear equation* $y = x^a$ *best fitted with a = 0.5841, wherein adjusted* $R^2$ *= -0.7747. Adding more terms could better the fit, at the "cost" of more parameters: power series equation:* $y = a \cdot x^b + c \cdot x^d$*: adjusted* $R^2$ *= 0.4272. A linear equation (which does not fit as well as the [aforementioned] supra-linear equation with same number of parameters):* $y = m \cdot x + c$*, where m = 0.02545, c = 14.42, and adjusted* $R^2$ *= 0.1445. When x and y are both logged, a linear equation fits well:* $log_{10}(y) = m \cdot log_{10}(x) + log_{10}(c)$*, where m = 0.1795,* $log_{10}(c)$ *= 1.115, and adjusted* $R^2$ *= 0.4485.*

**(*b*)** Supra-linear (negative convex) shaped plot.

Pearson correlation coefficient = -0.1182, one-tailed (alternative hypothesis is directional) p-value = 0.013497096.

Spearman's rank correlation coefficient = -0.8889, one-tailed p-value = 2.4789E-120.

*Using unlogged data, supra-linear equation* $y = a \cdot x^b$ *was curve-fitted, which best-fitted with a = 0.002611, b = -0.3404, wherein adjusted* $R^2$ *= 0.6261. A linear equation does not fit as well:* $y = m \cdot x + c$*, where m = -2.955e-006, c = 0.004900, and adjusted* $R^2$ *= 0.01114. When x and y are both logged, a linear equation fits well:* $log_{10}(y) = m \cdot log_{10}(x) + log_{10}(c)$*, where m = -0.2599,* $log_{10}(c)$ *= -2.537, and adjusted* $R^2$ *= 0.7391.*

**(*c*)** Supra-linear (negative convex) shaped plot.

Pearson correlation coefficient = -0.4158, one-tailed (alternative hypothesis is directional) p-value = 2.29546E-16.

Spearman's rank correlation coefficient = -0.6389, one-tailed p-value = 7.61166E-42.

*Using unlogged data, supra-linear equation* $y = a \cdot x^b$ *was curve-fitted, which best-fitted with a = 0.6999, b = -0.5248, wherein adjusted* $R^2$ *= 0.3727. A linear equation does not fit as well:* $y = m \cdot x + c$*, where m = -1104, c = 20.39, and adjusted* $R^2$ *= 0.1705. When x and y are both logged, a linear equation fits well:* $log_{10}(y) = m \cdot log_{10}(x) + log_{10}(c)$*, where m = -0.5763,* $log_{10}(c)$ *= -0.3556, and adjusted* $R^2$ *= 0.4222.*

For *all* the **BIRD** species that presently have adult body mass, specific basal metabolic rate, and maximal lifespan data in [1], which is 167 bird species: (*a*) maximal lifespan is proportional to adult body mass, which might be because across these same bird species: (*b*) specific basal metabolic rate is inversely proportional to adult body mass, and thence (*c*) maximal lifespan is inversely proportional to specific basal metabolic rate. The correlations in turn, all statistically significant, at least some incredibly so:

(***a***) Supra-linear (positive concave) shaped plot.

Pearson correlation coefficient = 0.2742, one-tailed (alternative hypothesis is directional) p-value = 0.000167739.

Spearman's rank correlation coefficient = 0.7021, one-tailed p-value = 2.04274E-26.

*Using unlogged data, supra-linear equation* $y = a \cdot x^b$ *was curve-fitted, which best-fitted with a = 27.65, b = 0.1903, wherein adjusted* $R^2$ *= 0.4928. A linear equation does not fit as well:* $y = m \cdot x + c$*, where m = 0.3777, c = 17.62, and adjusted* $R^2$ *= 0.06956. When x and y are both logged, a linear equation fits well:* $log_{10}(y) = m \cdot log_{10}(x) + log_{10}(c)$*, where m = 0.2094,* $log_{10}(c)$ *= 1.417, and adjusted* $R^2$ *= 0.5111.*

(***b***) Supra-linear (negative convex) shaped plot.

Pearson correlation coefficient = -0.1928, one-tailed (alternative hypothesis is directional) p-value = 0.006273745.

Spearman's rank correlation coefficient = -0.923, one-tailed p-value = 1.23038E-70.

*Using unlogged data, supra-linear equation* $y = a \cdot x^b$ *was curve-fitted, which best-fitted with a = 0.004587, b = -0.3104, wherein adjusted* $R^2$ *= 0.8298. A linear equation does not fit as well:* $y = m \cdot x + c$*, where m = -0.0001482, c = 0.01199, and adjusted* $R^2$ *= 0.03133. When x and y are both logged, a linear equation fits well:* $log_{10}(y) = m \cdot log_{10}(x) + log_{10}(c)$*, where m = 0.2094,* $log_{10}(c)$ *= 1.417, and adjusted* $R^2$ *= 0.5111.*

(***c***) Supra-linear (negative convex) shaped plot.

Pearson correlation coefficient = -0.5994, one-tailed (alternative hypothesis is directional) p-value = 5.74211E-18.

Spearman's rank correlation coefficient = -0.6382, one-tailed p-value = 8.76304E-21.

*Using unlogged data, supra-linear equation* $y = a \cdot x^b$ *was curve-fitted, which best-fitted with a = 1.712, b = -0.4939, wherein adjusted* $R^2$ *= 0.4300. A linear equation does not fit as well:* $y = m \cdot x + c$, *where m = -1074, c = 30.74, and adjusted* $R^2$ *= 0.3554. When x and y are both logged, a linear equation fits well:* $log_{10}(y) = m \cdot log_{10}(x) + log_{10}(c)$, *where m = -0.5321,* $log_{10}(c)$ *= 0.1086, and adjusted* $R^2$ *= 0.4474.*

Pooling the **MAMMAL AND BIRD** species data: For *all* the mammal and bird species that presently have adult body mass, specific basal metabolic rate, and maximal lifespan data in [1], which is 517 species: (*a*) maximal lifespan is proportional to adult body mass, which might be because across these same species: (*b*) specific basal metabolic rate is inversely proportional to adult body mass, and thence (*c*) maximal lifespan is inversely proportional to specific basal metabolic rate. The correlations in turn, all statistically significant, at least some incredibly so:

(***a***) Supra-linear (positive concave) shaped plot.

Pearson correlation coefficient = 0.3099, one-tailed (alternative hypothesis is directional) p-value = 2.84701E-13.

Spearman's rank correlation coefficient = 0.5651, one-tailed p-value = 2.9568E-45.

*Using unlogged data, supra-linear equation* $y = a \cdot x^b$ *was curve-fitted, which best-fitted with a = 17.78, b = 0.1376, wherein adjusted* $R^2$ *= 0.3205. A linear equation does not fit as well:* $y = m \cdot x + c$, *where m = 0.02498, c = 15.59, and adjusted* $R^2$ *= 0.09430. When x and y are both logged, a linear equation fits well:* $log_{10}(y) = m \cdot log_{10}(x) + log_{10}(c)$, *where m = 0.1482,* $log_{10}(c)$ *= 1.184, and adjusted* $R^2$ *= 0.3214. This latter plot is shown in Figure S3.*

(***b***) Supra-linear (negative convex) shaped plot.

Pearson correlation coefficient = -0.1138, one-tailed (alternative hypothesis is directional) p-value = 0.004808816.

Spearman's rank correlation coefficient = -0.8999, one-tailed p-value = 4.7379E-188.

*Using unlogged data, supra-linear equation* $y = a \cdot x^b$ *was curve-fitted, which best-fitted with a = 0.003119, b = -0.3567, wherein adjusted* $R^2$ *= 0.7080. A linear equation does not fit as well:* $y = m \cdot x + c$, *where m = -4.736e-006, c = 0.007170, and adjusted* $R^2$ *=*

*0.01103. When x and y are both logged, a linear equation fits well:* $log_{10}(y) = m \cdot log_{10}(x) + log_{10}(c)$*, where m = -0.3056,* $log_{10}(c)$ *= -2.488, and adjusted* $R^2$ *= 0.7738. This latter plot is shown in Figure S3.*

**(*c*)** Supra-linear (negative convex) shaped plot.

Pearson correlation coefficient = -0.3465, one-tailed (alternative hypothesis is directional) p-value = 2.47943E-16.

Spearman's rank correlation coefficient = -0.4505, one-tailed p-value = 1.66592E-27.

*Using unlogged data, supra-linear equation* $y = a \cdot x^b$ *was curve-fitted, which best-fitted with a = 1.907, b = -0.3854, wherein adjusted* $R^2$ *= 0.2347. A linear equation does not fit as well:* $y = m \cdot x + c$*, where m = -671.1, c = 20.78, and adjusted* $R^2$ *= 0.1183. When x and y are both logged, a linear equation fits well:* $log_{10}(y) = m \cdot log_{10}(x) + log_{10}(c)$*, where m = -0.3434,* $log_{10}(c)$ *= 0.3056, and adjusted* $R^2$ *= 0.2076. This latter plot is shown in Figure S3.*

The asymptotically exact harmonic mean p-value [76] (its post-published corrected method, as corrected on 7 October 2019), using the p-value for the Spearman's rank correlation coefficient for each of (*a*), (*b*), and (*c*), for the 517 mammal and bird species data set = **1.42137e-187**. Extremely significant.

Note that the supra-linear functions have parts that are nearly linear functions and so can be misclassified as linear functions if we only have data for one of those parts. For example, considering the (mammals and birds) maximal lifespan (*y*) vs. specific basal metabolic rate (*x*) plot, once the specific basal metabolic rate exceeds around 0.006 W/g it is nearly a linear relation. And so, if we only had *x* data above this value, we might incorrectly surmise that the relation is linear.

Using mammal data, if *y* values of specific basal metabolic rate and maximal lifespan are determined from *x* (adult body mass) using their respective (aforementioned) $y = a \cdot x^b$ best-fit line equations, and these calculated values of specific basal metabolic rate and maximal lifespan are then plotted, as *x* and *y* respectively, the plot is supra-linear (negative convex), with a best-fit line equation of $y = a \cdot x^b$, where *a* = 0.8272, *b* = -0.4874 (adjusted $R^2$ = 1), wherein these *a* and *b* values are not so dissimilar to those observed for this best-fit line equation fitted to the observed (rather than calculated) values plot of specific basal metabolic rate and maximal lifespan (aforementioned): $a$ = 0.6999, $b$ = -0.5248. A nice confirmatory result.

Homeotherm body mass isn't the only factor dictating how much metabolic heat generation per unit mass per unit time is required to maintain body temperature. There is also such a species' body shape (its surface-area, and how this is manipulated by behaviours such as curling, huddling, wing-folding, etc.), body insulation (innate such as fat tissue, fur, feathers, etc., which can be changeable, .e.g., with season, and external such as whether a nest is made with insulating materials, etc.), vascular control (congruent with body insulation, it setting how insulated the blood is), degree of blood counter-current architecture in body extremities such as limbs (wherein blood going outbound to an extremity is facilitated, by proximity, to transfer heat to blood coming inbound from that extremity), the typical ambient temperature (range) this species inhabits, etc. Don't underestimate the power of bodily insulation especially, which some species can change dramatically, which for example is largely what permits an arctic fox to be thermocomfortable at -40°C (!) [178, 317]. Moreover, not all homeotherm species maintain the same body temperature: e.g., naked mole rat body temperature is around 32°C [1]; e.g., some bat species don't endogenously maintain their body temperature during sleep (or hibernation), becoming poikilothermic then [199].

Curtailing the mammalian data set: **MAMMALS without bats**, which renders 318 mammal species remaining (*the absolute differences in brackets are as compared to when bats are included instead of excluded*):

***(a)*** Pearson correlation coefficient = 0.3968 (*i.e., |0.0134| more*).
Spearman's rank correlation coefficient = 0.77 (*i.e., |0.1006| more*).

***(b)*** Pearson correlation coefficient = -0.1159 (*i.e., |0.0023| less*).
Spearman's rank correlation coefficient = -0.8808 (*i.e., |0.0081| less*).

***(c)*** Pearson correlation coefficient = -0.4442 (i.e., |*0.0284*| *more*).
Spearman's rank correlation coefficient = -0.7238 (*i.e., |0.0849| more*).

Curtailing the mammalian data set: **MAMMALS without bats, naked mole rat, and marsupials** (these may be excluded on the grounds they are anomalous as discussed in the Discussion section of the main text), which renders 263 mammal species remaining (*the absolute differences in brackets are as compared to when bats, naked mole rat, and marsupials are included instead of excluded)*:

***(a)*** Pearson correlation coefficient = 0.4026 (*i.e., |0.0192| more*).

Spearman's rank correlation coefficient = **<u>0.8218</u>** (*i.e., |0.1524| more*), one-tailed (alternative hypothesis is directional) p-value = **5.20908E-66**.

*Equation* $y = a \cdot x^b$ *best-fitted with a = 14.87, b = 0.1733, wherein adjusted* $R^2$ *= 0.4896. When x and y are logged (*$log_{10}$*), equation* $log_{10}(y) = m \cdot log_{10}(x) + log_{10}(c)$ *best-fitted with m = 0.2149 (±0.0196; ≈* ***0.25****),* $log_{10}(c)$ *= 1.108 (±0.025), adjusted* $R^2$ *= 0.6400.*

***(b)*** Pearson correlation coefficient = -0.1256 (*i.e., |0.0074| more*).

Spearman's rank correlation coefficient = **<u>-0.8842</u>** (*i.e., |0.0047| less*), one-tailed (alternative hypothesis is directional) p-value = **1.45704E-88**.

*Equation* $y = a \cdot x^b$ *best-fitted with a = 0.002453, b = -0.3958, wherein adjusted* $R^2$ *= 0.7095. When x and y are logged (*$log_{10}$*), equation* $log_{10}(y) = m \cdot log_{10}(x) + log_{10}(c)$ *best-fitted with m =* ***-0.2668*** *(±0.0195; ≈* ***-0.25****),* $log_{10}(c)$ *= -2.515 (±0.024), adjusted* $R^2$ *= 0.7355.*

***(c)*** Pearson correlation coefficient = -0.4826 (i.e., *|0.0668| more*).

Spearman's rank correlation coefficient = **<u>-0.7877</u>** (*i.e., |0.1488| more*), one-tailed (alternative hypothesis is directional) p-value = **3.84467E-57**.

*Equation* $y = a \cdot x^b$ *best-fitted with a = 0.6802, b = -0.5344, wherein adjusted* $R^2$ *= 0.4483. When x and y are logged (*$log_{10}$*), equation* $log_{10}(y) = m \cdot log_{10}(x) + log_{10}(c)$ *best-fitted with m = -0.6780 (±0.0651),* $log_{10}(c)$ *= -0.6017 (±0.1631), adjusted* $R^2$ *= 0.6155. This plot is shown in Figure 12 in the Results section of the main text.*

The asymptotically exact harmonic mean p-value [76] (*its post-published corrected method, as corrected on 7 October 2019*) for the three p-values shown above for this 263 mammal species data set = 4.37112e-88. Extremely significant.

Curtailing the mammalian data set: **<u>MAMMALS without bats, naked mole rat, marsupials, human, and monotremes</u>** (human excluded by rationale given in Methods section of main text, and monotremes because they are such anomalous mammals [e.g., they lay eggs]), which renders 259 mammal species remaining:

***(a)*** Pearson correlation coefficient = 0.4801.

Spearman's rank correlation coefficient = **0.8214**.

***(b)*** Pearson correlation coefficient = -0.127.

Spearman's rank correlation coefficient = **-0.8857**.

***(c)*** Pearson correlation coefficient = -0.5322.

Spearman's rank correlation coefficient = **-0.7811**.

The specific basal metabolic rate value for a shrew (family of Soricidae) may not be a true specific basal metabolic rate value because it might not be possible to get such a value for shrews. Because they become hyperactive when postabsorptive [129]. And so postabsorptive and inactive, both requirements for a basal metabolic rate value, might be mutually exclusive in shrews. Curtailing the mammalian data set: **MAMMALS without bats, naked mole rat, marsupials, human, monotremes, and shrews**, which renders 249 mammal species remaining:

***(a)*** Pearson correlation coefficient = 0.4854.

Spearman's rank correlation coefficient = **0.8052**.

***(b)*** Pearson correlation coefficient = -0.1514.

Spearman's rank correlation coefficient = **-0.8736**.

***(c)*** Pearson correlation coefficient = -0.5767.

Spearman's rank correlation coefficient = **-0.7577**.

A criterion for a basal metabolic rate is that the animal should be postabsorptive but this might be a difficult state to achieve in ruminants because of their reliance on microbial fermentation of cellulose, which may go on for a long period beyond the ruminant's last meal. So, the specific basal metabolic rate values used for such species (of at least the suborder Ruminantia, of the suborder Tylopoda, and of the order Artiodactyla) herein may not be their true specific basal metabolic rate values, being higher, and are used with this caveat. The same issue may apply to pseudoruminants, which includes both foregut fermenters (e.g., of the family Macropodidae) and hindgut fermenters (e.g., of the order Lagomorpha). Curtailing the mammalian data set: MAMMALS without Artiodactyla, Macropodidae, Lagomorpha, which renders 320 mammal species remaining:

***(a)*** Pearson correlation coefficient = 0.3737.

Spearman's rank correlation coefficient = 0.6407.

***(b)*** Pearson correlation coefficient = -0.09046.

Spearman's rank correlation coefficient = -0.8855.

***(c)*** Pearson correlation coefficient = -0.4008.

Spearman's rank correlation coefficient = -0.61.

Table S12 below shows some of the (aforementioned) Pearson (R) and Spearman's rank (ρ) correlation coefficients of this section. The greatest (absolute) correlation coefficient between maximal lifespan and specific basal metabolic rate (-0.7877) is bolded.

[Table S12]

| | Set | | Adult body mass | Specific basal metabolic rate |
|---|---|---|---|---|
| MAXIMAL LIFESPAN | Mammals | *R* | 0.3834 | -0.4158 |
| | | ***ρ*** | 0.6694 | -0.6389 |
| | Birds | *R* | 0.2742 | -0.5994 |
| | | ***ρ*** | 0.7021 | -0.6382 |
| | Mammals + Birds | *R* | 0.3099 | -0.3465 |
| | | ***ρ*** | 0.5651 | -0.4505 |
| | Mammals - bats | *R* | 0.3968 | -0.4442 |
| | | ***ρ*** | 0.77 | -0.7238 |
| | Mammals – (bats, marsupials, naked mole rat) | *R* | 0.4026 | -0.4826 |
| | | ***ρ*** | 0.8218 | **-0.7877** |
| | Mammals – (bats, marsupials, naked mole rat, human, monotremes) | *R* | 0.4801 | -0.5322 |
| | | ***ρ*** | 0.8214 | -0.7811 |
| | Mammals – (bats, marsupials, naked mole rat, human, monotremes, shrews) | *R* | 0.4854 | -0.5767 |
| | | ***ρ*** | 0.8052 | -0.7577 |
| | Mammals – (Artiodactyla, Macropodidae, Lagomorpha) | *R* | 0.3737 | -0.4008 |
| | | ***ρ*** | 0.6407 | -0.61 |

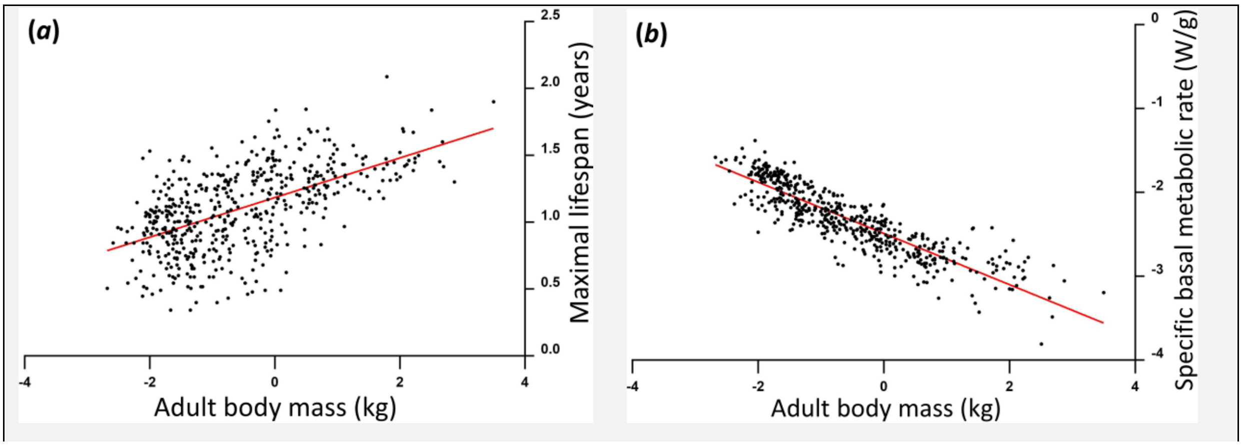

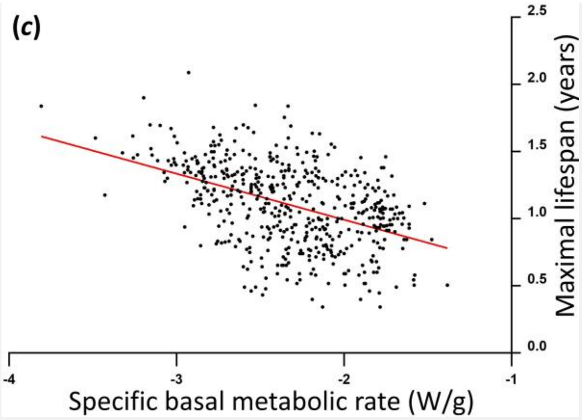

Figure S3 – $Log_{10}$-$log_{10}$ plots showing that, across **517** **MAMMAL AND BIRD** species:

**(*a*)** maximal lifespan (years) is proportional to adult body mass (kg),

**(*b*)** specific basal metabolic rate (W/g) is inversely proportional to adult body mass (kg),

**(*c*)** maximal lifespan (years) is inversely proportional to specific basal metabolic rate (W/g).

A variable can be standardized to be between 0 and 100 by the formula:

$$z_i = \left(\frac{x_i - \min(x)}{\max(x) - \min(x)}\right) \cdot 100 \qquad \text{[Eq. S1]}$$

Where $z_i$ is the $i^{th}$ standardized value in the variable, $x_i$ is the $i^{th}$ value in the variable, *min(x)* is the minimum value in the variable, and *max(x)* is the maximum value in the variable.

For 517 mammal and bird species: each of adult body mass (kg), specific basal metabolic rate (W/g), and maximal lifespan (years) were standardized in this way. Then those species with a value of 0 for one or more of these variables were removed (four species accordingly removed: *Trichechus manatus*, *Blarina brevicauda*, *Crocidura flavescens*, *Suncus etruscus*, leaving 513 species remaining), because each of these standardized variables was subsequently logged (base 10) and $log_{10}$ of zero (0) is undefined. These standardized and thereafter logged variables were then plotted: Figure S4 is a $log_{10}$-$log_{10}$ plot showing, across 513 mammal and bird species, that specific basal metabolic rate negatively, and maximal lifespan positively, scales with adult body mass. Wherein this data is herein interpreted to mean, at least across these species, adult body mass dictates specific basal metabolic rate, which in turn (at least in part) dictates maximal lifespan. Figure 13, in the main text, shows a subset of this data, showing only 259 mammals, with the aforementioned four mammals removed, with bats, naked mole rat, marsupials, and birds also absent.

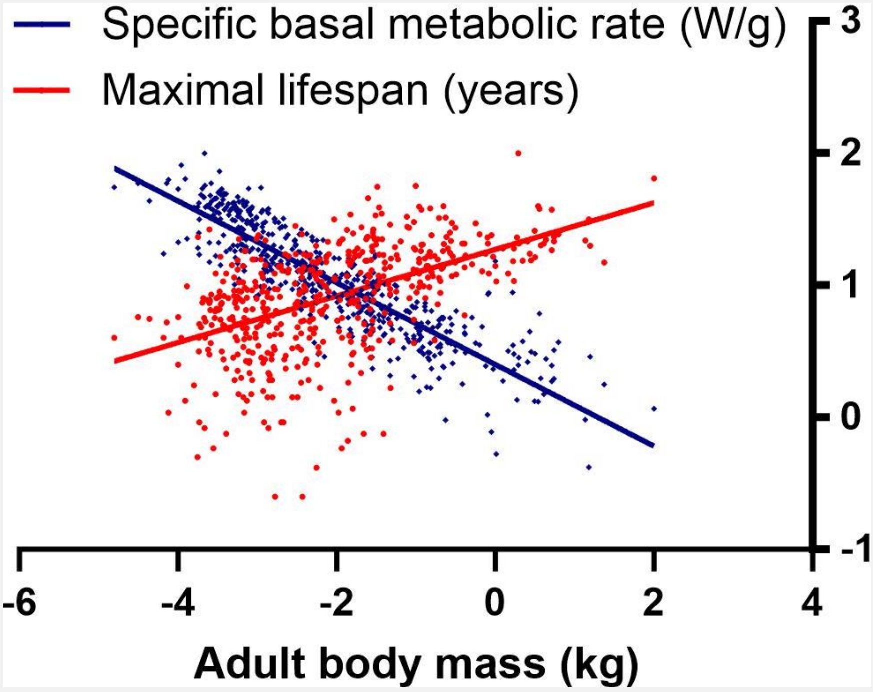

Figure S4 – $Log_{10}$-$log_{10}$ plot showing, across 513 mammal and bird species, that specific basal metabolic rate negatively, and maximal lifespan positively, scales with adult body mass. Data is standardized and logged as explained in corresponding text.

Across mammal species, maximal lifespan is inversely proportional to body surface-area per unit body mass *(shown using different data than earlier section with same title)*

This section is presented with the caveat that its values of adult body surface-area are estimated by calculation from adult body mass values sourced from [1] (*for those non-flying mammal species that have specific basal metabolic rate and maximal lifespan data therein*) using the Meeh equation ($S = k \cdot M^b$, where $S$ is body surface-area, $M$ is body mass) with generalized parameter values for non-flying mammals used, $k$ = 0.1 $m^2/kg^{0.67}$ and $b$ = 0.67 [128], rather than using experimentally determined values for each species. Only non-flying mammals considered (i.e., bats excluded: **MAMMALS without bats**, rendering 318 mammal species studied) as the Meeh parameters used are probably even less generalizable to flying mammals [128], which - because they have wings - tend to have a greater surface-area for their mass (all be it they fold their wings at rest). For the analysis of this section, units are maximal lifespan (years), adult body mass (kg), body surface-area per unit body mass ($m^2$/kg), and specific basal metabolic rate (W/g).

Using this data set it is observed that, across non-flying mammal species, (***a***) body surface-area is a supra-linear (positive concave) function of body mass, (***b***) body surface-area per unit body mass is a supra-linear (negative convex) function of body mass, (***c***) specific basal metabolic rate is a positive linear function of body surface-area per unit body mass, (***d***) maximal lifespan is a supra-linear (negative convex) function of body surface-area per unit body mass. It is herein proposed that (*a*) drives (*b*) drives (*c*) drives (*d*).

(***a***) Supra-linear (positive concave) shaped plot.

Pearson correlation coefficient = 0.9492.

Spearman's rank correlation coefficient = 1.

*Equation* $y = a \cdot x^b$ *best-fitted with a = 0.1, b = 0.67, wherein adjusted* $R^2 = 1$*. When x and y are logged (*$log_{10}$*), equation* $log_{10}(y) = m \cdot log_{10}(x) + log_{10}(c)$*, best-fitted with m = 0.67,* $log_{10}(c) = -1$*, adjusted* $R^2 = 1$*.*

(***b***) Supra-linear (negative convex) shaped plot.

Pearson correlation coefficient = -0.1671.

Spearman's rank correlation coefficient = -1.

*Equation* $y = a \cdot x^b$ *best-fitted with a = 0.1, b = -0.33, wherein adjusted* $R^2 = 1$*. When x and y are logged (*$log_{10}$*), equation* $log_{10}(y) = m \cdot log_{10}(x) + log_{10}(c)$ *best-fitted with m = -0.33,* $log_{10}(c) = -1$*, adjusted* $R^2 = 1$*.*

(***c***) Positive linear shaped plot.

Pearson correlation coefficient = 0.8069.

Spearman's rank correlation coefficient = 0.8808, one-tailed (alternative hypothesis is directional) p-value = 6.3692E-105.

*Equation* $y = m \cdot x + c$ *best-fitted with m = 0.02890, c = -0.0001065, adjusted* $R^2 = 0.6500$*.*

(***d***) Supra-linear (negative convex) shaped plot.

Pearson correlation coefficient = -0.5599.

Spearman's rank correlation coefficient = -0.77, one-tailed (alternative hypothesis is directional) p-value = 6.24113E-64.

*Equation* $y = a \cdot x^b$ *best-fitted with a = 3.973, b = -0.5498, wherein adjusted* $R^2 = 0.4811$*. When x and y are logged (*$log_{10}$*), equation* $log_{10}(y) = m \cdot log_{10}(x) + log_{10}(c)$ *best-fitted with m = -0.6344,* $log_{10}(c) = 0.4520$*, adjusted* $R^2 = 0.5765$*.*

Collinearity observed in multiple linear regression

In JASP software (version 0.14.1), multiple linear regression was performed for different, specified below, species sets. Wherein the response variable was maximal lifespan (years), whilst adult body mass (kg) and specific basal metabolic rate (W/g) were explanatory/predictor variables. All variables logged ($log_{10}$). With every species set, appreciable $R^2$ was observed. But high collinearity also, indicating redundancy/entwinning of explanatory variables. Which is consistent with, at least across these species, adult body mass dictating maximal lifespan because it determines specific basal metabolic rate. Where larger homeotherm species have lower specific basal metabolic rate as they need to generate less metabolic heat per unit mass per unit time to maintain their body temperature. Because (of the square-cube law of geometry) they have a lower body surface-area per unit body mass, and so lose their metabolically generated heat less readily. Whilst (to interpret) a metabolic rate (by-)product(s) drives aging.

So, across these species, adult body mass and specific basal metabolic rate are entwinned because larger adult body mass (because of the square-cube law) means a lower specific basal metabolic rate is required to maintain body temperature. Thence, because higher adult body mass inherently means lower specific basal metabolic rate, they aren't independent parameters. Which undermines how the slope regression coefficients below can be interpreted. Because the usual interpretation considers the change in the response variable for each additional unit increase in the predictor when the other predictor(s) is held constant. However, if predictors are entwinned, changing one predictor inherently changes the value(s) of the other(s).

In multiple linear regression, if the explanatory variables are not collinear then the Variance Inflation Factor (VIF) is one, and if they are collinear then VIF is greater than one, how much more depending upon the degree of collinearity, and a common rule of thumb is that if VIF is greater than four then this is of note. You can see below that in each case VIF exceeds four or is close to doing so.

Collinearity in a regression model can reduce the precision of the estimated coefficients: increasing their standard errors, which can decrease their statistical significance, possibly making important coefficients not statistically significant, and this increased error can even make a regression coefficient be of improper sign: negative when it should be positive, or vice-versa. Autocorrelation (explained later) can make a regression coefficient's standard errors smaller than they should be, which can confer unwarranted statistical significance, and it can inflate the F statistic, which can confer it unwarranted statistical significance. No collinearity and no autocorrelation are assumptions of multiple linear regression. All the multiple linear regressions below have collinearity. At least some have autocorrelation also. How autocorrelation and collinearity interact in a multiple linear regression model hasn't been formally studied that I know of. It might be that their combination is even more detrimental to the model than the sum of each alone: i.e., their detriments synergize.

To repeat, an assumption of multiple linear regression is that there is no collinearity. In the following multiple linear regression models, collinearity occurs, breaching this assumption, and corrupting them. Herein, the reason for including these models is to report this collinearity.

**[1]**

Below is a multiple regression model using data from all the **350 mammal species** with adult body mass, maximal lifespan, and specific basal metabolic rate data available in [1]:

Using JASP's default method, called "Enter", in which explanatory variables are forced into the regression model in a user-specified order, with adult body mass forced in first. Results are the same if specific basal metabolic rate is forced in first.

$R^2$ = 0.471, adjusted $R^2$ = 0.468, Unstandardized coefficients: $\beta_0$ (intercept) = 0.477 ± 0.172 (p-value = 0.006), $\beta_1$ = 0.114 ± 0.020 (regarding adult body mass. p-value = 5.091e-8. VIF = **3.844**), $\beta_2$ = -0.252 ± 0.068 (regarding specific basal metabolic rate. p-value = 2.374e-4. VIF = **3.844**).

*Standardized coefficients: $\beta_1$ = 0.426, $\beta_2$ = -0.284.*

*Durbin-Watson statistic, which has a possible range of 0-4, checks for correlations (at lag 1) between residuals (i.e., first order autocorrelation: serial correlation: when residual[i+1] is not independent of residual[i]), which can corrupt the test (linear, and multiple linear, regression requires independence of residuals. That the residuals are uncorrelated with each other. Assumption of independence of errors in the regression model). If below 2 (critically below 1), that indicates positive autocorrelation (next residual likely to be the same sign of that preceding it). If above 2 (critically above 3), that indicates negative autocorrelation (next residual likely to be the opposite sign of that preceding it). Between 1 and 3 is desirable, ideally around 2. Here, the Durbin-Watson statistic (is below 1!) = **0.837**. The source of the autocorrelation is the maximal lifespan variable because it is observed with linear regressions of maximal lifespan with adult body mass or* *specific basal metabolic rate, but not with linear regression of adult body mass and specific basal metabolic rate.*

*The higher the F statistic, the better the model. F statistic is significant: F (2, 347) = 154.565, p = 9.985e-49. Residuals vs. predicted plot shows a balanced random distribution of the residuals around the baseline suggesting that the assumption of homoscedasticity has not been violated. Quantile-Quantile (Q-Q) plot shows that the standardized residuals fit nicely along the diagonal suggesting that both assumptions of normality and linearity have not been violated.*

[Eq. S3]

$$log_{10}(maximal\ lifespan) = 0.477 + [0.114 \cdot log_{10}(adult\ body\ mass)] + [-0.252 \cdot log_{10}(specific\ basal\ metabolic\ rate)]$$

The same result is returned when JASP's "Backward" method is used. And it is the second result listed when its "Forward" or "Stepwise" method is used, while the first result only has adult body mass as a predictor, for which adjusted $R^2 = 0.449$.

When a linear regression is performed, with the response variable being maximal lifespan (years) and the sole predictor variable being specific basal metabolic rate (W/g), both logged ($log_{10}$), then adjusted $R^2 = 0.422$ (i.e., only 0.027 less than when adult body mass is the sole predictor). This linear regression result:

$R^2 = 0.424$, adjusted $R^2 = 0.422$, Unstandardized coefficients: $\beta_0$ (intercept) = -0.356 ± 0.09 (p-value = 8.873e-5), $\beta_1$ = -0.576 ± 0.036 (regarding specific basal metabolic rate. p-value = 1.412e-43). *Standardized coefficient: $\beta_1$ = -0.651. Durbin-Watson statistic = **0.865**. F (1, 348) = 256.001, p = 1.412e-43. Assumptions of homoscedasticity, normality, and linearity unviolated.*

[Eq. S4]

$$log_{10}(maximal\ lifespan) = -0.356 + [-0.576 \cdot log_{10}(specific\ basal\ metabolic\ rate)]$$

**[2]**

When the multiple linear regression above is repeated with basal metabolic rate (sourced from [1]) instead of specific basal metabolic rate (i.e., not per unit mass), the collinearity is even higher (VIF = **19.599**). Larger mammal species, with more cells, have a higher basal metabolic rate (all be it, as aforementioned, their basal metabolic rate per unit mass - i.e., their specific basal metabolic rate - is lower). Here the collinearity is so severe that the standard error in the regression coefficient for basal metabolic rate is so great that this coefficient is erroneously negative (highlighted by red coloured text below). Erroneous because maximal lifespan positively correlates (e.g., in a Pearson correlation) with basal metabolic rate. There is autocorrelation also: Durbin-Watson statistic = **0.888**.

Using JASP's method called "Enter" (results are the same whichever response variable is forced into the model first):

$R^2$ = 0.467, adjusted $R^2$ = 0.464, Unstandardized coefficients: $\beta_0$ (intercept) = 1.203 ± 0.030 (p-value = 8.134e-134), $\beta_1$ = 0.328 ± 0.046 (regarding adult body mass. p-value = 8.532e-12. VIF = **19.599**), $\beta_2$ = -0.214 ± 0.065 (regarding basal metabolic rate. p-value = 0.001. VIF = **19.599**). *Standardized coefficients: $\beta_1$ = 1.227, $\beta_2$ = -0.571. Assumptions of homoscedasticity, normality, and linearity unviolated. Durbin-Watson statistic = **0.888**. F (2, 347) = 151.863, p = 4.194e-48. The source of the autocorrelation is the maximal lifespan variable because it is observed with linear regressions of maximal lifespan with adult body mass or basal metabolic rate, but not with linear regression of adult body mass and basal metabolic rate.*

The same result is returned when JASP's "Backward" method is used instead. And it is the second result listed when its "Forward" or "Stepwise" method is used, the first being with adult body mass as sole predictor, for which adjusted $R^2$ = 0.449.

When linear regression is performed, with the response variable being maximal lifespan (years) and the predictor being basal metabolic rate (W), both logged ($log_{10}$), then adjusted $R^2$ = 0.388, with results as below (note how the slope coefficient is now correctly positive. Instead of erroneously negative as it was in the multiple linear regression above, caused by collinearity):

$R^2$ = 0.390, adjusted $R^2$ = 0.388, Unstandardized coefficients: $\beta_0$ (intercept) = 1.014 ± 0.014 (p-value = 4.857e-214), $\beta_1$ = 0.234 ± 0.016 (regarding basal metabolic rate. p-value = 3.096e-39). *Standardized coefficient: $\beta_1$ = 0.624. Durbin-Watson statistic = **0.793**. F (1, 348) = 222.416, p = 3.096e-39. Assumptions of homoscedasticity, normality, and linearity unviolated.*

[Eq. S5]

$$log_{10}(maximal\ lifespan) = 1.014 + [0.234 \cdot log_{10}(basal\ metabolic\ rate)]$$

**[3]**

Table S14 below reports multiple linear regression results, wherein maximal lifespan (years) is the response variable and adult body mass (kg) and specific basal metabolic rate (W/g) are explanatory variables, all logged ($log_{10}$). For different species sets, where *A* is 517 mammal and bird species, *B* is 350 mammal species (as used above), and *C* is 263 mammal species (excluding bats, naked mole rat, and marsupials). *Adj.* $R^2$ is adjusted $R^2$, *VIF* is the Variance Inflation Factor for each of the slope regression coefficients, *DW* is the Durbin-Watson statistic, *p_F* is the p-value for the F-statistic, $S\beta_1$ and $S\beta_2$ are the standardized regression coefficients for adult body mass and specific basal metabolic rate respectively, while $p\_\beta_0$, $p\_\beta_1$, and $p\_\beta_2$ are p-values for unstandardized regression coefficients for intercept, adult body mass and specific basal metabolic rate respectively. JASP's "Enter" method used, with adult body mass forced in first. For each, assumptions of homoscedasticity, normality, and linearity weren't violated.

[Table S14]

| Set | Adj. $R^2$ | VIF | DW | p_F | $S\_\beta_1$ | $S\_\beta_2$ | $p\_\beta_0$ | $p\_\beta_1$ | $p\_\beta_2$ |
|---|---|---|---|---|---|---|---|---|---|
| A | 0.328 | 4.429 | 0.764 | 1.524e-45 | 0.734 | 0.188 | 1.369e-24 | 2.064e-20 | 0.013 |
| B | 0.468 | 3.844 | 0.837 | 9.985e-49 | 0.426 | -0.284 | 0.006 | 5.091e-8 | 2.374e-4 |
| C | 0.675 | 3.796 | 1.170 | 1.068e-64 | 0.481 | -0.373 | 0.046 | 1.984e-11 | 1.262e-7 |

In Table S14 above, for species set *A*, because *VIF* is high at a value of 4.429 (collinearity), then the standard error in the regression coefficient for specific basal metabolic rate is so great that this coefficient is erroneously positive. Erroneous because maximal lifespan negatively correlates (e.g., in a Pearson correlation) with specific basal metabolic rate. Highlighted by use of red coloured text. So, collinearity is at least corrupting the results for species set *A*, and probably the (to some degree) results for the other species sets also. In each case, *DW* is, or is close to being, below 1. Indicating positive autocorrelation, which violates an assumption of regression. In each case, $p\_\beta_2 < 0.05$.

Table S15 below reports linear regression results, from using JASP's "Enter" method, where maximal lifespan (years) is the response variable and specific basal metabolic rate (W/g) is the explanatory variable, both logged (base 10). For species sets as specified above, wherein $S\beta_1$ is the standardized regression coefficient for specific basal metabolic rate, and $p\_\beta_1$ relates to specific basal metabolic rate. For each case, assumptions of homoscedasticity, normality, and linearity unviolated.

[Table S15]

| Set | Adj. $R^2$ | DW | p_F | $S\_\beta_1$ | $p\_\beta_0$ | $p\_\beta_1$ |
|---|---|---|---|---|---|---|
| A | 0.208 | 0.674 | 4.448e-28 | -0.457 | 1.251e-5 | 4.448e-28 |

| | | | | | | |
|---|---|---|---|---|---|---|
| B | 0.422 | 0.865 | 1.412e-43 | -0.651 | 8.873e-5 | 1.412e-43 |
| C | 0.616 | 1.240 | 2.538e-56 | -0.785 | 4.406e-12 | 2.538e-56 |

In Table S15 above, its *Adj.* $R^2$ values for linear regression, with specific basal metabolic rate as the sole explanatory variable, are not so different than those in earlier Table S14, which reported multiple linear regression results with both specific basal metabolic rate and adult body mass as explanatory variables. In each case, *DW* is, or is close to being, below 1. Indicating positive autocorrelation, which violates an assumption of regression. In each case here, specific basal metabolic rate is reported as a significant predictor of maximal lifespan ($p\_\beta_2 < 0.05$).

**[4]**

Table S16 below reports results from multiple linear regressions, where maximal lifespan (years) is the response variable and adult body mass (kg) and basal metabolic rate (W) are explanatory variables, all logged ($\log_{10}$). So, using basal metabolic rate instead of specific basal metabolic rate. Additional species groups included to try to match groups used in [6], where set *D* is the 350 mammal species set of set *A* but minus Artiodactyla, Macropodidae, Lagomorpha, and Soricidae, which renders 310 mammal species remaining, and set *E* is 167 bird species.

[Table S16]

| Set | Adj. $R^2$ | VIF | DW | p_F | $S\_\beta_1$ | $S\_\beta_2$ | $p\_\beta_0$ | $p\_\beta_1$ | $p\_\beta_2$ |
|---|---|---|---|---|---|---|---|---|---|
| A | 0.328 | 16.39 | 0.748 | 1.440e-45 | 0.213 | 0.366 | 2.376e-157 | 0.144 | 0.013 |
| B | 0.464 | 19.60 | 0.888 | 4.194e-48 | 1.227 | -0.571 | 8.134e-134 | 8.532e-12 | 0.001 |
| C | 0.654 | 18.39 | 1.187 | 5.018e-61 | 1.311 | -0.524 | 1.159e-118 | 2.8e-15 | 8.887e-4 |
| D | 0.411 | 16.69 | 0.892 | 2.067e-36 | 1.021 | -0.397 | 1.512e-177 | 2.490e-8 | 0.027 |
| E | 0.508 | 23.77 | 1.414 | 1.986e-26 | 0.693 | 0.025 | 7.127e-46 | 0.010 | 0.926 |

In Table S16 above, in each case *VIF* > 16. For species set *E*, *VIF* = 24. Note that for species sets *B*, *C*, and *D*: because *VIF* is so high (collinearity), the standard error in the regression coefficient for basal metabolic rate is so great that, given where it numerically is, this coefficient is erroneously negative. Erroneous because maximal lifespan positively correlates (e.g., in a Pearson correlation) with basal metabolic rate. Highlighted in the table by use of red text. In at least some of the cases, *DW* is, or is close to being, below 1. Indicating positive autocorrelation, which violates an assumption of regression.

So, we've seen above that collinearity can undermine multiple linear regression results. Because collinearity breaches a well-known assumption of multiple linear regression. That assumption being that there is no collinearity between predictors.

Partial and semi-partial correlations

Using Pearson: zero-order, partial, and semi-partial (part) correlations were performed in JASP software (version 0.18.1.0) for different species sets. Set *A* is 517 mammal and bird species, set *B* is 350 mammal species, and set *C* is 263 mammal species (excluding bats, naked mole rat, and marsupials). Results are reported in Table S17 below using the following capital letter symbols: maximal lifespan (*Y*), specific basal metabolic rate (*X*), and adult body mass (*Z*). All these variables being logged (base 10). Data was sourced from [1]. P-values are one-tailed because each alternative hypothesis is directional.

[Table S17]

Pearson correlation coefficients:

| Set | **Y.X** | Partial Y.X(Z) | Part Y.X(Z) | **Y.Z** | Partial Y.Z(X) | Part Y.Z(X) | **Z.X** |
|---|---|---|---|---|---|---|---|
| A | -0.457 | 0.110 | 0.090 | 0.568 | 0.392 | 0.349 | -0.880 |
| B | -0.651 | -0.196 | -0.145 | 0.671 | 0.287 | 0.217 | -0.860 |
| C | -0.785 | -0.319 | -0.191 | 0.801 | 0.399 | 0.247 | -0.858 |

Corresponding one-tailed p-values:

| Set | **Y.X** | Partial Y.X(Z) | Part Y.X(Z) | **Y.Z** | Partial Y.Z(X) | Part Y.Z(X) | **Z.X** |
|---|---|---|---|---|---|---|---|
| A | 2.41341E-28 | 0.9938372 | 0.9796015 | 8.42712E-46 | 9.74134E-21 | 1.48884E-16 | 6.1071E-169 |
| B | 7.3623E-115 | 0.0001121 75 | 0.00329 | 1.95838E-47 | 2.30888E-08 | 0.000021 | 5.2013E-104 |
| C | 1.64773E-56 | 6.1921E-08 | 0.000931 | 2.14531E-60 | 9.01273E-12 | 0.0000255 | 8.76144E-78 |

A zero-order correlation doesn't control for any other variable(s). A partial correlation is the correlation between a dependent variable and an independent variable after controlling for the influence of one or more other variables on *both* the dependent and independent variable. Partial correlation is that of residuals: e.g., conducting linear regression for $X$ vs. $Z$ and $Y$ vs. $Z$, and then finding the (e.g., Pearson) correlation between these two sets of residuals. Semi-partial correlation, also called part correlation, is the correlation between a dependent variable and an independent variable after controlling for the influence of one or more other variables on the dependent *or* the independent variable (herein the independent variable). A semi-partial correlation is that of a residual with a non-residual: e.g., conducting linear regression for $X$ vs. $Z$ and then calculating the (e.g., Pearson) correlation between $Y$ and these residuals (e.g., linear regression of specific basal metabolic rate as response variable and adult body mass as explanatory/predictor variable, collecting residuals, then finding the correlation of maximal lifespan to these residuals).

Partial correlation, which holds the other variable(s) constant for both $X$ and $Y$, and semi-partial correlation, which holds the other variable(s) constant for either $X$ or $Y$ (not both), can be adversely affected by collinearity. Because if variables are entwinned then changing one inherently changes the value(s) of another(s), and so it is a breached assumption to think that the other variable(s) can be held constant.

The equation for semi-partial (part) correlation is very similar to that for an unstandardized regression coefficient in multiple linear regression. The difference being that there is a square root in the denominator of the former, which curtails its value to be between -1 and +1. And so, just as collinearity can (by increasing standard error) make an unstandardized regression coefficient be of erroneous value, even of erroneous sign, collinearity can do the same for semi-partial correlation. Semi-partial correlation is the square root of the change in $R^2$ when the predictor is added to the multiple linear regression model. With collinearity there is high redundancy of predictors and so the change in $R^2$ is small and thence the semi-partial correlation is small. When there are only two predictors, the p-value for this change in $R^2$ is the same as the p-value for the corresponding partial correlation, which is the same as the p-value for the corresponding unstandardized regression coefficient, which can be made larger by collinearity, by it increasing the standard error of this unstandardized regression coefficient. So, to repeat, just as collinearity can corrupt multiple linear regression, it can corrupt partial and semi-partial correlations also.

We've already observed collinearity in presented multiple linear regressions, and we observe it here again in corresponding partial and semi-partial (part) correlations. In tables of this section, where a correlation is of the wrong sign it is shown in red text. In each such case there is also wrong sign for the corresponding unstandardized regression coefficient, in the corresponding multiple linear regression, caused by collinearity (e.g., compare Tables S17 and S14).

This collinearity, although corrupting these results, further shows that, at least across these species, adult body mass and specific basal metabolic rate are entwinned variables. Indeed, see in the results of Table S17 above how the partial and semi-partial correlations are lower than the zero-order correlations, not just for specific basal metabolic rate, but for adult body mass also. For species set *B*, the (absolute) difference between the zero-order and part correlations are 0.506 and 0.454 for specific basal metabolic rate and adult body mass respectively. Similar values. For species set *C*, the (absolute) difference between the zero-order and part correlations are 0.594 and 0.554 for specific basal metabolic rate and adult body mass respectively. Similar values again.

Despite the collinearity, all the results with correct sign in Table S17 are significant (at $p < 0.05$). Wherein, across species, maximal lifespan significantly (inversely) correlates with specific basal metabolic rate after the effect of adult body mass is statistically excluded.

Table S17 above used specific basal metabolic rate. By contrast, [6] used basal metabolic rate in their analysis (i.e., not per unit mass). For better comparison with [6], Table S18 below shows results when basal metabolic rate is used instead of specific basal metabolic rate. Maximal lifespan (*Y*), basal metabolic rate (*V*), and adult body mass (*Z*). All logged (base 10). Extra species sets *D* and *E* are as used earlier to generate Table S16. Data from [1]. P-values are one-tailed because each alternative hypothesis is directional.

[Table S18]

Pearson correlation coefficients:

| Set | **Y.V** | Partial Y.V(Z) | Part Y.V(Z) | **Y.Z** | Partial Y.Z(V) | Part Y.Z(V) | **Z.V** |
|---|---|---|---|---|---|---|---|

| | | | | | | | |
|---|---|---|---|---|---|---|---|
| A | 0.573 | 0.110 | 0.090 | 0.568 | 0.064 | 0.053 | 0.969 |
| B | 0.624 | -0.174 | -0.129 | 0.671 | 0.355 | 0.277 | 0.974 |
| C | 0.750 | -0.204 | -0.122 | 0.801 | 0.462 | 0.306 | 0.972 |
| D | 0.593 | -0.126 | -0.097 | 0.636 | 0.310 | 0.250 | 0.970 |
| E | 0.703 | 0.007 | 0.005 | 0.717 | 0.200 | 0.142 | 0.979 |

Corresponding one-tailed p-values:

| Set | **Y.V** | Partial Y.V(Z) | Part Y.V(Z) | **Y.Z** | Partial Y.Z(V) | Part Y.Z(V) | **Z.V** |
|---|---|---|---|---|---|---|---|
| A | 9.54735E-47 | 0.006163 | 0.0203985 | 8.42712E-46 | 0.0730875 | 0.1144835 | 0 |
| B | 1.80211E-39 | 0.9994595 | 0.9921288 | 1.95838E-47 | 3.90606E-12 | 6.92369E-08 | 8.6337E-227 |
| C | 4.59313E-49 | 0.9995621 | 0.9759493 | 2.14531E-60 | 1.29985E-15 | 2.08451E-07 | 1.7555E-166 |
| D | 3.97046E-31 | 0.9867343 | 0.9559014 | 7.64263E-37 | 1.24749E-08 | 0.000004 | 4.3667E-215 |
| E | 1.66119E-26 | 0.4642305 | 0.4744335 | 5.96633E-28 | 0.0047795 | 0.033584 | 3.4693E-116 |

In Table S18 above, collinearity corrupts at least the results shown in red text: conferring them the wrong sign. The species sets with red text in Table S18 are the same as those with red text in earlier Table S16.

Note in Table S18 above how greatly the correlation between maximal lifespan and adult body mass is diminished when controlling for basal metabolic rate, which is an aspect that [6] didn't report upon. Indeed, for species set *A*, which has the most species (=517), upon correcting for basal metabolic rate, maximal lifespan does ***not*** significantly correlate with adult body mass (!). By contrast, for species set *A*, upon correcting for adult body mass, maximal lifespan ***does*** significantly correlate with basal metabolic rate. Because, for species set *A*, the partial and part correlations for adult body mass are both lower than those for basal metabolic rate. Look in the last column, to see how highly basal metabolic rate correlates with adult body mass, when both variables are logged. To be expected because more body mass requires more oxygen per unit time (i.e., a higher metabolic rate). All be it, as I've said repeatedly herein, more homeothermic body mass permits a lower metabolic *per unit mass*: i.e., a lower specific basal metabolic rate.

From results corrupted by collinearity herein, I infer that the results of [6] are corrupted by collinearity too. Similarly, collinearity causing error, to the extent of incorrect sign(s), is probably why [318] (*erroneously*) reports that, across species, maximal lifespan negatively correlates with basal metabolic rate residuals. [6] and [318] didn't notice or appreciate this collinearity and therefore make erroneous conclusions.

So, we saw in the last section that collinearity can corrupt multiple linear regression results (because it breaches an assumption of the method), and we've seen in this section that collinearity can corrupt partial and semi-partial (part) correlations. Perhaps unsurprising given their interconnection with multiple linear regression, and that collinearity breaches an assumption of multiple linear regression.

<u>Hierarchical regression</u>

Using JASP software (version 0.18.1.0) for different species sets, where set *A* is 517 mammal and bird species, set *B* is 350 mammal species, and set *C* is 263 mammal species (excluding bats, naked mole rat, and marsupials), I performed hierarchical regression. Using data from [1], with maximal lifespan (years) as the response variable, and adult body mass (kg) and specific basal metabolic rate (W/g) as explanatory variables, all logged (base 10). In Table S19 below, when adult body mass is in the null model, the first column shows the change in $R^2$ when specific basal metabolic rate is added to the model. When specific basal metabolic rate is in the null model instead, the second column shows the change in $R^2$ when adult body mass is added to the model. P-values are shown italicized in brackets.

[Table S19]

| Set | Specific basal metabolic rate | Adult body mass |
|---|---|---|
| A | 0.008 (*0.013*) | 0.122 (*2.064e-20*) |
| B | 0.021 (*2.374e-4*) | 0.047 (*5.091e-8*) |
| C | 0.037 (*1.262e-7*) | 0.061 (*1.984e-11*) |

Table S20 below is as Table S19 above except that $\log_{10}$ basal metabolic rate (W) is used instead of $\log_{10}$ specific basal metabolic rate.

[Table S20]

| Set | Basal metabolic rate | Adult body mass |
|---|---|---|
| A | 0.008 (*0.013*) | 0.003 (*0.144*) |
| B | 0.017 (*0.001*) | 0.077 (*8.532e-12*) |
| C | 0.015 (*8.887e-4*) | 0.093 (*2.8e-15*) |

All the $R^2$ changes in Tables S19-S20 above are significant except, with species set *A*, adding adult body mass doesn't significantly improve $R^2$ when basal metabolic rate is already in the (null) model. All the $R^2$ changes are relatively small because of (aforementioned) collinearity.

<u>Adjusting for phylogeny</u>

An assumption of at least some statistical tests is that the data points are independent, which is arguably inherently breached in interspecies studies because all the different species are evolutionarily connected. Some closer than others. This section uses the Phylogenetically Independent Contrasts (PIC) method of [319], as implemented by PDAP module version 1.16 [320] of Mesquite version 3.81 (build 955) software [321], to try to account for this. With the caveat that we can't be sure of the phylogeny of any species. Phylogenetic relationships between the species were sourced from the timetree.org online database [322]. Allocated branch lengths in unit of time. Any species reported to have "Unresolved name" by [322] when it outputted the phylogeny was excluded: i.e., the phylogeny was sought again with this species absent. When conducting the phylogenetically independent contrasts method, best practices as taught by [323], which are built into PDAP by default, were followed: e.g., the Pearson correlation between the two phylogenetically independent contrasts is forced through the origin.

Analysis was conducted for the 517 mammal and bird species set used earlier but with the following 16 mammal species excluded because of the aforementioned "unresolved name" issue: *Microgale dobsoni*, *Erethizon dorsatus*, *Potorous tridactylus*, *Ailurus fulgens*, *Miniopterus schreibersii*, *Tatera indica*, *Cryptomys hottentotus*, *Proechimys semispinosus*, *Loris tardigradus*, *Eulemur fulvus*, *Saimiri sciureus*, *Rattus rattus*, *Eptesicus fuscus*, *Cercopithecus mitis*, *Isoodon obesulus*, *Dasyprocta azarae*. And with the following 22 bird species also excluded because of the same "unresolved name" issue: *Anas clypeata*, *Chordeiles minor*, *Botaurus lentiginosus*, *Anhinga anhinga*, *Bulweria bulwerii*, *Scolopax* *rusticola*, *Scolopax minor*, *Strix occidentalis*, *Aegolius acadicus*, *Pernis apivorus*, *Eremophila alpestris*, *Regulus regulus*, *Muscicapa striata*, *Sylvia curruca*, *Geococcyx californianus*, *Haemorhous cassinii*, *Larus occidentalis*, *Larus delawarensis*, *Sialia mexicana*, *Micrathene whitneyi*, *Phoebastria immutabilis*, *Phalacrocorax auritus*. Leaving 479 mammal and bird species remaining, which is species set *A* of this section (and of subsequent phylogenetic related sections). Species set *B* is just the 334 mammal species thereof. According to the phylogeny used, species set A has 20 polytomies and species set *B* has 14 polytomies.

The Phylogenetic Independent Contrasts (PIC) method requires compared traits to be linearly related [324]. The relation between maximal lifespan and each of specific basal metabolic rate and adult body mass is non-linear, becoming linear upon logging the variables. So, logged ($log_{10}$) data is used.

Using data from [1], in Table S21 below are Pearson correlation coefficients (not forced through the origin) for species set *A*, before phylogeny is considered, where *M* is $log_{10}$ adult body mass, *L* is $log_{10}$ maximal lifespan, *R* is $log_{10}$ basal metabolic rate, and *sR* is $log_{10}$ specific basal metabolic rate. P-values, italicized in brackets, are one-tailed because in each case the alternative hypothesis is directional.

[Table S21]

| | M | L | R | sR |
|---|---|---|---|---|
| M | 1 | | | |
| L | 0.5736 (*1.48028E-43*) | 1 | | |
| R | 0.9695 (*9.6488E-294*) | 0.577 (*3.67511E-44*) | 1 | |
| sR | -0.8807 (*3.3011E-157*) | -0.4658 (*1.79733E-27*) | -0.7654 (*1.18587E-93*) | 1 |

Table S22 below is as Table S21 above, but for species set *B* instead.

[Table S22]

| | M | L | R | sR |
|---|---|---|---|---|
| M | 1 | | | |

| | | | | |
|---|---|---|---|---|
| L | 0.6822 (*2.27188E-47*) | 1 | | |
| R | 0.9744 (*1.4509E-217*) | 0.634 (*2.97588E-39*) | 1 | |
| sR | -0.8609 (*9.326E-100*) | -0.6669 (*1.24965E-44*) | -0.7606 (*1.48404E-64*) | 1 |

Species sets *A* and *B* are referred to in Table S23 below, with orthogonal column headings: *C* is the (forced through origin [323, 325]) Pearson correlation coefficient between phylogenetically independent contrasts of $\log_{10}$ maximal lifespan and $\log_{10}$ specific basal metabolic rate (*and in the line below, labelled "P", the corresponding one-tailed [alternative hypothesis is directional] p-value thereof*), and subsequent columns are that but with a branch length adjustment made, that of Nee (page 416 of [326]), Pagel [327], Grafen [328], or wherein all the branch lengths are set to 1.0 (which are all options available in Mesquite). The latter transform was used in [6]. So, this table discloses phylogenetically adjusted Pearson correlation coefficients. Outputted by the PDAP module [320] of Mesquite [321].

[Table S23]

| | *C* | Nee | Pagel | Grafen | =1.0 |
|---|---|---|---|---|---|
| **A** | -0.32592391812554 | -0.3968607413260524 | -0.335866250576947 | -0.29535102899588583 | -0.42290935419591724 |
| *P* | 1.284547840406727E-13 | 8.066353290764338E-20 | 2.141979955383371E-14 | 2.1281195185864E-11 | 1.661808065251739E-22 |
| **B** | -0.3025067315490833 | -0.3798184420054401 | -0.30857627943944993 | -0.25220529255726704 | -0.41169655380702574 |
| *P* | 8.776279134052015E-9 | 3.6258369147 | 4.219470104559913E-9 | 1.5494210749931325E-6 | 2.1388406523682044E-15 |

| | | |
|---|---|---|
| | | 77282E-13 |

Table S24 below is as above but reporting, vs. $\log_{10}$ maximal lifespan, instead of $\log_{10}$ specific basal metabolic rate, for upper values of *A* and *B*, $\log_{10}$ basal metabolic rate. And for lower values of *A* and *B*, $\log_{10}$ adult body mass. Where the p-value is 0.0, this reflects that the p-value was so small that Mesquite rounded it down to 0.0. So, this table discloses phylogenetically adjusted Pearson correlation coefficients. Outputted by the PDAP module [320] of Mesquite [321].

[Table S24]

| | *C* | Nee | Pagel | Grafen | =1.0 |
|---|---|---|---|---|---|
| **A** | 0.316651980818457 86 | 0.41301785416501113 | 0.358838373432645 | 0.32001014666136346 | 0.44017607725323343 |
| *P* | 6.827871601444713E-13 | 0.0 | 2.220446049250313E-16 | 7.229772336359019E-13 | 0.0 |
| **B** | 0.39194936237949 45 | 0.3990169393799362 | 0.3367860556624948 | 0.28219599851402843 | 0.4362046181543429 |
| *P* | 5.728750807065808E-14 | 1.6986412276764895E-14 | 1.3298750989321206E-10 | 7.815518776599362E-8 | 0.0 |
| **A** | 0.36840603725645 55 | 0.44582023172027696 | 0.38812581066114527 | 0.34718341117278145 | 0.4745291419037174 |
| *P* | 0.0 | 0.0 | 0.0 | 2.55351295663786E-15 | 0.0 |

| | | | | | |
|---|---|---|---|---|---|
| **B** | 0.40526810858347806 | 0.4390037842693533 | 0.37103898261780976 | 0.31376863392164983 | 0.4787071205538979 |
| *P* | 6.8833827526759706E-15 | 0.0 | 1.2150280781497713E-12 | 2.2933249743672945E-9 | 0.0 |

Tables S23 and S24 above report that, for species set *A*, even after phylogeny adjustment, there is a statistically significant (negative) relation between $\log_{10}$ maximal lifespan and $\log_{10}$ specific basal metabolic rate (*for species set A, mean relation across the different columns = -0.3553823. But according to the diagnostic table, Table S26, disclosed later herein, only column "=1.0" should be considered = -0.4229094*). By contrast, for species set *A*, there is not a statistically significant relation shown between $\log_{10}$ maximal lifespan and $\log_{10}$ basal metabolic rate (*for species set A, mean relation across the different columns = 0.3697389. But according to the diagnostic table, Table S26, disclosed later, none should be considered*). And, for species set *A*, there is *not* a statistically significant relation shown between $\log_{10}$ maximal lifespan and $\log_{10}$ adult body mass (*for species set A, mean relation across the different columns = 0.4048129. But according to the diagnostic table, Table S26, disclosed later, none should be considered*).

Tables S23 and S24 above report that, for species set *B*, after phylogeny adjustment, there is a statistically significant (negative) relation between $\log_{10}$ maximal lifespan and $\log_{10}$ specific basal metabolic rate (*for species set B, mean relation across the different columns = -0.33096065987. But according to the diagnostic table, Table S26, disclosed later herein, only columns "Nee" and "=1.0" should be considered, and their mean = -0.3957574979*). And, for species set *B*, after phylogeny adjustment, a statistically significant relation is shown between $\log_{10}$ maximal lifespan and $\log_{10}$ basal metabolic rate (*for species set B, mean relation across the different columns = 0.36923059481. But according to the diagnostic table, Table S26, disclosed later, only column "Nee" should be considered = 0.3990169393799362*). And, for species set *B*, after phylogeny adjustment, a statistically significant relation is shown between $\log_{10}$ maximal lifespan and $\log_{10}$ adult body mass (*for species set B, mean relation across the different columns = 0.40155732598. But according to* *the diagnostic table, Table S26, disclosed later, only columns "C" and "Nee" should be considered, and their mean = 0.42213594642*).

In Table S25 below, for upper values of *A* and *B*: *C* is the (forced through origin [323, 325]) Pearson correlation coefficient between phylogenetically independent contrasts of $\log_{10}$ specific basal metabolic rate and $\log_{10}$ adult body mass (*and in the line below, labelled "P", the corresponding one-tailed [alternative hypothesis is directional] p-value thereof*), and subsequent columns are that but with a branch length adjustment made (with column headings as specified earlier). For the lower values of *A* and *B*: *C* is the (forced through origin [323, 325]) Pearson correlation coefficient between phylogenetically independent contrasts of $\log_{10}$ basal metabolic rate (i.e., not per unit mass) and $\log_{10}$ adult body mass (*and in the line below, labelled "P", the corresponding one-tailed [alternative hypothesis is directional] p-value thereof*), and subsequent columns are that but with a branch length adjustment made (with column headings as specified earlier). So, this table discloses phylogenetically adjusted Pearson correlation coefficients. Outputted by the PDAP module [323] of Mesquite [325]. Where the p-value is 0.0, this means that p-value was so small that Mesquite rounded it down to 0.0.

[Table S25]

| | *C* | Nee | Pagel | Grafen | =1.0 |
|---|---|---|---|---|---|
| **A** | -0.6242999490995295 | -0.759508686378264 | -0.7325334080788778 | -0.7093373093088843 | -0.7599918769809937 |
| *P* | 6.002801724936258E-53 | 3.6697773518676605E-90 | 5.850373545194245E-81 | 6.894713813737395E-74 | 2.448647108880704E-90 |
| **B** | -0.5944897631851525 | -0.7300534896255015 | -0.7031474752531311 | -0.6747677378428316 | -0.7360128954857933 |

| | | | | | |
|---|---|---|---|---|---|
| *P* | 2.453956894220469E-33 | 2.01188547159 61853E-56 | 8.174584363442112E-51 | 1.5512924347415462E-45 | 9.336700892665224E-58 |
| **A** | 0.88580770044897 52 | 0.9307238896594348 | 0.9232084735620868 | 0.9141045038277481 | 0.9310394172261127 |
| *P* | 0.0 | 0.0 | 0.0 | 0.0 | 0.0 |
| **B** | 0.88063835934708 63 | 0.9246938695228354 | 0.9155196967018939 | 0.9034315877807204 | 0.9271643024858973 |
| *P* | 0.0 | 0.0 | 0.0 | 0.0 | 0.0 |

In Table S25 above, one can see that even after phylogeny adjustment, across (at least these) species, there is a statistically significant (negative) relation between $\log_{10}$ specific basal metabolic rate and $\log_{10}$ adult body mass (*for species set A, mean relation across the different columns = -0.7171342. But according to the diagnostic table, Table S26, disclosed later, only column "Nee" should be considered = -0.7595087*). And a statistically significant relation between $\log_{10}$ basal metabolic rate and $\log_{10}$ adult body mass (*for species set A, mean relation across the different columns = 0.9169768. But according to the diagnostic table, Table S26, disclosed later, only column "Nee" should be considered = 0.9246939*).

Table S26 below is the (aforementioned) diagnostic table and it shows for both species sets *A* and *B*: for *Q* ($\log_{10}$ specific basal metabolic rate), *R* ($\log_{10}$ maximal lifespan), *S* ($\log_{10}$ basal metabolic rate), and *T* ($\log_{10}$ adult body mass): in its column *C*, the Pearson correlation coefficient (*not* forced through origin [323, 325]) of the absolute values of the standardized phylogenetically independent contrasts versus their standard deviations (i.e., the square root of the sum of their branch lengths), which is a diagnostic check of how well the topology and branch lengths of the phylogenetic tree (and assumed model of character evolution) adequately fit the tip data [323, 325], wherein no significant correlation is the desirable result (*and in the line below, labelled "P", the corresponding two-tailed p-value thereof*), and subsequent columns are that but with a branch length adjustment made as codified earlier. P-values greater than 0.05, which are desirable, are in red text. Table values were outputted by the PDAP module [320] of Mesquite [321].

[Table S26]

| Diagnostic table | | | | | | |
|---|---|---|---|---|---|---|
| | | *C* | Nee | Pagel | Grafen | =1.0 |
| **A** | **Q** | -0.1267024662294454 | -0.01909728598583698 | -0.21327647499110688 | -0.2781950814067795 | 0.08111725844153485 |
| | *P* | 0.005536182816942154 | 0.6770640667465346 | 2.539400307090789E-6 | 6.058579309147415E-10 | 0.07643422907229436 |
| | **R** | -0.21335942923711038 | -0.13803096228981837 | -0.2931069501492597 | -0.31216688576355045 | -0.06520429979637114 |
| | *P* | 2.51618572110318E-6 | 0.00249136857914845 | 6.336968653598267E-11 | 2.9030258027890894E-12 | 0.15463241820841045 |
| | **S** | -0.09883099251757707 | -0.003609442657171656 | -0.1906217350408323 | -0.2643640551279754 | 0.09951383056439922 |
| | *P* | 0.030741949437256735 | 0.9372651610668471 | 2.726064939951163E-5 | 4.372255288866673E-9 | 0.02960032044010874 |

| | | | | | | |
|---|---|---|---|---|---|---|
| | **T** | -0.020125637680331782 | 0.04951012935393999 | -0.15548872844252465 | -0.24220682994962628 | 0.1661117543997805 |
| | *P* | 0.660731476396867 | 0.2800179933686304 | 6.464155731262312E-4 | 8.249204684774928E-8 | 2.647014655077218E-4 |
| **B** | **Q** | -0.19211358351038854 | -0.034283579823641504 | -0.24156981299056562 | -0.28895795138361324 | 0.07696902441928963 |
| | *P* | 4.2275435792095905E-4 | 0.5329905253218005 | 8.278405577956139E-6 | 7.95697065158357E-8 | 0.16110913174188912 |
| | **R** | -0.09198830175133184 | -0.028077254317234145 | -0.2192183682885315 | -0.2659689425827684 | 0.03692553173784982 |
| | *P* | 0.09376329866617927 | 0.6096760936402202 | 5.471905143267537E-5 | 8.470537665520165E-7 | 0.5018874050961424 |
| | **S** | -0.14675926798725517 | 0.015974676005231298 | -0.18111967338647608 | -0.25584561411218687 | 0.11671882173361779 |

| | | | | | | |
|---|---|---|---|---|---|---|
| | *P* | 0.0073057779829497444 | 0.7714851056281382 | 8.9953907604735E-4 | 2.243587255703068E-6 | 0.03323837061305901 |
| | **T** | -0.06283841364566 | 0.08396352049223724 | -0.13627348466319902 | -0.2324326479366262 | 0.1990091702845919 |
| | *P* | 0.25282607196645956 | 0.126232138205578 | 0.012809264480624222 | 1.832590981426702E-5 | 2.5764206661049727E-4 |

This phylogeny analysis has made assumptions, one or more of which may not hold, which may or may not undermine/diminish this analysis, such as: the phylogeny's topology and initial branch lengths (in unit of time) are correct, the average evolution rate is the same throughout the phylogeny used, and there is no measurement error in the characters/trait values. To highlight: the phylogenetic topology might be incorrect, at least we cannot be sure that it is correct, wherein the topology can have a major impact on the results. Use of no branch length transformation or one of Nee, Pagel, or Grafen branch length transformations assumes character evolution behaves as Brownian motion (*which models evolution as purely random genetic drift with no selection. No natural selection or sexual selection. Although it may capture selection caused by randomly fluctuating environmental conditions. It might be inconsistent with at least some evolutionary theory/theories, for example, with the "punctuated equilibrium" concept [329], although even purely gradual Brownian motion models of evolutionary change will sometimes yield periods of rapid, almost punctuational, change*). A gradual model of evolution. By contrast, use of all branch lengths set to 1.0 (or equality at some other value) confers a "speciational" model of evolution [330], where all character change occurs in association with speciation events. Distinct from punctuated equilibrium (in which change only occurs in one of the daughter lineages) in that change occurs in *both* the daughter lineages.

<u>Other phylogenetic analysis</u>

The phylogenetic analysis above was constrained in requiring both compared variables/characters/traits to have the same branch length transformation, or lack thereof. In this section, following the teaching of [323, 325], analysis is performed without this constraint. By this teaching, distinct from analysis above, different continuous variables can be standardized with different branch lengths, by using different branch length transformations, or lack thereof.

Using species set *A*: raw phylogenetically independent contrasts were outputted from the PDAP module of Mesquite, and each was then standardized by dividing each contrast by the (also PDAP outputted) contrast's standard deviation (square root of sum of corrected branch lengths). The raw contrasts, and corresponding standard deviations to standardize them: for $\log_{10}$ adult body mass were sourced when no branch length transformation was used, for $\log_{10}$ maximal lifespan were sourced when branch lengths were set as 1.0, for $\log_{10}$ specific basal metabolic rate were sourced when the Nee branch length transformation was used, and for $\log_{10}$ basal metabolic rate were sourced when the Nee branch length transformation was used. So, each variable has a tailored branch length transformation, or lack thereof, whichever is best in each case (out of those I trialled) according to the earlier presented diagnostic table, Table S26.

The method above was repeated for species set *B*, wherein the best branch length transformations according to diagnostic Table S26 were used for the different variables: untransformed branch lengths for $\log_{10}$ adult body mass, Nee transformed branch lengths for each of $\log_{10}$ maximal lifespan, $\log_{10}$ specific basal metabolic rate, and $\log_{10}$ basal metabolic rate.

Table S27 below reports phylogenetically adjusted Pearson correlation coefficients (i.e., with any phylogeny effect factored out), wherein (distinct from earlier phylogenetic analysis herein) tailored branch length transformations were used for the different variables. It relates to logged ($\log_{10}$) variables, *A* and *B* refer to species sets *A* and *B* respectively, and it reports Pearson correlation coefficients (forced through the origin [323, 325]) between standardized (as specified earlier) phylogenetically independent contrasts, with one-tailed (alternative hypothesis is directional) p-values thereof in brackets. For species set *A*, as specified earlier, using untransformed branch lengths for $\log_{10}$ adult body mass, Nee transformed branch lengths for $\log_{10}$ specific basal metabolic rate and $\log_{10}$ basal metabolic rate, and branch lengths set as 1.0 for maximal lifespan. For species set *B*, as specified earlier, using untransformed branch lengths for $\log_{10}$ adult body mass, and Nee transformed branch lengths for each of $\log_{10}$ maximal lifespan, $\log_{10}$ specific basal metabolic rate, and $\log_{10}$ basal metabolic rate. How I performed a Pearson correlation coefficient forced through the origin is by using the data with the [331] statistical package to do a linear regression forced through the origin (i.e., intercept = 0), wherein the alongside reported Pearson correlation coefficient is also forced through the origin (which isn't the case with at least some more prominent statistical packages, as warned in [325]).

[Table S27]

| *All variables are logged ($\log_{10}$)* | | | | |
|---|---|---|---|---|
| | | Adult body mass | Specific basal metabolic rate | Basal metabolic rate |
| Maximal lifespan | A | 0.412 (2.3733E-21) | -0.3858 (9.49935E-19) | 0.4143 (1.36105E-21) |
| | B | 0.3929 (4.47621E-14) | -0.3798 (3.32922E-13) | 0.399 (1.6987E-14) |

Table S27 above shows that with phylogeny factored out, across at least the species of species sets *A* and *B*, maximal lifespan correlates better with basal metabolic rate than it does with adult body mass. The table shows that with phylogeny factored out, across at least these species, maximal lifespan *significantly* correlates with each of adult body mass, basal metabolic rate, and (inversely) with specific basal metabolic rate.

It might be that greater statistical significance may be found by using different, untested, valid (by diagnostic appraisal) branch length transformations and combinations thereof, because an exhaustive search of the different options available in the Mesquite software used wasn't performed, let alone the many further conceivable transformations and combinations. Moreover, greater statistical significance might be found if different branch length transformations, or lack thereof, were used in different parts of the phylogeny, which is permissible [325], and possible in Mesquite, which can be desirable, for example, if the average rate of evolution varies among clades and/or subclades, but this wasn't considered.

Be aware that using different branch lengths for different variables *assumes* that these variables evolve at a different rate. And use of branch lengths set as 1.0 *assumes* a different model of evolution (speciational change) than the other branch lengths trialled (gradual change). These assumptions might be unreasonable. For example, because of the earlier shown collinearity/entwinement between adult body mass and specific basal metabolic rate (*and even more between adult body mass and basal metabolic rate*), it might be unreasonable to assume, as was implicitly done with the different branch lengths employed for these variables, that they evolve at different rates. Arguably, given their entwinement, they might have co-evolved at the same rate. Presumably, to keep body temperature the same, a larger body size inexorably requires less metabolic heat generation per unit mass per unit time (and so a lower specific basal metabolic rate), because otherwise (*because, by the square-cube law of geometry, larger objects have less surface-area per unit object mass*) the body temperature will be more.

Adjusting for phylogeny and body mass

The following analysis further follows the teaching of [325] to find partial, and extrapolates its teaching to find semi-partial, (forced through origin [323, 325]) Pearson correlation coefficients using standardized phylogenetically independent contrasts. Although [325] is silent, and so equivocal, upon whether the following analysis is permissible with standardized phylogenetically independent contrasts generated with different branch length transformations. In the present section I present results from analysis with standardized phylogenetically independent contrasts generated with *different* branch lengths. In the next headed section, I present results from analysis with standardized phylogenetically independent contrasts generated with *same* branch lengths. The results are very different. Perhaps to be expected because whereas a (forced through the origin) Pearson correlation coefficient between phylogenetically independent contrasts isn't changed much by the contrasts being generated with different branch lengths, the slope of their linear regression can be [323]. And thence, to extrapolate, the slopes of a multiple linear regression also, which presumably means partial and semi-partial correlation coefficients can be very different. Indeed, this is observed in the results presented herein. So, the results of this present section must not be interpreted on their face, but with the different branch lengths used factored into their interpretation.

A linear regression of standardized (as specified earlier) phylogenetically independent contrasts for $\log_{10}$ specific basal metabolic rate (generated with Nee transformed branch lengths; regressed as dependent, *y*, variable) on standardized phylogenetically independent contrasts for $\log_{10}$ adult body mass (generated with untransformed branch lengths; regressed as independent, *x*, variable), with intercept set as zero, was performed using [331], and its residuals saved. Then a linear regression of standardized phylogenetically independent contrasts for $\log_{10}$ maximal lifespan (generated with branch lengths set to 1.0; regressed as dependent, *y*, variable) on standardized phylogenetically independent contrasts for $\log_{10}$ adult body mass (generated with untransformed branch lengths; regressed as independent, *x*, variable), with intercept set as zero, was performed using [331], and its residuals saved. Lastly, a linear regression with these two sets of residuals, with intercept set as zero, was conducted using [331]. Its alongside reported Pearson correlation coefficient (R) {when intercept = zero} is that between $\log_{10}$ maximal lifespan and $\log_{10}$ specific basal metabolic rate excluding the effects of both phylogeny and $\log_{10}$ adult body mass: R = -0.1652, one-tailed (alternative hypothesis is directional) p-value = 0.000143. When branch lengths were (instead of using the Nee transformation) set as 1.0 for $\log_{10}$ specific basal metabolic rate, wherein this is also a valid branch length transformation according to the diagnostic table, Table S26, presented earlier (because of non-significant p-value): R = -0.2186 (one-tailed p-value = 6.80284E-07). So, across (at least these) species, with the effects of phylogeny and adult body mass excluded, maximal lifespan significantly (negatively) correlates with specific basal metabolic rate. Repeating this analysis but with species set *B* instead, using (as before for species set *B*) untransformed branch lengths for $\log_{10}$ adult body mass, and Nee transformed branch lengths for both $\log_{10}$ maximal lifespan and $\log_{10}$ specific basal metabolic rate: R = -0.1809, one-tailed (alternative hypothesis is directional) p-value = 0.0004485.

Returning to species set *A*, the method above was emulated to report the Pearson correlation coefficient (R) {when intercept = zero} between $\log_{10}$ maximal lifespan and $\log_{10}$ basal metabolic rate (its standardized contrasts generated with Nee branch length transformation used), excluding the effects of both phylogeny and $\log_{10}$ adult body mass: R = 0.1276, one-tailed (alternative hypothesis is directional) p-value = 0.0025805. So, across (at least these) species, with the effects of phylogeny and adult body mass excluded, maximal lifespan significantly correlates with basal metabolic rate. Repeating this analysis with species set *B* instead, using (as before for species set *B*) untransformed branch lengths for $\log_{10}$ adult body mass, and Nee transformed branch lengths for both $\log_{10}$ maximal lifespan and $\log_{10}$ basal metabolic rate: R = 0.1284, one-tailed (alternative hypothesis is directional) p-value = 0.0094495.

Returning to species set *A*, replicating the earlier method but instead finding the relation between $log_{10}$ maximal lifespan and $log_{10}$ adult body mass when excluding the effects of phylogeny and $log_{10}$ specific basal metabolic rate (using, as before, untransformed branch lengths, branch lengths = 1.0, and Nee transformed branch lengths, for $log_{10}$ adult body mass, $log_{10}$ maximal lifespan, and $log_{10}$ specific basal metabolic rate respectively): R = 0.2262, one-tailed (alternative hypothesis is directional) p-value = 2.82324E-07. When instead excluding the effects of phylogeny and (with Nee transformed branch lengths) $log_{10}$ basal metabolic rate: R = 0.1186, one-tailed (alternative hypothesis is directional) p-value = 0.0046875. Repeating this analysis with species set *B* instead, using (as before for species set *B*) untransformed branch lengths for $log_{10}$ adult body mass, and Nee transformed branch lengths for each of $log_{10}$ maximal lifespan, $log_{10}$ specific basal metabolic rate, and $log_{10}$ basal metabolic rate: excluding the effects of phylogeny and $log_{10}$ specific basal metabolic rate: R = 0.21 (one-tailed [alternative hypothesis is directional] p-value = 0.000055); excluding the effects of phylogeny and $log_{10}$ basal metabolic rate: R = 0.1039 (one-tailed [alternative hypothesis is directional] p-value = 0.028923). The values of this paragraph are close to (and some are lower than) those in the prior paragraphs, when the effect of $log_{10}$ adult body mass, instead of $log_{10}$ specific basal metabolic rate or $log_{10}$ basal metabolic rate, was excluded. That the correlation between $log_{10}$ maximal lifespan and $log_{10}$ adult body mass is attenuated when (atop of excluding the effect of phylogeny) excluding the effect of $log_{10}$ specific basal metabolic rate or $log_{10}$ basal metabolic rate harks back to the earlier shown collinearity/entwinement between $log_{10}$ adult body mass and $log_{10}$ specific basal metabolic rate, and $log_{10}$ adult body mass and $log_{10}$ basal metabolic rate.

Not taught by [323, 325] and done as a further step here, using PDAP output (generated with branch length transformations, or lack thereof, as in prior paragraphs of this headed section): a linear regression of standardized phylogenetically independent contrasts for $log_{10}$ specific basal metabolic rate (as dependent, *y*, variable) on standardized phylogenetically independent contrasts for $log_{10}$ adult body mass (as independent, *x*, variable), with intercept set as zero, was performed using [331], and its residuals saved. A linear regression, with intercept set as zero, was then conducted, using [331], between these residuals (as independent, *x*, variable) and (PDAP output) standardized phylogenetically independent contrasts for $log_{10}$ maximal lifespan (as dependent, *y*, variable), and its alongside reported Pearson correlation coefficient (R) {when intercept = zero} was noted. This analysis adjusts all three variables for phylogeny but only excludes any effect of $\log_{10}$ adult body mass upon $\log_{10}$ specific basal metabolic rate and (unlike the earlier analysis) doesn't also exclude any effect of $\log_{10}$ adult body mass upon $\log_{10}$ maximal lifespan. For species set *A*: R = -0.1505, one-tailed (alternative hypothesis is directional) p-value = 0.0004825 (if branch lengths are instead set to 1.0 for $\log_{10}$ specific basal metabolic rate: R = -0.1992, one-tailed p-value = 0.0000055). For species set *B*: R = -0.1663, one-tailed (alternative hypothesis is directional) p-value = 0.0011475. Repeating this analysis but with $\log_{10}$ basal metabolic rate instead of $\log_{10}$ specific basal metabolic rate: for species set *A*: R = 0.1162, one-tailed (alternative hypothesis is directional) p-value = 0.0054615. For species set *B*: R = 0.1181, one-tailed (alternative hypothesis is directional) p-value = 0.015471.

A linear regression of standardized phylogenetically independent contrasts for $\log_{10}$ adult body mass (as dependent, *y*, variable) against standardized phylogenetically independent contrasts for $\log_{10}$ specific basal metabolic rate (as independent, *x*, variable), with intercept set as zero, was performed using [331], and its residuals saved. A linear regression, with intercept set as zero, was then conducted, using [331], between these residuals (as independent, *x*, variable) and (PDAP output) standardized phylogenetically independent contrasts for $\log_{10}$ maximal lifespan (as dependent, *y*, variable), and its alongside reported Pearson correlation coefficient (R) {when intercept = zero} was noted. This analysis adjusts all three variables for phylogeny but only excludes any effect of $\log_{10}$ specific basal metabolic rate upon $\log_{10}$ adult body mass and (unlike earlier analysis) doesn't also exclude any effect of $\log_{10}$ specific basal metabolic rate upon $\log_{10}$ maximal lifespan. For species set *A*: R = 0.2087, one-tailed (alternative hypothesis is directional) p-value = 0.000002 (if branch lengths are instead set to 1.0 for $\log_{10}$ specific basal metabolic rate: R = 0.1749, one-tailed p-value = 0.0000605). For species set *B*: R = 0.1943, one-tailed (alternative hypothesis is directional) p-value = 0.0001775. Repeating this analysis but with $\log_{10}$ basal metabolic rate instead of $\log_{10}$ specific basal metabolic rate: for species set *A*: R = 0.1079, one-tailed (alternative hypothesis is directional) p-value = 0.0090825. For species set *B*: R = 0.09523, one-tailed (alternative hypothesis is directional) p-value = 0.0411215.

Table S28 below consolidates phylogenetically adjusted partial and semi-partial (part), forced through the origin [323, 325], Pearson correlation coefficients disclosed earlier in this headed section, showing the highest value reported in each case, which are all statistically significant (one-tailed p-values shown italicized in brackets). Where the dependent (*y*) variable is standardized phylogenetically independent contrasts for $\log_{10}$ maximal lifespan, *M* is standardized phylogenetically independent contrasts for $\log_{10}$ adult body mass, *sR* is standardized phylogenetically independent contrasts for $\log_{10}$ specific basal metabolic rate, *R* is standardized phylogenetically independent contrasts for $\log_{10}$ basal metabolic rate, and the variable shown in a bracket is that whose effect is statistically excluded.

[Table S28]

| | | Set | M (-*sR*) | M (-*R*) | sR (-*M*) | R (-*M*) |
|---|---|---|---|---|---|---|
| Maximal Lifespan | Partial | A | 0.2262<br>*(2.82324E-07)* | 0.1186<br>*(0.0046875)* | -0.2186<br>*(6.80284E-07)* | 0.1276<br>*(0.0025805)* |
| | | B | 0.21<br>*(0.000055)* | 0.1039<br>*(0.028923)* | -0.1809<br>*(0.0004485)* | 0.1284<br>*(0.0094495)* |
| | Semi-Partial | A | 0.2087<br>*(0.000002)* | 0.1079<br>*(0.0090825)* | -0.1992<br>*(0.0000055)* | 0.1162<br>*(0.0054615)* |
| | | B | 0.1943<br>*(0.0001775)* | 0.09523<br>*(0.0411215)* | -0.1663<br>*(0.0011475)* | 0.1181<br>*(0.015471)* |

Table S28 above shows that, after adjusting for phylogeny and adult body mass, at least across these 479 mammal and bird species (species set *A*), or just the 334 mammal species thereof (species set *B*), maximal lifespan *significantly* (negatively) correlates with specific basal metabolic rate (and significantly correlates with basal metabolic rate). Comparing the values of Table S28 above with earlier Table S27, one can see how much adjusting for adult body mass attenuates the phylogenetically adjusted correlation coefficients. Symmetrically (qualitative), at least for these species, adjustment for specific basal metabolic rate (or basal metabolic rate) attenuates the phylogenetically adjusted correlation (coefficient) between maximal lifespan and adult body mass. The lowest correlation coefficient, and highest p-value, in the table is for maximal lifespan vs. adult body mass with the effect of basal metabolic rate statistically excluded.

Partial and semi-partial correlation coefficients and p-values thereof are interrelated with regression coefficients and p-values thereof of multiple linear regression. An assumption of multiple linear regression is that there is no collinearity between the explanatory/predictor variables. As shown in earlier sections, there *is* collinearity between $\log_{10}$ adult body mass and $\log_{10}$ specific basal metabolic rate (and even greater collinearity between $\log_{10}$ adult body mass and $\log_{10}$ basal metabolic rate), which presumably impacts the analysis of this present section also. It would certainly impact this present section if the regressions were done before instead of (as was done) after the phylogeny step. For example, if the $\log_{10}$ specific basal metabolic rate (or $\log_{10}$ basal metabolic rate) on $\log_{10}$ adult body mass, and $\log_{10}$ maximal lifespan on $\log_{10}$ adult body mass, regressions were done before, with neither forced through the origin, and the two sets of residuals thereof were inputted into the (PDAP conferred) phylogeny step. Whereas there might be (or might not be) at least some remedy in doing, as was done, the phylogeny step first, and regressions thereafter. *If* the phylogeny adjustment sufficiently reduces the collinearity between adult body mass and specific basal metabolic rate, and between adult body mass and basal metabolic rate, related variables. The next headed section reports that the phylogeny adjustment used to produce the results of Table S28 above, wherein the standardized phylogenetically independent contrasts for $\log_{10}$ adult body mass are generated with different branch lengths than the standardized phylogenetically independent contrasts for $\log_{10}$ specific basal metabolic rate and $\log_{10}$ basal metabolic rate, does reduce collinearity (quantified by presented VIF values). But some collinearity endures, especially pertaining to basal metabolic rate, and so collinearity is still impacting the results presented in Table S28 above. Moreover, as said at the beginning of this headed section, the use of different branch lengths for different variables means these results are not *prima facie* interpretable. Much easier to interpret are, as presented in the next headed section, results from when the same branch lengths are used for the different variables. But then there is less attenuation of collinearity, and thence more collinearity.

Multiple linear regression with standardized phylogenetically independent contrasts

Using standardized phylogenetically independent contrasts of the last headed section, Table S29 below reports multiple linear regression results, where standardized phylogenetically independent contrasts for $\log_{10}$ maximal lifespan is the response variable and standardized phylogenetically independent contrasts for $\log_{10}$ adult body mass and standardized phylogenetically independent contrasts for $\log_{10}$ specific basal metabolic rate are explanatory variables. In the table, $p\_\beta_1$ and $p\_\beta_2$ are p-values for unstandardized regression coefficients, and $S\beta_1$ and $S\beta_2$ are standardized regression coefficients, for these explanatory variables respectively. *Adj.* $R^2$ is adjusted $R^2$, *VIF* is the Variance Inflation Factor for each of the slope regression coefficients, *DW* is the Durbin-Watson statistic, and *p_F* is the p-value for the F-statistic. For species sets *A* and *B*. Regression was forced through the origin, following the teaching of [323, 325]. Conducted in JASP software (version 0.18.1.0), using its method called "Enter", wherein the adult body mass related predictor was forced in first. Assumptions of homoscedasticity, normality, and linearity weren't violated.

[Table S29]

| Set | Adj. $R^2$ | VIF | DW | p_F | S_$\beta_1$ | S_$\beta_2$ | p_$\beta_1$ | p_$\beta_2$ |
|---|---|---|---|---|---|---|---|---|
| A | 0.189 | 1.784 | 2.061 | 8.159e-23 | 0.279 | -0.201 | 5.820E-7 | 2.863E-4 |
| B | 0.177 | 1.702 | 2.108 | 3.631e-15 | 0.253 | -0.216 | 1.128E-4 | 9.155E-4 |

Table S30 below is as Table S29 above, except standardized phylogenetically independent contrasts for $\log_{10}$ basal metabolic rate were used in place of standardized phylogenetically independent contrasts for $\log_{10}$ specific basal metabolic rate. Greater *VIF* values (collinearity) observed. *VIF* values of 3.9 and 4. This is less than without any phylogeny adjustment (>16, as shown earlier) but is still notable collinearity.

[Table S30]

| Set | Adj. $R^2$ | VIF | DW | p_F | S_$\beta_1$ | S_$\beta_2$ | p_$\beta_1$ | p_$\beta_2$ |
|---|---|---|---|---|---|---|---|---|
| A | 0.180 | 3.916 | 2.052 | 1.191e-21 | 0.213 | 0.230 | 0.009 | 0.005 |
| B | 0.163 | 3.964 | 2.096 | 5.698e-14 | 0.189 | 0.235 | 0.058 | 0.019 |

Multiple linear regression in JASP software (version 0.18.1.0), if selected to, reports corresponding partial and semi-partial correlation coefficients. The corresponding partial and semi-partial correlation coefficients reported by JASP (version 0.18.1.0) for the multiple regressions of Tables S29 and S30 above are as shown in earlier Table S28, so reproducing those values.

Whereas a (forced through the origin) Pearson correlation coefficient between phylogenetically independent contrasts isn't changed much by the contrasts being generated with different branch lengths [323], as observed herein the slopes of a multiple linear regression can be, which means partial and semi-partial correlation coefficients can be. Multiple linear regression, and partial and semi-partial correlation results are presented below. Where, distinct from earlier, the *same* branch length transformation was used to generate all the variables (contrasts). The partial and semi-partial correlation results are meaningfully different than those earlier. This is probably at least in part because employing different branch length transformations to generate the adult body mass and specific basal metabolic rate (and the adult body mass and basal metabolic rate) related variables attenuated the collinearity between adult body mass and specific basal metabolic rate, and between adult body mass and basal metabolic rate, more. With a single branch length transformation used there was such attenuation, but less, such that the enduring collinearity was more, which is probably at least in part why the results differ. For the basal metabolic rate results especially, this enduring collinearity is substantial.

Tables S31 and S32 below are like Tables S29 and S30 above respectively, except that below a single branch length transformation was used to generate the different variables. For species set *A* this was setting branch lengths to 1.0, and for species set *B* this was the Nee branch length transformation. By the earlier diagnostic table, Table 26, the Nee branch length transformation is appropriate to use for all the different variables with species set *B*. But, referring to this table, for species set *A*, there is no single branch length transformation (or lack thereof) there that is appropriate for all four different variables. Branch lengths set as 1.0 is appropriate for two of the four variables (specific basal metabolic rate and maximal lifespan), and this was selected. This selection corresponds with the sole branch length transformation employed in [6]. Among the differences, the *VIF* values are higher in the tables below, the ratio between the standardized coefficients is very different in each case below than above, and standardized coefficients for basal metabolic rate are of the opposite sign below than above, highlighted by being in red text below. Assumptions of homoscedasticity, normality, and linearity weren't violated.

[Table S31]

| Set | Adj. $R^2$ | VIF | DW | p_F | $S\_\beta_1$ | $S\_\beta_2$ | $p\_\beta_1$ | $p\_\beta_2$ |
|---|---|---|---|---|---|---|---|---|
| A | 0.231 | 2.367 | 2.037 | 2.501e-28 | 0.362 | -0.147 | 7.982E-9 | 0.017 |
| B | 0.195 | 2.141 | 2.092 | 8.671e-17 | 0.345 | -0.127 | 2.252E-6 | 0.078 |

[Table S32]

| Set | Adj. $R^2$ | VIF | DW | p_F | $S\_\beta_1$ | $S\_\beta_2$ | $p\_\beta_1$ | $p\_\beta_2$ |
|---|---|---|---|---|---|---|---|---|
| A | 0.222 | 7.509 | 2.018 | 4.240e-27 | 0.485 | -0.012 | 1.366E-5 | 0.912 |
| B | 0.188 | 6.899 | 2.080 | 3.825e-16 | 0.482 | -0.048 | 2.289E-4 | 0.713 |

Table S33 below has the same format and headings as Table S28 in the last headed section. It shows partial and semi-partial (forced through origin [323, 325]) Pearson correlation coefficients correspondent with the multiple linear regression results presented in Tables S31 and S32 above. They are smaller than the partial and semi-partial correlation coefficients presented in earlier Table S28, and in the case of the *R(-M)* column are of the opposite sign (highlighted by red coloured text). This is probably at least in part related to the *VIF* values, shown in Tables S31 and S32 above, especially for basal metabolic rate (*VIF* ≥ 6.9). One-tailed (alternative hypothesis is directional) p-values shown italicized in brackets.

[Table S33]

| | | Set | M (-*sR*) | M (-*R*) | sR (-*M*) | R (-*M*) |
|---|---|---|---|---|---|---|
| Maximal Lifespan | Partial | A | 0.260<br>*(3.83901E-09)* | 0.197<br>*(0.000007)* | -0.109<br>*(0.0085055)* | -0.005<br>*(0.5434563)* |
| | | B | 0.256<br>*(0.000001)* | 0.201<br>*(0.000109)* | -0.097<br>*(0.038341)* | -0.020<br>*(0.642138)* |
| | Semi-Partial | A | 0.236<br>*(8.68555E-08)* | 0.177<br>*(0.000049)* | -0.096<br>*(0.0178455)* | -0.004<br>*(0.5347898)* |
| | | B | 0.237<br>*(0.000006)* | 0.184<br>*(0.0003635)* | -0.087<br>*(0.0562525)* | -0.018<br>*(0.6284517)* |

Table S33 above reports that, after adjusting for phylogeny and adult body mass, across 479 mammal and bird species (species set *A*), maximal lifespan *significantly* (negatively) correlates with specific basal metabolic rate.

Collinearity breaches an assumption of multiple linear regression, corrupts its results, and corrupts interrelated partial and semi-partial correlation results. It can even corrupt to the extent of making the sign of regression, partial and semi-partial correlation coefficients erroneous. Presumably, collinearity has corrupted the results presented in Table S33 above, especially those of its *R(-M)* column. The phylogenetic adjustment used attenuates but doesn’t cure the collinearity problem. Perhaps collinearity/entwinement between specific basal metabolic rate and adult body mass (*and even more so between basal metabolic rate and adult body mass*) precludes isolation of the specific effect (on maximal lifespan) of either. And phylogenetic adjustment can’t change this.

Incidentally, the Durbin-Watson statistic (*DW*) is higher in the tables of this section than in earlier Tables S14-S16. So, phylogeny might be contributory to the low *DW* values (positive autocorrelation) reported in earlier Tables S14-S16.

Principal Component Analysis (PCA)

I performed Principal Component Analysis (PCA) upon the last-used phylogenetically independent contrasts (*using the "factoextra" R package. R code used is in the supplementary materials part of this document. Code performs centering and scaling, in that order, before PCA. Maximum number of components allowable by the data retained. Rotation is orthogonal rather than oblique, making output components uncorrelated*). Performed for specific basal metabolic rate and adult body mass contrasts together. And for basal metabolic rate and adult body mass contrasts together. For species sets (*A* or *B*) separately. Not performed with maximal lifespan contrasts.

I intended to use these PCA-adjusted contrasts as predictors in principal component regression but abandoned this approach because, upon using the "pca_result$rotation" command in R to analyse the PCA output, it became clear that each two input variables, e.g., specific basal metabolic rate and adult body mass contrasts from species set *B*, weren't sufficiently separated across the two output variables, called PC1 and PC2, but were each reflected in both. Shown in Table S34 below, their loadings matrix, disclosing correlation coefficients between the original variables and principal components (PC1 and PC2) output.
[Table S34]

| | PC1 | PC2 |
|---|---|---|
| Specific basal metabolic rate contrasts | -0.7071068 | 0.7071068 |
| Adult body mass contrasts | 0.7071068 | 0.7071068 |

Using PCA-adjusted variables in principal component regression can be a fix for collinearity/multicollinearity in multiple linear regression. But here we see that the variables are so entwinned that PCA fails to discriminate the variables into distinct output variables, both the input variables being blended into both the output variables. Stymying the ability to do principal component regression.

Incidentally, when the contrasts used to generate Table S34 are used in Principle Component Analysis (PCA) in JASP (version 0.18.3.0), manually specifying for there to be two components, selecting rotation to be orthogonal instead of oblique, selecting to display loadings above 0 (rather than just above 0.4 as it is by default), the loadings matrix generated is shown in Table S35 below, again showing no separation of the inputs in the outputs. Shown diagrammatically in Figure S5.

[Table S35]

| | PC1 | PC2 |
|---|---|---|
| Specific basal metabolic rate contrasts | -0.938 | 0.347 |
| Adult body mass contrasts | 0.938 | 0.347 |

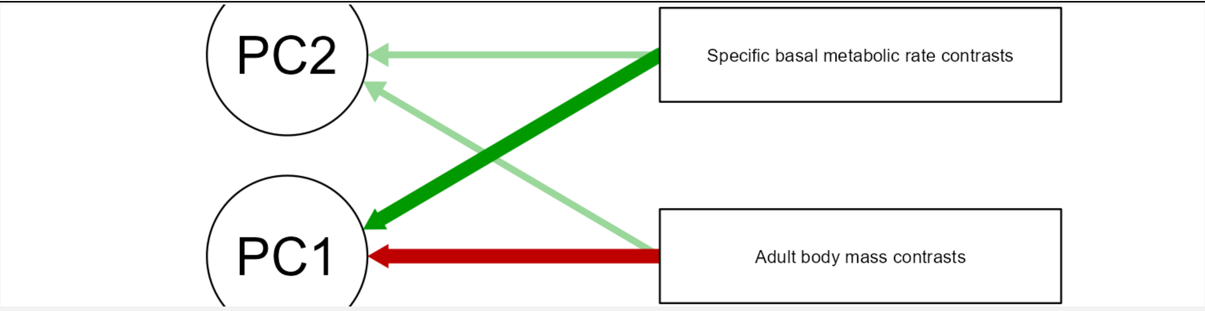

Figure S5 – Path diagram corresponding to the loadings matrix of Table S35. Generated by JASP (version 0.18.3.0).

Although, across mammals and birds, entwinement between specific basal metabolic rate and adult body mass, or their contrasts, stymies their simultaneous use as predictors in multiple linear regression, and even in principal component regression, there is information gained from this strong degree of entwinement. Interpreted to be that adult body mass, by the square-cube law of geometry, dictates metabolic heat generation per unit mass required/performed, and thence is a major determinant of specific basal metabolic rate.

Degrees of freedom

In the phylogenetic analysis herein, for a Pearson correlation forced through the origin, degrees of freedom (*df*) were calculated by $df = number\ of\ species - 2 = number\ of\ contrasts - 1$ [325]. This assumes that any polytomies are "hard", due to a multiway (rather than just a dichotomous) speciation event, and not "soft", being due to a lack of knowledge. An alternative equation is $df = number\ of\ species - 2 - number\ of\ soft\ polytomies$. If this alternative equation is used instead, assuming that all polytomies are soft, such that species sets *A* and *B* have 20 and 14 soft polytomies respectively, then no earlier presented significant p-value (for a Pearson correlation forced through the origin) loses statistical significance (at $p < 0.05$). Incidentally, I've discovered a bug in PDAP module version 1.16 [320] of Mesquite version 3.81 (build 955) [321], in that it doesn't subtract the number of polytomies from the degrees of freedom it uses, even when directed to through its Graphical User Interface (GUI).

Epigenetic clock "ticking" rate (*ΔM*) and phylogeny

*ΔM*, given by Eq. 5 of the main text, is the epigenetic clock's "ticking" rate. For the 50 species used to produce Figure 9 of the main text, the Pearson correlation coefficient between *ΔM* and specific basal metabolic rate is 0.7269, and that between *ΔM* and adult body mass is -0.2551. Do these correlations endure when phylogeny is considered?

The phylogenetic independent contrasts method requires compared traits to be linearly related [324]. Whereas the relation between *ΔM* and specific basal metabolic rate is linear, that between *ΔM* and adult body mass is not (and incidentally, nor is that between specific basal metabolic rate and adult body mass). When all these variables are logged ($\log_{10}$) their interrelations are linear. The tables of this headed section present results from using logged variables.

Phylogenetic relationships between the 50 species used to produce Figure 9 were sourced from [322]. Two species (*Eptesicus fuscus* and *Rattus rattus*) were then excluded because of an "unresolved name" issue, rendering a phylogeny for 48 different species, with no (zero) polytomies.

To compare with values below: without any phylogenetic adjustment, for these 48 species, when variables are logged ($\log_{10}$): the Pearson correlation coefficient between $\log_{10}$ *ΔM* and $\log_{10}$ specific basal metabolic rate is 0.7313, and that between $\log_{10}$ *ΔM* and $\log_{10}$ adult body mass is -0.6807.

For this species set, using specific basal metabolic rate data from [1] and *ΔM* data calculated by Eq. 5 from maximal lifespan data from [1], in Table S36 below, for line *A*: *C* is the (forced through origin [323, 325]) Pearson correlation coefficient between phylogenetically independent contrasts of $\log_{10} \Delta M$ and $\log_{10}$ specific basal metabolic rate (*and in the line below, labelled "P", the corresponding one-tailed [alternative hypothesis is directional] p-value thereof*), and subsequent columns on that row are that but with a branch length adjustment made (with column headings as codified earlier). For line *B*: *C* is the (forced through origin [323, 325]) Pearson correlation coefficient between phylogenetically independent contrasts of $\log_{10} \Delta M$ and $\log_{10}$ adult body mass (*and in the line below, labelled "P", the corresponding one-tailed [alternative hypothesis is directional] p-value thereof*), and subsequent columns on that row are that but with a branch length adjustment made (with column headings as codified earlier). So, this table discloses phylogenetically adjusted Pearson correlation coefficients. Outputted by the PDAP module [320] of Mesquite [321]. [Table S36]

| | *C* | Nee | Pagel | Grafen | =1.0 |
|---|---|---|---|---|---|
| **A** | 0.5586058420998354 | 0.6207192150029445 | 0.6141994229825142 | 0.602000610549938 | 0.5996456182513439 |
| *P* | 1.8445478996875764E-5 | 1.259558338739808E-6 | 1.7153475853071143E-6 | 3.0018795409425536E-6 | 3.335511383872891E-6 |
| **B** | -0.5475783527905894 | -0.5964005384833854 | -0.5904587422730392 | -0.5860946655630987 | -0.5759696796080356 |
| *P* | 2.812779428605605E-5 | 3.851568791873579E-6 | 4.9920878575452E-6 | 6.020124642648603E-6 | 9.200685099950125E-6 |

In Table S36 above, one can see that even with a phylogeny adjustment, across (at least these) species, there is a statistically significant relation between $\log_{10} \Delta M$ and $\log_{10}$ specific basal metabolic rate (*mean correlation across the different columns of line A = 0.59903414177. But according to the diagnostic table below, Table S37, only columns "C", "Nee" and "=1.0"* *should be considered, and their mean = 0.59299022511*), and there is a statistically significant relation between $\log_{10} \Delta M$ and $\log_{10}$ adult body mass (*mean correlation across the different columns of line B = -0.57930039574. But according to the diagnostic table below, Table S37, only columns "C", "Nee" and "=1.0" should be considered, and their mean = -0.57331619029*).

The diagnostic table below, Table S37, shows for $Q$ ($\log_{10}$ specific basal metabolic rate), $R$ ($\log_{10} \Delta M$), and $T$ ($\log_{10}$ adult body mass): in column $C$, the Pearson correlation coefficient (*not* forced through origin [323, 325]) of the absolute values of the standardized phylogenetically independent contrasts versus their standard deviations (i.e., the square root of the sum of their branch lengths), which is a diagnostic check of how well the topology and branch lengths of the phylogenetic tree (and assumed model of character evolution) adequately fit the tip data [323], wherein no significant correlation is the desirable result (*and in the line below, labelled "P", the corresponding two-tailed p-value thereof*), and subsequent columns are that but with a branch length adjustment made, as codified earlier. P-values greater than 0.05, which are desirable, are in red text. Table values were outputted by the PDAP module [320] of Mesquite [321].

[Table S37]

| Diagnostic table | | | | | |
|---|---|---|---|---|---|
| | *C* | Nee | Pagel | Grafen | =1.0 |
| **Q** | -0.027277197917221514 | -0.21514586718379863 | -0.3072774962152256 | -0.3457506254938258 | -0.18425530582321456 |
| *P* | 0.8555817276996744 | 0.1464132368326061 | 0.03564189888523807 | 0.01729153964426025 | 0.21504050021182963 |
| **R** | -0.03918731101624927 | -0.12414835126388174 | -0.2416318114169313 | -0.33405940038804544 | -0.11157930833622677 |

| | | | | | |
|---|---|---|---|---|---|
| *P* | 0.7936908716474487 | 0.40573464608788357 | 0.10177986256432536 | 0.021739355728941156 | 0.4552523841756628 |
| **T** | -0.13651624997104528 | -0.12164771108156246 | -0.26058212684101795 | -0.35800148149129973 | -0.10692227380370599 |
| *P* | 0.36019148513361665 | 0.41533035080490543 | 0.07688382713226168 | 0.013483250250432592 | 0.474400067884575 |

Following the teaching of [323, 321], raw phylogenetically independent contrasts were outputted from the PDAP module of Mesquite, and each was then standardized by dividing each contrast by the (also PDAP outputted) contrast's standard deviation (square root of sum of corrected branch lengths). The raw contrasts, and corresponding standard deviations to standardize them, for all the variables ($\log_{10}$ specific basal metabolic rate, $\log_{10}$ adult body mass, $\log_{10} \Delta M$) were sourced with no branch length transformation, as this (according to the diagnostic table, Table S37, above) is the best (highest p-value of those trialled) for each of these three variables.

Table S38 below shows results of multiple linear regression conducted in JASP software (version 0.18.1.0), forced through the origin [323, 321], with standardized phylogenetically independent contrasts, where such contrasts for $\log_{10} \Delta M$ is the response variable and such contrasts for $\log_{10}$ adult body mass and $\log_{10}$ specific basal metabolic rate are explanatory variables. In the table, $p\_\beta_1$ and $p\_\beta_2$ are p-values for unstandardized regression coefficients, and $S\beta_1$ and $S\beta_2$ are standardized regression coefficients, for these explanatory variables respectively. *Adj.* $R^2$ is adjusted $R^2$, *VIF* is the Variance Inflation Factor for each of the slope regression coefficients, *DW* is the Durbin-Watson statistic, and $p\_F$ is the p-value for the F-statistic. Used JASP's method called "Enter", wherein the $\log_{10}$ adult body mass related predictor was forced in first. Assumptions of homoscedasticity, normality, and linearity weren't violated.

[Table S38]

| Adj. $R^2$ | VIF | DW | p_F | S_$\beta_1$ | S_$\beta_2$ | p_$\beta_1$ | p_$\beta_2$ |
|---|---|---|---|---|---|---|---|
| 0.303 | 3.476 | 1.653 | 1.135e-4 | -0.267 | 0.338 | 0.250 | 0.147 |

Table S39 below presents corresponding JASP-outputted (version 0.18.1.0) partial and semi-partial correlation coefficients, where the response variable is standardized phylogenetically independent contrasts for $\log_{10}$ *ΔM*, *M* is standardized phylogenetically independent contrasts for $\log_{10}$ adult body mass, *sR* is standardized phylogenetically independent contrasts for $\log_{10}$ specific basal metabolic rate, and the variable shown in a bracket is that whose effect is statistically excluded. One-tailed (alternative hypothesis is directional) p-values shown italicized in brackets.

[Table S39]

| | | sR (-*M*) | M (-*sR*) |
|---|---|---|---|
| *ΔM* | Partial | 0.215<br>*(0.071117)* | -0.171<br>*(0.122602)* |
| | Semi-partial | 0.180<br>*(0.110432)* | -0.142<br>*(0.1678335)* |

Table S39 above reports that, after phylogeny adjustment, *ΔM* correlates better with specific basal metabolic rate, with the effect of adult body mass excluded, than with adult body mass, with the effect of specific basal metabolic rate excluded. Note, given the appreciable *VIF* value in Table S38, that there is collinearity. Wherein collinearity can corrupt multiple linear regression coefficients. And interrelated partial and semi-partial correlation coefficients.

Principal component analysis (PCA), using these contrasts of specific basal metabolic rate and adult body mass, was tried to try to circumnavigate this collinearity. But I hit the same problem as earlier, with the inputs blended (rather than separated) in the PC1 and PC2 outputs, and so this approach wasn't pursued further.

A drawback of this section's analysis is that it was conducted with $\log_{10}$ transformed variables, which washes out the non-linearity between *ΔM* and adult body mass (that distinct from the linearity between $\Delta M$ and specific basal metabolic rate), which is a salient aspect of the data. When unlogged data is used and so this non-linearity endures, breaching an assumption of the Phylogenetically Independent Contrasts (PIC) method [324], then after such PIC phylogenetic adjustment, there *is* a significantly significant relation between $\Delta M$ and specific basal metabolic rate, and *no* statistically significant relation between $\Delta M$ and adult body mass (not shown, improper analysis).

Specific $F_1F_0$ ATP hydrolysis rate and phylogeny

Across species, the negative correlation between maximal lifespan (data from [1], excluding human by rationale in Methods) and specific $F_1F_0$ ATP hydrolysis rate (data from [91]), reported in Table 3 of the main text, endures at statistical significance when phylogeny is accounted for. The (forced through origin [323, 325]) Pearson correlation coefficient between phylogenetically independent contrasts of $log_{10}$ maximal lifespan and $log_{10}$ specific $F_1F_0$ ATP hydrolysis rate, with no branch length adjustment, is -0.6168510956956115, one-tailed (alternative hypothesis is directional) p-value = 0.021611317854471027.

For comparison, across these same species, the (forced through origin [323, 325]) Pearson correlation coefficient between phylogenetically independent contrasts of $log_{10}$ maximal lifespan (data from [1]) and $log_{10}$ body mass (data from [91]), with no branch length adjustment, is 0.5581427898280978, one-tailed (alternative hypothesis is directional) p-value = 0.03718063405133687.

Using logged data, with no branch length adjustment, using the method used earlier herein, as taught in [323, 325] (conferring a partial correlation), the Pearson correlation coefficient (R) between $log_{10}$ maximal lifespan and $log_{10}$ specific $F_1F_0$ ATP hydrolysis rate, adjusting for $log_{10}$ body mass and phylogeny = -0.4838, one-tailed (alternative hypothesis is directional) p-value = 0.0658055. So, across at least these species, with the effects of phylogeny and $log_{10}$ body mass excluded, $log_{10}$ maximal lifespan inversely correlates with $log_{10}$ specific $F_1F_0$ ATP hydrolysis rate. Closely missing significance at $p < 0.05$, possibly because, although R is (absolutely) sizeable (=-0.4838), $n$ is small (*11 species*). This inverse correlation is greater than the positive correlation between $log_{10}$ maximal lifespan and $log_{10}$ body mass with the effects of phylogeny and $log_{10}$ specific $F_1F_0$ ATP hydrolysis rate excluded: Pearson = 0.3857, one-tailed (alternative hypothesis is directional) p-value = 0.1206925. So, at least across these species, when the effects of phylogeny and specific $F_1F_0$ ATP hydrolysis rate are excluded, maximal lifespan does *not* significantly correlate with body mass.

The last paragraph reported partial correlation coefficients. Corresponding semi-partial (part) correlation coefficients for $\log_{10}$ specific $F_1F_0$ ATP hydrolysis rate and $\log_{10}$ body mass are -0.401 and 0.304 respectively.

The (as earlier used) diagnostic, the Pearson correlation coefficient (not forced through origin) of the absolute values of the standardized phylogenetically independent contrasts versus their standard deviations (i.e., the square root of the sum of their branch lengths) [323, 325], is not significant, and so is good, for each of the logged variables, at least when there is no branch length transformation. As shown in Table S40 below, where *R* is the Pearson correlation coefficient and *P* is the two-tailed p-value.
[Table S40]

| Diagnostic table | | | |
|---|---|---|---|
| | Specific $F_1F_0$ ATP hydrolysis rate | Maximal lifespan | Body mass |
| R | -0.10766045579745312 | -0.054525326007089564 | -0.4819978237354271 |
| P | 0.7672035328687943 | 0.8810798206462368 | 0.15832696145578445 |

The phylogeny for this headed section, which had no (zero) polytomies, was sourced from [322].

The Phylogenetic Independent Contrasts (PIC) method requires compared traits to be linearly related [324]. The relation between maximal lifespan and each of specific $F_1F_0$ ATP hydrolysis rate and body mass is non-linear, becoming linear upon logging the variables. When unlogged variables are used, the non-linearity breaching an assumption of PIC, after PIC phylogenetic adjustment, the relationship between maximal lifespan and specific $F_1F_0$ ATP hydrolysis rate *is*, and that between maximal lifespan and body mass is *not*, statistically significant (not shown, improper analysis).

<u>P-value for semi-partial (part) correlations</u>

It may be that when calculating the p-value of a semi-partial (part) correlation the degrees of freedom should be less by *g*, the number of controlled variables (=1 here) [332-333], such that the degrees of freedom value is one less than would be used when typically calculating a p-value (or a soft polytomy or soft polytomies correction thereof). But this wasn't employed herein. If the reader wants to employ this method at their own initiative with semi-partial correlation coefficient values herein: for a two-tailed p-value, the following R statistical software [92] code can be used, where *p* is the p-value, *N* is the sample size, *g* is the number of controlled variables (1 for cases herein), and *R* is the semi-partial correlation coefficient:

```
df=N-2-g
Tog=R*sqrt(df/(1-R^2))
p=pt(Tog, df, lower = FALSE)*2
```

For a one-tailed p-value, the "*2" term is omitted, and "lower = FALSE" or "lower = TRUE" is used depending on the directionality of the alternative hypothesis.

<u>Ridge regression</u>

Using JASP software (version 0.18.1.0; using the regularized linear regression suite in its machine learning module) with earlier-defined species sets, where set *A* is 517 mammal and bird species, set *B* is 350 mammal species, and set *C* is 263 mammal species (excluding bats, naked mole rat, and marsupials), I performed ridge regression, which is less susceptible (but not impervious) to collinearity between predictors. Using data from [1], with maximal lifespan (years) as the response variable and adult body mass (kg) and specific basal metabolic rate (W/g) as predictors, all logged ($log_{10}$). In Table S41 below: $S\_\beta_0$, $S\_\beta_1$, and $S\_\beta_2$ are standardized regression coefficients for intercept, adult body mass, and specific basal metabolic rate respectively, *RMSE* is the root-mean-square error, and λ is the regularization parameter. In each case, returned values from three different optimization runs are shown and, in italics, the result when λ is fixed to equal one (1), which at least in some cases confers a higher RMSE and/or $R^2$ value. The higher the value of λ, the less susceptible the model is to collinearity, at the cost of greater bias in the model.

[Table S41]

| Set | RMSE | $R^2$ | λ | $S\_\beta_0$ | $S\_\beta_1$ | $S\_\beta_2$ |
|---|---|---|---|---|---|---|
| A | 0.77 | 0.367 | 0.053 | -5.716e-4 | 0.548 | 0.048 |
| | 0.838 | 0.287 | 0.060 | 0.020 | 0.554 | 0.030 |
| | 0.759 | 0.416 | 0.056 | 0.006 | 0.545 | 0.008 |
| | *0.792* | *0.273* | *1* | *-0.020* | *0.239* | *-0.131* |
| B | 0.696 | 0.517 | 0.063 | -0.005 | 0.374 | -0.309 |

| | | | | | | |
|---|---|---|---|---|---|---|
| | 0.741 | 0.426 | 0.069 | 0.030 | 0.370 | -0.329 |
| | 0.797 | 0.389 | 0.074 | 0.009 | 0.431 | -0.269 |
| | *0.794* | *0.409* | *1* | *-0.032* | *0.242* | *-0.225* |
| C | 0.549 | 0.669 | 0.078 | -0.031 | 0.487 | -0.320 |
| | 0.532 | 0.689 | 0.081 | 0.010 | 0.426 | -0.418 |
| | 0.628 | 0.518 | 0.084 | 0.010 | 0.466 | -0.390 |
| | *0.675* | *0.679* | *1* | *0.036* | *0.275* | *-0.271* |

Table S42 below is as Table S41 above except that $log_{10}$ basal metabolic rate (W) was used instead of $log_{10}$ specific basal metabolic rate.

[Table S42]

| Set | RMSE | $R^2$ | λ | S_$\beta_0$ | S_$\beta_1$ | S_$\beta_2$ |
|---|---|---|---|---|---|---|
| A | 0.784 | 0.314 | 0.076 | -0.027 | 0.251 | 0.332 |
| | 0.751 | 0.36 | 0.054 | -0.018 | 0.274 | 0.280 |
| | 0.859 | 0.369 | 0.082 | -0.009 | 0.223 | 0.306 |
| | *0.841* | *0.335* | *1* | *-0.010* | *0.188* | *0.196* |
| B | 0.715 | 0.31 | 0.072 | 0.003 | 0.536 | 0.106 |
| | 0.829 | 0.418 | 0.071 | 0.016 | 0.573 | 0.059 |
| | 0.85 | 0.413 | 0.067 | -0.039 | 0.508 | 0.135 |
| | *0.712* | *0.513* | *1* | *0.003* | *0.236* | *0.196* |
| C | 0.585 | 0.736 | 0.074 | 0.022 | 0.587 | 0.136 |
| | 0.564 | 0.697 | 0.072 | -5.106e-4 | 0.558 | 0.176 |
| | 0.686 | 0.576 | 0.084 | 0.001 | 0.619 | 0.147 |
| | *0.596* | *0.76* | *1* | *-0.032* | *0.277* | *0.228* |

Tables S43 and S44 below are generated using the same phylogenetically independent contrasts data used to produce earlier Tables S31 and S32, except here ridge regression is used. Forced through the origin [323, 325].

[Table S43]

| Set | RMSE | $R^2$ | λ | S_$\beta_1$ | S_$\beta_2$ |
|---|---|---|---|---|---|
| A | 0.991 | 0.264 | 0.149 | 0.255 | -0.187 |
| | 0.843 | 0.078 | 0.052 | 0.293 | -0.233 |
| | 0.907 | 0.292 | 0.150 | 0.264 | -0.202 |

| | | | | | |
|---|---|---|---|---|---|
| | *0.829* | *0.273* | *1* | *0.181* | *-0.141* |
| B | 0.996 | 0.178 | 0.043 | 0.362 | -0.058 |
| | 0.822 | 0.222 | 0.155 | 0.289 | -0.112 |
| | 1.014 | 0.156 | 0.202 | 0.221 | -0.208 |
| | *0.802* | *0.371* | *1* | *0.162* | *-0.119* |

[Table S44]

| Set | RMSE | $R^2$ | λ | $S\_\beta_1$ | $S\_\beta_2$ |
|---|---|---|---|---|---|
| A | 0.89 | 0.163 | 0.048 | 0.416 | 0.075 |
| | 0.981 | 0.266 | 0.055 | 0.310 | 0.119 |
| | 0.837 | 0.303 | 0.038 | 0.333 | 0.098 |
| | *0.985* | *0.141* | *1* | *0.179* | *0.142* |
| B | 1.167 | 0.067 | 0.042 | 0.385 | 0.029 |
| | 0.831 | 0.273 | 0.213 | 0.194 | 0.109 |
| | 0.944 | 0.297 | 0.165 | 0.203 | 0.107 |
| | *0.962* | *0.023* | *1* | *0.179* | *0.154* |

In Tables S41-S44 above, in each case, for at least some presented value(s) of λ, $S\_\beta_2$ is of appreciable absolute value as compared to $S\_\beta_1$. Unlike earlier multiple linear regression results (Tables S14, S16, and S32), here all the slope coefficients have the correct sign, except for instances with species set *A* in Table S41, which is rectified by a higher value of λ (=1). These ridge regression results are consistent with the statement: across species, specific basal metabolic rate is a predictor of maximal lifespan.

Generalized Linear Model (GLM)

Unlike multiple linear regression (and ridge regression), a Generalized Linear Model (GLM) doesn't require linearity between the response variable and predictors (if such linearity can be conferred by a link function in the GLM). So, it can be used with untransformed variables of adult body mass and specific basal metabolic rate, which are less collinear than their logged ($log_{10}$) versions, and so using GLM might solve the collinearity problem.

A GLM has a “systemic component” (“link”) that confers linearity between the response variable and predictors, and it has a “random component” that specifies an appropriate probability distribution for the response variable.

Using data from [1], I constructed a GLM in JASP software (version 0.18.1.0) with maximal lifespan (years) as the response variable, and adult body mass (kg) and specific basal metabolic rate (W/g) as predictors (all unlogged). I mandated that the response variable should be positive continuous, which restricted the random component options available in JASP to Gamma or Inverse Gaussian, with corresponding systemic component options available of Identity, Log, Inverse, and $1/\mu^2$. I tested combinations thereof, using data from 350 mammal species (species set *B*), to find which combination conferred the model with the lowest chi-squared p-value, which was Gamma Inverse.

Results are in Table S45 below, alongside results for species sets *A* and *C*, also retrieved with Gamma Inverse. Where *Q* is the model’s chi-squared ($\chi^2$) p-value, *VIF* is Variance Inflation Factor, *S* is the intercept coefficient (*S-p* is p-value thereof), *T* is the coefficient for adult body mass (*T-p* is p-value thereof), and *U* is the coefficient for specific basal metabolic rate (*U-p* is p-value thereof).

[Table S45]

| Set | Q | VIF | S | T | U | S-p | T-p | U-p |
|---|---|---|---|---|---|---|---|---|
| A | 6.890e-25 | 1.060 | 0.042 | -1.078e-5 | 3.758 | 6.701e-52 | 2.058e-6 | 1.021e-18 |
| B | 5.371e-33 | 1.084 | 0.031 | -8.351e-6 | 10.921 | 1.804e-18 | 6.212e-4 | 4.168e-24 |
| C | 4.240e-50 | 1.068 | 0.021 | -5.648e-6 | 14.792 | 2.433e-13 | 0.007 | 3.203e-37 |

In Table S45 above, the p-values for specific basal metabolic rate are highly significant, and smaller than for adult body mass. So, even with adult body mass included in the model, specific basal metabolic rate is a highly significant predictor of maximal lifespan.

*Diagnostic graphs for these GLMs: Quantile residuals vs. fitted (which is Quantile residuals vs. predicted values of the response variable) and Quantile residuals vs. predictor plots shouldn’t contain pattern(s) and/or trend(s), but there are some outlier data points which makes the other data points appear bunched in the graphs, which might be misconstrued as regularity. That aside, there might be minor non-randomness in these plots. The Q-Q plot should be linear and, some outliers aside, is linear, although it has some slight non-linearity at its ends. Working responses vs. linear predictor plot should be linear and (some outliers aside) it is linear except that it has a slightly non-linear part at one of its ends. Partial residuals vs. predictor plots should be linear, and (some outliers aside) they are, all be it vertically linear for adult body mass, which may reflect that adult body mass is included on an inappropriate scale. All the other systemic and random components trialled with these data sets, which converged to a result, had these same issues.*

*Maximal lifespan is a negative convex function of specific basal metabolic rate, and a positive concave function of adult body mass, wherein specific basal metabolic rate is a negative convex function of adult body mass (Figure 14). To speculate, perhaps the adverse diagnostics for these GLMs might be (at least in part) because this pattern of non-linearities is too complicated for them to model better. At least with the systemic and random component options available in JASP software.*

*To some extent, a problem for the GLM to model might be the different degrees of non-linearity between maximal lifespan and adult body mass, and maximal lifespan and specific basal metabolic rate. Table S46 below shows the exponent (b) values from the fitted-to-data-equation:* $maximal\ lifespan = a \cdot x^b$*, for the different species sets A, B, and C, wherein one can see that the non-linearity for specific basal metabolic rate is less than that for adult body mass (because, absolutely, its values of b are closer to 1). These b values are as disclosed earlier herein.*

*[Table S46]*

| | *b* | |
|---|---|---|
| *Set* | *x = adult body mass* | *x = specific basal metabolic rate* |
| *A* | *0.1376* | *-0.3854* |
| *B* | *0.1659* | *-0.5248* |
| *C* | *0.1733* | *-0.5344* |

*Or perhaps these adverse diagnostics are merely to be expected when dealing with the imperfection and complexity of real-world biological data. And can be disregarded. Indeed, their non-idealness is arguably relatively minor.*

When the link function isn't Inverse (or $1/\mu^2$), for example when it is Log instead (e.g., using Gamma Log instead of Gamma Inverse), the sign of *T* is positive, and the sign of *U* is negative (not shown).

Table S47 below shows results from when the GLM models (each Gamma Inverse) are simplified, with specific basal metabolic rate as the only predictor:
[Table S47]

| Set | Q | S | U | S-p | U-p |
|---|---|---|---|---|---|
| A | 5.220e-24 | 0.041 | 3.913 | 2.822e-50 | 9.615e-20 |
| B | 9.133e-33 | 0.029 | 11.329 | 2.667e-17 | 2.245e-25 |
| C | 1.263e-50 | 0.019 | 15.168 | 1.357e-12 | 6.402e-39 |

In Table S47 above, specific basal metabolic rate is a highly significant predictor of maximal lifespan. Compared with earlier Table S45, *Q* is lower for species set *C*, and *U-p* is lower for all the species sets.

Generalized Linear Model (GLM) with standardized data

Using data from 350 mammal species (species set *B*), I constructed a Generalized Linear Model (GLM) in JASP software (version 0.18.1.0) with maximal lifespan (years) as the response variable, and *standardized* (to be between 0-100 by Eq. S1) adult body mass (kg) and *standardized* (to be between 0-100 by Eq. S1) specific basal metabolic rate (W/g) as predictors (all unlogged). This standardization means the coefficients are comparable, reporting which predictor, adult body mass or specific basal metabolic rate, has greater influence upon maximal lifespan. The answer being specific basal metabolic rate, its coefficient being nearly nineteen (19) times larger than that for adult body mass. I mandated that the response variable should be positive continuous, which restricted the random component options available in JASP to Gamma and Inverse Gaussian, with corresponding systemic component ("link") options of Identity, Log, Inverse, and $1/\mu^2$. I selected the combination which conferred the lowest chi-squared ($\chi^2$) p-value, which was random component = "Gamma" and systemic component = "Inverse".

Results shown in Table S48 below. Alongside results for species sets *A* and *C*, also retrieved with Gamma Inverse. Where *Q* is the model's chi-squared ($\chi^2$) p-value, *VIF* is Variance

Inflation Factor, *S* is the intercept coefficient (*S-p* is p-value thereof), *T* is the coefficient for standardized adult body mass (*T-p* is p-value thereof), and *U* is the coefficient for standardized specific basal metabolic rate (*U-p* is p-value thereof).

[Table S48]

| Set | Q | VIF | S | T | U | S-p | T-p | U-p |
|---|---|---|---|---|---|---|---|---|
| A | 6.890e-25 | 1.013 | 0.043 | -3.424e-4 | 0.002 | 1.446e-54 | 2.058e-6 | 1.021e-18 |
| B | 5.371e-33 | 1.084 | 0.033 | -2.654e-4 | 0.005 | 2.147e-21 | 6.212e-4 | 4.168e-24 |
| C | 4.240e-50 | 1.068 | 0.023 | -1.795e-4 | 0.006 | 6.896e-17 | 0.007 | 3.203e-37 |

In Table S48 above, the p-values for specific basal metabolic rate are highly significant, and smaller than for adult body mass. For species set *C*, the coefficient for specific basal metabolic rate is >33 times greater than for adult body mass. So, even with adult body mass included in the model, specific basal metabolic rate is a highly significant predictor of maximal lifespan.

*Diagnostic graphs for these GLMs: Quantile residuals vs. fitted (which is Quantile residuals vs. predicted values of the response variable) and Quantile residuals vs. predictor plots shouldn't contain pattern(s) and/or trend(s), but there are some outlier data points which makes the other data points appear bunched in the graphs, which might be misconstrued as regularity. That aside, there might be minor non-randomness in these plots. The Q-Q plot should be linear and, some outliers aside, is linear, although it has some slight non-linearity at its ends. Working responses vs. linear predictor plot should be linear and (some outliers aside) it is linear except that it has a slightly non-linear part at one of its ends. Partial residuals vs. predictor plots should be linear, and (some outliers aside) they are, all be it vertically linear for adult body mass, which may reflect that adult body mass is included on an inappropriate scale (even though standardized data is used in this case). All the other systemic and random components trialled with these data sets, which converged to a result, had these same issues.*

When the link function isn't Inverse (or $1/\mu^2$), for example when it is Log instead (e.g., using Gamma Log instead of Gamma Inverse), the sign of *T* is positive, and the sign of *U* is negative (not shown).

Diagnostic graphs for the Generalized Linear Model (GLM) of Eq. 12 in the main text

*Diagnostic graphs for this GLM: Quantile residuals vs. fitted (which is Quantile residuals vs. predicted values of the response variable) and Quantile residuals vs. predictor plots shouldn't contain pattern(s) and/or trend(s), but in at least one of these graphs (vs. adult body mass) there is at least one outlier data point which makes the other data points appear bunched, which might be misconstrued as regularity. The Q-Q plot should be linear and is linear but with some slight non-linearity at its ends (no outliers). Working responses vs. linear predictor plot should be linear and (an outlier aside) it is linear. Partial residuals vs. predictor plots should be linear, and (some outliers aside) they are, all be it vertically linear for adult body mass, which may reflect that adult body mass is included on an inappropriate scale.*

*ΔM is a linear function of specific basal metabolic rate, and a negative convex function of adult body mass, wherein specific basal metabolic rate is a negative convex function of adult body mass (Figure 14). To speculate, the adverse diagnostics for this GLM might be (at least in part) because this pattern of non-linearities is too complicated for it to model better. At least with the systemic and random component options available in JASP software.*

Table S49 below shows output from the GLM model (Gamma Identity) when adult body mass is omitted, and specific basal metabolic rate is the only predictor. Where *Q* is the model's chi-squared ($\chi^2$) p-value, *S* is the intercept coefficient (*S-p* is p-value thereof), and *U* is the coefficient for standardized specific basal metabolic rate (*U-p* is p-value thereof).

[Table S49]

| Q | S | U | S-p | U-p |
|---|---|---|---|---|
| 1.277e-12 | 0.081 | 30.219 | 8.119e-10 | 1.123e-10 |

In Table S49 above, specific basal metabolic rate is a highly significant predictor of *ΔM*. Comparing with the model of Eq. 12 in the main text, *Q*, *S-p*, and *U-p* are lower when specific basal metabolic rate is the sole predictor.

Generalized Linear Model (GLM) for *ΔM* with standardized data

With data (from 50 eutherian species) used to generate Figure 10 in the main text, I constructed a Generalized Linear Model (GLM) in JASP software (version 0.18.1.0) with *ΔM* as the response variable, and *standardized* (to be between 0-100 by Eq. S1) adult body mass (kg) and *standardized* (to be between 0-100 by Eq. S1) specific basal metabolic rate (W/g) as predictors (all unlogged). This standardization means the coefficients are comparable, reporting which predictor, adult body mass or specific basal metabolic rate, has greater influence upon *ΔM*. The answer being specific basal metabolic rate, its coefficient being nearly eight (8) times larger than that for adult body mass.

I mandated that the response variable should be positive continuous, which restricted the random component options available in JASP to Gamma and Inverse Gaussian, with corresponding systemic component ("link") options of Identity, Log, Inverse, and $1/\mu^2$. I selected the combination which conferred the lowest chi-squared ($\chi^2$) p-value, which was random component = "Inverse Gaussian" and systemic component = "Identity".

Results shown in Table S50 below. Where *Q* is the model's chi-squared ($\chi^2$) p-value, *VIF* is Variance Inflation Factor, *S* is the intercept coefficient (*S-p* is p-value thereof), *T* is the coefficient for standardized adult body mass (*T-p* is p-value thereof), and *U* is the coefficient for standardized specific basal metabolic rate (*U-p* is p-value thereof).
[Table S50]

| Q | VIF | S | T | U | S-p | T-p | U-p |
|---|---|---|---|---|---|---|---|
| 7.107e-11 | 1.173 | 0.098 | -5.257e-4 | 0.004 | 5.751e-13 | 6.450e-4 | 4.248e-8 |

Table S50 above reports that, at least across these eutherian species, specific basal metabolic rate is a more significant predictor of epigenetic clock ticking rate (*ΔM*) than adult body mass. Also observed with the other random and systemic component combinations trialled (not shown). So, even with adult body mass included in the model, specific basal metabolic rate is a significant predictor of *ΔM*.

If instead random component = "Gaussian" and systemic component = "Identity":
[Table S51]

| Q | VIF | S | T | U | S-p | T-p | U-p |
|---|---|---|---|---|---|---|---|
| 1.851e-8 | 1.085 | 0.021 | -4.634e-4 | 0.004 | 2.552e-5 | 0.594 | 1.446e-8 |

Table S51 above reports that, at least across these eutherian species, specific basal metabolic rate *is*, whereas adult body mass is *not*, a significant predictor of epigenetic clock ticking rate

($\Delta M$). This is also observed when the random component is "Gaussian", and the systemic component is instead "Log" or "Inverse", and when the random component is "Gamma" with systemic component of "Inverse". In Table S51 above, the coefficient for specific basal metabolic rate is nearly nine (9) times larger than that for adult body mass.

*Diagnostic graphs for these GLMs: Quantile residuals vs. fitted (which is Quantile residuals vs. predicted values of the response variable) and Quantile residuals vs. predictor plots shouldn't contain pattern(s) and/or trend(s), but in at least one of these graphs (vs. adult body mass) there is at least one outlier data point that makes the other data points appear bunched, which might be misconstrued as regularity. The Q-Q plot should be linear and is linear but with some slight non-linearity at its ends (no outliers). Working responses vs. linear predictor plot should be linear and (an outlier aside) it is linear. Partial residuals vs. predictor plots should be linear, and (some outliers aside) they are, all be it vertically linear for adult body mass, which may reflect that adult body mass is included on an inappropriate scale (even though standardized data is used in this case).*

Generalized Additive Model (GAM)

A GAM is an alternative to a GLM, which (like GLM) doesn't require linearity between the response variable and predictors. So, like a GLM, it too can be used with untransformed variables of adult body mass and specific basal metabolic rate, which are less collinear than their logged ($\log_{10}$) versions. There is an argument that GAMs are superior to GLMs.

Using data from [1], for earlier defined species sets *A* (517 mammal and bird species), *B* (350 mammal species), and *C* (263 mammal species; with bats, naked mole rat, and marsupials omitted), I built GAM models with maximal lifespan in years ($y$) as the response variable, with specific basal metabolic rate in W/g ($x$) and adult body mass in kg ($z$) as predictors. No variables logged. Implemented in R code, presented in the supplementary methods section of this Supplementary Material document, which utilizes the Mixed GAM Computation Vehicle (mgcv) R package, version 1.9-1 [334].

For each GAM: family was Gaussian, link was identity, and their output is shown in Table S52 below. Where $A.R^2$ is adjusted $R^2$, *GCV* is Generalized Cross Validation, *Dev* is deviance explained, *Int* is intercept, *Int.p* is the two-tailed p-value for the intercept, s($x$) and s($z$) are the smooth terms, *edf* is effective degrees of freedom (measure of complexity: edf of 1 is equivalent to a straight line, edf of 2 is equivalent to a quadratic curve, and so on), *ref.df* is reference degrees of freedom, *P* is the two-tailed p-value for a smooth term, *K* is the smoothing parameter (when large, the model might not fit the data well. When small, the model might be overfitted to the data, and poorly generalizable to other data. *K* may be considered large when appreciably larger than the *edf* value, and small when not much larger than the *edf* value), and *CC* is concurvity ("worst" values shown; range is between 0 and 1; a subjective rule of thumb is that >0.8 is of concern; high concurvity, analogously to high collinearity, reflects redundancy/entwinement of predictors; high concurvity can inflate smooth term p-values).

[Table S52]

| Set | A.$R^2$ | GCV | Dev | Int | Int.p | | edf | ref.df | P | K | CC |
|---|---|---|---|---|---|---|---|---|---|---|---|
| A | 0.286 | 106.49 | 30% | 16.020 | <2e-16 | s(*x*) | 1.990 | 2.541 | 1.66e-05 | 9 | 0.5871208 |
| | | | | | | s(*z*) | 7.895 | 8.620 | <2e-16 | 9 | 0.58712085 |
| B | 0.449 | 83.103 | 46.6% | 15.0583 | <2e-16 | s(*x*) | 4.139 | 5.048 | <2e-16 | 9 | 0.5634203 |
| | | | | | | s(*z*) | 7.087 | 7.993 | <2e-16 | 9 | 0.56342032 |
| C | 0.539 | 82.266 | 55.7% | 15.7506 | <2e-16 | s(*x*) | 3.833 | 4.683 | <2e-16 | 9 | 0.5355484 |
| | | | | | | s(*z*) | 6.872 | 7.793 | <2e-16 | 9 | 0.53554845 |

In Table S52 above, the p-values for the intercept and both the smooth terms, including that for specific basal metabolic rate, are highly significant. So, even with adult body mass included in the model, specific basal metabolic rate is a significant predictor of maximal lifespan.

*Diagnostic graphs for these GAMs:*

*Model-predicted maximal lifespan vs. observed maximal lifespan plots are linear, each with a single outlier, which is human (rationale disclosed in the Methods section may explain why human is anomalous). Dispersion of values around linearity is less (i.e., plot is more faithfully linear) as one moves from species set A to C.*

*Residuals vs. predicted (values of response variable) plot shouldn't contain pattern(s) and/or trend(s), but there are some outlier data points which makes the other data points appear bunched in the graph, which might be misconstrued as regularity. There might be some minor non-randomness further to that. The Q-Q plot should be linear, and it is except for slight non-linearity at its ends, with a few outliers observed at the ends also. Working responses vs. linear predictor plot should be linear and it is, with no outliers present. For a GAM, partial residuals vs. predictor plots should appear random and evenly distributed around zero, which (some outliers aside) is arguably the case, all be it this isn't ideally clear cut, which is a potential issue. The Component-Plus-Residual (CPR) plot is, as it should be, data points scattered around linear, all be it there are some outliers to this.*

*The diagnostic graphs are increasingly better as one moves from species set A to C.*

*R code to generate these diagnostic graphs is provided in the supplementary methods section of this Supplementary Material document such that the reader can see these graphs for themselves and make their own judgements. Generally, I judge that the diagnostics for these GAM models are better than for the corresponding GLM models, that used to generate the data of earlier Table S45. All be it, there is some non-idealness in the diagnostic graphs here as well.*

*Unlike with GLMs earlier, plots using quantile residuals (quantile residuals vs. fitted/predicted values of the response variable. Quantile residuals vs. each predictor plots) are not investigated for GAMs herein, because I judge that sufficient diagnostic coverage is conferred without them. Indeed, they aren't commonly reported diagnostics for a GAM.*

Incidentally, when human is omitted from species set *C*, deviance explained (*Dev*) increases from 55.7 to 68.4%, and adjusted $R^2$ ($A.R^2$) increases from 0.549 to 0.667, and the p-values for the intercept and smooth terms are each still <2e-16 (not shown).

Table S53 below shows results from when the GAM models are simplified, such that specific basal metabolic rate is the only predictor (*N.B., human is included*):

[Table S53]

| | $A.R^2$ | GCV | Dev | Int | Int.p | | edf | ref.df | P | K |
|---|---|---|---|---|---|---|---|---|---|---|
| A | 0.217 | 116.4 | 23% | 16.0203 | <2e-16 | s(*x*) | 8.374 | 8.892 | <2e-16 | 9 |

| | | | | | | | | | | |
|---|---|---|---|---|---|---|---|---|---|---|
| B | 0.352 | 96.633 | 36.7% | 15.0583 | <2e-16 | s(*x*) | 7.901 | 8.657 | <2e-16 | 9 |
| C | 0.437 | 97.439 | 45% | 15.7506 | <2e-16 | s(*x*) | 6.235 | 7.287 | <2e-16 | 9 |

In Table S53 above, in all models, specific basal metabolic rate is a *significant* predictor of maximal lifespan. If human is excluded, the value of *Dev* is instead 23.6%, 41.7%, and 53.6% for species sets *A*, *B*, and *C* respectively, whilst in each case *Int.p* and *P* are <2e-16 (not shown).

Generalized Additive Model (GAM) with body surface-area per unit body mass instead of adult body mass

I removed bats from earlier defined species set *B* (350 mammal species), rendering a 318 species set. Using this set, with adult body mass values sourced from [1], I calculated body surface-area per unit body mass values, using body surface-area values calculated from body mass by the Meeh equation, with parameter values $k = 0.1\ m^2/kg^{0.67}$ and $b = 0.67$ [128]. The reason why bats were removed is because these parameter values aren't well generalizable to winged, flying species [128].

Using this data, and data from [1], I built a GAM model with maximal lifespan in years (*y*) as the response variable, with specific basal metabolic rate in W/g (*x*) and body surface-area per unit body mass in $m^2$/kg (*z*) as predictors. With the caveat that the latter is populated with estimates by calculation rather than actual experimental data. No variables logged. Incidentally, a corresponding multiple linear regression model, using logged variables, had collinearity (*VIF* = 3.694), and so a GAM model is more appropriate, as it can use unlogged variables, which are less collinear.

Using the same R code as prior, but with different data, family was Gaussian, link was identity, and output is shown in Table S54 below, with column headings as defined for earlier Table S52.

[Table S54]

| A.$R^2$ | GCV | Dev | Int | Int.p | | edf | ref.df | P | K | CC |
|---|---|---|---|---|---|---|---|---|---|---|
| 0.482 | 82.064 | 50.1% | 14.7654 | <2e-16 | s(*x*) | 7.756 | 8.565 | 0.00132 | 9 | 0.8209448 |

| | | | | | | | | | | |
|---|---|---|---|---|---|---|---|---|---|---|
| | | | | | s(*z*) | 3.774 | 4.797 | 6.04e-07 | 9 | 0.820 9448 |

In Table S54 above, even with body surface-area per unit body mass included in the model, specific basal metabolic rate is a significant predictor of maximal lifespan.

Although a possible problem with this model is concurvity, which, analogously to collinearity, indicates redundancy (entwinement) of predictors, and it can inflate smooth term p-values. Especially notable here because it is observed at appreciable magnitude, not just with the "worst" values, but with the "estimate" values also, as shown in Table S55 below. Refer to the mgcv R package documentation for guidance on these concurvity related terms ("worst", "observed", "estimate") if needed.

[Table S55]

| | Worst | Observed | Estimate |
|---|---|---|---|
| s(x) | 0.8209448 | 0.4566687 | 0.5190020 |
| s(z) | 0.8209448 | 0.7602801 | 0.7065754 |

This concurvity is interesting. It is consistent with body surface-area per unit body mass dictating maximal lifespan by dictating specific basal metabolic rate.

*Diagnostic graphs for this GAM:*

*Model-predicted maximal lifespan vs. observed maximal lifespan plot is linear, but with a single far-removed outlier, which is human.*

*Residuals vs. predicted (values of response variable) plot shouldn't contain pattern(s) and/or trend(s) and this is the case, all be it there is a single far-removed outlier. The Q-Q plot should be linear, and it is except for non-linearity at its ends, minor at one end, and much more at the other, which trends towards an outlier. Working responses vs. linear predictor plot should be linear and it is, with no outliers present. For a GAM, partial residuals vs. predictor plots should appear random and evenly distributed around zero, which is arguably the case, all be it there are outliers arranged in vertical and horizontal lines emanating away from the bunched rest, which is a concern. The Component-Plus-Residual (CPR) plot is, as it should be, data points scattered around linear.*

Omitting *Mustela frenata*, the long-tailed weasel, which has a very long tail that makes its surface-area anomalously large for its body mass, doesn't increase *Dev* or $A.R^2$ (not shown).

<u>Generalized Additive Model (GAM) for *ΔM*</u>

With data (from 50 eutherian species) used to generate Figure 10 in the main text, I constructed a GAM with *ΔM* as the response variable (*y*), and specific basal metabolic rate in W/g (*x*) and adult body mass in kg (*z*) as predictors. No variables logged. Family was Gaussian, link was identity, and its output is shown in Table S56 below, with column headings as defined for Table S52 above.

[Table S56]

| $A.R^2$ | GCV | Dev | Int | Int.p | | edf | ref.df | P | K | CC |
|---|---|---|---|---|---|---|---|---|---|---|
| 0.597 | 0.0070479 | 66% | 0.2044 | <2e-16 | s(*x*) | 6.622 | 7.724 | 8.99e-07 | 9 | 0.9249104 |
| | | | | | s(*z*) | 1.000 | 1.000 | 0.603 | 9 | 0.9249104 |

In Table S56 above, the p-value for adult body mass is *not* significant and that for specific basal metabolic rate is *highly* significant. So, even with adult body mass included in the model, specific basal metabolic rate is a significant predictor of *ΔM*.

*Diagnostic graphs for this GAM: Model-predicted ΔM vs. observed ΔM plot is linear. Residuals vs. predicted (values of response variable) plot shouldn't contain pattern(s) and/or trend(s), but there are some outlier data points which makes the other data points appear slightly bunched in the graph, which might be misconstrued as regularity. There might be some very minor non-randomness further to that. The Q-Q plot should be linear, and it is except for slight non-linearity at its ends, no outliers beyond that. Working responses vs. linear predictor plot should be linear and it is, with no outliers present. The Component-Plus-Residual (CPR) plot is, as it should be, data points scattered around linear, all be it there are some outliers to this. Partial residuals vs. predictor plots, aside from some outliers, are evenly distributed around zero, but linearity is observed, especially for specific basal metabolic rate, which is a concern. Unlike with a GLM, where linearity is desirable in partial residuals vs. predictor plots, in a GAM such linearity is concerning. However, it is perhaps to be expected in the specific basal metabolic rate case because the relation between specific basal metabolic rate and ΔM is linear. Not non-linear. But the vertical linearity observed with adult body mass is concerning, because ΔM is a non-linear function of adult body mass. The GAM model may not be capturing the non-linear relationship between ΔM and adult body mass well. Possibly because the linear relation between specific basal metabolic rate and ΔM is so sufficiently explanatory, that adult body mass is sidelined in the GAM model. This might be why adult body mass is not a statistically significant predictor of ΔM in the model.*

*High concurvity indicates redundancy/entwinement of predictors and can inflate smooth term p-values. In Table S56 above the concurvity values are high (0.9249104), although these values are the "worst". The "observed" and "estimate" values are much lower, shown in Table S57 below. Refer to the mgcv R package documentation for guidance on these concurvity related terms ("worst", "observed", "estimate") if needed.*

*[Table S57]*

| | *Worst* | *Observed* | *Estimate* |
|---|---|---|---|
| *s(x)* | *0.9249104* | *0.4059324* | *0.2469378* |
| *s(z)* | *0.9249104* | *0.3719547* | *0.1976479* |

*The results above are what the R code, which leverages the mgcv R package, returns by default. I did some very limited, somewhat arbitrary manual manipulation of the K parameter (e.g., setting it to 2 or 20) for s(x) and/or s(z) but couldn't find any value(s) thereof that could make adult body mass a significant predictor of ΔM. In each case trialled, specific basal metabolic rate was returned as a significant predictor of ΔM. Results not shown.*

<u>Generalized Additive Model (GAM) for *ΔM* using body surface-area per unit body mass</u>

With data from the 50 eutherian species used to generate Figure 10 in the main text, minus data for the 6 winged/flying (bat) species, rendering 44 species remaining, including human, I constructed a GAM with *ΔM* as the response variable (*y*), and specific basal metabolic rate in W/g (*x*) and (*calculated from body mass values sourced from [1] by the Meeh equation, with* $k = 0.1\ m^2/kg^{0.67}$ *and* $b = 0.67$ *[128]*) body surface-area per unit body mass in $m^2/kg$ (*z*) as predictors. No variables logged. Family was Gaussian, link was identity, and its output is shown in Table S58 below, with column headings as defined for Table S52 above.

[Table S58]

| $A.R^2$ | GCV | Dev | Int | Int.p | | edf | ref.df | P | K | CC |
|---|---|---|---|---|---|---|---|---|---|---|

| | | | | | | | | | | |
|---|---|---|---|---|---|---|---|---|---|---|
| 0.751 | 0.0049688 | 79.9% | 0.210847 | <2e-16 | s(*x*) | 7.284 | 8.275 | <2e-16 | 9 | 0.8999365 |
| | | | | | s(*z*) | 1 | 1 | 0.511 | 9 | 0.8999365 |

In Table S58 above, the p-value for body surface-area per unit body mass is not significant and that for specific basal metabolic rate is *highly* significant.

*Diagnostic graphs for this GAM: Model-predicted $\Delta M$ vs. observed $\Delta M$ plot is linear. Residuals vs. predicted (values of response variable) plot shouldn't contain pattern(s) and/or trend(s), and this is the case. The Q-Q plot should be linear, and it is except for slight non-linearity at its ends, with a single outlier, which is especially removed. Working responses vs. linear predictor plot should be linear and it is, with no outliers present. The Component-Plus-Residual (CPR) plot is, as it should be, data points scattered around linear, all be it there are some outliers to this. Partial residuals vs. predictor plots are not evenly distributed around zero (although not too far removed from this), and (concerningly) linearity is observed, diagonally for specific basal metabolic rate, and vertically for body surface-area per unit body mass.*

*High concurvity indicates redundancy/entwinement of predictors and can inflate smooth term p-values. In Table S58 above, concurvity values are high (*0.8999365*). All be it these values are the "worst". The "observed" and "estimate" values are much lower, shown in Table S59 below.*

*[Table S59]*

| | *Worst* | *Observed* | *Estimate* |
|---|---|---|---|
| *s(x)* | *0.8999365* | *0.5997681* | *0.4992866* |
| *s(z)* | *0.8999365* | *0.4137256* | *0.2750409* |

Table S60 below shows GAM model output, where the response variable is *$\Delta M$*, and there is only a single predictor (*x*). On the first line (*A*), the single predictor is body surface-area per unit body mass. On the second line (*B*), the single predictor is specific basal metabolic rate.

[Table S60]

| | A.$R^2$ | GCV | Dev | Int | Int.p | | edf | ref.df | P | K |
|---|---|---|---|---|---|---|---|---|---|---|

| | | | | | | | | | | |
|---|---|---|---|---|---|---|---|---|---|---|
| A | 0.333 | 0.011643 | 38.4% | 0.21085 | <2e-16 | s($x$) | 3.28 | 3.82 | 0.000937 | 9 |
| B | 0.756 | 0.0047517 | 79.7% | 0.210847 | <2e-16 | s($x$) | 7.345 | 8.314 | <2e-16 | 9 |

Table S60 above reports that body surface-area per unit body mass (estimated using Meeh calculation), and specific basal metabolic rate, are significant predictors of epigenetic clock ticking rate ($\Delta M$). Specific basal metabolic rate more so.

*Diagnostic graphs for the GAM with body surface-area per unit body mass as sole predictor: Model-predicted $\Delta M$ vs. observed $\Delta M$ plot is linear (but with many outliers, many of which are arranged in orthogonal linearity). Residuals vs. predicted (values of response variable) plot shouldn't contain pattern(s) and/or trend(s), but there is a trend of vertical linearity. The Q-Q plot should be linear, and it is except for non-linearity at its ends, with more pronounced non-linearity at one end, which trends non-linearly towards a single extreme outlier. Working responses vs. linear predictor plot should be linear and it is, with no outliers present. Partial residuals vs. predictor plot should be evenly distributed around zero and it, some outliers aside, approximately is, all be it there are some outliers especially removed from the other data.*

*Diagnostic graphs for the GAM with specific basal metabolic rate as sole predictor: Model-predicted $\Delta M$ vs. observed $\Delta M$ plot is linear. Residuals vs. predicted (values of response variable) plot shouldn't contain pattern(s) and/or trend(s), and it doesn't. The Q-Q plot should be linear, and it is except for slight non-linearity at its ends, a single outlier present at one of the ends, which isn't too far removed. Working responses vs. linear predictor plot should be linear and it is, with no outliers present. Partial residuals vs. predictor plot should be evenly distributed around zero and it approximately is, but (perhaps concerningly) diagonal linearity is observed.*

<u>Multiple non-linear regression</u>

The following equation, Eq. S6, was fitted to data (sourced from [1]) for maximal lifespan in years ($y$), specific basal metabolic rate in W/g ($x$), and adult body mass in kg ($z$), all variables unlogged, independently for each of species sets $A$ (517 mammal and bird species), $B$ (350 mammal species), and *C* (263 mammal species; with bats, naked mole rat, and marsupials omitted), using R code shown in the supplementary methods section of this Supplementary Material document, which utilizes the Non-Linear Regression with Brute Force (nls2) R package, version 0.3-3 [335]. [Eq. S6]

$$y = (B \cdot x^{C}) + (D \cdot z^{E})$$

Results are shown in Table S61 below, where *p.B*, *p.C*, *p.D*, and *p.E* are two-tailed p-values for fitted parameters *B*, *C*, *D*, and *E* respectively.

[Table S61]

| Set | B | C | D | E | *p.B* | *p.C* | *p.D* | *p.E* |
|---|---|---|---|---|---|---|---|---|
| A | 4 | -0.1 | 10 | 0.2 | *0.0734* | *0.3513* | *0.0179* | *6.78e-05* |
| B | 1 | -0.4 | 4 | 0.3 | *0.0219* | *1.24e-10* | *0.0418* | *6.89e-06* |
| C | 1 | -0.4 | 4 | 0.3 | *0.0461* | *5.17e-9* | *0.0627* | *3.49e-5* |

Shown in Table S61 above, for species set *B*, all the parameters are significant at two-tailed $p < 0.05$. And for species set *C*, all the parameters are significant except for parameter *D*, which relates to the adult body mass equation component. The fitted parameters are the same for species sets *B* and *C*.

The fitted parameter values are highly dependent on the parameter space brute-forced, and the specified step-size in each parameter, which was the same for each of species sets *A*, *B*, and *C*. I did these optimizations on a laptop, so didn't have much compute available, and did a very delimited search with relatively large step-sizes specified. It is possible that better parameter values could be obtained if a broader parameter space was searched and/or with smaller parameter step-sizes specified. It might be that a different equation form can fit the data better.

Comparing maximal lifespan values predicted by Eq. S6 (with parameters optimized for species set *C*) with maximal lifespan values for species set *C*: Pearson correlation coefficient = 0.7239, one-tailed (alternative hypothesis is directional) p-value = 2.8408E-44. Comparing maximal lifespan values predicted by Eq. S6 (with parameters optimized for species set *C*) with maximal lifespan values for species set *A* (so testing the model's predictive power for a bigger and more varied species set than its parameters were optimized upon): Pearson = 0.5505, one-tailed p-value = 1.3442E-42.

For each species set, the model-predicted maximal lifespan vs. observed maximal lifespan plot is linear. But with a single outlier, which is human (rationale disclosed in the Methods section may explain why human is anomalous). Dispersion of values around linearity is less (i.e., plot is more faithfully linear) as one moves from species set *A* to *C*.

<u>Generalized Linear Model (GLM) with phylogenetically independent contrasts</u>

The Phylogenetically Independent Contrasts (PIC) method requires interlinear variables, which in this case could be, and was, conferred by logging ($\log_{10}$) each of the inputted variables. And so (as observed earlier) its outputted contrasts inherit the collinearity problem, all be it somewhat attenuated by the PIC method. So, no remedy was expected in using these contrasts with GLM rather than multiple linear regression. And, indeed, no remedy was observed. GLM and GAM conferred remedy earlier, to the collinearity problem, by being useable with unlogged (non-interlinear) variables, which doesn't apply here because the variables were all (as they needed to be) logged prior to the PIC step.

In JASP (version 0.18.3.0): PIC contrasts, generated (as earlier) with the same branch length transformation (*all branch lengths set to 1.0 for species set A, all branch lengths being Nee transformed for species set B. Species sets A and B are as used earlier for phylogeny related study of predictors of maximal lifespan*), were used with GLM (forced through the origin [323, 325]), it using Gaussian Identity (the only combination of one of Gamma, Inverse Gaussian, or Gaussian, with one of Identity, Log, Inverse, or $1/\mu^2$ that converged), and results are in Tables S62 and S63 below, with column headings as used when presenting GLM results earlier (all be it there is no intercept here). With maximal lifespan contrasts as the response variable, Table S62 reports results when specific basal metabolic rate and adult body mass contrasts are predictors, and Table S63 reports results when basal metabolic rate and adult body mass contrasts are predictors. Coefficients of erroneous sign are written in red.

[Table S62]

| Set | Q | VIF | T | U | T-p | U-p |
|---|---|---|---|---|---|---|
| A | 2.501e-28 | 2.367 | 0.116 | -0.138 | 7.982e-9 | 0.017 |
| B | 8.671e-17 | 2.141 | 0.101 | -0.107 | 2.252e-6 | 0.078 |

[Table S63]

| Set | Q | VIF | T | U | T-p | U-p |
|---|---|---|---|---|---|---|
| A | 4.240e-27 | 7.509 | 0.155 | -0.005 | 1.366e-5 | 0.912 |
| B | 3.825e-16 | 6.899 | 0.141 | -0.019 | 2.289e-4 | 0.713 |

The used GLM here, with random component = "Gaussian" and systemic component = "Identity", is essentially just typical multiple linear regression, and so the results of Tables S62 and S63 above correspond with earlier Tables S31 and S32, all be it the former show unstandardized and the latter show standardized coefficients.

*Diagnostic graphs for these GLMs: Quantile residuals vs. fitted (which is Quantile residuals vs. predicted values of the response variable) and Quantile residuals vs. predictor plots shouldn't contain pattern(s) and/or trend(s), and they don't. The Q-Q plot should be linear, and it is, although with slight non-linearity at its ends. Working responses vs. linear predictor plot should be linear and it is linear all be it not tightly so and there is a broad width of dispersion. Partial residuals vs. predictor plots should be linear and are, all be it horizontally linear for basal metabolic rate contrasts, and slightly negatively linear for specific basal metabolic rate contrasts.*

<u>Does maximal lifespan significantly (inversely) correlate with specific basal metabolic rate across mammal and bird species when the effects of adult body mass and phylogeny are statistically excluded?</u>

Both Phylogenetically Independent Contrasts (PIC) and multiple linear regression methods require interlinear variables, which in this case was conferred by logging ($\log_{10}$) each of maximal lifespan, specific basal metabolic rate, and adult body mass. As shown herein, PIC reports that maximal lifespan significantly (inversely) correlates with specific basal metabolic rate across mammal and bird species when the effect of phylogeny is statistically excluded. But when statistically excluding the effect of adult body mass, atop or not of excluding phylogeny, using partial and/or semi-partial correlation analysis, which interrelate with multiple linear regression, high collinearity between specific basal metabolic rate and adult body mass (or phylogenetically independent contrasts thereof) corrupts the results to an extent proportional to the *VIF* values. Incidentally, most of these results *do* report that maximal lifespan significantly (inversely) correlates with specific basal metabolic rate when the effect of adult body mass is statistically excluded, including when the effect of phylogeny is simultaneously excluded. Ridge regression results, which are less susceptible (but not impervious) to collinearity, are supportive also. Generalized Linear Model (GLM), Generalized Additive Model (GAM), and Multiple Non-Linear Regression (MNLR) methods don't require inter-linearity and so can use untransformed variables, which are less collinear, and these methods report that specific basal metabolic rate is a highly significant explanatory variable of maximal lifespan, even when adult body mass is accounted for (*all be it equivocally with MNLR as full exploration couldn't be conducted because of computational intensiveness*). PIC requires interlinear (so in this case logged) variables input, and so the contrasts it outputs inherit the collinearity problem, and so there is no remedy in using them with one or more of GLM, GAM, and MNLR.

I think in totality, combining the results herein, sourced from different methods and sub-methods, it can be concluded that maximal lifespan significantly (inversely) correlates with specific basal metabolic rate across mammal and bird species when the effects of adult body mass and phylogeny are statistically excluded.

Moreover, refer to new variable *λ* herein, which specific basal metabolic rate is componentry to, for which it is unequivocally shown later that maximal lifespan inversely correlates with it, even when the effects of adult body mass and phylogeny are both, simultaneously, statistically excluded.

Correlations basis to Figure 14

At least some of the following correlations have been shown earlier in this present paper (its main section and/or this Supplementary Material), but they are presented again below, to disclose all these correlations in a single place, in some cases shown below with different data than used prior (demonstrating robustness, and to try to use less data sources to speed this workflow).

For Table S64 below, all the data used is from mammal species, wherein *N* is the number of species used (constrained in each case by the number of species that overlap between the compared data sets), *R* is the Pearson correlation coefficient, Rho (*ρ*) is the Spearman's rank correlation coefficient, *P* is the one-tailed (alternative hypothesis is directional) p-value, *A* is adult body mass (data from [1]), *B* is body surface-area per unit body mass (data from [128]), *C* is specific IF1 protein activity (data from [91]), *D* is specific $F_1F_0$ ATP hydrolysis rate (data from [91]), *E* is specific basal metabolic rate (data from [1], and as specified later in this paragraph), *F* is heart rate (data from [91]), *G* is specific Reactive Oxygen Species (ROS) generation rate (mean of superoxide and hydrogen peroxide data, for species for which both are provided, from [95]), *H* is specific DNA damage rate (total of SBS1, SBSB, and SBSC substitution rates, data from [100]), *I* is specific mtDNA damage rate (data from [100]), *J* is (inferred, as explained in the main text) specific protein damage rate (data from [102]), *K* is specific lipid damage rate (data from [101]), *L* is specific epigenetic damage rate (data calculated for eutherian species probed by the mammalian methylation array [127], curtailed to those species thereof which have maximal lifespan data in [1]), and *M* is maximal lifespan (data from [1]).

Where "specific" refers to per unit mass or otherwise controlled for body mass. If human data was present, it wasn't omitted when calculating any of these correlations.

From [1], only adult body mass, maximal lifespan, and specific basal metabolic rate data for the 350 mammal species with data for *all* these three variables was sourced. Although, adult body mass and maximal lifespan data was sourced atop from [1] for *Felis catus*, *Ovis aries*, *Canis lupus familiaris*, wherein specific basal metabolic rate data for these species, absent in [1], was sourced elsewhere: for cat from [289], and for sheep {*Ovis aries*} and dog {*Canis lupus familiaris*} as given in earlier Table S2, with sources disclosed in its corresponding text.

Variable *K* is (actually) amount of damage accumulated after a period, not rate of damage. But it is abstractly compared to other variables as if it were a rate of damage. Because rate=amount/period, where period is the same for all its data, from different species, and so cancels out in comparison with other variable(s). Although the period isn't the same chronologically, it is instead *approximately* the same percentage (15-30% [101]) of maximal lifespan in each species case. So, an abstractive drag on accuracy.

The resultant data set comprised data from 384 mammal species. I curtailed this to the 353 mammal species that had specific basal metabolic rate data. So, Figure 14 schematizes correlation data from 353 mammal species. All be it not all these 353 mammal species have data for all the variables, which is clear in the *N* values in Table S64 below.

[Table S64]

| Pair | N | R | P | Rho (ρ*)* | P |
|---|---|---|---|---|---|
| A-B | 27 | -0.2034 | 0.15444 | -0.9573 | 2.76069E-15 |
| A-B-*2* | 26 | -0.2859 | 0.078407 | -0.9603 | 4.12791E-15 |
| A-C | 10 | 0.2178 | 0.2727635 | 0.4909 | 0.074832 |
| A-D | 10 | -0.3221 | 0.1820345 | -0.697 | 0.012544 |
| A-E | 353 | -0.1184 | 0.0130585 | -0.8891 | 1.7713E-121 |
| A-F | 10 | -0.4345 | 0.104779 | -0.9515 | 0.0000115 |
| A-G | 7 | -0.6537 | 0.0556335 | -0.9286 | 0.0012585 |
| A-H | 11 | -0.1852 | 0.2928155 | -0.4182 | 0.1002745 |
| A-I | 11 | -0.2407 | 0.2379335 | -0.4091 | 0.105767 |
| A-J | 10 | -0.4475 | 0.0973475 | -0.5758 | 0.0407625 |
| A-K | 7 | -0.2935 | 0.2614595 | -0.75 | 0.0260905 |
| A-L | 53 | -0.2387 | 0.042598 | -0.7002 | 2.682E-09 |
| A-M | 353 | 0.3818 | 5.38715E-14 | 0.6736 | 2.57715E-48 |
| B-C | 8 | -0.6967 | 0.027424 | -0.4286 | 0.144684 |
| B-D | 8 | 0.7416 | 0.017603 | 0.619 | 0.0508835 |
| B-E | 27 | 0.5149 | 0.002996 | 0.7527 | 0.000003 |
| B-E-*2* | 26 | 0.7987 | 5.0128E-07 | 0.734 | 0.00001 |
| B-E-*3* | 26 | 0.8161 | 1.86762E-07 | 0.734 | 0.00001 |
| B-F | 8 | 0.9431 | 0.0002205 | 0.9762 | 0.0000165 |
| B-F-*2* | 8 | 0.9768 | 0.0000155 | 0.9762 | 0.0000165 |
| B-G | 6 | 0.9141 | 0.0053755 | 0.9429 | 0.002399 |
| B-H | 7 | 0.9138 | 0.001999 | 0.9286 | 0.0012585 |
| B-I | 7 | 0.9385 | 0.0008715 | 0.8929 | 0.0034005 |
| B-J | 5 | 0.8139 | 0.046818 | 0.8 | 0.1 |
| B-K | 8 | 0.9608 | 0.000073 | 0.619 | 0.0508835 |
| B-L | 20 | 0.8608 | 5.55773E-07 | 0.7292 | 0.000132 |
| B-M | 27 | -0.4377 | 0.011206 | -0.8127 | 1.30408E-07 |

| | | | | | |
|---|---|---|---|---|---|
| B-M-*2* | 26 | -0.6265 | 0.000308 | -0.8142 | 2.09129E-07 |
| C-D | 10 | -0.8182 | 0.0019065 | -0.6727 | 0.0165255 |
| C-E | 10 | -0.715 | 0.0100585 | -0.4061 | 0.122116 |
| C-F | 10 | -0.7849 | 0.003579 | -0.4667 | 0.086952 |
| C-G | 7 | -0.7413 | 0.0282755 | -0.5714 | 0.0901145 |
| C-H | 6 | -0.8396 | 0.0182645 | -0.4286 | 0.198233 |
| C-I | 6 | -0.8718 | 0.0117995 | -0.4286 | 0.198233 |
| C-J | 6 | -0.8972 | 0.0076545 | -0.8857 | 0.009425 |
| C-K | 7 | -0.8046 | 0.014544 | -0.5 | 0.126585 |
| C-L | 10 | -0.7669 | 0.0048215 | -0.4061 | 0.122116 |
| C-M | 10 | 0.2615 | 0.2327535 | 0.4424 | 0.100225 |
| D-E | 10 | 0.6414 | 0.022814 | 0.7091 | 0.0108315 |
| D-E-*2* | 10 | 0.7465 | 0.0065655 | 0.7091 | 0.0108315 |
| D-F | 10 | 0.8011 | 0.0026725 | 0.6727 | 0.0165255 |
| D-G | 7 | 0.9238 | 0.001477 | 0.7857 | 0.018122 |
| D-H | 6 | 0.9155 | 0.0052045 | 0.4286 | 0.198233 |
| D-I | 6 | 0.906 | 0.0064195 | 0.4286 | 0.198233 |
| D-J | 6 | 0.9465 | 0.0021085 | 0.9429 | 0.002399 |
| D-K | 7 | 0.8122 | 0.0132285 | 0.9286 | 0.0012585 |
| D-L | 10 | 0.8324 | 0.0014025 | 0.7939 | 0.003052 |
| D-M | 10 | -0.4241 | 0.110953 | -0.8424 | 0.0011105 |
| E-F | 10 | 0.9422 | 0.000023 | 0.9515 | 0.0000115 |
| E-F-*2* | 10 | 0.9852 | 1.03093E-07 | 0.9515 | 0.0000115 |
| E-G | 7 | 0.7198 | 0.0340865 | 0.7143 | 0.0356675 |
| E-H | 11 | 0.87 | 0.000249 | 0.5364 | 0.044464 |
| E-I | 11 | 0.8778 | 0.0001905 | 0.4909 | 0.062606 |
| E-J | 10 | 0.7394 | 0.007264 | 0.8424 | 0.0011105 |
| E-K | 7 | 0.958 | 0.0003395 | 0.75 | 0.0260905 |
| E-L | 53 | 0.6857 | 7.32024E-09 | 0.7711 | 7.1315E-12 |
| E-L-*2* | 52 | 0.7596 | 3.32706E-11 | 0.7578 | 3.91602E-11 |
| E-M | 353 | -0.4183 | 1.09821E-16 | -0.6418 | 1.14365E-42 |
| F-G | 7 | 0.8314 | 0.0102135 | 0.8214 | 0.011729 |

| | | | | | |
|---|---|---|---|---|---|
| F-H | 6 | 0.9595 | 0.0012135 | 0.8857 | 0.009425 |
| F-I | 6 | 0.9592 | 0.0012315 | 0.8857 | 0.009425 |
| F-J | 6 | 0.9065 | 0.0063525 | 0.7714 | 0.036207 |
| F-K | 7 | 0.9291 | 0.001237 | 0.75 | 0.0260905 |
| F-L | 10 | 0.9341 | 0.000038 | 0.8182 | 0.0019065 |
| F-M | 10 | -0.5012 | 0.0700045 | -0.8303 | 0.00147 |
| G-H | 4 | 0.9541 | 0.02295 | 1 | 0 |
| G-I | 4 | 0.9626 | 0.0187 | 1 | 0 |
| G-J | 5 | 0.9069 | 0.01681 | 0.7 | 0.09406 |
| G-K | 6 | 0.874 | 0.011407 | 0.7714 | 0.036207 |
| G-L | 7 | 0.8267 | 0.010911 | 0.6786 | 0.0468655 |
| G-M | 7 | -0.8436 | 0.0085235 | -0.75 | 0.0260905 |
| H-I | 14 | 0.9322 | 6.05138E-07 | 0.7846 | 0.000445 |
| H-J | 5 | 0.9422 | 0.008268 | 0.6 | 0.1423785 |
| H-K | 5 | 0.9429 | 0.008119 | 0.7 | 0.09406 |
| H-L | 15 | 0.926 | 3.66809E-07 | 0.8357 | 0.0000525 |
| H-M | 11 | -0.57 | 0.0335675 | -0.7727 | 0.002651 |
| I-J | 5 | 0.9482 | 0.007021 | 0.8 | 0.052044 |
| I-K | 5 | 0.973 | 0.002652 | 0.7 | 0.09406 |
| I-L | 14 | 0.8719 | 0.000024 | 0.8242 | 0.000144 |
| I-M | 11 | -0.4846 | 0.06544 | -0.7273 | 0.0056 |
| J-K | 4 | 0.7644 | 0.1178 | 0.8 | 0.1 |
| J-L | 12 | 0.9326 | 0.000005 | 0.7972 | 0.00095 |
| J-M | 10 | -0.5069 | 0.0674165 | -0.8061 | 0.002429 |
| K-L | 8 | 0.8336 | 0.0050645 | 0.6905 | 0.028989 |
| K-M | 7 | -0.8302 | 0.010389 | -0.8571 | 0.0068535 |
| L-M | 53 | -0.6573 | 4.48189E-08 | -0.9989 | 9.35751E-70 |

The asymptotically exact harmonic mean p-value [76] (*its post-published corrected method, as corrected on 7 October 2019*) for the p-values in Table S64 above, when selecting from each line whichever is the smaller p-value (for Pearson or Spearman's rank correlation coefficient), and where there are multiple lines for a pair only using the first line (*e.g., for B-E, B-E-2, and B-E-3: only using B-E*), and substituting the two p-values of value zero with an arbitrarily small value (1e-100) to make it calculable = 1.381614e-119; if the two p-values of zero are omitted instead = 1.346188e-119 = **$1*10^{-119}$**.

When graphed, all the relations in Table S64 are linear (positive or negative) except those where one of the pair is *A* or *M* (including A-M), which are supra-linear (convex shape if positive relation, concave shape if negative relation).

For all the correlations against *A*, and for all the correlations against *M* (except G-M): their Spearman's rank correlation coefficient is greater their Pearson correlation coefficient because these relations are *not* linear, but instead supra-linear: if a positive correlation, convex, and if a negative correlation, concave. Why is G-M (i.e., a correlation against *M*) different? Well, by sight, I interpret it as being supra-linear too (concave), my interpreting that there are some outliers, compounded by a very low number of data points (7), and further compounded by a disproportionate number of these data points being in the relatively linear section of the supra-linear plot.

A-B-*2*, B-E-*2*, and B-M-*2* use the same data as A-B, B-E, and B-M respectively except that outlier species *Mustela frenata* (the long-tailed weasel, which has a very long tail that makes its surface-area anomalously large for its body mass) is omitted. B-E-*3*, D-E-*2*, and E-F-*2* differ from B-E-*2*, D-E, and E-F respectively in using, for *Mus musculus*, a specific basal metabolic rate value of 0.010858546 W/g, which is a mean of values in the literature (from [1], [284], [179]) as specified earlier, in Table S3 and its accompanying text, instead of 0.015055556 W/g from [1]. B-F-*2* differs from B-F in using a heart rate value for rat (*Rattus norvegicus*) of 396.6 bpm [331] instead of 481 bpm [91]. E-L-*2* uses the same data as E-L except that outlier species *Sorex araneus* (Eurasian shrew) is omitted (as aforementioned, shrews become hyperactive when postabsorptive [129], and so they don't have a specific basal metabolic rate value).

A correlation's Spearman's rank correlation coefficient can be greater than its Pearson correlation coefficient if the correlation is supra-linear and/or when there are one or more outliers. Because of one or more outliers for each of B-E, D-E, E-F, E-L, their Spearman's rank correlation coefficient is greater than their Pearson correlation coefficient, which is remedied by the (aforementioned) outlier treatment conferring B-E-*2* (and B-E-*3*), D-E-*2*, E-F-*2*, and E-L-*2* respectively, which each have a greater Pearson than Spearman's rank correlation coefficient. To my eye and reasoning, B-G, B-H, D-K, E-J, G-H, G-I are linear relations too (indeed, the Pearson of 4 of these 6 pairs is >0.9 and that for G-I is 0.99) but outlier(s), compounded by a low number of data points, make each have a greater Spearman's rank than Pearson correlation coefficient.

All the correlations are *significant* (at $P < 0.05$, even despite very low values of $N$ in many cases) with Pearson and/or Spearman's rank correlation coefficients, *except* A-C, A-H, A-I, C-M, and J-K, which all have sizeable correlations (all >|0.4|), but which all have small values of $N$. Arguably underpowered. E.g., J-K has a Pearson correlation coefficient of 0.7644, which doesn't confer statistical significance as its $N$ is very low (=4).

As aforementioned, out of the data set comprising data from 384 mammal species, I curtailed to using data from the 353 mammal species that had specific basal metabolic rate data. This was to omit 29 species from the data set that had data for one or more of variables *H*, *I*, *J*, *K*, *L*, but for which I didn't have specific basal metabolic rate data (atop, I didn't have adult body mass and/or maximal lifespan data for some of these). To revisit correlations that didn't reach significance in Table S64 above with this extra species data: Table S65 below shows A-H and A-I if this extra data is also used. J-K isn't shown because there is no overlap of extra species for these variables, and so the correlation is the same.

[Table S65]

| Pair | N | R | P | Rho (ρ) | P |
|---|---|---|---|---|---|
| A-H | 15 | -0.2884 | 0.1486065 | -0.4179 | 0.1261 |
| A-I | 14 | -0.2633 | 0.1715225 | -0.5121 | 0.025494 |

In Table S65 above, A-H doesn't, but A-I does, reach significance. Whereas A-O and A-Q both reach significance, wherein *O* is different specific DNA damage data (data from [96]), and *Q* is different specific mtDNA damage data (data from [99], for this comparison an additional [to the 353 species] adult body mass value was sourced from [1], that of *Equus caballus*):

[Table S66]

| Pair | N | R | P | Rho (ρ) | P |
|---|---|---|---|---|---|
| A-O | 6 | -0.7143 | 0.0553885 | -0.9429 | 0.002399 |
| A-Q | 8 | -0.6622 | 0.0368 | -0.9048 | 0.001003 |

Variable *D* divided by variable *C* correlates with variable *A* with Spearman's rank correlation coefficient of -0.6727, and with Variable *M* with value of -0.7091, with one-tailed (alternative hypothesis is directional) p-values of 0.0165255 and 0.0108315 respectively.

Incidentally, the correlation between maximal lifespan and adult body mass for *all* the mammal species that have data for both these variables in [1]: *N* = 1023, Pearson = 0.5227 (one-tailed [alternative hypothesis is directional] p-value = 4.22874E-73), Spearman = 0.6958 (one-tailed p-value = 3.1744E-149). Spearman is greater than Pearson because this is a supra-linear (positive concave) relation.

Parameter values for Eq. 16

The data used to generate Table S64 in the last section was used to generate the following Table S67. No omission of outliers here. In Table S67 below, *N* is number of data points, Adj. $R^2$ is adjusted $R^2$, for rows *A*, *B*, *C*, *D*, *E*, *F*, *G*, *H*, *I*, *J, K*: *x* is respectively: body surface-area/mass, specific IF1 protein activity, specific $F_1F_0$ ATP hydrolysis rate, specific basal metabolic rate, heart rate, specific Reactive Oxygen Species (ROS) generation rate, specific DNA damage rate, specific mtDNA damage rate, specific protein damage rate, specific lipid damage rate, and specific epigenetic damage rate. Parameters determined using OriginPro 8 software (OriginLab Corporation, Northampton, MA, USA. Using its "Fit Linear" functionality, which uses linear regression, with least squares estimation).

[Table S67]

| *x* | N | Adj. $R^2$ | $\log_{10}(a)$ | *b* | N | Adj. $R^2$ | $\log_{10}(c)$ | *d* |
|---|---|---|---|---|---|---|---|---|
| A | 27 | 0.83313 | -0.90609 ±0.03982 | -0.3392 ±0.02966 | 27 | 0.53967 | 0.60207 ±0.10198 | -0.49456 ±0.08814 |
| B | 10 | 0.46745 | 0.23552 ±0.08802 | 0.16457 ±0.05516 | 10 | 0.51726 | 0.79069 ±0.15354 | 1.04804 ±0.32124 |
| C | 10 | 0.57518 | 0.31533 ±0.05357 | -0.12191 ±0.03357 | 10 | 0.50746 | 1.5094 ±0.15258 | -1.52632 ±0.47622 |
| D | 353 | 0.73962 | -2.53543 ±0.01006 | -0.25869 ±0.00818 | 353 | 0.42421 | -0.36066 ±0.08941 | -0.57906 ±0.03589 |
| E | 10 | 0.96015 | 2.38548 ±0.02845 | -0.26312 ±0.01783 | 10 | 0.70454 | 3.37834 ±0.47582 | -1.00822 ±0.21274 |

| | | | | | | | | |
|---|---|---|---|---|---|---|---|---|
| F | 7 | 0.89307 | -0.10708 ±0.04368 | -0.20096 ±0.02811 | 7 | 0.63309 | 0.81646 ±0.09272 | -0.86645 ±0.25715 |
| G | 11 | 0.0607 | 2.37542 ±0.1229 | -0.09677 ±0.07542 | 11 | 0.80324 | 3.732 ±0.37776 | -1.04951 ±0.16228 |
| H | 11 | 0.19152 | -1.4181 ±0.10781 | -0.12143 ±0.06616 | 11 | 0.64607 | -0.20969 ±0.35638 | -1.01009 ±0.23019 |
| I | 10 | 0.30069 | -0.54394 ±0.02263 | -0.03532 ±0.016 | 10 | 0.65701 | -1.44854 ±0.60508 | -4.77243 ±1.11744 |
| J | 7 | 0.58896 | 1.68192 ±0.06034 | -0.11181 ±0.03609 | 7 | 0.76624 | 3.27601 ±0.49018 | -1.37642 ±0.30277 |
| K | 53 | 1 | 0 ±0 | 1 ±0 | 53 | 0.99353 | 0.12656 ±0.01289 | -1.43966 ±0.01611 |

In Table S68 below, for its rows *L* and *M*: *x* is respectively specific DNA damage (data from [96]) and specific mtDNA damage (data from [99], with an additional adult body mass value sourced from [1], that of *Equus caballus*):

[Table S68]

| *x* | N | Adj. $R^2$ | $\log_{10}(a)$ | *b* | N | Adj. $R^2$ | $\log_{10}(c)$ | *d* |
|---|---|---|---|---|---|---|---|---|
| L | 6 | 0.85415 | 1.4491 ±0.06376 | -0.24213 ±0.044 | 6 | 0.81676 | 3.04711 ±0.39431 | -1.397 ±0.2895 |
| M | 8 | 0.63455 | 0.67921 ±0.03745 | -0.07578 ±0.02089 | 8 | 0.54339 | 2.42751 ±0.42839 | -2.0889 ±0.68386 |

Using the same data, and A-K letter code, as Table S67 but with the exception that in Table S69 below, A-D has outlier *Mustela frenata* omitted from A, and D-K has outlier *Sorex araneus* omitted from D (why these are outliers is as aforementioned):

[Table S69]

| *x* | *q* | N | Adj. $R^2$ | *e* | *g* |
|---|---|---|---|---|---|
| A | B | 8 | 0.39968 | 0.32089 ±0.09091 | -0.06515 ±0.02739 |
| A | C | 8 | 0.47499 | -0.08329 ±0.08512 | 0.1175 ±0.04339 |

| | | | | | |
|---|---|---|---|---|---|
| A | D | 26 | 0.62276 | 2.47564E-4 ±0.02507 | 34.07725 ±5.24133 |
| A | E | 8 | 0.87105 | -0.02054 ±0.02762 | 6.27102E-4 ±9.02467E-5 |
| A | F | 6 | 0.79451 | -0.07454 ±0.05831 | 0.31031 ±0.06882 |
| A | G | 7 | 0.80209 | -0.06169 ±0.04405 | 5.1783E-4 ±1.02917E-4 |
| A | H | 7 | 0.85704 | -0.0271 ±0.03239 | 2.87026 ±0.47206 |
| A | I | 5 | 0.54995 | -0.43574 ±0.25296 | 2.04559 ±0.84302 |
| A | J | 8 | 0.9104 | -0.12173 ±0.03243 | 0.00558 ±6.57003E-4 |
| A | K | 20 | 0.72661 | -0.05539 ±0.02095 | 0.76988 ±0.10728 |
| B | C | 10 | 0.6282 | 5.07247 ±0.63891 | -1.17747 ±0.29248 |
| B | D | 10 | 0.45018 | 3.79165 ±0.49763 | -245.48638 ±84.8579 |
| B | E | 10 | 0.56815 | 4.154 ±0.49529 | -0.00587 ±0.00164 |
| B | F | 7 | 0.45939 | 4.61882 ±0.98741 | -2.4533 ±0.99343 |
| B | G | 6 | 0.63112 | 4.64594 ±0.72913 | -0.00495 ±0.0016 |
| B | H | 6 | 0.70009 | 4.4157 ±0.59216 | -28.67818 ±8.05637 |
| B | I | 6 | 0.75612 | 0.39228 ±0.02349 | -0.03925 ±0.00966 |
| B | J | 7 | 0.57679 | 5.10226 ±0.83256 | -0.04886 ±0.01613 |
| B | K | 10 | 0.53668 | 4.60555 | -7.43106 |

| | | | | | |
|---|---|---|---|---|---|
| | | | | ±0.62969 | ±2.19848 |
| C | D | 10 | 0.33778 | 1.31478<br>±0.37952 | 153.01768<br>±64.71633 |
| C | E | 10 | 0.59702 | 0.97147<br>±0.33248 | 0.00416<br>±0.0011 |
| C | F | 7 | 0.82403 | 0.4995<br>±0.3692 | 2.00368<br>±0.37145 |
| C | G | 6 | 0.79767 | 0.76001<br>±0.27147 | 0.00272<br>±5.96798E-4 |
| C | H | 6 | 0.7761 | 0.92677<br>±0.25722 | 14.98289<br>±3.49947 |
| C | I | 6 | 0.86976 | -2.23216<br>±0.82035 | 15.15478<br>±2.58421 |
| C | J | 7 | 0.59159 | 0.60516<br>±0.46567 | 0.02808<br>±0.00902 |
| C | K | 10 | 0.65449 | 0.56936<br>±0.37787 | 5.60484<br>±1.31929 |
| D | E | 10 | 0.87378 | -6.87094E-4<br>±7.79916E-4 | 2.0522E-5<br>±2.57928E-6 |
| D | F | 7 | 0.42168 | -5.3884E-4<br>±0.00286 | 0.00668<br>±0.00288 |
| D | G | 11 | 0.72984 | -6.63111E-4<br>±0.00103 | 1.51909E-5<br>±2.87005E-6 |
| D | H | 11 | 0.74498 | -2.70434E-4<br>±9.41435E-4 | 0.08803<br>±0.01602 |
| D | I | 10 | 0.48999 | -0.01065<br>±0.00522 | 0.05385<br>±0.01734 |
| D | J | 7 | 0.9014 | -0.00402<br>±0.00129 | 1.87098E-4<br>±2.50345E-5 |
| D | K | 352 | 0.56852 | -8.4136E-5<br>±6.13262E-4 | 0.02208<br>±0.00267 |
| E | F | 7 | 0.62948 | 15.09039<br>±98.00387 | 329.88603<br>±98.60121 |

| | | | | | |
|---|---|---|---|---|---|
| E | G | 6 | 0.90082 | -29.91053 ±51.76472 | 0.77526 ±0.1138 |
| E | H | 6 | 0.90018 | 15.29797 ±46.7743 | 4320.16643 ±636.36761 |
| E | I | 6 | 0.77716 | -705.21392 ±241.35439 | 3264.65678 ±760.29517 |
| E | J | 7 | 0.83593 | -104.22125 ±72.08886 | 7.84652 ±1.39653 |
| E | K | 10 | 0.85671 | -63.33182 ±46.83028 | 1210.47624 ±163.50073 |
| F | G | 4 | 0.86535 | -0.00122 ±0.19241 | 0.0016 ±3.54823E-4 |
| F | H | 4 | 0.88984 | 0.18116 ±0.14244 | 8.13811 ±1.62009 |
| F | I | 5 | 0.76318 | -1.60482 ±0.71752 | 8.10653 ±2.17507 |
| F | J | 6 | 0.7048 | 0.04347 ±0.21799 | 0.01441 ±0.00401 |
| F | K | 7 | 0.62006 | -0.03917 ±0.30142 | 2.96529 ±0.90264 |
| G | H | 14 | 0.85805 | 43.91192 ±34.12617 | 5535.32326 ±620.48142 |
| G | I | 5 | 0.85044 | -1114.71561 ±308.88727 | 5111.68637 ±1049.00501 |
| G | J | 5 | 0.85217 | -86.8735 ±113.19265 | 9.88189 ±2.01471 |
| G | K | 15 | 0.84659 | -21.88416 ±38.89158 | 1622.40313 ±183.39973 |
| H | I | 12 | 0.85662 | -0.18822 ±0.04888 | 0.85826 ±0.16598 |
| H | J | 5 | 0.929 | -0.03673 ±0.01488 | 0.00193 ±2.64764E-4 |
| H | K | 14 | 0.74016 | -0.00487 | 0.25594 |

| | | | | | |
|---|---|---|---|---|---|
| | | | | ±0.00905 | ±0.0415 |
| I | J | 4 | 0.37635 | 0.20466 ±0.06879 | 0.00179 ±0.00107 |
| I | K | 12 | 0.85662 | 0.19375 ±0.01175 | 0.37568 ±0.04599 |
| J | K | 8 | 0.64406 | 10.33029 ±10.40129 | 140.59265 ±38.03136 |

Data basis to Figure 15

All the data used to generate this figure is from mammal species: adult body mass data from [1], body surface-area per unit body mass data from [128], specific IF1 protein activity data from [91], specific $F_1F_0$ ATP hydrolysis rate data from [91], specific basal metabolic rate data from [1] (for those 350 mammal species with specific basal metabolic rate data, plus adult body mass and maximal lifespan data, there. Specific basal metabolic rate values, absent in [1], were sourced for cat {*Felis catus*}, by value from [289], and for sheep {*Ovis aries*} and dog {*Canis lupus familiaris*} as given in earlier Table S2, with sources disclosed in its corresponding text), heart rate data from [91], specific Reactive Oxygen Species (ROS) generation rate data (mean of superoxide and hydrogen peroxide data, for species for which both are provided) from [95], specific DNA substitution (total of SBS1, SBSB, and SBSC types) rate data from [100], specific DNA indel (somatic insertions and deletions) rate data from [100], specific DNA damage rate data from [96], specific mtDNA damage rate data from [100], specific mtDNA damage rate data from [99], (inferred, as explained in the main text) specific protein damage rate data from [102], specific lipid damage rate data from [101], specific epigenetic damage rate data (data calculated for eutherian species probed by the mammalian methylation array [127], curtailed to those species thereof which have maximal lifespan data available in [1]), and maximal lifespan data from [1].

Where "specific" here refers to per unit mass or otherwise controlled for body mass. Where human data was present, it wasn't omitted.

Mentioned above, each of specific lipid damage rate data from [101], specific DNA damage data from [96], and specific mtDNA damage data from [99], are (actually) amount of damage accumulated after a period, not rate of damage. But each is abstractly compared to other variables as if each were a rate of damage. Because rate=amount/period, where period is the same within (but not necessarily between) each of these variables, for different species' values, and so cancels out in comparison with other variable(s). In the mtDNA case the period is approximately the same chronologically (0.8-2.5 years [99]), whereas in the other two cases the period is approximately the same percentage (15-30% [101]; 20%, except pig at 4% [96]) of maximal lifespan in each species case. So, two different methods, which presumably can't both be equally valid. All this is an abstractive and approximating drag on accuracy. Accuracy might be improved by calculating rates explicitly for these variables where data availability permits, but this wasn't done. Although reassuringly the Pearson correlation coefficient between DNA damage rate data (*explicitly* rate data; total of SBS1, SBSB, and SBSC substitution rates) from [100] and DNA damage rate data (*abstractly* rate data) from [96] is 0.9517. And the Pearson correlation coefficient between mtDNA damage rate data (*explicitly* rate data) from [100] and mtDNA damage rate data (*abstractly* rate data) from [99] is 0.9864.

From [1], only adult body mass, maximal lifespan, and specific basal metabolic rate data for the 350 mammal species with data for *all* these three variables was sourced. Although, adult body mass and maximal lifespan data was sourced atop from [1] for (because datum/data for each of these species is in at least one of the other aforementioned data sets used in this section) *Felis catus*, *Ovis aries*, *Canis lupus familiaris*, *Equus caballus*, *Giraffa camelopardalis*, *Mustela furo* (*Mustela putorius*), *Castor canadensis*, *Megaptera novaeangliae*, *Balaena mysticetus*, *Lemur catta*, *Macaca mulatta*, *Lontra canadensis*, *Capra hircus*, *Tursiops truncates*, *Delphinapterus leucas*, *Balaenoptera acutorostrata*, *Phoca hispida*, *Halichoerus grypus*, *Felis silvestris*, *Delphinus delphis*, *Lagenorhynchus acutus*, *Balaenoptera physalus*, *Balaenoptera musculus*, *Physeter macrocephalus*. When sourcing data from [128], data for "Peromyscus" and "rhinoceros" in [128] was ignored because these terms are ambiguous. "Cavia" in [128] was *assumed* to refer to *Cavia porcellus*. Data in [128] for the following species was ignored because these species don't have adult body mass and/or maximal lifespan values in [1]: *Lagenorhynchus albirostris, Mesoplodon bidens, Antilope rubricata, Mustela foina, Phoca groenlandica, Microtus pennsylvanicus, Clethrionomys gapperi, Ondatra zibethicus*.

Data from 374 mammal species was in this data set. This data set is broader than that used for Figure 14 because it uses more data sources as in some cases it uses multiple data sources for the same/similar thing, e.g., nuclear DNA damage, and because it uses less stringent species data exclusion criteria. For inclusion, distinct from before, here not requiring a species to have a specific basal metabolic rate value in [1]. But still excluding species from the full set of 384 species when they don't have both a maximal lifespan and adult body mass value in [1], which excluded (*the previously listed*) 8 species, and excluding atop the (*previously listed*) 2 species of ambiguous identity (so we don't know if they have such data in [1] or not), rendering 374 mammal species.

Considering the variables shown in Figure 14: terming adult body mass as the "input variable", maximal lifespan as the "output variable", and the other variables as "internal variables".

Specific $F_1F_0$ ATP hydrolysis rate data and specific IF1 protein activity data were collapsed into a single variable, of the former divided by the latter, and they were both substituted for this combination variable. Replacing two variables with one variable. This "internal variable", like specific $F_1F_0$ ATP hydrolysis rate, and unlike specific IF1 protein activity, positively correlates with the other internal variables. When counting internal variables herein, this combination variable is counted as a single internal variable.

The collective data set was curtailed to data from species that had data for 3 or more of the 13 internal variables, which was 25 species. For this 25 species subset only, each of the internal variables was standardized. Standardized to be within the range of 0-100, using Eq. S1. To repeat, only data for these 25 species was used in the standardization step, only data for these 25 species determined values of min($x$) and max($x$) in Eq. S1. Then for each of these 25 species, the mean of its standardized internal variables was calculated. Such means were used to populate a new variable, of 25 data points, called $\lambda$. Which is plotted in Figure 15 of the main text.

A flaw is that not all these 25 species have data for all the internal variables, and there is a lot of variation in how many, and which, internal variables these 25 species have data for (162 internal variable data points across 25 species = mean of 6.48 internal variable data points per species). Incredibly, despite this, because the internal variables are so highly linearly correlated, and because averaging divides by the number of data points, this analysis still returns a tractable result.

Shown in Figure S6 (which also shows $\log_{10}$-$\log_{10}$ plots), equivalent results were returned when instead curtailing to data for those species that had data for 2 or more of the 13 internal variables, which was 80 species (*after omitting outlier Sorex Araneus, the Eurasian shrew, because species in family of Soricidae don't have a true specific basal metabolic rate [129]. And after omitting outlier Mustela frenata, the long-tailed weasel, which has a very long tail, compared to its body length, that makes its surface-area anomalously large for its body mass*). Here, only data for these 80 species was used in the standardization step, only data for these 80 species determined values of min($x$) and max($x$) in Eq. S1. In this case, 272 internal variable data points across 80 species = mean of 3.4 internal variable data points per species.

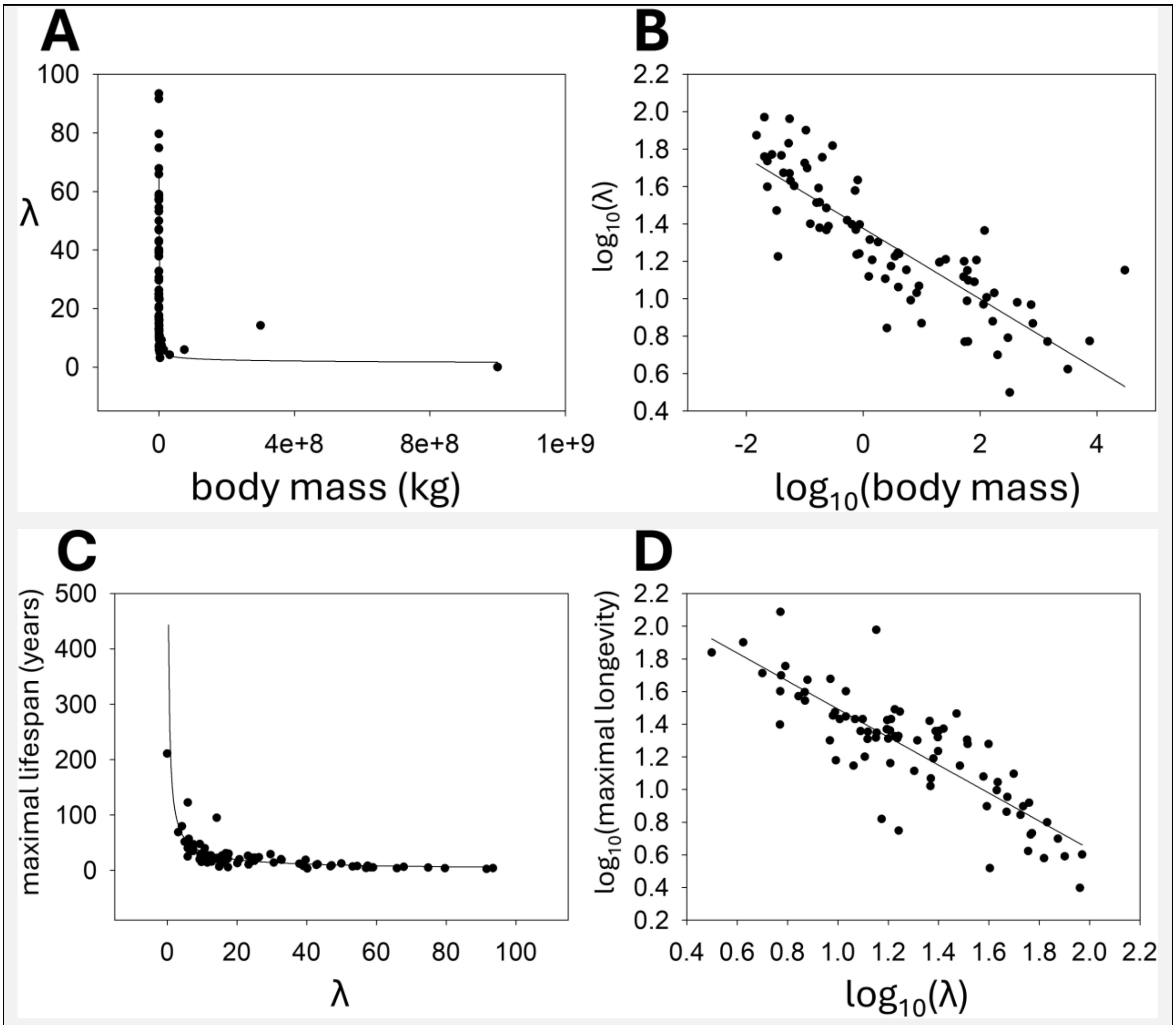

Figure S6 – **(A)** $\lambda$ vs. body mass (kg), where $\lambda$ incorporates data from 80 species that have data for 2 or more of the internal variables. Best-fit line equation: $\lambda = 25.0758 \cdot$

$(body\ mass)^{-0.2339}$. Adjusted $R^2$ = 0.6799. **(B)** $Log_{10}$-$log_{10}$ plot of $\lambda$ vs. body mass (kg). Best-fit line equation: $log_{10}(\lambda) = 1.3768 + -0.1889 \cdot log_{10}(body\ mass)$. Adjusted $R^2$ = 0.7018. **(C)** Maximal lifespan (years) vs. $\lambda$. Best-fit line equation: $maximal\ lifespan = 201.7743 \cdot \lambda^{-0.7805}$. Adjusted $R^2$ = 0.5528. **(D)** $Log_{10}$-$log_{10}$ plot of maximal lifespan (years) vs. $\lambda$. Best-fit line equation: $log_{10}(maximal\ lifespan) = 2.3509 + -0.8577 \cdot log_{10}(\lambda)$. Adjusted $R^2$ = 0.7040.

*Plots made, and best-fit lines found, using SigmaPlot 15.0 software (Systat Software, Inc., San Jose, California, USA). Human data included. Bowhead whale excluded from the $log_{10}$-$log_{10}$ plots because its value of $\lambda$ is negative (-3.38414E-08; not precisely zero because of rounding error) and so can't be logged.*

Comparing Figures 15 and S6 (more precisely, 15A-B with S6A and S6C), their best-fit line equation parameters are relatively close to each other. Their exponent for $\lambda$ vs. body mass is the same to two significant figures (≈*-0.25, quarter-power scaling*).

[Table S70]

| | **Figure 15** $\lambda$ vs. body mass | Figure S6 $\lambda$ vs. body mass | **Figure 15** longevity vs. $\lambda$ | Figure S6 longevity vs. $\lambda$ |
|---|---|---|---|---|
| Multiplicator | 28.9609 (±4.2354) | 25.0758 (±1.7717) | 335.3174 (±60.9121) | 201.7743 (±37.9390) |
| Exponent | -0.2344 (±0.0519) | -0.2339 (±0.0241) | -1.0438 (±0.1007) | -0.7805 (±0.0846) |

Because of the observed non-linearity between maximal lifespan and $\lambda$ (Figures 15 and S6), formalized in the best-fit line equation (Eq. 18), an absolute decrease amount in $\lambda$ corresponds with a greater absolute increase amount in maximal lifespan the smaller that $\lambda$ is. Although this equation was derived from interspecies data, it might have utility for intraspecies use. For example, to predict a new higher maximal lifespan if $\lambda$ is successfully reduced to a lower value by a drug intervention (e.g., by a drug that selectively inhibits $F_1F_0$ ATP hydrolysis, *without inhibiting $F_1F_0$ ATP synthesis* [18-20]). Interestingly, if applicable in this way, this equation predicts that a certain absolute decrease in $\lambda$ corresponds with, for example, a greater increase in maximal lifespan in a human than in a mouse.

Equivalent results were returned when specific $F_1F_0$ ATP hydrolysis rate data and specific IF1 protein activity data weren't (as aforementioned) combined and, specific IF1 protein activity data was omitted instead (data not shown).

Figure S7A is a plot of $\lambda$ (on the $x$-axis) against componentry data to $\lambda$. For the 80 species set (species with data for 2 or more internal variables set). For these species, Figure S7B is a plot of *predicted* maximal lifespan, predicted from $\lambda$ by Eq. 18 (in main text), against *observed* maximal lifespan. The parameters of Eq. 18 were fitted on a smaller species set, of 25 species. So, here tests Eq. 18 in predicting values that were, but also many others that weren't, used in setting its parameters. It performs well.

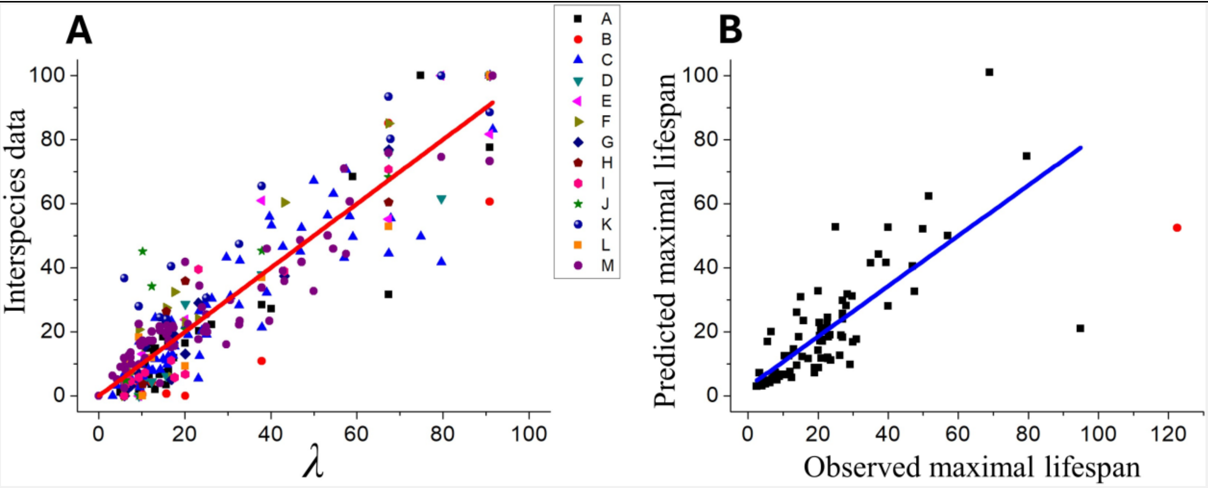

Figure S7 – (**A**) Plot of $\lambda$ (on $x$-axis) against componentry data to $\lambda$. For the 80 species set (species with data for 2 or more internal variables set). $\lambda$ was generated as described in the corresponding text, with data sources given there. This graph wasn't used to generate $\lambda$. It visualizes the result. Referring to the key panel in the graph: *A* is body surface-area per unit body mass, *B* is specific $F_1F_0$ ATP hydrolysis rate divided by specific IF1 protein activity, *C* is specific basal metabolic rate, *D* is heart rate, *E* is specific Reactive Oxygen Species (ROS) generation rate, *F* is specific DNA substitution rate, *G* is specific DNA indel rate, *H* is specific DNA damage rate, *I* is specific mtDNA damage rate, *J* is specific mtDNA damage rate, *K* is (inferred) specific protein damage rate, *L* is specific lipid damage rate, and *M* is (calculated from data) specific epigenetic damage rate. Best-fit line shown is for *all* the data (pooled). Its slope equals 1 (±0.02539), its intercept is close to zero (8.16804e-11±0.94437), and its adjusted $R^2$ is 0.85122. (**B**) For these 80 species, plot of *predicted* maximal lifespan, predicted from $\lambda$ by Eq. 18 in the main text, against *observed* maximal lifespan. Bowhead whale omitted because its value of $\lambda$ (-3.38414E-08, not zero because of rounding error), when inputted into Eq. 18, returns a complex number. So, data from 79 species is used to generate the plot. Human data point is a red circle, as opposed to the others, which are black squares. Human was omitted from calculating the best-fit line, by rationale given in the Methods section. The slope of this best-fit line is 0.7872 (±0.07601), its intercept is 2.7928 (±2.15844), and adjusted $R^2$ is 0.57987. If the other outlier, humpback whale, is omitted also, the best-fit line slope is 1.0240, with adjusted $R^2$ = 0.7571. Including human and humpback whale, Pearson correlation coefficient = 0.7361, one-tailed (alternative hypothesis is directional) p-value = 5.41271E-15. Excluding human: Pearson = 0.7651. Excluding human and humpback whale: Pearson = 0.872.

Figure 11 in the main text is a plot for 52 of these 80 species. Those species that I have specific basal metabolic rate and specific epigenetic damage rate *($\Delta M$)* data for. It is a plot of maximal lifespan vs. these variables when they are standardized by Eq. S1. Note the overlap of plots.

I'd like to repeat the analysis of this section in the future when more complete data becomes available, especially because the standardization step standardizes a variable value according to, and so is sensitive to, what the other values are in the variable. So, when the different internal variables don't all have data from the same plurality of species, plus can greatly vary in how many species they have data for, the average of internal variables for some species can be adversely skewed compared to others, and the results sub-optimal. Which is a problem made worse the lower the stipulation of how many internal variables a species must have data for to be included in the analysis (e.g., 2 is worse than 3).

Despite this, similar results were returned when this issue was made even worse, by instead curtailing to the data for those species that had data for just 1 or more of the 13 internal variables, which was 363 species (*after omitting all species present from the family of Soricidae, which don't have a true specific basal metabolic rate value [129]. And omitting outlier Mustela frenata, the long-tailed weasel, which has a very long tail, compared to its body length, that makes its surface-area anomalously large for its body mass*). Shown in Figure S8. Here, only data for these 363 species was used in the standardization step, only data for these 363 species determined values of min($x$) and max($x$) in Eq. S1. Out of this 363 species data set, 275 species only had data for specific basal metabolic rate, and 8 species only had data for body surface-area per unit body mass. 555 internal variable data points across 363 species = mean of 1.53 internal variable data points per species.

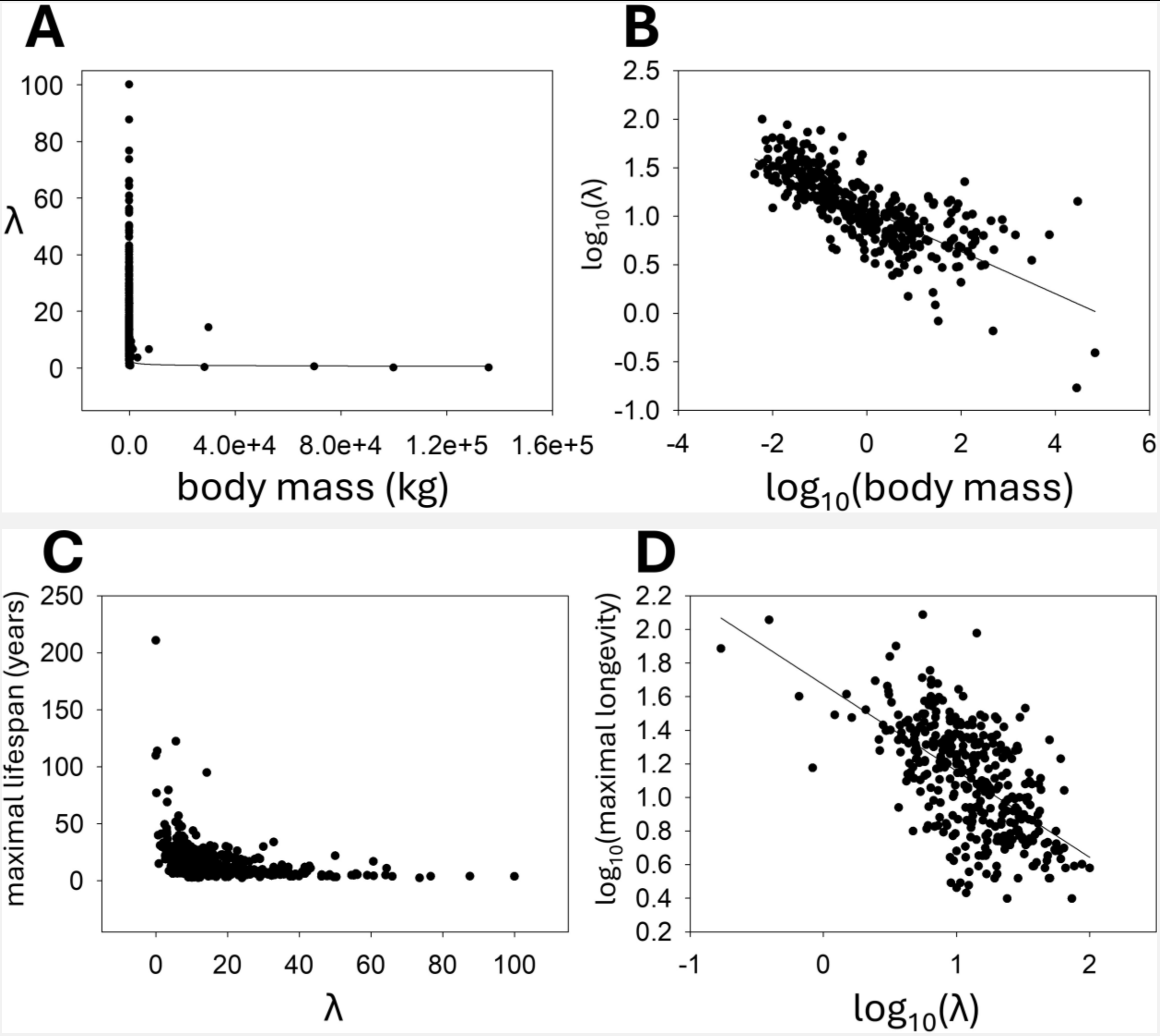

Figure S8 – **(A)** $\lambda$ vs. body mass (kg), where $\lambda$ incorporates data from 363 species that have data for 1 or more of the internal variables. Best-fit line equation: $\lambda = 12.5931 \cdot (body\ mass)^{-0.2577}$. Adjusted $R^2$ = 0.5283. **(B)** $Log_{10}$-$log_{10}$ plot of $\lambda$ vs. body mass (kg). Best-fit line equation: $log_{10}(\lambda) = 1.0724 + -0.2175 \cdot log_{10}(body\ mass)$. Adjusted $R^2$ = 0.5741. **(C)** Maximal lifespan (years) vs. $\lambda$. Program used couldn't converge to a best-fit line equation, reporting instead "Did not converge, inner loop failure. Parameters may not be valid. Overflow in partial derivatives." The equation that I tried to fit was $y = m \cdot x^a$, with initial parameters: $m$ = 200, $a$ = -1. **(D)** $Log_{10}$-$log_{10}$ plot of maximal lifespan (years) vs. $\lambda$.

Best-fit line equation: $log_{10}(maximal\ lifespan) = 1.6274 + -0.5144 \cdot log_{10}(\lambda)$.
Adjusted $R^2$ = 0.3565.
*Plots made, and best-fit lines found, using SigmaPlot 15.0 software (Systat Software, Inc., San Jose, California, USA). Human data included. Bowhead whale (Balaena mysticetus) and blue whale (Balaenoptera musculus) excluded from the $log_{10}$-$log_{10}$ plots because their value of $\lambda$ is zero, or close to zero and negative, and so can't be logged.*

Comparing best-fit line equation $\{log_{10}(y) = log_{10}(a) + b \cdot log_{10}(x)\}$ parameters for data sets used to generate Figures 15, S6, and S8, which each curtail to data for those species that had data for 3, 2, and 1 of the internal variables respectively: 25, 80, and 363 species respectively: *A*, *B*, and *C* respectively in Table S71 below: the parameters for *A* and *B* overlap within their standard errors:
[Table S71]

| | $Log_{10}$-$log_{10}$ ($\lambda$ vs. body mass) | | $Log_{10}$-$log_{10}$ (maximal lifespan vs. $\lambda$) | |
|---|---|---|---|---|
| | $log_{10}(a)$ | $b$ | $log_{10}(a)$ | $b$ |
| A | 1.3702 (±0.0659) | -0.2024 (±0.0418) | 2.3440 (±0.0940) | -0.8470 (±0.0750) |
| B | 1.3768 (±0.0223) | -0.1889 (±0.0139) | 2.3509 (±0.0837) | -0.8577 (±0.0628) |
| C | 1.0724 (±0.0130) | -0.2175 (±0.0099) | 1.6724 (±0.0420) | -0.5144 (±0.0363) |

<u>Alternative equation relating maximal lifespan to $\lambda$</u>
The following is the best-fit line equation of Figure S9A: [Eq. S7]

$$\frac{ln(Maximal\ lifespan)}{Maximal\ lifespan} = (0.0032 \cdot \lambda) + 0.0788$$

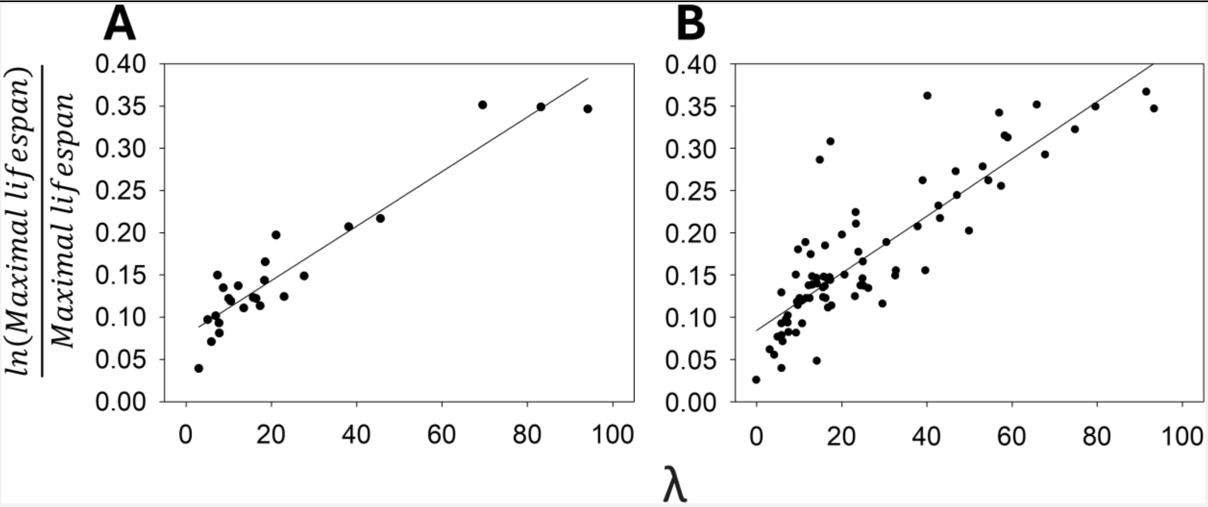

<u>Figure S9</u> – **(A)** $(ln[Maximal\ lifespan])/Maximal\ lifespan$ vs. $\lambda$, where $\lambda$ incorporates data from 25 species that have data for 3 or more of the internal variables. Best-fit line equation: $(ln[Maximal\ lifespan])/Maximal\ lifespan = (0.0032 \cdot \lambda) + 0.0788$. Adjusted $R^2$ = 0.8986. Pearson correlation coefficient = 0.9502, one-tailed (alternative hypothesis is directional) p-value = 1.94744E-13.

**(B)** $(ln[Maximal\ lifespan])/Maximal\ lifespan$ vs. $\lambda$, where $\lambda$ incorporates data from 80 species that have data for 2 or more of the internal variables. Best-fit line equation: $(ln[Maximal\ lifespan])/Maximal\ lifespan = (0.0034 \cdot \lambda) + 0.0843$. Adjusted $R^2$ = 0.7250. Pearson correlation coefficient = 0.8535, one-tailed (alternative hypothesis is directional) p-value = 4.35429E-24.

The parameters/constants of the two best-fit line plots are the same within their standard errors: Plot A = 0.0032 (±0.0002), 0.0788 (±0.0074); Plot B = 0.0034 (± 0.0002), 0.0843 (±0.0079).

Plots done, and their best-fit line equations found, using SigmaPlot 15.0 software (Systat Software, Inc., San Jose, California, USA). Human data among that used.

Figure S6 and S9B use slightly incorrect data, which I can't fix now as I no longer have access to SigmaPlot software. They use $\lambda$ data in which the specific $F_1F_0$ ATP hydrolysis rate divided by specific IF1 protein activity variable was accidentally omitted for rat and mouse, which makes these species' values of $\lambda$ incorrect by 2.46% and 1.51% respectively. Small percentage changes, with 2 of 80 species. This impacts best-fit lines calculated from those plots and any comparisons made thereto. Such as in earlier Tables S70 and S71.

Repeating the linear regression of Eq. 21 but with different $\lambda$

Eq. 21 used the 25 species set (species with data for 3 or more internal variables set). If $\log_{10} \lambda$ instead incorporates data from the 80 species that have data for 2 or more of the internal variables, but omitting bowhead whale because its value of $\lambda$ is negative (-3.38414E-08, not precisely zero because of rounding error in Microsoft Excel spreadsheet software) and so can't be logged, rendering 79 species included in the model, including human: adjusted $R^2$ is 0.704. The p-values for unstandardized $\beta_0$ and $\beta_1$ are lower at 3.215e-42 and 2.792e-22 respectively, and the p-value for the F-statistic is lower at 2.792e-22. The Pearson correlation coefficient between $\log_{10}$ maximal lifespan and $\log_{10} \lambda$ = -0.8414, with one-tailed (alternative hypothesis is directional) p-value = 1.41034e-22.

As shown in the next headed section, if $\log_{10}$ adult body mass (kg) is added as a second predictor, the adjusted $R^2$ is 0.707 (i.e., only increased by 0.003).

Multiple linear regressions with $\lambda$

In JASP (version 0.18.1.0), I conducted multiple linear regressions with maximal lifespan (*years; data sourced from [1]*) as the response variable, with adult body mass (*kg; data sourced from [1]*) and $\lambda$ as predictors. All logged ($\log_{10}$).

In the first model, model *A*, $\lambda$ incorporates data from 25 species that have data for 3 or more of the internal variables. In the second model, model *B*, $\lambda$ incorporates data from 80 species that have data for 2 or more of the internal variables: 79 species after omitting bowhead whale because its value of $\lambda$ is negative (-3.38414E-08) and so can't be logged. Human included in both.

Results shown in Table S72 below. Where *Adj.* $R^2$ is adjusted $R^2$, *VIF* is the Variance Inflation Factor for each of the slope regression coefficients, *DW* is the Durbin-Watson statistic, *p_F* is the p-value for the F-statistic, $S\beta_1$ and $S\beta_2$ are the standardized regression coefficients for adult body mass and $\lambda$ respectively, while $p\_\beta_0$, $p\_\beta_1$, and $p\_\beta_2$ are p-values for unstandardized regression coefficients for intercept, adult body mass and $\lambda$ respectively. JASP's "Enter" method used, with adult body mass forced in first. For each, assumptions of homoscedasticity, normality, and linearity weren't violated.

[Table S72]

| | Adj. $R^2$ | VIF | DW | p_F | S_$\beta_1$ | S_$\beta_2$ | p_$\beta_0$ | p_$\beta_1$ | p_$\beta_2$ |
|---|---|---|---|---|---|---|---|---|---|
| A | 0.833 | 2.018 | 1.415 | 1.050e-9 | -0.012 | -0.929 | 2.931e-13 | 0.918 | 8.028e-8 |
| B | 0.707 | 3.393 | 1.494 | 2.164e-21 | 0.144 | -0.721 | 4.927e-22 | 0.207 | 1.265e-8 |

Table S72 above shows that $\lambda$ *is*, and adult body mass is *not*, a significant predictor of maximal lifespan. So, even with adult body mass included in the model, $\lambda$ is a significant predictor of maximal lifespan.

In Table S73 below are corresponding (to the multiple linear regressions of Table S72 above, also JASP outputted) partial and semi-partial (part) Pearson correlation coefficients to $\log_{10}$ maximal lifespan. One-tailed (alternative hypothesis is directional) p-values are italicized in brackets.

[Table S73]

| *Variables are logged ($log_{10}$). Pearson correlation coefficients.* | | | | |
|---|---|---|---|---|
| | | Zero-order | Partial | Part |
| A | $\lambda$ | -0.9205 *(3.64435E-11)* | -0.859 *(1.93925E-08)* | -0.654 *(0.000195636)* |
| | Adult body mass | 0.6476 *(0.000233)* | -0.022 *(0.5406359)* | -0.009 *(0.5166461)* |
| B | $\lambda$ | -0.8414 *(1.41034e-22)* | -0.591 *(4.90234E-09)* | -0.391 *(0.000183293)* |
| | Adult body mass | 0.749 *(1.02732E-15)* | 0.144 *(0.102737)* | 0.078 *(0.247218)* |

Table S73 above shows that statistically excluding the effect of $\log_{10}$ adult body mass doesn't significantly reduce the Pearson correlation coefficient between $\log_{10}$ maximal lifespan and $\log_{10} \lambda$. By contrast, statistically excluding the effect of $\log_{10} \lambda$ significantly reduces the Pearson correlation coefficient between $\log_{10}$ maximal lifespan and $\log_{10}$ adult body mass. Collinearity, measured by *VIF*, can corrupt partial and part correlations, but for *A* at least, its *VIF* value isn't so high (=2.018; as shown in earlier Table S72).

Generalized Additive Models (GAM) with $\lambda$

Using the same R code as earlier (which leverages the mgcv R package [334]) but with different data: I built GAM models with maximal lifespan in years ($y$; data sourced from [1]) as the response variable, with adult body mass in kg ($z$; data sourced from [1]) and $\lambda$ ($x$) as predictors. No variables logged. In the first model, model $A$, $\lambda$ incorporates data from 25 species that have data for 3 or more of the internal variables. In the second model, model $B$, $\lambda$ incorporates data from 80 species that have data for 2 or more of the internal variables. Human included in both. For each GAM: family was Gaussian, link was identity, and their output is shown in Table S74 below, with column headings as defined for earlier Table S52.

[Table S74]

| | A.$R^2$ | GCV | Dev | Int | Int.p | | edf | ref.df | P | K | CC |
|---|---|---|---|---|---|---|---|---|---|---|---|
| A | 0.83 | 192 | 91.3% | 28.301 | 4.17e-09 | s($x$) | 8.650 | 8.953 | 0.000181 | 9 | 0.8671001 |
| | | | | | | s($z$) | 3.116 | 3.725 | 0.181739 | 9 | 0.8671001 |
| B | 0.848 | 149.24 | 86.6% | 26.274 | <2e-16 | s($x$) | 6.891 | 7.953 | <2e-16 | 9 | 0.7934631 |
| | | | | | | s($z$) | 2.755 | 3.036 | <2e-16 | 9 | 0.7934631 |

In Table S74 above, $\lambda$ is a highly significant predictor of maximal lifespan in both the GAM models, while adult body mass is only significant in the second GAM model ($B$). Note the high values of adjusted $R^2$ ($A.R^2$) and deviance explained ($Dev$). So, even with adult body mass included in the model, $\lambda$ is a significant predictor of maximal lifespan.

*Diagnostic graphs for these GAMs: Model-predicted maximal lifespan vs. observed maximal lifespan plots are linear, each with a single outlier, which is human (rationale disclosed in the Methods section may explain why human is anomalous).*

*Residuals vs. predicted (values of response variable) plot shouldn't contain pattern(s) and/or trend(s), but there are some outlier data points which makes the other data points appear bunched in the graph, which might be misconstrued as regularity. There might be some minor non-randomness further to that. The Q-Q plot should be linear, and it is except for non-linearity at its ends, with a single far-removed outlier observed. Working responses vs. linear predictor plot should be linear and it is, with no outliers present. For a GAM, partial residuals vs. predictor plots should appear random and evenly distributed around zero, which (some outliers aside) is arguably the case, all be it this isn't ideally clear cut, there arguably being pattern(s) and/or trend(s), which is a potential issue. The Component-Plus-Residual (CPR) plot is, as it should be, data points scattered around linear, all be it there are some far-removed outliers.*

*High concurvity indicates redundancy/entwinement of predictors and can inflate smooth term p-values. The first GAM model (A) has quite high concurvity in its presented "worst" values, but less in its "observed" and "estimate" values, especially less in its "estimate" values, as shown in Table S75 below.*

*[Table S75]*

| | *Worst* | *Observed* | *Estimate* |
|---|---|---|---|
| *s(x)* | *0.8671001* | *0.6390933* | *0.3008846* |
| *s(z)* | *0.8671001* | *0.3368205* | *0.3092758* |

When human is omitted: for the first GAM model (*A*), deviance explained (*Dev*) increases from 91.3 to 92.6%, and adjusted $R^2$ ($A.R^2$) increases from 0.83 to 0.865; and for the second GAM model (*B*), *Dev* increases from 86.6 to **95.4%**, and $A.R^2$ increases from 0.848 to **0.943**. These are high values. With human omitted, the p-values for the intercept and both the smooth terms are significant (at $p < 0.05$) in both GAM models, *A* and *B*.

Table S76 below shows results from when the GAM models are simplified, such that $\lambda$ is the only predictor (*N.B., human is included*):

[Table S76]

| | $A.R^2$ | GCV | Dev | Int | Int.p | | edf | ref.df | P | K |
|---|---|---|---|---|---|---|---|---|---|---|
| A | 0.764 | 210.88 | 84.7% | 28.301 | 1.91e-09 | s(*x*) | 8.481 | 8.913 | 7.96e-05 | 9 |
| B | 0.7 | 289.97 | 73.2% | 26.274 | <2e-16 | s(*x*) | 8.509 | 8.932 | <2e-16 | 9 |

In Table S76 above, in both models, $\lambda$ is a *significant* predictor of maximal lifespan.

$\lambda$ and phylogeny

The Phylogenetic Independent Contrasts (PIC) method requires compared traits to be linearly related [324], and so the following PIC analysis was conducted upon logged variables of maximal lifespan (data sourced from [1]), adult body mass (data sourced from [1]), and $\lambda$. Where $\lambda$ incorporates data from 80 species, which have data for 2 or more of the internal variables, or at least this was the starting point. But the following 4 species were omitted, leaving 76 species remaining in $\lambda$: *Balaena mysticetus* (bowhead whale), *Potorous tridactylus*, *Eptesicus fuscus*, and *Rattus rattus*. Omitting bowhead whale because its value of $\lambda$ is negative (-3.38414E-08, not precisely zero because of rounding error) and so can't be logged. And the other species were omitted because of an "unresolved name" issue when sourcing the phylogeny (which has no polytomies) from [322].

The diagnostic table below, Table S77, shows for $Q$ ($\log_{10}\lambda$), $R$ ($\log_{10}$ maximal lifespan), and $T$ ($\log_{10}$ adult body mass): in column $C$, the Pearson correlation coefficient (*not* forced through origin [323, 325]) of the absolute values of the standardized phylogenetically independent contrasts versus their standard deviations (i.e., the square root of the sum of their branch lengths), which is a diagnostic check of how well the topology and branch lengths of the phylogenetic tree (and assumed model of character evolution) adequately fit the tip data [323], wherein no significant correlation is the desirable result (*and in the line below, labelled "P", the corresponding two-tailed p-value thereof*), and subsequent columns are that but with a branch length adjustment made, as codified earlier. P-values greater than 0.05, which are desirable, are in red text. Table values were outputted by the PDAP module [320] of Mesquite [321].

[Table S77]

| Diagnostic table | | | | | |
|---|---|---|---|---|---|
| | *C* | Nee | Pagel | Grafen | =1.0 |
| **Q** | 0.4855995758204129 | -0.21846110120165887 | -0.36471494525695697 | -0.4151211821819699 | -0.13844495814996974 |
| *P* | 9.04960995029569E-10 | 0.05970514884521479 | 0.0012951344909916745 | 2.128465305369673E-4 | 0.23619738683069658 |
| **R** | 0.4260924530900193 | -0.0183782095 | -0.204892 | -0.3249839391353377 | 0.08832675568001609 |

| | | | | | |
|---|---|---|---|---|---|
| | | 7206389 | 76453873748 | | |
| *P* | 3.466329963286441E-6 | 0.8756389921207685 | 0.07783637813081218 | 0.0044435333176292635 | 0.4511140944730112 |
| **T** | 0.5135019280022866 | -0.08836621215235003 | -0.2594555927454398 | -0.34775401364160224 | -0.024583584617693232 |
| *P* | 6.366240867805573E-11 | 0.45091112738509753 | 0.024585380271517212 | 0.0022350574555563413 | 0.8341704617339946 |

Following the teaching of [323, 325], raw phylogenetically independent contrasts were outputted from the PDAP module of Mesquite, and each was then standardized by dividing each contrast by the (also PDAP outputted) contrast's standard deviation (square root of sum of corrected branch lengths). The raw contrasts, and corresponding standard deviations to standardize them, for all three variables ($\log_{10} \lambda$, $\log_{10}$ adult body mass, $\log_{10}$ maximal lifespan) were sourced with branch lengths = 1.0, as this (according to the diagnostic table, Table S77, above) is a valid single branch length transformation for all three. Their having the same branch length transformation eases interpretability. These standardized phylogenetically independent contrasts are used in the multiple linear regression of the next paragraph.

Table S78 below reports upon multiple linear regression, where standardized phylogenetically independent contrasts for $\log_{10}$ maximal lifespan is the response variable and standardized phylogenetically independent contrasts for $\log_{10}$ adult body mass and standardized phylogenetically independent contrasts for $\log_{10} \lambda$ are explanatory variables. In the table, $p\_\beta_1$ and $p\_\beta_2$ are p-values for unstandardized regression coefficients, and $S\beta_1$ and $S\beta_2$ are standardized regression coefficients, for these explanatory variables respectively. *Adj.* $R^2$ is adjusted $R^2$, $p\_F$ is the p-value for the F-statistic, *VIF* is the Variance Inflation Factor for each of the slope regression coefficients, and *DW* is the Durbin-Watson statistic.

Regression was forced through the origin, following the teaching of [323, 325]. Conducted in JASP software (version 0.18.1.0), using its method called "Enter", wherein the adult body mass related predictor was forced in first. Assumptions of homoscedasticity, normality, and linearity weren't violated.

[Table S78]

| Adj. $R^2$ | VIF | DW | p_F | $S\_\beta_1$ | $S\_\beta_2$ | $p\_\beta_1$ | $p\_\beta_2$ |
|---|---|---|---|---|---|---|---|
| 0.431 | 1.701 | 1.939 | 4.184e-10 | 0.256 | -0.476 | 0.027 | 7.969e-5 |

In Table S78 above, which uses phylogenetically adjusted data: even with adult body mass in the model, *λ is* a highly significant predictor of maximal lifespan.

In Table S79 below are corresponding zero-order, partial, and semi-partial (part) Pearson correlation coefficients. Forced through the origin [323, 325]. One-tailed (alternative hypothesis is directional) p-values are italicized in brackets.

[Table S79]

| | Zero-order | Partial | Part |
|---|---|---|---|
| *λ* | -0.7140 *(2.23391E-13)* | -0.440 *(0.000035)* | -0.364 *(0.000614)* |
| Adult body mass | 0.5604 *(7.02677E-08)* | 0.255 *(0.013105)* | 0.196 *(0.0448605)* |

In Table S79 above, its zero-order correlations are PDAP [320] outputted Pearson correlation (forced through the origin [323, 325]) coefficients, adjusted to account for phylogeny, which are statistically significant. The partial and part correlations (forced through the origin [323, 325]) are JASP output. Statistically excluding the effects of both phylogeny and adult body mass doesn't significantly reduce the correlation between maximal lifespan and *λ*. Statistically excluding the effects of both phylogeny and *λ* reduces the correlation between maximal lifespan and adult body mass, nearly significantly so. Collinearity, measured by *VIF*, can corrupt partial and part correlations, but the corresponding *VIF* value isn't so high (=1.701; as shown in earlier Table S78).

<u>Across mammal species, higher adult body mass dictates higher maximal lifespan through *λ*</u>

Utilizing logged ($\log_{10}$) variables, conferring inter-linearity, mediation analysis reports that adult body mass *only* affects maximal lifespan through *λ*, with *no* direct effect upon maximal lifespan. Here $\lambda$ incorporates data from species that have data for 2 or more of the internal variables, which is 79 species after omitting bowhead whale because its value of $\lambda$ is negative (-3.38414E-08, not precisely zero because of rounding error) and so can't be logged. Conducted in JASP software (version 0.18.1.0), selecting its bootstrap method called "Percentile", run with 10,000 replications, utilizing its standardized estimates option, which makes each variable mean = 0 and standard deviation = 1. Result shown in Table S80 below, where zero (0) is *not* within the 95% Confidence Interval (CI) for an indirect effect, but it *is* for a direct effect.

[Table S80]

| | | | 95% Confidence Interval (CI) | | | 95% Confidence Interval (CI) | |
|---|---|---|---|---|---|---|---|
| α | β | **Indirect effect (α·β)** | Lower | Upper | **Direct effect (τ)** | Lower | Upper |
| -0.55 | -0.69 | 0.381 | 0.205 | 0.608 | 0.099 | -0.132 | 0.279 |

<u>Instrumental variable analysis reports causality between lower $\lambda$ and higher maximal lifespan</u>

The mediation analysis of the last headed section reported that adult body mass only affects maximal lifespan through $\lambda$, with *no* direct effect upon maximal lifespan. This teaches adult body mass as a suitable instrumental variable to explore causality between $\lambda$ and maximal lifespan.

Table S81 below reports such analysis with $\log_{10}$ maximal lifespan as the dependent variable, $\log_{10}$ $\lambda$ as the independent variable, and $\log_{10}$ adult body mass as the instrumental variable. Its result infers causality between lower $\lambda$ and higher maximal lifespan.

The coefficient for the effect of the independent variable on the dependent variable, $\beta$, is a Two-Stage Least Squares (TSLS) estimate. *E* is the standard error in $\beta$. *P* is the one-tailed (alternative hypothesis is directional) p-value for $\beta$. Causality can be inferred from it being significant (at $P < 0.05$). *A* is the p-value for the Anderson-Rubin test (under F distribution) and *C* is its 95% confidence interval. *A2* is the p-value for the conditional likelihood ratio test (under Normal approximation) and *C2* is its 95% confidence interval. R code used is in the supplementary methods section of this Supplementary Material document.

[Table S81]

| $\beta$ | E | P | A | C | A2 | C2 |
|---|---|---|---|---|---|---|

| | | | | | | |
|---|---|---|---|---|---|---|
| -0.90978 | 0.07 512 | **<2e-16** | 1.998 4e-15 | -1.06488668906218, -0.762354458397254 | 1.998 4e-15 | -1.06488654380149, -0.762354589629884 |

In Table S81 above, the value of *P*, at <2e-16, is smaller than the smallest value reportable by the R statistical software used (2e-16). Highly statistically significant.

Alternative standardization equation

As aforementioned, earlier presented Eq. S1 is herein used to standardize, which standardizes a variable's values to be in the range 0-100. An alternative standardization equation is below, which standardizes a variable's values such that their mean is zero and standard deviation is one.

$$z_i = \frac{x_i - \bar{x}}{x_\sigma} \quad \text{[Eq. S8]}$$

Where $z_i$ is the $i^{\text{th}}$ standardized value in the variable, $x_i$ is the $i^{\text{th}}$ value in the variable, $\bar{x}$ is the mean value of the variable, and $x_\sigma$ is the standard deviation of the variable.

To make the mean be zero, one or more of the equation's output values are negative, which prohibits their logging. An equation variant, not shown, has a sufficiently large additive constant to make all its output values positive. Incidentally, such an additive constant can be applied to Eq. S1 also, not shown and not used, so that no value of zero (which can't be logged) is in its output: e.g., with value one (1), such that all its standardized values outputted are between 1-101.

Across mammal species, maximal lifespan is inversely proportional to the specific damage rate of DNA, mtDNA, proteins, lipids, and the epigenome, which all linearly correlate with each other, and with Reactive Oxygen Species (ROS) generation rate

This section is a focus upon the lower part of Figure 14, from specific Reactive Oxygen Species (ROS) generation rate, which I interpret drives the damage rates thereunder.

In Table S82 below, [A] is specific mitochondrial ROS detected rate (mean of superoxide and hydrogen peroxide data. That used to produce columns F and K in the main text's Table 11, which reports correlations to mitochondrial ROS detected per unit mass per unit time); data from [95]. Incidentally, the Pearson correlation coefficient between this superoxide and hydrogen peroxide data is 0.6579 (one-tailed, because the alternative hypothesis is directional, p-value is 0.0540995). To remind, these ROS data sets are from reduced systems: Sub-Mitochondrial Particles (SMPs) and isolated mitochondria respectively. Reporting ROS detected (not necessarily ROS generated), after the interdicting action of any antioxidant enzyme(s) present in the experimental preparation (e.g., in the mitochondrial matrix in the case of isolated mitochondria). All be it at many points in this paper, when referring to this data, I do refer to "ROS generated" instead of "ROS detected". Abstractive language, which is hereby notice of. [B] is mean urinary excretion amount of repair products of oxidative DNA damage: 8-oxoGua, 8-oxodG, and 5-HMUra when each animal is aged around 20% of their maximal lifespan (except for pig at 4%), standardized for body mass by dividing by creatinine amount in the urine; data from [96]. [C] is DNA substitution rate (total of SBS1, SBSB, and SBSC substitutions per genome per year); data from [100]. [D] is mtDNA mutations per mtDNA copy per year; data from [100]. [E] is DNA indel rate (somatic insertions and deletions per genome per year); data from [100]. [F] is mean (across heart and brain, calculated as specified in the main text) amount (per amount of deoxyguanosine) of ROS damage product, 8-oxodG, in mtDNA, when each animal is aged around 0.8-2.5 years old; data from [99]. [G] is amount (per unit tissue mass) of lipid peroxidation product MDA when each animal is aged within 15-30% of their maximal lifespan, data from [101]. [H] is median protein degradation rate constant (median $K_{deg}$), which I *assume* to indirectly report upon (be a function of) the rate of ROS damage of proteins (*hypothesizing* that greater rate of ROS damage to proteins requires/means a greater protein degradation rate); data from [102] (N.B., median $K_{deg}$ for bowhead whale is a mean because only two bowhead whale individuals were studied in [102], prohibiting calculation of a median). [I] is *calculated* (from fit to experimental data, not directly experimental data itself) average epigenetic change per year, $\Delta M$ (defined in the main text, Eq. 5). [I] is calculated for each eutherian species probed by the mammalian methylation array [127], curtailed to those species thereof which have maximal lifespan data in [1] and that I have at least one data point for in any of variables [A] to [H].

Correlations of some variables to the data of one or more of variables [A], [B], [C], [D], [E], [F], [G], [H], [I], has been disclosed earlier in this report. Their correlations to each other are in Table S82 below (Pearson correlation coefficients. Corresponding p-values are in the lower table section, each one-tailed because each alternative hypothesis is directional).

[Table S82]

| **Pearson correlation coefficient (R):** | | | | | | | | | |
|---|---|---|---|---|---|---|---|---|---|
| | [A] | [B] | [C] | [D] | [E] | [F] | [G] | [H] | [I] |
| [A] | 1 | | | | | | | | |
| [B] | 0.9797 | 1 | | | | | | | |
| [C] | 0.9541 | 0.9517 | 1 | | | | | | |
| [D] | 0.9626 | 0.9449 | 0.9322 | 1 | | | | | |
| [E] | 0.9693 | 0.9498 | 0.9707 | 0.98 | 1 | | | | |
| [F] | 0.8745 | 0.808 | 0.9789 | 0.9864 | 0.9896 | 1 | | | |
| [G] | 0.874 | 0.9696 | 0.9429 | 0.973 | 0.9497 | 0.8324 | 1 | | |
| [H] | 0.9938 | 0.8863 | 0.9422 | 0.9482 | 0.9579 | 0.9272 | 0.7644 | 1 | |
| [I] | 0.8267 | 0.915 | 0.926 | 0.8719 | 0.9122 | 0.7549 | 0.8336 | 0.9326 | 1 |
| **p-value (one-tailed):** | | | | | | | | | |
| | [A] | [B] | [C] | [D] | [E] | [F] | [G] | [H] | [I] |
| [A] | 1 | | | | | | | | |
| [B] | 0.01015 | 1 | | | | | | | |
| [C] | 0.02295 | 0.006325 | 1 | | | | | | |
| [D] | 0.0187 | 0.007699 | 6.05E-07 | 1 | | | | | |
| [E] | 0.01535 | 0.0067 | 9.92E-10 | 4.42E-10 | 1 | | | | |
| [F] | 0.011319 | 0.096 | 0.001834 | 0.00095 | 0.000636 | 1 | | | |
| [G] | 0.011407 | 0.0152 | 0.008119 | 0.002652 | 0.02515 | 0.010069 | 1 | | |
| [H] | 0.0031 | 0.153267 | 0.008268 | 0.007021 | 0.005152 | 0.0364 | 0.1178 | 1 | |
| [I] | 0.010911 | 0.005265 | 3.67E-07 | 0.8719 | 2.5E-06 | 0.024903 | 0.005065 | 0.000005 | 1 |

In Table S82 above, all the correlations are significant (at $p < 0.05$). The asymptotically exact harmonic mean p-value [76] (*its post-published corrected method, as corrected on 7 October 2019*) for all p-values in Table S82 above = 1.099757E-08. Very significant. Even despite some extremely low values of *n* (i.e., underpowered) in some cases. Note how high the values of R are in the table. These interrelations, and others, are shown explicitly or implicitly in Figure 14.

In Figure 14, and in other places in this paper, specific (i.e., per unit mass) damage rates are referred to but wherein not all these variables [B] to [I] are strictly per unit mass measures. Although they are each standardized in a way that controls for the different adult body masses of the different species: e.g., each species entry in variable [D] is not a per organism measure but instead a per mtDNA copy measure. So, some self-lexicography here to give "specific" a slightly broader meaning in this context than is typical: it here meaning per unit mass, or otherwise controlled for body mass. A lexicography used often herein, which is hereby notice of.

Each of the variables correlate with the calculated rate of epigenetic change, [I], such that monitoring epigenetic change (e.g., by an epigenetic clock) is likely a good surrogate/proxy for monitoring them all.

The high degree of correlation here is consistent with ROS, abstracted by variable [A], driving (at least in part) each of the other variables: [B], [C], [D], [E], [F], [G], [H], [I].

I interpret that variables [B], [C], [D], [E], [F], [G], [H], and [I] are (at least partially, directly and/or indirectly) ROS damage (*N.B., ROS can cause DNA deletions and insertions [337-340]; ROS damage may increase other forms of damage by being damage of, and/or taking resource(s) {e.g., energy} away from, their repair/replacement/mitigation processes*). And that they each correlate with each of specific $F_1F_0$ ATP hydrolysis rate, specific basal metabolic rate, and variable [A] (≈specific ROS generation rate) because they are all (at least partially) driven by these three variables. Wherein specific $F_1F_0$ ATP hydrolysis rate is a drive of specific basal metabolic rate, which in turn is a drive of variable [A] (≈specific ROS generation rate), which (at least partially) drives variables: [B], [C], [D], [E], [F], [G], [H], [I] (Figure 14). Such that they might *all* be reduced by a selective $F_1F_0$ ATP hydrolysis inhibitor drug (*that doesn't inhibit $F_1F_0$ ATP synthesis*): e.g., a drug of one or more of my international (PCT) patent applications [18-20]).

To theorize, such inhibition of $F_1F_0$ ATP hydrolysis slows aging. Perhaps even reverses it, because of extant repair and degradation/turnover processes (e.g., protein turnover). Where (I argue) decreasing ROS generation and damage rate, by selectively inhibiting $F_1F_0$ ATP hydrolysis, is a better strategy than increasing repair/replacement processes. Because the latter requires ATP energy, which inherently generates more ROS (if this ATP is made aerobically), thence actually requiring greater repair/turnover, which subtracts from (and possibly even detrimentally exceeds) any benefit from the increased repair/turnover. Moreover, as a general principle when something is being repaired its use is typically impaired/prevented.

Combination variable $\Delta RD$

Using the data set used to produce Table S82 above, I omitted species that didn't have data for at least two of variables [A] to [I]. Then, using this 24 species subset, each of variables

[A] to [I] were standardized (to be within the range of 0-100) using Eq. S1. Then a new variable was made, populated with, for each species used, the mean of standardized [A] to [I].

In this 24 species subset, many species don't have data available for many of the variables [A] to [I], but the use of standardized data and averaging thereof (which divides by the number of data points) makes this analysis possible, all be it with the enduring caveat that it utilizes sparse data. I call this new variable $\Delta RD$, which I interpret as combined specific ROS generation and damage rates.

For these same species, I sourced adult body mass, maximal lifespan, and specific basal metabolic rate data from [1]. Specific basal metabolic rate data absent in [1], for *Felis catus* was sourced from [289], and for *Ovis aries* and *Canis lupus familiaris* was sourced from earlier Table S2 herein, with sources disclosed in its corresponding text. I couldn't find specific basal metabolic rate data (although I could find adult body mass and maximal lifespan data in [1]) for *Equus caballus*, *Giraffa camelopardalis*, *Mustela furo* (i.e., domesticated *Mustela putorius*), *Castor canadensis*, *Megaptera novaeangliae*, *Balaena mysticetus*, and *Lemur catta*. So, for any calculation using $\Delta RD$ and specific basal metabolic rate, these species are absent.

Variable $\Delta RD$ is a positive linear function of specific basal metabolic rate (*Pearson = 0.8393, one-tailed {alternative hypothesis is directional} p-value = 0.0000125. Spearman's = 0.7132*). Variable $\Delta RD$ is a supra-linear (negative convex) function of adult body mass (body masses sourced from [1]. *Pearson = -0.2598. Spearman's = -0.7043, one-tailed {alternative hypothesis is directional} p-value = 0.000061*). So, variable $\Delta RD$ correlates better with specific basal metabolic rate than it does negatively with adult body mass. Maximal lifespan is a supra-linear (negative convex) function of $\Delta RD$ (*Pearson = -0.5126. Spearman's = -0.8983, one-tailed {alternative hypothesis is directional} p-value = 1.2881E-09. Spearman's coefficient is greater than Pearson here because Spearman's, unlike Pearson, doesn't penalize supra-linearity in its coefficient. With human excluded: Spearman's = -0.8844*). These interrelations, and others, are shown explicitly or implicitly in Figure 14.

Table S83 below shows the result of mediation analysis between specific basal metabolic rate and maximal lifespan, with Mediator variable $\Delta RD$. Conducted in JASP software (version 0.18.1.0), selecting its bootstrap method called "Percentile", run with 10,000 replications, utilizing its (aforementioned) standardized estimates option, which makes each variable mean = 0 and standard deviation = 1. The upper line is with human excluded, and the lower line is with human included (*for maximal lifespan, and for the underlying calculation of [I], which uses maximal lifespan as input, wherein [I] is componentry to* Δ*RD*).

[Table S83]

| | | | 95% Confidence Interval (CI) | | | 95% Confidence Interval (CI) | |
|---|---|---|---|---|---|---|---|
| α | β | **Indirect effect (α·β)** | Lower | Upper | **Direct effect (τ)** | Lower | Upper |
| 233 | -0.93 | -216.936 | -491.5 | -83.299 | 17.576 | -273.101 | 178.694 |
| 238 | -0.67 | -160.793 | -574.49 | -52.942 | 44.003 | -145.694 | 336.571 |

Zero (0) is not within the 95% Confidence Interval (CI) for an indirect effect, but it is for a direct effect, which is consistent with specific basal metabolic rate dictating, via its dictation of Δ*RD*, maximal lifespan.

Simplification of Figure 14

Figure S10 is a simplification of Figure 14, wherein variables in Figure 14 that I interpret relate to metabolic damage (specific DNA damage rate, specific mtDNA damage rate, specific protein damage rate, specific lipid damage rate, specific epigenetic damage rate) are collapsed into a single variable called "specific metabolic damage rate", symbolized as *ΔD* in this corresponding text.

Unlike earlier variable *ΔRD*, *ΔD* doesn't utilize (aforementioned) variable [A]. Using the data set used to produce Table S82 above, I omitted species that didn't have data for at least two variables from the selection of variables [B] through to [I]. Then, using this 24 species subset, each of variables [B] to [I] were standardized (to be within the range of 0-100) using Eq. S1. Thereafter a new variable was made, populated with, for each species used, the mean of standardized [B] to [I]. This variable being *ΔD*.

For these same 24 species, I sourced maximal lifespan, adult body mass, and specific basal metabolic rate data from [1], but the specific basal metabolic rate value used for *Felis catus* (absent in [1]) was from [289], and the specific basal metabolic rate values used for *Ovis aries* and *Canis lupus familiaris* (absent in [1]) were as in earlier Table S2 herein, with sources disclosed in its corresponding text. I couldn't find specific basal metabolic rate data (although I could find maximal lifespan and adult body mass data in [1]) for *Equus caballus*,

*Giraffa camelopardalis*, *Mustela furo* (i.e., domesticated *Mustela putorius*), *Castor canadensis*, *Megaptera novaeangliae*, *Balaena mysticetus*, and *Lemur catta*. So, for any calculation using $\Delta D$ and specific basal metabolic rate, these species were omitted.

Maximal lifespan vs. $\Delta D$ is plotted in Figure S11. I hypothesize that $\Delta D$ *is* aging rate. Where $\Delta D$ is the mean of standardized specific metabolic damage rates, to biological macromolecules/information such as DNA, mtDNA, lipids, proteins, and the epigenome. And possibly to other biological macromolecules/information that I don't have data for, but which (I predict) could be, after equivalent standardization, integrated into $\Delta D$ without changing the form of its interrelations shown in Figure S10. Such as metabolic damage to RNA, carbohydrates (e.g., glycans), telomeres, etc. Because, to predict, they have the same interrelations. For predictive example, across species, maximal lifespan is a supra-linear (negative convex) function of each of them also.

Indeed, for telomere shortening rate, across species, this relation form to maximal lifespan has already been observed [341]. [341] reports, across species, that maximal lifespan is a supra-linear (negative convex) function of telomere shortening rate (base pairs/year). Similarly, across species, maximal lifespan is also such a shaped function of $\Delta D$ (Figure S11), and of each of its components (Figure 14). But (unfortunately) there weren't enough species of overlap for me to use the data of [341] herein.

ROS are a known drive to telomere shortening [342-346] (and senescence [347-356]). So, selectively inhibiting $F_1F_0$ ATP hydrolysis (*without inhibiting $F_1F_0$ ATP synthesis*) - e.g., by a drug in one or more of my international (PCT) patent applications [18-20] – is herein predicted to slow telomere shortening. And delay senescence.

Reducing the number of senescent cells by reducing their generation is likely preferable to killing them. Because there isn't consensus on what constitutes a senescent cell, and whether selectively killing them is *net* beneficial or harmful, or even whether such selectivity is possible. Refer to later Table S124 for more on this.

To summarize/formalize: [Eq. S9]

$$\Delta D = \frac{A + B + C + D + E \ldots}{n}$$

Where *ΔD* is (standardized) specific metabolic damage rate, which is the mean of standardized (to be between 0-100, by Eq. S1) specific metabolic damage rates, where *A* is standardized specific DNA damage rate, *B* is standardized specific mtDNA damage rate, *C* is standardized specific lipid damage rate, *D* is standardized specific protein damage rate, and *E* is standardized specific epigenetic damage rate, *n* is the number of terms in the numerator, and the equation is open-ended so that it can include further standardized specific metabolic damage rates. It can accommodate more than one data set for one or more of the standardized specific metabolic damage rate types (as was the case here: e.g., variables [D] and [F] both relate to mtDNA damage, and both were componentry to *ΔD*).

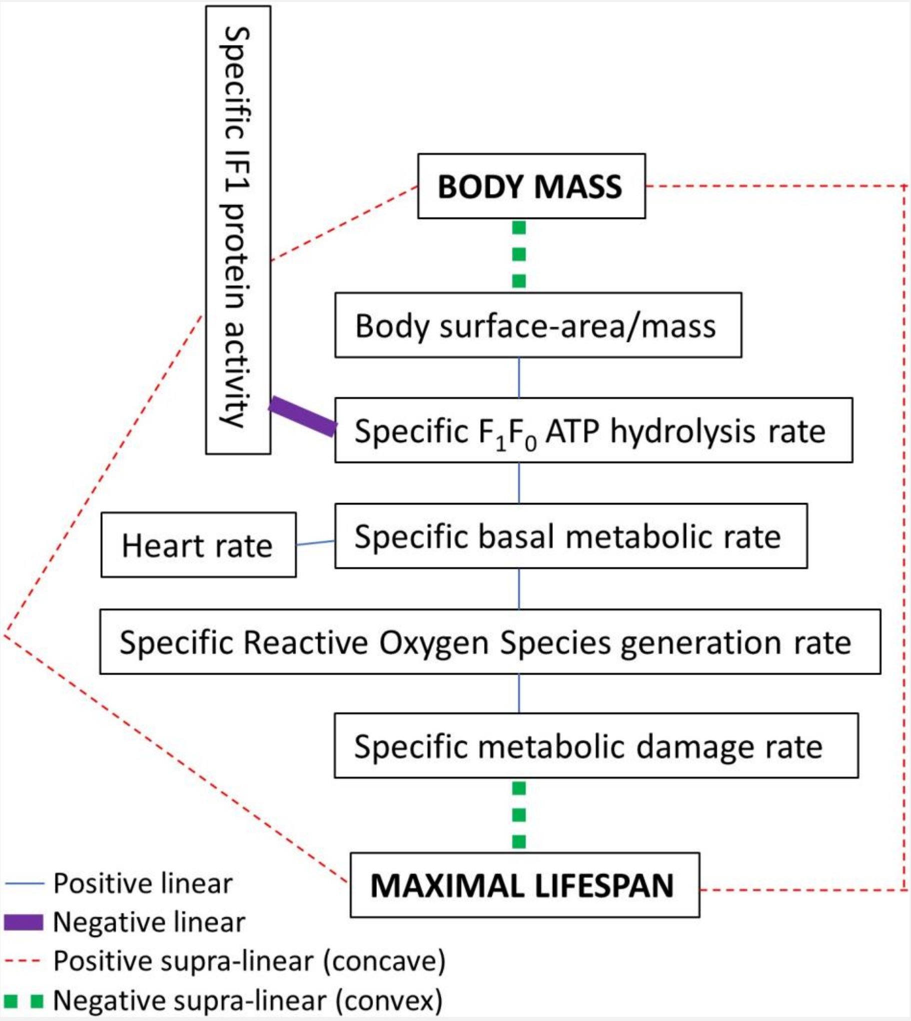

Figure S10 – Where specific metabolic damage rate, symbolized as *ΔD* in accompanying text, is combination of at least specific DNA damage rate, specific mtDNA damage rate, specific protein damage rate, specific lipid damage rate, and specific epigenetic damage rate (*and*

*possibly of other specific metabolic damage rates, to further biological macromolecules/information, which aren't included in ΔD herein because of lack of data*). "Specific" in the figure means per unit mass, or otherwise controlled for body mass. "Body surface-area/mass" is body surface-area per unit body mass. For each interrelation shown, the higher box is the independent (*x*-axis), and the lower box is the dependent (*y*-axis), variable. Each supra-linear function can be made linear by logging the data of the *x*- and *y*-axis. I interpret this pattern as each higher box causes/determines its lower box, wherein the highest box's determination of the second is because of the square-cube law of geometry, from which the rest cascades, wherein a (by-)product(s) of metabolic rate causes damage (*ΔD*), which *is* aging.

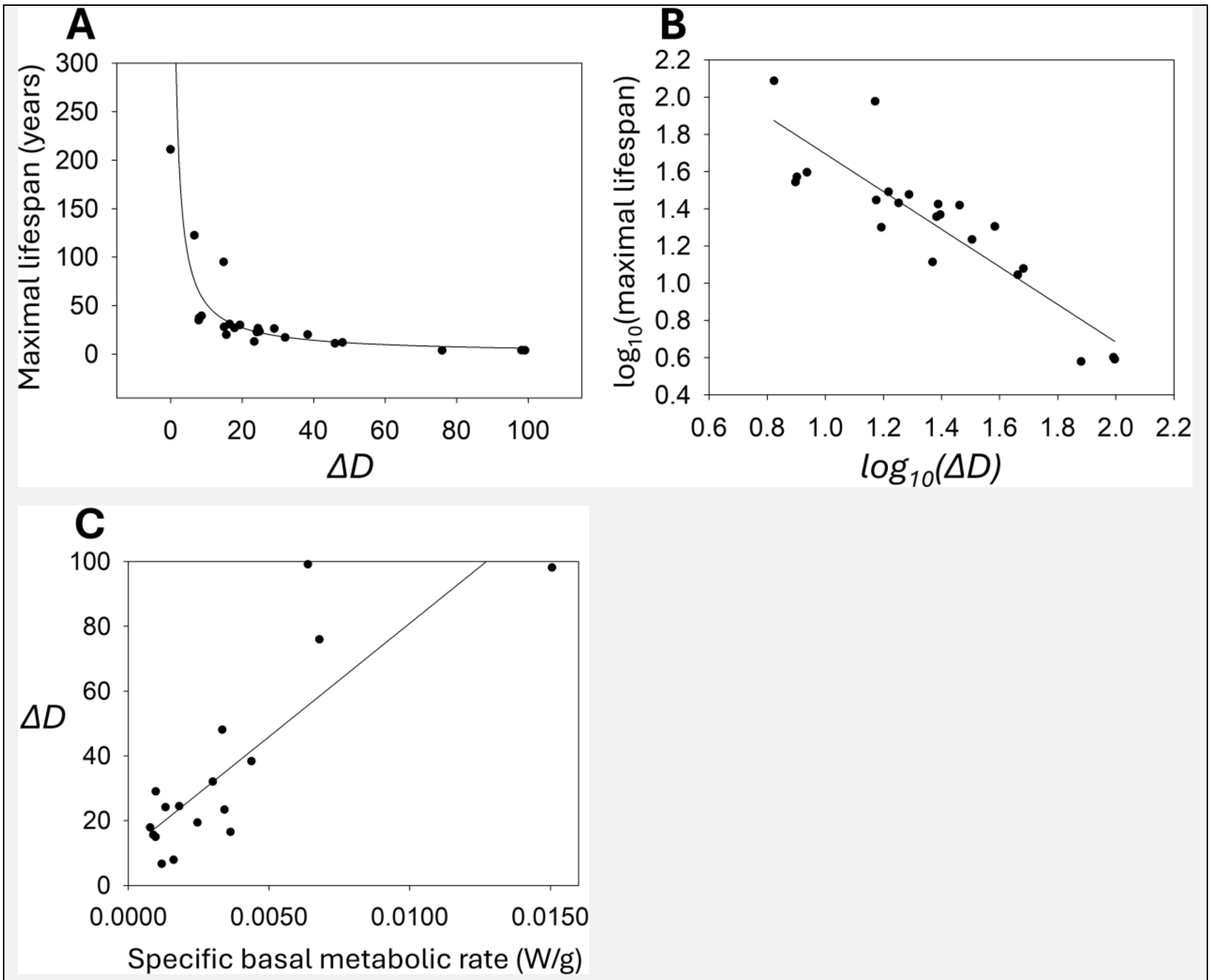

Figure S11 – **(A)** Maximal lifespan (years) vs. *ΔD*. Adjusted $R^2$ = 0.5194. Best-fit line: $maximal\ lifespan = 439.8046 \cdot \Delta D^{-0.9282}$. **(B)** $\log_{10}$-$\log_{10}$ plot of maximal lifespan (years) vs. *ΔD*. Adjusted $R^2$ = 0.7747. Best-fit line: $log_{10}(maximal\ lifespan) = 2.7075 +$

$-1.0118 \cdot log_{10}(\Delta D)$. **(C)** *ΔD* vs. specific basal metabolic rate (W/g). Adjusted $R^2 = 0.6971$. Best-fit line: $\Delta D = 10.9579 + 6998.1933 \cdot (specific\ basal\ metabolic\ rate)$.

Plots and best-fit lines made in SigmaPlot 15.0 software (Systat Software, Inc., San Jose, California, USA). Human data included. Bowhead whale data excluded from $log_{10}$-$log_{10}$ plot because its value of *ΔD* is negative (-4.46038E-08, not precisely zero because of rounding error) and so can't be logged.

*P.S. an error with plots (A) and (B) above is that the data point for Equus caballus was accidentally omitted from these plots, which I can't fix now as I no longer have access to SigmaPlot software. Its inclusion wouldn't change the plots much. For example, replicating plot (B) in OriginPro 8.0 software instead, with Equus caballus data included, has adjusted $R^2$ value of 0.78624 (i.e., higher) with best-fit line: $log_{10}(maximal\ lifespan) = 2.68696 + -0.99886 \cdot log_{10}(\Delta D)$, which has the best-fit line of (B) above within its standard errors (2.68696 ± 0.15347, -0.99886 ± 0.11036). Equus caballus is not an outlier: it very nearly lies upon the best-fit line.*

Correlations basis to Figure S10

Only correlations to *ΔD* are shown in Table S84 below because the other correlations have been presented earlier for Figure 14. The (aforementioned) full data set comprising data from 384 mammal species is used here. *N* is the number of species used (constrained in each case by the number of species that overlap between the compared data sets), *R* is the Pearson correlation coefficient, Rho ($\rho$) is the Spearman's rank correlation coefficient, *P* is the one-tailed (alternative hypothesis is directional) p-value, *A* is adult body mass (data from [1]), *B* is body surface-area per unit body mass (data from [128]), *C* is specific IF1 protein activity (data from [91]), *D* is specific $F_1F_0$ ATP hydrolysis rate (data from [91]), *E* is specific basal metabolic rate (data from [1], plus value for *Felis catus* from [289], and for *Ovis aries* and *Canis lupus familiaris* as given in earlier Table S2, with sources disclosed in its corresponding text), *F* is heart rate (data from [91]), *G* is specific Reactive Oxygen Species (ROS) generation rate (mean of superoxide and hydrogen peroxide data, for species for which both are provided, from [95]), and *M* is maximal lifespan (data from [1]). If human data was present, it wasn't omitted when calculating any of these correlations. Specifically, human was included for maximal lifespan, and for the underlying calculation of the epigenetic clock ticking rate, variable [I], component to Δ*D*.

[Table S84]

| Pair | N | R | P | Rho (ρ) | P |
|---|---|---|---|---|---|
| $\Delta D$-A | 24 | -0.2596 | 0.110284 | -0.7043 | 0.000061 |
| $\Delta D$-B | 11 | 0.9415 | 0.0000075 | 0.7455 | 0.0042245 |
| $\Delta D$-C | 10 | -0.8598 | 0.000711 | -0.4788 | 0.0807555 |
| $\Delta D$-D | 10 | 0.9091 | 0.0001335 | 0.7455 | 0.0066605 |
| $\Delta D$-E | 17 | 0.8462 | 0.0000095 | 0.7108 | 0.000691 |
| $\Delta D$-F | 10 | 0.9351 | 0.000036 | 0.9152 | 0.000102 |
| $\Delta D$-G | 7 | 0.9473 | 0.000595 | 0.9643 | 0.000227 |
| $\Delta D$-M | 24 | -0.5128 | 0.005198 | -0.9165 | 1.60465E-10 |

All the correlations are *significant* (at $P < 0.05$) with Pearson and/or Spearman's rank correlation coefficients. Unlike the others, which are linear, $\Delta D$-A and $\Delta D$-M are supra-linear relations (negative concave), which is why their Spearman's rank correlation coefficient is greater their Pearson correlation coefficient.

Parameter values for $\Delta D$ as $x$ in Eq. 16

The data used to generate Table S84 in the last section was used to generate the following tables (same A-G letter code is used), where $N$ is number of data points and Adj. $R^2$ is adjusted $R^2$. For logged data, bowhead whale data excluded because its value of $\Delta D$ is negative (-4.46038E-08, not precisely zero because of rounding error) and so can't be logged. Parameters determined using OriginPro 8 software (OriginLab Corporation, Northampton, MA, USA. Using its "Fit Linear" functionality, which uses linear regression, with least squares estimation).

[Table S85]

| $x$ | N | Adj. $R^2$ | $\log_{10}(a)$ | $b$ | N | Adj. $R^2$ | $\log_{10}(c)$ | $d$ |
|---|---|---|---|---|---|---|---|---|
| $\Delta D$ | 23 | 0.36575 | 1.47979 ±0.06729 | -0.13746 ±0.03716 | 23 | 0.78624 | 2.68696 ±0.15347 | -0.99886 ±0.11036 |

[Table S86]

| $x$ | $q$ | N | Adj. $R^2$ | $e$ | $g$ |
|---|---|---|---|---|---|
| $\Delta D$ | B | 11 | 0.87391 | 13.53601 ±3.9139 | 210.39758 ±25.09263 |

| | | | | | |
|---|---|---|---|---|---|
| *ΔD* | C | 10 | 0.70659 | 98.68538 ±13.08545 | -19.85674 ±4.17005 |
| *ΔD* | D | 10 | 0.80469 | -15.29836 ±10.69484 | 30.21278 ±4.89599 |
| *ΔD* | E | 17 | 0.69711 | 10.95788 ±5.48931 | 6998.19294 ±1137.8818 |
| *ΔD* | F | 10 | 0.85873 | 5.73289 ±6.5424 | 0.16149 ±0.02164 |
| *ΔD* | G | 7 | 0.87675 | -2.96324 ±9.92377 | 65.98831 ±9.98426 |

<u>Instrumental variable analysis reports causality between lower *ΔD* and higher maximal lifespan</u>

Shown in Table S87 below: result of instrumental variable analysis with maximal lifespan as the dependent variable, *ΔD* as the independent variable, and specific basal metabolic rate as the instrumental variable. Which reports causality between lower *ΔD* and higher maximal lifespan.

The coefficient for the effect of the independent variable on the dependent variable, *β*, is a Two-Stage Least Squares (TSLS) estimate. *E* is the standard error in *β*. *P* is the one-tailed (alternative hypothesis is directional) p-value for *β*. Causality can be inferred from it being significant (at $P < 0.05$). *A* is the p-value for the Anderson-Rubin test (under F distribution) and *C* is its 95% confidence interval. *A2* is the p-value for the conditional likelihood ratio test (under Normal approximation) and *C2* is its 95% confidence interval. R code used is in the supplementary methods section of this Supplementary Material document. Human data excluded by rationale in the Methods.

[Table S87]

| *β* | E | P | A | C | A2 | C2 |
|---|---|---|---|---|---|---|
| -0.29599 | 0.05265 | 0.0000315 | 0.0018419 | -0.411565647573926, -0.16862443940339 | 0.0018419 | -0.411565837533968, -0.168624208703323 |

Is specific basal metabolic rate an appropriate instrumental variable? Mediation analysis reports yes. Mediation analysis with the data used to generate Table S87 above, its results are in Table S88 below, reports that specific basal metabolic rate only affects maximal lifespan through *ΔD*, with *no* direct effect upon maximal lifespan. This mediation analysis was conducted in JASP software (version 0.18.1.0), selecting its bootstrap method called "Percentile", run with 10,000 replications, utilizing its standardized estimates option, which makes each variable mean = 0 and standard deviation = 1.

[Table S88]

| | | | 95% Confidence Interval (CI) | | | 95% Confidence Interval (CI) | |
|---|---|---|---|---|---|---|---|
| α | β | **Indirect effect (α·β)** | Lower | Upper | **Direct effect (τ)** | Lower | Upper |
| 235 | -0.96 | -225.098 | -501.619 | -80.323 | 25.738 | -268 | 189.117 |

Zero (0) is not within the 95% Confidence Interval (CI) for an indirect effect, but it is for a direct effect. This reports that specific basal metabolic rate only affects maximal lifespan through *ΔD*, with *no* direct effect upon maximal lifespan. So, to repeat, specific basal metabolic rate *is* an appropriate instrumental variable here.

Linear regression of maximal lifespan on *ΔD*

Linear regression was performed, with the response variable being maximal lifespan (years; data from [1]) and the predictor variable being *ΔD*. Both logged ($log_{10}$). Human included. Bowhead whale excluded because its value of *ΔD* is negative (-4.46038E-08, not precisely zero because of rounding error) and so can't be logged.

$R^2 = 0.796$, adjusted $R^2 = 0.786$, Unstandardized coefficients: $\beta_0$ (intercept) = 2.687 ± 0.153 (p-value = 5.268e-14), $\beta_1$ = -0.999 ± 0.110 (p-value = 1.078e-8). *Standardized coefficient:* $\beta_1 = -0.892$. *Durbin-Watson statistic = 2.172. F (1, 21) = 81.920, p = 1.078e-8. Assumptions of homoscedasticity, normality, and linearity unviolated.*

[Eq. S10]

$$log_{10}(maximal\ lifespan) = 2.687 + [-0.999 \cdot log_{10}(\Delta D)]$$

Pearson correlation coefficient between $log_{10}$ maximal lifespan and $log_{10}$ *ΔD* = -0.8922, one-tailed (alternative hypothesis is directional) p-value = 5.36871E-09.

If $log_{10}$ adult body mass (kg; data from [1]) is added as a second predictor, the adjusted $R^2$ becomes 0.806 (i.e., only 0.02 more), wherein $log_{10}$ *ΔD* is a highly significant predictor and

$log_{10}$ adult body mass is *not*, with unstandardized regression coefficient p-values of 3.947e-6 and 0.090 respectively (*VIF* = 1.652).

<u>Structural Equation Model (SEM) of Figure 14</u>

I built a SEM model of Figure 14 in JASP software (version 0.18.1.0; JASP utilizes the lavaan R package [94] for SEM). Earlier herein, when investigating correlation basis to Figure 14, data from 384 mammal species was sourced. But then data from species that don't have data available for each of adult body mass, maximal lifespan, and specific basal metabolic rate in [1] was dismissed (350 mammal species have *all* such data in [1]), with the exception that cat, sheep, and dog weren't excluded, and specific basal metabolic rate values for them were sourced elsewhere. Rendering data from 353 mammal species used. When building this SEM model of Figure 14, there was no such data omission, and so data from 384 mammal species was used. Not all 384 species have data for all the variables. Full-Information Maximum Likelihood (FIML) estimation was used in the SEM model to account for missing data. Using logged ($log_{10}$) variables in the SEM model so that *all* the interrelations are linear, a requisite of SEM. The SEM model only contains manifest, and no latent, variables, and it is in lavaan [94] notation (~) below, which must be and was inputted into JASP in this order (dependent variable first):

maximal lifespan ~ specific epigenetic damage rate
maximal lifespan ~ specific DNA damage rate
maximal lifespan ~ specific mtDNA damage rate
maximal lifespan ~ specific lipid damage rate
maximal lifespan ~ specific protein damage rate
specific epigenetic damage rate ~ specific Reactive Oxygen Species (ROS) generation rate
specific DNA damage rate ~ specific Reactive Oxygen Species (ROS) generation rate
specific mtDNA damage rate ~ specific Reactive Oxygen Species (ROS) generation rate
specific lipid damage rate ~ specific Reactive Oxygen Species (ROS) generation rate
specific protein damage rate ~ specific Reactive Oxygen Species (ROS) generation rate
specific Reactive Oxygen Species (ROS) generation rate ~ specific basal metabolic rate
heart rate ~ specific basal metabolic rate
specific basal metabolic rate ~ specific $F_1F_0$ ATP hydrolysis rate
specific $F_1F_0$ ATP hydrolysis rate ~ specific IF1 protein activity
specific IF1 protein activity ~ body surface-area/mass

body surface-area/mass ~ body mass

Table S89 below shows unstandardized regression coefficients (*β*; standard error in brackets), p-values thereof (*β_p*), 95% Confidence Intervals thereof (*Lower* and *Upper* values), and standardized regression coefficients (*s_β*). A-B, A-C, A-D, A-E, A-F, B-G, C-G, D-G, E-G, F-G, G-H, H-I, I-J, J-K, K-L, and L-M, each relate to a line of the model above, where A-B maps to the top line thereof, A-C the second line down, and so on. P-value of 0 means the p-value is so small that JASP rounded it down to 0.

[Table S89]

| | β | β_p | Lower | Upper | s_β |
|---|---|---|---|---|---|
| A-B | -1.335 *(0.043)* | 0 | -1.419 | -1.252 | -0.944 |
| A-C | 0.011 *(0.040)* | 0.780 | -0.067 | 0.089 | 0.011 |
| A-D | -0.047 *(0.040)* | 0.235 | -0.126 | 0.031 | -0.044 |
| A-E | -0.066 *(0.065)* | 0.308 | -0.193 | 0.061 | -0.033 |
| A-F | -0.027 *(0.190)* | 0.888 | -0.399 | 0.346 | -0.005 |
| B-G | 0.690 *(0.077)* | 0 | 0.539 | 0.840 | 0.914 |
| C-G | 0.965 *(0.149)* | 8.278e-11 | 0.674 | 1.257 | 0.906 |
| D-G | 0.877 *(0.142)* | 7.318e-10 | 0.598 | 1.156 | 0.890 |
| E-G | 0.385 *(0.132)* | 0.003 | 0.127 | 0.643 | 0.712 |
| F-G | 0.185 *(0.037)* | 6.529e-7 | 0.112 | 0.258 | 0.845 |
| G-H | 0.625 *(0.073)* | 0 | 0.481 | 0.768 | 0.773 |
| H-I | 0.914 *(0.073)* | 0 | 0.770 | 1.058 | 0.969 |
| I-J | 1.174 *(0.098)* | 0 | 0.981 | 1.367 | 0.946 |
| J-K | -0.618 *(0.046)* | 0 | -0.708 | -0.528 | -0.967 |
| K-L | -0.878 *(0.076)* | 0 | -1.026 | -0.729 | -0.949 |
| L-M | -0.375 *(0.016)* | 0 | -0.407 | -0.344 | -0.952 |

For the model, chi-squared ($\chi^2$) for the baseline test = 338.124 (df = 61), p-value = 5.058e-40 (so model fits significantly better than the baseline model). Goodness of Fit Index (GFI) = 0.518.

In this model, for damage types, the ordering of their standardized regression coefficients (*s_β*) for negative effect on maximal lifespan, from most to least relative effect: specific epigenetic damage rate ($s\_\beta$ = -0.944, $\beta$ = -1.335), specific mtDNA damage rate ($s\_\beta$ = -0.044, $\beta$ = -0.047), specific lipid damage rate ($s\_\beta$ = -0.033, $\beta$ = -0.066), specific protein damage rate ($s\_\beta$ = -0.005, $\beta$ = -0.027), specific DNA damage rate ($s\_\beta$ = 0.011, $\beta$ = 0.011). Of these, specific epigenetic damage rate is the only statistically significant effector upon maximal lifespan. If specific epigenetic damage rate is omitted from the model, this ordering (drastically) changes: specific DNA damage rate ($s\_\beta$ = -0.547, $\beta$ = -0.584), specific protein damage rate ($s\_\beta$ = -0.233, $\beta$ = -1.161), specific lipid damage rate ($s\_\beta$ = -0.255, $\beta$ = -0.466), and specific mtDNA damage rate ($s\_\beta$ = -0.022, $\beta$ = -0.022). Of these, specific DNA damage rate is the only statistically significant effector upon maximal lifespan, and GFI = 0.958. If specific DNA damage rate is then omitted also, the ordering (drastically) changes again: specific mtDNA damage rate ($s\_\beta$ = -0.376, $\beta$ = -0.402), specific lipid damage rate ($s\_\beta$ = -0.363, $\beta$ = -0.640), and specific protein damage rate ($s\_\beta$ = -0.290, $\beta$ = -1.446). Of these, specific mtDNA damage rate is the only statistically significant effector upon maximal lifespan, and GFI = 0.957. When each of these damage types are the only damage type in the model, each is *highly* significant, with $s\_\beta$, $\beta$, and $\beta\_p$ values, with model *GFI* and $\chi^2$ *p-value* values, as follows: specific epigenetic damage rate ($s\_\beta$ = -0.997, $\beta$ = -1.415, $\beta\_p$ = 0, *GFI* = 0.514, $\chi^2$ *p-value* = 7.469e-34), specific mtDNA damage rate ($s\_\beta$ = -0.878, $\beta$ = -0.889, $\beta\_p$ = 0, *GFI* = 0.962, $\chi^2$ *p-value* = 1.070e-32), specific lipid damage rate ($s\_\beta$ = -0.888, $\beta$ = -1.460, $\beta\_p$ = 1.377e-14, *GFI* = 0.955, $\chi^2$ *p-value* = 8.015e-31), specific protein damage rate ($s\_\beta$ = -0.845, $\beta$ = -4.248, $\beta\_p$ = 0, *GFI* = 0.958, $\chi^2$ *p-value* = 9.207e-34), specific DNA damage rate ($s\_\beta$ = -0.914, $\beta$ = -0.948, $\beta\_p$ = 0, *GFI* = 0.970, $\chi^2$ *p-value* = 6.865e-32).

The preceding paragraph showed that the unstandardized regression coefficient ($\beta$) of each of the damage rate variables changes according to which other damage rate variable(s) is included in the model, indicating multicollinearity among these variables. For example, to list the $\beta$ values for specific DNA damage rate mentioned in that paragraph: 0.011, -0.584, -0.948. So, multicollinearity stymies discerning the relative detriment of the different damage types. But some indication can be obtained by comparing standardized regression coefficient values ($s\_\beta$) sourced when each of these damage types are the only damage type in the model. The ordering of which, with $s\_\beta$ values in brackets: specific epigenetic damage rate (-0.997), specific DNA damage rate (-0.914), specific mtDNA damage rate (-0.878), specific lipid damage rate (-0.888), and specific protein damage rate (-0.845). The largest and smallest $s\_\beta$ values of which are -0.997 and -0.845 respectively, a difference of 0.152. Not such a great difference. Consistent with their collective being important, rather than any one or some.

To interpret, all these damage rate variables are contributory to curtailing maximal lifespan, as a collective. Considering a smaller number thereof or just any one as responsible, a reductionist aim, is (I argue) overly reductionist. Teaching that a better model is with these different variables consolidated into a single variable, which is the model of Figure S10, which (indeed, shown in the next section) fits the data better, with a much higher GFI value (0.967 instead of 0.518), with *all* (not only some of) its regression coefficient p-values being *extremely* small. Indeed, all so small that JASP rounds them down to zero (0), except one which is small at 1.643e-14.

A reductionism presently popular among leaders in the longevity field is to place epigenetic change as the sole cause of aging (e.g., [46]), ignoring DNA damage with age/aging, mitochondrial DNA damage with age/aging, etc.

Incidentally, the multicollinearity of these logged ($\log_{10}$) damage rate variables can be observed in the high VIF values when they are regressed upon, in multiple linear regression, with the response variable being $\log_{10}$ maximal lifespan (years). For example, with logged explanatory variables of specific DNA damage rate, specific mtDNA damage rate, and specific protein damage rate, VIF is 10.263, 29.023, and 11.609 respectively. When specific epigenetic damage rate is added too, all the VIFs become NAN (Not a Number), which is consistent with extreme multicollinearity. Incidentally, this also happens when $\log_{10}$ specific basal metabolic rate is added instead. Or $\log_{10}$ adult body mass.

<u>Structural Equation Model (SEM) of Figure S10</u>

Using the same data as the prior SEM model but substituting the damage rate variables with the (aforementioned) single consolidatory damage rate variable, $\Delta D$, I built a SEM model of Figure S10 in JASP software (version 0.18.1.0). Using logged ($\log_{10}$) variables so that *all* the interrelations are linear, which is a requisite of SEM. Bowhead whale omitted because its value of $\Delta D$ is negative (-4.46038E-08, not precisely zero because of rounding error), which can't be logged. Full-Information Maximum Likelihood (FIML) estimation was used to account for missing data. The SEM model only contains manifest, and no latent, variables, and it is in lavaan [94] notation (~) below, which must be and was inputted into JASP in this order (dependent variable first):

maximal lifespan ~ specific metabolic damage rate (*ΔD*)
*ΔD* ~ specific Reactive Oxygen Species (ROS) generation rate
specific Reactive Oxygen Species (ROS) generation rate ~ specific basal metabolic rate
heart rate ~ specific basal metabolic rate
specific basal metabolic rate ~ specific $F_1F_0$ ATP hydrolysis rate
specific $F_1F_0$ ATP hydrolysis rate ~ specific IF1 protein activity
specific IF1 protein activity ~ body surface-area/mass
body surface-area/mass ~ body mass

In Table S90 below, A-B, B-C, C-D, D-E, E-F, F-G, G-H, and H-I, each relate to a line of the model above, where A-B maps to the top line thereof, B-C the second line down, and so on. Column headings as earlier for Table S89.
[Table S90]

| | β | β_p | Lower | Upper | s_β |
|---|---|---|---|---|---|
| A-B | -0.941 *(0.077)* | 0 | -1.093 | -0.789 | -0.890 |
| B-C | 0.973 *(0.115)* | 0 | 0.749 | 1.198 | 0.937 |
| C-D | 0.663 *(0.086)* | 1.643e-14 | 0.494 | 0.833 | 0.842 |
| D-E | 0.885 *(0.072)* | 0 | 0.744 | 1.027 | 0.966 |
| E-F | 1.174 *(0.099)* | 0 | 0.979 | 1.368 | 0.945 |
| F-G | -0.617 *(0.046)* | 0 | -0.708 | -0.527 | -0.967 |
| G-H | -0.875 *(0.076)* | 0 | -1.024 | -0.726 | -0.948 |
| H-I | -0.375 *(0.016)* | 0 | -0.406 | -0.343 | -0.952 |

In Table S90 above, the regression coefficients are of expected sign, highly significant, and with no zero (0) within any 95% confidence intervals. P-value of 0 means the p-value is so small that JASP rounded it down to 0. For the model, chi-squared ($\chi^2$) for the baseline test = 209.745 (df=27), p-value = 3.420e-30 (so model fits significantly better than the baseline model). Goodness of Fit Index (GFI) = 0.967 (i.e., close to 1). This SEM model fits the data very well. Conferring further statistical basis to Figure S10, and its parent Figure 14.

Why are p-values of 0 reported? JASP employs the lavaan R package [94] to perform SEM and, given the R program's default double-precision floating-point arithmetic, its smallest reportable p-value is 2.22e-16. JASP and/or R is rounding down values smaller than that to 0. Incidentally, some R packages use a workaround (various strategies available) to be able to report smaller values, but lavaan isn't one of them. Other R packages (e.g., ivmodel and mgcv) don't round down to 0 but instead report such a small p-value as <2e-16, which is observed frequently herein. At many points, in this and other sections herein, the p-value is smaller than the smallest reportable by the R statistical software. *Highly* statistically significant.

<u>SEM of Figure S10 part II</u>

In the preceding section, the SEM of Figure S10 was built in the SEM module of JASP software (version 0.18.1.0). Because it only contains manifest and no latent variables, its construction could be replicated in the Process (beta version) module of a later version of JASP software (version 0.18.3.0), again using logged ($log_{10}$) data, with FIML to account for missing data, setting all "process types" as "direct" in this module's settings. Reported unstandardized regression coefficients were the same. The reported p-value for the model pathway from adult body mass to maximal lifespan was 0 (also for pathway of adult body mass to heart rate): i.e., so small that JASP rounded down to 0. Distinctly, this other module reports intercept values. The intercept values all had a p-value of zero, except those for specific IF1 protein activity and specific ROS generation rate, which had p-values of 2.494e-6 and 1.168e-11 respectively. Shown below, using the unstandardized regression coefficient and intercept values, the equation chain could be reproduced, with the output of one equation being the input of the next, with adult body mass (kg) as the initial input, and maximal lifespan (years) as the ultimate output (heart rate was in the SEM model but is omitted from the chain below as it isn't on the principal pathway, refer Figure S10).

[Eq. S11]

$$log_{10}\left(\frac{body\ surface\ area}{body\ mass}\right) = -0.375 \cdot log_{10}(body\ mass) + (-0.902);$$

$$log_{10}(specific\ IF1\ protein\ activity) = -0.875 \cdot log_{10}\left(\frac{body\ surface\ area}{body\ mass}\right) + (-0.583);$$

$$log_{10}(specific\ F_1F_0\ ATP\ hydrolysis\ rate) = -0.617 \cdot log_{10}(specific\ IF1\ protein\ activity) + 0.431;$$

$$log_{10}(specific\ basal\ metabolic\ rate) = 1.174 \cdot log_{10}(specific\ F_1F_0\ ATP\ hydrolysis\ rate) + (-2.889);$$

$$log_{10}(specific\ ROS\ generation\ rate) = 0.663 \cdot log_{10}(specific\ basal\ metabolic\ rate) + 1.449;$$

$$log_{10}(\Delta D) = 0.973 \cdot log_{10}(specific\ ROS\ generation\ rate) + 1.768;$$

$$log_{10}(maximal\ lifespan) = -0.941 \cdot log_{10}(\Delta D) + 2.567$$

*Table S91 below shows slope and intercept values, with their standard errors, correspondent to the equation chain (Eq. S11) above.*

*[Table S91]*

| *Slope* | *Standard error* | *Intercept* | *Standard error* |
|---|---|---|---|
| *-0.375* | *0.014* | *-0.902* | *0.027* |
| *-0.875* | *0.107* | *-0.583* | *0.124* |
| *-0.617* | *0.069* | *0.431* | *0.033* |
| *1.174* | *0.142* | *-2.889* | *0.045* |
| *0.663* | *0.084* | *1.449* | *0.214* |
| *0.973* | *0.110* | *1.768* | *0.043* |
| *-0.941* | *0.078* | *2.567* | *0.113* |

Using this equation chain, maximal lifespan (years) is predicted from inputted adult body mass (kg). Replicated in R code below (with a required initial logging, and ultimate de-logging, step added):

x = *input adult body mass (kg) value here*

x=log10(x)

x=-0.375*x+-0.902

x=-0.875*x+-0.583

x=-0.617*x+0.431

x=1.174*x+-2.889

x=0.663*x+1.449

x=0.973*x+1.768

x=-0.941*x+2.567

x=10^(x)

x # *maximal lifespan (years) prediction outputted*

Using an R code variant to the above, which can take multiple adult body mass (kg) values as input, yielding multiple corresponding maximal lifespan (years) predictions as output, presented in the supplementary methods section (of this Supplementary Material document), maximal lifespan was predicted for 1023 mammal species, which is all the mammal species with adult body mass and maximal lifespan values available in [1]. This is a larger mammal species set than the 384 mammal species set whose data was used to generate this SEM model. So, (partially) testing its ability to predict data that wasn't used to generate the model. Figure S12 shows model predicted maximal lifespan values against observed maximal lifespan values. There is good concordance, so good (predominantly) out-of-sample prediction of biological data (*despite most of the species' maximal lifespan values being derived from very small sample sizes, which likely introduces noise, and the sparseness of the data that the model was built from*). Conferring further statistical basis to Figure S10, and its Figure 14 parent.

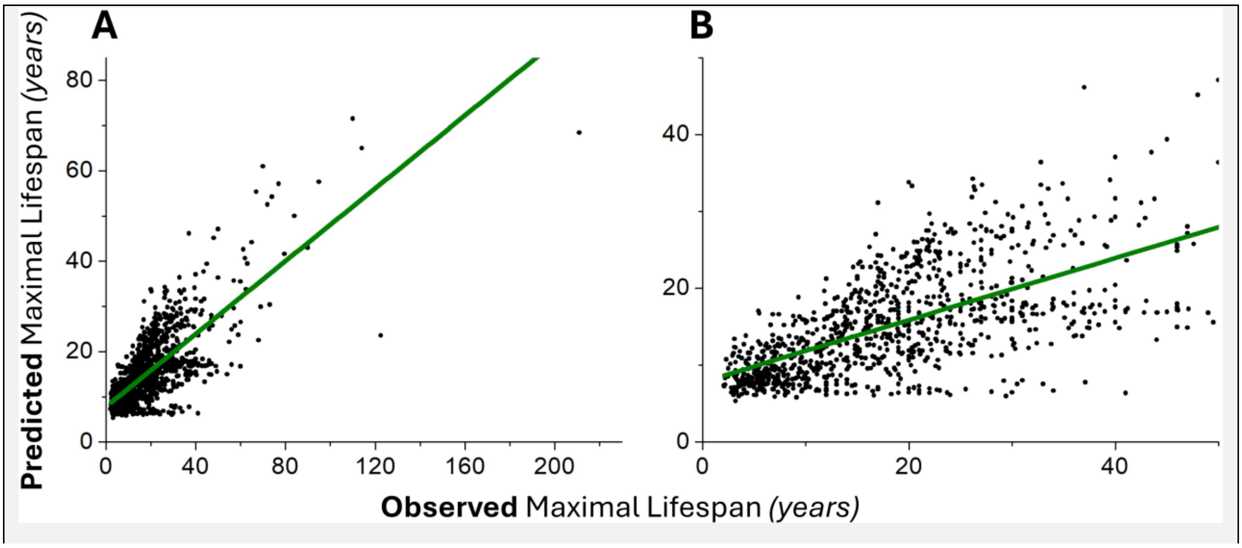

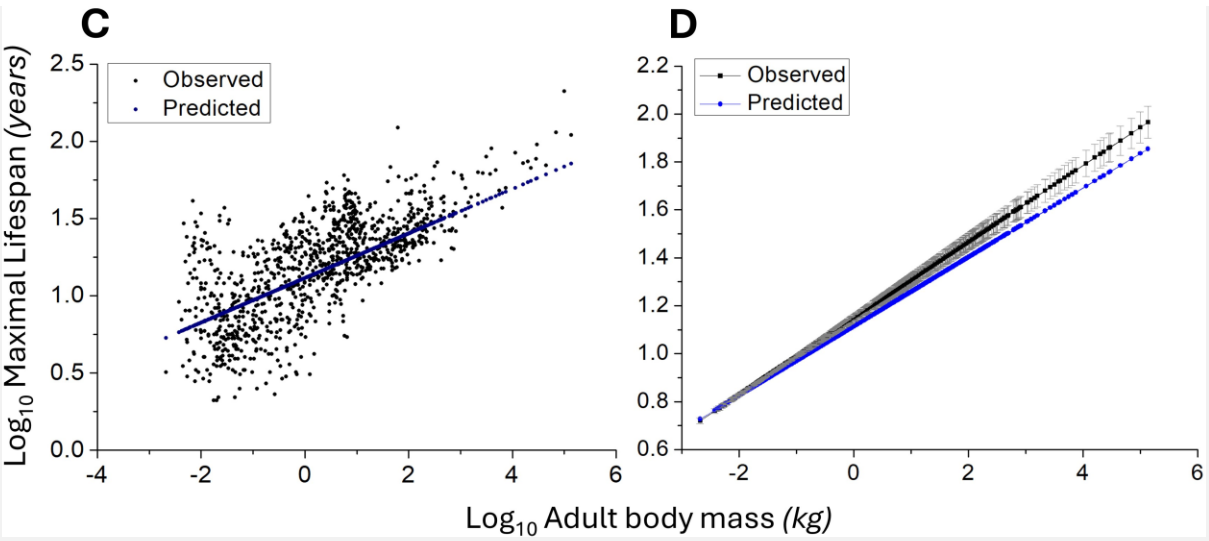

Figure S12 – **(A)** Predicted maximal lifespan (years) by Eq. S11 vs. observed maximal lifespan (years) for 1,023 mammal species. Testing against a much larger species set than the 384 species used to set model parameters. Adult body mass (model input) and maximal lifespan data sourced from [1]. All mammal species with such data available in [1] were used. Best-fit line slope = 0.40236 (±0.01145), intercept = 7.84228 (±0.28901), adjusted $R^2$ = 0.54686, Root Mean Squared Error (RMSE) = 5.7196 years, Mean Absolute Error (MAE) = 7.24517 years. Pearson correlation coefficient = 0.7398, one-tailed (alternative hypothesis is directional) p-value = $\mathbf{3.2661*10^{-178}}$. So, good (predominantly) out-of-sample prediction, conferring further statistical basis to Figure S10, and its parent Figure 14. **(B)** Part of (A) at higher resolution. At least some of the dispersion is likely due to noise in observed maximal lifespans, as most of these values are from extremely small sample sizes. **(C)** $Log_{10}$-$Log_{10}$ plot of observed and model predicted maximal lifespan (years) vs. observed adult body mass (kg). **(D)** $Log_{10}$-$Log_{10}$ plot of observed best-fit line (*y=mx+c, m=0.1594 (±0.00501), c=1.1473 (±0.00725), its error bars are shown in grey*) and model-predicted maximal lifespan (years) vs. observed adult body mass (kg). Model predicted plot (*y=0.14428*x+1.11388*) is nearly completely within the error bars of the observed best-fit line plot. The plots might completely overlap, within their standard errors, if standard errors in the model's parameters were considered also (not done).

*Incidentally, not shown, in-sample predicted vs. observed maximal lifespan, using data from 374 of the 384 species used to build the model, those 374 species with data for both adult body mass and maximal lifespan: best-fit line slope = 0.3798 (±0.01447), intercept = 7.51143 (±0.3658), adjusted* $R^2$ *= 0.64829, RMSE = 5.12583, MAE = 6.69327, and Pearson correlation coefficient = 0.8057. So, comparing RMSE and/or MAE values, the model's* *predominantly out-of-sample prediction is nearly as good as its completely in-sample prediction. The model has good generalisability.*

*Not shown, in-sample prediction when omitting bats, marsupials, naked mole rat, conferring 263 mammal species: best-fit line slope = 0.33018 (±0.02052), intercept = 8.31862 (±0.41832), adjusted $R^2$ = 0.49613, RMSE = 4.30792, MAE = 5.54615, and Pearson correlation coefficient = 0.7057.*

For these 1023 mammal species, how does maximal lifespan predicted by Eq. S11 compare to maximal lifespan predicted by (also ultimately Figure 14 derived) Eq. 20 in the main text? Their Pearson correlation coefficient is 0.9656 (Spearman's rank correlation coefficient = 1, p-value = 0). Best-fit straight line for Eq. S11 predicted maximal lifespan (on *y*-axis) vs. Eq. 20 predicted maximal lifespan (on *x*-axis): slope = 0.47161 (±0.00398), intercept = 8.19694 (±0.0944), adjusted $R^2$ = 0.93226.

Which model is better? For predicted maximal lifespan versus observed maximal lifespan, for Eq. 20, its slope is closer to 1, its adjusted $R^2$ and Pearson correlation coefficient is higher, and it is simpler. Although Eq. S11 has lower RMSE and MAE, and perhaps even lower could be achieved if its parameters were optimized within their standard errors (not done). But most distinctly, Eq. S11 confers (at least easier) access to variables, such as specific IF1 protein activity, which allows us to do interesting experiments *in silico*.

<u>Increased specific IF1 protein activity increases maximal lifespan *in silico*</u>

Below is the earlier presented R code but modified to include a multiplicative constant *q* to modulate specific IF1 protein activity, with de-logging before it is applied and re-logging thereafter. This new component is written in blue text.

```
x = input adult body mass (kg) value here
q = input multiplicative constant to modulate specific IF1 protein activity
x=log10(x)
x=-0.375*x+-0.902
x=-0.875*x+-0.583
x=10^(x)
x=x*q
x=log10(x)
x=-0.617*x+0.431
x=1.174*x+-2.889
```

```
x=0.663*x+1.449
x=0.973*x+1.768
x=-0.941*x+2.567
x=10^(x)
x       # maximal lifespan (years) prediction outputted
```

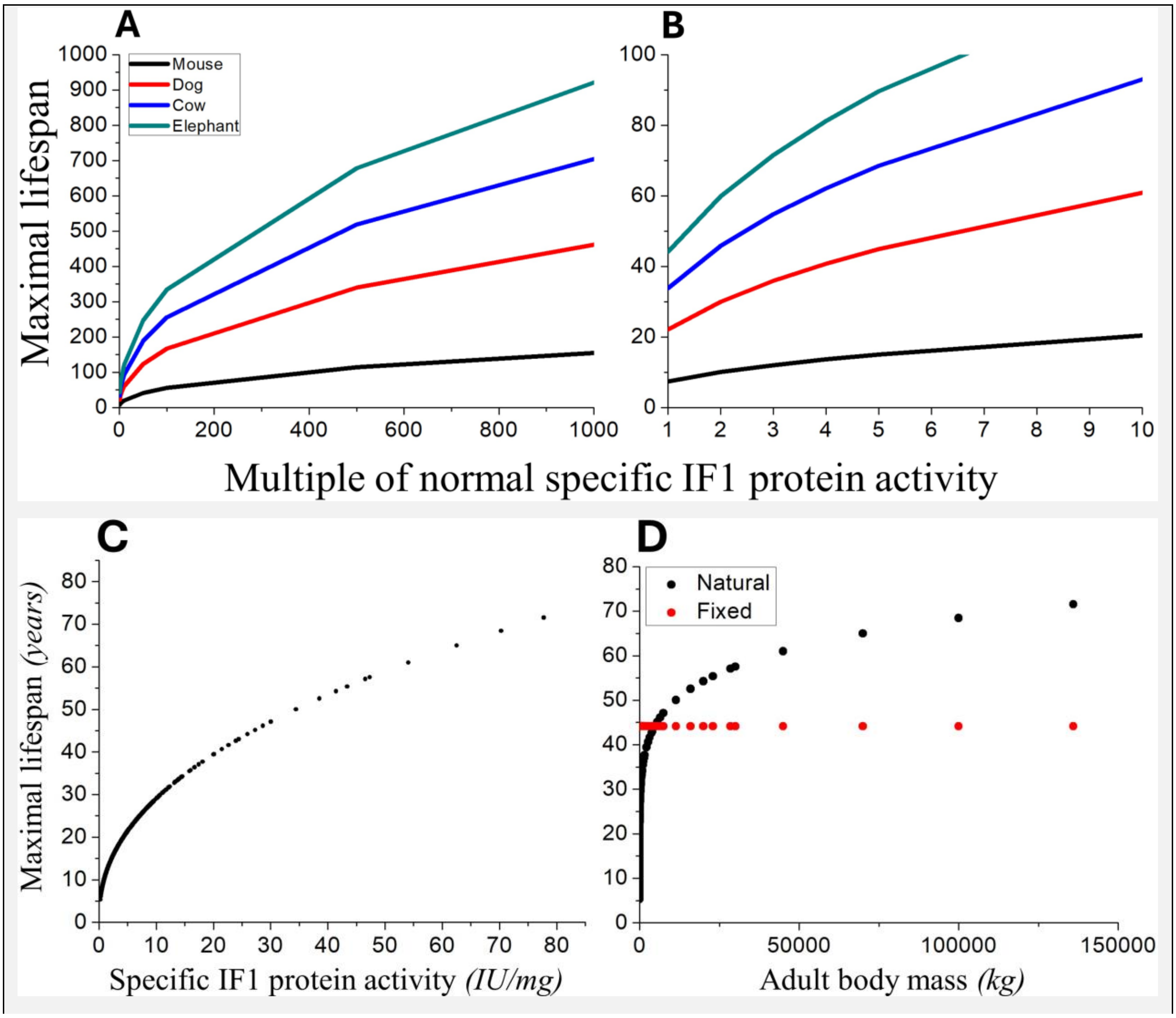

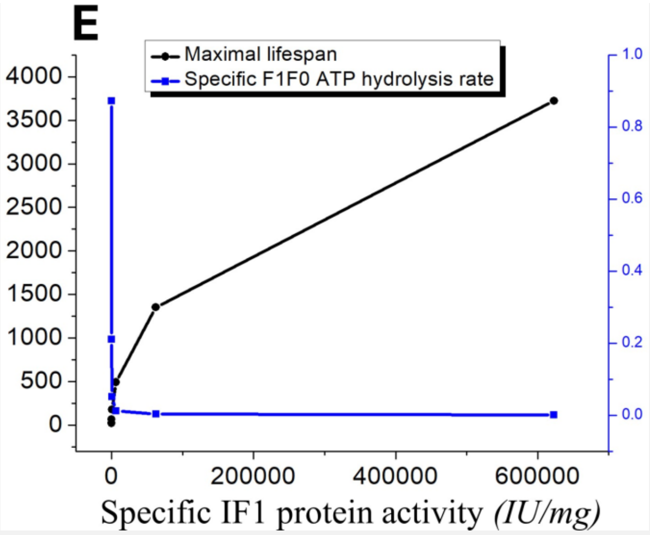

Figure S13 – Model data. **(A)** Increased specific IF1 protein activity increases maximal lifespan *in silico*. Predicted maximal lifespan (years) of a mouse (0.0205 kg), dog (40 kg), cow (750 kg), and African bush elephant (4,800 kg) for different multiples of their normal specific IF1 protein activity (*i.e., different values of multiplicative constant q in the R code*). The gradient is greater for longer-living species because they have more specific IF1 protein activity and so each multiple change of that is greater. The specific IF1 protein activity of mouse at 10x and 60x multiple confers it similar predicted maximal lifespan to a dog and elephant respectively. **(B)** Part of (A) at higher resolution. **(C)** For the 1023 mammal species in the model, their maximal lifespan (years) versus their specific IF1 protein activity (*IU/mg, where 1 Inhibitory Unit, IU, is the amount of inhibitor that inhibits one international unit of enzyme activity: i.e., which reduces $F_1F_0$ ATP hydrolysis by 1 µmol ATP hydrolysed per minute [91]*). **(D)** Predicted maximal lifespan (years) versus adult body mass (kg) for the 1023 mammal species. The black "Natural" plot is when all the 1023 mammal species have their naturally occurring specific IF1 protein activity. The red "Fixed" plot is when all these species are made to have the same specific IF1 protein activity as the African bush elephant, 44.15872 IU/mg, and accordingly all the species have the same maximal lifespan as the African bush elephant (=44 years). **(E)** With two *y*-axes, showing greater specific IF1 protein activity (IU/mg), causing lower specific $F_1F_0$ ATP hydrolysis rate (µmol/min/mg), causes longer maximal lifespan (years). Presenting a computational exploration beyond what is observed in nature (in the diversity of mammals). Finding specific IF1 protein activity values conferring, at least *in silico*, extremely long maximal lifespans. Adult body mass used = 62 kg. Furthest right data point in the graph: specific IF1 protein activity = 622842.9 IU/mg, specific $F_1F_0$ ATP hydrolysis rate = 0.000717577 µmol/min/mg, and maximal lifespan =

3,724.402 years. The model didn't break down with any input and *q* values trialled. Even when very large, even much beyond what is shown in the graph. A robust model.
*N.B., this model disregards any thermoregulatory consequence of modifying specific IF1 protein activity, or any non-consequence when ambient temperature and/or bodily insulation offsets any reduced metabolic heat generation.*

Incorporating postnatal growth rate into the SEM model

To interpret, as briefly discussed in the main text, resting specific metabolic rate isn't only a drive of metabolic damage (≈aging) during adulthood, but during development/growth also, wherein (presumably) a higher growth rate inextricably requires a higher resting metabolic rate per unit mass, and so maximal lifespan should be inversely proportional to growth rate, e.g., postnatal growth rate. Observed in the following SEM model.

The model was constructed in the SEM module of JASP (version 0.18.3.0), again all variables are logged and utilizing FIML, with two extra lines at the top here, pertaining to postnatal growth rate. Postnatal growth rate (days$^{-1}$) data was sourced from [1], which sourced it from [357]. What [1] calls postnatal growth rate, which is consistently referred to as such herein, is the growth rate constant ($K$, days$^{-1}$) in a Gompertz function fitted to body weight vs. age plot (i.e., body weight vs. time plot) during growth. Smaller mammal species tend to have higher $K$ values.

maximal lifespan ~ postnatal growth rate
postnatal growth rate ~ adult body mass
maximal lifespan ~ specific metabolic damage rate ($\Delta D$)
$\Delta D$ ~ specific Reactive Oxygen Species (ROS) generation rate
specific Reactive Oxygen Species (ROS) generation rate ~ specific basal metabolic rate
heart rate ~ specific basal metabolic rate
specific basal metabolic rate ~ specific $F_1F_0$ ATP hydrolysis rate
specific $F_1F_0$ ATP hydrolysis rate ~ specific IF1 protein activity
specific IF1 protein activity ~ body surface-area/mass
body surface-area/mass ~ body mass

In Table S92 below, A-B, B-C, C-D, D-E, E-F, F-G, G-H, H-I, I-J, and J-K, each relate to a line of the model above, where A-B maps to the top line thereof, B-C the second line down, and so on. Column headings as earlier for Table S89.

[Table S92]

| | β | β_p | Lower | Upper | s_β |
|---|---|---|---|---|---|
| A-B | -0.317 *(0.036)* | 0 | -0.387 | -0.247 | -0.443 |
| B-C | -0.293 *(0.015)* | 0 | -0.322 | -0.264 | -0.851 |
| C-D | -0.694 *(0.069)* | 0 | -1.828 | -0.559 | -0.663 |
| D-E | 0.980 *(0.126)* | 8.216e-15 | 0.733 | 1.227 | 0.932 |
| E-F | 0.496 *(0.086)* | 6.493e-9 | 0.329 | 0.664 | 0.632 |
| F-G | 0.924 *(0.074)* | 0 | 0.780 | 1.069 | 0.968 |
| G-H | 1.168 *(0.100)* | 0 | 0.972 | 1.363 | 0.943 |
| H-I | -0.615 *(0.046)* | 0 | -0.705 | -0.524 | -0.966 |
| I-J | -0.866 *(0.076)* | 0 | -1.015 | -0.717 | -0.948 |
| J-K | -0.377 *(0.016)* | 0 | -0.409 | -0.345 | -0.951 |

In Table S92 above, the regression coefficients are of expected sign, highly significant, and with no zero (0) within any 95% confidence intervals. P-value of 0 means the p-value is so small that JASP rounded it down to 0. For the model, chi-squared ($\chi^2$) for the baseline test = 201.618 (df=34), p-value = 1.068e-25 (so model fits significantly better than the baseline model). Goodness of Fit Index (GFI) = 0.965 (i.e., close to 1).

Replicating this model in the Process module (beta) of JASP version 0.18.3.0 permitted obtaining of corresponding intercepts. The intercept values all had a p-value of zero, except those for specific IF1 protein activity and specific ROS generation rate, which had p-values of 3.136e-6 and 6.673e-6 respectively. The reported p-value for the model pathway from adult body mass to maximal lifespan (via the pathway shown in Figure S10) was 1.102e-12. The p-value for the model pathway from adult body mass to heart rate (as shown in Figure S10) was 0. The p-value for the model pathway from adult body mass to maximal lifespan via postnatal growth rate was 2.220e-14.

Shown below, using the unstandardized regression coefficient and intercept values, the equation chain was reproduced, with the output of one equation being the input of the next (*with the exception of the first equation: its output is input to the last equation*), with adult body mass (kg) as the initial input, and maximal lifespan (years) as the ultimate output (heart rate was in the SEM model but is omitted from the chain below as it isn't on the principal pathway, refer Figure S10).

[Eq. S12]

$$log_{10}(postnatal\ growth\ rate) = -0.293 \cdot log_{10}(body\ mass) + (-1.769);$$

$$log_{10}\left(\frac{body\ surface\ area}{body\ mass}\right) = -0.377 \cdot log_{10}(body\ mass) + (-0.903);$$

$$log_{10}(specific\ IF1\ protein\ activity) = -0.866 \cdot log_{10}\left(\frac{body\ surface\ area}{body\ mass}\right) + (-0.576);$$

$$log_{10}(specific\ F_1F_0\ ATP\ hydrolysis\ rate) = -0.615 \cdot log_{10}(specific\ IF1\ protein\ activity) + 0.431;$$

$$log_{10}(specific\ basal\ metabolic\ rate) = 1.168 \cdot log_{10}(specific\ F_1F_0\ ATP\ hydrolysis\ rate) + (-2.885);$$

$$log_{10}(specific\ ROS\ generation\ rate) = 0.496 \cdot log_{10}(specific\ basal\ metabolic\ rate) + 0.969;$$

$$log_{10}(\Delta D) = 0.980 \cdot log_{10}(specific\ ROS\ generation\ rate) + 1.759;$$

$$log_{10}(maximal\ lifespan) = -0.694 \cdot log_{10}(\Delta D) + 1.584 + \{-0.317 \cdot log_{10}(postnatal\ growth\ rate)\}$$

*Table S93 below shows slope and intercept values, with their standard errors, correspondent to the equation chain (Eq. S12) above.*

*[Table S93]*

| *Slope* | *Standard error* | *Intercept* | *Standard error* |
|---|---|---|---|
| *-0.293* | *0.015* | *-1.769* | *0.021* |
| *-0.377* | *0.016* | *-0.903* | *0.027* |
| *-0.866* | *0.076* | *-0.576* | *0.124* |
| *-0.615* | *0.046* | *0.431* | *0.033* |
| *1.168* | *0.1* | *-2.885* | *0.045* |
| *0.496* | *0.086* | *0.969* | *0.215* |

| | | | |
|---|---|---|---|
| *0.980* | *0.126* | *1.759* | *0.048* |
| *-0.694* | *0.069* | *1.584* | *0.155* |
| *-0.317* | *0.036* | | |

Using this equation chain, maximal lifespan (years) is predicted from inputted adult body mass (kg). Replicated in R code below (with a required initial logging, and ultimate de-logging, step added):

```
x = input adult body mass (kg) value here
x=log10(x)
y=-0.293*x+-1.769
x=-0.377*x+-0.903
x=-0.866*x+-0.576
x=-0.615*x+0.431
x=1.168*x+-2.885
x=0.496*x+0.969
x=0.980*x+1.759
x=-0.694*x+1.584+(-0.317*y)
x=10^(x)
x    # maximal lifespan (years) prediction outputted
```

Using an R code variant to the above, replicated in the supplementary methods section of this present document, which can take multiple adult body mass (kg) values as input, yielding multiple corresponding maximal lifespan (years) predictions as output, maximal lifespan was predicted for 1023 mammal species. Which is all the mammal species with adult body mass and maximal lifespan values available in [1]. This is a larger mammal species set than the 384 mammal species set whose data was used to generate this SEM model. So, (partially) testing its ability to predict data that wasn't used to generate the model. Figure S14 shows model predicted against observed maximal lifespan values. Comparing Figures S14 and S12, the predictive power of this SEM model, which incorporates postnatal growth rate, is slightly better than that without it.

Decreasing postnatal growth rate increases maximal lifespan *in silico*

Key maximal lifespan determining levers in the model are specific IF1 protein activity and postnatal growth rate. Higher specific IF1 protein activity increases, and higher postnatal growth rate decreases, maximal lifespan. To interpret, higher specific IF1 protein activity decreases, and (during juvenility) higher postnatal growth rate increases, resting specific metabolic rate, and thereby specific metabolic damage rate.

Below is the earlier presented R code but modified to include a multiplicative constant *k* to modulate postnatal growth rate, with de-logging before it is applied and re-logging thereafter. And with a multiplicative constant *q* to (if desired: as well or instead) modulate specific IF1 protein activity, with de-logging before it is applied and re-logging thereafter. New components are in blue text.

```
x = input adult body mass (kg) value here
k = input here multiplicative constant to modulate postnatal growth rate
q = input here multiplicative constant to modulate specific IF1 protein activity
x=log10(x)
y=-0.293*x+-1.769
y=10^(y)
y=y*k
y=log10(y)
x=-0.377*x+-0.903
x=-0.866*x+-0.576
x=10^(x)
x=x*q
x=log10(x)
x=-0.615*x+0.431
x=1.168*x+-2.885
x=0.496*x+0.969
x=0.980*x+1.759
x=-0.694*x+1.584+(-0.317*y)
x=10^(x)
x       # maximal lifespan (years) prediction outputted
```

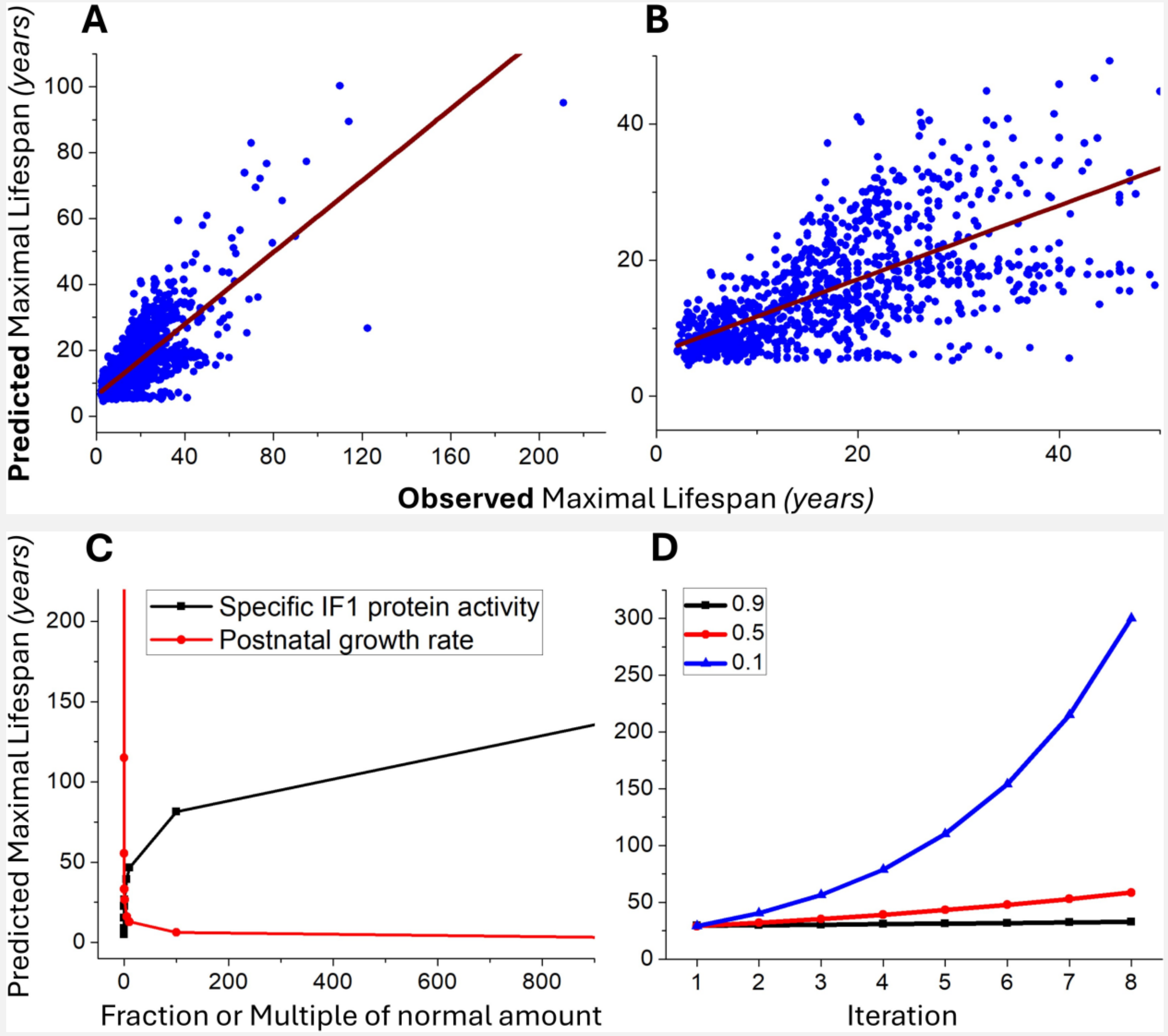

Figure S14 – **(A)** Predicted maximal lifespan (years) by Eq. S12 vs. observed maximal lifespan (years) for 1,023 mammal species. Testing against a much larger species set than the 384 species used to set model parameters. Adult body mass (model input) and maximal lifespan data sourced from [1]. All mammal species with such data available in [1] were used. Best-fit line slope = 0.5437 (±0.01512), intercept = 6.29139 (±0.38162), adjusted $R^2$ = 0.55829, Root Mean Squared Error (RMSE) = 7.55246 years, Mean Absolute Error (MAE) = 7.06259 years. Pearson correlation coefficient = 0.7475, one-tailed (alternative hypothesis is directional) p-value = $\mathbf{6.7845*10^{-184}}$. So, good (predominantly) out-of-sample prediction. **(B)** Part of (A) at higher resolution. **(C)** With body mass of 62 kg, *in silico* exploration of increasing by a multiple, or decreasing by a fraction, the normal amount of either specific IF1 protein activity or postnatal growth rate. The lines cross over when value of one (1) is on the *x*-axis. Reducing postnatal growth rate to fraction thereof, to values between 0 and 1, can steeply increase maximum lifespan. **(D)** The default model mirrors the data in that smaller mammal species have a higher postnatal growth rate. But the model allows us to explore settings not observed within the present diversity of mammals. In the model, decreased body mass can correlate with increased maximal lifespan when occurring with a proportional or super-proportional decrease in postnatal growth rate. Inverting the relationship between body mass and lifespan. Such that, as body mass decreases, maximal lifespan increases. Starting with body mass = 100 kg (number 1 on the *x*-axis) and reducing body mass and postnatal growth rate by multiplying each by the same decimal fraction (0.9, 0.5, or 0.1) for 7 iterations, numbered 2 to 8 on the *x*-axis, increasingly increases maximal lifespan. At the end, for 0.9, 0.5, and 0.1: body mass is 47.82969, 0.78125, and 0.00001 kg respectively, and postnatal growth rate is 0.002621478, 0.0001429561, and 4.965923e-08 days$^{-1}$ respectively. Perhaps relevantly, dwarf mice, which are longer-lived than normal mice [353], have a lower postnatal growth rate [359-360]. Lower postnatal growth rate might explain at least some intra-species inversions of the inter-species positive relation between lifespan and body mass.

Further interrelations discovered

Figure S15 summarizes a further pattern of interrelations I've discovered across mammal species studied, wherein those on its left-hand-side weren't shown in earlier Figure 14. Unlike Figure 14, its boxes aren't necessarily arranged as per (*predicted*) causality/determination (e.g., I don't consider specific basal metabolic rate to dictate maximal lifespan via heart rate). For each interrelation shown, the higher box (*or the equally high box on the right-hand side*) is the independent (*x*-axis), and the lower box (*or the equally low box on the left-hand side*) is the dependent (*y*-axis), variable.

When each of specific IF1 protein activity, specific $F_1F_0$ ATP hydrolysis rate, specific basal metabolic rate, and heart rate (*specific ROS generation rate and specific damage rate also, but these aren't shown*) are multiplied by body mass, each of these new variables (e.g., one being basal metabolic rate: i.e., not per unit mass) is a linear function of body surface-area, and a supra-linear (negative convex) function, of body surface-area per unit body mass. Figure S15 schematizes these, and further, interrelations.

To illustrate an implicit feature of this schema, by using three imaginary illustrating variables called *v1*, *v2* and *v3*: if variable *v1* is some function (one of the four types shown in the schema) of another variable *v2*, wherein *v2* is a positive linear function of a third variable *v3*, then *v1* is the same form of function (that it is of *v2*) of *v3* also. For example, Body surface-area/mass is a negative supra-linear (convex) function of Body mass, wherein Specific $F_1F_0$ ATP hydrolysis rate is a positive linear function of Body surface-area/mass, meaning Specific $F_1F_0$ ATP hydrolysis rate is a negative supra-linear (convex) function of Body mass also. Or in another example, Specific basal metabolic rate is a positive linear function of Specific $F_1F_0$ ATP hydrolysis rate, wherein Heart rate is a positive linear function of Specific basal metabolic rate, meaning Heart rate is a positive linear function of Specific $F_1F_0$ ATP hydrolysis rate. Or in a further example, which crosses the gap in the middle of the schema, $F_1F_0$ ATP hydrolysis rate is a negative supra-linear (convex) function of Body surface-area/mass, wherein $F_1F_0$ ATP hydrolysis rate is a positive linear function of Body surface-area, meaning Body surface-area is a negative supra-linear (convex) function of Body surface-area/mass also. Or in a last example, Maximal lifespan is a positive supra-linear (concave) function of Heart rate*Body mass, wherein IF1 protein activity is a positive linear function of the latter (connected to it by a chain of positive linear relations), and so Maximal lifespan is a positive supra-linear (concave) function of IF1 protein activity.

So, because of this implicit character to the schema, the number of interrelations that this schema discloses is greater than the relatively small number of connecting lines it shows.

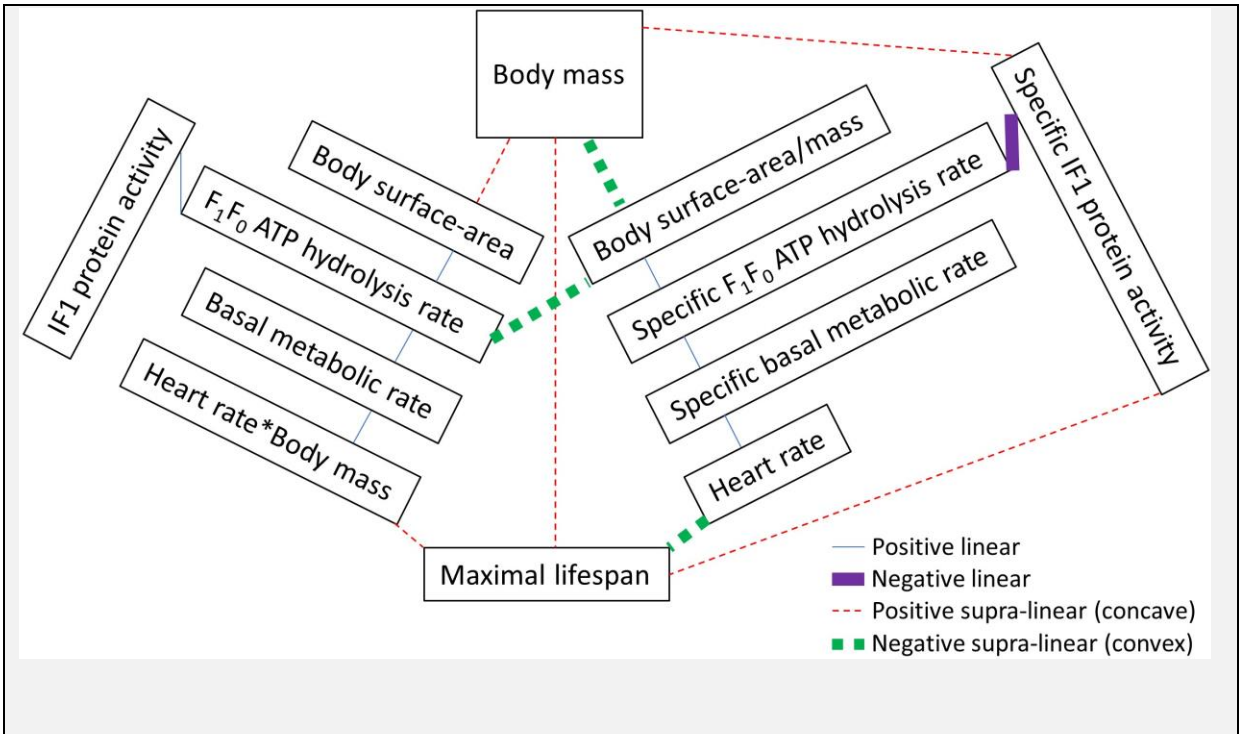

Figure S15 – Using data from 377 mammal species, this pattern of interrelations is observed across mammal species. "Specific" in the figure means per unit mass. For each interrelation shown, the higher box is the independent (*x*-axis), and the lower box is the dependent (*y*-axis), variable. When two boxes have the same height in the schema, the box on the right is the independent (*x*-axis), and the box on the left is the dependent (*y*-axis), variable. Each supra-linear function can be made linear by logging the *x*- and *y*-axis data.

Correlations basis to Figure S15

Only correlations not presented earlier for Figure 14 are shown below. *N* is the number of species used (constrained in each case by the number of species that overlap between the compared data sets), *R* is the Pearson correlation coefficient, Rho ($\rho$) is the Spearman's rank correlation coefficient, *P* is the one-tailed (alternative hypothesis is directional) p-value, *A* is body surface-area (data from [128]), *B* is IF1 protein activity (specific IF1 protein activity data from [91] multiplied by body mass data from [91]), *C* is $F_1F_0$ ATP hydrolysis rate (specific $F_1F_0$ ATP hydrolysis rate data from [91] multiplied by body mass data from [91]), *D* is basal metabolic rate (data from [1], for the 350 mammal species that have such data, plus adult body mass and maximal lifespan data, there. To get basal metabolic rate values for *Felis catus*, *Ovis aries*, and *Canis lupus familiaris*, which are absent in [1], each of their specific basal metabolic rate values, sourced as specified later in this paragraph, were multiplied by their respective adult body mass values from [1]), *E* is heart rate*body mass (heart rate data from [91] multiplied by body mass data from [91]. A contrived variable), *F* is adult body mass (data from [1]), *G* is maximal lifespan (data from [1]), *H* is body surface-area per unit body mass (data from [128]), *I* is specific IF1 protein activity (data from [91]), *J* is specific $F_1F_0$ ATP hydrolysis rate (data from [91]), *K* is specific basal metabolic rate (data from [1], for those 350 mammal species that have such data, plus adult body mass and maximal lifespan data, in [1]. For the following species, specific basal metabolic rate values absent in [1] were sourced elsewhere: *Felis catus* by value from [289], and for *Ovis aries* {sheep} and *Canis lupus familiaris* {dog} as given in earlier Table S2, with sources disclosed in its corresponding text), and *L* is heart rate (data from [91]).

To populate variables *F* and G, data from [1] was pulled for those 350 mammal species that have data for all three of specific basal metabolic rate, adult body mass and maximal lifespan there. As available, adult body mass and maximal lifespan data was then pulled for 24 additional identifiable species (ignoring the 2, as earlier explained, ambiguous species in [128]) from [1], species that have data in variables *A* and *H* (data from [128]), which are (presently) missing one or more of specific basal metabolic rate, maximal lifespan, and adult body mass data in [1]. If human data was present, it wasn't omitted when calculating any of these correlations. Figure S15 schematizes data from 377 mammal species. All be it not all the 377 mammal species have data for all the variables, which is clear in the *N* values in Table S94 below.

[Table S94]

| Pair | N | R | P | Rho (ρ) | P |
|---|---|---|---|---|---|
| A-B | 8 | 0.9958 | 9.23157E-08 | 0.9762 | 0.0000165 |
| A-C | 8 | 0.9927 | 4.83607E-07 | 0.9524 | 0.00013 |
| A-D | 27 | 0.9603 | 1.12851E-15 | 0.922 | 4.25019E-12 |
| A-E | 8 | 0.9967 | 6.96124E-12 | 0.9762 | 0.0000165 |
| B-C | 10 | 0.9968 | 2.28519E-10 | 0.9636 | 0.0000035 |
| B-D | 10 | 0.9669 | 2.52326E-06 | 0.9636 | 0.0000035 |
| B-E | 10 | 0.9918 | 9.79403E-09 | 1 | 0 |
| C-D | 10 | 0.9616 | 0.0000045 | 0.9636 | 0.0000035 |
| C-E | 10 | 0.9849 | 1.1167E-07 | 0.9636 | 0.0000035 |
| D-E | 10 | 0.9867 | 6.73524E-08 | 0.9636 | 0.0000035 |
| F-A | 46 | 0.9575 | 1.08848E-25 | 0.9594 | 4.05858E-26 |
| F-B | 10 | 0.9968 | 2.28519E-10 | 0.9879 | 4.62216E-08 |
| F-B-*2* | 9 | 0.9515 | 0.0000395 | 0.9833 | 0.000001 |
| F-C | 10 | 0.993 | 5.20801E-09 | 0.9515 | 0.0000115 |
| F-C-*2* | 9 | 0.844 | 0.002112 | 0.9333 | 0.000118 |
| F-D | 353 | 0.9782 | 5.3877E-242 | 0.975 | 1.1214E-231 |
| F-D-*2* | 352 | 0.8313 | 1.37391E-91 | 0.9748 | 2.0008E-230 |
| F-E | 10 | 0.9817 | 2.3995E-07 | 0.9879 | 4.62216E-08 |
| G-A | 43 | 0.8374 | 1.28439E-12 | 0.828 | 3.70172E-12 |
| G-A-*2* | 41 | 0.5925 | 0.0000225 | 0.8017 | 1.50175E-10 |
| G-B | 10 | 0.103 | 0.3885315 | 0.8061 | 0.002429 |
| G-C | 10 | 0.1181 | 0.3726145 | 0.7939 | 0.003052 |
| G-D | 353 | 0.3769 | 1.16961E-13 | 0.6339 | 2.2836E-41 |

| | | | | | |
|---|---|---|---|---|---|
| G-E | 10 | 0.1858 | 0.3036525 | 0.8061 | 0.002429 |
| H-A | 53 | -0.1534 | 0.136398 | -0.9242 | 2.8686E-23 |
| H-B | 8 | -0.4238 | 0.1476975 | -0.9762 | 0.0000165 |
| H-C | 8 | -0.4047 | 0.159985 | -1 | 0 |
| H-D | 27 | -0.2989 | 0.064949 | -0.931 | 9.63646E-13 |
| H-E | 8 | -0.4798 | 0.1144535 | -0.9762 | 0.0000165 |
| I-A | 8 | 0.2111 | 0.3078945 | 0.3095 | 0.2278405 |
| I-B | 10 | 0.2383 | 0.253663 | 0.4667 | 0.086952 |
| I-C | 10 | 0.2268 | 0.2643085 | 0.4667 | 0.086952 |
| I-D | 10 | 0.3153 | 0.187428 | 0.503 | 0.0691805 |
| I-E | 10 | 0.2774 | 0.218883 | 0.4667 | 0.086952 |
| J-A | 8 | -0.3883 | 0.1709055 | -0.6667 | 0.035484 |
| J-B | 10 | -0.3593 | 0.1539395 | -0.7212 | 0.009288 |
| J-C | 10 | -0.332 | 0.1743225 | -0.6242 | 0.0268695 |
| J-D | 10 | -0.4887 | 0.075889 | -0.6364 | 0.0239475 |
| J-E | 10 | -0.4352 | 0.104371 | -0.7212 | 0.009288 |
| K-A | 8 | -0.4299 | 0.143873 | -0.9048 | 0.001003 |
| K-B | 10 | -0.3899 | 0.132678 | -0.9636 | 0.0000035 |
| K-C | 10 | -0.376 | 0.142125 | -0.903 | 0.000172 |
| K-D | 353 | -0.1159 | 0.0147335 | -0.8131 | 8.63837E-85 |
| K-E | 10 | -0.4561 | 0.0926055 | -0.9636 | 0.0000035 |
| L-A | 8 | -0.5283 | 0.089158 | -0.9762 | 0.0000165 |
| L-B | 10 | -0.475 | 0.082672 | -0.9758 | 7.28668E-07 |
| L-C | 10 | -0.4604 | 0.0902865 | -0.9758 | 7.28668E-07 |
| L-D | 10 | -0.6171 | 0.028674 | -0.9394 | 0.0000275 |
| L-E | 10 | -0.5511 | 0.0493535 | -0.9758 | 7.28668E-07 |

The asymptotically exact harmonic mean p-value [76] (*its post-published corrected method, as corrected on 7 October 2019*) for the p-values in Table S94 above, when selecting from each line whichever is the smaller p-value (for Pearson or Spearman's rank correlation coefficient), and where there are multiple lines for a pair only using the first line (e.g., using F-B and not F-B-*2*), and substituting the two p-values of value zero with an arbitrarily small value (1e-100) to make it calculable = 2.424465e-240; if the two p-values of zero are omitted instead = 2.316711e-240 = **$2*10^{-240}$**.

Table S94 above excludes correlations shown in Figure S15 that have already been reported upon for Figure 14. The asymptotically exact harmonic mean p-value [76] (*its post-published corrected method, as corrected on 7 October 2019*) of that for Figure 14 ($1*10^{-119}$) and the above ($2*10^{-240}$) = $4*10^{-240}$.

F-B-2 and F-C-*2* use the same data as F-B and F-C respectively except that "leverage point" for *Bos taurus* is omitted. F-D-*2* uses the same data as F-D except that leverage point for *Elephas maximus* is omitted. G-A-*2* uses the same data as G-A except that leverage points for *Balaenoptera physalus* and *Balaenoptera musculus* are omitted. Whereas for F-B, F-C, F-D, and G-A, their Pearson is greater than their Spearman's rank correlation coefficient, the inverse is true for F-B-*2*, F-C-*2*, F-D-*2*, and G-A-*2*. For B-E and C-D, their Spearman's rank is greater than their Pearson correlation coefficient, but not by much, and their Pearson correlation coefficient is very high (0.9918 and 0.9616 respectively), and their Pearson plot appears linear by eye. Aside from these mentioned exceptions (F-B, F-C, F-D, G-A, B-E, C-D), for the correlations in Table S94: when their Pearson is greater than their Spearman's rank correlation coefficient, the relation is linear, and when not, the relation is supra-linear (if a positive correlation, convex, and if a negative correlation, concave).

All the correlations are *significant* (at $P < 0.05$, even despite very low values of $N$ in many cases) with Pearson and/or Spearman's rank correlation coefficients, *except* I-A and I-D, which have sizeable correlations (>0.3), but small values of $N$ (≤10). Arguably underpowered. Variable $I$ is common to both. Refer to the earlier section (headed "IF1 protein content") where the sub-optimal assay used to generate variable $I$ is discussed, teaching that the assay used to generate variable $J$ captures more biology. Variable $I$ correlation to somewhat related (to $A$ and $D$) variables, variables $H$ and $K$, *does* hit statistical significance (shown, but with different letter coding, earlier with Figure 14).

Figure S15 in equations

Figure S15 expressed mathematically (omitting relations already shown by Figure 14): [Eq. S13]:

$$log_{10}(y) = log_{10}(a) + b \cdot log_{10}(x)$$

$$log_{10}(maximal\ lifespan) = log_{10}(c) + d \cdot log_{10}(y)$$
$$y = e + g \cdot q$$

Incidentally, these equations without any logging:

$$y = a \cdot x^b$$
$$maximal\ lifespan = c \cdot y^d$$
$$y = e + g \cdot q$$

Where these scaffold equations are populated with different variables to get the full set of equations. Iterating, *y* is selected in turn from body surface-area, IF1 protein activity, $F_1F_0$ ATP hydrolysis rate, basal metabolic rate, heart rate*body mass, and *x* is (iterating, each different option of *x* selected in turn) selected from body mass, body surface-area/mass, specific IF1 protein activity, specific $F_1F_0$ ATP hydrolysis rate, specific basal metabolic rate, heart rate, *q* is a different selection of *y* (iterating, each different option of *y* selected in turn), and *a*, *b*, *c*, *d*, *e*, and *g* are constants, which can be different for different selections of *y*, *x*, and *q*. When *x* is body mass or specific IF1 protein activity, *b* is positive, otherwise *b* is negative. Note that the first three equations are of the well-known straight-line form: y=c+mx.

The data used to generate Table S94 in the last section was used to generate the following tables (with same A-L letter code used), where *N* is number of data points and Adj. $R^2$ is adjusted $R^2$. Parameters determined using OriginPro 8 software (OriginLab Corporation, Northampton, MA, USA. Using its "Fit Linear" functionality, which uses linear regression, with least squares estimation).

[Table S95]

| For 1st equation | | | | | |
|---|---|---|---|---|---|
| *y* | *x* | N | Adj. $R^2$ | $\log_{10}(a)$ | *b* |
| A | F | 46 | 0.94153 | -1.07745 ±0.04705 | 0.62785 ±0.02331 |
| A | H | 53 | 0.79133 | -2.50796 ±0.16706 | -1.61154 ±0.11447 |
| A | I | 8 | 0.29511 | -1.39199 ±0.47849 | 1.88864 ±0.95262 |

| | | | | | |
|---|---|---|---|---|---|
| A | J | 8 | 0.50353 | 0.03795 ±0.34802 | -3.43944 ±1.20852 |
| A | K | 8 | 0.584 | -5.53965 ±0.78794 | -1.87511 ±0.30621 |
| A | L | 8 | 0.95049 | 4.59099 ±0.45735 | -2.3835 ±0.20486 |
| B | F | 10 | 0.96983 | 0.19367 ±0.10803 | 1.15357 ±0.0677 |
| B | H | 8 | 0.96001 | -2.86897 ±0.32786 | -3.45842 ±0.266 |
| B | I | 10 | 0.62228 | -0.47616 ±0.50488 | 4.20254 ±1.05635 |
| B | J | 10 | 0.68846 | 2.47843 ±0.45111 | -6.43496 ±1.40796 |
| B | K | 10 | 0.93797 | -9.37472 ±0.89638 | -4.01107 ±0.34258 |
| B | L | 10 | 0.97562 | 10.5046 ±0.50806 | -4.31725 ±0.22715 |
| C | F | 10 | 0.98802 | 0.27347 ±0.05074 | 0.8671 ±0.0318 |
| C | H | 8 | 0.99164 | -2.18013 ±0.11484 | -2.68589 ±0.09317 |
| C | I | 10 | 0.40975 | -0.05225 ±0.4705 | 2.65028 ±0.98443 |
| C | J | 10 | 0.5121 | 1.85428 ±0.42086 | -4.24547 ±1.31354 |
| C | K | 10 | 0.92448 | -6.80559 ±0.73732 | -2.9712 ±0.28179 |
| C | L | 10 | 0.96484 | 7.93033 ±0.45487 | -3.20269 ±0.20337 |
| D | F | 353 | 0.94931 | 0.41325 ±0.01058 | 0.69825 ±0.0086 |
| D | H | 27 | 0.75913 | -1.09488 | -1.77752 |

| | | | | | |
|---|---|---|---|---|---|
| | | | | ±0.22581 | ±0.19518 |
| D | I | 10 | 0.48942 | 0.25448 ±0.35647 | 2.31413 ±0.74584 |
| D | J | 10 | 0.62489 | 1.92958 ±0.3006 | -3.75199 ±0.93821 |
| D | K | 353 | 0.57669 | -4.24311 ±0.20592 | -1.81189 ±0.08265 |
| D | L | 10 | 0.97009 | 6.8251 ±0.34179 | -2.61516 ±0.15281 |
| E | F | 10 | 0.99043 | 2.34363 ±0.03792 | 0.72588 ±0.02377 |
| E | H | 8 | 0.99289 | 0.3634 ±0.08705 | -2.20857 ±0.07062 |
| E | I | 10 | 0.451 | 2.04247 ±0.37945 | 2.30015 ±0.79393 |
| E | J | 10 | 0.61987 | 3.73273 ±0.31064 | -3.83877 ±0.96956 |
| E | K | 10 | 0.92489 | -3.57683 ±0.61493 | -2.4851 ±0.23501 |
| E | L | 10 | 0.95871 | 8.7305 ±0.41221 | -2.67066 ±0.1843 |

[Table S96]

| For 2nd equation (maximal lifespan equation) | | | | |
|---|---|---|---|---|
| $y$ | N | Adj. $R^2$ | $\log_{10}(c)$ | $d$ |
| A | 43 | 0.723 | 1.39718 ±0.03125 | 0.29941 ±0.02847 |
| B | 10 | 0.69256 | 0.92935 ±0.09706 | 0.22932 ±0.04972 |
| C | 10 | 0.62051 | 0.90025 ±0.11321 | 0.29374 ±0.07409 |
| D | 353 | 0.39266 | 1.01454 ±0.01374 | 0.2338 ±0.01546 |

| | | | | |
|---|---|---|---|---|
| E | 10 | 0.6256 | 0.15376 ±0.26694 | 0.35246 ±0.08801 |

[Table S97]

| For 3rd equation | | | | | |
|---|---|---|---|---|---|
| *y* | *q* | N | Adj. $R^2$ | *e* | *g* |
| A | B | 8 | 0.99017 | 0.17224 ±0.08001 | 0.00336 ±1.26407E-4 |
| A | C | 8 | 0.98292 | 0.22567 ±0.10431 | 0.00883 ±4.39417E-4 |
| A | D | 27 | 0.91912 | 0.00891 ±0.07543 | 0.01717 ±9.97206E-4 |
| A | E | 8 | 0.99237 | 0.0088 ±0.07317 | 2.35631E-4 ±7.80389E-6 |
| B | C | 10 | 0.99285 | 9.54847 ±16.03507 | 2.63156 ±0.07439 |
| B | D | 10 | 0.92681 | 18.6681 ±9.24783 | 0.17302 ±0.01614 |
| B | E | 10 | 0.98159 | 776.48661 ±372.72967 | 14.26131 ±0.65038 |
| C | D | 10 | 0.91529 | -34.21255 ±22.90699 | 2.03564 ±0.20538 |
| C | E | 10 | 0.96637 | -21.1126 ±13.86874 | 0.02595 ±0.00161 |
| D | E | 10 | 0.97022 | 8.57818 ±6.16509 | 0.01228 ±7.15795E-4 |

Numerical prediction observed

Kleiber's law is that an adult animal's basal metabolic rate scales to $M^{0.75}$, where M is the animal's adult body mass [288-289]. Thence an adult animal's specific (per unit mass) basal metabolic rate scales to $M^{0.75}/M = M^{-0.25}$. Therefore, the $\log_{10}$-$\log_{10}$ plot of specific basal metabolic rate (*y*-axis), and adult body mass (*x*-axis), has a gradient of -0.25. Indeed, this is observed in the 2nd panel of Figure 3 in the main text (refer to the corresponding table of its best fit line equations, Table 1). Moreover, its 3rd and 4th panels also conform to "quarter-power scaling". In which its heart rate and maximal lifespan data both scale to body mass with the same exponents as (is typical) in the literature. $M^{-0.25}$ and $M^{0.25}$ respectively [363-364]. However, in Figure 3, it is notable that specific $F_1F_0$ ATP hydrolysis rate, which I suggest is a drive to specific basal metabolic rate (which scales to $M^{-0.25}$), does *not* have the -0.25 exponent itself. Its exponent is instead -0.13213 (±0.03077). This suggests that something might be missing. Specific $F_1F_0$ ATP hydrolysis rate data in prior figures (e.g., in Figure 3 in the main text) is from Sub-Mitochondrial Particles [SMPs] (data from [91]). In which all the intricate internal structure of mitochondria is lost. I suggest that this is what is missing in the exponent.

In intact mitochondria, across different species, paper [361] reports that [mitochondrial inner membrane surface area/mitochondrial matrix volume] ($cm^2$/µl) scales to $M^{-0.102}$.

Selecting data from [361], for species that I have specific $F_1F_0$ ATP hydrolysis rate data for (from SMPs, from [91]), the product, *g*, of [specific $F_1F_0$ ATP hydrolysis rate (from SMPs)], multiplied by [mitochondrial inner membrane surface area/mitochondrial matrix volume], scales to **precisely** $M^{-0.25}$. Shown in Figure S16.

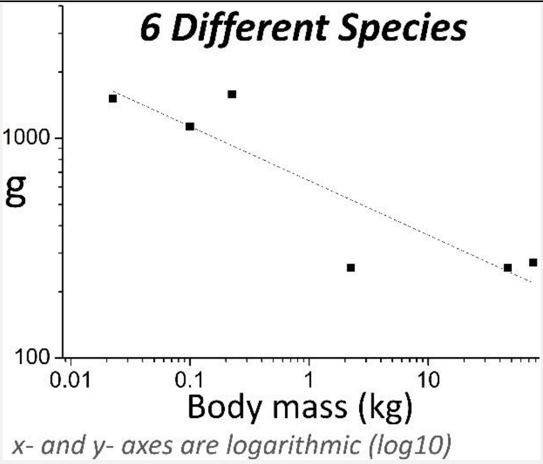

Figure S16 - On the *y*-axis, *g* is

$$g = h \cdot (s/v)$$

wherein *h* is specific $F_1F_0$ ATP hydrolysis rate in Sub-Mitochondrial Particles [SMPs] (µmol/min/mg; mg is of mitochondrial protein), *s* is mitochondrial inner membrane surface area ($cm^2$), *v* is mitochondrial matrix volume (µl). Data for *h* is from [91] (and has been used in prior figures: e.g., Figure 3 in the main text). Data for *s* and *v* is from [361]. In this figure, each species' body mass value is the mean of pooled body mass values, for the species, from [91] and [361].

Across species, *g* decreases with body mass (M). With an **exponent of -0.25**.

The best fit line in Figure S16 is:

$\log_{10}(y) = \log_{10}(a) + b \cdot \log_{10}(x)$, where $b$ is gradient: [Eq. S14]

[Table S98]

| $\log_{10}(a)$ | $b$ | Adjusted $R^2$ |
|---|---|---|
| 2.80659 (±0.08226) | -0.24725 (±0.06192) ≈ ***-0.25*** | 0.74928 |

Pearson correlation coefficient (R), and its p-value (one-tailed, because the alternative hypothesis is directional), for the $\log_{10}$-$\log_{10}$ data of Figure S16:

[Table S99]

| | | $\log_{10}$(g) |
|---|---|---|
| R | $\log_{10}$(body mass) | -0.8941 |
| $p$ | | 0.0081 |

I suggest that *g*, mathematically defined in the Figure S16 legend, can be (abstractly) interpreted as being specific $F_1F_0$ ATP hydrolysis rate in mitochondria (μmol/min/mg·cm$^2$/μl). So, whereas *h* is specific $F_1F_0$ ATP hydrolysis rate in Sub-Mitochondrial Particles [SMPs] *in vitro, g* is (interpreted to correspond to) specific $F_1F_0$ ATP hydrolysis rate in intact mitochondria *in vivo*. Wherein *g* and *h* are interrelated by the equation in the legend of Figure S16.

More mitochondrial inner membrane surface area, per mitochondrial matrix volume, is to say the mitochondrial inner membrane has more and/or larger invaginations (cristae). Sacrificing mitochondrial matrix volume for greater mitochondrial inner membrane surface area. This greater surface area of the mitochondrial inner membrane means there is more proton leak across it.

As aforementioned, Figure S16 shows that specific $F_1F_0$ ATP hydrolysis rate combines with mitochondrial inner membrane surface area (per mitochondrial matrix volume) to confer, *precisely*, the expected -0.25 exponent. That this relation is because of proton leak is suggested by specific $F_1F_0$ ATP hydrolysis rate combining with the rate of proton leak to confer, within the margin of slope error, a -0.25 exponent also. As shown in Figure S17.

In overlapping support, [361] uses data from more species than are represented in Figure S17 and finds body mass and proton leak related with an exponent of -0.129 (±0.049). Which, when added to the (aforementioned) exponent relating body mass and specific $F_1F_0$ ATP hydrolysis rate (-0.13213±0.03077), yields an exponent of -0.26, with -0.25 within its margin of error.

Pearson correlation coefficient (R), and its p-value (one-tailed, because the alternative hypothesis is directional), between *l*, the rate of proton leak ([nmol of $H^+$]/min/mg protein) across the inner mitochondrial membrane {when the membrane potential across it is -170 mV, with ATP synthase blocked by oligomycin}, and (*s*/*v*); where *s* is mitochondrial inner membrane surface area ($cm^2$) and *v* is mitochondrial matrix volume (μl); data from [361]: [Table S100]

| | | $(s/v)$ |
|---|---|---|
| R | $l$ | 0.861 |
| $p$ | | 0.014 |

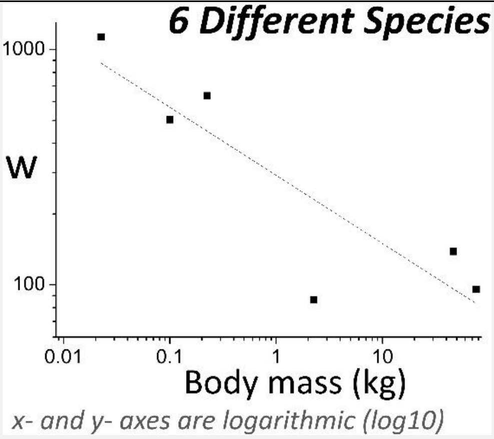

Figure S17 - On the *y*-axis, *w* is

$$w = h \cdot l$$

wherein *h* is specific $F_1F_0$ ATP hydrolysis rate in Sub-Mitochondrial Particles [SMPs] (μmol/min/mg; mg is of mitochondrial protein), *l* is proton leak rate across the inner

mitochondrial membrane, when the membrane potential across it is -170 mV, and ATP synthase is blocked by oligomycin ([nmol of $H^+$]/min/mg protein). Data for $h$ is from [91] (and has been used in prior figures: e.g., Figure 3 in the main text). Data for $l$ is from [361]. In this figure, each species' body mass value is the mean of pooled body mass values, for the species, from [91] and [361].
Across species, $w$ decreases with increasing body mass (M). With (within the margin of slope error) an exponent of -0.25.

The best fit line in Figure S17 is:

$\log_{10}(y) = \log_{10}(a) + b \cdot \log_{10}(x)$, where $b$ is gradient: [Eq. S15]

[Table S101]

| $\log_{10}(a)$ | $b$ | Adjusted $R^2$ |
|---|---|---|
| 2.46545 (±0.10233) | -0.28943 (±0.07704) ≈ *-0.25* | 0.72399 |

Pearson correlation coefficient (R), and its p-value (one-tailed, because the alternative hypothesis is directional), for the $\log_{10}$-$\log_{10}$ data of Figure S17:

[Table S102]

| | | $\log_{10}$(w) |
|---|---|---|
| R | $\log_{10}$(body mass) | -0.8827 |
| $p$ | | 0.0099 |

Pearson correlation coefficients (R), and their p-values (one-tailed, because each alternative hypothesis is directional), between $h$, $g$, and $w$ (variables as defined in the legends of Figures S16 and S17):

[Table S103]

| | | $g$ | $w$ |
|---|---|---|---|
| R | $h$ | 0.8323 | 0.6779 |
| $p$ | | 0.020 | 0.069 |
| R | | $g$ | 0.9036 |
| $p$ | | | 0.0067 |

Pearson correlation (R) coefficients, and their p-values (one-tailed, because each alternative hypothesis is directional):

[Table S104]

| | | Specific basal metabolic rate | Heart rate | Maximal lifespan |
|---|---|---|---|---|
| R | ***h*** | **0.7016** | **0.8007** | **-0.7192** |
| $p$ | | 0.0054965 | 0.000877 | 0.006307 |
| R | ***g*** | **0.8816** | **0.9466** | **-0.7942** |
| $p$ | | 0.010099 | 0.002101 | 0.029586 |
| R | ***w*** | **0.9691** | **0.9442** | **-0.6952** |
| $p$ | | 0.0007085 | 0.002292 | 0.062598 |

The correlations with $h$, where $h$ is specific $F_1F_0$ ATP hydrolysis rate (from [91]), are as presented in Table 3 in the main text. Its correlations are calculated with more species, and so a higher value of $n$, than the correlations to $g$ and $w$. And thence its correlations have a smaller p-value for any given value of R.
Looking at R values only, the correlations of $g$ are superior to those of $h$. Those of $w$ are superior to those of $g$. Except for the correlation to maximal lifespan.
When $h$ is correlated to the same, smaller data set as $g$ and $w$, its correlation to maximal lifespan doesn't worsen but improves (-0.7842), and its correlation to specific basal metabolic rate and heart rate is 0.7318 and 0.7706 respectively.

Comparing Table S105 below, to Table S104 above, one can see that the correlations of each of $g$ and $w$, are *much* superior to those of $(s/v)$ and $l$ (variables as defined in legends of Figures S16 and S17). To specific basal metabolic rate, heart rate, and maximal lifespan (data for these variables is presented in Table S2). R is the Pearson correlation coefficient and p-values are one-tailed, because each alternative hypothesis is directional. Incidentally, proton leak, $l$, negatively correlates with maximal lifespan (at least across these species). Although not significantly so.
[Table S105]

| | | $h$ | Specific basal metabolic rate | Heart rate | Maximal lifespan |
|---|---|---|---|---|---|
| $(s/v)$ | R | 0.4445 | 0.7588 | 0.8193 | -0.5301 |
| | p | 0.188581 | 0.040125 | 0.023015 | 0.1396655 |
| $l$ | R | 0.4159 | 0.8794 | 0.8283 | -0.496 |
| | p | 0.20606 | 0.01047 | 0.020845 | 0.158506 |

Considered in totality, the data of this present section (encompassing Figures S16 and S17) suggests that it might be the combination of $F_1F_0$ ATP hydrolysis *and* proton leak, which largely sets specific basal metabolic rate (setting heart rate), which dictates maximal lifespan.

Why might $F_1F_0$ ATP hydrolysis, and proton leak, *both* be important?

Oxidative phosphorylation synthesizes ATP. $F_1F_0$ ATP hydrolysis hydrolyses a fraction of this ATP. At a level constrained by IF1 protein activity. Wherein smaller, shorter living species have less specific IF1 protein activity. This $F_1F_0$ ATP hydrolysis pumps protons out of the mitochondrial matrix, across the inner mitochondrial membrane, into the mitochondrial intermembrane space. A fraction of these protons return to the mitochondrial matrix via proton leak. Wherein smaller, shorter living species tend to have greater proton leak because of a greater mitochondrial inner membrane surface area (per volume of mitochondrial matrix). And because of their greater (per unit mass) rate of $F_1F_0$ ATP hydrolysis, pumping more protons per unit time into the mitochondrial intermembrane space. A fraction of the ATP synthase proton conduits in the mitochondrial inner membrane, actively pumping protons energetically "uphill", instead of passing protons energetically "downhill", can explain how so much proton leak can occur. Why so many protons cross the mitochondrial inner membrane at a point of higher resistance. Dissipating their potential energy as heat. Rather than just passing through the lower resistance conduit, ATP synthase, which can capture (a fraction of) their potential energy loss in the chemical energy of ATP. *(But these ATP synthase complexes are available for "forward", proton passing, ATP synthesizing [instead of "reverse", ATP consuming, proton pumping] action if cellular ATP demand increases).*

To theorize, this proton leaking may be especially pronounced when there is particularly high [ATP] in the mitochondrial matrix, which may happen when cellular energy demand is low. Thereby, when the cell is hydrolysing much ATP to perform much useful work, inherently generating heat as a by-product (2$^{nd}$ Law of Thermodynamics), [ATP] in the mitochondrial matrix is lower and $F_1F_0$ ATP hydrolysis, and proton leak, is lower. But when the cell isn't performing much work, [ATP] in the mitochondrial matrix is higher and so $F_1F_0$ ATP hydrolysis, and proton leak, is higher. Which generates heat to keep the body at 37°C. So, when useful work can generate heat, the spurious processes are less. But when not, they are more. For emphasis, this is recast in the next paragraph.

When the body is resting and so isn't performing much work, $F_1F_0$ ATP hydrolysis (and its drive to $F_1F_0$ ATP synthesis and metabolic rate) confers endogenous heat production. But when the body is active and performing appreciable work, ATP is consumed to do this work (inherently generating heat as a by-product, 2nd Law of Thermodynamics), and so there is less ATP available to $F_1F_0$ ATP hydrolysis, which is reduced as a result. So, when performing work, which inherently generates heat, the futile (no work) process to generate heat is decreased, affording more energy to perform work, and reducing the chance that the body will overheat. So, in the resting state, heat generation is (fractionally) by a futile process, until more work is required, after which the heat generation by the actual performing of work substitutes for a reduction in heat generation by the futile process.

Quarter-power scaling

It is interesting to contemplate that the observed quarter-power scaling, between novel variable, *w*, and body mass, might be the fundamental drive to the quarter-power scaling observed (e.g., in Figure 3 in the main text) between body mass and each of specific basal metabolic rate (-0.25), heart rate (-0.25), and maximal lifespan (+0.25). Indeed, the basis might be at the cellular or sub-cellular level because the specific metabolic rate of liver slices, sourced from different sized species, adheres to quarter-power scaling (-0.25) *in vitro* [362]. Curiously, *w* has a geometric component, via its proton leak rate, which is a function of inner mitochondrial membrane surface area, which is a *non-Euclidean* shape. Because of its invaginations (sacrifices some volume for greater surface area). A longstanding mystery is why, across species, where M is adult body mass, basal metabolic rate scales with $M^{0.75}$ (Kleiber's law), instead of $M^{0.67}$, which is what is expected given how surface area scales with volume for a *Euclidean* object [363]. "One cannot but wonder why the power formula, in general, and the mass exponents of 3/4, 1/4 and -1/4, in particular, are so effective in describing biological phenomenon" [363].

Mitochondrial membrane potential, proton leak, and maximal lifespan

Table S106 below shows correlations (R is the Pearson correlation coefficient; p-values are one-tailed, because each alternative hypothesis is directional) to the *absolute* value of the resting mitochondrial membrane potential, $|\Psi_{IM}|$. $\Psi_{IM}$ data (mV) is from [365], from isolated hepatocytes.

[Table S106]

| | | | | | | | Specific basal | | Maximum |
|---|---|---|---|---|---|---|---|---|---|
| | | *(s/v)* | *l* | *h* | *g* | *w* | metabolic rate | Heart rate | lifespan |
| \|ΨIM\| | R | -0.6482 | -0.7588 | -0.7818 | -0.9759 | -0.9615 | -0.8956 | -0.899 | 0.8712 |
| | p | 0.081938 | 0.0401 | 0.05913 | 0.01205 | 0.01925 | 0.0199265 | 0.018971 | 0.027202 |

At least across these species, the less hyperpolarised the mitochondrial membrane potential (i.e., the smaller the value of $|\Psi_{IM}|$), the shorter the maximal lifespan.

At least across these species, the greater the proton leak, *l*, the smaller the value of $|\Psi_{IM}|$, i.e., the less hyperpolarised the mitochondrial membrane potential. Proton leak is attenuating to the proton motive force (pmf), and its $\Psi_{IM}$ component. As is futile cycling of $F_1F_0$ ATP synthesis (pmf dissipation) and $F_1F_0$ ATP hydrolysis (pmf creation): because no energy conversion can be 100% efficient (2$^{nd}$ Law of Thermodynamics).

Proton leak is not only an (attenuating) input into $\Psi_{IM}$, but also a function of it. Wherein greater $\Psi_{IM}$ is an Ohmic, or supra-Ohmic, drive to greater proton leak. The proton leak variable used herein, *l*, is the proton leak when $\Psi_{IM}$ is fixed at a value of -170 mV (and when ATP synthase is blocked by oligomycin). But, as referred to earlier, the different species don't have the same (resting) value of $\Psi_{IM}$. And so, then there is a question mark over the fidelity of *l* to the actual proton leaks of these species. So, use of *l* herein is with this caveat.

Table S107 below shows correlations to *l* multiplied by *absolute* resting $\Psi_{IM}$, which might better capture the proton leak across these species. At least across these species, maximal lifespan is inversely proportional to this variable. But this doesn't reach statistical significance (at $p < 0.05$), because although R is sizeable, *n* is small (=4).

[Table S107]

| | | | | | | | Specific basal | | Maximum |
|---|---|---|---|---|---|---|---|---|---|
| | | *(s/v)* | *l* | *h* | *g* | *w* | metabolic rate | Heart rate | lifespan |
| *l**\|ΨIM\| | R | 0.8497 | 0.9898 | 0.6511 | 0.7613 | 0.9617 | 0.9616 | 0.8902 | -0.7568 |
| | p | 0.016094 | 8E-05 | 0.17445 | 0.11935 | 0.01915 | 0.0192 | 0.0549 | 0.1216 |

<u>USING LOGGED DATA: Instrumental variable analysis reports causality between lower specific $F_1F_0$ ATP hydrolysis rate and higher maximal lifespan</u>

The following differs from that presented in the main text in that it uses logged ($\log_{10}$) variables, but its results are consistent. Analysis with $\log_{10}$ maximal lifespan as the dependent variable, $log_{10}$ specific $F_1F_0$ ATP hydrolysis rate as the independent variable, and $log_{10}$ specific IF1 protein activity as the instrumental variable, reports causality between lower specific $F_1F_0$ ATP hydrolysis rate and higher maximal lifespan.

[Table S108]

| *β* | E | P | A | C | A2 | C2 |
|---|---|---|---|---|---|---|
| -2.2010 | 0.9717 | 0.0249 | 0.00069013 | (-Infinity, -1.05040841895407) union (15.8214097099972, Infinity) | 0.00069013 | (-Infinity, -1.05040745763014) union (15.8211738664685, Infinity) |

<u>USING LOGGED DATA: Instrumental variable analysis reports causality between lower specific basal metabolic rate and higher maximal lifespan</u>

The following differs from that presented in the main text in that it uses logged ($log_{10}$) variables, but its results are consistent. Analysis with $log_{10}$ maximal lifespan as the dependent variable, $log_{10}$ specific basal metabolic rate as the independent variable, and $log_{10}$ specific $F_1F_0$ ATP hydrolysis rate as the instrumental variable, reports causality between lower specific basal metabolic rate and higher maximal lifespan.

[Table S109]

| *β* | E | P | A | C | A2 | C2 |
|---|---|---|---|---|---|---|
| -0.7549 | 0.2066 | 0.00264 | 0.024597 | -1.34710285048184, -0.191076194529155 | 0.024597 | -1.34710165010201, -0.191077282465823 |

<u>USING LOGGED DATA: Instrumental variable analysis reports causality between lower *ΔD* and higher maximal lifespan</u>

The following differs from that presented earlier in that it uses logged ($log_{10}$) variables, but its results are consistent. Analysis with $log_{10}$ maximal lifespan as the dependent variable, $log_{10}$ *ΔD* as the independent variable, and $log_{10}$ specific basal metabolic rate as the instrumental variable, reports causality between lower *ΔD* and higher maximal lifespan.

[Table S110]

| *β* | E | P | A | C | A2 | C2 |
|---|---|---|---|---|---|---|
| -1.0950 | 0.1693 | 0.0000074 | 0.00037588 | -1.60545542806554, -0.76074833068146 | 0.00037588 | -1.60545303631843, -0.76074935601888 |

In Table S111 below, mediation analysis reports that $log_{10}$ specific basal metabolic rate only affects $log_{10}$ maximal lifespan through $log_{10}$ *ΔD*, with *no* direct effect upon $log_{10}$ maximal lifespan. Conducted in JASP software (version 0.18.1.0), selecting its bootstrap method called "Percentile", run with 10,000 replications, utilizing its standardized estimates option, which makes each variable mean = 0 and standard deviation = 1.

[Table S111]

| | | | 95% Confidence Interval (CI) | | | 95% Confidence Interval (CI) | |
|---|---|---|---|---|---|---|---|
| α | β | **Indirect effect** | Lower | Upper | **Direct effect (τ)** | Lower | Upper |
| 2.1 | -0.74 | -1.54 | -2.746 | -0.501 | -0.58 | -1.399 | 0.48 |

<u>Across ectotherm species, maximal lifespan is inversely proportional to specific basal metabolic rate</u>

For *all* the <u>**ECTOTHERM**</u> species that presently have adult body mass, specific basal metabolic rate, and maximal lifespan data available in [1], which is 29 species, 16 of Amphibia and 13 of Reptilia: (*a*) maximal lifespan is proportional to adult body mass, which might be because across these same ectotherm species: (*b*) specific basal metabolic rate is inversely proportional to adult body mass, and thence (*c*) maximal lifespan is inversely proportional to specific basal metabolic rate. The correlations in turn for (*a*), (*b*), and (*c*) are listed below:

***(a)*** Pearson correlation coefficient = 0.6373, one-tailed (alternative hypothesis is directional) p-value = 0.0001005.

Spearman's rank correlation coefficient = 0.5987, one-tailed p-value = 0.0003005.

*Using unlogged data with GraphPad Prism 10.0.3 software with its default settings (using least square regression. With chosen starting values: a = 1, b = 0.25): supra-linear equation* $y = a \cdot x^b$ *was curve-fitted, which best-fitted with a = 31.90, b = 0.1352, wherein adjusted* $R^2$ *= 0.4521. Adding more terms could better the fit, at "cost" of more parameters: power series equation:* $y = a \cdot x^b + c \cdot x^d$*: adjusted* $R^2$ *= 0.4722. A linear equation (which does not fit as well as the [aforementioned] supra-linear equation with same number of parameters):* $y = m \cdot x + c$*, where m = 0.3672, c = 22.13, and adjusted* $R^2$ *= 0.3842. When x and y are both logged, linear equation:* $log_{10}(y) = m \cdot log_{10}(x) + log_{10}(c)$*, where m = 0.1430,* $log_{10}(c)$ *= 1.456, and adjusted* $R^2$ *= 0.3144.*

**(b)** Pearson correlation coefficient = -0.2417, one-tailed (alternative hypothesis is directional) p-value = 0.103264.

Spearman's rank correlation coefficient = -0.5562, one-tailed p-value = 0.000865.

*Using unlogged data, supra-linear equation* $y = a \cdot x^b$ *was curve-fitted, which best-fitted with a = 0.0002609, b = -0.1012, wherein adjusted* $R^2$ *= 0.1673. A linear equation does not fit as well:* $y = m \cdot x + c$*, where m = -1.902e-006, c = 0.0003568, and adjusted* $R^2$ *= 0.02356. When x and y are both logged, linear equation:* $log_{10}(y) = m \cdot log_{10}(x) + log_{10}(c)$*, where m = -0.1579,* $log_{10}(c)$ *= -3.747, and adjusted* $R^2$ *= 0.2629.*

**(c)** Pearson correlation coefficient = -0.5203, one-tailed (alternative hypothesis is directional) p-value = 0.001906.

Spearman's rank correlation coefficient = -0.5684, one-tailed p-value = 0.000648.

*Using unlogged data, supra-linear equation* $y = a \cdot x^b$ *was curve-fitted, which best-fitted with a = 1.110, b = -0.3715, wherein adjusted* $R^2$ *= 0.2437. A linear equation does not fit as well:* $y = m \cdot x + c$*, where m = -38094, c = 37.76, and adjusted* $R^2$ *= 0.2437. When x and y are both logged, linear equation:* $log_{10}(y) = m \cdot log_{10}(x) + log_{10}(c)$*, where m = -0.4821,* $log_{10}(c)$ *= -0.4218, and adjusted* $R^2$ *= 0.3071.*

Table S112 below shows multiplicator (*a*) and exponent (*b*) values from the fitted-to-data-equation: maximal lifespan $= \mathrm{a} \cdot \mathrm{x}^{\mathrm{b}}$. For species sets *A* (517 homeotherm/endotherm species: mammal and bird species), *B* (350 mammal species), *C* (263 mammal species, with bats, naked mole rat, and marsupials excluded), and *E* (29 ectotherm species). These values are as disclosed earlier herein.

[Table S112]

| Set | *x* = adult body mass | | *x* = specific basal metabolic rate | |
|---|---|---|---|---|
| | a | b | a | b |
| *A* | 17.78 | 0.1376 | 1.907 | -0.3854 |
| *B* | 15.02 | 0.1659 | 0.6999 | -0.5248 |
| *C* | 14.87 | 0.1733 | 0.6802 | -0.5344 |
| *E* | 31.90 | 0.1352 | 1.110 | -0.3715 |

Table S113 below shows multiplicator (*a*) and exponent (*b*) values from the fitted-to-data-equation: specific basal metabolic rate $= \mathrm{a} \cdot (\text{adult body mass})^{\mathrm{b}}$. For species sets *A*, *B*, *C*, and *E*. Values as disclosed earlier herein.

[Table S113]

| Set | a | b |
|---|---|---|
| A | 0.003119 | -0.3567 |
| *B* | 0.002611 | -0.3404 |
| *C* | 0.002453 | -0.3958 |
| *E* | 0.0002609 | -0.1012 |

In Table S112 above, comparing values for the 517 endotherm species of species set *A* with the 29 ectotherm species of species set *E*: maximal lifespan tends to be higher for a given adult body mass in ectotherms than endotherms, which might be related to (as shown in Table S113 above) specific basal metabolic rate tending to be lower for a given adult body mass in ectotherms than endotherms (presumably because they aren't conducting deliberate metabolic heat generation). Table S112 shows that maximal lifespan tends to be slightly higher for a given specific basal metabolic rate in endotherms than ectotherms, but that in ectotherms too, maximal lifespan is inversely proportional to (as a negative convex function of) specific basal metabolic rate. So, to interpret, how larger ectotherm species live longer is the same as how larger endotherm species live longer, by a lower specific basal metabolic rate.

Consistent with this correlative result: lower ambient temperature, which (*because chemical reaction rates are temperature dependent [366-367]*) reduces their metabolic rate, makes ectotherms live longer. At least in worms [23], flies [24-26], wasps [27], and fish [28].

Larger adult body mass, lower specific basal metabolic rate, and higher maximal lifespan across ectotherm species

Smaller ectotherm species may require greater non-basal metabolic rate per unit mass, (at least) because moving a smaller body is more energetically expensive per unit mass [368-372]. Generally, this greater non-basal metabolic rate per unit mass requires greater mitochondrial membrane surface-area per unit mass, which means (all other factors being constant) greater proton leak rate per unit mass (*ectotherms do have proton leak [373-374]; greater mitochondrial membrane surface-area means greater proton leak [361]*), and thence greater specific basal metabolic rate. Indeed, smaller species have greater mitochondrial membrane surface-area per unit mass. Observed (as aforementioned) with endotherms [361]. And with ectotherms [375].

In [375], which reports on ectotherms, refer to its Fig. 5, and the furthest-right column of its Table 1. To consider, for example, the first entry of that column:
$mitochondrial\ membrane\ surface\ area\ (m^2) = 0.98 \cdot W^{0.64}$, where $W$ is weight (g). Deductively, the corresponding mitochondrial surface area per unit mass ($m^2/g$) equation includes term $W^{0.64}/W$, which corresponds to $W^{-0.36}$. So, mitochondrial surface area per unit mass is less, the larger the value of $W$.

So, as ectotherms *do* have proton leak [373-374], why aren't they endothermic? I suggest (at least) because they do *not* have appreciable $F_1F_0$ ATP hydrolysis, which generates heat by futile cycling of $F_1F_0$ ATP synthesis and $F_1F_0$ ATP hydrolysis and by (by it pumping protons back into the intermembrane space) greatly increasing proton leak.

<u>Linear regressions with 29 ectotherm species data</u>

Table S114 below, in its first data line, reports linear regression, with maximal lifespan (years) as the response variable, and specific basal metabolic rate (W/g) as the explanatory variable. Both variables logged ($log_{10}$). To generate the results of the second line, $log_{10}$ basal metabolic rate (W) was used in place of $log_{10}$ specific basal metabolic rate. All data sourced from [1]. Column headings: $S\beta_1$ is the standardized regression coefficient, $p\_\beta_1$ is p-value for the unstandardized regression coefficient, *Adj.* $R^2$ is adjusted $R^2$, *p_F* is the p-value for the F-statistic, and *DW* is the Durbin-Watson statistic. Conducted in JASP software (version 0.18.1.0), using its method called "Enter". Assumptions of homoscedasticity, normality, and linearity weren't violated.

[Table S114]

| Adj. $R^2$ | DW | p_F | $S\_\beta_1$ | $p\_\beta_1$ |
|---|---|---|---|---|
| 0.307 | 1.945 | 0.001 | -0.576 | 0.001 |
| 0.116 | 2.057 | 0.040 | 0.384 | 0.040 |

In Table S114 above, specific basal metabolic rate is a significant predictor of maximal lifespan, and basal metabolic rate is a significant predictor of maximal lifespan, across these ectotherm species.

<u>Multiple linear regressions with 29 ectotherm species data</u>

Table S115 below, in its first data line, reports multiple linear regression, where maximal lifespan (years) is the response variable, and specific basal metabolic rate (W/g) and adult body mass (kg) are explanatory variables. All variables logged ($log_{10}$). To generate the results of the second line, $log_{10}$ basal metabolic rate (W) was used in place of $log_{10}$ specific basal metabolic rate. All data sourced from [1]. Column headings: $p\_\beta_1$ and $p\_\beta_2$ are p-values for unstandardized regression coefficients, and $S\beta_1$ and $S\beta_2$ are standardized regression coefficients, for these explanatory variables, wherein $\beta_1$ relates to the $log_{10}$ adult body mass predictor. *Adj.* $R^2$ is adjusted $R^2$, *p_F* is the p-value for the F-statistic, *VIF* is the Variance Inflation Factor for each of the slope regression coefficients, and *DW* is the Durbin-Watson statistic. Conducted in JASP software (version 0.18.1.0), using its method called "Enter", wherein the $log_{10}$ adult body mass predictor was forced in first. Assumptions of homoscedasticity, normality, and linearity weren't violated in either case.

[Table S115]

| Adj. $R^2$ | VIF | DW | p_F | $S\_\beta_1$ | $S\_\beta_2$ | $p\_\beta_1$ | $p\_\beta_2$ |
|---|---|---|---|---|---|---|---|
| 0.393 | 1.407 | 1.647 | 5.820e-4 | 0.383 | -0.370 | 0.037 | 0.044 |
| 0.387 | 4.952 | 1.857 | 6.527e-4 | 1.186 | -0.676 | 0.001 | 0.050 |

In Table S115 above, even with adult body mass in the model, specific basal metabolic rate is a significant predictor of maximal lifespan across these ectotherm species.

The results with basal metabolic rate appear corrupted by collinearity (*VIF* = 5), conferring the standardized regression coefficient for basal metabolic rate, written in red text, with erroneous sign. Erroneous because, as can be seen in Table S116 below (in its zero-order column), maximal lifespan positively correlates with basal metabolic rate.

In Table S116 below are corresponding zero-order, partial, and semi-partial (part) Pearson correlation coefficients. JASP outputted. One-tailed (alternative hypothesis is directional) p-values are italicized in brackets.

[Table S116]

| | Zero-order | Partial | Part |
|---|---|---|---|
| *MAXIMAL LIFESPAN vs.* | | | |
| Specific basal metabolic rate | -0.576 *(0.000538)* | -0.384 *(0.019865)* | -0.312 *(0.049708)* |

| | | | |
|---|---|---|---|
| Adult body mass | 0.5821 *(0.000462)* | 0.395 *(0.0169745)* | 0.323 *(0.043721)* |
| *MAXIMAL LIFESPAN vs.* | | | |
| Basal metabolic rate | 0.3835 *(0.0200055)* | -0.374 *(0.9750377)* | -0.304 *(0.942113)* |
| Adult body mass | 0.5821 *(0.000462)* | 0.577 *(0.000525)* | 0.533 *(0.0014555)* |

In Table S116 above, maximal lifespan is a significant correlate of specific basal metabolic rate, even when the effect of adult body mass is statistically excluded.

When statistically excluding the effect of specific basal metabolic rate, the correlation between maximal lifespan and adult body mass is appreciably less.

For basal metabolic rate, its partial and part correlation coefficients, written in red text, are of erroneous sign. Probably because of the earlier shown collinearity, which made the regression coefficient for basal metabolic rate of erroneous sign.

Generalized Additive Model (GAM) for ectotherms

Using the same R code as earlier, but with different data, I built a GAM model for the 29 ectotherm species with maximal lifespan in years ($y$) as the response variable, with adult body mass in kg ($z$) and specific basal metabolic rate in W/g ($x$) as predictors. No variables logged. Data sourced from [1]. Family was Gaussian, link was identity, and output is shown in Table S117 below, with column headings as defined for earlier Table S52.

[Table S117]

| A.R$^2$ | GCV | Dev | Int | Int.p | | edf | ref.df | P | K | CC |
|---|---|---|---|---|---|---|---|---|---|---|
| 0.55 | 147.81 | 62.4% | 24.669 | 1.34e-11 | s($x$) | 3.384 | 4.105 | 0.04245 | 9 | 0.8554514 |
| | | | | | s($z$) | 1.252 | 1.441 | 0.00142 | 9 | 0.8554514 |

In Table S117 above, even with adult body mass included in the model, specific basal metabolic rate is a significant predictor of maximal lifespan.

*Diagnostic graphs for this GAM: Model-predicted maximal lifespan vs. observed maximal lifespan plot is linear, but with appreciable scatter of points around this linearity.*

*Residuals vs. predicted (values of response variable) plot shouldn't contain pattern(s) and/or trend(s) and this is the case, but with one outlier. The Q-Q plot should be linear, and it is except for non-linearity at its ends. Working responses vs. linear predictor plot should be linear and it is, with no outliers present. For a GAM, partial residuals vs. predictor plots should appear random and evenly distributed around zero, which is the case for x, but z has some vertical linearity to its plot. The Component-Plus-Residual (CPR) plot is, as it should be, data points scattered around linear, all be it there is one outlier.*

*High concurvity indicates redundancy/entwinement of predictors and can inflate smooth term p-values. The GAM model has high concurvity in its presented "worst" values, but less in its "observed" and "estimate" values, especially less in its "estimate" values, as shown in Table S118 below.*

*[Table S118]*

| | *Worst* | *Observed* | *Estimate* |
|---|---|---|---|
| *s(x)* | *0.8554514* | *0.6468042* | *0.3900225* |
| *s(z)* | *0.8554514* | *0.2593360* | *0.2737324* |

Table S119 below shows results when the GAM model is simplified, such that specific basal metabolic rate is the only predictor:

[Table S119]

| $A.R^2$ | GCV | Dev | Int | Int.p | | edf | ref.df | P | K |
|---|---|---|---|---|---|---|---|---|---|
| 0.244 | 214.84 | 27.1% | 24.669 | 5.35e-10 | s(*x*) | 1 | 1 | 0.00381 | 9 |

In Table S119 above, specific basal metabolic rate is a significant predictor of maximal lifespan.

<u>Across ectotherm species, higher maximal lifespan significantly correlates with lower specific basal metabolic rate, even with phylogeny and adult body mass statistically excluded</u>

A phylogeny for the 29 ectotherm species was sought from [322], all be it *Trachemys scripta* was omitted because of an "unresolved name" issue, rendering 28 species remaining, in a phylogeny with no polytomies. Data for species adult body mass (kg), maximal lifespan (years), specific basal metabolic rate (W/g), and basal metabolic rate (W) was sourced from [1]. The Phylogenetic Independent Contrasts (PIC) method requires compared traits to be linearly related [324], and so the following PIC analysis was conducted upon logged variables.

The diagnostic table below, Table S120, shows for $Q$ ($\log_{10}$ specific basal metabolic rate), $R$ ($\log_{10}$ maximal lifespan), $S$ ($\log_{10}$ basal metabolic rate), and $T$ ($\log_{10}$ adult body mass): in column $C$, the Pearson correlation coefficient (*not* forced through origin [323, 325]) of the absolute values of the standardized phylogenetically independent contrasts versus their standard deviations (i.e., the square root of the sum of their branch lengths), which is a diagnostic check of how well the topology and branch lengths of the phylogenetic tree (and assumed model of character evolution) adequately fit the tip data [323], wherein no significant correlation is the desirable result (*and in the line below, labelled "P", the corresponding two-tailed p-value thereof*), and subsequent columns are that but with a branch length adjustment made, as codified earlier. P-values greater than 0.05, which are desirable, are in red text. Table values were outputted by the PDAP module [320] of Mesquite [321]. [Table S120]

| Diagnostic table | | | | | |
|---|---|---|---|---|---|
| | *C* | Nee | Pagel | Grafen | =1.0 |
| **Q** | -0.16418410026374758 | 0.002078855211505119 | -0.14208073597361584 | -0.3264836695472011 | -0.014120377611371946 |
| *P* | 0.4131676057146124 | 0.9917892028128117 | 0.479604966504968 | 0.09649529175103316 | 0.9442708095815049 |
| **R** | -0.4964624151856449 | -0.11154524803662727 | -0.22901684844338474 | -0.3346449657160886 | -0.1494405669696904 |
| *P* | 0.008437865949246896 | 0.5796381112902009 | 0.250535740940109 | 0.08797526690812338 | 0.45689750666756557 |

| | | | | | |
|---|---|---|---|---|---|
| **S** | -0.44461730725714016 | -0.2896099926895699 | -0.4270500417801416 | -0.5259680527052257 | -0.3878913253483443 |
| *P* | 0.020144258910349797 | 0.14284894906110512 | 0.026307001689366682 | 0.0048343849310328955 | 0.04558390917994332 |
| **T** | -0.13549805229599723 | -0.016409813461495837 | -0.12054051879538694 | -0.29626830569569185 | -0.053551992087170854 |
| *P* | 0.5003931943057311 | 0.9352523857541619 | 0.5492399667615651 | 0.1334764981621771 | 0.7907880587689944 |

Following the teaching of [323, 325], raw phylogenetically independent contrasts were outputted from the PDAP module of Mesquite, and each was then standardized by dividing each contrast by the (also PDAP outputted) contrast's standard deviation (square root of sum of corrected branch lengths). The raw contrasts, and corresponding standard deviations to standardize them, for all four variables ($log_{10}$ adult body mass, $log_{10}$ maximal lifespan, $log_{10}$ specific basal metabolic rate, $log_{10}$ basal metabolic rate) were sourced with the Nee branch length transformation. As this (according to the diagnostic table, Table S120, above) is a valid single branch length transformation for all four. Their having the same branch length transformation eases interpretability. These standardized phylogenetically independent contrasts are used in the multiple linear regressions of the next paragraph.

Table S121 below, on its first line, reports upon multiple linear regression, where standardized phylogenetically independent contrasts for $log_{10}$ maximal lifespan is the response variable and standardized phylogenetically independent contrasts for $log_{10}$ adult body mass and standardized phylogenetically independent contrasts for $log_{10}$ specific basal metabolic rate are explanatory variables. In the table, $p\_\beta_1$ and $p\_\beta_2$ are p-values for unstandardized regression coefficients, and $S\beta_1$ and $S\beta_2$ are standardized regression coefficients, for these explanatory variables respectively. *Adj.* $R^2$ is adjusted $R^2$, *p_F* is the p-value for the F-statistic, *VIF* is the Variance Inflation Factor for each of the slope regression coefficients, and *DW* is the Durbin-Watson statistic. Regression was forced through the origin, following the teaching of [323, 325]. Conducted in JASP software (version 0.18.1.0), using its method called "Enter", wherein the adult body mass related predictor was forced in first. Assumptions of homoscedasticity, normality, and linearity weren't violated. On the table's second line is as aforementioned, except that standardized phylogenetically independent contrasts for $\log_{10}$ basal metabolic rate are used in place of those for $\log_{10}$ specific basal metabolic rate.

[Table S121]

| Adj. $R^2$ | VIF | DW | p_F | $S\_\beta_1$ | $S\_\beta_2$ | $p\_\beta_1$ | $p\_\beta_2$ |
|---|---|---|---|---|---|---|---|
| 0.356 | 1.121 | 2.190 | 0.002 | 0.425 | -0.376 | 0.015 | 0.041 |
| 0.243 | 3.597 | 1.943 | 0.012 | 0.666 | -0.144 | 0.046 | 0.643 |

In Table S121 above, which uses phylogenetically adjusted data, even with adult body mass in the model, specific basal metabolic rate is a significant predictor of maximal lifespan across these ectotherm species.

The results with basal metabolic rate appear corrupted by collinearity (*VIF* = 3.597), conferring the standardized regression coefficient for basal metabolic rate, written in red text, with erroneous sign. Erroneous because, as can be seen in Table S122 below (in its zero-order column), maximal lifespan contrasts positively correlate with basal metabolic rate contrasts.

In Table S122 below are corresponding zero-order, partial, and semi-partial (part) Pearson correlation coefficients. Forced through the origin [323, 325]. One-tailed (alternative hypothesis is directional) p-values are italicized in brackets.

[Table S122]

| | Zero-order | Partial | Part |
|---|---|---|---|
| *MAXIMAL LIFESPAN vs.* | | | |
| Specific basal metabolic rate | -0.491 *(0.003968)* | -0.395 *(0.018749)* | -0.332 *(0.042171)* |
| Adult body mass | 0.541 | 0.462 | 0.403 |

| | | | |
|---|---|---|---|
| | *(0.001464)* | *(0.006661)* | *(0.016737)* |
| *MAXIMAL LIFESPAN vs.* | | | |
| Basal metabolic rate | 0.419 *(0.013318)* | -0.093 *(0.677736)* | -0.079 *(0.652355)* |
| Adult body mass | 0.541 *(0.001464)* | 0.388 *(0.020665)* | 0.352 *(0.033106)* |

In Table S122 above, its zero-order correlations are PDAP [320] outputted Pearson correlation (forced through the origin [323, 325]) coefficients, adjusted to account for phylogeny, which are statistically significant. The partial and part correlation coefficients (forced through the origin [323, 325]) are JASP output. For basal metabolic rate, its partial and part correlation coefficients, written in red text, are of erroneous sign. Probably because of the earlier shown collinearity, which made the regression coefficient for basal metabolic rate of erroneous sign. Collinearity, which breaches an assumption of multiple linear regression, and interrelated methods of partial and semi-partial (part) correlations, makes these basal metabolic rate results hard to interpret, possibly uninterpretable. But the specific basal metabolic rate related results don't have collinearity (*VIF*=1.121).

Table S122 above reports: specific basal metabolic rate is a significant predictor of maximal lifespan across these ectotherm species, even with the effects of phylogeny and adult body mass statistically excluded.

<u>Consolidation</u>

Interpreting all the data herein, lower specific basal metabolic rate universally confers higher maximal lifespan. Across endotherm and ectotherm species.

<u>Eq. 8 from Eq. 5</u>

Start with Eq. 5, which is of the form:

$$y = \frac{\ln{(x+2)}}{x}$$

Rearranging:

$$\ln(x+2) = yx$$

$$e^{\ln{(x+2)}} = e^{yx}$$

Because $e^{\ln(x)} = x+2$

$$x + 2 = e^{yx}$$

Solving above:

$$x = -\frac{W\left(-\frac{y}{e^{2y}}\right) + 2y}{y}$$

<u>Eq. 9 from Eq. 6</u>

Start with Eq. 6, which is of the form:

$$y = \frac{\ln(x)}{x}$$

Rearranging:

$$\ln(x) = yx$$

$$e^{\ln(x)} = e^{yx}$$

Because $e^{\ln(x)} = x$

$$x = e^{yx}$$

Solving above:

$$x = -\frac{W(-y)}{y}$$

<u>Supplementary content for the Results section titled "Anti-cancer"</u>

Consolidated absolute excess, and enantiomeric excess, data:

[Table S123]

| | | | | | | | |
|---|---|---|---|---|---|---|---|
| [From MassLynx 4.1 software on | | | \| | | | | |
| the Chiral SFC-MS machine used] | | | \| | ***[SAME data in different software: Mestrelab ]*** | | | |
| | **Chiral SFC** | | \| | **Chiral SFC** | | | |
| | **(UV chromatogram)** | | \| | **(UV chromatogram)** | | ***Chiral SFC-MS*** | |
| | Absolute(%) | ee(%) | \| | Absolute(%) | ee(%) | Absolute(%) | ee(%) |
| **6a** | 98.79 | 97.58 | \| | 98.83 | 97.66 | 99.64 | 99.28 |
| **6b** | 99.02 | 98.04 | \| | 98.23 | 96.46 | 97.9 | 95.8 |
| **7a** | 98.71 | 97.42 | | *Done on a different machine (other broke) ** | | | |
| **7b** | 100 | 100 | \| | 100 | 100 | *Can't calculate #* | |
| | | **Enantiomeric excess (ee) = (Absolute excess-50)*2** | | | | | |
| Wavelength (λ) for UV chromatogram = 220 nm | | | | | | | |
| * This other machine *only* has Chiral SFC capability. It is *not* a Chiral SFC-MS machine. | | | | | | | |
| Its shown data is *not* from MassLynx 4.1, but from Empower 3 software instead. | | | | | | | |
| This data won't open in Mestrelab (incompatibility). | | | | | | | |
| Chiral SFC-MS machine used = Thar analytical SFC with UV & mass detectors. | | | | | | | |
| Chiral SFC machine used = Waters Acquity UPC2 with PDA detector. | | | | | | | |
| # Without equivalent Chiral-SFC data for 7a, it isn't possible to calculate from the Chiral | | | | | | | |
| SFC-MS data of 7b. Because the Retention Time (RT) of 7a on this machine is unknown. | | | | | | | |

As shown earlier, the mean of 7a and 7b median anti-cancer activity (63.30) is slightly greater (0.54) than the mean of 6a and 6b median anti-cancer activity (62.76). Similarly, the mean of 7a and 7b mean anti-cancer activity (62.58) is slightly greater (0.85) than the mean of 6a and 6b mean anti-cancer activity (61.73). These differences are small enough (<1) to be within a margin of experimental error. This is their most likely explanation. But, alternatively, perhaps, they might reflect real features of the data. Because (*i*) 7b has slightly greater enantiomeric excess than 6b, (*ii*) 7a has slightly less enantiomeric excess than 6a. Meaning, for absolute excess values, (7b-7a) is 1.06 greater than (6b-6a). Wherein values 0.54, 0.85, and 1.06, all round to 1.

**SUPPLEMENTARY DISCUSSION**

Body heat balance equation [210, 376-377] (by 1st Law of Thermodynamics)

$$S = M - W - E - C - K - R \quad \text{[Eq. S16]}$$

Wherein,

*S* = Storage of heat in body (= 0 = heat balance: i.e., no change in body temperature. Positive = increased body temperature, negative = decreased body temperature).

*M* = Metabolic heat production (always positive in living organisms).

*W* = Work (positive = useful work accomplished, negative = mechanical work absorbed by the body).

*E* = Evaporative heat transfer (positive = transfer to environment).

*C* = Convective heat transfer (positive = transfer to environment).

*K* = Conductive heat transfer (positive = transfer to environment).

*R* = Radiant heat exchange (positive = transfer to environment).

*Units selected from consistently W/kg or W/m$^2$, where kg and m$^2$ relate to body mass and body surface-area respectively.*

From this equation, one can see that less *M* (e.g., by drug inhibition of $F_1F_0$ ATP hydrolysis) can be offset, such that *S* remains 0 (i.e., no change in body temperature), by more body insulation (e.g., more/better clothing) and/or higher ambient temperature, such that *E, C, K,* and *R* are less positive. One can also see that when ambient temperature = body temperature, as *E*, *C*, *K*, and *R* are all then 0, then *M* (and *W*) can be 0, and *S* is still 0: i.e., if the ambient temperature is 37°C, no metabolic heat generation is required for the body to be at 37°C: e.g., a mammal can even be dead and if the ambient temperature is 37°C its body temperature will be 37°C.

Future mouse lifespan study and (cheap, quick) derisking pre-experiments

There is interventional experimental evidence for causality between (*i*) lower specific $F_1F_0$ ATP hydrolysis rate and lower specific metabolic rate (Figure 2), and (*ii*) between lower specific metabolic rate and longer lifespan [21-22]. Combining this experimental data, causality can be inferred between lower $F_1F_0$ ATP hydrolysis rate and longer lifespan. Motivating, funding permitting, a future mouse lifespan (and/or surrogate endpoint/biomarker [378]) study. With the mice administered a selective $F_1F_0$ ATP hydrolysis inhibiting drug (*that doesn't inhibit $F_1F_0$ ATP synthesis*): e.g., from one or more of my international (PCT) patent applications [18-20]. At least compound 6b in Figure 18A is highly orally bioavailable, at least in rats [99], and so can be administered via chow, which facilitates a lifespan experiment.

Such a drug permits interventional power to investigate the inverse relationship observed within, and across many, mammal species between metabolic rate per unit mass and lifespan. What happens to lifespan if a mouse's metabolic rate per unit mass is drug reduced to that of a whale? When the mouse's ambient temperature is 37°C, meaning any metabolic heat generation is superfluous because the mouse's body temperature can be 37°C merely by equilibration with the ambient temperature. We might reveal - if greatly extending a mouse's lifespan thereby - that the lifespan difference between mice and whales is (at least in part) because a mouse performs more metabolic heat generation per unit mass, as it is smaller, so (by the square-cube law of geometry) has a larger surface-area-to-volume ratio, meaning it loses metabolically generated heat more readily.

A mouse has a heart rate of 618 beats per minute (bpm) [91] and a blue whale has a heart rate range of 2 to 37 bpm (lower end when diving, upper end when at the surface) [379]. So, a mouse has a very fast, and a blue whale a very slow, heart rate. I predict this disparity, with the mouse's large excess of bpm as compared to the whale, is (at least) mostly because of the mouse's greater metabolic rate per unit mass, because of its greater metabolic heat generation per unit mass per unit time: i.e., because of its greater specific $F_1F_0$ ATP hydrolysis rate. Such that drug inhibiting $F_1F_0$ ATP hydrolysis (without inhibiting $F_1F_0$ ATP synthesis) can reduce mouse heart rate to that of a whale. Wherein this is safe when in an ambient temperature of

37°C because then mice don't require their excess (to whales) of bpm, as they don't require any metabolic heat generation when in an ambient temperature of 37°C. Like this, when a mouse has a low heart rate, like a whale, I predict that its lifespan will be of the order of a whale.

Heart rate is an easily and quickly measurable physiological parameter. Assuming the ambient temperature is 37°C, I predict the amount of drug-conferred lifespan extension is predictable from the degree to which the heart rate is slowed by an administration routine of the drug: rearranging the plot equation from the 3rd (bottom) panel in Figure 6, gives an equation for maximal lifespan (*years*), *x*, in terms of heart rate (beats per minute, bpm), *y*:
[Eq. S17]

$$x = 10^{(\{log_{10}(y) - 3.26704\}/-0.91411)}$$

N.B. the constants of that equation have associated error bars, reported in the table of best fit lines of Figure 6 (Table 4; replicated here: 3.26704±0.247 and -0.91411 ±0.2212), which can be used to attribute error bars to *x* if desired.

Assessing the degree to which the drug can safely reduce mouse heart rate in an ambient temperature of 37°C is an incredibly cheap and quick experiment, which confers incredible predictive value for a mouse lifespan study. Using Eq. S17 to predict the higher maximal lifespan obtainable from the lower heart rate achieved.

In an experiment variant, it is found what a mouse's heart rate can be safely drug-reduced to, and maximal lifespan extension predicted therefrom using Eq. S17, when it is housed at typical room temperature (~20°C [171]). But in this case using Eq. S17 as a part, and not the totality, of a predictive equation. That also accounts, atop, for any lifespan extension due to any body temperature drop. Chemical reactions run slower at lower temperature and since aging is at least one chemical reaction, presumably aging is slower at lower temperature. The well-known $Q_{10}$ temperature coefficient equation can be found (for example) in [380-381], wherein it has two componentry reaction rates: $R_1$ (at temperature $T_1$) and $R_2$ (at temperature $T_2$), temperatures are in °C:
[Eq. S18]

$$Q_{10} = \left(\frac{R_2}{R_1}\right)^{10/(T_2 - T_1)}$$

$$R_2 = R_1 \cdot {Q_{10}}^{(T_2-T_1)/10}$$

Presumably, although aging rate increases with temperature, its inverse, maximum longevity, decreases with temperature and this is captured in the following derivative equation (*derived by swopping* $R_2$ *and* $R_1$*. Corresponding, in the second equation there is division instead of multiplication by the* $Q_{10}$ *containing term*):

[Eq. S19]

$$Q_{10} = \left(\frac{R_1}{R_2}\right)^{10/(T_2-T_1)}$$

$$R_2 = \frac{R_1}{{Q_{10}}^{(T_2-T_1)/10}}$$

Substituting earlier Eq. S17 into Eq. S19 above yields the equation below, which gives maximal lifespan (*years*), *x*, in terms of heart rate (*beats per minute, bpm*), *y*, accounting for any (safe) body temperature drop to new temperature $T_2$ from normal $T_1$ (temperatures in °C):

[Eq. S20]

$$x = \frac{10^{(\{log_{10}(y)-3.26704\}/-0.91411)}}{{Q_{10}}^{(T_2-T_1)/10}}$$

Where $Q_{10}$ = 2.125, justified later herein.

Incidentally, Eq. S20 above can correctly predict the maximal lifespan of the naked mole rat (31 years [1]) when its heart rate of 256 bpm [382] and its body temperature of 32°C [1] are inputted (the latter as $T_2$), and the constants are both increased to certain values within their (aforementioned) error bars.

Optionally, in pre-studies to any lifespan study (at any ambient temperature), first ensuring the drug dose to be used can decrease ROS generation in the body (expected possible given Figure 5). Or at least can decrease the rate of ROS generation increase with age (ROS generation rate increases with age [40-43]). Assayed by exhalation of ethane and/or pentane (lipid peroxidation products [383-396]). And/or assayed, probably more accurately, by one or more F2-Isoprostane species (lipid peroxidation products [42, 397-404]) in one or more bodily fluids, such as plasma (preferred) or urine. The production of all these lipid peroxidation species increases with age, a biomarker of aging, wherein the drug is expected to at least shallow their gradient of increase. In addition, or instead, using blood samples drawn over a sufficient time-period, assaying if such a drug's administration can slow the rate of "ticking" rate of the epigenetic clock, optionally using clock of [105].

These biomarker-of-aging experiments can be pre-studies to a mouse lifespan study (e.g., to optimize the dose(s) to be used) or perhaps conducted as stand-alone investigations, if one trusts such a biomarker(s) of aging to be a faithful read-out of aging itself.

Optionally, a drug-lifespan study can be started with already aged mice. Or with a mouse model of premature aging. For example, mice without SOD2, which die within 10 to 18 days from birth [228, 251], assaying if the drug can extend their lifespan to some statistically significant point beyond this. Alternatively, one could use a shorter-living species than mice: e.g., the forest shrew (*Myosorex varius*), which has a maximal lifespan of 2.1 years [1].

<u>A relevant mouse lifespan experiment may have already been conducted</u>

I hypothesize that the reason for the lower metabolic heat generation, and lower body temperature thereby, of the transgenic mice in [21] is their brain instructing cells of their body to upregulate activity of their natural, selective $F_1F_0$ ATP hydrolysis inhibitor (*that doesn't inhibit $F_1F_0$ ATP synthesis*): IF1 protein. If so, this would be a direct interventional connection already shown between higher IF1 protein activity, and lower $F_1F_0$ ATP hydrolysis rate, and longer lifespan.

IF1 protein, in mouse and human, has a phosphorylation control switch serine: S14 in the mature (without mitochondrial import sequence) IF1 protein sequence. IF1 protein is inactivated by said phosphorylation, thereby increasing $F_1F_0$ ATP hydrolysis rate. Phosphorylated by cyclic AMP (cAMP)-dependent Protein Kinase A (PKA) [405]. My hypothesis is that the transgenic and control mice differ in what fraction of their IF1 protein is phosphorylated at this S14 position and/or differ in IF1 protein amount, conferring lower specific $F_1F_0$ ATP hydrolysis rate in the transgenic mice.

The first author of [21] says (in personal communication) that he still has cryopreserved sperm for this transgenic mouse line, and so (at least for now) it is still possible to explore this. Leveraging mouse lifespan data already in hand.

Interventional *in vitro* test that $F_1F_0$ ATP hydrolysis is a drive of epigenetic clock ticking rate

In the main text, I interpret data as teaching that specific $F_1F_0$ ATP hydrolysis rate is a drive, via its dictating of specific basal metabolic rate and thence specific ROS generation rate, to the ticking rate of the epigenetic clock. Specific $F_1F_0$ ATP hydrolysis rate generating (most of) the metabolic heat generation per unit mass, in accordance with body surface-area per unit body mass.

An interventional test of this teaching would be to assay whether a selective $F_1F_0$ ATP hydrolysis inhibiting drug (*that doesn't inhibit $F_1F_0$ ATP synthesis*. E.g., from one or more of my international (PCT) patent applications [18-20]) can slow the ticking rate of the epigenetic clock of human tissue *in vitro*. In the assay used in [406-407].

In this assay the human tissue is kept at an ambient temperature of 37°C and so it doesn't require any metabolic heat generation to be at 37°C. So, all metabolic heat generation is redundant. Thence this drug can substantially reduce metabolic heat generation, substantially reducing metabolic rate thereby, whilst the human tissue temperature is maintained at 37°C by the ambient temperature. Thereby reporting metabolic rate's effect (if any) on the ticking rate of the epigenetic clock, a biomarker of aging.

Interventional *in vivo* test that $F_1F_0$ ATP hydrolysis is a drive of epigenetic clock ticking rate

Another possible experiment is to assay whether such a selective $F_1F_0$ ATP hydrolysis inhibiting drug (*that doesn't inhibit $F_1F_0$ ATP synthesis*) can slow the ticking of the epigenetic clock of mice, assaying this by analysing blood samples regularly drawn over a sufficient time-period, using the epigenetic clock of [105]. Testing the following (in part or totality) quantitative prediction: that drug-conferred slowing of a mouse's heart rate by an amount selected (by administered drug dose) from 1, 10, 20, 50, 100, 300, 500, 600, 617 beats per minute (bpm) will slow the ticking rate of their epigenetic clock by 0.1, 1.1, 2.2, 5.6, 11.4, 36.9, 69.6, 93.6 and 99.6% respectively. This prediction comes from substituting Eq. S17 into Eq. 5 to produce:

[Eq. S21]

$$\Delta M = \frac{ln\left(\left[10^{([log_{10}(y)-3.26704]/-0.91411)}\right]+2\right)}{10^{([log_{10}(y)-3.26704]/-0.91411)}}$$

Where *ΔM* is the average change in *M* per year (i.e., is the ticking rate of the epigenetic clock), *M* is a linear combination of DNAm (=β) values for selected DNA cytosines of the pan-eutherian epigenetic clock of [105], and *y* is heart rate (in beats per minute, bpm). So, for eutherians, this equation relates epigenetic clock ticking rate to heart rate. And it can be used to quantitatively predict how much the former slows for a given $F_1F_0$ ATP hydrolysis inhibiting drug (*that doesn't inhibit $F_1F_0$ ATP synthesis*) conferred slowing of the latter. By, to illustrate for a mouse:

$$\% = 100 - \left\{ \left( \frac{\Delta M \ when \ y = 618 - z}{\Delta M \ when \ y = 618} \right) \cdot 100 \right\} \qquad \text{[Eq. S22]}$$

Where *% is* percentage change in *ΔM* when heart rate (*y*) is *z* less, 618 is the normal heart rate of a mouse in bpm [91], and *z* is the amount in bpm that the heart rate is drug-conferred less.

High drug doses, conferring large drops in metabolic heat generation, metabolic rate, and heart rate, will likely require the mice to be kept at a higher than typical room temperature accordingly. Lower drug doses, conferring just small drops in heart rate, probably won't.

Eq. S21 assumes no body temperature drop. Combining Eq. S21 with earlier Eq. S18 gives Eq. S23 below that accounts for any safe body temperature drop from $T_1$ (normal body temperature) to $T_2$, each in °C, all be it the $Q_{10}$ value applicable here is presently unknown: [Eq. S23]

$$\Delta M = \left\{ \frac{ln\left(\left[10^{([log_{10}(y) - 3.26704]/-0.91411)}\right] + 2\right)}{10^{([log_{10}(y) - 3.26704]/-0.91411)}} \right\} \cdot {Q_{10}}^{(T_2 - T_1/10)}$$

For epigenetic clock analysis the mice blood samples can be sent to the non-profit Clock Foundation (https://clockfoundation.org/), which can use the clock of [105], or there are some companies offering a similar service (e.g., Zymo Research).

Optionally, this experiment can be done as a pre-experiment to a mouse lifespan study to optimize the drug dose(s) to be used for it. Maximal lifespan predicted from drug-decreased *ΔM* value using earlier Eq. 7.

Already shown to reduce a biomarker of aging

The ticking of the epigenetic clock is a correlate, a "biomarker", of aging. Less $F_1F_0$ ATP hydrolysis has already been shown to safely reduce a biomarker of aging in mice. Reducing intracellular Reactive Oxygen Species concentration (Figure 5), which is a biomarker of aging because it increases with age [40-43]. As both epigenetic and [ROS] change correlate with aging, but neither are (*presently*) unequivocally causal to aging, they are (*presently*) biomarkers of aging of equal standing.

Recently shown, gene therapy mediated partial reprogramming, with a decrease in epigenetic change rate observed, increases lifespan, with modest effect size (*study atypically reported increase in "median remaining lifespan" instead of median lifespan. Increase in the latter is 7%. Study started with aged mice*) [409]. In multiple studies, interventional decrease of ROS by antioxidants increases lifespan [389-391, 483-484, 500-520], and interventional decrease of metabolic rate increases lifespan [56-63], and so ROS is a biomarker of aging with more supporting interventional lifespan data. Including data from the Interventions Testing Program (ITP) [279].

Cosmetic testing

Probably the most used assay for testing a cosmetic's anti-aging capability is subjective assessment, optionally collected by questionnaire, from users of the cosmetic. Before and/or after it is on the market. *E.g., conducted by a Contract Research Organization (CRO) such as International Research Services, Inc, Port Chester, NY, USA.*

A less common but industry-recognized, more objective, assay of the anti-aging utility of a cosmetic is to test whether it can reduce the amount of Reactive Oxygen Species (ROS) inside skin cells or skin tissue *in vitro*. Less $F_1F_0$ ATP hydrolysis has ***already*** been shown to greatly decrease intracellular ROS safely *in vivo* (Figure 5). Therefore, a $F_1F_0$ ATP hydrolysis inhibitor compound (*that doesn't inhibit $F_1F_0$ ATP synthesis*) [18-20] is expected to succeed in this assay. Thereby demonstrating valuable cosmetic utility. Relatively cheaply and quickly. Potentially, soon bringing the fundamental discovery of why different mammal species age at different rates into first commercial application.

Reactive Oxygen Species (ROS) increase with age/aging [40-43], which is a biomarker of aging, like the ticking of the epigenetic clock for example, but its decrease by a drug might be detected faster, possibly in hours to days, rather than the months used in [406-407] for the *in vitro* epigenetic clock assay.

Options include assaying the amount of one or more of superoxide ($O_2^{\bullet-}$), hydrogen peroxide ($H_2O_2$), hydroxyl radical ($^{\bullet}OH$), and/or the amount of one or more lipid peroxidation products, such as malondialdehyde (MDA) and 4-hydroxynonenal (4-HNE), and/or protein carbonyls, and/or DNA oxidation products (e.g., 8-hydroxy-2'-deoxyguanosine). For non-limiting example, assayed by one or more of fluorescent probes, chemiluminescent substrates, colorimetric assays. Optionally using one or more of cultured skin cells (e.g., one or more of fibroblasts, keratinocytes, and melanocytes. *E.g., obtained from Coriell Institute for Medical Research, Camden, NJ or MatTek Life Science, Ashland, MA, USA*), and/or a three-dimensional (3D) skin model (reconstructed human skin model. *E.g., [409]*), and/or skin explants (excised from humans and maintained *ex vivo* in culture conditions. *E.g., obtained from ZenBio, Research Triangle, NC, USA*). Optionally conducted by a CRO (*e.g., www.dermaclaim.com*).

Even just with the supporting data already in hand, pending cosmetic safety tests, a cosmetic containing a $F_1F_0$ ATP hydrolysis inhibitor compound (*that doesn't inhibit $F_1F_0$ ATP synthesis*) [18-20] would be extremely differentiated in the cosmetics marketplace. Most cosmetics have little to no supporting data. Superior supporting data than competitor products might confer a niche in the multi-hundred-billion US dollars cosmetics market.

Humans with lower specific basal metabolic rate have a longer lifespan

The present paper interprets that metabolic rate, that is oxygen consumption rate, enables life, but also causes damaging byproducts and accumulating damage, which is aging, that ultimately takes life away. In this present section, consistent human data from the literature is discussed (with some supporting data from other species). In immediately subsequent sections, consistent data in the literature from some non-human species is discussed.

In the Baltimore Longitudinal Study of Aging, people with lower specific basal metabolic rate at entry to the study had *significantly* lower mortality over the subsequent >40 years of monitoring [29]. Above a specific basal metabolic rate (BMR) of 36.4 kcal/m$^2$/h at study entry, "the risk of death increased significantly as a squared function of BMR" [29]. The transform of [410] can convert a standardized mortality ratio difference into a corresponding difference in life expectancy ("a 1% decrease (or increase) in the standardized mortality ratio (SMR) will result in 0.1373 years increased (or decreased) life expectancy based on white male data for the US population"). Using this transform with data (already adjusted/controlled for age and various other risk factors) from Figure 2 of [29], humans with a specific basal metabolic rate in the range of 31.3-33.9 kcal/m$^2$/h have 3.90 and >7.32 years (e.g., 13.28 years) greater life expectancies than those with higher specific basal metabolic rates in the range of 33.9-36.4 kcal/m$^2$/h (*i.e. 7.82% more*) and >36.4 kcal/m$^2$/h (*i.e. >11.66% more*) respectively.

Humans with lower resting heart rate, which might indicate lower specific basal metabolic rate, tend to live longer over subsequent years of observation, in multiple different studies: e.g., [411-416]. Limited physical exercise (i.e., with sufficient time for the body to rest, recover, and adapt), which might prolong lifespan [417-420] (or not, perhaps artefactual [421]), lowers resting heart rate in humans [422-424], which might indicate lowered specific basal metabolic rate. Indeed, at least in mice, data suggests that exercise (*perhaps so not as much food is required despite the exertion of exercise, that completed and if repeated in the future*) causes lower specific basal metabolic rate, at least in part by reducing metabolic heat generation, and body temperature drops accordingly [187], wherein there is (in part) consistent data in humans [425]. So, to speculate, physical exercise itself may *not* confer benefit, but instead the body's response/adaptation to it. And if physical exercise is such that the body doesn't have the time to make and/or benefit from such a response it might be detrimental to health. For example, when doing long, hard, forced labour every day: e.g., in a gulag. Perhaps any benefit from exercise should be reframed as benefit from "exercise and sufficient rest", to discriminate from exercise and insufficient rest, which might be harmful. Perhaps consistently, kestrels whose brood size is manipulated smaller (less work and energy expenditure) live longer, and those whose brood size is manipulated larger (more work and energy expenditure) live shorter [426]. Bees with weights attached to them to make flying more energetically costly live shorter [427], and houseflies restricted from flying (housed in small jars), decreasing their energy expenditure, live three times longer [428].

Something to keep in mind when reading the literature: it is conceivably possible for *decreased* specific basal metabolic rate (i.e., per unit mass) to correspond with *increased* basal metabolic rate upon weight gain: e.g., of muscle mass.

Tentatively extrapolated to be interventional evidence, that lifespan is inversely proportional to specific basal metabolic rate, is that calorie restriction slows metabolic rate (beyond the effect of weight loss. Decreasing oxidative damage) in humans [33-34], wherein calorie restriction can extend lifespan in rodents and primates [31-32] (*no study has yet explored the effect, if any, of calorie restriction upon human lifespan*).

At least some species interventionally made to have a lower metabolic rate live longer

Chemical reactions run slower at lower temperature (*the exponential dependence of chemical reaction rate upon temperature can be seen in the Arrhenius [366] and Eyring [367] equations {the latter having an additional pre-exponential temperature term}*). The body temperature of cold-blooded animals is that of ambient temperature. So, the metabolic rate (which is comprised of chemical reactions) of cold-blooded animals can be reduced by reducing ambient temperature. Lower ambient temperature, reducing their metabolic rate, makes cold-blooded animals live longer. At least as observed in *D. melanogaster* (fruit fly) [24-25], *Calliphora stygia* (a blowfly) [26], *C. elegans* (nematode worm) [23], *Trichogramma platneri* (a wasp) [27], and *Cynolebias adloffi* (a fish) [28].

But a complication to these cold-blooded animal studies is that the extended lifespan is concordant with two different variables changed. Lower body temperature and lower metabolic rate. And so, we can't be certain which the extended lifespan is because of. I suggest it is because of both. Because of:

(i) Lower metabolic rate (i.e., lower oxygen consumption per unit time) meaning less Reactive Oxygen Species (ROS) generation per unit time.

(ii) And the lower body temperature slowing ROS generation, ROS-caused molecular damage, and repair thereof also, but since there is *net* damage over repair (damage outruns repair), there is *net* benefit to lower temperature.

*Indeed, since aging must comprise at least one net chemical reaction, and (by a well-known law of chemistry) chemical reactions run slower at lower temperature (e.g., why food perishes slower in a fridge), deductively aging should be slower at lower temperature.*

Lower metabolic rate extends the lifespan of mice

Interventional evidence for causality between lower specific basal metabolic rate and higher maximum lifespan is that an interventional decrease in metabolic rate can extend lifespan. As shown in the work of [21-22]. Where mice with less metabolic heat generation (consuming less $O_2$/kg/hour than control mice: i.e., with a lower metabolic rate), and lower mean body temperature (average of 0.3°C and 0.34°C less in males and females respectively), had an increased median lifespan (by 12% and 20% in males and females respectively). Wherein females had greater mean body temperature drop, and greater median lifespan extension.

This lower metabolic rate was conferred by overexpressing uncoupling protein UCP2 in hypocretin neurons of the hypothalamus. Selectively making this brain region hotter, so that the nearby preoptic area (POA) was also hotter, fooling it into thinking the rest of the body was warmer than it was, so that it directed less metabolic heat generation.

But there is an ambiguity here, because the lifespan increase could be due to the decreased metabolic heat generation (lower specific metabolic rate) and/or the lower body temperature caused thereby. I estimate it is due to both, as argued now.

The transgenic mice of [21-22], which have a longer median lifespan, have (*a*) less metabolic heat generation (and a lower metabolic rate specifically because of this) and (*b*) a lower body temperature (and a lower metabolic rate specifically because of this). In the previous section relating to cold-blooded animals there was only (*b*), and no (*a*). I use this disparity to try to estimate how much of the lifespan extension of these mice is due to (*a*).

Using the transgenic female mice data (*sourced from the supplementary information of [21]. Assuming the wild-type mean body temperature is 37°C [1]*), the $Q_{10}$ temperature coefficient [380-381] for their lower mean body temperature (0.34°C lower) and longer median lifespan (+20%) is, using Eq. S18, repeated here, $(R_2/R_1)^{10/(T_2-T_1)} = (662/550)^{10/([37-0.34]-37)} = 0.00429033474$. A fraction because higher temperature decreases (rather than increases) the temperature-dependent variable, median lifespan. If the $Q_{10}$ equation is modified (swopping around $R_2$ and $R_1$; earlier Eq. S19) to instead make $Q_{10}$ >1 in this case: $(550/662)^{10/([37-0.34]-37)} = 233.08204613$ (for a temperature range of 36.66-37°C).

For ectotherms (which don't have metabolic heat generation), the $Q_{10}$ value for ambient temperature (which sets their body temperature) and mean lifespan is, using this modification of the $Q_{10}$ equation (Eq. S19): 2 (for fruit fly *Drosophila melanogaster*, for a temperature range of 9-34°C) [25] or 2.25 (for blowfly *Calliphora stygia*, for a temperature range of 12-34°C) [26]: mean = 2.125.

I assume what is observed in ectotherms teaches us the order of magnitude of a lifespan effect when there is no aforementioned ($a$), and only ($b$). And so, the remainder (overwhelming majority. $100 - \{(2.125/233.08204613) \cdot 100\} = 99.09\%$) must be due to ($a$) [*actually more because of the different temperature ranges of the $Q_{10}$ values, wherein $Q_{10}$ is less with higher temperature ranges*]: i.e., due to lower metabolic heat generation (lower metabolic rate thereby). And so is not conditional upon any body temperature drop. It can cause lower body temperature, but presumably only if the ambient temperature is conducive to that.

To summarize this analysis: majority of the lifespan increase is because of less metabolic heat generation (and lower metabolic rate thereby), which doesn't necessarily cause a body temperature drop because it can be compensated for by increased body insulation and/or ambient temperature. But lower body temperature, if it does occur, and is within safe limits, can conceivably extend lifespan further. Indeed, as presented in the next section, humans with lower body temperatures tend to live longer.

Slightly lower body temperature corresponds with a much longer lifespan in humans

Resting body temperature varies across different people in a 2.5°C range, from 35.2 to 37.7°C [168, 429]. People with lower resting body temperature have a longer healthspan [183-184] and lifespan [30, 168, 181-182].

In a set of humans, mean body temperature ranged (2.2°C) from 35.2 to 37.4°C in [429]. And (2.4°C) from 35.3 to 37.7°C, across 35,488 human subjects, in [168]. And human rectal temperature data ranged (3.4°C) from 34.4 to 37.8°C in [430] (this paper consolidates data from multiple studies). In humans, in the Baltimore Longitudinal Study of Aging, people with a body temperature in the bottom half of the observed body temperatures (at study entry) had lower mortality over the subsequent 25 years of monitoring [30]. Analysis of 677,423 human body temperature measurements, spanning 157 years (1860–2017), showed that mean body temperature has decreased 0.03°C per birth decade (men born in the early 19th century had a body temperature 0.59°C higher than men today), alongside an increase in median lifespan over the same time-period [181]. In a 18,630 people cohort (ages in the range of 20-98 years old): >60 years old, the proportion of people with a mean body temperature <35.56°C was more, whilst the proportion of people with a mean body temperature >36.67°C was less, possibly reflecting increased survival of individuals with lower mean body temperature (and/or a decrease of mean body temperature with age) [182]. In a large cohort of (35,488) humans, higher mean body temperature was linked to higher one year mortality [168]. In humans, lower early morning body temperature correlates with better physical (e.g., faster walking speed) functioning in aged individuals [183]. And lower body temperature correlates with better mental functioning in aged humans [184].

Calorie restriction slows aging and extends lifespan in model organisms, such as rodents and primates [31-32]. It causes a lower body temperature [35], by as much as 5°C in some mouse strains [36]. Calorie restriction decreases body temperature in humans [37-39]. Calorie restriction is the best intervention for life extension that we have found to date: e.g., increasing the mean and maximal lifespans of mice by 65% and 51% respectively [31]. Rapamycin drug, which can slow aging and extend lifespan in mice [162], can decrease the body temperature of mice [431]. Ames dwarf mice have a 1.6°C lower mean body temperature than normal sized mice [432], with a mean lifespan ≥68% longer in females, and 49% longer in males [358]. At least in mice, lower resting body temperature is observed after exercise [187].

Difference in human life expectancy between higher and lower body temperature groups

Using data from the Baltimore Longitudinal Study of Aging, Fig. 1D in [30] plots cumulative survival vs. time (over 25 years; corrected for initial subject age). With two plots: one for men with a body temperature below, and the other for men with a body temperature above, the median body temperature of men in the study (recorded upon entry into the study; there was no significant difference in subject age between the two groups at the start of the study; subject ages ranged from 18 to 91 upon study entry; 324 men died over the 25 years of observation). I used (aforementioned) WebPlotDigitizer to pull representative data points from one of the plots. And then did the same for the other plot. I then opened the data for the lower body temperature group in OriginPro 8 software (OriginLab Corporation, Northampton, MA, USA), plotted it as a scatter plot, and then fitted a polynomial curve to it. Using this fitted polynomial equation $\{y = 0.99534 + (0.00101 \cdot x) + ([-5.12587 \cdot 10^4] \cdot x^2)\}$ I generated further data points to extrapolate this plot to y = 0 (all subjects dead). Assuming the curve follows the same trajectory until all subjects die. I then integrated this plot. Its mathematical area (under the curve) was 29.765031340107, which I then divided by the y-axis value at the beginning of the time period (1), which returned 29.765031340107. Which is the life expectancy of that group in years (from the start of the study period, not from birth). A notional life expectancy because the x-axis isn't subject age but time since start of observation. [433] guided this technique I used, to get (notional) life expectancy from a survival plot (dimensionally correct because the y-axis has the dimension of persons, the x-axis has the dimension of years, so the area under the curve has the dimension of person years, and when divided by the number of persons alive at the beginning, the dimension is years). I then repeated this methodology for the other plot. That for the higher body temperature group. Which returned {its polynomial plot: $y = 0.99673 + ([4.39509 \cdot 10^{-4}] \cdot x) + ([-5.87422 \cdot 10^{-4}] \cdot x^2)$} a notional life expectancy of 27.394083088806 years. So, a difference in life expectancy of 2.3709482513 (=2.4) years, i.e. men with a lower body temperature have a greater life expectancy. A significant difference (to at least $p < 0.05$), inheriting significance from that shown for this data by [30]. Although acknowledging that my calculation is assumptive and abstractive.

Hereafter I tentatively extrapolate this data (from a sample of the US population) to the entire US population. If the half of men with higher than median body temperature, and 2.4 years less life expectancy, were to maintain an effective amount of a selective $F_1F_0$ ATP hydrolysis inhibiting drug (*that doesn't inhibit $F_1F_0$ ATP synthesis*) in their system, by regular oral administration, such that their body temperature was lower than the original median body temperature (with the same distribution of values below it than the other half of men), then the overall life expectancy of all men would be increased by 1.2 years. Assuming this effect could be replicated for woman also, the overall life expectancy for both genders would be increased by 1.2 years. Economists calculate that a slowdown in aging, which increases US life expectancy by 1 year, is worth $38 Trillion US dollars [434].

To extrapolate further, an even greater increase in life expectancy might be conferred if everyone's resting body temperature was bought down to, not just below the original median resting body temperature, but to the lowest values naturally observed within the human population. Or possibly even lower. As aforementioned, mice with 0.34°C lower mean body temperature have 20% greater median lifespan [21]. Wherein the body temperature of many people could be reduced by at least 0.34°C and their body temperature would still be within the range of resting mean body temperature observed within the human population (which is 35.2 to 37.7°C: i.e., a 2.5°C range [168, 429]).

A drug-conferred body temperature difference of 0.34°C is anticipated to be imperceptible to a human because our body temperature imperceptibly varies by more than that (0.7 to 1.6°C) during a day (following the circadian rhythm) [188]. And more than that over the course of a month in women, whose body temperature varies with their menstrual cycle (0.3 to 0.7°C body temperature difference between the luteal and follicular phases) [435]. Human body temperature can vary with season, being around 0.2°C cooler in winter [436]. Human body temperature is imperceptibly lower during calorie restriction. For example, mean human body temperature is 0.2°C lower during calorie restriction (0.3°C lower in a calorie restriction and exercise group) in [37-39], and 0.5-0.8°C lower during semi-starvation in [39]. Conceivably, administration of a $F_1F_0$ ATP hydrolysis inhibiting drug (*that doesn't inhibit $F_1F_0$ ATP synthesis*) allows one to have the cooler body temperature of calorie restriction, which might be componentry to how calorie restriction extends lifespan [185]. Thereby acting as a calorie restriction mimetic.

To put such increased life expectancy in perspective: curing/eliminating all cancers would increase US life expectancy (from birth) by 3.2 years [437].

In humans, the variation in resting body temperature [30, 168, 181-184, 429-430], which correlates with the variation in life expectancy [30, 168, 181-182], might be due (at least in part) to variation in specific $F_1F_0$ ATP hydrolysis rate (due to variation in specific IF1 protein activity, per the specific amount of ATP synthase). The lower body temperature of humans undergoing calorie restriction [37-39] might be due (at least in part) to lower specific $F_1F_0$ ATP hydrolysis rate (due to greater specific IF1 protein activity, per the specific amount of ATP synthase).

<u>Alzheimer's disease increases body temperature, which accelerates/exacerbates its pathology</u>
People with Alzheimer's disease have a higher body temperature [438], which (at least in an Alzheimer's mouse model) accelerates/exacerbates the disease, wherein lower body temperature slows Alzheimer's disease progression, slowing its amyloid plaque build-up in the brain and, critically, its cognitive decline [439]. Perhaps to be expected because Alzheimer's disease progression comprises at least one chemical reaction and, by the laws of

chemistry, chemical reactions run *exponentially* faster at higher temperatures. So, lower (e.g., not elevated, but normal) body temperature can slow Alzheimer's disease progression without us having to know what causes the disease. A selective $F_1F_0$ ATP hydrolysis inhibitor drug (*that doesn't inhibit $F_1F_0$ ATP synthesis*) can (*if ambient temperature is sufficiently low: e.g., at typical room temperature: ~20°C [171]*) dose-dependently lower body temperature (shown in mice in Figure 2). Slight decrease of body temperature is known to be safe because it occurs during calorie restriction [35-39] (which extends lifespan [31-32]). Rectifying the body temperature of Alzheimer's disease patients to normality could be the endpoint, or the surrogate endpoint (for the accelerated approval program [440]), of such a drug's clinical trial(s) for FDA approval.

In a mouse model of Alzheimer's disease (AD), raising body temperature (by 0.50–0.79°C) results in enhanced BACE1 expression, more amyloid beta (Aβ) generation, greater Aβ plaque area, greater amount and hyperphosphorylation of tau, and worse memory (in a maze task. "Our study highlights an impact of high body temperature on the progression of AD pathologies, providing a mechanistic insight for its prevention") [439]. From early in disease progression, mice of a mouse model of Alzheimer's disease have greater mean body temperature than control mice (0.2-0.4°C more, statistically significant), which cannot be lowered by ibuprofen (an antipyretic) administration [441]. Body temperature is increased (at statistical significance), by around 0.1°C, in people with Alzheimer's disease (from a meta-analysis, summarizing six independent studies, with a total of 90 Alzheimer's disease patients and 78 healthy controls [438]). Perhaps (at least in part) because of thermogenesis by Aβ aggregation [442]. Where, *in vitro*, greater temperature promotes greater Aβ production [438] and greater Aβ aggregation [442, 444-447]. Thence positive feedback is possible: Aβ aggregation generates heat, driving more Aβ aggregation, which generates more heat, in a runaway loop.

Systemic administration of a selective $F_1F_0$ ATP hydrolysis inhibiting drug (*that doesn't inhibit $F_1F_0$ ATP synthesis*), which (*if ambient temperature is sufficiently low, e.g. at typical room temperature: ~20°C [171]*) dose-dependently lowers body temperature, might interdict this runaway pathology (e.g., the drug of Figure 2 in the main text). Returning body temperature to within the normal range, optionally to the low end of this range, or even lower. This should reduce Aβ production and aggregation. Conferring the benefit that it is likely to simultaneously slow all the steps up to Aβ plaque formation, for all the different lengths of Aβ peptides, inside and outside cells.

Particularly, this drug should reduce ROS generation, as per the teaching of Figure 5 in the main text. Note that Aβ monomers are antioxidants and might be produced in such pathological, aggregating quantities in response to high [ROS] [448-457]. Their amount dictated by the amount of BACE1 [458-463], whose gene promoter senses [ROS] [458]. But when the Aβ concentration gets too high, Aβ aggregates and thereby some report [449] it becomes pro-oxidising. ROS might be the cause of Alzheimer's disease [459, 464-492]. Which might be why drugs operating to the "amyloid hypothesis" [493-495] have underwhelmed to date, because they are merely acting upon a response to the cause, which may even be attenuating to the cause, at least at lower response levels.

[ROS] in the body increases with age [40-42]. Much data suggests that oxidative damage could be an early, arguably the earliest, event in Alzheimer's disease [496-517]. In mice models of Alzheimer's disease, interventional decrease of oxidative stress decreases BACE1 expression and activity, decreases the amount of Aβ and plaques thereof, decreases tau amount and phosphorylation, decreases neurofibrillary tangles, improves memory, and increases lifespan [518-521]. Whereas interventional increase of oxidative stress increases the amount of Aβ and plaques thereof, increases tau phosphorylation, and accelerates the behavioural pathology [518, 522-528]. Two different antioxidants, one of them being Vitamin E (locates to cell membranes), were each separately shown to modestly slow the decline in performance of activities of daily living and the need for care, but not the decline in cognitive test scoring, in human Alzheimer's disease patients in a double-blind, placebo controlled, randomized, multicentre clinical trial [528]. Mice studies indicate that the result might have been better if this clinical trial had utilized patients earlier in the course of the disease [527]. Human epidemiological studies show dietary and/or supplemental antioxidants decrease the risk of Alzheimer's disease [528-530]. The flawed approach of ROS mitigation (antioxidants) was contrasted with the merit of decreasing ROS generation (by selective $F_1F_0$ ATP hydrolysis inhibition, Figure 5) in the main text.

A selective $F_1F_0$ ATP hydrolysis inhibiting drug (*that doesn't inhibit $F_1F_0$ ATP synthesis*) is anticipated to slow aging, as it can (*if ambient temperature is sufficiently low, e.g. at typical room temperature: ~20°C [171]*) dose-dependently lower body temperature (Figure 2) and slightly lower body temperature corresponds with a much longer lifespan in mice [21] and humans [30, 168, 181-182] (and a much longer healthspan in humans [183-184]). Which is relevant here if Alzheimer's disease correlates with age/aging because aging is a causal drive of it. Alzheimer's disease may correlate with age/aging because aging is a drive to (and of) it. For clinical trialling, a simple, quick clinical endpoint can be used. Assaying whether this drug can reduce the pathologically high body temperature of Alzheimer's disease [438] to normality. If it can, this should be sufficient for clinical approval, for helping patients with Alzheimer's disease (especially because data suggests that higher body temperature accelerates Alzheimer's disease progression [439]). Optionally screening a population of Alzheimer's disease patients and (by setting the inclusion/exclusion criteria as such) only admitting those with the highest body temperatures into the clinical trial.

Aducanumab is a human monoclonal antibody drug that binds aggregated Aβ. Weekly, intraperitoneal (i.p.) administration of (a murinized analogue of) aducanumab to Tg2576 mice (a mouse model of Alzheimer's disease), starting at 9.5 months old, assessed at 15.5 months old, at the highest dose tested (30 mg/kg), reduced Aβ plaque amount in the cortex and hippocampus by up to 63% (Figure 4c-d in [532]. When instead starting drug administration at 18 months old, assessed at 24 months old, there was failure to reduce Aβ plaque amount [533]). To contrast, in mice of the same mouse model of Alzheimer's disease (Tg2576, starting at 4 months old, assessed at 17 months old), those with a body temperature of 0.50–0.79°C (*middle of this range is 0.65°C, which is less than the variation in resting body temperature across different humans [168, 429], and is less than the intraday variation in resting body temperature in humans [188]*) lower had up to 50% less Aβ plaque amount in their cortex and hippocampus (Figure 3a in [439]). Distinctly, whilst there is no published data showing that aducanumab can improve cognition in a mouse model of Alzheimer's disease, there is such data for a slightly lower body temperature, showing that it confers better memory [439]. So, at least in mice, slightly lower body temperature reduces Aβ plaque amount by nearly as much as the highest pre-clinical dose (30 mg/kg i.p.) of aducanumab. Less Aβ plaque amount (measured by PET imaging) is a surrogate endpoint for "accelerated approval" for Alzheimer's disease by the United States Food and Drug Administration (FDA) [440]. As conferred upon aducanumab, which is now on the US market, with the caveat that a further "post-marketing" (Phase IV) trial be conducted to get further data for it, especially pertaining to efficacy against cognitive decline [5334-536]. If this isn't positive, its clinical approval will be rescinded. In two Phase III trials to date, restricting entry to patients early in the course of the disease (but that do have Aβ plaque burden observed by PET imaging), although aducanumab certainly reduced Aβ plaque amount, whether it slowed cognitive decline (assayed by Clinical Dementia Rating scale Sum of Boxes, CDR-SB, scoring at 18 months) was equivocal in one of these trials. UPDATE (written in 2024): since writing this paper section (in 2022) Biogen has discontinued aducanumab.

Given the (aforementioned) implication of ROS in Alzheimer's disease, a good surrogate endpoint (that could be proposed to the FDA for their accelerated approval program) would be to test if a candidate drug can decrease ROS generation in the body. Assayed by less exhalation of ethane and/or pentane (lipid peroxidation products [383-396]). And/or assayed, probably more accurately, by less of one or more F2-Isoprostane species (lipid peroxidation products [42, 397-402]) in one or more bodily fluids, such as plasma, urine, or cerebrospinal fluid (CSF). These molecular species are elevated in Alzheimer's disease patients as compared to control subjects [537-538], correlating with disease severity [539]. They're also elevated in patients with Mild Cognitive Impairment (MCI), which is prodromal to Alzheimer's disease, as compared to control subjects [517].

My interpretation (of the data in hand to date) is that decreasing Aβ plaque area, as aducanumab does, is acting upon a downstream effect, whilst reducing ROS is to intervene at the upstream cause of the disease. Where Alzheimer's disease is caused, as aging is caused, by ROS. Indeed, arguably Alzheimer's disease *is* aging. Subclinical in many elderly people, who ultimately succumb to another facet of an aging body.

In two different mouse models of Alzheimer's disease, calorie restriction reduces Aβ plaque number and size [540]. Perhaps (at least in part) because calorie restriction can decrease body temperature [35-39]. However, calorie restriction is arguably dangerous in Alzheimer's disease patients because of the following. Some Alzheimer's disease patients, despite such patients tending to have equivalent or more/hyperphagic food intake [541-545], tend to lose body weight [541-543, 546-556] (*incidentally, such weight loss was first reported by Alois Alzheimer himself*), an increasing amount as their disease progresses, wherein those Alzheimer's disease patients that instead maintain or gain body weight have lower mortality [555-556] (perhaps relatedly, multiple mouse models of Alzheimer's disease exhibit hypermetabolism [557-560] and at least one of these models has an elevated body temperature [441]). Transgenic mice with less metabolic heat generation, and so slightly lower body temperature, have higher body weight (with the same food consumption as control mice) [21]. Presumably because of greater metabolic efficiency, with less of the chemical energy of food dissipated as heat. A selective $F_1F_0$ ATP hydrolysis inhibiting drug (*that doesn't inhibit $F_1F_0$ ATP synthesis*) increases metabolic efficiency, decreasing the fraction of food energy dissipated as heat (Figure 2), which is expected to counteract weight loss. Counteracting (e.g., slowing of) weight loss is a very objective, tractable clinical trial endpoint/surrogate endpoint.

There is presently no proven treatment for Alzheimer's disease ("for disease-modifying therapies (DMTs), the failure rate has been 100%" [561]). A huge unmet medical need. The number of people with the disease is increasing every year as populations age [562]. Its financial burden to the USA alone, which was $259 billion per year in 2019, is projected to exceed $1 trillion per year by 2050 [561] ($9.1 trillion per year worldwide [563]). "Alzheimer's disease poses a grave and growing challenge to our Nation" [564]. An efficacious drug for Alzheimer's disease could become the best-selling drug (by sales revenue) on the market. Exceeding the present leader, Humira (for rheumatoid arthritis), which had $20.4B of sales in 2020 alone [565]. Because, for one reason, the market size for Alzheimer's disease is a multiple of that for rheumatoid arthritis, probably a double-digit multiple [564, 566].

To summarize this section: Alzheimer's disease patients can have higher body temperature [438], where higher body temperature has been shown to accelerate/exacerbate the disease in a mouse model of Alzheimer's disease (e.g., causing greater Aβ plaque area and worse memory) [439]. A selective $F_1F_0$ ATP hydrolysis inhibiting drug (*that doesn't inhibit $F_1F_0$ ATP synthesis*) can (*if ambient temperature is sufficiently low: e.g., at typical room temperature: ~20°C [171]*) dose-dependently lower body temperature (shown in mice, Figure 2) and thereby is predicted to delay/slow/combat Alzheimer's disease. Moreover, Alzheimer's disease correlates with age/aging, probably because aging is (at least in part) causal to it, and slightly lower body temperature corresponds with much slower aging in mice and humans [21, 168, 30, 181-184]. Furthermore, many papers, and much data, suggests that ROS are causal to Alzheimer's disease [87, 40-41, 448-492, 496-516, 518-531]. And selective inhibition of $F_1F_0$ ATP hydrolysis (*without $F_1F_0$ ATP synthesis inhibition*) decreases ROS generation (Figure 5).

Potential clinical utility of local administration

There is a thesis that neurodegenerative diseases, such as Alzheimer's (which can cause dementia) and Parkinson's disease, correlate with aging because they are caused by aging. Indeed, caused by the oxidative stress of aging [459, 464-492]. "Mitochondrial dysfunction and oxidative stress occur early in all major neurodegenerative diseases, and there is strong evidence that this dysfunction has a causal role in disease pathogenesis" [475]. More broadly, many further diseases have an exponentially increased risk of incidence with age. And can be said to be diseases of aging. Many, if not all, diseases of aging might be caused by a body part(s) aging, and declining in function, faster than the rest of the body. So, such diseases might be prevented/ameliorated by locally slowing the aging of that body part(s). So that it better matches the aging rate of the rest of the body. Or is even slower. This can better match healthspan to lifespan. For example, dopamine neurons in the pars compacta, in the substantia nigra, decline in number by 5-10% per decade, presumably by dying, which is a predisposing drive to Parkinson's disease [567-569]. Speculatively extrapolating from the data herein, specifically reducing $F_1F_0$ ATP hydrolysis (*without reducing $F_1F_0$ ATP synthesis*) in these neurons will slow their aging rate. Where their resulting reduction in metabolic heat production will be offset by heat transfer from other cell types and body regions, especially via blood flow. Another postulated use case is topical administration of a selective $F_1F_0$ ATP hydrolysis inhibitor drug (*that doesn't inhibit $F_1F_0$ ATP synthesis*) to the eye(s), to avert/slow/combat Age-related Macular Degeneration (AMD). There is presently no treatment for dry AMD, which more than 150 million people have now, projected to increase to nearly 300 million people by 2040 [570].

Some further medical applications?

As taught by this work, a systemically administered selective $F_1F_0$ ATP hydrolysis inhibitor drug (*that doesn't inhibit $F_1F_0$ ATP synthesis*) can reduce body temperature to a value controlled by the intersection of drug dosage and ambient temperature. Even at maximum possible effect, the drug cannot make body temperature fall below, only towards/to, ambient temperature. So, at high (systemically administered) drug dose, body temperature can be controlled by controlling ambient temperature.

This might have medical applications. For example, emergency grade hyp*er*thermia is an extremely dangerous aspect to many Emergency Room (ER) admissions. For example, in some trauma patients. And there are many medical conditions in which the body can get too (dangerously) hot: e.g., fever, sepsis, malignant hyperthermia, neuroleptic malignant syndrome, adverse drug reaction, hypermetabolism, etc.

It may also enable controlled hypothermia for a surgical/protective purpose (called "Targeted Temperature Management, TTM", "therapeutic hypothermia", or "protective hypothermia"), for which there are many applications: e.g., neuroprotection after a stroke, cardioprotection after a heart attack, deep hypothermic circulatory arrest for surgery (e.g., brain surgery: e.g. brain tumour removal, where body temperature is made 20-25°C, currently induced crudely), or for slowing the process of dying after one or more of injury, poisoning, and infection, giving more time to be saved. For example, after a gunshot wound, conferring longer than the "golden hour" to get an injured soldier to hospital.

<u>Historical odds of progressing into clinical trials</u>

The compound used to produce Figure 2 has therein shown efficacy in mice. For seeking FDA approval, its next step is testing in "Non-clinical toxicology" studies (also called "IND-enabling studies") in two mammalian species, one of which must be a non-rodent. For example, rat and dog. If passing those it can proceed into human clinical trials. It is interesting to look at historical data to see how often drug candidates succeed in passing this Non-clinical toxicology step, to make it into human clinic trialling. Analysing the pooled small-molecule pipelines of four pharmaceutical companies, over ten years, looking at pre-clinical compounds nominated for further pursuit [571]: for 386 compounds that made it into the clinic, 211 compounds failed Non-clinical toxicology (legend of Figure 6 in [571]): 100-((211/(211+386))*100) = 64.66% penetrance for this Non-clinical toxicology step. Some or all of the other 145 compounds that were dismissed pre-clinically, for a reason unrelated to Non-clinical toxicology (Figure 2c of [571]. *For example, 75 compounds were dismissed because of "rationalization of company portfolio", and 26 compounds were dismissed because of "commercial" reason*), might have passed Non-clinical toxicology beforehand. If we assume all of them did, then the success rate is instead: 100-((211/(211+386+145))*100) = 71.56%. 60 pre-clinical compounds are labelled "progressing" in Figure 1 of [571], which might mean that all these compounds had already passed Non-clinical toxicology, and so were ready to progress into the clinic, or that was only the case for zero or more and one or more others were still undergoing (or yet to undergo) Non-clinical toxicology. If we assume the former, then the success rate is instead: 100-((211/(211+386+145+60))*100) = 73.69%. Alternatively, more conservatively, these "progressing" pre-clinical compounds are dismissed (by assuming they have the same proportional outcomes as the other pre-clinical compounds). In which case the historical penetrance for Non-clinical toxicology (IND-enabling studies) into the clinic is somewhere in the range 64.66-71.56%. Splitting the difference = 68.11% = 68%. The probability can be projected as higher still for this present compound (>68%) because if it is found to be only systemically toxic its use can be restricted to local administration, as Unity Biotechnology did with its "senolytic" drugs [196]. Such local administration is still anticipated to have much utility.

Hypothesis

The body can repair/replace damage caused by metabolic rate. But damage outruns repair/replacement, conferring *net* damage, which is aging. Reducing metabolic rate reduces rate of damage, slowing aging, and reversing aging if the rate of damage can be made less than the rate of repair/replacement.

The rate of damage caused by Reactive Oxygen Species (ROS) is a linear function of [ROS]. Whilst the repair/replacement of ROS damage is done by enzymes and so its rate is a Michaelis–Menten function of [ROS] [145, 572]. So, with less [ROS] there is less damage *and* repair/replacement. But given their different gradients, there is a *net* reduction in damage. If [ROS] is decreased enough, and the amount of repair/replacement enzyme(s) is sufficient, then the rate of repair/replacement of ROS damage can exceed the rate of ROS damage, which means prior ROS damage (*assuming* it can still be recognized as damage) can get repaired/replaced. In this case, the rate of ROS damage doesn't just slow, but is reversed. Illustrated, for the case of DNA damage in Figure S18. But equally applicable to other forms of ROS damage. Such as (hypothesized herein) ROS-driven "ticking" of the epigenetic clock. In experiment, selectively inhibiting $F_1F_0$ ATP hydrolysis (*without inhibition of* $F_1F_0$ *ATP synthesis*) very substantially decreases [ROS] (Figure 5). A drug that reverses, rather than just slows, damage only needs to be administered intermittently. For example, administered for a limited period each year.

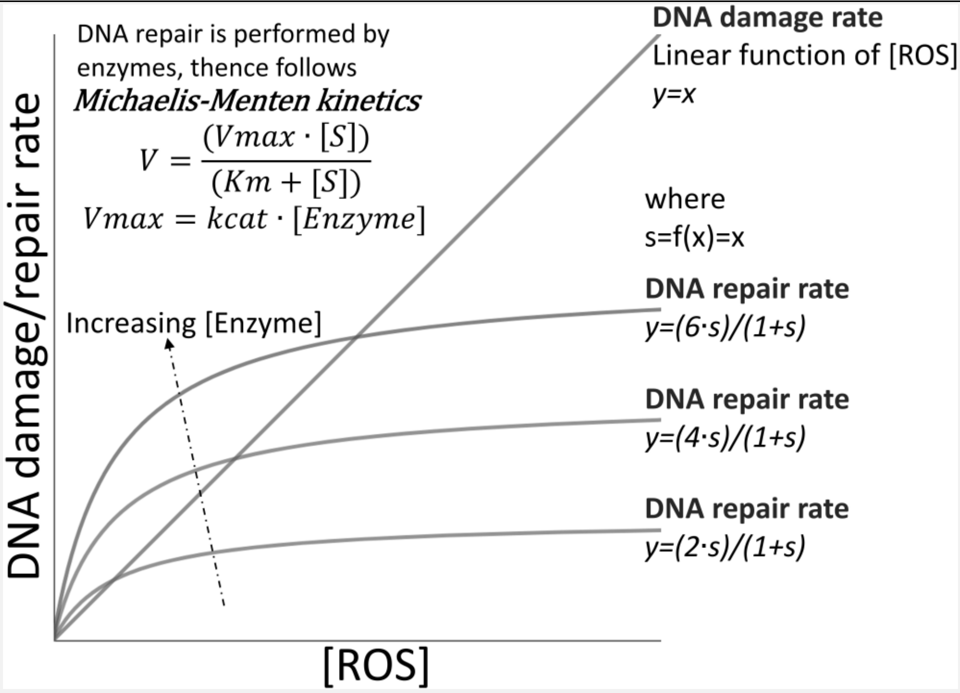

Figure S18 - Diagram: Damage is linear, whilst repair is a Michaelis-Menten, function of [ROS]. With less [ROS], there is less DNA damage *and* repair, but given their different gradients, there is a *net* reduction in DNA damage. If [ROS] falls further, DNA repair can exceed DNA damage, *if* there is excess DNA damage from a prior time when DNA damage exceeded DNA repair (*if* this DNA damage can still be recognized as damage by the DNA repair system), and *if* [Enzyme] is sufficient.

In the figure's definition of s, f(x) is approximated to x, such that s=x. But a more faithful simulacrum is provided in the next headed section. Where s=x+z, wherein x is [ROS] (=DNA damage it causes, because plot of DNA damage, y, is y=x) and z is any accumulated DNA damage due to the balance of past DNA damage and repair due to any past [ROS].

"Increasing [Enzyme]" in the figure, at least alone, is a constrained approach because (typically) when something is being repaired its use is prevented/impaired. A better approach is less [ROS], and so less ROS damage in the first place. In this DNA case, more repair would require more repair enzymes "sliding" (1D instead of 3D diffusion [573-574]) along the DNA looking for damage, further disrupting gene transcription. More repair requires more ATP energy, which can mean (via more oxidative phosphorylation) more ROS generation and damage, requiring more repair.

Conceivably, slight body temperature drop can assist repair/replacement vs. damage. Because repair/replacement is, and damage isn't, enzyme catalysed. Wherein enzymes follow macromolecular rate theory (MMRT), which can have (near their optimum temperature, $T_{opt}$) a shallower gradient of reaction rate decline with declining temperature [575-578]: e.g., refer to Figure 1B in [578].

If possible, leveraging the body's own breadth of repair/replacement mechanisms, as hypothesized above, is probably easier than us trying to execute repair/replacement, for all the many different types of molecular damage (that characterize/are aging), in parallel, *de novo*.

To theorize further, such aging reversal, by this mechanism, might be what happens naturally in a zygote (reprogramming). Indeed, Reactive Oxygen Species (ROS) might be a differentiation signal (e.g., [579-581]), and so much less ROS may be a de-differentiation signal, whilst simultaneously conferring aging reversal as per the above. Not such a large reduction in ROS may confer partial reprogramming, wherein there is aging reversal without de-differentiation.

<u>A better plot of DNA repair rate for substitution into Figure S18</u>

This section should be read with Figure S18 and its legend. It contributes a more faithful simulacrum of DNA repair rate, *y*. In the DNA repair rate equation below, the *s* variable is DNA damage concentration, [DNA damage], which is a function of not only present [ROS], *x*, shown on the x-axis of Figure S18, but *also* any past [ROS]. For example, where *t* is present time, (*t*-Δ*t*) is a very short time (*Δt*) before the present time (*t*), *t*=0 is time at the start of this DNA, with surrounding [ROS], system, *Δt* is the step-size in time to use in the calculation, wherein a smaller value of *Δt* confers more accuracy, $t_i$ is the latest interim time in the calculation of *z*, *x* is [ROS] (=DNA damage it causes, because plot of DNA damage, *y*, is y=x), wherein this equation system has a Michaelis-Menten form at various points within it, wherein its selected values are merely illustrative, and not restricting:

plot of DNA repair rate, *y*, wherein *z* is any accumulated DNA damage due to the balance of past DNA damage and repair due to past [ROS], in the past time period (t=0)→(t-Δt):

$$y = \left(\frac{2 \cdot s}{1 + s}\right)$$

$$s = f(x) = x + z$$

$$z = \begin{cases} \sum_{(t=0)\rightarrow(t-\Delta t)} \begin{cases} \left(x - \left(\frac{2 \cdot x}{1 + x}\right)\right) & if\ x > \left(\frac{2 \cdot x}{1 + x}\right) \\ 0 & if\ x = \left(\frac{2 \cdot x}{1 + x}\right) \\ 0 & \begin{cases} if\ x < \left(\frac{2 \cdot x}{1 + x}\right) \\ and\ if \left(\sum_{(t=0)\rightarrow t_i} x\right) \leq \left(\sum_{(t=0)\rightarrow t_i} \left(\frac{2 \cdot x}{1 + x}\right)\right) \end{cases} \\ \left(x - \left(\frac{2 \cdot x}{1 + x}\right)\right) = v & \begin{cases} if\ x < \left(\frac{2 \cdot x}{1 + x}\right) \\ and\ if \left(\sum_{(t=0)\rightarrow t_i} x\right) > \left(\sum_{(t=0)\rightarrow t_i} \left(\frac{2 \cdot x}{1 + x}\right)\right) \\ wherein\ used\ step\ size\ in\ time\ (t)\ is\ such: \\ |v| \leq \left(\left(\sum_{(t=0)\rightarrow t_i} x\right) - \left(\sum_{(t=0)\rightarrow t_i} \left(\frac{2 \cdot x}{1 + x}\right)\right)\right) \end{cases} \end{cases} \\ 0 & if\ (z < 0) \end{cases}$$

$$if \left(\left(\frac{2 \cdot s}{1 + s}\right) > x\right) \left\{\left[s = \frac{x + z}{q}\right], where\ q\ is\ set\ such\ that \left(y = \left(\frac{2 \cdot s}{1 + s}\right) = x\right)\right\}$$

[Eq. S24]

The last line is "catch code" ensuring that total DNA damage repair cannot exceed total DNA damage.

The above uses idiosyncratic sigma notation. Which I think is clearer. If more conventional notation is wanted, please substitute the sigma notation accordingly. For the first, most principal, sigma symbol, please substitute in:

$$\sum_{n=b}^{p}$$

Where *b* is an "arbitrarily small" number, *p* is $t$-$\Delta t$ and every instance of *x* falling under this sigma should instead be $x_n$. There are six daughter sigma systems (which too should each use $x_n$ instead of *x*), which can each use the same sigma notation:

$$\sum_{n=b}^{g}$$

Where *g* is $t_i$.

Comparison with "senolytics"

[Table S124]

| | **Less senescent cells** | **Slightly lower body temperature** |
|---|---|---|
| Mouse | Transgenic mice (males and females) with less senescent cells had 27% (*or 24% in a different mouse strain*) greater median lifespan. Published in Nature [582]. | Transgenic mice with 0.34°C lower mean body temperature had 20% greater median lifespan (*females. Males had 0.3°C lower, conferring 12% extension*). Published in Science [21]. |
| Human | **<u>No data</u>** before senolytic drugs entered clinical trials.<br>Since then, there has been the Phase II clinical failure of a senolytic drug (Unity Biotechnology's UBX0101 *<which is (according to its Phase II clinical study protocol) nutlin-3a in [583], which was developed by the pharmaceutical company Hoffmann-La Roche>* injected into knees with osteoarthritis) [584]. At the time of writing (in 2022), clinical trials ongoing for UBX1325, fisetin and for the Dasatinib+Quercetin combination [585]. | Humans with lower body temperature have a longer lifespan [168, 30, 181-182] and healthspan [183-184].<br>*Men with body temperature below the median have greater life expectancy (shown herein using data from [30]).* |
| Corresponding drug success in mice? | Drugs successfully shown to decrease the number of senescent cells.<br>*All be it only locally (because of systemic toxicity) for the first senolytic drugs that entered clinical trials, from Unity* | Drug successfully shown to dose-dependently lower body temperature (Figure 2).<br>*Drug administered systemically.*<br>Simple correspondence (*drug lowers body temperature of mice*) with the |

| | | |
|---|---|---|
| | *Biotechnology. Shown with senescent cells caused by injury rather than aging [196]. What distinguishes a senescent from a normal cell still isn't agreed [584].* | transgenic mice (*which have lower body temperature*). |
| Tractability of clinical trialling | At the time of writing, the FDA doesn't classify aging as a disease [586]. But it can grant drug approval for treatment of an age-related disease. Unity Biotechnology showed that UBX0101 could decrease senescent cell number after knee injury in mice (but this couldn't be replicated in rats or dogs) and extrapolated that UBX0101 would treat osteoarthritis [196]. Endpoint for an osteoarthritis clinical trial is typically patient self-reporting pain (e.g., using the WOMAC scoring scale questionnaire), for which the placebo effect, especially for knee injections of saline [587], can be hard to surpass. Unity Biotechnology also modelled aging with injury, in its pre-clinical testing in the eye, for its follow-up drug UBX1967 [196]. | Alzheimer's disease can cause a higher body temperature in humans [438]. Higher body temperature accelerates disease progression in a mouse model of Alzheimer's disease [439]. A clinical trial might use the objective endpoint of body temperature (as an endpoint itself. Or as a surrogate endpoint for the FDA's accelerated approval program [440]). Can this drug reduce the higher body temperature of Alzheimer's disease patients to normality? |

| | | |
|---|---|---|
| Corresponding drug extends mouse lifespan? | At the time of writing (2024), years after its IPO, none of Unity Biotechnology's drugs have been shown to extend lifespan in any organism. Extended lifespan has been shown with a different senolytic treatment, starting with already aged mice (quicker result), with systemic administration of fisetin [588] or a Dasatinib+Quercetin combination [589]. Although the Interventions Testing Program (ITP) [590] has tested fisetin (at the same dose as [588]) and it has no lifespan effect [279, 591]. | No such data to date. |
| Companies | Dozens [584]. Including Unity Biotechnology, which raised $222M in venture capital and then $85M in an IPO on NASDEQ, which valued the company at (pre-money) $700M (since had up to $80M of debt financing available). | None.<br>UPDATE, since October 2022: Biophysical Therapeutics, Inc. |
| Market size | At least some senolytic drugs, e.g. the failed and presently extant clinical drug candidates of Unity Biotechnology, are too toxic for systemic use, which restricts them to localized use [196]. | Systemic drug administration anticipated possible in humans. So, (conceivably) able to slow aging throughout the body, delaying/slowing/ameliorating/treating ***all*** age-related diseases (simultaneously) thereby. Economists |

| | | |
|---|---|---|
| | | calculate that a slowdown in aging, which increases US life expectancy by one year, is worth $38T [434]. And further years accorded multiple thereof. Local administration anticipated possible also: e.g., to the knee or eye. Potential market is everyone (all adults). Because everyone ages and no one wants to. |
| Potential | The result in transgenic mice, given that *many/most* senolytic cells were eliminated [582], is probably near or at the fullest lifespan extending potential of this approach. | There is reason herein to believe that the result in transgenic mice [21] is not at, or anywhere near, the full lifespan extending potential of this (reducing metabolic heat generation: i.e. reducing metabolic rate) approach. |
| Criticism(s) | At least some senescent cells (or type[s] thereof) might be useful, not least for performing the tissue function when they are an appreciable proportion of it. "Senolytics are likely to accelerate tissue pathology, exacerbate clinical disease, and result in early morbidity and mortality" [592]. "We found that removing senescent cells, however, may be worse than leaving them to accumulate, as this can have systemic and liver-debilitating consequences"; "we show here that there are abundant | To be seen. |

| | | |
|---|---|---|
| | p16[High] senescent cell types in an aging organism that are structurally and functionally important, while their removal has detrimental consequences"; "defined p16[High] senescent cell types are indispensable for mouse healthspan" [593]. What differentiates senescent from normal cells is contentious and so there isn't consensus about how they can be specifically and safely targeted ("we are far from being able to define something as senescence-specific") [584]. Studies often *assume* that only senescent cells express p16 [196], which might not be correct. There might be different types of senescent cells [584]. Killing senescent cells may cause them to release their contents, which are posited to be harmful, and so making things worse, not better. Senolytic drug action is often pre-clinically assayed against senescent cells caused by injury rather than aging [196] - but are these the same? | |
| Anticancer utility | At least some senolytic drugs are repurposed pre- | Selective $F_1F_0$ ATP hydrolysis inhibiting drugs (*that don't inhibit $F_1F_0$ ATP synthesis*) have anticancer activity. |

| | | |
|---|---|---|
| | clinical/clinical anticancer drugs. | This is a new finding. Patents filed (e.g. [18-20]) and so commercially viable to pursue. Clinical trials for cancer have well-defined, objective endpoints (e.g., delay of death), and surrogate endpoints (e.g., less tumour growth), that can be shown relatively quickly (especially in cancers that {tragically} tend to have poor prognosis, where patients only live months after diagnosis). There is a great unmet medical need, especially for a cancer drug without horrific side-effects (which often cause patients to rescind consent to treatment). Cancer cachexia (wasting/weight loss) is a huge unmet medical need. Relevantly, transgenic mice with slightly lower body temperature have higher body weight (with the same food consumption as control mice) [21]. Presumably because of greater metabolic efficiency, with less of the chemical energy of food dissipated as heat. Alzheimer's disease patients also suffer weight loss [541-543, 546-556], which their mortality is a function of [555-556]. Counteracting (e.g., slowing of) weight loss is a very objective, tractable clinical trial endpoint/surrogate endpoint. |

Caveat

For expediency of explanation, at some points in the paper I write as though all homeothermic species maintain a body temperature around 37°C. Although this is true for many homeothermic mammals (e.g., mice and humans [1]), it certainly isn't true for all. For example, the naked mole rat (*Heterocephalus glaber*) maintains a body temperature around 32°C instead [1].

#### SUPPLEMENTARY METHODS

<u>Synthesis of 6a and 6b</u> (*6b administered to mice in Figure 2; anti-cancer data in Figure 18*)

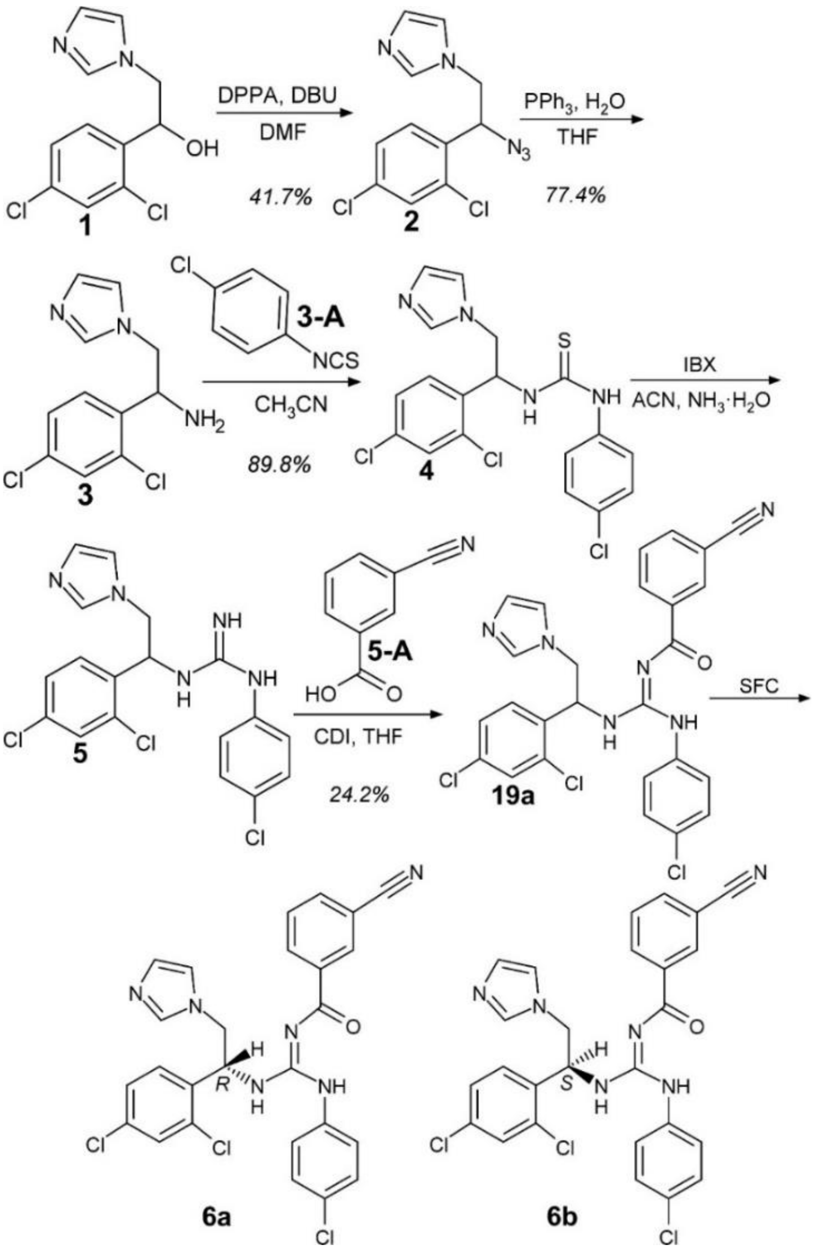

**6a** {enantiomeric excess (ee) ≥97%}: Liquid Chromatography-Mass Spectrometry (LC-MS): Liquid Chromatography (LC) Retention Time (RT) = 2.516 minutes, Mass spectrometry (MS; electrospray ionization, positive mode): *m/z* 537.1 $[M+H]^+$, 559.1 $[M+Na]^+$, 269.1 $[M+2H]^{2+}$.

$^1$H NMR (400 MHz, DMSO-*d6*) δ (ppm) 11.43 (s, 1H), 8.30 (s, 1H), 8.21 (d, J = 7.9 Hz, 1H), 7.93 (d, J = 7.8 Hz, 2H), 7.67 – 7.56 (m, 3H), 7.50 (t, J = 6.5 Hz, 4H), 7.20 (s, 1H), 7.09 (d, J = 8.2 Hz, 2H), 6.97 (s, 1H), 5.96 (s, 1H), 4.34 (s, 1H), 4.19 (s, 1H). {*NMR probe temperature = 298.15 K*}.

**6b** {enantiomeric excess (ee) ≥97%}: Liquid Chromatography-Mass Spectrometry (LC-MS): Liquid Chromatography (LC) Retention Time (RT) = 2.516 minutes, Mass Spectrometry (MS; electrospray ionization, positive mode): *m/z* 537.1 $[M+H]^+$, 559.1 $[M+Na]^+$, 269.1 $[M+2H]^{2+}$.
$^1$H NMR (400 MHz, DMSO-*d6*) δ (ppm) 11.43 (s, 1H), 8.30 (s, 1H), 8.21 (d, J = 7.9 Hz, 1H), 7.93 (d, J = 7.8 Hz, 2H), 7.68 – 7.56 (m, 3H), 7.56 – 7.46 (m, 4H), 7.20 (s, 1H), 7.09 (d, J = 8.1 Hz, 2H), 6.97 (s, 1H), 5.96 (s, 1H), 4.33 (s, 1H), 4.19 (s, 1H). {*NMR probe temperature = 298.15 K*}.

The presented Nuclear Magnetic Resonance (NMR) peaks are from using the "Auto Assignment" algorithm in MestReNova version 12 software (Mestrelab Research, Santiago de Compostela, Spain). Which picks peaks in an inputted NMR spectrum, influenced by an inputted structure, which was structure 19a in the reaction scheme above.

<u>Performed synthesis of 6a and 6b in detail</u>
In this section, numbers in square brackets are CAS numbers.

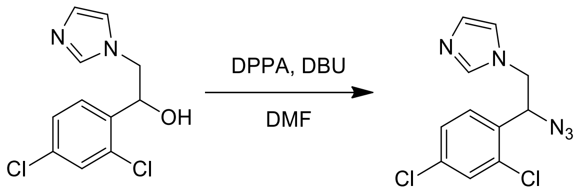

**1** **2**

1. Charged Compound 1 [24155-42-8] (10.0 g, 1.00 X by weight) and DPPA [26386-88-9] (10.7 g, 1.07 X by weight) in DMF [68-12-2] (50 mL, 5.00 X by volume).
2. Charged DBU [6674-22-2] (5.90 g, 0.59 X by weight) into the mixture, under nitrogen atmosphere at 0°C, then stirred for about 15 minutes.
3. After that, took the reaction mixture to 20-30°C and stirred for 24 hours.
4. Thin-Layer Chromatography (TLC; DCM/MeOH = 10/1, $R_f$ = 0.6) showed raw material remained, LC-MS (Starting Material [SM]: RT = 0.729 minutes, Product: RT = 0.822 minutes) indicated reaction mixture had desired product.

5. The reaction mixture was extracted with EtOAc (100 mL x 3).
6. The combined organic phase was washed with aq.NaCl (100 mL), dried with $Na_2SO_4$, filtered and concentrated in vacuum to get the residue.
7. The residue was purified by column chromatography (DCM/MeOH = 1/0 to 100/1).
8. Compound 2 (4.82 g, 41.7%, 95% purity) was obtained as a yellow oil, which was confirmed by $^{H}$NMR and HPLC (Retention Time (RT) = 1.919 minutes).
Compound 2: $^1$H NMR ($CDCl_3$, 400 MHz) δ (ppm) 7.47 (d, J = 2.2 Hz, 1H), 7.41 (s, 1H), 7.33 - 7.28 (m, 1H), 7.26 - 7.23 (m, 1H), 7.06 (s, 1H), 6.92 (s, 1H), 5.26 (m,1H), 4.23 (m, 1H), 4.02 (m, 1H).

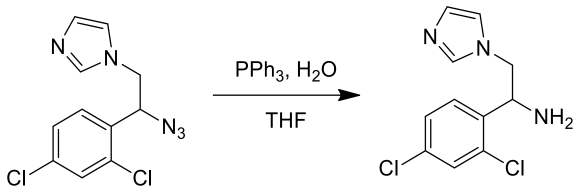

2 3

1. Charged Compound 2 (3.00 g, 1.00 X by weight) and $PPh_3$ [603-35-0] (8.40 g, 2.80 X by weight) in $H_2O$ (1 mL, 0.33 X by volume) and THF [109-99-9] (10 mL, 3.33 X by volume).
2. Stirred at 70°C for 24 hours.
3. Checked the reaction by LC-MS (SM: RT = 0.822 minutes, Product: RT = 0.299 minutes), it showed that Compound 2 was consumed completely, and that the desired mass was detected.
4. The reaction mixture was concentrated under reduced pressure to remove THF.
5. The reaction mixture was added with $H_2O$ (30 mL), adjusted pH to 3 with con.HCl and washed with EtOAc (30 mL).
6. After that, collected water phase, adjusted pH to 13 with 1N aq.NaOH, then exacted with EtOAc (30 mL x 2).
7. The combined organic phase was washed with aq.NaCl (30 mL), dried with $Na_2SO_4$, filtered, and concentrated in vacuum to obtain the product.
8. Compound 3 (2.23 g, 77.4%, 94.5% purity) was obtained as a yellow liquid, which was confirmed by $^{H}$NMR and HPLC (Retention Time = 0.658 minutes).
Compound 3: $^1$H NMR ($CDCl_3$, 400 MHz) δ (ppm) 7.48 - 7.37 (m, 3H), 7.30 - 7.26 (m, 1H), 7.06 (s, 1H), 6.91 (s, 1H), 4.72 (m, 1H), 4.21 (m, 1H), 3.91 (m, 1H).

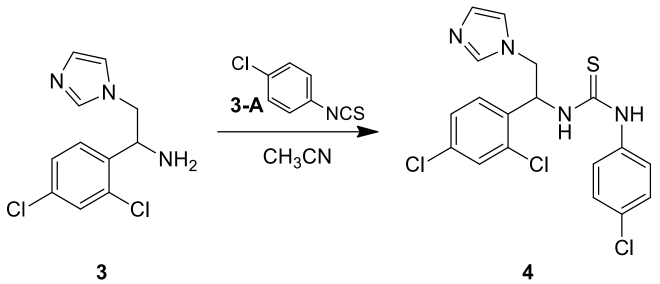

3 4

1. Charged Compound 3 (2.00 g, 1.00 X by weight) and Compound 3-A [2131-55-7] (1.30 g, 0.65 X by weight) in $CH_3CN$ [75-05-8] (30 mL, 15.00 X by volume).
2. Stirred at 20-30°C for 12 hours.
3. Checked the reaction by TLC (DCM/MeOH = 10/1, $R_f$ = 0.7) and LC-MS (SM: RT = 0.299 minutes, Product: RT = 0.949 minutes), desired product was detected, TLC indicated Compound 3 was consumed completely.
4. The reaction mixture was concentrated under reduced pressure to remove $CH_3CN$ to obtain the residue.
5. The residue was purified by column chromatography (DCM/MeOH = 1/0 to 50/1).
6. Compound 4 (2.95 g, 89.8%, 99.7% purity) was obtained as a white solid, which was confirmed by $^{H}$NMR and HPLC (Retention Time = 2.521 minutes).
Compound 4: $^{1}$H NMR (DMSO-*d6*, 400 MHz) δ (ppm) 9.81 (s, 1H), 8.54 (d, J = 7.5 Hz, 1H), 7.64 (s, 1H), 7.55 (s, 1H), 7.53 - 7.48 (m, 1H), 7.45 - 7.41 (m, 1H), 7.38 (d, J = 8.6 Hz, 2H), 7.28 (d, J=8.4 Hz, 2H), 7.11 (s, 1H), 6.94 (s, 1H), 6.03 (s, 1H), 4.46 - 4.34 (m, 1H), 4.31 - 4.22 (m, 1H).

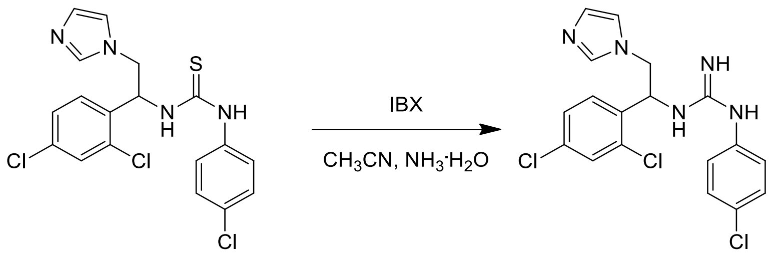

4 5

1. Charged Compound 4 (1.63 g, 1.00 X by weight) and IBX [61717-82-6] (1.18 g, 0.75 X by weight) in $CH_3CN$ [75-05-8] (16 mL, 10.00 X by volume) and $NH_3.H_2O$ [1336-21-6] (8 mL, 5.00 X by volume).
2. Stirred at 20-30°C for 18 hours.

3. Checked the reaction by LC-MS (SM: RT = 0.949 minutes, Product: RT = 1.050 minutes) and HPLC (SM: RT = 2.535 minutes, Product: RT = 1.757 minutes), desired product was detected.

4. The reaction mixture was extracted with EtOAc (20 mL x 3).

5. The combined organic phase was washed with aq.NaCl (20 mL), dried with $Na_2SO_4$, filtered and concentrated *in vacuo* to obtain the crude product (1.0 g, white solid), which was used into next step directly.

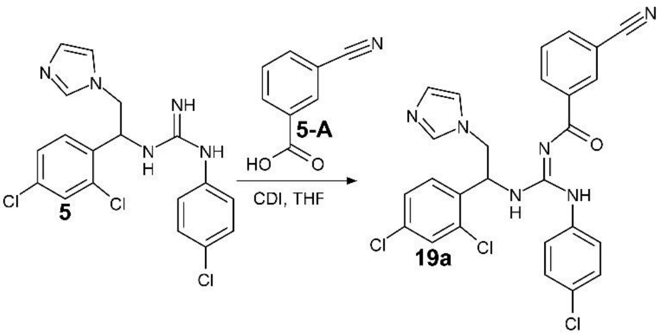

1. Charged Compound 5-A [1877-72-1] (0.36 g, 0.36 X by weight) and CDI [530-62-1] (0.40 g, 0.36 X by weight) in THF [109-99-9] (20 mL, 20.00 X by volume), stirred at 20-30°C for 3 hours.

2. Charged Compound 5 into the reaction mixture, stirred at 20-30°C for 14 hours.

3. Checked the reaction by LC-MS (SM: RT = 1.050 minutes, Product: RT = 1.347 minutes) and TLC (DCM/MeOH = 10/1, $R_f$ = 0.75), desired mass was detected. TLC indicated the raw material was consumed completely.

4. The reaction mixture was concentrated under reduced pressure to remove THF.

5. The reaction mixture was extracted with EtOAc (20 mL x 3)

6. The combined organic phase was washed with aq.NaCl (20 mL), dried with $Na_2SO_4$, filtered and concentrated *in vacuo* to obtain the residue.

7. The residue was purified by column chromatography (DCM/MeOH = 1/0 to 100/1) and chiral SFC {column: AD (250 mm*50 mm, 10 µm); mobile phase: [0.1%$NH_3H_2O$ IPA]; B%: 28%-28%, 3.5 minutes} separating the stereoisomers of Compound 19a: 6b (0.17 g, SFC RT = 2.35 minutes) was obtained as a white solid, 6a (0.15 g, SFC RT = 2.69 minutes) was obtained as a white solid.

*Abbreviations used: Ph=phenyl, MeOH=methanol, CDI=1,1'-carbonyldiimidazole, DCM=dichloromethane, DBU=1,8-diazabicyclo[5,4,0]undec-7-ene, DMF=N,N-dimethylformamide, DPPA=diphenylphosphoryl azide, EtOAc=ethyl acetate, IBX=2-*

*iodoxybenzoic acid, IPA=isopropanol, $R_f$=retention value, RT=retention time, THF=tetrahydrofuran, TLC=thin-layer chromatography, SM=starting material, SFC=chiral supercritical fluid chromatography, LC-MS=Liquid Chromatography Mass Spectrometry, HPLC=High-performance liquid chromatography, NMR=Nuclear magnetic resonance, aq.=aqueous, con.=concentrated, MeCN=methyl cyanide (acetonitrile), DMSO=dimethyl sulfoxide.*

Spectral data for 6b

Its expected/calculated *m/z*, from its structure (Figure 2), using the "exact molecular weight" calculator in MarvinSketch software (Chemaxon, Budapest, Hungary), which uses the mass of the *most prevalent* isotope of each element in the molecule:
537.08 $[M+H]^+$, 559.06 $[M+Na]^+$, 269.04 $[M+2H]^{2+}$, 575.03 $[M+K]^+$).

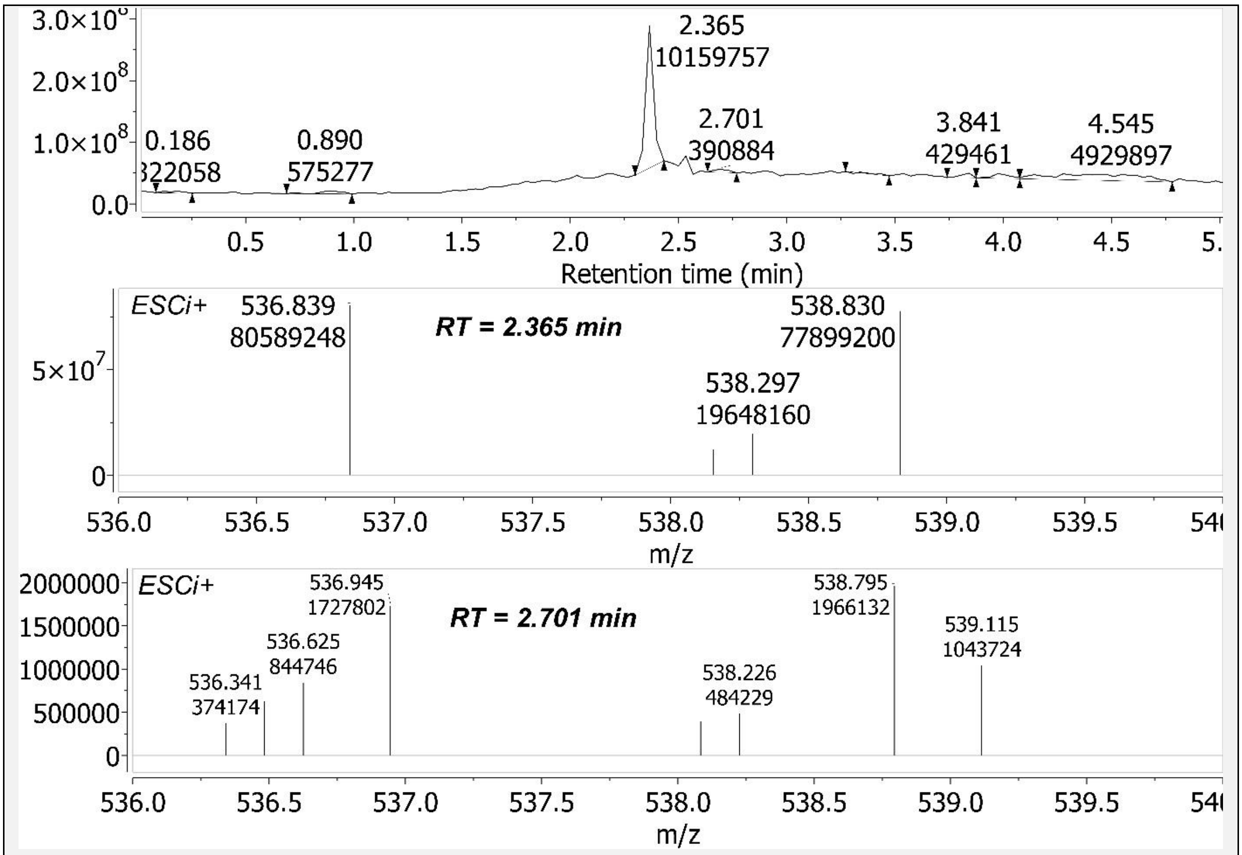

Figure S19 - Chiral Supercritical Fluid Chromotography Mass Spectrum (chiral SFC-MS) of 6b sample. *Upper panel*: Chiral-SFC annotated with Retention Time (RT, minutes), and area under peak. *S* stereoisomer is found in the [RT=2.365] peak. *Middle panel*: Component of the Mass Spectrum (MS) of the [RT=2.365] peak. Therein is *m/z* 536.839 $[M+H]^+$ (*0.19 removed from Expected. Better concordance in subsequent spectra*). Y-axis presents intensity. ESCi™

is multi-mode ionization (PCT/US03/16892), which switches at very high speed between Electrospray Ionization (ESI) and Atmospheric Pressure Chemical Ionization (APCI) ionization methods. ESCi+ refers to using ESCi in its positive ionization mode. Polarity positive. *Bottom panel*: Component of the Mass Spectrum (MS) of the [RT=2.701] peak.

Using this present figure, *S* stereoisomer excess (%) in this sample = (80589248/(80589248+1727802))*100 = 97.9%; enantiomeric excess (ee, %) = (97.9-50)*2 = 95.8%. Calculating the ee directly, rather than via the stereoisomer excess, ee = ((80589248-1727802)/(80589248+1727802))*100 = 95.8%. Working back from this ee to the stereoisomer excess: (95.8/2)+50 = 97.9%.

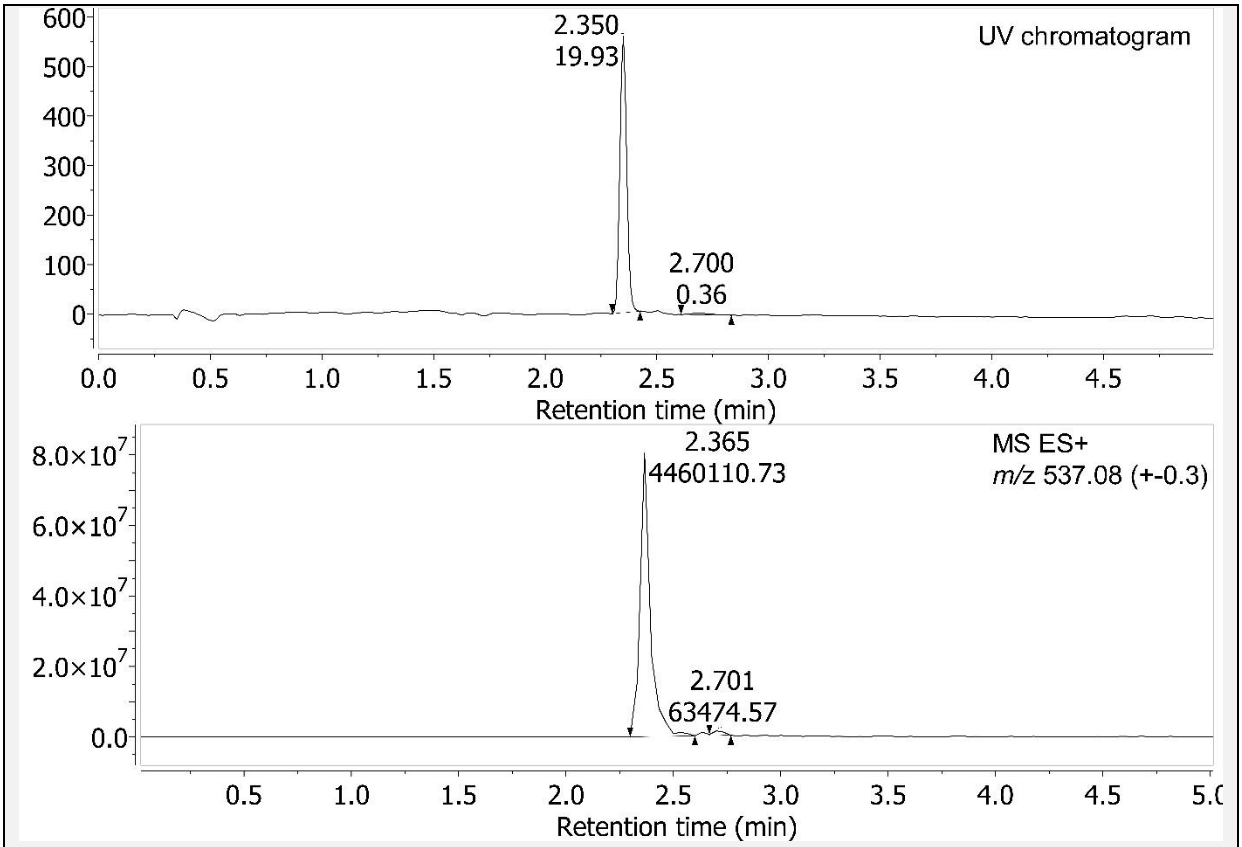

Figure S20 – Chiral Supercritical Fluid Chromotography (chiral-SFC) of 6b sample. Using the UV Chromatogram component, *S* stereoisomer excess (%) in this 6b sample = (19.93/(19.93+0.36))*100 = 98.23%; enantiomeric excess (ee) = (98.23-50)*2 = 96.46%.

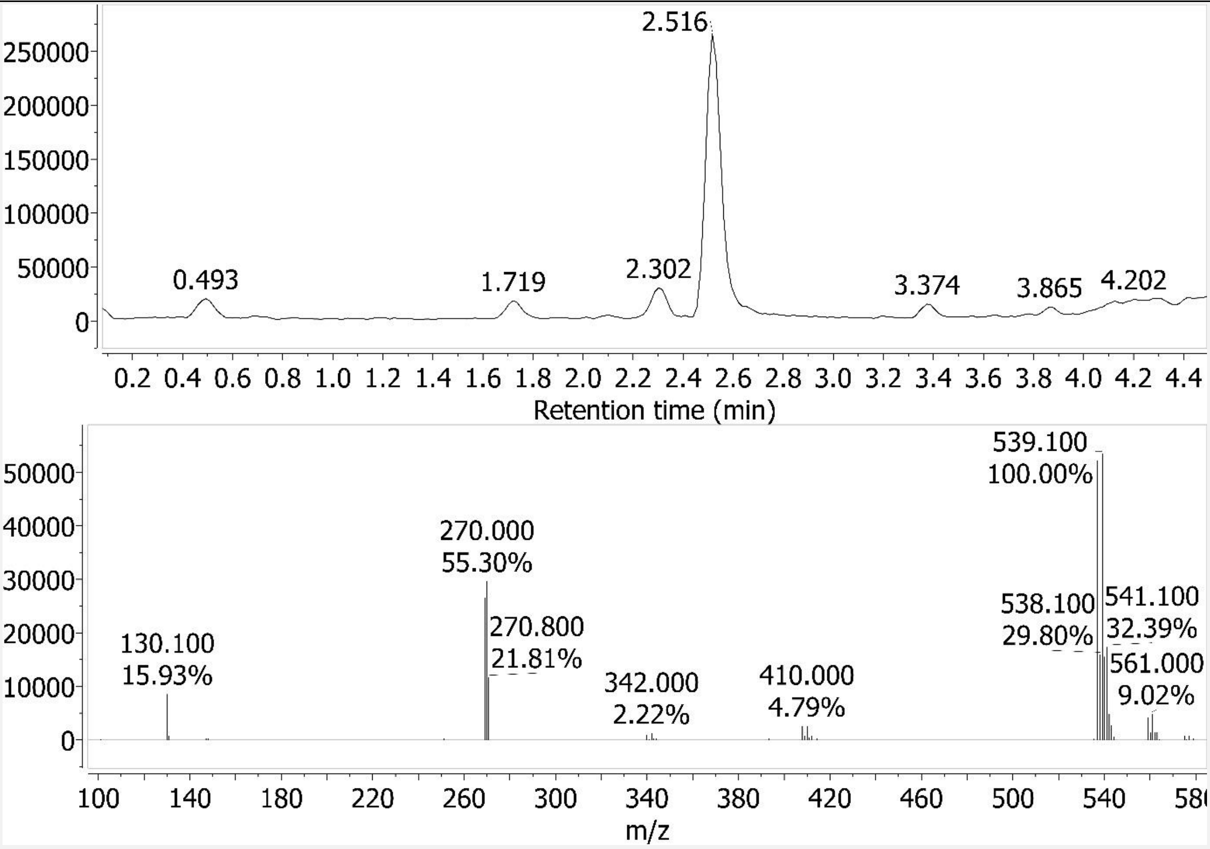

Figure S21 - Liquid Chromatography–Mass Spectrometry (LC-MS) spectrum for 6b sample. *Upper panel*: Liquid Chromatography (LC) Retention Time (RT) = 2.516 minutes. *Lower panel*: Mass Spectrum (MS) of the [RT peak=2.516]. Shows positively charged species: "Ionization mode = API-ES" and "Polarity = Positive". Wherein API is "Atmospheric Pressure Ionization". And ES is Electrospray. Percentages are relative abundance.

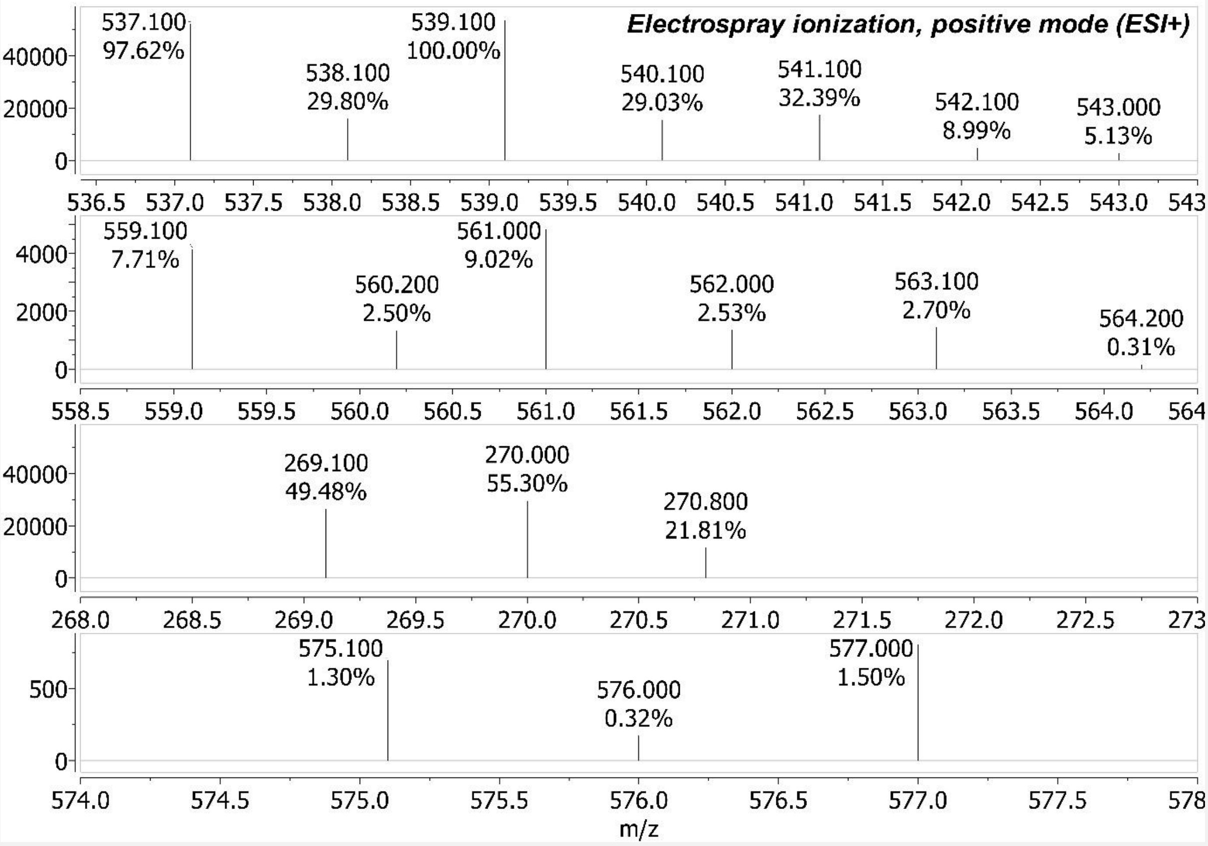

Figure S22 – Some Mass Spectrum (MS) components of Figure S21. *Upper panel*: *m/z* 537.1 $[M+H]^+$. *2nd panel*: *m/z* 559.1 $[M+Na]^+$. *3rd panel*: *m/z* 269.1 $[M+2H]^{2+}$. *Bottom panel*: *m/z* 575.1 $[M+K]^+$. All these Observed *m/z* are within 0.07 of Expected.

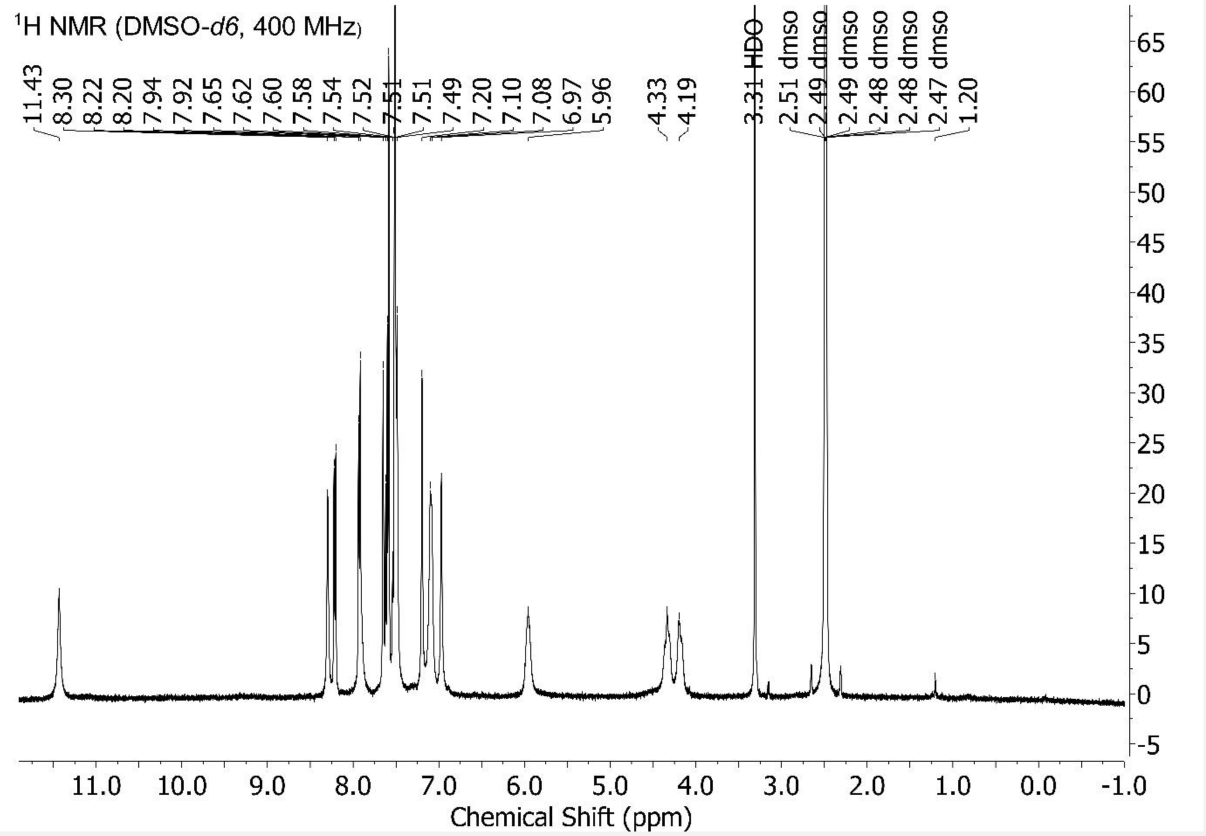

Figure S23 - $^{1}$H NMR {*NMR probe temperature = 298.15 K*} for 6b sample. Scaled to the highest compound peak. Peaks identified by the "Auto peak picking" algorithm (based on Global Spectral Deconvolution, GSD) of MestReNova version 12 software. With default program settings used. And no chemical structure used/inputted to guide/constrain this peak picking.

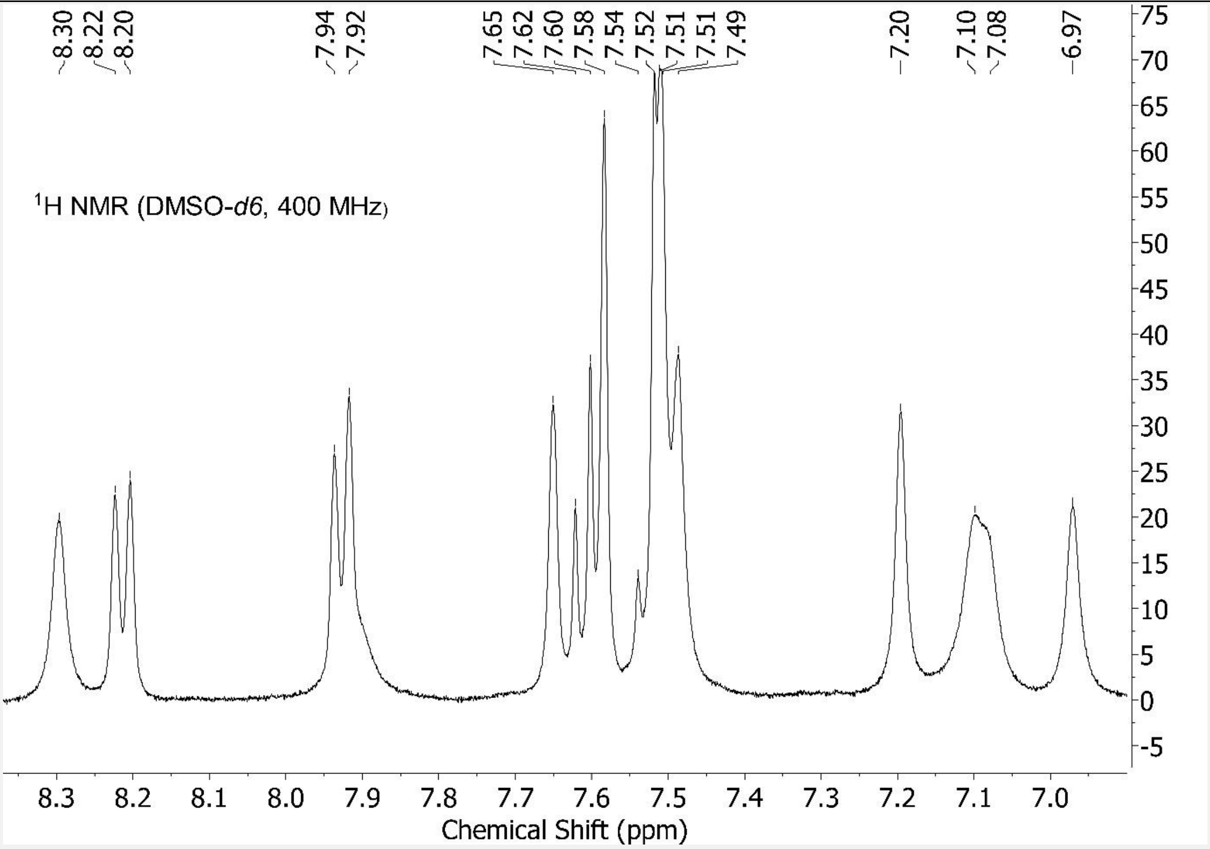

Figure S24 - Aromatic region of Figure S23 expanded.

Spectral data for 6a

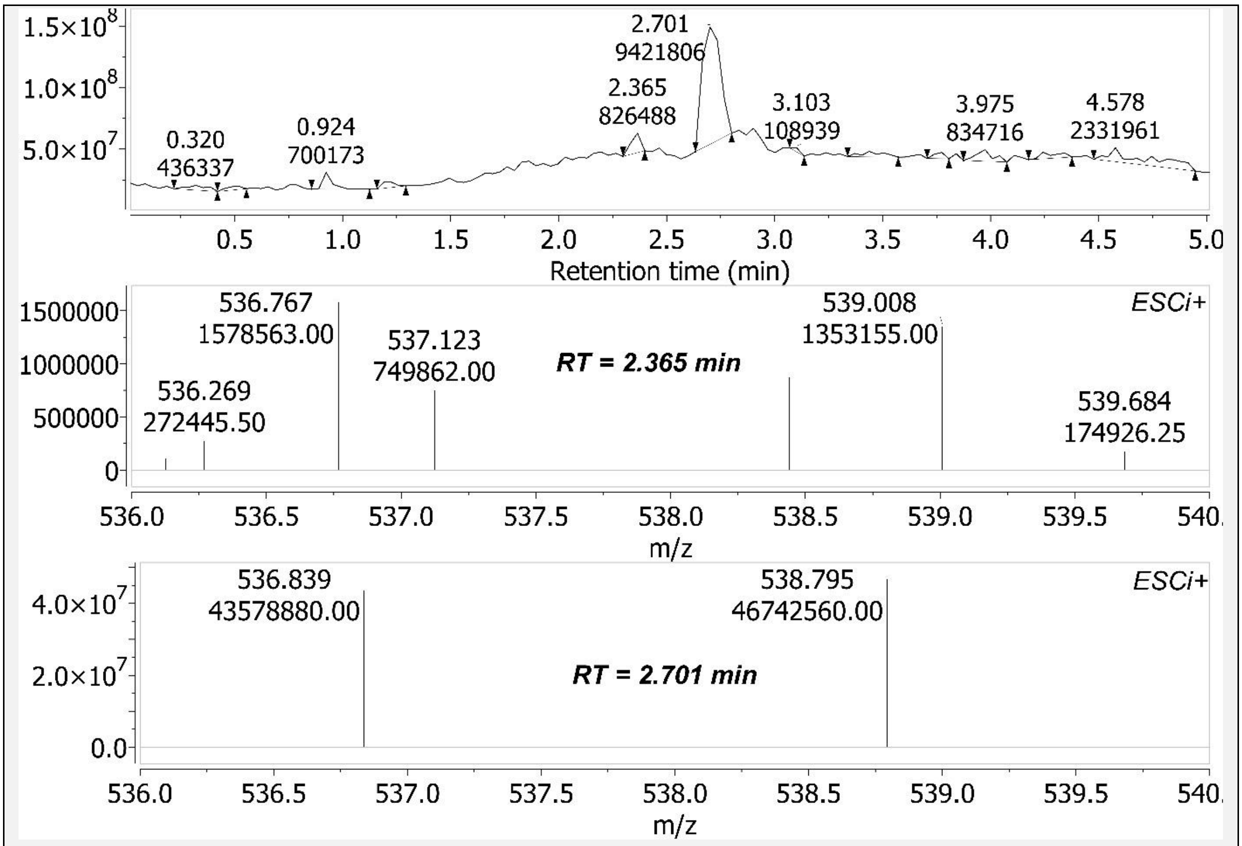

Figure S25 - Chiral SFC-MS of 6a sample. *Upper panel*: Chiral-SFC annotated with Retention Time (RT, minutes), and area under peak. *R* stereoisomer is found in the [RT=2.701] peak. *Middle panel*: Component of the Mass Spectrum (MS) of the [RT=2.365] peak. *Bottom panel*: Component of the Mass Spectrum (MS) of the [RT=2.701] peak. Therein is *m/z* 536.839 $[M+H]^+$ (*0.19 removed from Expected. Better concordance in subsequent spectra*). Using this present figure, *R* stereoisomer excess (%) in this sample = (435788880/(435788880+1578563))*100 = 99.64%; enantiomeric excess (ee) = (99.64-50)*2 = 99.28%.

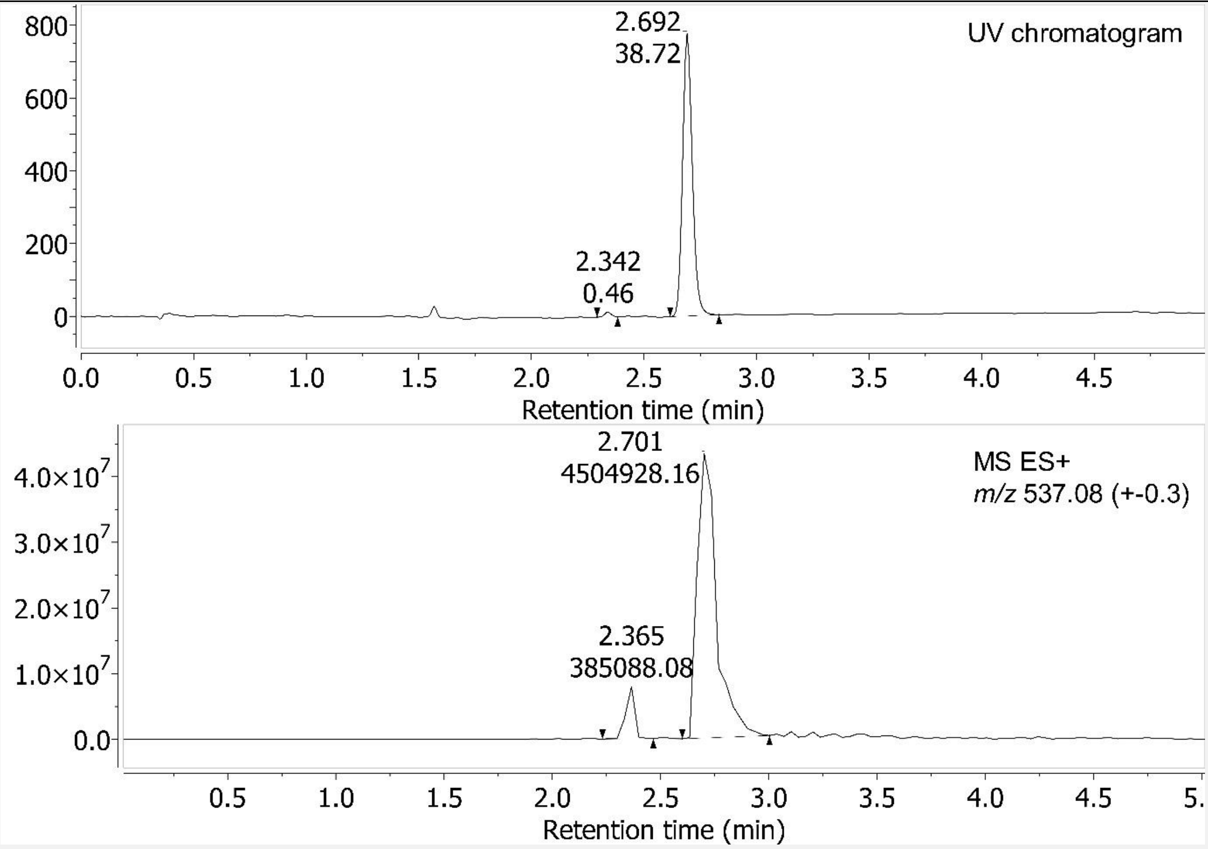

Figure S26 – Chiral-SFC of 6a sample. Using the UV Chromatogram component, *R* stereoisomer excess (%) in this 6a sample = (38.72/(38.72+0.46))*100 = 98.83%; enantiomeric excess (ee) = (98.83-50)*2 = 97.66%.

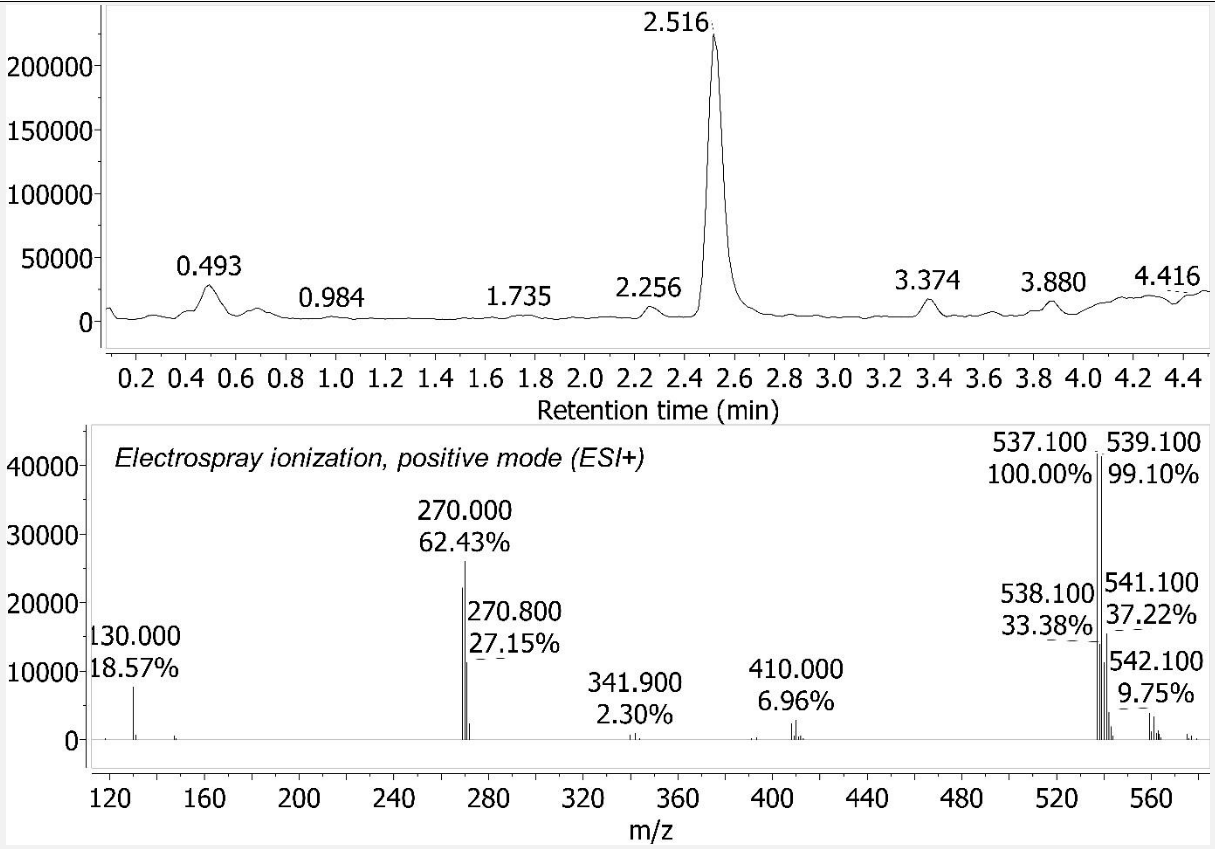

Figure S27 – LC-MS spectrum for 6a sample. *Upper panel*: Liquid Chromatography (LC) Retention Time (RT) = 2.516 minutes. *Lower panel*: Mass Spectrum (MS) of the [RT peak=2.516].

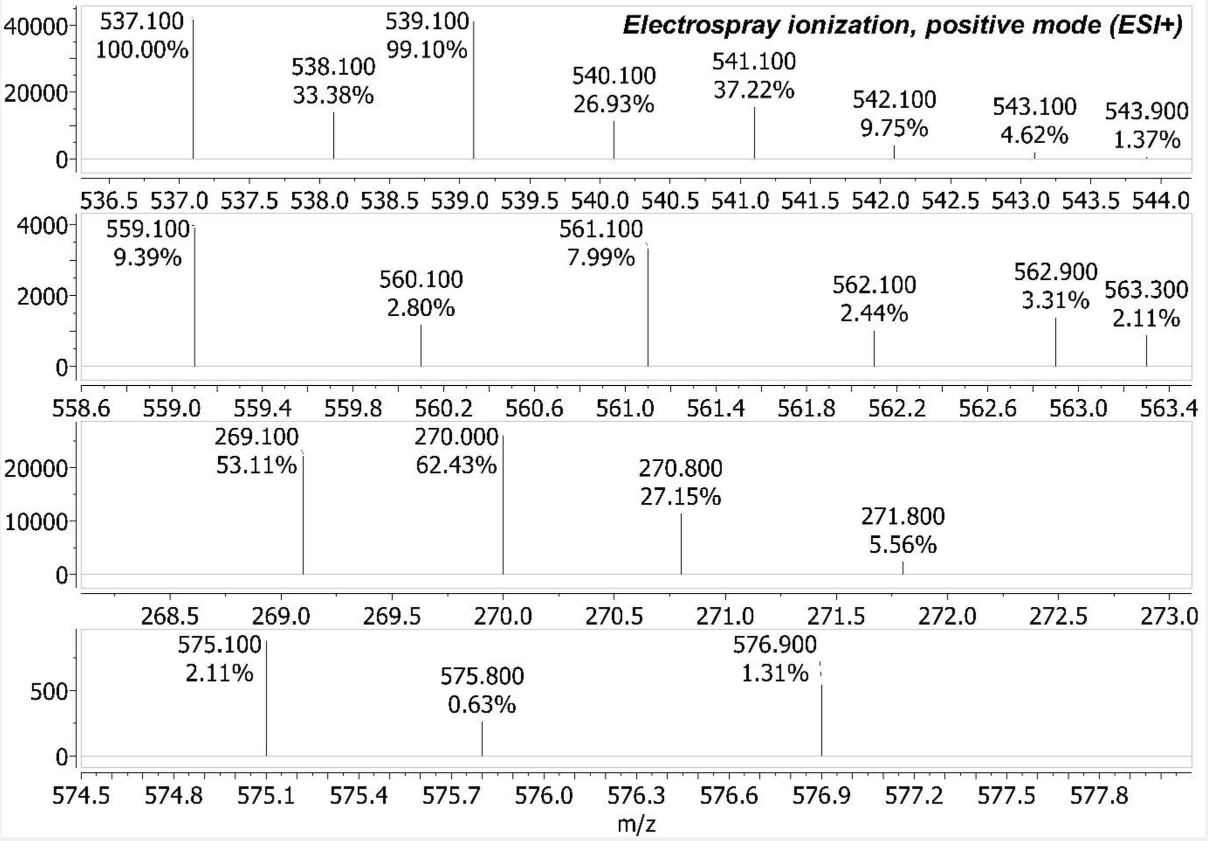

Figure S28 – Some Mass Spectrum (MS) components of Figure S27. Upper panel: *m/z* 537.1 $[M+H]^+$. 2nd panel: *m/z* 559.1 $[M+Na]^+$. 3rd panel: *m/z* 269.1 $[M+2H]^{2+}$. Bottom panel: *m/z* 575.1 $[M+K]^+$. All these Observed *m/z* are within 0.07 of Expected.

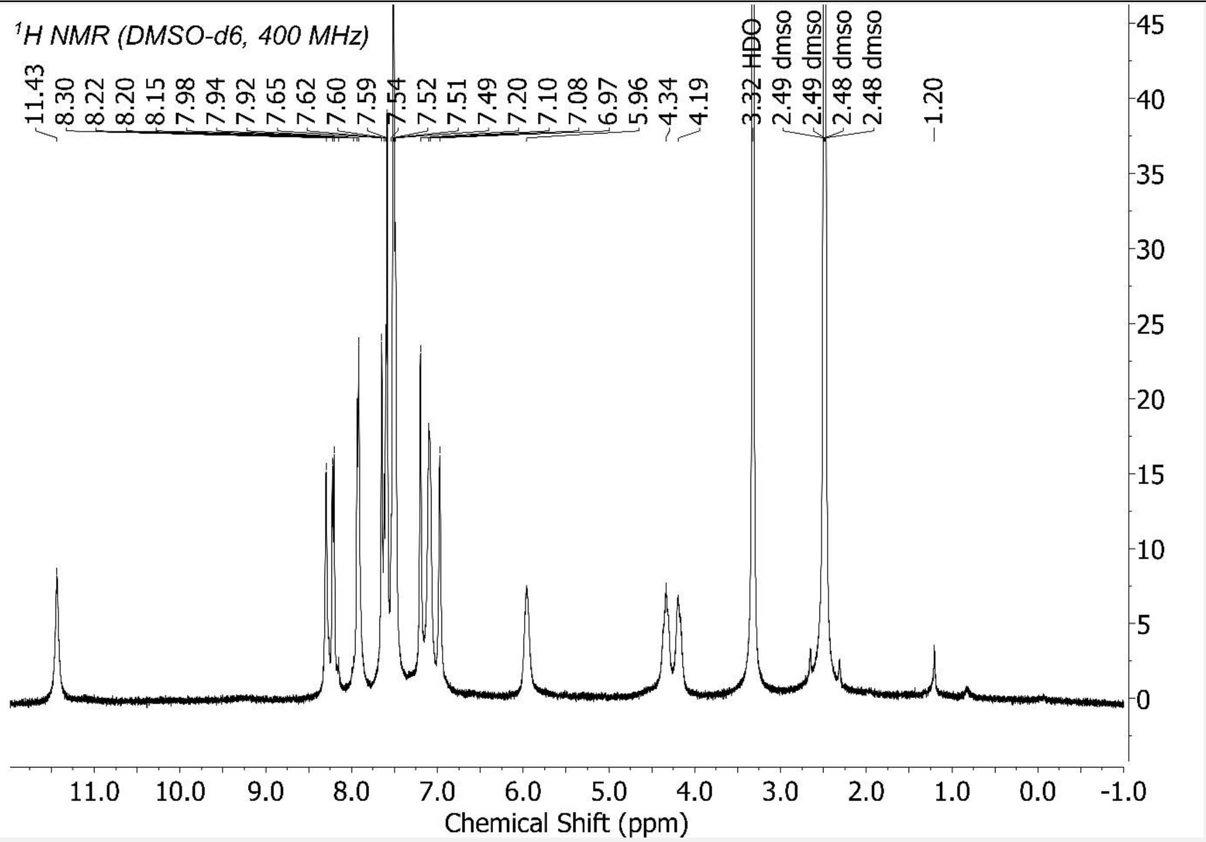

Figure S29 – $^1$H NMR {*NMR probe temperature = 298.15 K*} for 6a sample. Scaled to the highest compound peak. Peaks identified by the "Auto peak picking" algorithm (based on Global Spectral Deconvolution, GSD) of MestReNova version 12 software. With default program settings used. And no chemical structure used/inputted to guide/constrain this peak picking.

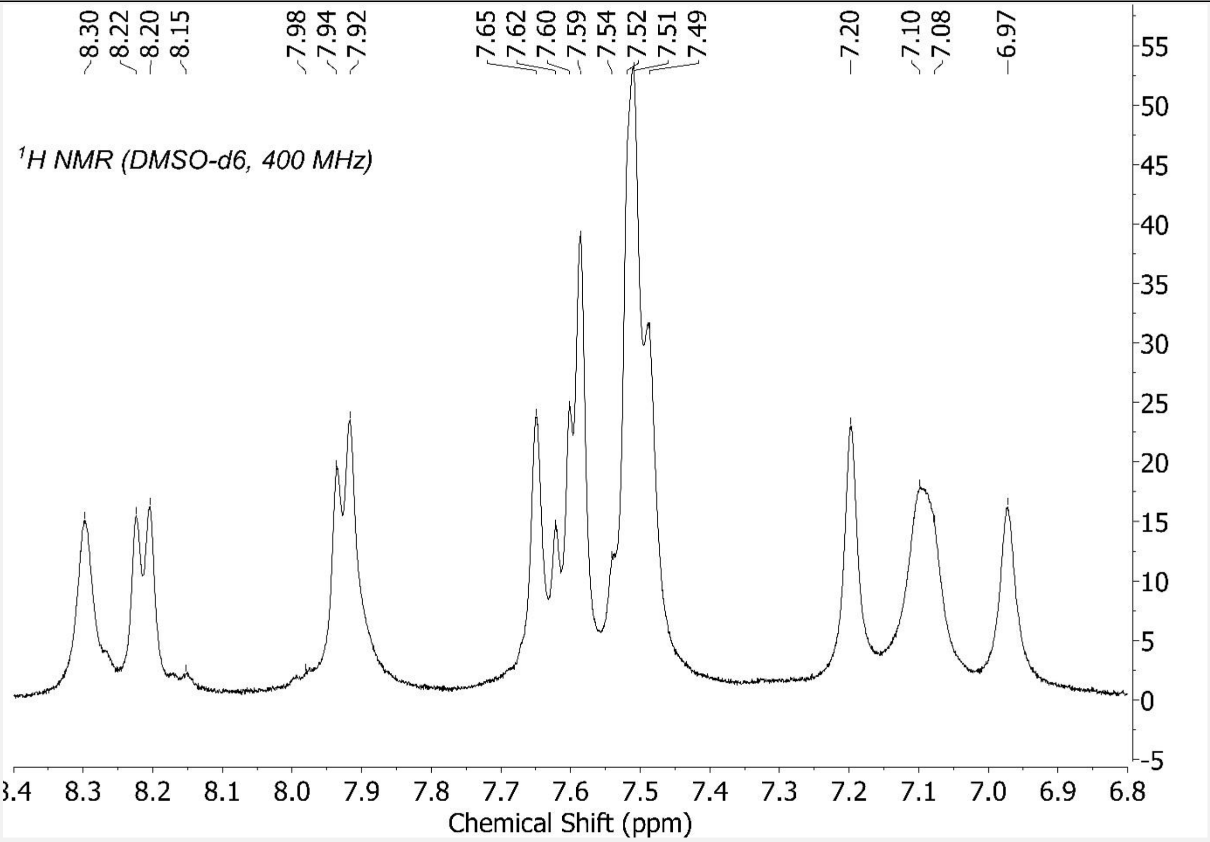

Figure S30 - Aromatic region of Figure S29 expanded.

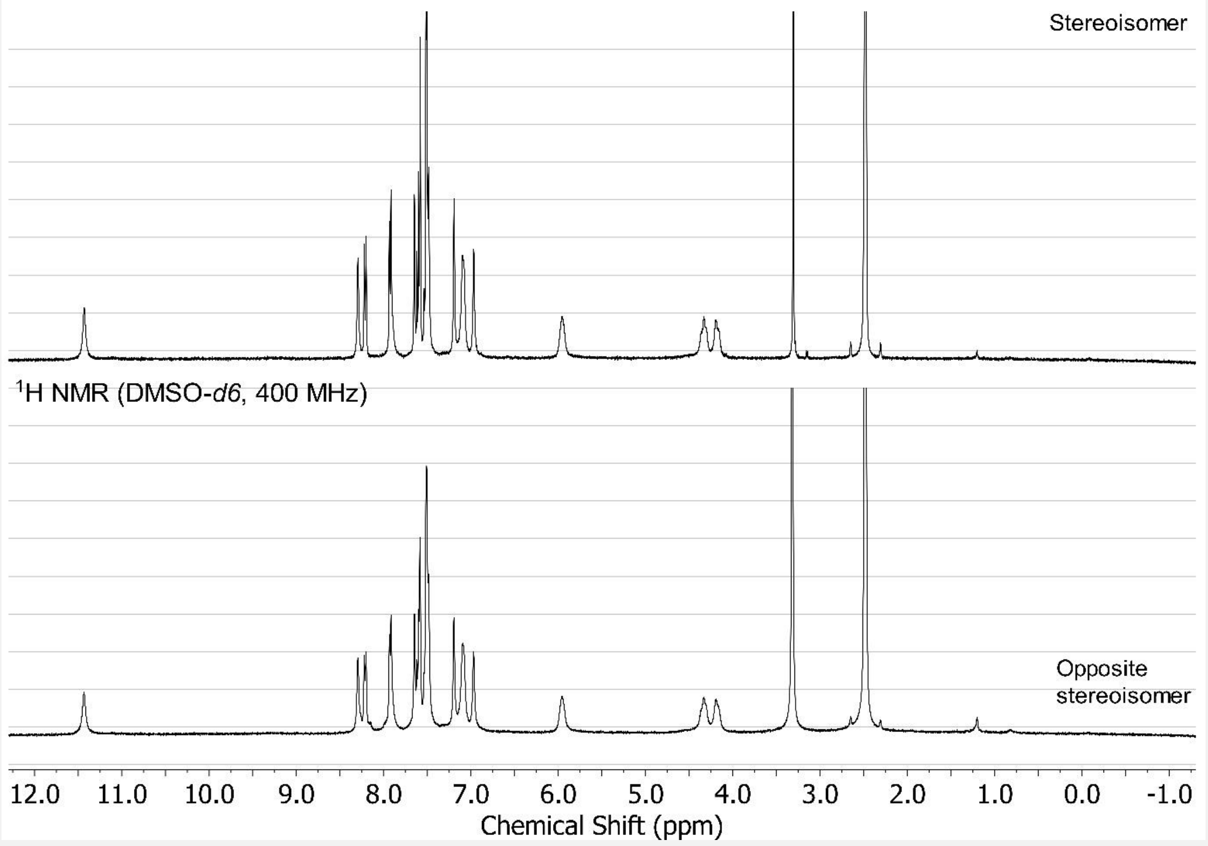

Figure S31 - 6b data in upper panel, 6a data in lower panel.

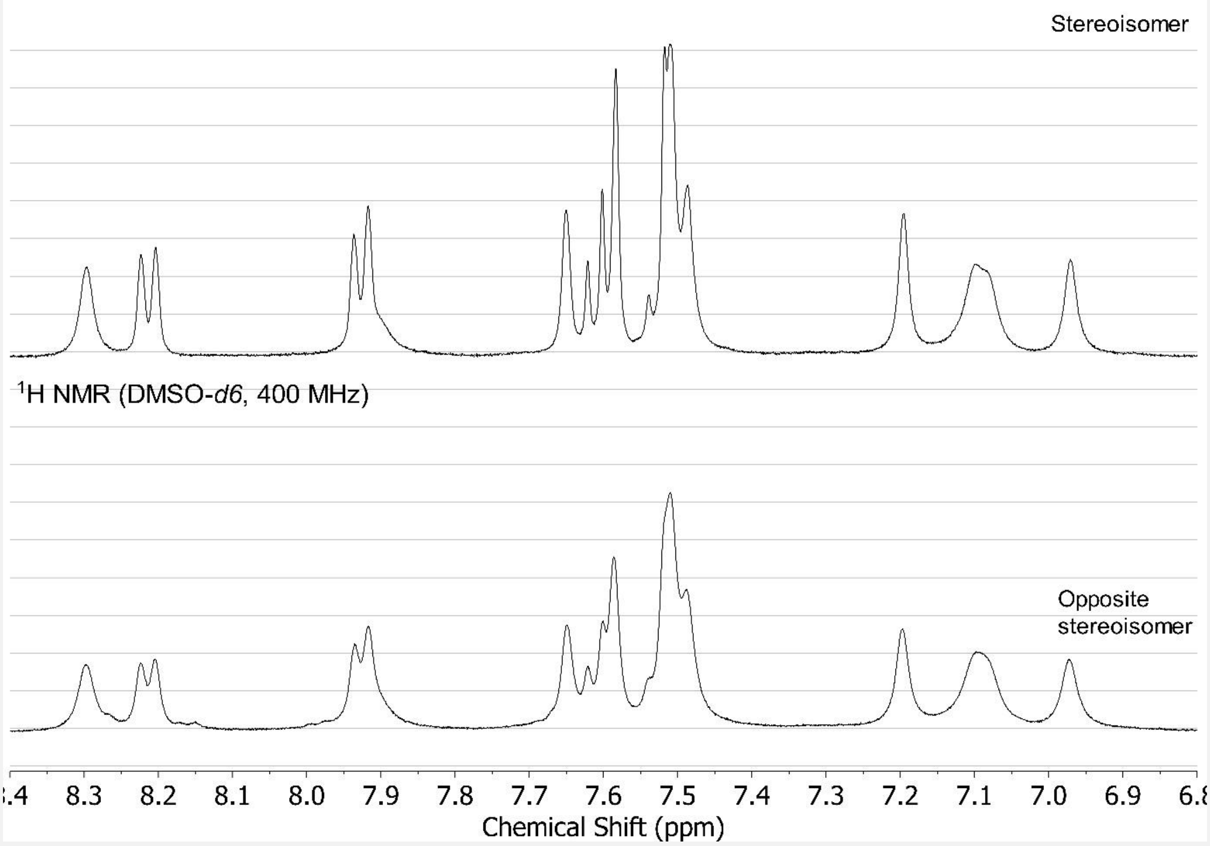

Figure S32 - Aromatic region: 6b data in upper panel, 6a data in lower panel.

Synthesis of 7a and 7b (*anti-cancer activity in Figure 19*)

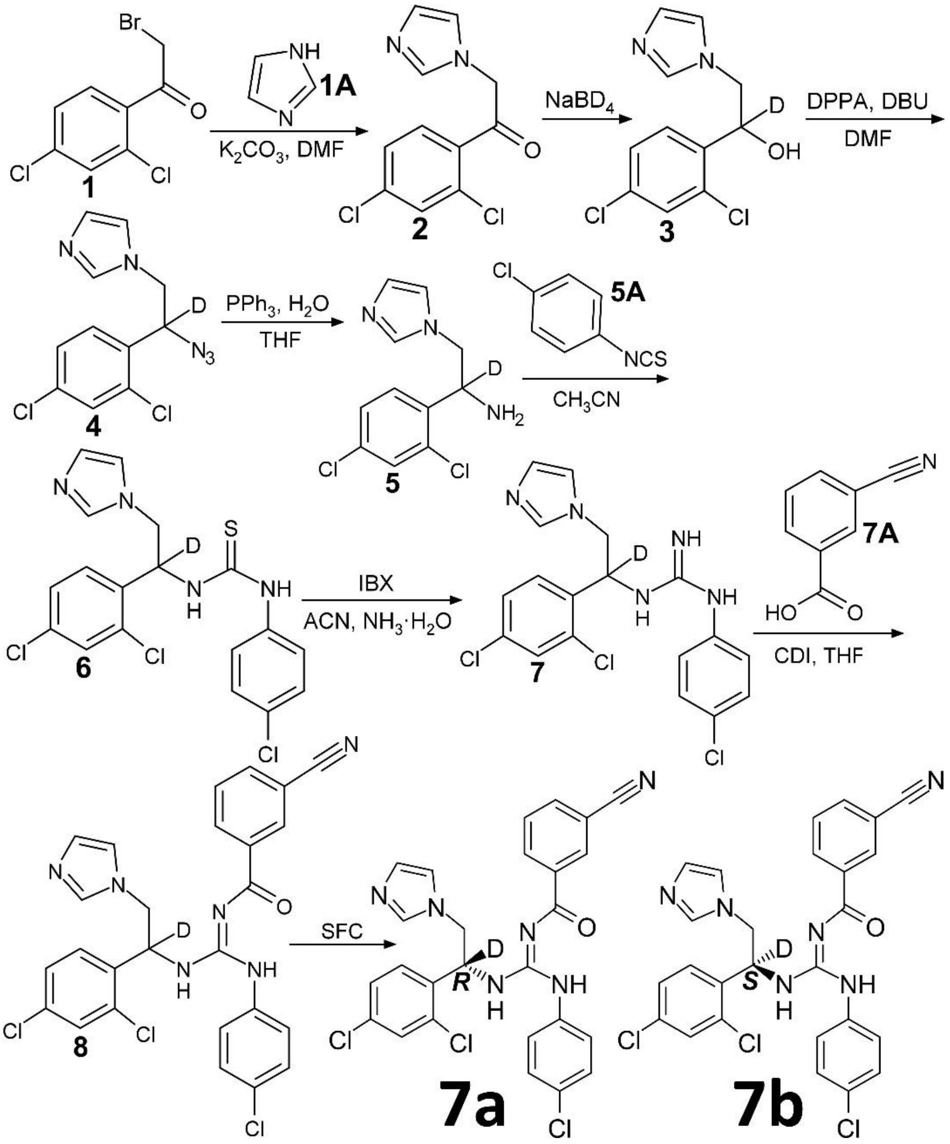

**7a** (enantiomeric excess (ee) ≥97%): Liquid Chromatography-Mass Spectrometry (LC-MS): LC Retention Time (RT) = 2.685 minutes, MS (electrospray ionization, positive mode): *m/z* 538.1 $[M+H]^+$, 560.1 $[M+Na]^+$, 269.6 $[M+2H]^{2+}$. High Resolution Mass Spectrometry (HRMS): Liquid Chromatography-Time of Flight (LC-TOF) MS (electrospray ionization, positive mode): LC Retention Time (RT) = 0.163 minutes, *m/z* 538.0727757864 $[M+H]^+$, *m/z* 560.0513502753 $[M+Na]^+$, *m/z* 576.0327248583 $[M+K]^+$, molar percent deuterium incorporation at chiral carbon = 99.14%.

$^1H$ NMR (400 MHz, DMSO-*d6*) δ 11.46 (s, 1H), 8.32 (s, 1H), 8.24 (d, J = 7.9 Hz, 1H), 7.95 (d, J = 7.7 Hz, 2H), 7.70 – 7.59 (m, 3H), 7.59 – 7.49 (m, 4H), 7.21 (s, 1H), 7.12 (d, J = 7.9 Hz, 2H), 6.99 (s, 1H), 4.36 (d, J = 13.9 Hz, 1H), 4.22 (s, 1H). {*NMR probe temperature = 300.7 K*}.

**7b** (enantiomeric excess (ee) ≥97%): LC-MS: LC Retention Time (RT) = 2.685 minutes, MS (electrospray ionization, positive mode): *m/z* 538.1 $[M+H]^+$, 560.1 $[M+Na]^+$, 269.6 $[M+2H]^{2+}$. HRMS: LC-TOF MS (electrospray ionization, positive mode): LC RT = 0.166 minutes, *m/z* 538.0745928061 $[M+H]^+$, *m/z* 560.0600137508 $[M+Na]^+$, *m/z* 576.0250917093 $[M+K]^+$, molar percent deuterium incorporation at chiral carbon = 99.13%.
$^1$H NMR (400 MHz, DMSO-*d6*) δ 11.48 (s, 1H), 8.33 (s, 1H), 8.25 (dt, J = 7.8, 1.5 Hz, 1H), 7.96 (dt, J = 7.7, 1.5 Hz, 1H), 7.92 (s, 1H), 7.71 – 7.60 (m, 3H), 7.60 – 7.49 (m, 4H), 7.23 (s, 1H), 7.14 (s, 2H), 7.00 (s, 1H), 4.38 (d, J = 14.1 Hz, 1H), 4.23 (s, 1H). {*NMR probe temperature = 301 K*}.

Presented NMR peaks are from using the "Auto Assignment" algorithm in MestReNova version 12 software (Mestrelab Research, Santiago de Compostela, Spain). Which picks peaks in an inputted NMR spectrum, influenced by an inputted structure, which was structure 8 in the reaction scheme above. However, after automatic assignment, I manually made a single change for 7a: I changed the number of hydrogens at 7.95 ppm from 1H to 2H. I executed this change by manually extending the integration line (this manipulation automatically changes the MestReNova program's integration calculation method from "peak" to "sum") in this 7.95 ppm region for 7a, making it the same length as the integration line in the same region for 7b. Which then rendered/integrated this region with 2H as compared to 1H. The same as for 7b in this region (by "Auto Assignment" algorithm: 7b has peaks at 7.96 [1H] and 7.92 [1H] = 2H in 7.95 ppm region). This change in integration increases the nuclide count to the correct number for the structure (18H). This manipulation is best explained in pictures and so please refer to Figures S46 and S47. Which both show 7b data in the upper panel, 7a data in the lower panel. Wherein Figure S46 shows how this region was automatically integrated. And Figure S47 shows how I manually extended the integration line for 7a, to match the length of that for 7b. Such that both integrated to 2H in this region.

<u>Performed synthesis of 7a and 7b in detail</u>

In this section, numbers in square brackets are CAS numbers.

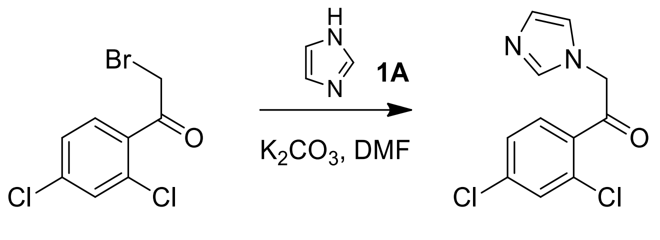

1 2

1. Charged Compound 1 [170894-53-8] (15.0 g, 55.9 mmol) and DMF [68-12-2] (150 mL) into the reactor at 25°C under $N_2$.

2. Charged $K_2CO_3$ [584-08-7] (23.2 g, 167 mmol) into the reactor at 25°C.

3. Charged imidazole [288-32-4] (11.4 g, 167 mmol) into the reactor at 25°C.

4. The mixture was stirred at 25°C for 3 hours.

5. TLC (petroleum ether:ethyl acetate = 5:1, $R_f$ = 0.0) showed raw material was consumed.

6. The mixture was quenched with water (100 mL).

7. The aqueous phase was extracted with ethyl acetate (100 mL x 2). The combined organic phase was washed with brine (50 mL), dried with anhydrous $Na_2SO_4$, filtered and concentrated in vacuum.

8. The residue was purified by silica gel chromatography (column height: 250 mm, diameter: 100 mm, 100-200 mesh silica gel, petroleum ether/ethyl acetate=20/1, 3/1) to give Compound 2 (13.0 g, 38.8 mmol, 69.3% yield, 76.1% purity) as yellow solid.

Compound 2: $^1$H NMR ($CDCl_3$, 400 MHz) δ (ppm) 7.55 (d, J = 8.0 Hz, 1H), 7.50 (s, 1H), 7.35-7.37 (dd, J = 8.0 Hz, 8.0 Hz, 1H), 7.09 (d, J = 8.0 Hz, 1H), 6.93 (s, 1H), 5.32 (s, 2H).

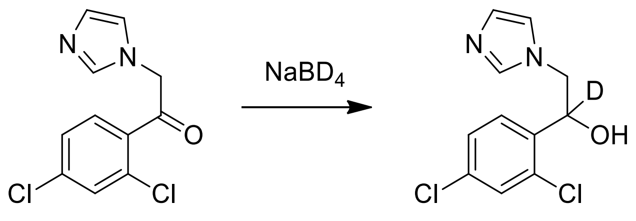

2 3

1. Charged Compound 2 (8.00 g, 31.3 mmol) and MeOH (40 mL) into the reactor at 25°C under $N_2$.

2. Charged $NaBD_4$ [15681-89-7] (1.31 g, 34.5 mmol) to the solution.

3. The mixture was stirred at 50°C for 3 hours.

4. LC-MS (product: RT= 0.914 min) showed raw material was consumed.

5. The reaction was quenched with water (100 mL).

6. The aqueous phase was extracted with ethyl acetate (50 mL x 2). The combined organic phase was washed with brine (50 mL), dried with anhydrous $Na_2SO_4$, filtered and concentrated in vacuum.

7. The residue was purified by silica gel chromatography (column height: 250 mm, diameter: 100 mm, 100-200 mesh silica gel, petroleum ether/ethyl acetate=10/1, 0/1) to give Compound 3 (5.50 g, 21.3 mmol, 67.9% yield) as white solid.

Compound 3: $^1H$ NMR ($CDCl_3$, 400 MHz) δ (ppm) 7.57 (d, J = 8.0 Hz, 1H), 7.39 (m, 1H), 7.28-7.30 (dd, J = 8.0 Hz, 8.0 Hz, 1H), 6.88 (d, J = 8.0 Hz, 2H), 4.20 (d, J = 12 Hz, 1H), 3.84-3.88 (d, J = 16 Hz, 1H).

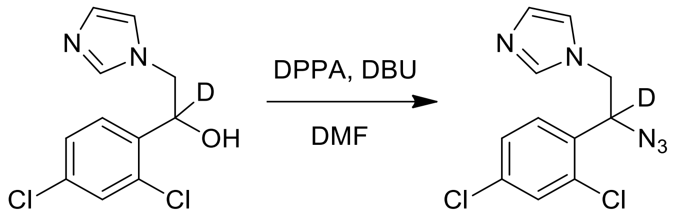

3 4

1. Set up a reactor R-1. (Note: R-1 is a 100 mL three-necked bottle)

2. Charged Compound 3 (4.50 g, 17.4 mmol) and DPPA [26386-88-9] (4.80 g, 17.4 mmol, 3.78 mL), DMF (22.5 mL) into the reactor R-1 at 25-30°C.

3. Charged DBU [6674-22-2] (2.65 g, 17.4 mmol, 2.63 mL) into the mixture under nitrogen atmosphere at 0-5°C, then the mixture was stirred for about 15 minutes.

4. After that, the reaction mixture was stirred for 24 hours at 20-30°C.

5. TLC (DCM/MeOH = 10/1, SM $R_f$ = 0.55, product $R_f$ = 0.72) showed raw material was consumed, LC-MS (product: RT = 0.764 min) and HPLC (product: RT = 1.893 min) indicated reaction mixture has desired product.

6. The reaction mixture was extracted with EtOAc (40 mL x 3).

7. The combined organic phase was washed with brine (40 mL), dried with $Na_2SO_4$, filtered and concentrated in vacuum to get the residue.

8. The residue was purified by column chromatography ($SiO_2$, dichloromethane: methanol = 1/0 to 100/1) to give Compound 4 (3.80 g, crude) as yellow oil.

Compound 4: $^1H$ NMR ($CDCl_3$, 400 MHz) δ (ppm) 7.50 (s, 1H), 7.44 (s, 1H), 7.35 (d, J = 2.0 Hz, 1H), 7.33 (d, J = 2.0 Hz, 1H), 7.29-7.30 (m, 1H), 6.94-7.10 (m, 1H), 4.25 (d, J = 14.4 Hz, 1H), 4.04 (d, J = 14.4 Hz, 1H).

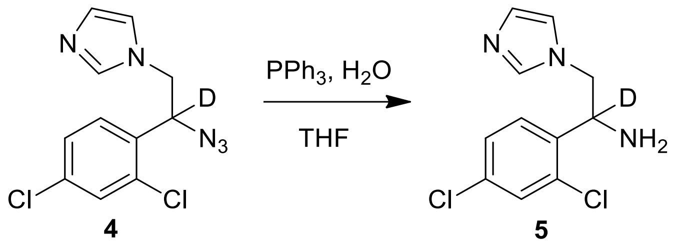

1. Set up a reactor R-1. (Note: R-1 is a 100 mL three-necked bottle)

2. Charged Compound 4 (3.80 g, 13.4 mmol), $PPh_3$ [603-35-0] (10.5 g, 40.2 mmol), THF [109-99-9] (12.6 mL) and $H_2O$ (1.26 mL) into the reactor R-1 at 25-30°C.

3. Stirred at 65-70°C for 16 hours.

4. Checked the reaction mixture by TLC (dichloromethane: methanol=10:1, SM: $R_f$ = 0.54, product: $R_f$ = 0.45) and LC-MS (product: RT = 1.253min) showed that reactant was consumed completed.

5. The reaction mixture was concentrated under reduced pressure to remove THF.

6. The reaction mixture was added with $H_2O$ (38 mL), adjusted pH to 3 with con. HCl and washed with EtOAc (38 mL).

7. The water phase was adjusted pH to 13 with 1N aq. NaOH, then exacted with EtOAc (38 mL x 2).

8. The combined organic phase was washed with brine (38 mL), dried with $Na_2SO_4$, filtered and concentrated in vacuum to give Compound 5 (2.60 g, crude) as a yellow liquid.

Compound 5: $^1$H NMR ($CDCl_3$, 400 MHz) δ (ppm) 7.44 (d, J = 14 Hz, 1H), 7.41 (s, 1H), 7.29 (d, J = 2.0 Hz, 1H), 7.27 (d, J = 1.2 Hz, 1H), 7.07 (s, 1H), 6.92 (s, 1H), 4.21 (d, J = 14 Hz, 1H), 3.92 (d, J = 14 Hz, 1H).

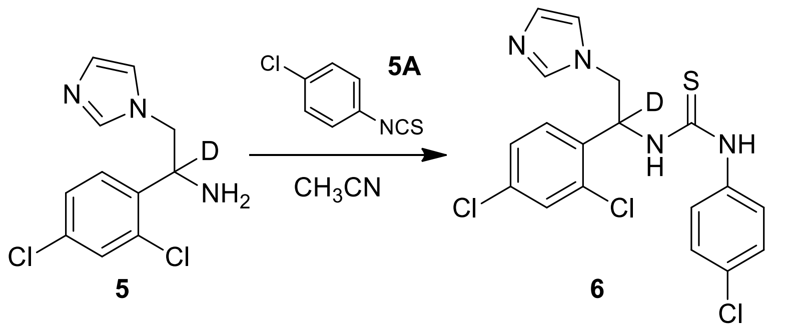

1. Charged Compound 5 (2.50 g, 9.72 mmol), Compound 5A [2131-55-7] (1.65 g, 9.72 mmol) and MeCN (37.5 mL) into the reactor at 25°C.

2. The mixture was stirred at 25°C for 12 hours.

3. TLC (dichloromethane: methanol=10:1, SM: $R_f$ = 0.5, product: $R_f$ = 0.2) showed raw material was consumed completed.
4. The mixture was concentrated in vacuum.
5. The residue was purified by silica gel chromatography (column height: 250 mm, diameter: 100 mm, 100-200 mesh silica gel, petroleum ether/ethyl acetate=3/1, 0/1) to give Compound 6 (3.60 g, 8.44 mmol, 86.7% yield) as white solid.
Compound 6: $^{1}H$ NMR ($CDCl_3$, 400 MHz) δ (ppm) 9.08 (s, 1H), 7.48 (d, J = 2.0 Hz, 1H), 7.33 (d, J = 8.0 Hz, 1H), 7.25-7.28 (m, 1H), 7.16 (d, J = 2.0 Hz, 1H), 7.00 (s, 1H), 6.88 (d, J = 8.0 Hz, 1H), 6.72 (s, 1H), 4.46 (dd, J = 14.4 Hz, 14 Hz, 1H), 1.84 (s, 1H).

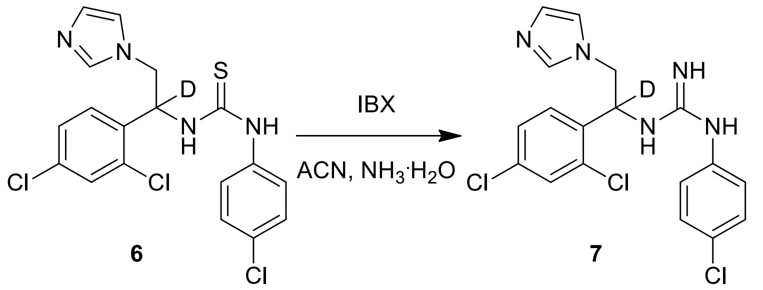

1. Charged Compound 6 (3.00 g, 7.03 mmol) and MeCN (30 mL) into the reactor at 25°C.
2. Charged IBX [61717-82-6] (2.17 g, 7.73 mmol) into the reactor at 25°C.
3. Charged $NH_3.H_2O$ [1336-21-6] (13.6 g, 116 mmol, 15 mL, 30% purity) into the reactor at 25°C.
4. The mixture was stirred at 25°C for 12 hours.
5. LC-MS (product: RT = 0.717 min) showed raw material was consumed completed.
6. The mixture was filtered and the filter cake was concentrated in vacuum to give Compound 7 (2.50 g, 6.10 mmol, 86.8% yield) as white solid.
Compound 7: $^{1}H$ NMR ($CDCl_3$, 400 MHz) δ (ppm) 7.87 (d, J = 8.0 Hz, 1H), 7.67 (s, 1H), 7.55-7.60 (m, 2H), 7.37-7.41 (m, 2H), 7.23 (s, 1H), 7.07-7.09 (m, 1H), 6.92 (s, 1H), 4.33 (dd, J = 14.4 Hz, 14 Hz, 1H).

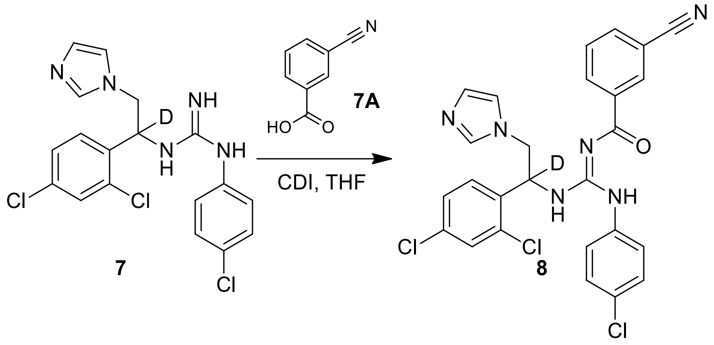

1. Charged Compound 7 (538 mg, 3.66 mmol), CDI [530-62-1] (593 mg, 3.66 mmol) and THF [109-99-9] (30 mL) into the reactor at 25°C.
2. The solution was stirred at 25°C for 3 hours.
3. Charged compound 7A [1877-72-1] (1.50 g, 3.66 mmol) into the reactor at 25°C.
4. The mixture was stirred at 25°C for 12 hours.
5. LC-MS (SM: RT = 0.730 min, product: RT = 1.030 min) showed the raw material was consumed/completed.
6. The mixture was concentrated in vacuum.
7. Separating stereoisomers of Compound 8: the residue was purified with Prep-HPLC (column: DAICEL CHIRALPAKAD-H (250 mm*30 mm, 5 µm); mobile phase: [0.1% $NH_3H_2O$ IPA]; B%: 44%-44%, 6.45 min) to give 7b (0.025 g, 45.8 µmol, 1.25% yield, 98.8% purity) as white solid and 7a (0.025 g, 45.5 µmol, 1.24% yield, 98.1% purity) as white solid.

<u>Spectral data for 7b</u>

Its expected/calculated *m/z*, from its structure (Figure 19A), using the "exact molecular weight" calculator in MarvinSketch software (Chemaxon, Budapest, Hungary), which uses the mass of the *most prevalent* isotope of each element in the molecule:
538.08 $[M+H]^+$, 560.06 $[M+Na]^+$, 269.54 $[M+2H]^{2+}$, 576.04 $[M+K]^+$).

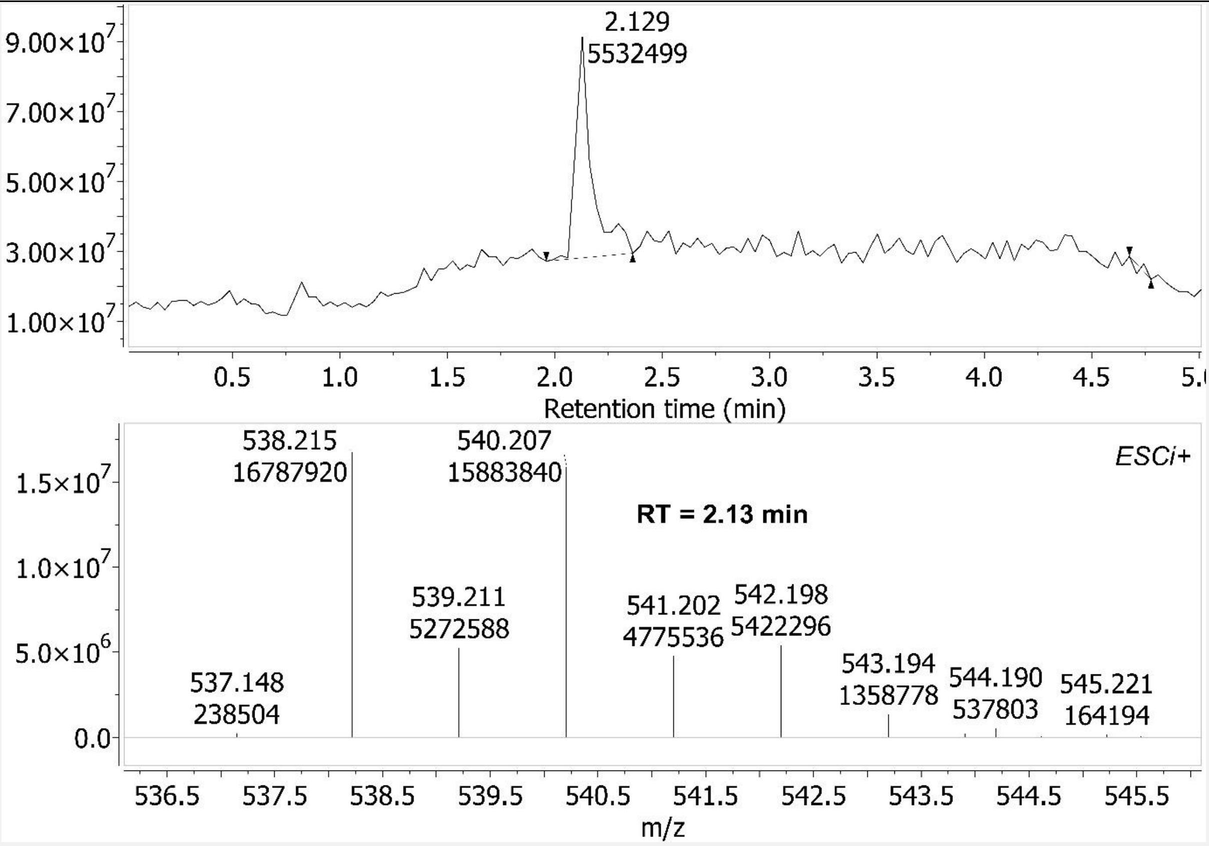

Figure S33 – Chiral SFC-MS for 7b sample. *Upper panel*: Chiral-SFC annotated with Retention Time (RT, minutes) and area under peak. *S* stereoisomer is found in [RT=2.13] peak. *Lower panel*: component of mass spectrum of [RT=2.13] peak. And therein is *m/z* 538.215 $[M+H]^+$ (*0.135 removed from Expected. Better concordance in subsequent spectra*).

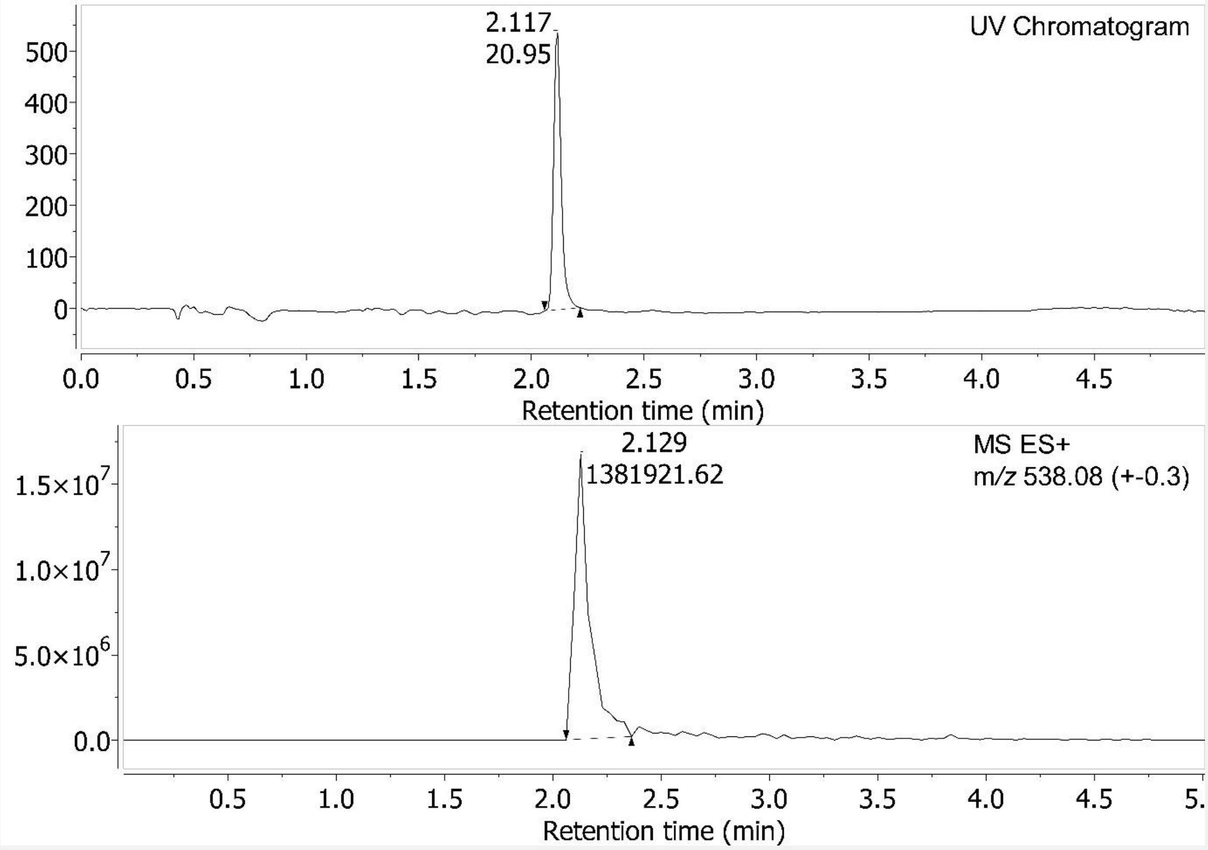

Figure S34 – Chiral-SFC of 7b sample. *S* stereoisomer excess = 100% (no other peak in the UV chromatogram). Enantiomeric excess (ee) = (100-50)*2 = 100%.

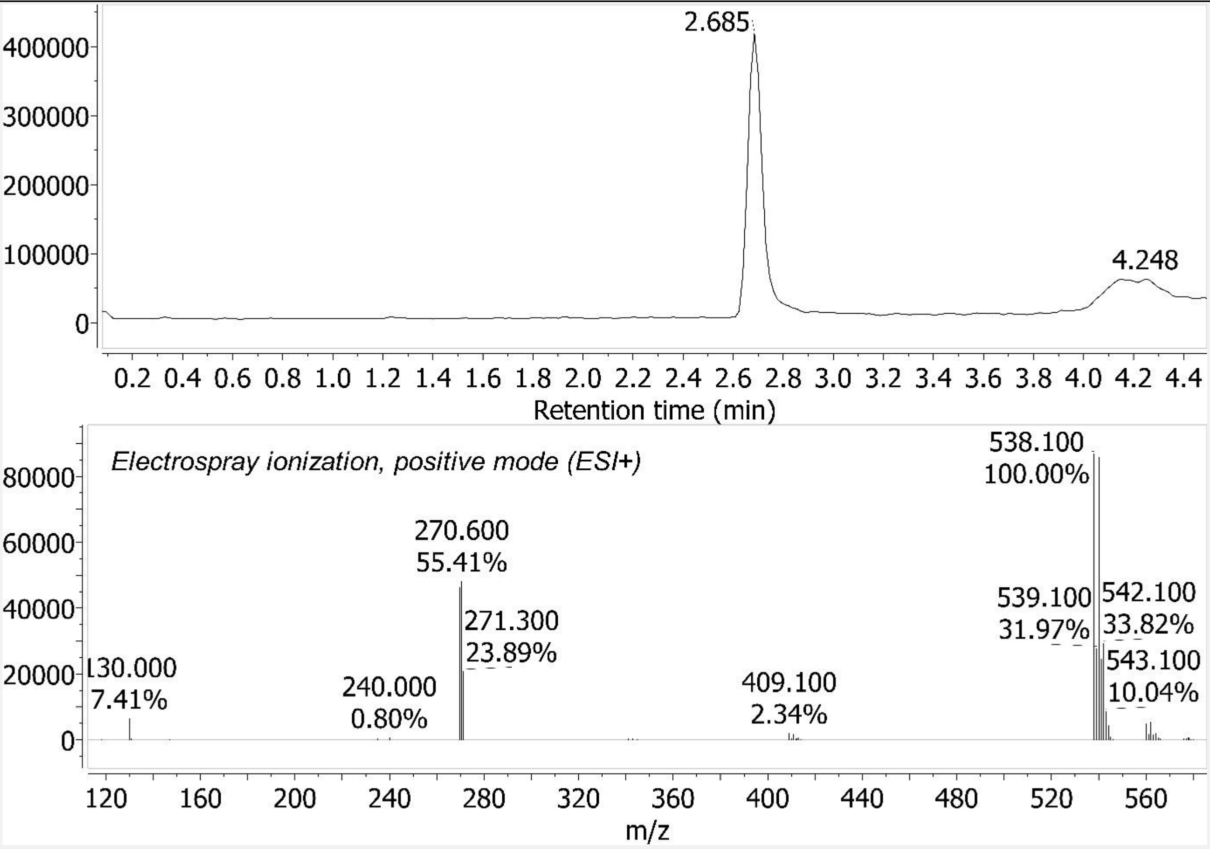

Figure S35 – LC-MS spectrum for 7b sample. *Upper panel*: Liquid Chromatography (LC) Retention Time (RT) = 2.685 minutes. *Bottom panel*: Mass Spectrum (MS) of [RT=2.685 peak]. Shows positively charged species (API-ES positive). Percentages are relative abundance.

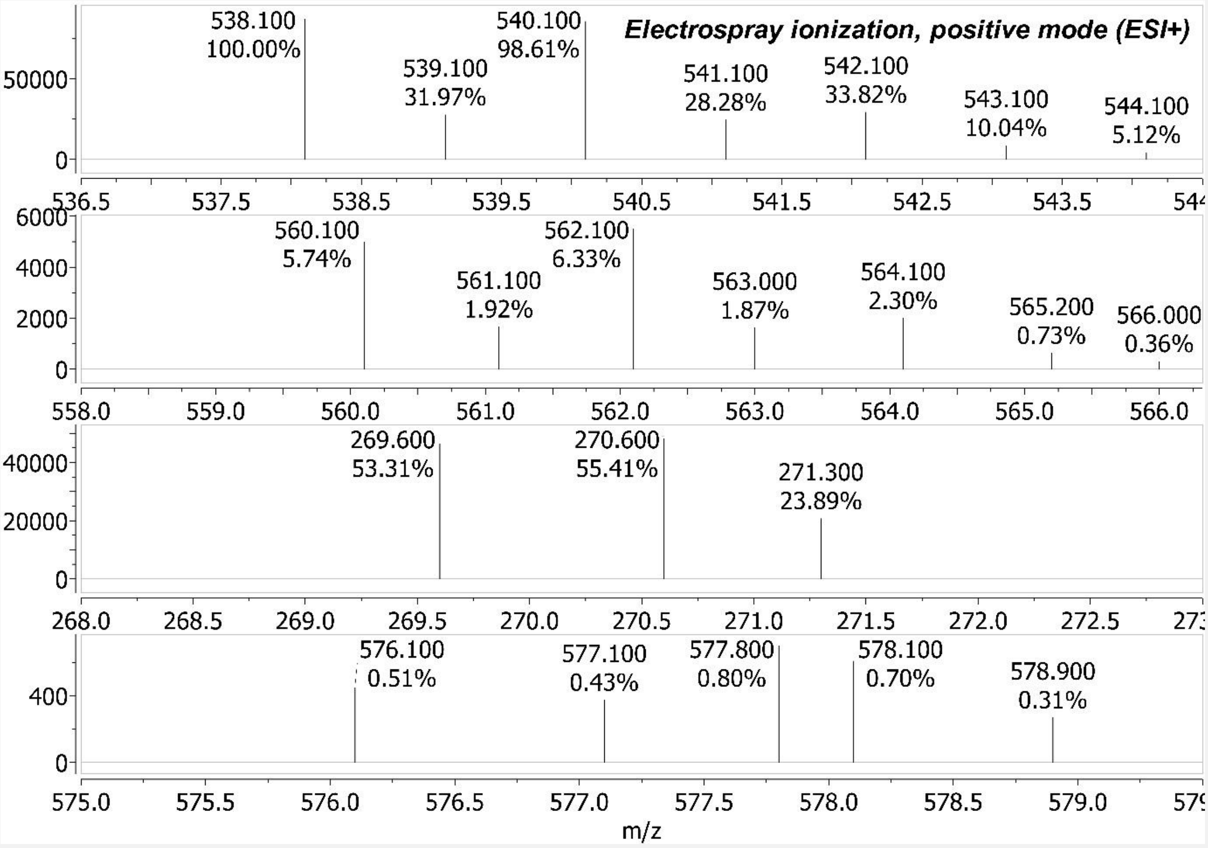

Figure S36 – Some Mass Spectrum (MS) components of Figure S35. <u>*Upper panel*</u>: *m/z* 538.1 $[M+H]^+$. <u>*2nd panel*</u>: *m/z* 560.1 $[M+Na]^+$. <u>*3rd panel*</u>: *m/z* 269.6 $[M+2H]^{2+}$. <u>*Bottom panel*</u>: *m/z* 576.1 $[M+K]^+$. All these Observed *m/z* are within 0.06 of Expected.

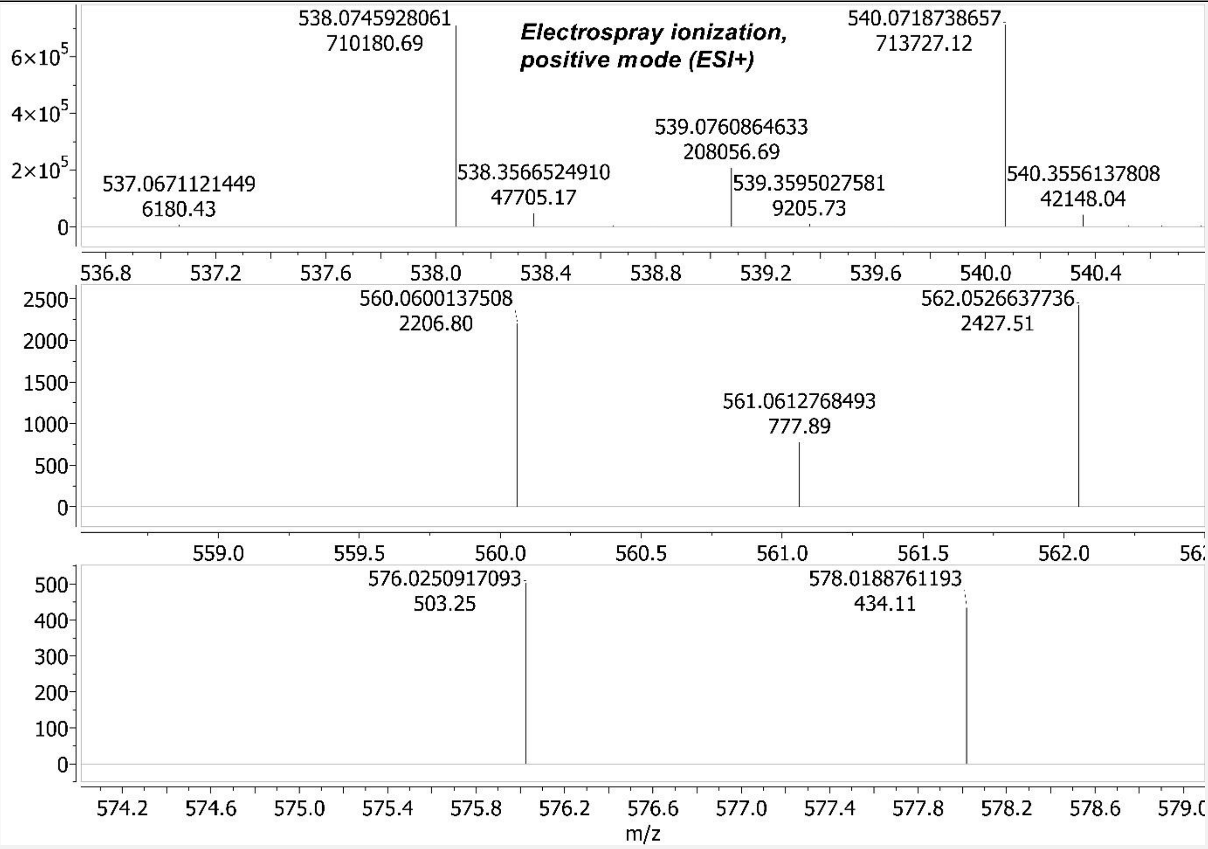

Figure S37 – High Resolution Mass Spectrometry (HRMS): Liquid Chromatography-Time of Flight (LC-TOF) Mass Spectrometry (MS) for 7b sample. LC Retention Time (RT) = 0.166 minutes (LC data not shown). _Upper panel_: OBSERVED *m/z* 538.0745928061 $[M+H]^+$. CALCULATED from structure *m/z* 538.0821456 $[M+H]^+$. So, Observed mass is only 0.0075527939 Daltons (0.7% the mass of a proton) removed from calculation. The species with *m/z* 537.0671121449 $[M+H]^+$ has hydrogen instead of deuterium on the chiral carbon. And indeed is the mass of a neutron (1.008664 Daltons) lighter: (538.0745928061-537.0671121449) = 1.0074806612 Daltons mass difference (0.12% removed from being the exact mass of a neutron difference). Absolute abundance is shown on the y-axis. And the molar percent deuterium incorporation for 7b sample at the chiral carbon is 100% - ((6180.43/710180.69)*100=0.87%) = 99.13%. _2$^{nd}$ panel_: OBSERVED *m/z* 560.0600137508 $[M+Na]^+$. CALCULATED from structure *m/z* 560.0640898 $[M+Na]^+$. So, Observed mass is only 0.0040760492 Daltons (0.4% the mass of a proton) removed from calculation. The species with hydrogen instead of deuterium on chiral carbon is too low abundance to be observed here. _Bottom panel_: OBSERVED *m/z* 576.0250917093 $[M+K]^+$. CALCULATED from structure *m/z* 576.0380270 $[M+K]^+$. So, Observed mass is only 0.0129352907 Daltons (1.3% the mass of a proton) removed from calculation. The species with hydrogen instead of deuterium on chiral carbon is too low abundance to be observed here.

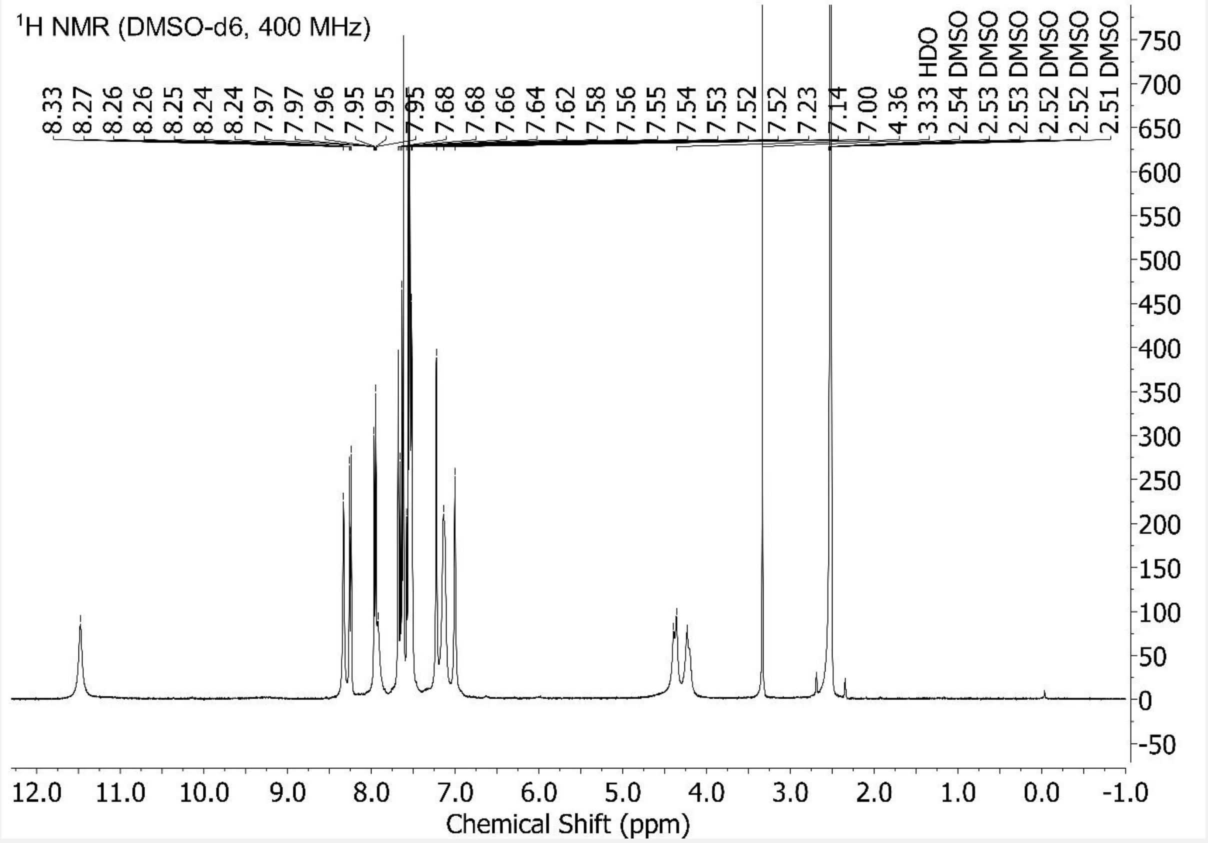

Figure S38 – $^{1}$H NMR {*NMR probe temperature = 301 K*} for 7b sample. Scaled to highest compound peak. Peaks identified by "Auto peak picking" algorithm of MestReNova v12 software. With default program settings used. And no chemical structure used/inputted to guide/constrain this peak picking.

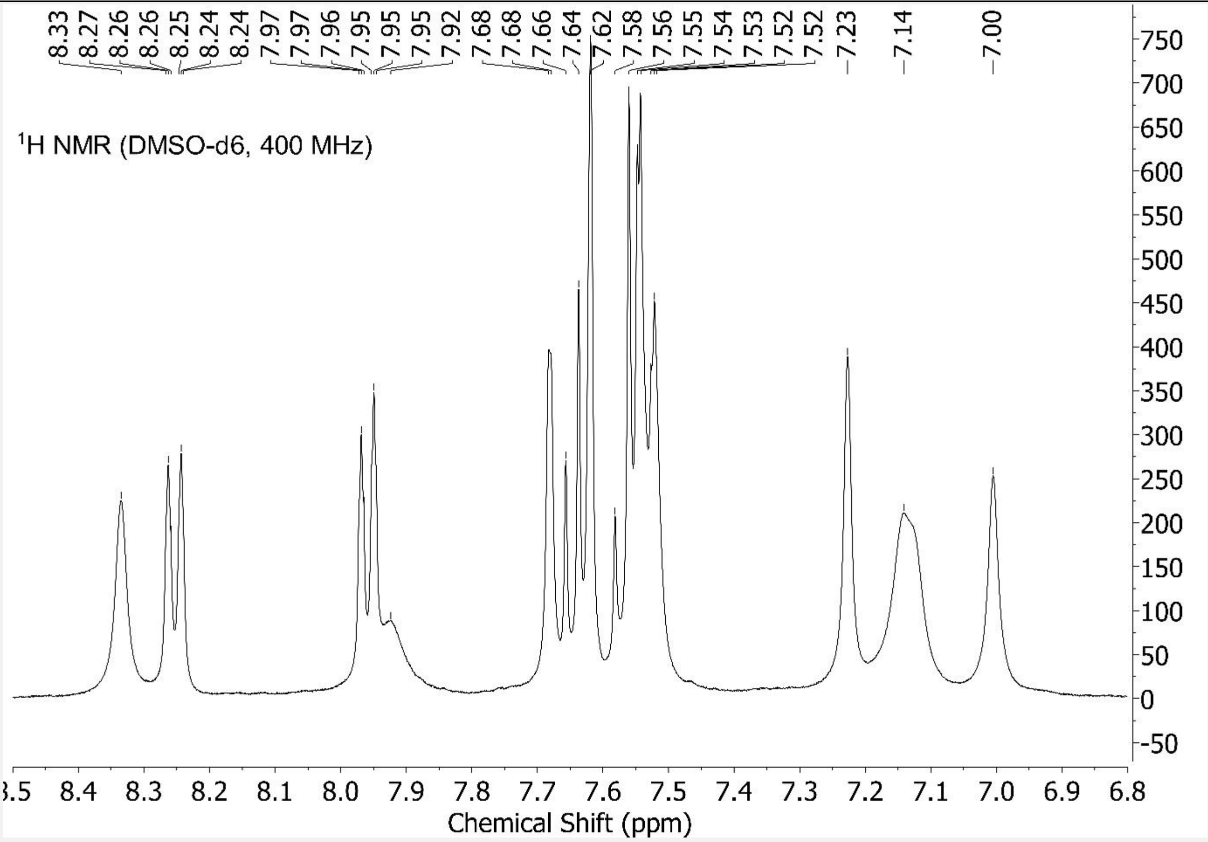

Figure S39 – Aromatic region of Figure S38 expanded.

Spectral data for 7a

Note that for 7a, its LC-MS, HRMS and NMR data are not all correspondent. To explain, LC-MS, HRMS and $^{1}$H NMR were performed for 7a. And its $^{1}$H NMR showed a high amount of impurities (spectrum not shown). Further mass-based purification steps were taken as a result. And the $^{1}$H NMR was repeated (spectrum shown herein). And $^{13}$C NMR performed. All showing a much lower, acceptable level of impurity. And it is this purer form that was entered into anti-cancer testing at the National Cancer Institute (Figure 19B). However, LC-MS and HRMS were not repeated. But the LC-MS and HRMS are clear enough to identify the [M+ion] species needed. All be it with some additional species/impurities also observed, which aren't in the LC-MS and HRMS spectra for 7b (e.g., compare Figures S37 and S43).

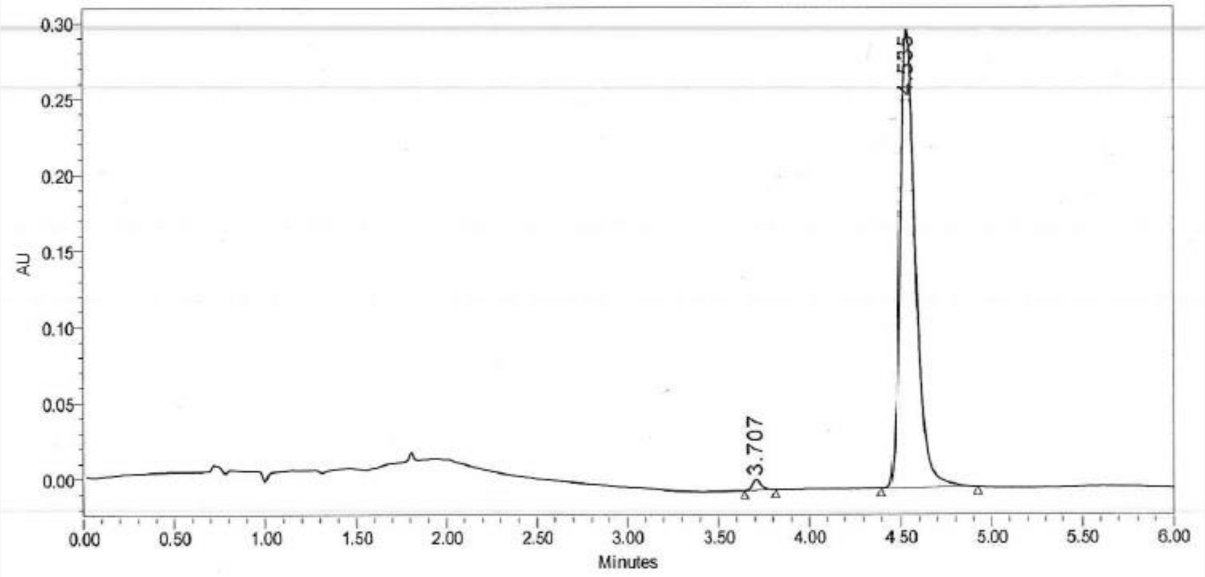

| | RT | Area | % Area | Height |
|---|---|---|---|---|
| 1 | 3.707 | 23021 | 1.29 | 6747 |
| 2 | 4.535 | 1765686 | 98.71 | 301165 |

Figure S40 – The chiral SFC-MS machine broke down. And so a different machine, with only chiral-SFC capability, had to be used. So, only chiral-SFC was performed for the 7a sample. Retention Time (RT) = 4.535 min.

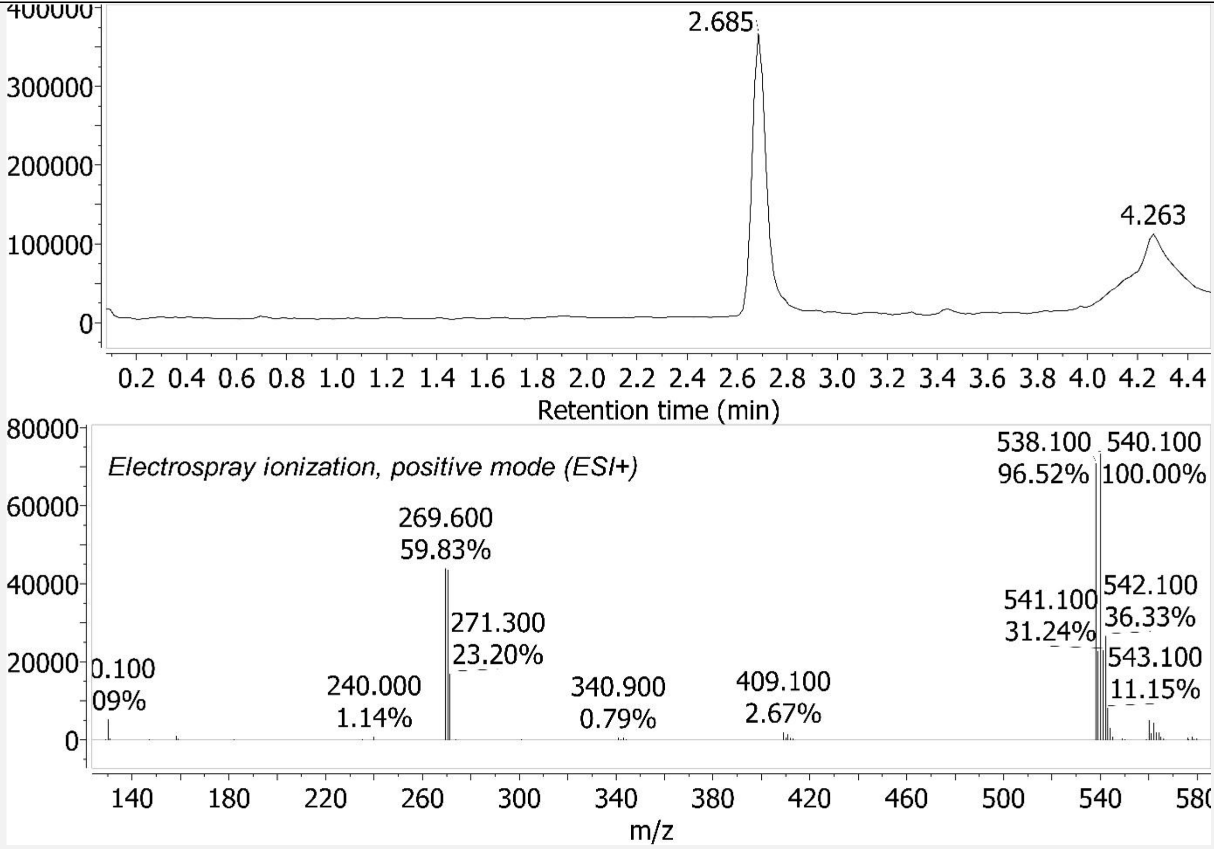

Figure S41 – LC-MS spectrum for 7a sample. Retention Time (RT) = 2.685 minutes.

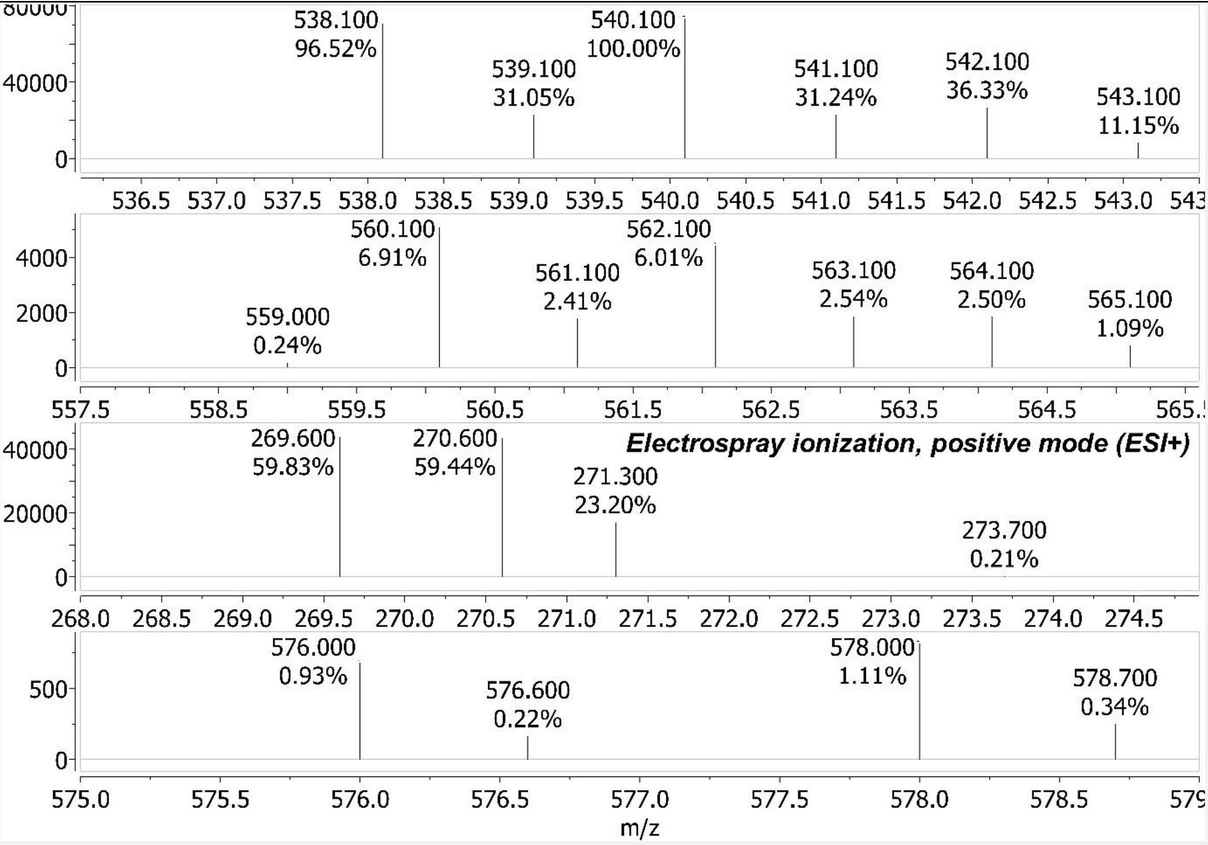

Figure S42 – Some Mass Spectrum (MS) components of Figure S41. _Upper panel_: *m/z* 538.1 $[M+H]^+$. _2nd panel_: *m/z* 560.1 $[M+Na]^+$. _3rd panel_: *m/z* 269.6 $[M+2H]^{2+}$. _Bottom panel_: *m/z* 576.0 $[M+K]^+$. All these Observed *m/z* are within 0.06 of Expected.

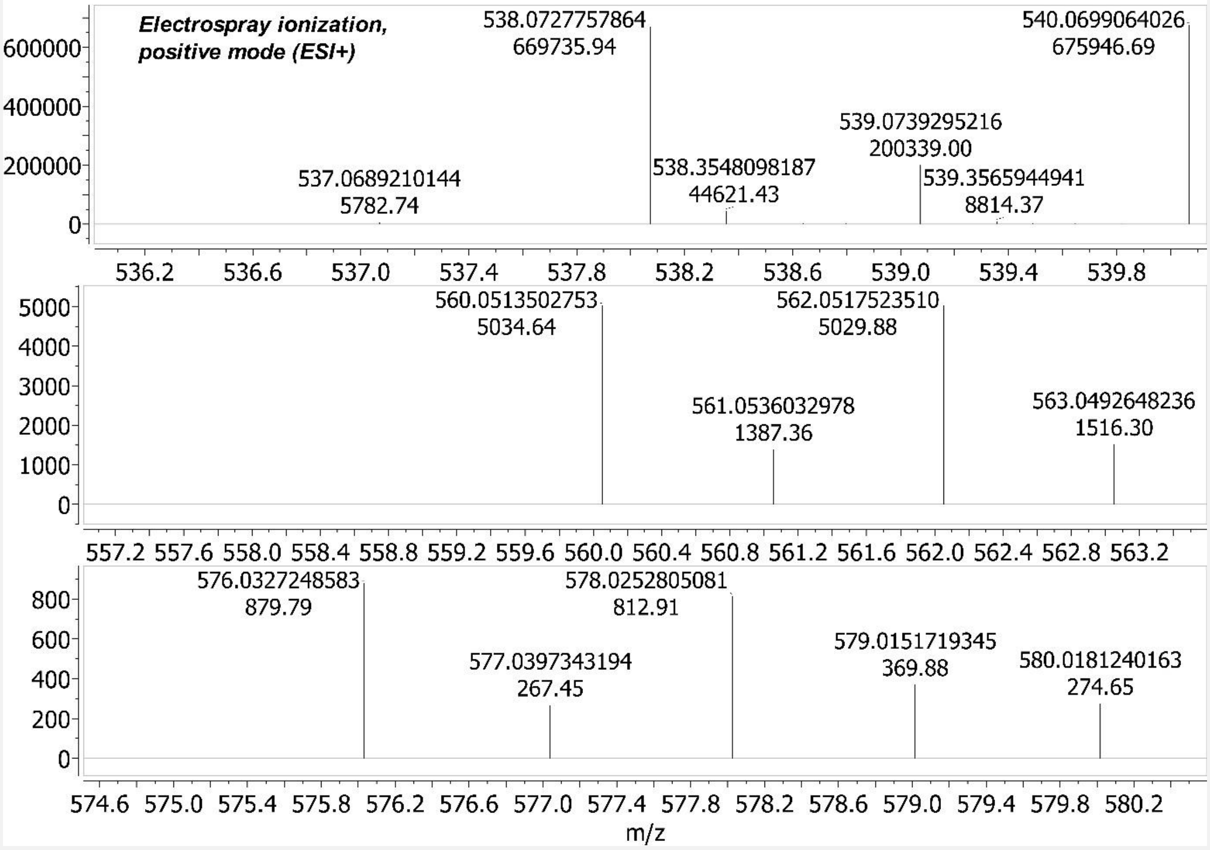

Figure S43 – HRMS: LC-TOF MS for 7a sample. LC Retention Time (RT) = 0.163 minutes (LC data not shown). *Upper panel*: *m/z* 538.0727757864 $[M+H]^+$. The species with *m/z* 537.0689210144 $[M+H]^+$ has hydrogen instead of deuterium on the chiral carbon. Absolute abundance is shown on the y-axis. And the molar percent deuterium incorporation for 7a at the chiral carbon is 100% - ((5782.74/669735.94)*100=0.86%) = 99.14%. *2nd panel*: *m/z* 560.0513502753 $[M+Na]^+$. *Bottom panel*: *m/z* 576.0327248583 $[M+K]^+$.

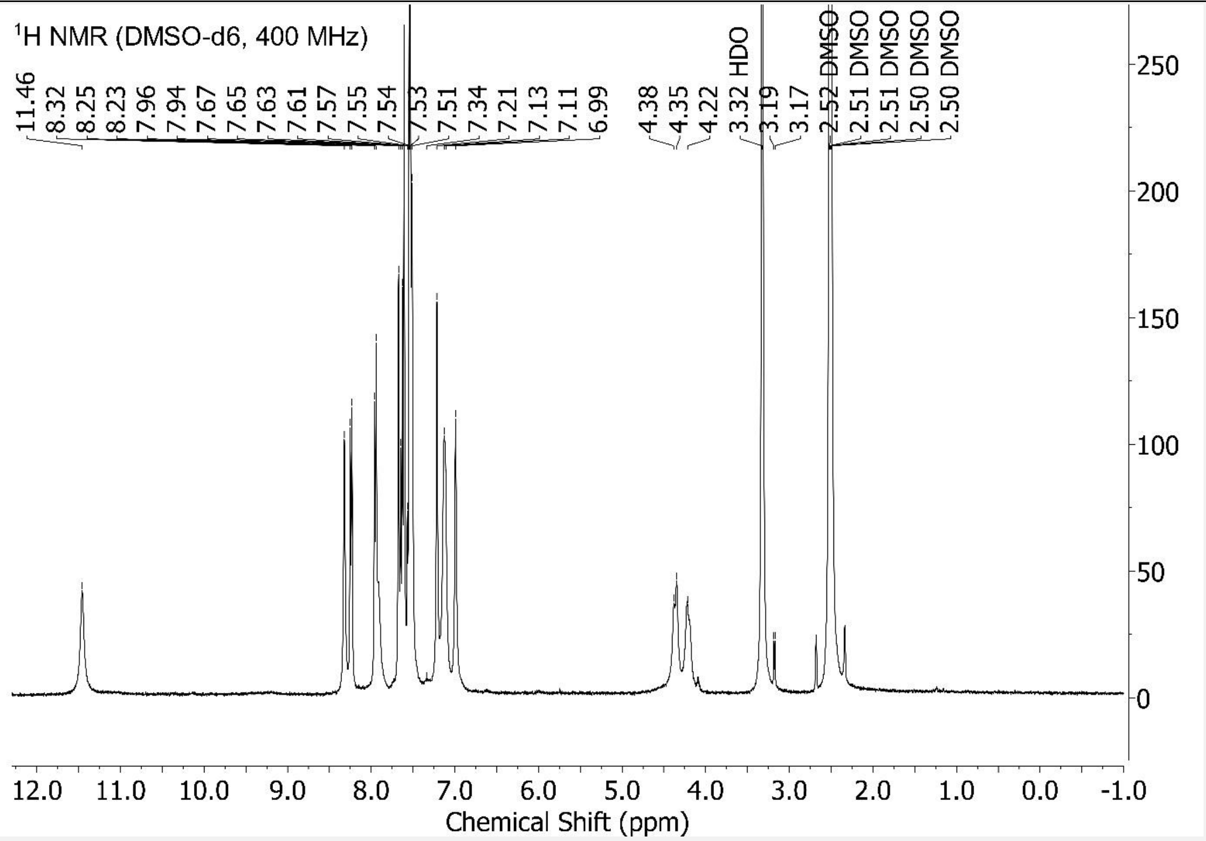

Figure S44 – $^1$H NMR {*NMR probe temperature = 300.7 K*} for 7a sample. Scaled to highest compound peak. "Auto peak picking" algorithm in MestReNova v12 used to pick peaks shown. With default program settings used. And no chemical structure used/inputted to guide/constrain this peak picking.

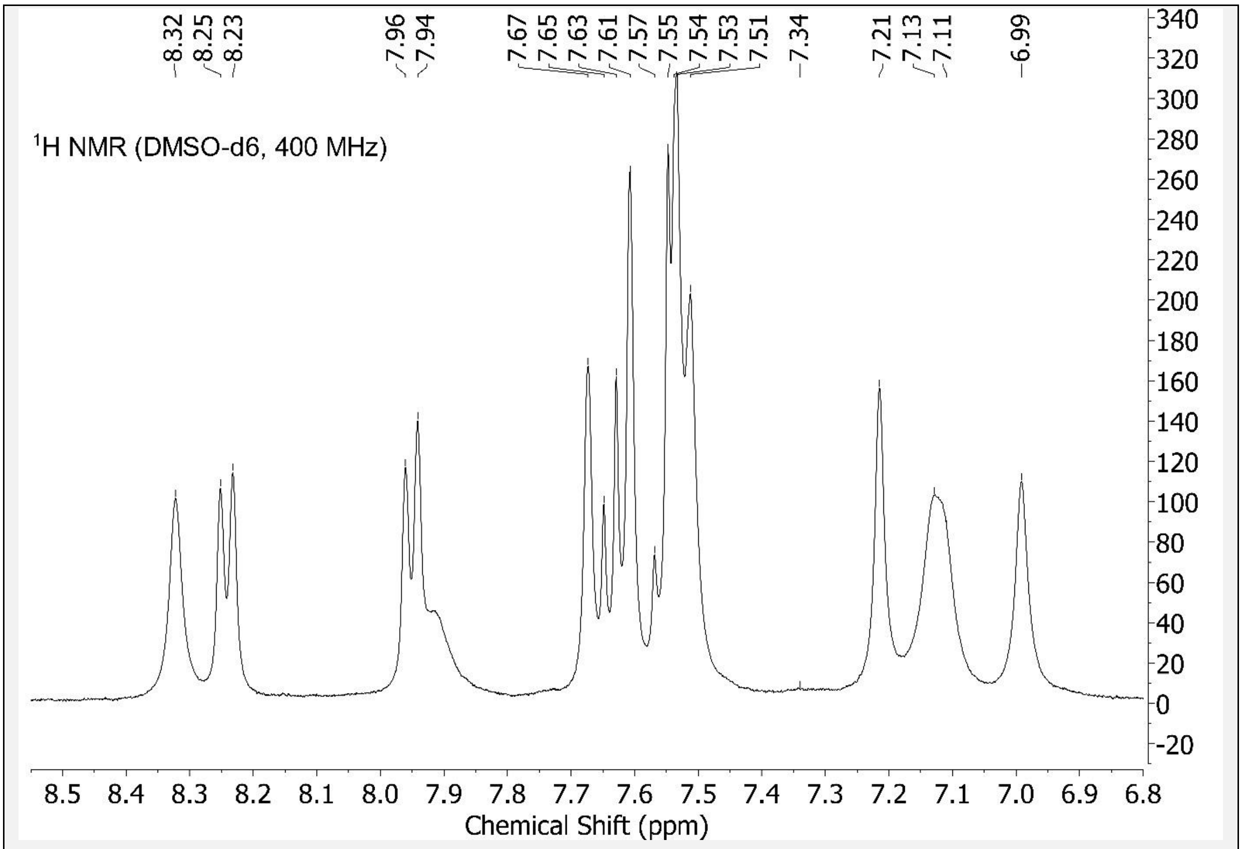

Figure S45 – Aromatic region of Figure S44 expanded.

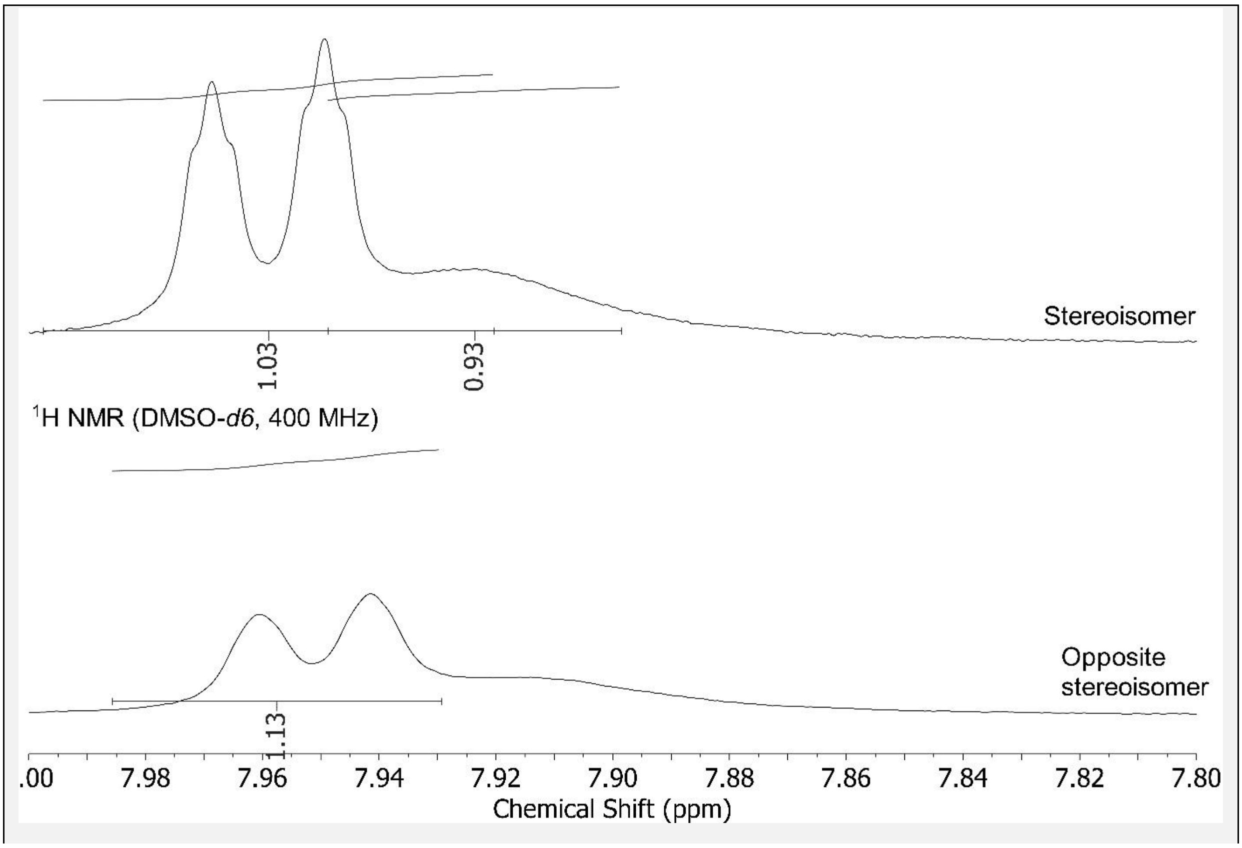

Figure S46 – 7b data in upper panel, 7a data in lower panel. Figure shows integration values in this ppm region as automatically assigned by the MestReNova v12 software used. 7b has 2H assigned (1.03+0.93=1.96, rounds to 2), whilst 7a has 1H assigned (1.13, rounds to 1), in this ppm region.

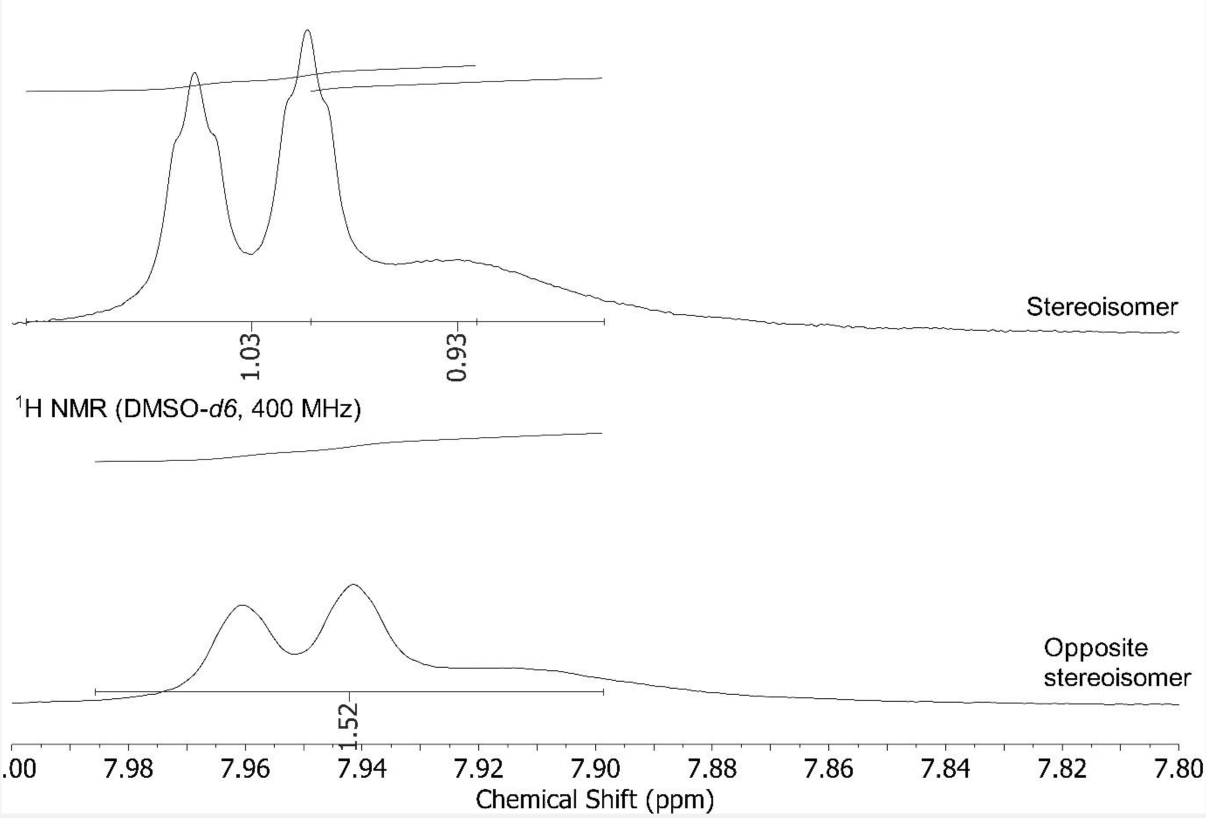

Figure S47 – Shown here, I manually extended the integration line (N.B., this automatically changes the MestReNova v12 integration calculation method from “peak” to “sum”) for 7a to be the same length as that for 7b. Which changes its integration value from 1.13 (prior) to 1.52, which rounds to 2H. This improves the 7a assignment because now it has the correct number of hydrogen nuclides (18H), whereas before it had (incorrectly) 17H. It is true that 1.52 only narrowly crosses the threshold for 2H. But interestingly, the integration value is also very low in this ppm region for the hydrogen on chiral carbon stereoisomers (integration value at 7.93 ppm = 1.51 for 6b, 1.57 for 6a. As assigned by “Auto Assignment” algorithm of MestReNova v12).

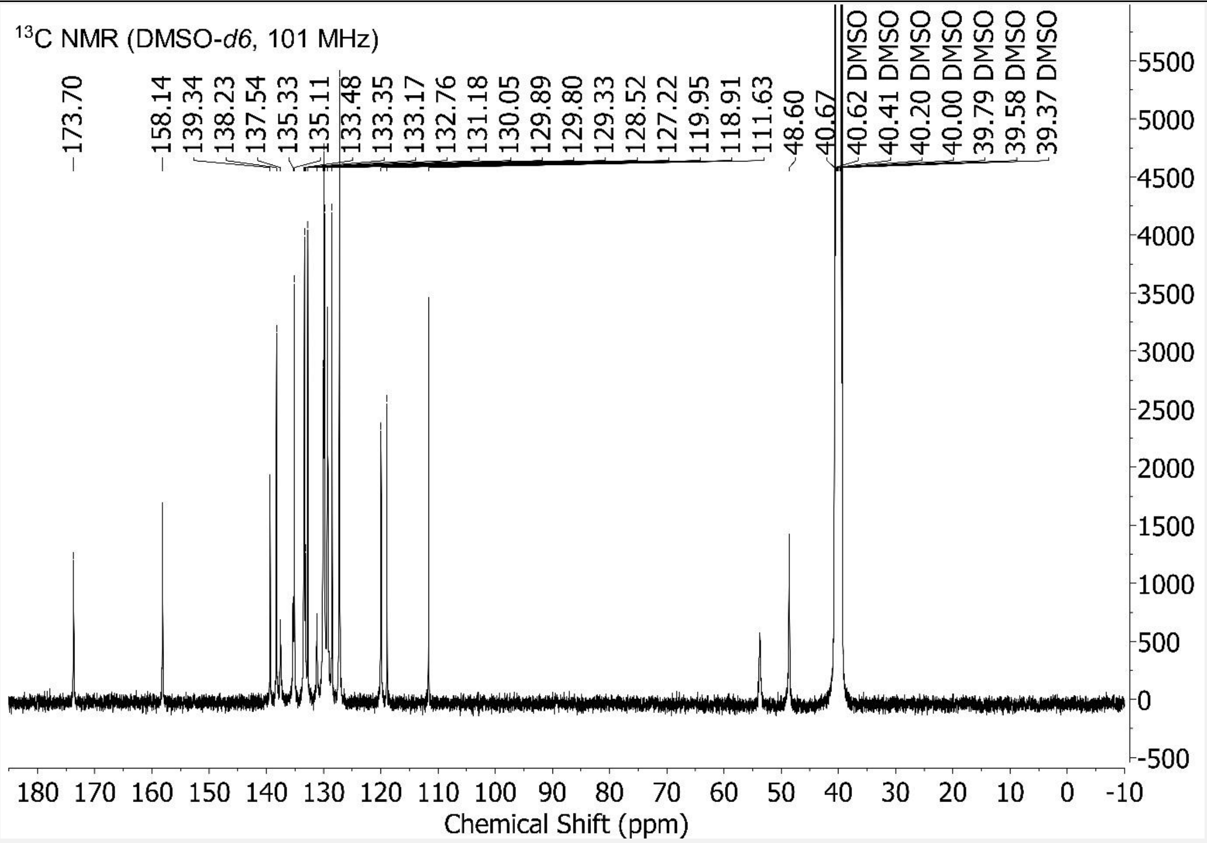

Figure S48 – $^{13}$C NMR {*NMR probe temperature = 311.9 K*} for 7a. Scaled to highest compound peak. "Auto peak picking" algorithm in MestReNova v12 used to pick peaks shown. With default program settings used. And no chemical structure used/inputted to guide/constrain this peak picking.

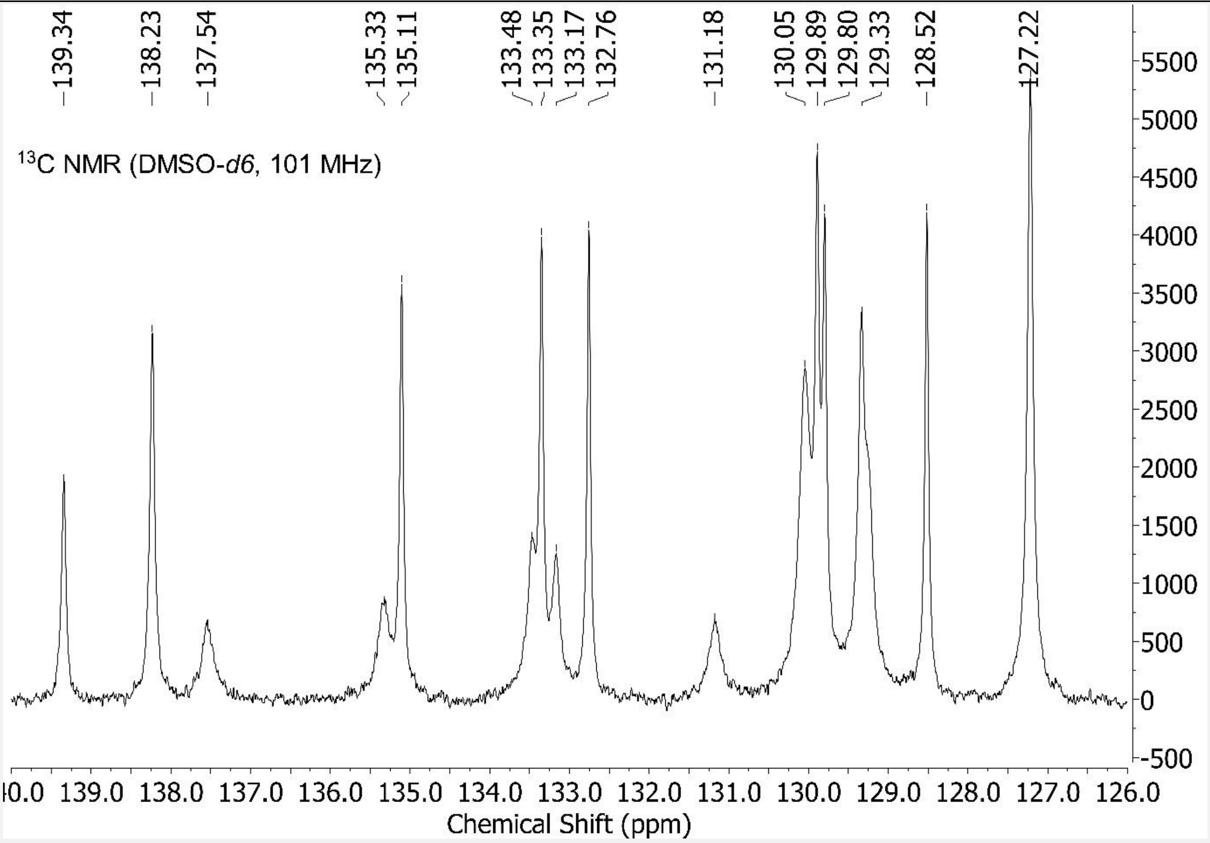

[Figure S49] – Region of Figure S48 expanded.

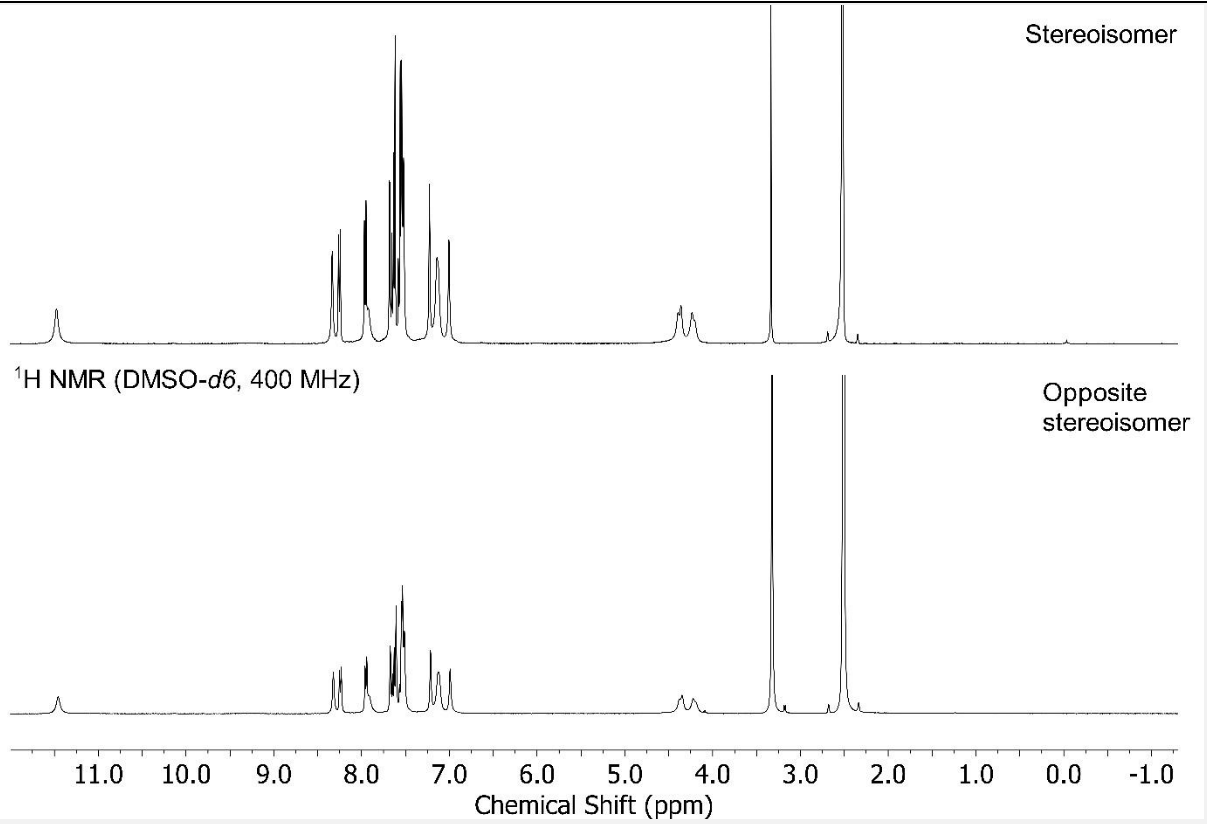

[Figure S50] – 7b data in upper panel, 7a data in lower panel.

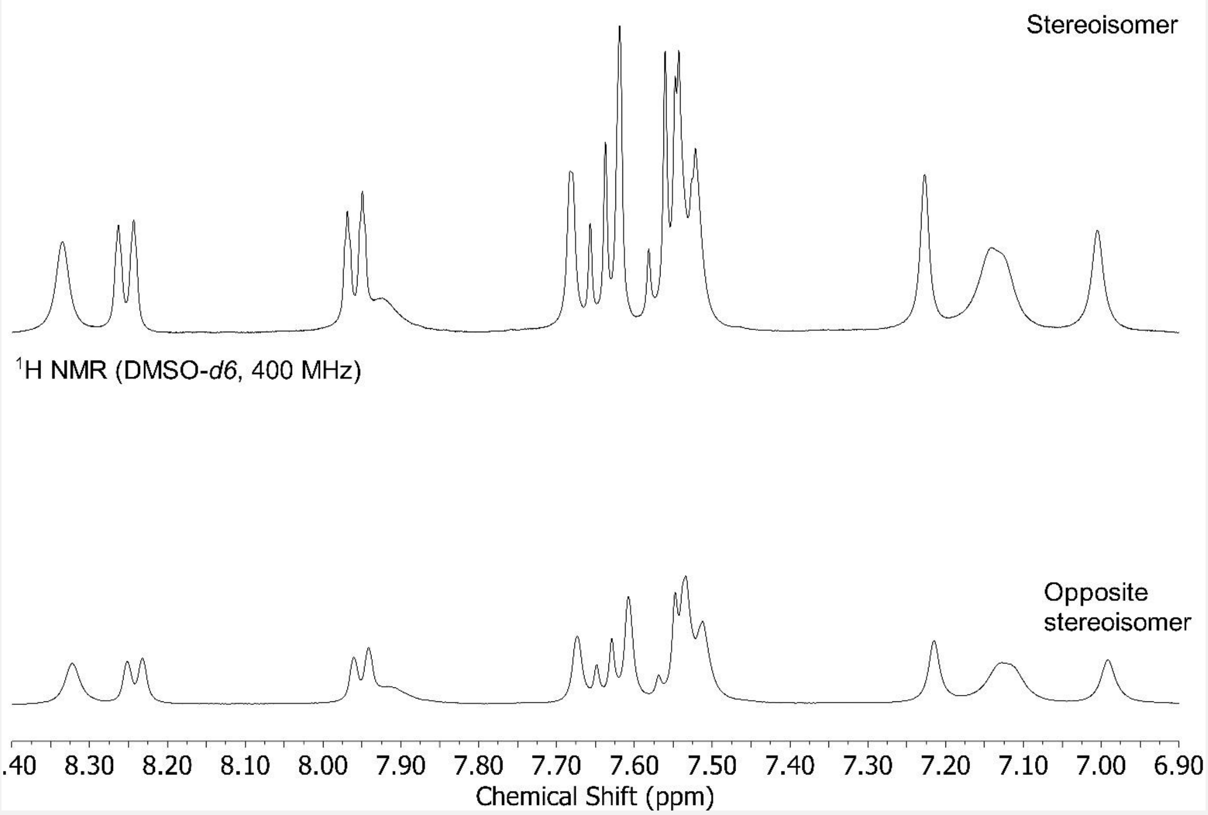

Figure S51 – Aromatic region. 7b data in upper panel, 7a data in lower panel.

Hydrogen vs. deuterium on chiral carbon

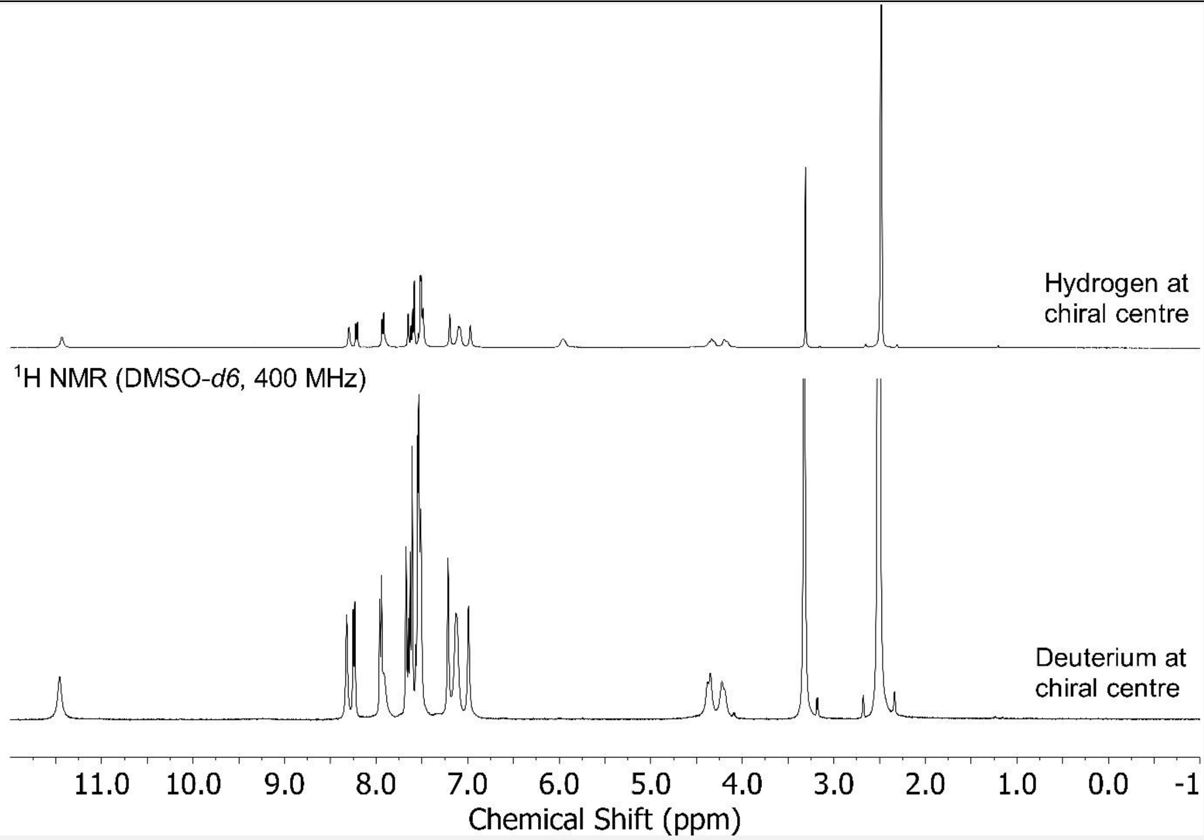

Figure S52 – *Upper panel*: $^1$H NMR spectrum for 6b. *Lower panel*: $^1$H NMR spectrum for 7a. The upper has a peak at 5.96 ppm from the hydrogen attached to its chiral carbon. The lower has this hydrogen replaced with a deuterium, and thence this 5.96 ppm peak is absent in its $^1$H NMR spectrum.

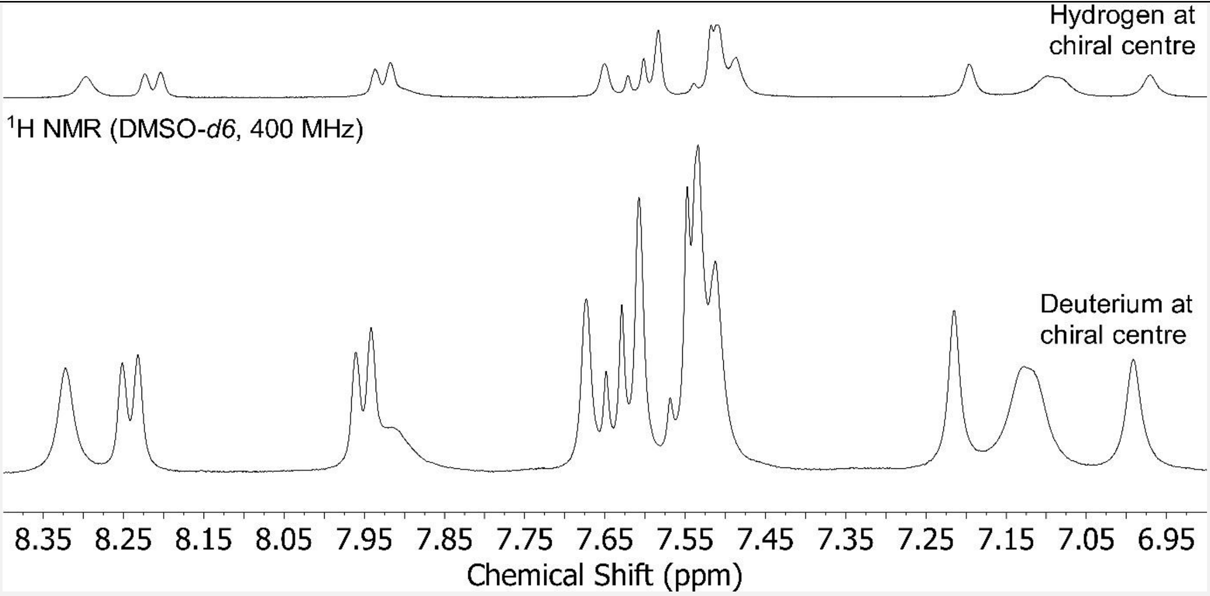

Figure S53 – Aromatic region. *Upper panel*: $^1$H NMR spectrum for 6b. *Lower panel*: $^1$H NMR spectrum for 7a.

R code used to generate Table 9

```
# R code
# Modified from code of Kang et al. paper “ivmodel: An R Package for Inference and Sensitivity Analysis of Instrumental Variables Models with One Endogenous Variable”
# Prerequisite to this code is having the ivmodel package installed in R (and any package dependencies thereof).
# Y is the dependent variable, D is the independent variable, and Z is the instrumental variable. The X covariate variable in the original code has been removed.
# Instrumental variable (specific F1F0 ATP hydrolysis rate) analysis reports causality between lower specific basal metabolic rate and longer maximal lifespan.
# Data is from Table S2 of this Supplementary Material. Human data excluded by rationale given in Methods section.
# N.B. a two-tailed p-value output can be made one-tailed by dividing by 2.
# When copying and pasting this code you may inadvertently copy and paste a page number, which must be removed before entering this code into the R console.
# You may need to press return after copying and pasting the code into the R console.

library(ivmodel)

Y <- c(20, 27, 22.8, 24, 9, 17.5, 17.7, 12, 3.8, 3.9, 4)
D <- c(0.000798458, 0.000771, 0.001319925, 0.001804232, 0.002887816, 0.002418352,
0.004975053, 0.003740752, 0.00601463, 0.006377, 0.010858546)
Z <- c(1.32, 0.81, 1.49, 1, 1.12, 2.89, 2.66, 2.71, 3.02, 3.9, 2.73)

card.model1IV = ivmodel(Y=Y,D=D,Z=Z)
card.model1IV

coefOther(card.model1IV)

card.modelFormula = ivmodelFormula(Y ~ D | Z)
card.modelFormula
```

R code used to generate Table 10

```
# R code
# Modified from code of Kang et al. paper "ivmodel: An R Package for Inference and Sensitivity Analysis of Instrumental Variables Models with One Endogenous Variable"
# Prerequisite to this code is having the ivmodel package installed in R (and any package dependencies thereof).
# Y is the dependent variable, D is the independent variable, and Z is the instrumental variable. The X covariate variable in the original code has been removed.
# Instrumental variable (specific IF1 protein activity) analysis reports causality between lower specific F1F0 ATP hydrolysis rate and longer maximal lifespan.
# Maximal lifespan data from [1] and other data is from [91]. Human data excluded by rationale given in Methods section.
# N.B. a two-tailed p-value output can be made one-tailed by dividing by 2.
# When copying and pasting this code you may inadvertently copy and paste a page number, which must be removed before entering this code into the R console.
# You may need to press return after copying and pasting the code into the R console.

library(ivmodel)

Y <- c(20, 27, 22.8, 24, 9, 17.5, 17.7, 12, 3.8, 3.9, 4)
D <- c(1.32, 0.81, 1.49, 1, 1.12, 2.89, 2.66, 2.71, 3.02, 3.9, 2.73)
Z <- c(3.49, 3.32, 3.34, 3.9, 5.06, 4.88, 4.44, 3.45, 0.65, 0.72, 0.81)

card.model1IV = ivmodel(Y=Y,D=D,Z=Z)
card.model1IV

coefOther(card.model1IV)

card.modelFormula = ivmodelFormula(Y ~ D | Z)
card.modelFormula
```

<u>R code used to generate Table 21</u>

```
# R code
# Modified from R code of Kang et al. paper "ivmodel: An R Package for Inference and Sensitivity Analysis of Instrumental Variables Models with One Endogenous Variable"
```

### Prerequisite to using this code is having the ivmodel package installed in R (and any package dependencies thereof).
### Y is the dependent variable, D is the independent variable, and Z is the instrumental variable. The X covariate variable in the original code has been removed.
### Instrumental variable (specific F1F0 ATP hydrolysis rate) analysis reports causality between specific basal metabolic rate and the ticking rate of the epigenetic clock (ΔM).
### Data is from Table S2 of this Supplementary Material, with ΔM calculated by Eq. 5 from maximal lifespan values in [1]. Human data excluded by rationale given in Methods section.
### N.B. a two-tailed p-value output can be made one-tailed by dividing by 2.
### When copying and pasting this code you may inadvertently copy and paste a page number, which must be removed before entering this code into the R console.
### You may need to press return after copying and pasting the code into the R console.

```
library(ivmodel)

Y <- c(0.154552123, 0.12471466, 0.140826476, 0.135754022, 0.266432808, 0.219921444,
0.462594189, 0.455115987, 0.447939867)
D <- c(0.000798458, 0.000771, 0.001319925, 0.001804232, 0.002887816, 0.003740752,
0.00601463, 0.006377, 0.010858546)
Z <- c(1.32, 0.81, 1.49, 1, 1.12, 2.71, 3.02, 3.9, 2.73)

card.model1IV = ivmodel(Y=Y,D=D,Z=Z)
card.model1IV

coefOther(card.model1IV)

card.modelFormula = ivmodelFormula(Y ~ D | Z)
card.modelFormula
```

R code used to generate Table S81

### R code
### Modified from code of Kang et al. paper “ivmodel: An R Package for Inference and Sensitivity Analysis of Instrumental Variables Models with One Endogenous Variable”

```r
# Prerequisite to this code is having the ivmodel package installed in R (and any package
dependencies thereof).
# Y is the dependent variable, D is the independent variable, and Z is the instrumental
variable. The X covariate variable in the original code has been removed.
# Instrumental variable (log10 adult body mass) analysis reports causality between lower log10
λ and higher log10 maximal lifespan.
# N.B. a two-tailed p-value output can be made one-tailed by dividing by 2.
# When copying and pasting this code you may inadvertently copy and paste a page number,
which must be removed before entering this code into the R console.
# You may need to press return after copying and pasting the code into the R console.

library(ivmodel)
Y <- c(1.278753601, 1.301029996, 1.45331834, 1.361727836, 1.431363764, 1.328379603,
1.311753861, 1.447158031, 1.419955748, 1.431363764, 1.31386722, 1.431363764,
1.096910013, 1.677606953, 1.309630167, 1.465382851, 0.954242509, 1.320146286,
1.359835482, 1.278753601, 1.348304863, 0.73239376, 0.698970004, 0.819543936,
1.178976947, 1.397940009, 1.201397124, 1.161368002, 1.068185862, 1.113943352,
0.84509804, 1.354108439, 0.748188027, 1.672097858, 1.357934847, 1.544068044,
1.301029996, 2.088136089, 1.900913068, 1.190331698, 1.491361694, 1.079181246,
1.235528447, 0.51851394, 0.591064607, 0.72427587, 1.021189299, 0.897627091,
0.919078092, 0.86332286, 0.995635195, 0.799340549, 0.602059991, 0.579783597,
0.62324929, 1.146128036, 0.897627091, 1.326335861, 1.146128036, 1.372912003,
1.305351369, 1.838849091, 0.397940009, 1.474216264, 1.357934847, 1.425289616,
1.477121255, 1.755874856, 1.596597096, 1.045322979, 1.369215857, 1.977723605,
1.571708832, 1.602059991, 1.431363764, 1.318063335, 1.712649702, 1.602059991,
1.69897000)
D <- c(1.515863084, 0.968353456, 0.980403656, 1.207186082, 1.007408842, 1.240480176,
1.199851427, 1.031373886, 1.364855052, 1.098452798, 1.236351357, 1.211234765,
1.698579647, 0.970402117, 1.117760113, 1.471877955, 1.673367431, 1.39741027,
1.400988909, 1.598662046, 1.15423023, 1.771300307, 1.874223695, 1.173913576,
0.992067042, 0.769926424, 1.106749129, 1.208121037, 1.369506183, 1.303390977,
1.725934804, 1.119280998, 1.241479123, 0.879450555, 1.388634544, 0.869392662,
1.316238787, 0.771773513, 0.623957491, 1.37983839, 1.226094696, 1.578522323,
1.398293766, 1.604022674, 1.90122943, 1.766139393, 1.368099338, 1.736446004,
```

```
1.759692602, 1.670458888, 1.623478846, 1.831536505, 1.958572748, 1.828539863,
1.756278312, 1.485206601, 1.591957358, 1.22741882, 1.06232791, 1.419894529,
1.513726937, 0.49911155, 1.961874192, 0.98867858, 1.090216566, 1.196235036,
1.246053737, 0.791560023, 0.868260697, 1.634894363, 1.194113108, 1.15277103,
0.843464988, 1.031807585, 1.068177474, 1.151071549, 0.699969722, 0.771098345,
0.774350899)
Z <- c(-0.744727495, 2.875061263, 2.63748973, 1.939519253, 2.113943352, 0.61612878,
1.728353782, 2.243038049, 2.07809415, 1.799340549, -0.110138279, 1.412460547, -
0.957424488, 2.06069784, 1.720159303, -1.478861916, -1.36251027, -0.059483515, -
0.903089987, -1.638272164, 0.740362689, -1.560667306, -1.823908741, 0.477121255,
0.812913357, 1.740362689, 0.379305518, 0.155336037, -0.124938737, 0.255272505, -1,
0.096910013, -0.066006836, 2.21747868, -0.59311933, 0.997823081, 0.113943352,
1.792636787, 3.502153893, -0.744727495, -1.455931956, -0.137868621, -0.192126868, -
1.180456064, -0.978810701, -1.397940009, -0.627087997, -1.638272164, -1.688246139, -
1.259637311, -1.244125144, -1.274088368, -1.688246139, -0.522878745, -0.698970004, -
0.628932138, -0.762707662, 0.544068044, 0.602059991, -0.273272791, -0.795880017,
2.507855872, -1.257274869, 1.77815125, 1.903089987, 1.31386722, 0.591064607,
2.477121255, 2.903089987, -0.092051478, 1.306425028, 4.477121255, 0.407390904,
0.915663604, 0.954242509, 1.785329835, 2.301029996, 3.155336037, 3.875061263)
card.model1IV = ivmodel(Y=Y,D=D,Z=Z)
card.model1IV
coefOther(card.model1IV)
card.modelFormula = ivmodelFormula(Y ~ D | Z)
card.modelFormula
```

<u>R code used to generate Table S87</u>

```
# R code
# Modified from code of Kang et al. paper “ivmodel: An R Package for Inference and
Sensitivity Analysis of Instrumental Variables Models with One Endogenous Variable”
# Prerequisite to this code is having the ivmodel package installed in R (and any package
dependencies thereof).
# Y is the dependent variable, D is the independent variable, and Z is the instrumental
variable. The X covariate variable in the original code has been removed.
```

### Instrumental variable (specific basal metabolic rate) analysis reports causality between lower ΔD and longer maximal lifespan.
### Human data excluded by rationale given in Methods section.
### N.B. a two-tailed p-value output can be made one-tailed by dividing by 2.
### When copying and pasting this code you may inadvertently copy and paste a page number, which must be removed before entering this code into the R console.
### You may need to press return after copying and pasting the code into the R console.

```
library(ivmodel)

Y <- c(20, 27, 28, 26.3, 13, 35, 31, 12, 17.2, 3.9, 4, 3.8, 20.2, 22.8, 26.625, 30)
D <- c(15.59559661, 17.87501179, 14.96911115, 28.99329722, 23.39744412, 7.896615237,
16.50265574, 48.07412358, 32.02595596, 99.14473303, 98.15119314, 75.95983356,
38.34156448, 24.13779014, 24.45232192, 19.41510748)
Z <- c(0.000884063, 0.000771481, 0.000965102, 0.000970696, 0.003411135, 0.001603694,
0.003626062, 0.003332812, 0.002999771, 0.006377079, 0.015055556, 0.006785887,
0.004372197, 0.001319925, 0.00180432, 0.00245358)

card.model1IV = ivmodel(Y=Y,D=D,Z=Z)
card.model1IV

coefOther(card.model1IV)

card.modelFormula = ivmodelFormula(Y ~ D | Z)
card.modelFormula
```

<u>Confirmatory check: R code that replicates the β coefficient value (and standard error and p-value thereof) of Table 9 using a different R package</u>

### R code
### Prerequisite to this code is having the AER package installed in R (and any package dependencies thereof). AER stands for “Applied Econometrics with R”.
### AER package described in Kleiber C, Zeileis A (2008) Applied Econometrics with R. Springer-Verlag, New York.

```
# Y is the dependent variable, D is the independent variable, and Z is the instrumental variable.
# Instrumental variable (specific F1F0 ATP hydrolysis rate) analysis reports causality between lower specific basal metabolic rate and longer maximal lifespan.
# Data is from Table S2 of this Supplementary Material. Human data excluded by rationale given in Methods section.
# N.B. a two-tailed p-value output can be made one-tailed by dividing by 2.
# When copying and pasting this code you may inadvertently copy and paste a page number, which must be removed before entering this code into the R console.
# You may need to press return after copying and pasting the code into the R console.

library(AER)

# Define variables
Y <- c(20, 27, 22.8, 24, 9, 17.5, 17.7, 12, 3.8, 3.9, 4)
D <- c(0.000798458, 0.000771, 0.001319925, 0.001804232, 0.002887816, 0.002418352, 0.004975053, 0.003740752, 0.00601463, 0.006377, 0.010858546)
Z <- c(1.32, 0.81, 1.49, 1, 1.12, 2.89, 2.66, 2.71, 3.02, 3.9, 2.73)

# Specify instrumental variable regression model
iv_model <- ivreg(Y ~ D | Z)

# Diagnostic tests
summary(iv_model)

# Coefficients
coef(iv_model)

# Standard errors
sqrt(diag(vcov(iv_model)))
```

<u>Confirmatory check: R code that replicates the β coefficient value (and standard error and p-value thereof) of Table 10 using a different R package</u>

```
# R code
```

### Prerequisite to this code is having the AER package installed in R (and any package dependencies thereof). AER stands for "Applied Econometrics with R".
### AER package described in Kleiber C, Zeileis A (2008) Applied Econometrics with R. Springer-Verlag, New York.
### Y is the dependent variable, D is the independent variable, and Z is the instrumental variable.
### Instrumental variable (specific IF1 protein activity) analysis reports causality between lower specific F1F0 ATP hydrolysis rate and longer maximal lifespan.
### Maximal lifespan data from [1] and other data is from [91]. Human data excluded by rationale given in Methods section.
### N.B. a two-tailed p-value output can be made one-tailed by dividing by 2.
### When copying and pasting this code you may inadvertently copy and paste a page number, which must be removed before entering this code into the R console.
### You may need to press return after copying and pasting the code into the R console.

```
library(AER)

# Define variables
Y <- c(20, 27, 22.8, 24, 9, 17.5, 17.7, 12, 3.8, 3.9, 4)
D <- c(1.32, 0.81, 1.49, 1, 1.12, 2.89, 2.66, 2.71, 3.02, 3.9, 2.73)
Z <- c(3.49, 3.32, 3.34, 3.9, 5.06, 4.88, 4.44, 3.45, 0.65, 0.72, 0.81)

# Specify instrumental variable regression model
iv_model <- ivreg(Y ~ D | Z)

# Diagnostic tests
summary(iv_model)

# Coefficients
coef(iv_model)

# Standard errors
sqrt(diag(vcov(iv_model)))
```

Confirmatory check: R code that replicates the β coefficient value (and standard error and p-value thereof) of Table 21 using a different R package

```
# R code
# Prerequisite to this code is having the AER package installed in R (and any package dependencies thereof). AER stands for "Applied Econometrics with R".
# AER package described in Kleiber C, Zeileis A (2008) Applied Econometrics with R. Springer-Verlag, New York.
# Y is the dependent variable, D is the independent variable, and Z is the instrumental variable.
# Instrumental variable (specific F1F0 ATP hydrolysis rate) analysis reports causality between specific basal metabolic rate and the ticking rate of the epigenetic clock (ΔM).
# Data is from Table S2 of this Supplementary Material, with ΔM calculated by Eq. 5 from maximal lifespan values in [1]. Human data excluded by rationale given in Methods section.
# N.B. a two-tailed p-value output can be made one-tailed by dividing by 2.
# When copying and pasting this code you may inadvertently copy and paste a page number, which must be removed before entering this code into the R console.
# You may need to press return after copying and pasting the code into the R console.

library(AER)

# Define variables
Y <- c(0.154552123, 0.12471466, 0.140826476, 0.135754022, 0.266432808, 0.219921444, 0.462594189, 0.455115987, 0.447939867)
D <- c(0.000798458, 0.000771, 0.001319925, 0.001804232, 0.002887816, 0.003740752, 0.00601463, 0.006377, 0.010858546)
Z <- c(1.32, 0.81, 1.49, 1, 1.12, 2.71, 3.02, 3.9, 2.73)

# Specify instrumental variable regression model
iv_model <- ivreg(Y ~ D | Z)

# Diagnostic tests
summary(iv_model)

# Coefficients
```

```
coef(iv_model)

# Standard errors
sqrt(diag(vcov(iv_model)))
```

Confirmatory check: R code that replicates the β coefficient value (and standard error and p-value thereof) of Table S81 using a different R package

```
# R code
# Prerequisite to this code is having the AER package installed in R (and any package dependencies thereof). AER stands for “Applied Econometrics with R”.
# AER package described in Kleiber C, Zeileis A (2008) Applied Econometrics with R. Springer-Verlag, New York.
# Y is the dependent variable, D is the independent variable, and Z is the instrumental variable.
# Instrumental variable (log10 adult body mass) analysis reports causality between lower log10 λ and higher log10 maximal lifespan.
# N.B. a two-tailed p-value output can be made one-tailed by dividing by 2.
# When copying and pasting this code you may inadvertently copy and paste a page number, which must be removed before entering this code into the R console.
# You may need to press return after copying and pasting the code into the R console.

library(AER)

# Define variables
Y <- c(1.278753601, 1.301029996, 1.45331834, 1.361727836, 1.431363764, 1.328379603,
1.311753861, 1.447158031, 1.419955748, 1.431363764, 1.31386722, 1.431363764,
1.096910013, 1.677606953, 1.309630167, 1.465382851, 0.954242509, 1.320146286,
1.359835482, 1.278753601, 1.348304863, 0.73239376, 0.698970004, 0.819543936,
1.178976947, 1.397940009, 1.201397124, 1.161368002, 1.068185862, 1.113943352,
0.84509804, 1.354108439, 0.748188027, 1.672097858, 1.357934847, 1.544068044,
1.301029996, 2.088136089, 1.900913068, 1.190331698, 1.491361694, 1.079181246,
1.235528447, 0.51851394, 0.591064607, 0.72427587, 1.021189299, 0.897627091,
0.919078092, 0.86332286, 0.995635195, 0.799340549, 0.602059991, 0.579783597,
0.62324929, 1.146128036, 0.897627091, 1.326335861, 1.146128036, 1.372912003,
```

```
1.305351369, 1.838849091, 0.397940009, 1.474216264, 1.357934847, 1.425289616,
1.477121255, 1.755874856, 1.596597096, 1.045322979, 1.369215857, 1.977723605,
1.571708832, 1.602059991, 1.431363764, 1.318063335, 1.712649702, 1.602059991,
1.69897000)
D <- c(1.515863084, 0.968353456, 0.980403656, 1.207186082, 1.007408842, 1.240480176,
1.199851427, 1.031373886, 1.364855052, 1.098452798, 1.236351357, 1.211234765,
1.698579647, 0.970402117, 1.117760113, 1.471877955, 1.673367431, 1.39741027,
1.400988909, 1.598662046, 1.15423023, 1.771300307, 1.874223695, 1.173913576,
0.992067042, 0.769926424, 1.106749129, 1.208121037, 1.369506183, 1.303390977,
1.725934804, 1.119280998, 1.241479123, 0.879450555, 1.388634544, 0.869392662,
1.316238787, 0.771773513, 0.623957491, 1.37983839, 1.226094696, 1.578522323,
1.398293766, 1.604022674, 1.90122943, 1.766139393, 1.368099338, 1.736446004,
1.759692602, 1.670458888, 1.623478846, 1.831536505, 1.958572748, 1.828539863,
1.756278312, 1.485206601, 1.591957358, 1.22741882, 1.06232791, 1.419894529,
1.513726937, 0.49911155, 1.961874192, 0.98867858, 1.090216566, 1.196235036,
1.246053737, 0.791560023, 0.868260697, 1.634894363, 1.194113108, 1.15277103,
0.843464988, 1.031807585, 1.068177474, 1.151071549, 0.699969722, 0.771098345,
0.774350899)
Z <- c(-0.744727495, 2.875061263, 2.63748973, 1.939519253, 2.113943352, 0.61612878,
1.728353782, 2.243038049, 2.07809415, 1.799340549, -0.110138279, 1.412460547, -
0.957424488, 2.06069784, 1.720159303, -1.478861916, -1.36251027, -0.059483515, -
0.903089987, -1.638272164, 0.740362689, -1.560667306, -1.823908741, 0.477121255,
0.812913357, 1.740362689, 0.379305518, 0.155336037, -0.124938737, 0.255272505, -1,
0.096910013, -0.066006836, 2.21747868, -0.59311933, 0.997823081, 0.113943352,
1.792636787, 3.502153893, -0.744727495, -1.455931956, -0.137868621, -0.192126868, -
1.180456064, -0.978810701, -1.397940009, -0.627087997, -1.638272164, -1.688246139, -
1.259637311, -1.244125144, -1.274088368, -1.688246139, -0.522878745, -0.698970004, -
0.628932138, -0.762707662, 0.544068044, 0.602059991, -0.273272791, -0.795880017,
2.507855872, -1.257274869, 1.77815125, 1.903089987, 1.31386722, 0.591064607,
2.477121255, 2.903089987, -0.092051478, 1.306425028, 4.477121255, 0.407390904,
0.915663604, 0.954242509, 1.785329835, 2.301029996, 3.155336037, 3.875061263)

# Specify instrumental variable regression model
iv_model <- ivreg(Y ~ D | Z)
```

### Diagnostic tests
summary(iv_model)

### Coefficients
coef(iv_model)

### Standard errors
sqrt(diag(vcov(iv_model)))

<u>Confirmatory check: R code that replicates the β coefficient value (and standard error and p-value thereof) of Table S87 using a different R package</u>

### R code
### Prerequisite to this code is having the AER package installed in R (and any package dependencies thereof). AER stands for "Applied Econometrics with R".
### AER package described in Kleiber C, Zeileis A (2008) Applied Econometrics with R. Springer-Verlag, New York.
### Y is the dependent variable, D is the independent variable, and Z is the instrumental variable.
### Instrumental variable (specific basal metabolic rate) analysis reports causality between lower ΔD and longer maximal lifespan.
### Human data excluded by rationale given in Methods section.
### N.B. a two-tailed p-value output can be made one-tailed by dividing by 2.
### When copying and pasting this code you may inadvertently copy and paste a page number, which must be removed before entering this code into the R console.
### You may need to press return after copying and pasting the code into the R console.

library(AER)

### Define variables
Y <- c(20, 27, 28, 26.3, 13, 35, 31, 12, 17.2, 3.9, 4, 3.8, 20.2, 22.8, 26.625, 30)
D <- c(15.59559661, 17.87501179, 14.96911115, 28.99329722, 23.39744412, 7.896615237, 16.50265574, 48.07412358, 32.02595596, 99.14473303, 98.15119314, 75.95983356, 38.34156448, 24.13779014, 24.45232192, 19.41510748)

```
Z <- c(0.000884063, 0.000771481, 0.000965102, 0.000970696, 0.003411135, 0.001603694,
0.003626062, 0.003332812, 0.002999771, 0.006377079, 0.015055556, 0.006785887,
0.004372197, 0.001319925, 0.00180432, 0.00245358)

# Specify instrumental variable regression model
iv_model <- ivreg(Y ~ D | Z)

# Diagnostic tests
summary(iv_model)

# Coefficients
coef(iv_model)

# Standard errors
sqrt(diag(vcov(iv_model)))
```

R code used to generate Generalized Additive Model (GAM)

```
# Need to have the mgcv package installed (and any package dependencies thereof).
# When copying and pasting this code you may inadvertently copy and paste a page number,
which must be removed before entering this code into the R console.
# You may need to press return after copying and pasting the code into the R console.
# Be aware that the R console has a maximum width for R code, which the data variables for
species sets A and B exceeds, and so their data must be broken up with at least one line break
in each variable (if this isn't done, when trying to run the code you will get an error message,
which doesn't explicitly specify that this width issue is the reason for the error, but it is, and
when solved the R code runs fine). Code below is presented with data for species set C,
which is a smaller species set, and doesn't face this width problem.

library(mgcv)

# Data for species set C: 263 mammals (without bats, naked mole rat, and marsupials)
x <- c(0.006443299, 0.003264605, 0.00706278, 0.005522727, 0.001340037, 0.001121538,
0.001465613, 0.000884063, 0.001170855, 0.00148949, 0.002789277, 0.001450438,
```

0.001710766, 0.00239881, 0.002147813, 0.001345146, 0.000552283, 0.00129513,
0.002155674, 0.00167806, 0.001995506, 0.001407765, 0.000771481, 0.001617805,
0.003022594, 0.00085331, 0.001913803, 0.002789249, 0.001561719, 0.003772571,
0.002129167, 0.002785755, 0.00279706, 0.002997839, 0.002323154, 0.002222398,
0.001606665, 0.001654095, 0.001451944, 0.001422925, 0.002494255, 0.000965102,
0.001238472, 0.000970696, 0.001325968, 0.001153571, 0.004237875, 0.003679214,
0.002034118, 0.000931603, 0.001463189, 0.002682692, 0.002308814, 0.003670098,
0.002501181, 0.0025104, 0.003703052, 0.004347826, 0.001506516, 0.010167331,
0.005973333, 0.004124242, 0.001673556, 0.001303279, 0.002674818, 0.002763281,
0.001834605, 0.001424713, 0.001852459, 0.002231034, 0.002153567, 0.000702899,
0.000892647, 0.001534826, 0.002465568, 0.001685859, 0.00377807, 0.001537838,
0.000992952, 0.001363623, 0.000842613, 0.0003738, 0.001010345, 0.00125977, 0.00184,
0.002354443, 0.002005768, 0.002120452, 0.001395349, 0.00201546, 0.00318, 0.002800796,
0.003411135, 0.003648275, 0.008550459, 0.006236559, 0.005877551, 0.005981132,
0.007487179, 0.001874211, 0.000799931, 0.000547997, 0.000929127, 0.000971487,
0.000475261, 0.001319191, 0.001432857, 0.002732641, 0.002454818, 0.004463158,
0.005411021, 0.0058, 0.005293415, 0.002228774, 0.001603694, 0.001986, 0.001664851,
0.003626667, 0.001676471, 0.00445481, 0.005961538, 0.002611978, 0.002988584,
0.001182571, 0.001115821, 0.001785519, 0.003548544, 0.002514085, 0.001332624,
0.002004956, 0.00380531, 0.000636302, 0.003183549, 0.004435995, 0.003300518,
0.00128327, 0.001673453, 0.003332812, 0.004574534, 0.001392988, 0.002510612,
0.002999771, 0.001563586, 0.006473073, 0.013194444, 0.003424583, 0.008080986,
0.010646766, 0.0133875, 0.006377079, 0.01468, 0.0156621, 0.010205479, 0.008506224,
0.012298387, 0.014511628, 0.004138514, 0.004088889, 0.00774359, 0.008638743,
0.005680851, 0.007902439, 0.008238095, 0.00955157, 0.010682927, 0.01, 0.009274924,
0.01, 0.009350649, 0.005438374, 0.006798246, 0.001673547, 0.002733437, 0.003235579,
0.003067834, 0.006866667, 0.00557554, 0.008390805, 0.003514056, 0.003569659,
0.00156189, 0.001990249, 0.003903553, 0.004685315, 0.003807339, 0.005859375,
0.01493617, 0.007653061, 0.006821192, 0.004886578, 0.006445498, 0.006542553,
0.00709205, 0.007176471, 0.008333333, 0.001165929, 0.006142857, 0.012897527,
0.011092437, 0.004258912, 0.006815021, 0.009539906, 0.007979798, 0.004343434,
0.004635879, 0.004184332, 0.0033, 0.008425926, 0.004664537, 0.026447368, 0.016125,
0.015055556, 0.007801858, 0.006197368, 0.003238532, 0.006785887, 0.006581197,
0.004520202, 0.004308176, 0.004856157, 0.003571471, 0.007537313, 0.004023495,

0.004799599, 0.004445325, 0.001924783, 0.005430393, 0.003874199, 0.002119932, 0.009206349, 0.006142433, 0.003234783, 0.004954637, 0.002008382, 0.001394717, 0.003607908, 0.004686364, 0.0053, 0.007079108, 0.009073661, 0.007354281, 0.002620833, 0.004004796, 0.003358562, 0.002790698, 0.004372197, 0.004243902, 0.004796623, 0.000328264, 0.000155568, 0.01645933, 0.00746988, 0.014188034, 0.013240741, 0.016231884, 0.026031746, 0.0205, 0.041428571, 0.02625, 0.010151134, 0.01255102, 0.007875, 0.000714063)

z <- c(0.18, 0.18, 0.0378, 0.05, 0.225, 0.9, 46.1, 750, 164.5, 175.333, 5, 90, 70.275, 6.25, 13, 500, 434, 140, 21.6667, 200, 87, 101.25, 130, 20.2, 2.85, 4.325, 13.25, 10.25, 6.5, 26.5, 5.2, 2.5, 2.4, 4.1317, 1.25, 1.54, 53.5, 8.8, 3.6667, 13.35, 8.6, 175, 81.15, 119.7, 63, 7, 0.5, 0.8, 0.776, 40, 10, 0.6, 4.5, 25.85, 16.3333, 6.75, 0.884, 1.3, 13, 0.1103, 0.1506, 0.925, 8, 1.9, 115, 0.9, 3.75, 4.75, 3, 6.27, 6, 100, 12.25, 2.25, 1.82, 3.1, 52.5, 0.84, 3, 5.5, 4.85, 33, 1.5, 1.5, 0.5, 0.719, 0.75, 0.342, 0.188, 3.6, 4.175, 4.175, 1.8, 0.9, 0.1, 0.052, 0.048, 0.058, 0.04, 1.25, 3.5, 7.5, 164.998, 6.25, 28.5, 4.5, 4.5, 0.8, 7, 0.2552, 0.124, 0.4993, 0.925, 9, 9.95, 7.75, 18, 0.38, 0.1692, 0.1922, 0.06, 1.0945, 1.3, 62.035, 5, 2.485, 0.2575, 0.2382, 0.8905, 0.9199, 0.1192, 3178, 0.18, 0.1325, 0.181, 4.6833, 1.5, 0.728, 0.3379, 55, 0.95, 0.6425, 6, 0.12, 0.008, 0.5067, 0.066, 0.04, 0.07, 0.105, 0.046, 0.0275, 0.0516, 0.04, 0.0255, 0.0208, 0.209, 0.236, 0.022, 0.022, 0.042, 0.0165, 0.025, 0.023, 0.0205, 0.014, 0.025, 0.0234, 0.032, 0.0722, 0.185, 9, 2.65, 3.265, 0.775, 0.055, 0.134, 0.018, 0.5, 0.3735, 3.9, 8.6, 0.1767, 0.1155, 0.125, 0.024, 0.0273, 0.0187, 0.019, 0.116, 0.065, 0.042, 0.057, 0.008, 0.0201, 24, 0.0452, 0.0294, 0.0234, 0.144, 0.0545, 0.0748, 0.03, 0.0185, 0.05, 0.037, 0.85, 0.0532, 0.0476, 0.006, 0.0072, 0.0205, 0.035, 0.13, 0.115, 0.3, 0.2, 0.051, 0.065, 0.1467, 1.5, 0.106, 0.0474, 0.235, 0.098, 3.5, 0.105, 0.1576, 1.125, 0.218, 0.0654, 0.3265, 0.1727, 3.5, 4, 0.459, 0.533, 0.217, 0.0441, 0.096, 0.2, 0.15, 0.1633, 0.588, 0.65, 0.16, 0.2, 0.35, 480, 322, 0.0216, 0.045, 0.011, 0.0116, 0.0077, 0.0045, 0.015, 0.009, 0.0021, 0.0459, 0.0553, 0.09, 60)

y <- c(19, 2.7, 5.6, 5.8, 14.1, 8.7, 17, 20, 24.3, 30, 17.3, 20.8, 20.6, 15.9, 9.3, 26.1, 28.4, 28.9, 17.5, 31.5, 23, 21.8, 27, 31.5, 14, 19, 21.8, 16.7, 12.7, 17, 16.3, 15.8, 15.9, 21.3, 16.3, 21.4, 20.5, 28.2, 24, 22.4, 32.3, 28, 28, 26.3, 27, 18.6, 12.6, 16.6, 20.6, 25, 20, 10.5, 22.3, 27, 19.5, 18.2, 17.8, 18.2, 18.6, 12.5, 8.8, 11.4, 25, 21, 47.6, 24, 26.4, 23.7, 38.4, 19, 21, 33.3, 27, 16.1, 21.3, 24.5, 20.4, 16.2, 25.2, 22.3, 22.1, 15, 36.8, 12.5, 11.4, 7, 11.7, 7.6, 6.3, 14.8, 11.8, 18, 13, 7.8, 7, 9.3, 7.2, 7.9, 8.7, 22.6, 49.5, 41.2, 47, 41, 31, 16, 19, 30.1, 24, 22.8, 18.6, 20.5, 30.2, 37.8, 35, 30, 37.5, 29, 16.6, 17.1, 13.4, 22.7, 20, 122.5, 31, 35.5, 13, 21.6, 25.8, 32.4, 16, 79.6, 15.5, 11, 11.2, 13.8, 13.5, 12, 3.5, 15.1, 12.9, 17.2, 10.8, 2.5, 3.3, 3.6, 3.3, 4.1, 3.3, 3.9, 4.8, 4.8, 3.9, 5.3, 3.8, 4.9, 9.5, 10.5, 8.4, 4.6, 5.5, 7.7, 7.4, 7.9, 8.3, 5.5, 5.4, 3.9, 7.4, 5.5,

5.2, 16.3, 11.8, 17.8, 14.8, 7.3, 5.5, 5.6, 5.8, 7, 26.6, 23.4, 12, 6.1, 8.7, 6.9, 5.3, 8.3, 7.1, 8.6, 8.3, 9.7, 9.9, 8.3, 5.8, 23.1, 5.9, 4.5, 6.3, 4.5, 6.1, 5.7, 5.9, 4.5, 5.5, 8.2, 7.3, 6.3, 4.7, 3.8, 4.3, 4, 6.4, 5.3, 4.4, 3.8, 4.2, 4.5, 7.3, 7, 8.4, 6.3, 4.5, 14, 10.8, 20, 11.2, 10.4, 15, 10, 19, 8.7, 7.9, 21.2, 14, 7.2, 23.6, 6.7, 10, 9.5, 9.8, 7.8, 8.9, 11.5, 4.8, 20.2, 12.4, 11.5, 40, 69, 2.2, 2.2, 2.9, 4, 2.8, 3.5, 3.1, 3.2, 3.2, 3.2, 2.5, 6.2, 29.8)

```
# Create a data frame with the variables
data <- data.frame(x = x, z = z, y = y)

# Fit a GAM model
gam_model <- gam(y ~ s(x) + s(z), data = data)

# Summary of the GAM model
summary(gam_model)
```

<u>R code to run some diagnostics for GAM model</u>

```
gam.check(gam_model)
concurvity(gam_model, full = TRUE)
concurvity(gam_model, full = FALSE)
```

<u>R code to generate Q-Q plot for GAM model (*should be linear*)</u>

```
# Extract residuals from the fitted model
residuals <- residuals(gam_model)

# Create a Q-Q plot of the residuals
qqnorm(residuals)
qqline(residuals)
```

<u>R code to plot residuals vs. predicted values (of the response variable) of a GAM model (*residuals should be randomly distributed around zero with no discernible pattern, and there should be a spread of residuals across the range of predicted values*)</u>

```
# Extract predicted values and residuals
predicted_values <- predict(gam_model)
residuals <- residuals(gam_model)
```

```
# Plot residuals versus predicted values
plot(predicted_values, residuals, xlab = "Predicted Values", ylab = "Residuals",
     main = "Residuals vs Predicted Values Plot", pch = 20)
abline(h = 0, col = "red", lty = 2)  # Add a horizontal line at y = 0
```

<u>R code to plot working responses vs. linear predictor plot for a GAM model (*should be linear*)</u>

```
# Extract working responses and linear predictor
working_responses <- predict(gam_model, type = "response")
linear_predictor <- predict(gam_model, type = "link")

# Create scatter plot
plot(linear_predictor, working_responses,
     xlab = "Linear Predictor", ylab = "Working Responses",
     main = "Working Responses vs. Linear Predictor",
     col = "blue", pch = 16)
abline(0, 1, col = "red")  # Add a reference line for linearity
```

<u>R code to plot partial residuals vs. *x* predictor plot for a GAM model (*ideally should appear random and evenly distributed around zero*)</u>

```
# Extract residuals and predictor variable
residuals <- residuals(gam_model)
predictor_variable <- x

# Calculate partial residuals
partial_residuals <- residuals + predict(gam_model, type = "terms")[, "s(x)"]  # Replace 's(x)'
with the term you want to plot

# Plot partial residuals against predictor variable
plot(predictor_variable, partial_residuals,
     xlab = "Predictor Variable", ylab = "Partial Residuals",
     main = "Partial Residual Plot",
     col = "blue", pch = 16)
```

R code to plot partial residuals vs. *z* predictor plot for a GAM model (*ideally should appear random and evenly distributed around zero*)

```
# Extract residuals and predictor variable
residuals <- residuals(gam_model)
predictor_variable <- z

# Calculate partial residuals
partial_residuals <- residuals + predict(gam_model, type = "terms")[, "s(z)"]

# Plot partial residuals against predictor variable
plot(predictor_variable, partial_residuals,
    xlab = "Predictor Variable", ylab = "Partial Residuals",
    main = "Partial Residual Plot",
    col = "blue", pch = 16)
```

R code to plot a Component-Plus-Residual (CPR) plot for a GAM (*ideally, the points should be scattered randomly around the diagonal line. It should exhibit randomness around the diagonal line*)

```
# Predict smoothed components and residuals
smoothed_components <- predict(gam_model, type = "terms", se.fit = FALSE)
residuals <- residuals(gam_model, type = "response")

# Create the CPR plot
plot(smoothed_components[, "s(x)"] + residuals, smoothed_components[, "s(z)"] + residuals,
    xlab = "Smoothed Component + Residual (s(x))", ylab = "Smoothed Component + 
Residual (s(z))",
    main = "Component-Plus-Residual (CPR) Plot")
```

R code to plot model predicted *y* vs. actual *y* (e.g., model-predicted maximal lifespan vs. actual maximal lifespan)

```
# Obtain predicted values
predicted_values <- predict(gam_model, type = "response")
```

```
# Plot predicted vs. actual
plot(y, predicted_values,
     xlab = "Actual y", ylab = "Predicted y",
     main = "Predicted vs. Actual")
abline(a = 0, b = 1, col = "red")  # Add a 45-degree line for reference
```

<u>R code used to generate Eq. S6</u>

### The nls2 function with the “brute-force” algorithm will systematically try all combinations of starting values within the specified grid to find the best fit for the model parameters. This can be computationally intensive, especially for larger grids or more complex models.
### Need to have the nls2 package installed (and any package dependencies thereof).
### When copying and pasting this code you may inadvertently copy and paste a page number, which must be removed before entering this code into the R console.
### You may need to press return after copying and pasting the code into the R console.
### Be aware that the R console has a maximum width for R code, which the data variables for species sets *A* and *B* exceeds, and so their data must be broken up with at least one line break in each variable (if this isn’t done, when trying to run the code you will get an error message, which doesn’t explicitly specify that this width issue is the reason for the error, but it is, and when solved the R code runs fine). Code below is presented with data for species set *C*, which is a smaller species set, and doesn’t face this width problem.

```
# Load the nls2 package
library(nls2)

# Define the equation
custom_function <- function(x, z, B, C, D, E) {
  (B * x^C) + (D * z^E)
}

# Define the variables
# Data for species set C: 263 mammals (without bats, naked mole rat, and marsupials)
x <- c(0.006443299, 0.003264605, 0.00706278, 0.005522727, 0.001340037, 0.001121538,
0.001465613, 0.000884063, 0.001170855, 0.00148949, 0.002789277, 0.001450438,
0.001710766, 0.00239881, 0.002147813, 0.001345146, 0.000552283, 0.00129513,
```

0.002155674, 0.00167806, 0.001995506, 0.001407765, 0.000771481, 0.001617805, 0.003022594, 0.00085331, 0.001913803, 0.002789249, 0.001561719, 0.003772571, 0.002129167, 0.002785755, 0.00279706, 0.002997839, 0.002323154, 0.002222398, 0.001606665, 0.001654095, 0.001451944, 0.001422925, 0.002494255, 0.000965102, 0.001238472, 0.000970696, 0.001325968, 0.001153571, 0.004237875, 0.003679214, 0.002034118, 0.000931603, 0.001463189, 0.002682692, 0.002308814, 0.003670098, 0.002501181, 0.0025104, 0.003703052, 0.004347826, 0.001506516, 0.010167331, 0.005973333, 0.004124242, 0.001673556, 0.001303279, 0.002674818, 0.002763281, 0.001834605, 0.001424713, 0.001852459, 0.002231034, 0.002153567, 0.000702899, 0.000892647, 0.001534826, 0.002465568, 0.001685859, 0.00377807, 0.001537838, 0.000992952, 0.001363623, 0.000842613, 0.0003738, 0.001010345, 0.00125977, 0.00184, 0.002354443, 0.002005768, 0.002120452, 0.001395349, 0.00201546, 0.00318, 0.002800796, 0.003411135, 0.003648275, 0.008550459, 0.006236559, 0.005877551, 0.005981132, 0.007487179, 0.001874211, 0.000799931, 0.000547997, 0.000929127, 0.000971487, 0.000475261, 0.001319191, 0.001432857, 0.002732641, 0.002454818, 0.004463158, 0.005411021, 0.0058, 0.005293415, 0.002228774, 0.001603694, 0.001986, 0.001664851, 0.003626667, 0.001676471, 0.00445481, 0.005961538, 0.002611978, 0.002988584, 0.001182571, 0.001115821, 0.001785519, 0.003548544, 0.002514085, 0.001332624, 0.002004956, 0.00380531, 0.000636302, 0.003183549, 0.004435995, 0.003300518, 0.00128327, 0.001673453, 0.003332812, 0.004574534, 0.001392988, 0.002510612, 0.002999771, 0.001563586, 0.006473073, 0.013194444, 0.003424583, 0.008080986, 0.010646766, 0.0133875, 0.006377079, 0.01468, 0.0156621, 0.010205479, 0.008506224, 0.012298387, 0.014511628, 0.004138514, 0.004088889, 0.00774359, 0.008638743, 0.005680851, 0.007902439, 0.008238095, 0.00955157, 0.010682927, 0.01, 0.009274924, 0.01, 0.009350649, 0.005438374, 0.006798246, 0.001673547, 0.002733437, 0.003235579, 0.003067834, 0.006866667, 0.00557554, 0.008390805, 0.003514056, 0.003569659, 0.00156189, 0.001990249, 0.003903553, 0.004685315, 0.003807339, 0.005859375, 0.01493617, 0.007653061, 0.006821192, 0.004886578, 0.006445498, 0.006542553, 0.00709205, 0.007176471, 0.008333333, 0.001165929, 0.006142857, 0.012897527, 0.011092437, 0.004258912, 0.006815021, 0.009539906, 0.007979798, 0.004343434, 0.004635879, 0.004184332, 0.0033, 0.008425926, 0.004664537, 0.026447368, 0.016125, 0.015055556, 0.007801858, 0.006197368, 0.003238532, 0.006785887, 0.006581197, 0.004520202, 0.004308176, 0.004856157, 0.003571471, 0.007537313, 0.004023495, 0.004799599, 0.004445325, 0.001924783, 0.005430393, 0.003874199, 0.002119932,

0.009206349, 0.006142433, 0.003234783, 0.004954637, 0.002008382, 0.001394717, 0.003607908, 0.004686364, 0.0053, 0.007079108, 0.009073661, 0.007354281, 0.002620833, 0.004004796, 0.003358562, 0.002790698, 0.004372197, 0.004243902, 0.004796623, 0.000328264, 0.000155568, 0.01645933, 0.00746988, 0.014188034, 0.013240741, 0.016231884, 0.026031746, 0.0205, 0.041428571, 0.02625, 0.010151134, 0.01255102, 0.007875, 0.000714063)

z <- c(0.18, 0.18, 0.0378, 0.05, 0.225, 0.9, 46.1, 750, 164.5, 175.333, 5, 90, 70.275, 6.25, 13, 500, 434, 140, 21.6667, 200, 87, 101.25, 130, 20.2, 2.85, 4.325, 13.25, 10.25, 6.5, 26.5, 5.2, 2.5, 2.4, 4.1317, 1.25, 1.54, 53.5, 8.8, 3.6667, 13.35, 8.6, 175, 81.15, 119.7, 63, 7, 0.5, 0.8, 0.776, 40, 10, 0.6, 4.5, 25.85, 16.3333, 6.75, 0.884, 1.3, 13, 0.1103, 0.1506, 0.925, 8, 1.9, 115, 0.9, 3.75, 4.75, 3, 6.27, 6, 100, 12.25, 2.25, 1.82, 3.1, 52.5, 0.84, 3, 5.5, 4.85, 33, 1.5, 1.5, 0.5, 0.719, 0.75, 0.342, 0.188, 3.6, 4.175, 4.175, 1.8, 0.9, 0.1, 0.052, 0.048, 0.058, 0.04, 1.25, 3.5, 7.5, 164.998, 6.25, 28.5, 4.5, 4.5, 0.8, 7, 0.2552, 0.124, 0.4993, 0.925, 9, 9.95, 7.75, 18, 0.38, 0.1692, 0.1922, 0.06, 1.0945, 1.3, 62.035, 5, 2.485, 0.2575, 0.2382, 0.8905, 0.9199, 0.1192, 3178, 0.18, 0.1325, 0.181, 4.6833, 1.5, 0.728, 0.3379, 55, 0.95, 0.6425, 6, 0.12, 0.008, 0.5067, 0.066, 0.04, 0.07, 0.105, 0.046, 0.0275, 0.0516, 0.04, 0.0255, 0.0208, 0.209, 0.236, 0.022, 0.022, 0.042, 0.0165, 0.025, 0.023, 0.0205, 0.014, 0.025, 0.0234, 0.032, 0.0722, 0.185, 9, 2.65, 3.265, 0.775, 0.055, 0.134, 0.018, 0.5, 0.3735, 3.9, 8.6, 0.1767, 0.1155, 0.125, 0.024, 0.0273, 0.0187, 0.019, 0.116, 0.065, 0.042, 0.057, 0.008, 0.0201, 24, 0.0452, 0.0294, 0.0234, 0.144, 0.0545, 0.0748, 0.03, 0.0185, 0.05, 0.037, 0.85, 0.0532, 0.0476, 0.006, 0.0072, 0.0205, 0.035, 0.13, 0.115, 0.3, 0.2, 0.051, 0.065, 0.1467, 1.5, 0.106, 0.0474, 0.235, 0.098, 3.5, 0.105, 0.1576, 1.125, 0.218, 0.0654, 0.3265, 0.1727, 3.5, 4, 0.459, 0.533, 0.217, 0.0441, 0.096, 0.2, 0.15, 0.1633, 0.588, 0.65, 0.16, 0.2, 0.35, 480, 322, 0.0216, 0.045, 0.011, 0.0116, 0.0077, 0.0045, 0.015, 0.009, 0.0021, 0.0459, 0.0553, 0.09, 60)

y <- c(19, 2.7, 5.6, 5.8, 14.1, 8.7, 17, 20, 24.3, 30, 17.3, 20.8, 20.6, 15.9, 9.3, 26.1, 28.4, 28.9, 17.5, 31.5, 23, 21.8, 27, 31.5, 14, 19, 21.8, 16.7, 12.7, 17, 16.3, 15.8, 15.9, 21.3, 16.3, 21.4, 20.5, 28.2, 24, 22.4, 32.3, 28, 28, 26.3, 27, 18.6, 12.6, 16.6, 20.6, 25, 20, 10.5, 22.3, 27, 19.5, 18.2, 17.8, 18.2, 18.6, 12.5, 8.8, 11.4, 25, 21, 47.6, 24, 26.4, 23.7, 38.4, 19, 21, 33.3, 27, 16.1, 21.3, 24.5, 20.4, 16.2, 25.2, 22.3, 22.1, 15, 36.8, 12.5, 11.4, 7, 11.7, 7.6, 6.3, 14.8, 11.8, 18, 13, 7.8, 7, 9.3, 7.2, 7.9, 8.7, 22.6, 49.5, 41.2, 47, 41, 31, 16, 19, 30.1, 24, 22.8, 18.6, 20.5, 30.2, 37.8, 35, 30, 37.5, 29, 16.6, 17.1, 13.4, 22.7, 20, 122.5, 31, 35.5, 13, 21.6, 25.8, 32.4, 16, 79.6, 15.5, 11, 11.2, 13.8, 13.5, 12, 3.5, 15.1, 12.9, 17.2, 10.8, 2.5, 3.3, 3.6, 3.3, 4.1, 3.3, 3.9, 4.8, 4.8, 3.9, 5.3, 3.8, 4.9, 9.5, 10.5, 8.4, 4.6, 5.5, 7.7, 7.4, 7.9, 8.3, 5.5, 5.4, 3.9, 7.4, 5.5, 5.2, 16.3, 11.8, 17.8, 14.8, 7.3, 5.5, 5.6, 5.8, 7, 26.6, 23.4, 12, 6.1, 8.7, 6.9, 5.3, 8.3, 7.1, 8.6,

```
8.3, 9.7, 9.9, 8.3, 5.8, 23.1, 5.9, 4.5, 6.3, 4.5, 6.1, 5.7, 5.9, 4.5, 5.5, 8.2, 7.3, 6.3, 4.7, 3.8, 4.3,
4, 6.4, 5.3, 4.4, 3.8, 4.2, 4.5, 7.3, 7, 8.4, 6.3, 4.5, 14, 10.8, 20, 11.2, 10.4, 15, 10, 19, 8.7, 7.9,
21.2, 14, 7.2, 23.6, 6.7, 10, 9.5, 9.8, 7.8, 8.9, 11.5, 4.8, 20.2, 12.4, 11.5, 40, 69, 2.2, 2.2, 2.9,
4, 2.8, 3.5, 3.1, 3.2, 3.2, 3.2, 2.5, 6.2, 29.8)

# Define a grid of starting values for each parameter
start_grid <- expand.grid(
  B = seq(1, 10, by = 1),
  C = seq(-0.1, -1, by = -0.1),
  D = seq(1, 10, by = 1),
  E = seq(0.1, 1, by = 0.1)
)

# Fit the model using nls2 with multiple starting values
fit <- nls2(y ~ custom_function(x, z, B, C, D, E),
            start = start_grid,
            algorithm = "brute-force")

# Print the summary of the fitted model
summary(fit)
```

<u>SEM model in R code, which outputs maximal lifespan values for one or more adult body masses inputted</u>

### Be aware that the R console has a maximum width for R code and so if a lot of data is inputted into the following code it must be broken up with line breaks so the width doesn’t get too great (if this isn’t done, when trying to run the code you will get an error message, which doesn’t explicitly specify that this width issue is the reason for the error, but it is, and when solved the R code runs fine).

```
# Define initial values for x
x_values <- c(1, 2, 3, 4, 5) # Add adult body mass input data into here

# Note that there is a logging step before the equations and a de-logging step after them
# Iterate through each equation and update x values
```

```
for (i in 1:length(x_values)) {
  x_values[i] <- log10(x_values[i])
  x_values[i] <- -0.375 * x_values[i] - 0.902
  x_values[i] <- -0.875 * x_values[i] - 0.583
  x_values[i] <- -0.617 * x_values[i] + 0.431
  x_values[i] <- 1.174 * x_values[i] - 2.889
  x_values[i] <- 0.663 * x_values[i] + 1.449
  x_values[i] <- 0.973 * x_values[i] + 1.768
  x_values[i] <- -0.941 * x_values[i] + 2.567
  x_values[i] <- 10^(x_values[i])
}

# Create a data frame with the updated values of x
output_data <- data.frame(x = x_values)

# Write the data frame to a CSV file (will be put in whatever working directory is set for R, which can be set in its GUI). Output will all be in a single column when this file is opened in Excel.
write.csv(output_data, file = "output.csv", row.names = FALSE)

# Printing it in the console as well as writing it to a CSV file: Belt and Braces delivery
print(output_data)
```

<u>SEM model (that includes postnatal growth rate) in R code, which outputs maximal lifespan values for one or more adult body masses inputted</u>

```
# Be aware that the R console has a maximum width for R code and so if a lot of data is inputted into the following code it must be broken up with line breaks so the width doesn't get too great (if this isn't done, when trying to run the code you will get an error message, which doesn't explicitly specify that this width issue is the reason for the error, but it is, and when solved the R code runs fine).

# Define initial values for x and y
x_values <- c(1, 2, 3, 4, 5)
y_values <- rep(NA, length(x_values))
```

### Note that there is a logging step before the equations and a de-logging step after them
### Iterate through each equation and update x and y values
```
for (i in 1:length(x_values)) {
  x_values[i] <- log10(x_values[i])
  y_values[i] <- -0.293 * x_values[i] - 1.769
  x_values[i] <- -0.377 * x_values[i] - 0.903
  x_values[i] <- -0.866 * x_values[i] - 0.576
  x_values[i] <- -0.615 * x_values[i] + 0.431
  x_values[i] <- 1.168 * x_values[i] - 2.885
  x_values[i] <- 0.496 * x_values[i] + 0.969
  x_values[i] <- 0.980 * x_values[i] + 1.759
  x_values[i] <- -0.694 * x_values[i] + 1.584 + (-0.317 * y_values[i])
  x_values[i] <- 10^(x_values[i])
}

# Create a data frame with the updated values of x
output_data <- data.frame(x = x_values)

# Write the data frame to a CSV file
write.csv(output_data, file = "output.csv", row.names = FALSE)

# Printing it in the console as well as writing it to a CSV file: Belt and Braces delivery
print(output_data)
```

<u>R code used to perform Principal Component Analysis (PCA)</u>

```
# Load required libraries. Need to have R package factoextra installed and any R package dependencies thereof
library(factoextra)

# Read your dataset (replace 'data.csv' with your file path)
data <- read.csv("data.csv")

# Center and scale the data
```

```
scaled_and_centered_data <- scale(data, center = TRUE, scale = TRUE)

# Perform PCA
pca_result <- prcomp(scaled_and_centered_data)

# Extract principal components
principal_components <- as.data.frame(pca_result$x)

# Write transformed variables to a CSV file (replace 'transformed_data.csv' with desired file path)
write.csv(principal_components, "transformed_data.csv", row.names = FALSE)
```

#### SUPPLEMENTARY MATERIAL REFERENCES

[296] Pamplona R, Jové M, Mota-Martorell N, Barja G (2021) Is the NDUFV2 subunit of the hydrophilic complex I domain a key determinant of animal longevity? The FEBS Journal.

[307] Rouslin W, Broge CW, Grupp IL (1990) ATP depletion and mitochondrial functional loss during ischemia in slow and fast heart-rate hearts. American Journal of Physiology-Heart and Circulatory Physiology. 259(6):H1759-H1766.
[308] Rouslin W, Broge CW (1989) Regulation of mitochondrial matrix pH and adenosine 5′-triphosphatase activity during ischemia in slow heart-rate hearts: Role of Pi/H+ symport. Journal of Biological Chemistry. 264(26):15224-15229.
[309] Satre M, De Jerphanion MB, Huet J, Vignais PV (1975) ATPase inhibitor from yeast mitochondria. Purification and properties. Biochimica et Biophysica Acta (BBA)-Bioenergetics. 387(2):241-55.
[310] Hill AB (1965) The environment and disease: association or causation?
[311] Fisher RA (1958) Cancer and smoking. Nature. 182(4635):596-.
[312] Damuth J (1981) Population density and body size in mammals. Nature. 290(5808):699-700.
[313] Hutchinson GE, MacArthur RH (1959) A theoretical ecological model of size distributions among species of animals. The American Naturalist. 93(869):117-25.
[314] Brown JH, Nicoletto PF (1991) Spatial scaling of species composition: body masses of North American land mammals. The American Naturalist. 138(6):1478-512.
[315] Bakker VJ, Kelt DA (2000) Scale-dependent patterns in body size distributions of neotropical mammals. Ecology. 81(12):3530-47.
[316] Promislow DE (1991) Senescence in natural populations of mammals: a comparative study. Evolution. 45(8):1869-87.
[317] Scholander PF, Hock R, Walters V, Johnson F, Irving L (1950) Heat regulation in some arctic and tropical mammals and birds. The Biological Bulletin. 99(2):237-58.
[318] White CR, Seymour RS (2004) Does basal metabolic rate contain a useful signal? Mammalian BMR allometry and correlations with a selection of physiological, ecological, and life-history variables. Physiological and Biochemical Zoology. 77(6):929-41.
[319] Felsenstein J (1985) Phylogenies and the comparative method. The American Naturalist. 125(1):1-5.
[320] Midford PE, Garland T Jr, Maddison WP (2005) PDAP Package of Mesquite. Sourced from https://mesquiteproject.org/pdap_mesquite/index.html.
[321] Maddison WP and Maddison DR. Mesquite: a modular system for evolutionary analysis. Version 3.81. Downloaded from http://www.mesquiteproject.org.
[322] Kumar S, Suleski M, Craig JE, Kasprowicz AE, Sanderford M, Li M, Stecher G, Hedges SB (2022) TimeTree 5: An Expanded Resource for Species Divergence Times.

Molecular Biology and Evolution. DOI: 10.1093/molbev/msac174. Available on the web at: https://timetree.org/.
[323] Garland Jr T, Harvey PH, Ives AR (1992) Procedures for the analysis of comparative data using phylogenetically independent contrasts. Systematic biology. 41(1):18-32.
[324] Quader S, Isvaran K, Hale RE, Miner BG, Seavy NE (2004) Nonlinear relationships and phylogenetically independent contrasts. Journal of Evolutionary Biology. 17(3):709-15.
[325] Accessed in December 2023 from https://biologyfiles.ucr.edu/people/faculty/Garland/PDTREE_Mesquite.doc.
[326] Purvis A (1995) A composite estimate of primate phylogeny. Philosophical Transactions of the Royal Society of London. Series B: Biological Sciences. 348(1326):405-21.
[327] Pagel MD (1992) A method for the analysis of comparative data. Journal of theoretical Biology. 156(4):431-42.
[328] Grafen A (1989) The phylogenetic regression. Philosophical Transactions of the Royal Society of London. B, Biological Sciences. 326(1233):119-57.
[329] Gould SJ, Eldredge N (1972) Punctuated equilibria: an alternative to phyletic gradualism. Models in paleobiology. 82-115.
[330] Garland Jr T, Dickerman AW, Janis CM, Jones JA (1993) Phylogenetic analysis of covariance by computer simulation. Systematic biology. 42(3):265-92.
[331] Accessed in December 2023: https://www.statskingdom.com/linear-regression-calculator.html.
[332] Kim S (2015) ppcor: an R package for a fast calculation to semi-partial correlation coefficients. Communications for statistical applications and methods. 22(6):665.
[333] Kim S (2022) P-value calculation methods for semi-partial correlation coefficients. Communications for statistical applications and methods. 29(3):397.
[334] Wood SN (2011) Fast stable restricted maximum likelihood and marginal likelihood estimation of semiparametric generalized linear models. Journal of the Royal Statistical Society (B) 73(1):3-36.
[335] Grothendieck G (2022) nls2: Non-Linear Regression with Brute Force. R package version 0.3-3, <https://CRAN.R-project.org/package=nls2>.
[336] Storlund RL, Rosen DA, Trites AW (2021) Electrocardiographic scaling reveals differences in electrocardiogram interval durations between marine and terrestrial mammals. Frontiers in Physiology. 12:690029.

[363] Savage VM, Gillooly JF, Woodruff WH, West GB, Allen AP, Enquist BJ, Brown JH (2004) The predominance of quarter-power scaling in biology. Functional Ecology. 18(2):257-282.
[364] Prinzinger R (2005) Programmed ageing: the theory of maximal metabolic scope: How does the biological clock tick? EMBO reports. 6(S1):S14-S19.
[365] Porter RK, Brand MD (1995) Causes of differences in respiration rate of hepatocytes from mammals of different body mass. American Journal of Physiology-Regulatory, Integrative and Comparative Physiology. 269(5):R1213-R1224.
[366] Laidler KJ (1984) The development of the Arrhenius equation. Journal of chemical Education. 61(6):494.
[367] Eyring H (1935) The activated complex in chemical reactions. The Journal of Chemical Physics. 3(2):107-15.
[368] Taylor CR, Schmidt-Nielsen K, Raab JL (1970) Scaling of energetic cost of running to body size in mammals. American Journal of Physiology-Legacy Content. 219(4):1104-7.
[369] White CR, Alton LA, Crispin TS, Halsey LG (2016) Phylogenetic comparisons of pedestrian locomotion costs: confirmations and new insights. Ecology and evolution. 6(18):6712-20.
[370] Taylor CR, Heglund NC, Maloiy GM (1982) Energetics and mechanics of terrestrial locomotion. I. Metabolic energy consumption as a function of speed and body size in birds and mammals. Journal of Experimental Biology. 97(1):1-21.
[371] Alexander RM (2005) Models and the scaling of energy costs for locomotion. Journal of Experimental Biology. 208(9):1645-52.
[372] Schmidt-Nielsen K (1972) Locomotion: energy cost of swimming, flying, and running. Science. 177(4045):222-8.
[373] Brand MD, Couture P, Else PL, Withers KW, Hulbert AJ (1991) Evolution of energy metabolism. Proton permeability of the inner membrane of liver mitochondria is greater in a mammal than in a reptile. Biochemical Journal. 275(1):81-6.
[374] Brookes PS, Buckingham JA, Tenreiro AM, Hulbert AJ, Brand MD (1998) The proton permeability of the inner membrane of liver mitochondria from ectothermic and endothermic vertebrates and from obese rats: correlations with standard metabolic rate and phospholipid fatty acid composition. Comparative Biochemistry and Physiology Part B: Biochemistry and Molecular Biology. 119(2):325-34.

[375] Else PL, Hulbert AJ (1985) An allometric comparison of the mitochondria of mammalian and reptilian tissues: the implications for the evolution of endothermy. Journal of Comparative Physiology B. 156:3-11.
[376] Parsons K (2014) Human thermal environments: the effects of hot, moderate, and cold environments on human health, comfort, and performance. CRC press.
[377] Bligh J, Johnson KG (1973) Glossary of terms for thermal physiology. Journal of Applied Physiology. 35(6):941-61.
[378] Peto MV, De la Guardia C, Winslow K, Ho A, Fortney K, Morgen E (2017) MortalityPredictors.org: a manually-curated database of published biomarkers of human all cause mortality. Aging (Albany NY). 9(8):1916.
[379] Goldbogen JA, Cade DE, Calambokidis J, Czapanskiy MF, Fahlbusch J, Friedlaender AS, Gough WT, Kahane-Rapport SR, Savoca MS, Ponganis KV, Ponganis PJ (2019) Extreme bradycardia and tachycardia in the world's largest animal. Proceedings of the National Academy of Sciences. 116(50):25329-32.
[380] Mundim KC, Baraldi S, Machado HG, Vieira FM (2020) Temperature coefficient (Q10) and its applications in biological systems: Beyond the Arrhenius theory. Ecological Modelling. 431:109127.
[381] Sterratt DC (2015) Q10: the effect of temperature on ion channel kinetics. In Encyclopedia of Computational Neuroscience. 2551-2552. Springer New York.
[382] Grimes KM, Voorhees A, Chiao YA, Han HC, Lindsey ML, Buffenstein R (2014) Cardiac function of the naked mole-rat: ecophysiological responses to working underground. American Journal of Physiology-Heart and Circulatory Physiology. 306(5):H730-7.
[383] Riely CA, Cohen G, Lieberman M (1974) Ethane evolution: a new index of lipid peroxidation. Science. 183(4121):208-210.
[384] Kneepkens CF, Lepage G, Roy CC (1994) The potential of the hydrocarbon breath test as a measure of lipid peroxidation. Free Radical Biology and Medicine. 17(2):127-160.
[385] Topp H, Vangala M, Kritzler K, Schöch G (1995) Assessment of lipid peroxidation in rats of different body weight by determining expired ethane. Biological Chemistry Hoppe-Seyler. 376(11):691-694.
[386] Sagai M, Ichinose T (1980) Age-related changes in lipid peroxidation as measured by ethane, ethylene, butane and pentane in respired gases of rats. Life sciences. 27(9):731-738.
[387] Zarling EJ, Mobarhan S, Bowen P, Kamath S (1993) Pulmonary pentane excretion increases with age in healthy subjects. Mechanisms of ageing and development. 67(1-2):141-147.

[400] Ward WF, Qi W, Remmen HV, Zackert WE, Roberts LJ, Richardson A (2005) Effects of age and caloric restriction on lipid peroxidation: measurement of oxidative stress by F2-isoprostane levels. The Journals of Gerontology Series A: Biological Sciences and Medical Sciences. 60(7):847-51.
[401] Pratico D, Barry OP, Lawson JA, Adiyaman M, Hwang SW, Khanapure SP, Iuliano L, Rokach J, FitzGerald GA (1998) IPF2α-I: an index of lipid peroxidation in humans. Proceedings of the National Academy of Sciences. 95(7):3449-54.
[402] Montuschi P, Barnes PJ, Roberts LJ (2004) Isoprostanes: markers and mediators of oxidative stress. The FASEB journal. 18(15):1791-800.
[403] Milne GL, Yin H, Brooks JD, Sanchez S, Roberts II LJ, Morrow JD (2007) Quantification of F2-isoprostanes in biological fluids and tissues as a measure of oxidant stress. Methods in enzymology. 433:113-26.
[404] Morrow JD, Roberts II LJ (1999) Mass spectrometric quantification of F2-isoprostanes in biological fluids and tissues as measure of oxidant stress. Methods in enzymology. 300: 3-12.
[405] García-Bermúdez J, Sánchez-Aragó M, Soldevilla B, del Arco A, Nuevo-Tapioles C, Cuezva JM (2015) PKA phosphorylates the ATPase inhibitory factor 1 and inactivates its capacity to bind and inhibit the mitochondrial H+-ATP synthase. Cell reports. 12(12):2143-55.
[406] Horvath S, Lu AT, Cohen H, Raj K (2019) Rapamycin retards epigenetic ageing of keratinocytes independently of its effects on replicative senescence, proliferation and differentiation. Aging (Albany NY). 11(10):3238.
[407] Lujan C, Tyler EJ, Ecker S, Webster AP, Stead ER, Miguel VEM, ... & Bjedov I (2020) A CellAgeClock for expedited discovery of anti-ageing compounds. bioRxiv:803676.
[408] Macip CC, Hasan R, Hoznek V, Kim J, Lu YR, Metzger IV LE, Sethna S, Davidsohn N (2024) Gene Therapy-Mediated Partial Reprogramming Extends Lifespan and Reverses Age-Related Changes in Aged Mice. Cellular Reprogramming. 26(1):24-32.
[409] Pennacchi PC, de Almeida ME, Gomes OL, Faião-Flores F, de Araújo Crepaldi MC, Dos Santos MF, de Moraes Barros SB, Maria-Engler SS (2015) Glycated reconstructed human skin as a platform to study the pathogenesis of skin aging. Tissue Engineering Part A. 21(17-18):2417-25.
[410] Tsai SP, Hardy RJ, Wen CP (1992) The standardized mortality ratio and life expectancy. American journal of epidemiology. 135(7):824-31.

[436] Harding C, Pompei F, Bordonaro SF, McGillicuddy DC, Burmistrov D, Sanchez LD (2019) The daily, weekly, and seasonal cycles of body temperature analyzed at large scale. Chronobiology International. 36(12):1646-57.
[437] Olshansky SJ, Carnes BA, Cassel C (1990) In search of Methuselah: estimating the upper limits to human longevity. Science. 250(4981):634-40.
[438] Klegeris A, Schulzer M, Harper DG, McGeer PL (2007) Increase in core body temperature of Alzheimer's disease patients as a possible indicator of chronic neuroinflammation: a meta-analysis. Gerontology. 53(1):7-11.
[439] Jung CG, Kato R, Zhou C, Abdelhamid M, Shaaban EI, Yamashita H, Michikawa M (2022) Sustained high body temperature exacerbates cognitive function and Alzheimer's disease-related pathologies. Scientific Reports. 12(1):1-1.
[440] https://www.fda.gov/drugs/information-health-care-professionals-drugs/accelerated-approval-program (accessed on 31 July 2022).
[441] Knight EM, Brown TM, Gümüsgöz S, Smith JC, Waters EJ, Allan SM, Lawrence CB (2013) Age-related changes in core body temperature and activity in triple-transgenic Alzheimer's disease (3xTgAD) mice. Disease models & mechanisms. 6(1):160-70.
[442] Chung CW, Stephens AD, Konno T, Ward E, Avezov E, Kaminski CF, Hassanali AA, Kaminski Schierle GS (2022) Intracellular Aβ42 aggregation leads to cellular thermogenesis. Journal of the American Chemical Society.
[443] Noorani AA, Yamashita H, Gao Y, Islam S, Sun Y, Nakamura T, Enomoto H, Zou K, Michikawa M (2020) High temperature promotes amyloid β-protein production and γ-secretase complex formation via Hsp90. Journal of Biological Chemistry. 295(52):18010-22.
[444] Naiki H, Nakakuki K (1996) First-order kinetic model of Alzheimer's beta-amyloid fibril extension in vitro. Laboratory investigation; a journal of technical methods and pathology. 74(2):374-83.
[445] Kusumoto Y, Lomakin A, Teplow DB, Benedek GB (1998) Temperature dependence of amyloid β-protein fibrillization. Proceedings of the National Academy of Sciences. 95(21):12277-82.
[446] Sabaté R, Gallardo M, Estelrich J (2005) Temperature dependence of the nucleation constant rate in β amyloid fibrillogenesis. International journal of biological macromolecules. 35(1-2):9-13.
[447] Levine Iii H (2004) Alzheimer's β-peptide oligomer formation at physiologic concentrations. Analytical biochemistry. 335(1):81-90.

[482] Forster MJ, Dubey A, Dawson KM, Stutts WA, Lal H, Sohal RS (1996) Age-related losses of cognitive function and motor skills in mice are associated with oxidative protein damage in the brain. Proceedings of the National Academy of Sciences. 93(10):4765-4769.
[483] Halliwell B (2001) Role of free radicals in the neurodegenerative diseases. Drugs & aging. 18(9):685-716.
[484] Abou-Sleiman PM, Muqit MM, Wood NW (2006) Expanding insights of mitochondrial dysfunction in Parkinson's disease. Nature Reviews Neuroscience. 7(3):207-219.
[485] Halliwell B (1992) Reactive oxygen species and the central nervous system. Journal of neurochemistry. 59(5):1609-1623.
[486] Radi Isola RA, Castro Peyronel LA, Rodríguez Rey MI, Cassina Gómez AM, Thomson Garibotti ML, Flint Beal M, ... & Bodis-Wollner I (1997) Free radical damage to mitochondria. Mitochondria and Free Radicals in Neurodegenerative Diseases.
[487] Pope S, Land JM, Heales SJ (2008) Oxidative stress and mitochondrial dysfunction in neurodegeneration; cardiolipin a critical target? Biochimica et Biophysica Acta (BBA)-Bioenergetics. 1777(7-8):794-799.
[488] Bender A, Krishnan KJ, Morris CM, Taylor GA, Reeve AK, Perry RH, ... & Turnbull DM (2006) High levels of mitochondrial DNA deletions in substantia nigra neurons in aging and Parkinson disease. Nature genetics. 38(5):515-517.
[489] Massaad CA, Pautler RG, Klann E (2009) Mitochondrial superoxide: a key player in Alzheimer's disease. Aging (Albany NY). 1(9):758.
[490] Massaad CA, Washington TM, Pautler RG, Klann E (2009) Overexpression of SOD-2 reduces hippocampal superoxide and prevents memory deficits in a mouse model of Alzheimer's disease. Proceedings of the National Academy of Sciences. 106(32):13576-13581.
[491] Perry G, Nunomura A, Hirai K, Zhu X, Prez M, Avila J, ... & Smith MA (2002) Is oxidative damage the fundamental pathogenic mechanism of Alzheimer's and other neurodegenerative diseases? Free Radical Biology and Medicine. 33(11):1475-1479.
[492] Harman D (2002) Alzheimer's disease: role of aging in pathogenesis. Annals of the New York Academy of Sciences. 959(1):384-395.
[493] Hardy JA, Higgins GA (1992) Alzheimer's disease: the amyloid cascade hypothesis. Science. 256(5054):184-5.
[494] Makin S (2018) The amyloid hypothesis on trial. Nature. 559(7715):S4-.
[495] Thambisetty M, Howard R, Glymour MM, Schneider LS (2021) Alzheimer's drugs: Does reducing amyloid work? Science. 374(6567):544-5.

[517] Praticò D, Clark CM, Liun F, Lee VY, Trojanowski JQ (2002) Increase of brain oxidative stress in mild cognitive impairment: a possible predictor of Alzheimer disease. Archives of neurology. 59(6):972-6.
[518] Dumont M, Wille E, Stack C, Calingasan NY, Beal MF, Lin MT (2009) Reduction of oxidative stress, amyloid deposition, and memory deficit by manganese superoxide dismutase overexpression in a transgenic mouse model of Alzheimer's disease. The FASEB Journal. 23(8):2459-66.
[519] Mao P, Manczak M, Calkins MJ, Truong Q, Reddy TP, Reddy AP, Shirendeb U, Lo HH, Rabinovitch PS, Reddy PH (2012) Mitochondria-targeted catalase reduces abnormal APP processing, amyloid β production and BACE1 in a mouse model of Alzheimer's disease: implications for neuroprotection and lifespan extension. Human Molecular Genetics. 21(13):2973-90.
[520] McManus MJ, Murphy MP, Franklin JL (2011) The mitochondria-targeted antioxidant MitoQ prevents loss of spatial memory retention and early neuropathology in a transgenic mouse model of Alzheimer's disease. Journal of Neuroscience. 31(44):15703-15.
[521] Young ML, Franklin JL (2019) The mitochondria-targeted antioxidant MitoQ inhibits memory loss, neuropathology, and extends lifespan in aged 3xTg-AD mice. Molecular and Cellular Neuroscience. 101:103409.
[522] Lee HP, Pancholi N, Esposito L, Previll LA, Wang X, Zhu X, Smith MA, Lee HG (2012) Early induction of oxidative stress in mouse model of Alzheimer disease with reduced mitochondrial superoxide dismutase activity. PloS one. 7(1):e28033.
[523] Leuner K, Schütt T, Kurz C, Eckert SH, Schiller C, Occhipinti A, Mai S, Jendrach M, Eckert GP, Kruse SE, Palmiter RD (2012) Mitochondrion-derived reactive oxygen species lead to enhanced amyloid beta formation. Antioxidants & redox signaling. 16(12):1421-33.
[524] Li F, Calingasan NY, Yu F, Mauck WM, Toidze M, Almeida CG, Takahashi RH, Carlson GA, Flint Beal M, Lin MT, Gouras GK (2004) Increased plaque burden in brains of APP mutant MnSOD heterozygous knockout mice. Journal of neurochemistry. 89(5):1308-12.
[525] Melov S, Adlard PA, Morten K, Johnson F, Golden TR, Hinerfeld D, Schilling B, Mavros C, Masters CL, Volitakis I, Li QX (2007) Mitochondrial oxidative stress causes hyperphosphorylation of tau. PloS one. 2(6):e536.
[526] Esposito L, Raber J, Kekonius L, Yan F, Yu GQ, Bien-Ly N, Puoliväli J, Scearce-Levie K, Masliah E, Mucke L (2006) Reduction in mitochondrial superoxide dismutase modulates

Alzheimer's disease-like pathology and accelerates the onset of behavioral changes in human amyloid precursor protein transgenic mice. Journal of Neuroscience. 26(19):5167-79.
[527] Sung S, Yao Y, Uryu K, Yang H, Lee VM, Trojanowski JQ, Praticò D (2004) Early vitamin E supplementation in young but not aged mice reduces Aβ levels and amyloid deposition in a transgenic model of Alzheimer's disease. The FASEB journal. 18(2):323-5.
[528] Sano M, Ernesto C, Thomas RG, Klauber MR, Schafer K, Grundman M, Woodbury P, Growdon J, Cotman CW, Pfeiffer E, Schneider LS (1997) A controlled trial of selegiline, alpha-tocopherol, or both as treatment for Alzheimer's disease. New England Journal of Medicine. 336(17):1216-22.
[529] Zandi PP, Anthony JC, Khachaturian AS, Stone SV, Gustafson D, Tschanz JT, Norton MC, Welsh-Bohmer KA, Breitner JC, Cache County Study Group (2004) Reduced risk of Alzheimer disease in users of antioxidant vitamin supplements: the Cache County Study. Archives of neurology. 61(1):82-8.
[530] Morris MC, Evans DA, Bienias JL, Tangney CC, Wilson RS (2002) Vitamin E and cognitive decline in older persons. Archives of neurology. 59(7):1125-32.
[531] Engelhart MJ, Geerlings MI, Ruitenberg A, van Swieten JC, Hofman A, Witteman JC, Breteler MM (2002) Dietary intake of antioxidants and risk of Alzheimer disease. Jama. 287(24):3223-9.
[532] Sevigny J, Chiao P, Bussière T, Weinreb PH, Williams L, Maier M, Dunstan R, Salloway S, Chen T, Ling Y, O'Gorman J (2016) The antibody aducanumab reduces Aβ plaques in Alzheimer's disease. Nature. 537(7618):50-6.
[533] Kastanenka KV, Bussiere T, Shakerdge N, Qian F, Weinreb PH, Rhodes K, Bacskai BJ (2016) Immunotherapy with aducanumab restores calcium homeostasis in Tg2576 mice. Journal of Neuroscience. 36(50):12549-58.
[534] Tagliavini F, Tiraboschi P, Federico A (2021) Alzheimer's disease: The controversial approval of Aducanumab. Neurological Sciences. 42(8):3069-70.
[535] Howard R, Liu KY (2020) Questions EMERGE as Biogen claims aducanumab turnaround. Nature Reviews Neurology. 16(2):63-4.
[536] Knopman DS, Jones DT, Greicius MD (2021) Failure to demonstrate efficacy of aducanumab: An analysis of the EMERGE and ENGAGE trials as reported by Biogen, December 2019. Alzheimer's & Dementia. 17(4):696-701.
[537] Montine TJ, Markesbery WR, Morrow JD, Roberts III LJ (1998) Cerebrospinal fluid F2-isoprostane levels are increased in Alzheimer's disease. Annals of neurology. 44(3):410-3.

[551] Wolf-Klein GP, Silverstone FA, Lansey SC, Tesi D, Ciampaglia C, O'Donnell M, Galkowski J, Jaeger A, Wallenstein S, Leleiko NS (1995) Energy requirements in Alzheimer's disease patients. Nutrition (Burbank, Los Angeles County, Calif.). 11(3):264-8.
[552] Wang PN, Yang CL, Lin KN, Chen WT, Chwang LC, Liu HC (2004) Weight loss, nutritional status and physical activity in patients with Alzheimer's disease. Journal of neurology. 251(3):314-20.
[553] Wolf-Klein GP, Silverstone FA, Levy AP (1992) Nutritional patterns and weight change in Alzheimer patients. International Psychogeriatrics. 4(1):103-18.
[554] Cova I, Clerici F, Rossi A, Cucumo V, Ghiretti R, Maggiore L, Pomati S, Galimberti D, Scarpini E, Mariani C, Caracciolo B (2016) Weight loss predicts progression of mild cognitive impairment to Alzheimer's disease. PloS one. 11(3):e0151710.
[555] White H, Pieper C, Schmader K (1998) The association of weight change in Alzheimer's disease with severity of disease and mortality: a longitudinal analysis. Journal of the American Geriatrics Society. 46(10):1223-7.
[556] White HK, McConnell ES, Bales CW, Kuchibhatla M (2004) A 6-month observational study of the relationship between weight loss and behavioral symptoms in institutionalized Alzheimer's disease subjects. Journal of the American Medical Directors Association. 5(2):89-97.
[557] Knight EM, Verkhratsky A, Luckman SM, Allan SM, Lawrence CB (2012) Hypermetabolism in a triple-transgenic mouse model of Alzheimer's disease. Neurobiology of aging. 33(1):187-93.
[558] Vloeberghs E, Van Dam D, Franck F, Serroyen J, Geert M, Staufenbiel M, De Deyn PP (2008) Altered ingestive behavior, weight changes, and intact olfactory sense in an APP overexpression model. Behavioral neuroscience. 122(3):491.
[559] Morgan D, Gordon MN (2008) Amyloid, hyperactivity, and metabolism: theoretical comment on Vloeberghs et al.(2008).
[560] Pugh PL, Richardson JC, Bate ST, Upton N, Sunter D (2007) Non-cognitive behaviours in an APP/PS1 transgenic model of Alzheimer's disease. Behavioural brain research. 178(1):18-28.
[561] Cummings J, Feldman HH, Scheltens P (2019) The "rights" of precision drug development for Alzheimer's disease. Alzheimer's research & therapy. 11(1):1-14.
[562] Mather M, Jacobsen LA, Pollard KM (2015) Aging in the United States: population bulletin. Population Reference Bureau.

[563] Tahami Monfared AA, Byrnes MJ, White LA, Zhang Q (2022) The Humanistic and Economic Burden of Alzheimer's Disease. Neurology and Therapy. 1-27.
[564] Alzheimer's Study Group. A national Alzheimer's strategic plan: The report of the Alzheimer's study group.
[565] Urquhart L (2021) Top companies and drugs by sales in 2020. Nature reviews. Drug discovery. 20(4):253.
[566] Birnbaum H, Pike C, Kaufman R, Maynchenko M, Kidolezi Y, Cifaldi M (2010) Societal cost of rheumatoid arthritis patients in the US. Current medical research and opinion. 26(1):77-90.
[567] Naoi M, Maruyama W (1999) Cell death of dopamine neurons in aging and Parkinson's disease. Mechanisms of ageing and development. 111(2):175-88.
[568] Bender A, Krishnan KJ, Morris CM, Taylor GA, Reeve AK, Perry RH, Turnbull DM (2006) High levels of mitochondrial DNA deletions in substantia nigra neurons in aging and Parkinson disease. Nature genetics. 38(5):515-517.
[569] Kraytsberg Y, Kudryavtseva E, McKee AC, Geula C, Kowall NW, Khrapko K (2006) Mitochondrial DNA deletions are abundant and cause functional impairment in aged human substantia nigra neurons. Nature genetics. 38(5):518-520.
[570] Wong WL, Su X, Li X, Cheung CM, Klein R, Cheng CY, Wong TY (2014) Global prevalence of age-related macular degeneration and disease burden projection for 2020 and 2040: a systematic review and meta-analysis. The Lancet Global Health. 2(2):e106-16.
[571] Waring MJ, Arrowsmith J, Leach AR, Leeson PD, Mandrell S, Owen RM, Pairaudeau G, Pennie WD, Pickett SD, Wang J, Wallace O (2015) An analysis of the attrition of drug candidates from four major pharmaceutical companies. Nature reviews Drug discovery. 14(7):475-86.
[572] Meyer AS, McCain MD, Fang Q, Pegg AE, Spratt TE (2003) O 6-Alkylguanine− DNA Alkyltransferases Repair O 6-Methylguanine in DNA with Michaelis−Menten-like Kinetics. Chemical research in toxicology. 16(11):1405-1409.
[573] Blainey PC, van Oijen AM, Banerjee A, Verdine GL, Xie XS (2006) A base-excision DNA-repair protein finds intrahelical lesion bases by fast sliding in contact with DNA. Proceedings of the National Academy of Sciences. 103(15):5752-5757.
[574] Gorman J, Chowdhury A, Surtees JA, Shimada J, Reichman DR, Alani E, Greene EC (2007) Dynamic basis for one-dimensional DNA scanning by the mismatch repair complex Msh2-Msh6. Molecular cell. 28(3):359-370.

[575] Hobbs JK, Jiao W, Easter AD, Parker EJ, Schipper LA, Arcus VL (2013) Change in heat capacity for enzyme catalysis determines temperature dependence of enzyme catalyzed rates. ACS chemical biology. 8(11):2388-2393.
[576] Jones HB, Crean RM, Matthews C, Troya AB, Danson MJ, Bull SD, ... & Pudney CR (2018) Uncovering the relationship between the change in heat capacity for enzyme catalysis and vibrational frequency through isotope effect studies. ACS Catalysis. 8(6):5340-5349.
[577] Arcus VL, Mulholland AJ (2020) Temperature, dynamics, and enzyme-catalyzed reaction rates. Annual review of biophysics. 49:163-180.
[578] Arcus VL, Prentice EJ, Hobbs JK, Mulholland AJ, Van der Kamp MW, Pudney CR, ... & Schipper LA (2016) On the temperature dependence of enzyme-catalyzed rates. Biochemistry. 55(12):1681-1688.
[579] Owusu-Ansah E, Banerjee U (2009) Reactive oxygen species prime Drosophila haematopoietic progenitors for differentiation. Nature. 461(7263):537-41.
[580] Theopold U (2009) A bad boy comes good. Nature. 461(7263):486-7.
[581] Gough NR. Differentiated with ROS. Science Signaling. 2(90):ec321-.
[582] Baker DJ, Childs BG, Durik M, Wijers ME, Sieben CJ, Zhong J, Van Deursen JM (2016) Naturally occurring p16 Ink4a-positive cells shorten healthy lifespan. Nature. 530(7589):184-189.
[583] Vassilev LT, Vu BT, Graves B, Carvajal D, Podlaski F, Filipovic Z, Kong N, Kammlott U, Lukacs C, Klein C, Fotouhi N (2004) In vivo activation of the p53 pathway by small-molecule antagonists of MDM2. Science. 303(5659):844-8.
[584] Dolgin E (2020) Send in the senolytics. Nature Biotechnology. 38(12):1371-1378.
[585] https://clinicaltrials.gov (accessed on 25 July 2022).
[586] Hall SS (2015) The man who wants to beat back aging. Science. 16.
[587] Previtali D, Merli G, Di Laura Frattura G, Candrian C, Zaffagnini S, Filardo G (2021) The long-lasting effects of "placebo injections" in knee osteoarthritis: a meta-analysis. Cartilage. 13(1_suppl):185S-96S.
[588] Yousefzadeh MJ, Zhu YI, McGowan SJ, Angelini L, Fuhrmann-Stroissnigg H, Xu M, Ling YY, Melos KI, Pirtskhalava T, Inman CL, McGuckian C (2018) Fisetin is a senotherapeutic that extends health and lifespan. EBioMedicine. 36:18-28.
[589] Xu M, Pirtskhalava T, Farr JN, Weigand BM, Palmer AK, Weivoda MM, Inman CL, Ogrodnik MB, Hachfeld CM, Fraser DG, Onken JL (2018) Senolytics improve physical function and increase lifespan in old age. Nature medicine. 24(8):1246-56.

[590] Miller RA, Harrison DE, Astle CM, Floyd RA, Flurkey K, Hensley KL, Javors MA, Leeuwenburgh C, Nelson JF, Ongini E, Nadon NL (2007) An aging interventions testing program: study design and interim report. Aging cell. 6(4):565-75.
[591] https://www.youtube.com/watch?v=42PzfNs9egA (accessed on 2 August 2022).
[592] Fossel M (2019) Cell senescence, telomerase, and senolytic therapy. OBM Geriatrics. 3(1):1-.
[593] Grosse L, Wagner N, Emelyanov A, Molina C, Lacas-Gervais S, Wagner KD, Bulavin DV (2020) Defined $p16^{High}$ senescent cell types are indispensable for mouse healthspan. Cell Metabolism. 32(1):87-99.
